# Expression genome-wide association study and differential methylome profiling reveal upstream regulators of drought memory genes in *Arabidopsis thaliana*

**DOI:** 10.64898/2026.01.28.702454

**Authors:** Debankona Marik, Rishabh Kumar, Ayan Sadhukhan

**Affiliations:** Department of Bioscience and Bioengineering, Indian Institute of Technology Jodhpur, Jodhpur, Rajasthan, India

**Keywords:** Expression GWAS, recurrent drought, transcriptional memory, whole-genome bisulfite sequencing

## Abstract

Plants adapt to recurrent drought through transcriptional memory, yet the upstream regulators remain largely unknown. This study integrated expression genome-wide association study (eGWAS) across 115 *Arabidopsis thaliana* ecotypes with differential methylome profiling to identify these regulators. Focusing on the memory genes *LKR*, *HIS1-3*, and *DREB1A*, eGWAS identified signaling and epigenetic loci involved in ABA/JA responses and DNA methylation. Methylome profiling by whole-genome bisulfite sequencing of ecotypes contrasting in drought tolerance, as well as in superinduction of memory genes, revealed significantly greater methylation variation during the second drought (D2) than during the first (D1), highlighting the role of epigenetic reprogramming in memory maintenance. Functional validation using T-DNA mutants demonstrated specific modulation of the D2/D1 induction ratio without affecting initial drought responses. Mutants of *LKR* eGWAS-delineated genes AT1G56660, AT2G19120, AT4G16490, and *DEG3*, those of *HIS1-3* eGWAS genes AT1G14220, AT2G24960, AT3G10845, AT3G19340, *CNGC10*, *EMB2770*, *GRF7*, and *RPP2A*, and *DREB1A* eGWAS genes AT1G67000, AT3G61610, AT5G62110, *HK2*, *JMJ12*, and *LUP5* abolished respective memory gene induction. The eGWAS and methylome approaches converged on DNA repair, chromatin modification, vesicular transport, and proteostasis as core memory hubs. These findings reveal a genetic-epigenetic interplay that coordinates transcriptional memory, priming plants for rapid reactivation of stress pathways during recurrent drought.

## 1. Introduction

Plant growth is severely affected by drought, resulting in substantial agricultural yield losses (Marik et al. 2025), a situation further complicated by recurrent events. Plants’ ability to adapt to recurrent drought relies on their ‘memory’ of past events, which triggers a heightened response during subsequent drought episodes (Sadhukhan et al. 2022). Transcriptional memory involves short-term changes in gene expression that enhance stress responses over time. Transgenerational memory transmits stress tolerance across generations through heritable epigenetic changes, such as DNA methylation. Recurrent drought experiments, in which well-watered plants (W) undergo an initial drought treatment (D1), followed by a recovery phase (R) and then another drought (D2), and continued in this manner, led to the discovery of four categories of drought memory genes: +/+ genes associated with enhanced abscisic acid (ABA) biosynthesis and chaperones for protecting cellular structures from dehydration; −/+ genes related to the restoration of photosynthesis and the electron transport chain; +/− genes responsible for modulating phytohormone signaling and ion homeostasis; and −/− genes involving the reduction of chloroplast function during repeated drought events (Ding et al. 2013). In these experiments, the first +/− indicates a higher/lower fold-change in expression in D1 compared to W, and the second in D2 compared to D1. The memory of drought stress is not only mediated by hormone levels but also by stable chromatin remodeling, which involves DNA methylation and histone modifications, as well as the regulatory roles of non-coding RNAs. These allow stress-responsive genes to remain poised for reactivation. ABA and jasmonic acid (JA) signaling potentially help recruit chromatin remodelers near drought-responsive genes, thereby reinforcing a memory imprint, such as maintaining open chromatin and permissive methylation patterns, or facilitating the reactivation of stalled RNA polymerase II during D2 (Sadhukhan et al. 2022).

Plant DNA methylation occurs in CpG, CHG, and CHH sequence contexts through linked pathways involving maintenance methyltransferases, such as MET1, CMT3, and CMT2, as well as *de novo* RNA-directed DNA methylation (RdDM) mechanisms (Leichter et al. 2022). Conventionally, methylation silences a gene and demethylation activates it, both of which are profoundly altered by drought stress in plants. DNA methylation varies widely between genotypes and is influenced by nucleotide sequence variations in methylase genes, as identified through genome-wide association studies (GWAS) (Kawakatsu et al. 2016). Sensitive rice genotypes exhibited higher levels of methylation than tolerant genotypes (Kou et al. 2021). In a comparison of two fava bean cultivars, DNA methylation in root tissues decreased in the drought-resistant cultivar upon drought exposure, whereas it increased in the drought-sensitive variety (Abid et al. 2017). Drought stress triggers genotype-specific DNA methylation changes in chickpea: tolerant varieties exhibit broad hypomethylation alongside small RNA-directed CHH hypermethylation targeted at transposable elements, whereas sensitive varieties predominantly display hypermethylation. These contrasting methylation profiles control unique sets of drought-responsive genes and pathways (Gayacharan and Joel 2013; Gupta and Garg 2025). Repeated cycles of mild drought followed by re-watering modify the rice methylome in conjunction with transcriptional memory genes involved in hormone signaling pathways, secondary metabolism, and defense responses. More than 5,000 drought-memory transcripts correlate strongly with differentially methylated regions, underscoring the role of DNA methylation in regulating drought memory through enhanced gene expression during recurrent stress (Li et al. 2019).

Our understanding of transcriptional drought memory remains limited by the lack of identification of upstream regulators, including mechanisms of RNA polymerase II stalling and reactivation, and the unknown modulators of phytohormone signaling and the interplay between genetic and epigenetic factors (Sadhukhan et al. 2022). *Arabidopsis thaliana* acts as a key model organism for core plant biology research (Yaschenko et al. 2024). GWAS spanning hundreds of its ecotypes allows statistical pinpointing of complex quantitative traits (Uffelmann et al. 2021; He et al. 2023; Marik et al. 2026). Expression GWAS (eGWAS), which leverages natural variation in gene expression, is a powerful tool for identifying the upstream transcriptional regulation of stress-responsive genes (Sadhukhan et al. 2021; Yang et al. 2024). We previously employed a hydroponic phenotyping platform for *A. thaliana* using PEG-6000 to faithfully simulate drought conditions. Under these conditions, GWAS identified drought tolerance genes, which were subsequently validated by reverse genetic analysis in hydroponic assays (Marik et al. 2026). These hydroponic conditions were further employed in this study to identify transcriptional memory genes associated with drought. Next, we identified natural variation in transcriptional drought memory in natural *A. thaliana* ecotypes and employed eGWAS to identify upstream signaling genes involved in both genetic and epigenetic regulation of drought memory, some of which were validated by reverse genetics. In addition, identification of DNA methylation patterns by whole-genome bisulfite sequencing of ecotypes contrasting in drought memory behavior revealed differentially methylated genes under recurrent drought conditions, which functionally overlap and are interconnected with the eGWAS-identified genes in co-expression and protein-protein interaction networks. These results shed light on the interplay between genetic and epigenetic factors that regulate the transcriptional memory of drought.

## 2. Materials and Methods

### 2.1 Plant materials, growth conditions, and stress treatments

We maintained 115 *Arabidopsis thaliana* worldwide ecotypes, sourced from the Nottingham Arabidopsis Stock Centre (NASC, UK) and RIKEN Bioresource Research Centre (BRC, Japan), via single-seed descent (Marik et al. 2026) and freshly harvested seeds were used for eGWAS. The seeds were subjected to imbibition and stratification in deionized water at 4°C for three days to synchronize germination and were then pipetted onto nylon meshes mounted in photographic frames, which were floated on ¼-Hoagland’s growth medium, pH 5.8 (Hoagland and Arnon, 1938; Kobayashi et al. 2016; Marik et al. 2026). The growth room was maintained at 22 24°C and 55 60% humidity. T8 surface-mount LED tubes were used to supply a 12 h light period at a photosynthetic photon flux density of 40 μmol m ². The seedlings were grown under these conditions for 12 days, with the medium replaced every 2 days to maintain consistent nutrient levels. After 12 days, they were harvested and flash frozen in liquid nitrogen; this sample was designated as watered (W). More 12-day-old seedlings were subjected to the first drought stress (D1) by transferring them to ¼-Hoagland’s solution supplemented with 5% polyethylene glycol 6000 (PEG-6000) for 3 days. At the end of D1, samples were harvested and flash frozen in liquid nitrogen. Following D1, seedlings underwent a non-stressful recovery phase (R) of 3 days in ¼-Hoagland’s solution without PEG. After recovery, a second drought stress (D2) was applied by re-exposing the seedlings to 5% PEG-6000 in ¼-Hoagland’s solution for 3 days. Samples were collected at the end of D2 and rapidly frozen in liquid nitrogen.

### 2.2 RNA isolation, transcriptome, and quantitative PCR analysis

RNA was extracted from the samples using the CTAB method, with three biological replicates per treatment. The cDNA library was prepared using the NEBNext Ultra II RNA library kit (New England Biolabs, New Delhi, India), prior to sequencing on the Illumina NovaSeq 6000 V1.5 (Illumina Inc., Gurgaon, India). Raw sequencing reads underwent quality control, adapter trimming, and alignment to the *A. thaliana* reference genome (Araport 11) using standard pipelines. Differential gene expression was assessed using edgeR version 4.1.2 (Chen et al. 2025) and analyzed with Fisher’s exact test, adjusted with Benjamini-Hochberg FDR. The transcriptome data were further validated in-house using reverse transcription-quantitative polymerase chain reaction (RT-qPCR). For this, cDNA synthesis used the RevertAid Reverse Transcriptase Kit (Thermo Scientific, India) with random hexamer primers. RT-qPCR employed TB Green^®^ Premix Ex Taq™ (RR420; Takara, India) on a QuantStudio™ 5 Real-Time PCR System (Thermo Scientific). Primers were designed using the Primer3 web tool (Kõressaar et al. 2018), and relative gene expression levels were calculated using the ΔΔC_t_ method, with *AtUBQ1* as the internal reference gene. ΔC_t_ was calculated by subtracting the Ct values of *AtUBQ1* from those of the genes. ΔΔC_t_ was calculated by subtracting the value of ΔC_t_ (D1) from ΔC_t_ (D2), followed by 2^-ΔΔCt^ to analyse the fold change in gene expression.

### 2.3 Expression genome-wide association study (eGWAS)

A GWAS was performed in TASSEL version 5.0 (Bradbury et al. 2007) by utilising the relative fold change (RFC) in expression level (D2/D1) for selected memory genes in 115 *A. thaliana* ecotypes as the quantitative phenotype data and genotype data comprising of 105,856 single nucleotide polymorphisms (SNPs) sourced from the 1001 genomes database (https://1001genomes.org/), and applying a mixed linear model (MLM) keeping into account kinship and population structure (Sadhukhan et al. 2017; Marik et al. 2026). A quantile-quantile (Q-Q) plot comparing the observed and expected *P*-value distributions was generated using TASSEL (Marik et al. 2026). Ridge regression assessed the combined effects of top eGWAS SNPs using the glmnet R package (Friedman et al. 2010) with 5-fold cross-validation over 100 iterations. We plotted the mean correlation coefficient (*r*^2^) between observed and predicted phenotypes as a function of the number of top eGWAS SNPs. Haploblocks surrounding GWAS SNPs were defined by linkage disequilibrium (LD; *r*^2^ ≥ 0.6) of adjacent SNPs within a 10 kb LD decay window using TASSEL; genes within these blocks were retrieved from TAIR coordinates (https://www.arabidopsis.org/) via custom Excel macros. Denser polymorphism data for the ecotypes used in eGWAS were extracted using the POLYMORPH 1001 tool (http://tools.1001genomes.org/polymorph/). The promoter and 5’-untranslated regions (5’-UTRs) up to 1 kb upstream of the start codon, as well as the coding sequences (CDSs), were analyzed using GENETYX version 11. For CDSs, introns were removed from translation to amino acid sequences. A local association study (LAS) in TASSEL assessed the phenotypic associations of local genomic region polymorphisms in eGWAS-identified genes across 80 available ecotypes (out of 115) using both generalized linear models (GLMs) and MLMs (Sadhukhan et al. 2021).

### 2.4 Functional analysis of eGWAS-identified genes

Functional enrichment analysis of genes was conducted using ShinyGO 0.85.1 (Ge et al. 2020). The STRING database (https://string-db.org/) was utilized to conduct network analysis on the genes identified by eGWAS (Szklarczyk et al. 2023). We selected up to 50 first-shell interactors for GWAS-identified genes using a high-confidence STRING score cutoff of 0.7, leveraging coexpression data and experimentally validated protein-protein interactions to construct the network. The outcomes were represented using Cytoscape version 3.10.3 (Shannon et al. 2003). The hub genes were identified using Cytohubba, which employs the maximal clique centrality (MCC) algorithm (Ono et al. 2025). Subsequently, cluster analysis was conducted using the Molecular Complex Detection (MCODE) plugin, version 2.0.3, to identify hub genes with a node degree (i.e., the number of interacting partners) of 2 or more. The OMICS visualizer plugin facilitated the visualization of the functional enrichment analysis of the networking genes, depicted as colored rings encircling the nodes (Legeay et al. 2020).

### 2.5 Validation of homozygosity and phenotyping T-DNA insertion mutants

T-DNA insertion mutants for eGWAS-identified genes were identified via the Arabidopsis Gene Mapping Tool (http://signal.salk.edu/cgi-bin/tdnaexpress) and procured from NASC. Seeds were germinated hydroponically for 2 weeks, then transplanted to pots with a 1:2:1 soil:soilrite:vermiculite mix using Arasystem modules (https://www.arasystem.com/) for propagation. Leaf samples were frozen in liquid nitrogen, then bead-beaten, and genomic DNA was isolated using the CTAB method. Homozygosity of T-DNA lines and WT (Col-0) was confirmed by two PCR amplifications using genomic DNA as a template, and one with left and right genomic primers (LP and RP), and the other with RP and T-DNA left border primer (LB), as recommended by SALK (http://signal.salk.edu/tdnaprimers.2.html). Homozygous lines were propagated by single-seed descent and freshly harvested for phenotyping (Marik et al. 2026). Reduction in gene expression levels, i.e., gene knockdown (KD) in promoter, UTR, and intronic T-DNA insertion lines, was assessed by RT-qPCR after a 5-day drought on 10-day-old seedlings, comparing expression levels between mutants and WT. Independent knockout (KO; with T-DNA inserted in exons) and KD lines were grown in hydroponics for 12 days and subjected to the exact same stress regime used in the eGWAS. Samples were harvested at W, D1, and D2 and analyzed for gene expression by RT-qPCR. Relative fold changes between mutants and wild type (WT) were calculated, followed by one-way ANOVA and Tukey’s honestly significant difference (HSD) post-hoc test to identify significant differences among groups (*P* < 0.05).

### 2.6 Whole-genome bisulfite sequencing and identification of differentially methylated genes

High-quality genomic DNA was extracted from the samples and treated with sodium bisulfite using the Zymo-Seq WGBS Library Kit (Zymo Research, Irvine, CA, USA), converting unmethylated cytosines to uracils while preserving methylated cytosines. The converted DNA underwent purification, second-strand synthesis to generate a double-stranded template, tagmentation-based fragmentation, adapter tagging with bead-linked transposomes, and PCR amplification. Libraries were purified using magnetic beads and the Nextera kit (Illumina, San Diego, CA, USA) to eliminate adapter dimers and contaminants, then assessed for fragment size distribution with an Agilent TapeStation (Agilent Technologies, Santa Clara, CA, USA) and quantified via Qubit (Thermo Fisher Scientific, Waltham, MA, USA) prior to Illumina platform sequencing for whole genome bisulfite sequencing. Raw reads were trimmed with AdapterRemoval v2.3.2 (Q30 threshold), yielding clean data. High-quality reads were aligned, deduplicated, and processed for methylation extraction with Bismark v0.24.2 (Krueger & Andrews, 2011), followed by differential methylation analysis of CpG/CHG/CHH contexts with methylKit (R v4.3.1; Akalin et al. 2012). Methylated sites were annotated using HOMER v5 (Heinz et al. 2010).

## 3. Results

### 3.1 Transcriptional memory of drought stress in hydroponically grown *Arabidopsis thaliana*

Twelve-day-old *Arabidopsis thaliana* seedlings from W were subjected to D1 by transferring to PEG-infused media for three days, and their transcriptomes were analyzed by RNA sequencing. In D1, 1,100 genes were up-regulated (log_2_fold-change > 1, *P* < 0.05), and 486 genes were down-regulated (log_2_FC < 1, *P* < 0.05), compared to W. This was followed by a recovery period, R, of three days, during which 1,098 genes were up-regulated, and 526 were down-regulated. A second three-day drought episode, D2, resulted in the up-regulation of 1,207 genes and the down-regulation of 479 genes, compared to W. Compared to D1, 147 genes were up-regulated, and 51 were down-regulated in D2 (Fig. 1A, Table S1). From the transcriptome analysis, four types of memory genes emerged: 12 [+/+] primed/super-induced (W > D1 > D2), 38 [−/+] late-induced (W > D1 < D2), six [+/−] late-supressed (W < D1 > D2), and five [−/−] de-primed/super-supressed (W < D1 < D2) genes, following Ding et al. (2013) criteria (Table S2). A gene ontology (GO) functional enrichment analysis of the genes from each category revealed several enriched biological processes (Fig. 1B). The [+/+] genes were responsive to water deprivation, secondary metabolism, nucleosome assembly and chromosome condensation, L-lysine metabolism, autophagosome assembly, intracellular sterol transport, response to 1-aminocyclopropane-1-carboxylic acid, nucleoside and amino acid metabolism, plant secondary cell wall biosynthesis, and positive regulation of gravitropism. This indicates plant strategies for enhanced drought tolerance during D2, including rapid osmolyte accumulation, nutrient recycling, reinforced cell walls, root water foraging, membrane stabilization, and epigenetic priming. The [−/+] genes were responsive to water deprivation, salt and oxidative stresses, light stimulus, and ABA, and involved in transcriptional regulation, signal transduction, protein folding, photosynthesis and chlorophyll biosynthesis, skotomorphogenesis, ion transport, salicylic acid (SA) and jasmonate (JA) response, L-pipecolic acid biosynthes, oxylipin metabolism, and phosphoenolpyruvate transport, linked to drought stress responses. These signify transcriptional memory that delays but amplifies drought responses during D2, thereby enhancing cross-stress tolerance through hormone signaling, metabolic reprogramming, ion homeostasis readjustment, and restoration of photosynthesis, ultimately improving survival. The [+/−] genes were involved in defense responses and wounding, and included two chloroplast genes of the photosynthetic light-harvesting system of photosystem I (*psaA* and *psaB*). Additionally, the RNA polymerase beta subunit (*rpoC1*) gene, which is involved in transcriptional elongation, also belongs to this category. These genes represent transcriptional memory that triggers strong but non-sustained activation during D1, followed by repression in D2, thereby facilitating photoprotection and conserving resources for survival. The [−/−] genes were involved in defense, JA, light, and hypoxia responses, as well as the centrosome cycle and spindle assembly (Fig. 1B). These genes indicate hormonal signaling, metabolic adjustments, and mitotic stability during recurrent stress while avoiding unnecessary gene expression that could impair growth recovery. Following a stricter cutoff (*FDR* < 0.05) for each comparison, six [+/+], one [−/−], and 19 [−/+] genes were obtained. From these, we selected two [+/+] genes, *lysine-ketoglutarate reductase* (*LKR*; AT4G33150) and *histone H1-3* (*HIS1-3*; AT2G18050), along with one [−/+] gene, *dehydration response element-binding 1A* (*DREB1A*; AT4G25480), for further analysis after validation by qPCR (Fig. 1C).

**Fig. 1.**
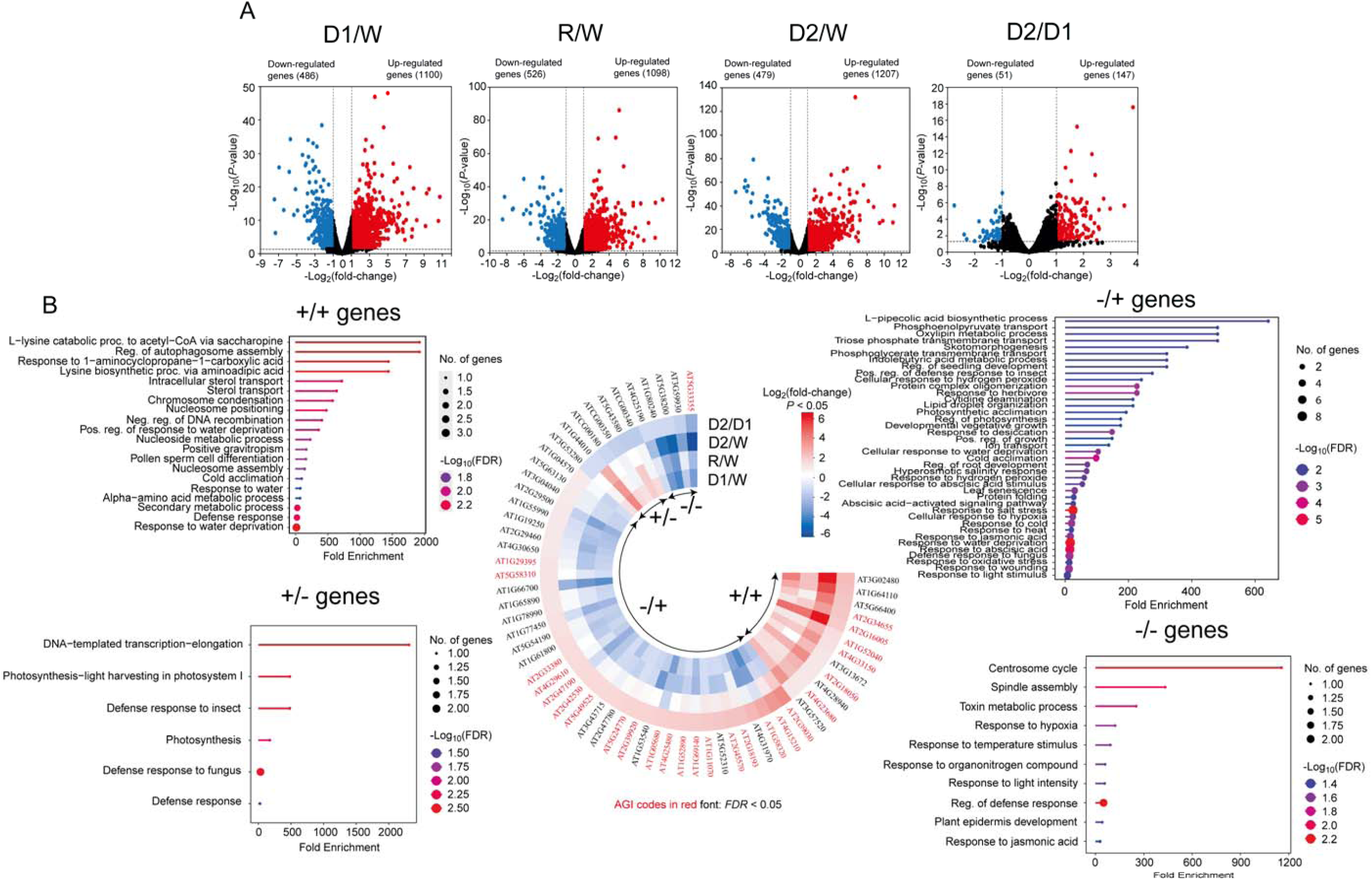
Transcriptome analysis of *Arabidopsis thaliana* seedlings under recurrent drought stress to identify memory genes. **(A)** Volcano plots of log_10_(*P*-value) versus log_2_FC for four comparisons: first drought (D1) vs. watered (W), recovery (R) vs W, D2 vs. W, and second drought (D2) vs. D1 are shown, where blue dots represent downregulated genes with log_2_(fold change) < −1, *P*-value < 0.05, and red dots symbolize upregulated genes with log_2_FC > 1, *P*-value < 0.05. *A. thaliana* ecotype Col-0 seeds were grown in ¼^-^Hoagland’s solution for 12 days (W), then subjected to 3 days of drought stress in ¼^-^Hoagland’s solution supplemented with 5% PEG-6000 (D1), followed by a recovery period of 3 days in ¼^-^Hoagland’s solution (R) and again treated with PEG for 3 days (D2). Whole seedlings were sampled from each condition for RNA sequencing. (**B)** A Circos plot displays the memory genes of four categories, with concentric semicircles indicating the four conditions and a color gradient from blue to red representing the log_2_fold change (log_2_FC) for each gene. Four types of memory genes were identified based on their log_2_FC values, namely +/+, −/+, +/−, and −/−, where the first sign indicates up (+) or down (−) regulation of genes in D1 vs. W, and the second sign indicates that in D2 vs. D1. The genes in red font passed the criteria: *false discovery rate* (*FDR*) < 0.05 for the comparisons D1 vs. W and D2 vs. D1. The lollipop plots display functional enrichment analysis, conducted using ShinyGO (https://bioinformatics.sdstate.edu/go/) for the four categories of memory genes. The circle color indicates −log_10_(*FDR*), and the circle size represents the number of genes in each Gene Ontology (GO) biological process category.

### 3.2 EGWAS under recurrent droughts unravels upstream regulators of memory genes

We observed natural variation in superinduction (D2/D1) of *LKR* and *HIS1-3*, as well as late induction (D2/D1) of *DREB1A*, in 115 *A. thaliana* ecotypes. The log_2_FC values for *LKR* ranged from −1.4 to 5.8, *HIS1-3* from −3.4 to 4.5, and *DREB1A* from −3.9 to 4.8 (Fig. 2A, F, K). To unravel the genetic regulation of transcriptional memory, eGWAS analyses were conducted for each memory gene, revealing SNPs that were significantly associated with the expression phenotype. Although few SNPs reached the stringent Bonferroni threshold for one million genome-wide SNPs (*P* < 4.7 × 10 ^7^), several exceeded acceptable suggestive cutoffs (*P* < 10) established for *A. thaliana* GWAS using permutation-based methods (Sadhukhan et al. 2021). Given the modest population size (115 ecotypes) and the application of MLM kinship correction, the Q-Q plot tail deviations at *P* < 10 ^3^–10 ^4^ suggest true associations (Fig. 2B, G, L). In each eGWAS, the top 200 ranking SNPs cumulatively account for ∼80% of the natural variation, as evident from the ridge regression analyses (Fig. 2C, H, M). The nearest genes within haploblocks, located within 20 Kb LD (*r^2^* > 0.6) windows of the eGWAS-identified SNPs, were identified, revealing 10, 14, and six genes at *P* < 10^-4^ (Fig. 2D, I, N; Manhattan plots) and 120, 111, and 94 genes at *P* < 10^-3^ for *LKR*, *HIS1-3*, and *DREB1A* eGWASs, respectively (Tables S3–S5). Genes with *P* < 10 ³ were subjected to GO enrichment analysis, revealing several processes shared by the three eGWASs. These included signal transduction, responses to ABA and JA, oxidative stress response, RNA splicing, nucleotide metabolism, and DNA methylation. Several uniquely enriched processes, including the ubiquitin-dependent endoplasmic reticulum (ER)-associated degradation (ERAD) pathway, the nuclear exosome targeting, transcription-dependent tethering of RNA polymerase II gene DNA at the nuclear periphery, positive regulation of histone H3-K4 methylation, and regulation of miRNA metabolism (Fig. 2E, J, O), suggest that our eGWAS identified upstream signaling loci that may play a role in both genetic and epigenetic regulation of transcriptional memory during reiterated drought stress.

**Fig. 2.**
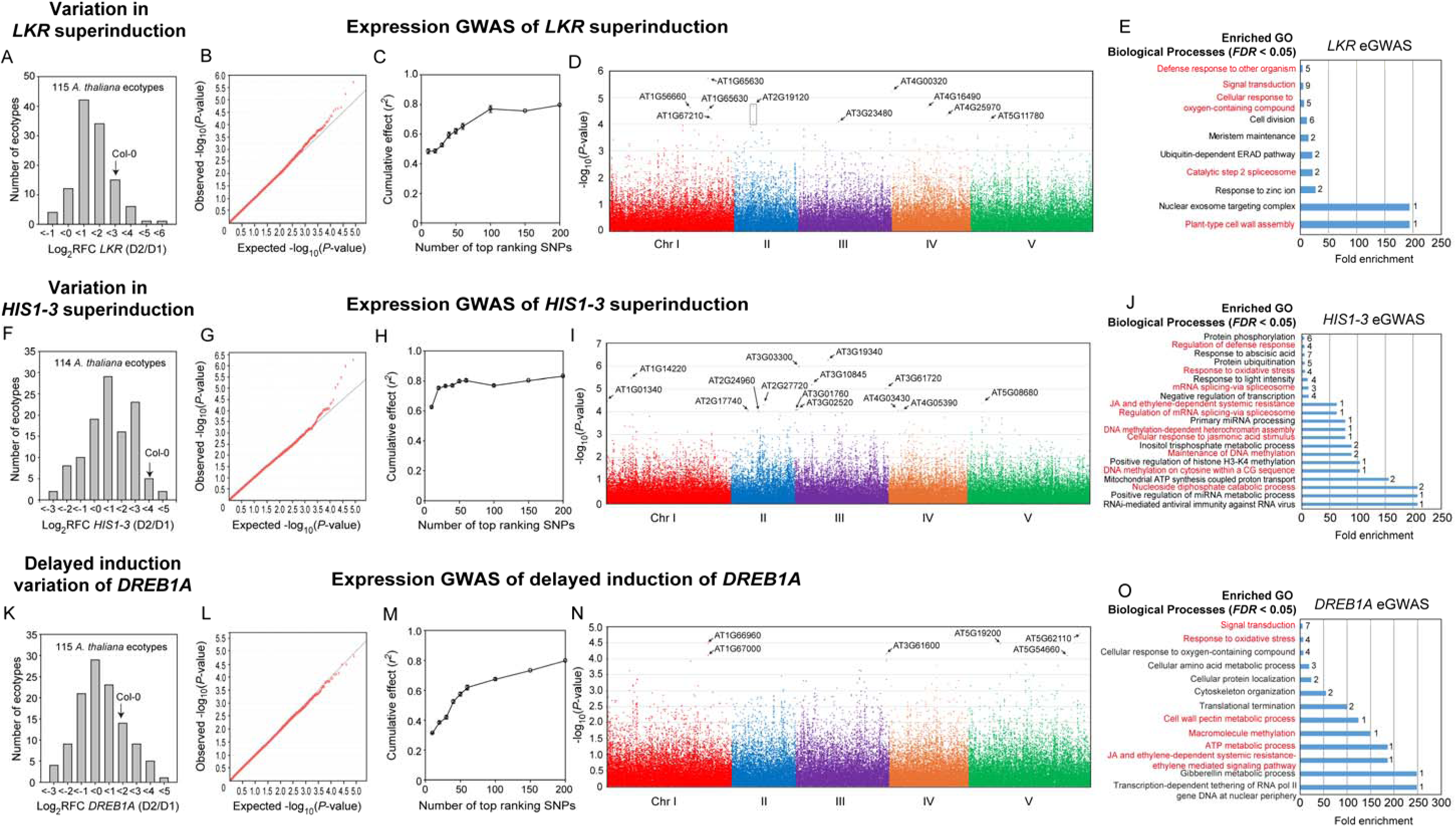
Expression genome-wide association studies of three transcriptional memory genes of *Arabidopsis thaliana* under recurrent droughts. One hundred and fifteen *A. thaliana* ecotypes were grown in ¼-Hoagland’s solution for 12 days (W), then subjected to 3 days of drought stress in ¼-Hoagland’s solution supplemented with 5% PEG 6000 (D1), followed by a recovery period of 3 days in ¼-Hoagland’s solution and again treated with PEG for 3 days (D2). Seedlings were harvested at D1 and D2, followed by quantitative PCR for the selected candidate memory genes *HIS1-3*, *LKR*, and *DREB1A*. Fold changes in gene expression were normalized with the expression levels of *AtUBQ1* from three biological replicates, each with three technical replicates. An expression GWAS (eGWAS) was performed using the D2/D1 ratio of expression levels for each of the three memory genes as the phenotype, with a genome-wide number of 105,856 SNPs, applying a mixed linear model (MLM) that accounted for kinship and population structure. Histogram plots of the relative fold change (RFC) distribution of *LKR* (A), *HIS1-3* (**F)**, and *DREB1A* (**K**) in 115 *A. thaliana* ecotypes are shown. The arrow indicates the reference genotype Col-0. Quantile-quantile (Q-Q) plots for *LKR* (B), *HIS1-3* (**G)**, and *DREB1A* (**L**) show the observed versus expected *P*-value distribution of SNPs in the GWAS. The black straight lines indicate the null distribution of *P*-values. Ridge regression analyses for *LKR* (C), *HIS1-3* (**H)**, and *DREB1A* (**M**), representing the mean correlation coefficient (*r^2^*) between predicted and observed RFCs with standard deviation for 100 iterations plotted against the number of top-ranking SNPs. Results of eGWAS are presented as Manhattan plots, depicting the significance of association (log_10_*P*-value) in relation to physical locations on five *A. thaliana* chromosomes for *LKR* (D), *HIS1-3* (**I)**, and *DREB1A* (**N**). Arrows mark the positions of the protein-coding genes closest to the suggestive SNPs. Enriched Gene Ontology (GO) biological processes (*false discovery rate* < 0.05) of genes within haploblocks of the most significant SNPs (MLM *P* < 10^3^) are shown for *LKR* (E), *HIS1-3* (**J)**, and *DREB1A* (**O**). The numbers to the right of each bar denote the number of genes in each enriched category.

The *LKR* eGWAS detected genes involved in enriched processes, including signal transduction, cell division, ubiquitin-dependent ERAD pathway, lignin biosynthetic process, plant-type cell wall assembly, nucleoid organisation, and nuclear exosome targeting complex. The genes included a *Rab5-interacting family protein* (AT2G29020), involved in endocytosis, *regulator of chromosome condensation 1* (*RCC1*; AT3G1543), responsible for nuclear transport and mitotic spindle formation, and AT5G04020 encoding a calmodulin-binding protein (Ebina et al. 2015; Reddy et al. 2002). Among the cell division-related proteins, AT2G24970 codes for spindle/kinetochore-associated protein (SKA2), involved in cell division. AT5G16210 encoding Trans-Golgi network (TGN)-associated protein 1 (TGNAP1) that acts as a scaffold/tether between the TGN and the microtubule cytoskeleton and is required for both (stress-responsive) secretory and endocytic traffic homeostasis (Renna et al. 2018). The ubiquitin-dependent ERAD pathway removes misfolded proteins from the ER, involving the eGWAS-identified gene *SAY1* (AT4G11740) (Christian et al. 2012). *Pinoresinol reductase 2* (*PRR2*; AT4G13660) encodes a NADPH-dependent enzyme critical for lignan biosynthesis, reducing (-)-pinoresinol to (-)-lariciresinol, a key step in the lignan pathway which contributes to drought tolerance mainly through cell wall reinforcement and antioxidant defense (Nakatsubo et al. 2008). *Galacturonosyltransferase-like 5* (*GATL5*; AT1G02720) encodes a key enzyme in pectin biosynthesis (Kong et al. 2013), helping plants tolerate drought by tuning cell wall flexibility, hydration, and hydraulic conductivity.

Enriched processes among the *HIS1-3* eGWAS-detected genes include response to ABA, negative regulation of transcription, nucleoside diphosphate catabolic process, mitochondrial ATP synthesis coupled proton transport, and maintenance of DNA methylation. Seven genes were ABA responsive; among them, *global transcription factor group E 11* (*GTE11*; AT3G01770) mediates ABA responses, while *thioglucoside glucohydrolase 1* (*TGG1*; AT5G26000), a myrosinase involved in glucosinolate breakdown, contributes to stomatal closure in guard cells (Qian et al. 2024; Islam et al. 2009). Four genes encoded transcriptional repressors, including *ringlet 1* (*RLT1*; AT1G28420), which modulates nucleosome spacing for target gene expression, *damaged DNA-binding protein 1A* (*DDB1A*; AT4G05420), a core component of an E3 ubiquitin ligase complex targeting misfolded proteins for degradation, and *increased level of polyploidy1-1D* (*ILP1*; AT5G08550), a repressor of CYCLINA2 that controls endoreduplication (Yoshizumi et al. 2006). *Variant in methylation 5* (*VIM5*; AT1G57800) is implicated in CpG methylation-dependent transcriptional regulation, whereas *dicer-like 2* (*DCL2*; AT3G03300) cleaves double-stranded RNA precursors, contributing to miRNA maturation and RNA interference mechanisms (Kraft et al. 2008; You et al. 2019).

The *DREB1A* eGWAS detected seven genes classified under the enriched process, signal transduction; four under response to oxidative stress; two genes each under translational termination and cytoskeletal reorganisation; and one gene each under transcription-dependent tethering of RNA polymerase II gene DNA at the nuclear periphery and macromolecule methylation. Among oxidative stress-responsive genes, *phytocystatin 6* (*CYS6*; AT3G12490) encodes a cysteine protease inhibitor that stabilizes the NADPH oxidase RBOHD, and its overexpression confers high tolerance to drought and salt (Tan et al. 2017). AT1G12920 and AT1G62850, which encode eukaryotic release factors that recognise stop codons in mRNAs during protein synthesis (Zhou et al. 2009). Eukaryotic release factors may contribute to transcriptional memory of drought by modulating translation termination, mRNA stability, and nonsense mediated decay, thereby influencing the persistence of stress responsive transcripts and associated chromatin states that underlie faster and stronger re induction of drought memory genes upon recurring dehydration. *Nucleoporin 155* (*NUP155*; AT1G14850), which encodes a major component of the nuclear pore complex (NPC), is involved in the enriched process: transcription-dependent tethering of RNA polymerase II gene DNA to the nuclear periphery (Kleine et al. 2012). The “trainable” drought memory genes remain associated with RNA polymerase II that is stalled at the promoter (with Ser5 phosphorylated Pol II and high H3K4me3) even during recovery between two droughts, which poises them for rapid reactivation during the second drought. *NUP155*’s role in tethering Pol II transcribed genes to the periphery suggests that similar NPC associated gene positioning might help “bookmark” drought responsive genes, keeping them in a nuclear environment that is permissive for rapid transcriptional re initiation during recurring drought, and might be a key determinant of transcriptional memory (Ding et al. 2012). *Apex1-like protein* (*APE1L*; AT3G48425) encodes a DNA lyase involved in DNA demethylation. APE1L processes 3’-blocking ends, such as 3’-phosphor-α,β-unsaturated aldehyde (3’-PUA) and 3’-phosphate groups, generated by ROS1 glycosylase/lyase during base excision repair. It converts these to 3’-OH termini via phosphodiesterase and phosphatase activities, enabling DNA polymerase and ligase to complete demethylation and fill gaps. Mutants like *ape1l-1* exhibit altered methylation at thousands of genomic loci, confirming its necessity (Li et al. 2015).

The results of eGWAS point to the key upstream processes regulating the transcriptional memory under recurrent droughts. We also observed that some of the eGWAS-delineated genes, e.g., AT3G48020 showed [−/+], AT5G19190 [+/−], and AT1G14240 and AT5G23575 [+/+] themselves display memory behavior, at a lower fold-change cutoff of expression (−0.6 > log_2_FC > 0.6, *P* < 0.05) (Table S6).

### 3.3 EGWAS-identified genes show polymorphisms associated with transcriptional memory

We analyzed the phenotypic associations of dense polymorphisms in CDSs and upstream regulatory regions of genes with the top 10 eGWAS-detected SNPs for 81 of 120 ecotypes available in the 1001 genome database using LAS. Among the *LKR* eGWAS-detected genes, two genes had 5’-UTR and promoter polymorphism significant in GLM and MLM, while four genes showed moderate-impact amino acid substitutions, and four genes showed high-impact polymorphisms, like stop codon gains or losses, frameshift variants, or splice acceptor variants (Table S7). Two ecotypes from the low-induction group (LIG) showed stop codon gains at *Chr1:24406598* and *Chr1:24407370* in the coding regions of *DEG3* (AT1G65630). In *HIS1-3* eGWAS-detected genes, four genes showed 5’-UTR variants, ten genes showed promoter variants significant in GLM or MLM or both, eight genes showed moderate-impact amino acid polymorphisms, and six genes showed high-impact drastic polymorphisms (Table S8). Stop codon gain variants at *Chr3:3397443*, *Chr3:3397147,* and frameshift variants at *Chr3:3396135* corresponded to lower D2/D1 induction levels of the gene AT3G10845. In *DREB1A* eGWAS, six genes showed 5’-UTR variants, ten genes showed promoter variants, eight genes showed amino acid substitutions, and five genes showed high-impact drastic polymorphisms (Table S9). The stop codon gain polymorphism at *Chr1:25006285*, and the frameshift at *Chr1:25006272* in gene AT1G67000 were found to be highly associated with low D2/D1 induction levels. We analyzed expression-level polymorphisms in genes with promoter polymorphisms significantly associated with the D2/D1 induction phenotype in the LAS (Tables S7, S8, and S9) in a representative set of 20 ecotypes by RT-qPCR (Figs S5, S6, and S7) using primers listed in Table S10. Altogether, we identified various polymorphisms in the coding and regulatory regions of eGWAS-delineated genes that are significantly associated with variation in D2/D1 induction among *A. thaliana* ecotypes.

### 3.4 Memory gene induction is abolished or attenuated during the second drought in T-DNA insertion mutants of eGWAS-delineated genes

We assessed expression of memory genes *LKR*, *HIS1-3*, and *DREB1A* in T-DNA insertion knockout/knockdown lines of genes closest to the top 10 eGWAS SNPs (MLM *P* < 10^−4^) under control (C), mild drought (D1), and severe drought (D2) conditions. The wild type (WT) Col-0 showed an average D1/C induction of 1.5 ± 0.5-fold and a D2/D1 induction of 2.8 ± 0.1-fold in the expression levels of *LKR*. The mutant lines of the genes AT2G19120, AT1G56660, AT3G23480, and *DEG3* (AT1G65630), in which T-DNA was inserted in the exons, i.e., *at2g19120* (SALK_032807C), *at1g56660* (SAIL_163_D09), *at3g23480* (SALK_073503C) and *deg3* (WiscDsLoxHs138_01E) showed 0.8, 0.9, 0.3 and 0.9-fold decrease in the D2/D1 induction of the genes, respectively, compared to WT. All lines with T-DNA inserted into the promoter, UTR, or introns were screened for gene expression levels to confirm gene knockdown by RT-qPCR using primers listed in Table S10. A knockdown line of the gene AT4G16490, *at4g16490* (SALK_121505C), having 0.5-fold suppression in gene expression due to T-DNA insertion in the 5’-UTR (Fig. S1, S4), showed a 0.9-fold decrease in D2/D1 induction of *LKR* compared to WT. On the other hand, except *at1g56660*, other mutants did not show any difference in D1/C induction compared to the WT (Fig. 3).

**Fig. 3.**
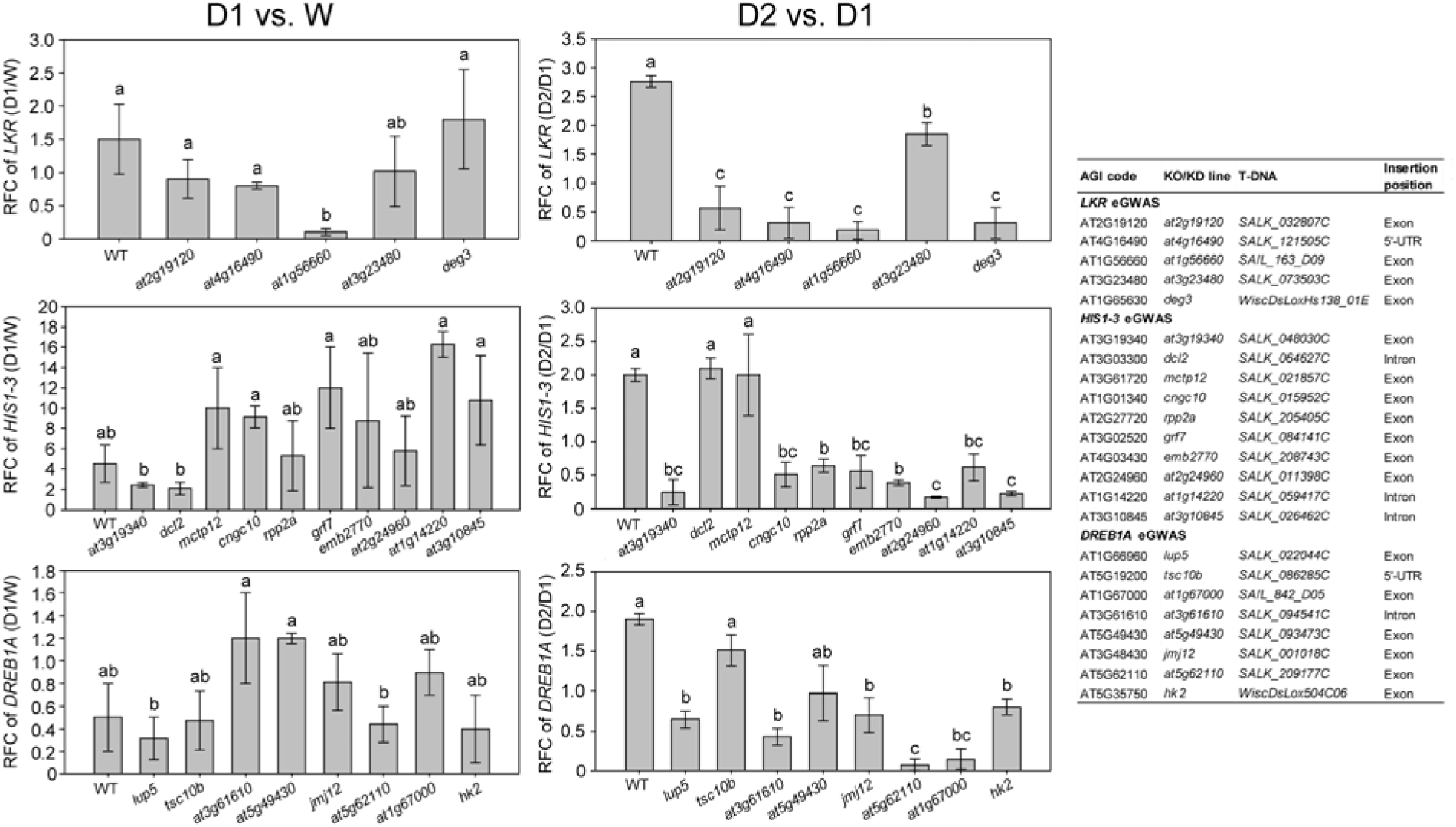
Reverse genetic validation of eGWAS-delineated genes. The genes nearest to the top 10 most-significant single-nucleotide polymorphisms (SNPs) in the *LKR*, *HIS1-3*, and DREB1A eGWASs were selected for reverse genetics analysis, and homozygous T-DNA insertion lines were isolated (see Fig. S1-3). The expression levels of the drought memory genes LKR, HIS1-3, and DREB1A were assessed in watered (W), first drought (D1), and second drought (D2) conditions. The relative fold change (RFC) of gene expression in D2 vs. D1 is shown as the average for three independent homozygous lines, with standard error. Different letters above the bars indicate significant differences (*P* < 0.05; Tukey’s HSD test; *N* = 3). The details of the T-DNA insertion lines are tabulated, including gene names and the positions of T-DNA insertions. The T-DNA insertions at the exon are designated as knockout (KO) while those at the intron, promoter, and 5’-UTR are designated as knockdown (KD), based on expression levels (see Fig. S4).

The WT displayed an average D1/C induction of *HIS1-3* of 4.5 ± 1.8-fold and a D2/D1 induction of 2.0 ± 0.5-fold. T-DNA mutants of the genes AT3G19340, *CNGC10* (AT1G01340), *RPP2A* (AT2G27720), *GRF7* (AT3G02520), *EMB2770/STA1* (AT4G03430) and AT2G24960, i.e., *at3g19340* (SALK_048030C), *cngc10* (SALK_015952C), *rpp2a* (SALK_205405C), *grf7* (SALK_084141C), *emb2770* (SALK_208743C) and *at2g24960* (SALK_011398C), having T-DNA insertions in exons, showed a 0.9, 0.7, 0.7, 0.7, 0.8 and 0.9-fold decrease in the D2/D1 induction, respectively. Knock-down lines *at1g14220* (SALK_059417C) and *at3g10845* (SALK_026462C), which have intronic T-DNA insertions (Fig. 3, S2) and exhibit 0.9-fold expression suppression (Fig. S4), showed 0.7-and 0.9-fold decreases in D2/D1 induction of *HIS1-3* compared to the WT. The mutants of *DCL2* (AT3G03300), and *MCTP12* (AT3G61720), i.e., *dcl2* (SALK_064627C) and *mctp12* (SALK_021857C) showed similar D2/D1 induction of *HIS1-3* compared to the WT. On the other hand, all mutant lines except *at3g19340* showed D1/C induction similar to that of the WT.

A 0.5 ± 0.3-fold D1/C and 1.9 ± 0.1-fold D2/D1 induction of *DREB1A* was recorded in the WT. Compared to that, mutants of the genes *LUP5* (AT1G66960), AT1G67000, *JMJ12* (AT3G48430) AT5G62110, and *HK2* (AT5G35750), i.e., *lup5* (SALK_022044C), *at1g*67000 (SAIL_842_D05), *jmj12* (SALK_001018C), *at5g62110* (SALK_209177C) and *hk2* (WiscDsLox504C06), with exonic T-DNA insertions, showed a 0.7, 0.8, 0.6, 0.96, 0.9 and 0.6-fold decrease in D2/D1 induction of *DREB1A*, respectively. Alongside that, knockdown line of AT3G61610, *at3g61610* (SALK_094541C) with T-DNA in the intron (causing 0.5-fold suppression) (Fig. S3, S4) showed a 0.5-fold decrease in the D2/D1 induction of *DREB1A*. The mutant of AT5G49430, *at5g49430* (SALK_093473C), showed no significant change in D2/D1 induction. No significant suppression of D1/C induction of *DREB1A* was observed in any of the mutants (Fig. 3). These results clearly indicate that while most mutants of eGWAS-detected genes did not affect D1/C induction, they abolished or attenuated the D2/D1 induction of memory genes, demonstrating their role in regulating transcriptional drought memory.

### 3.5 Differential methylation of genes underlie the transcriptional memory of recurrent droughts

We selected four drought-tolerant *A. thaliana* ecotypes that also exhibited high D2/D1 induction for bisulfite sequencing. These were designated as the high induction group (HIG: Na-1, Kin-0, Li-2-1, and Col-0). In contrast, we selected four drought-sensitive ecotypes with low D2/D1 induction, designated as the low induction group (LIG: KNO-18, Ru3.1-31, Sq-8, and Pla-3) (Fig. 4A, 4B). We analyzed differentially methylated regions (DMRs) in HIG versus LIG (Table S11, Fig. 4C) and in D2 versus D1 (Table S12, Fig. S8) across CpG, CHG, and CHH contexts. DMRs were identified using a methylation difference (MD) cutoff of 25% and *q*-values < 0.01 (Fig. 4C). Regions with MD > 25 (*q* < 0.01) in HIG vs. LIG were designated as hyper-DMRs, while those with MD < −25 (*q* < 0.01) were designated as hypo-DMRs (in both D1 and D2). DMRs were found in gene bodies as well as promoters of protein-coding genes, intergenic regions, pseudogenes, novel transcribed regions, transposable elements, long non-coding RNAs, antisense lncRNAs, antisense RNAs, miRNAs, pre-tRNAs, rRNAs, snRNAs, and snoRNAs (Fig. 4D, 4E). In D1, seven genes were identified as differentially methylated between HIG and LIG, whereas in D2, 1,321 genes were identified, indicating greater methylation variation during the second drought (Table S11). Similarly, six genes were differentially methylated between D2 and D1 in HIG, whereas 208 genes were differentially methylated between D2 and D1 in LIG, indicating greater methylation in sensitive ecotypes during D2 (Table S12). Among the differentially methylated genes, several showed transcriptional memory under recurrent droughts. Among these, *IGMT5* (AT1G76790) showed [+/+], the ribosomal protein *EL41X* (AT3G08520) [+/−], RmlC-like cupins superfamily protein (AT5G39120), *responsive to desiccation 20* (*RD20*; AT2G33380), AT4G12900, *alpha-glucan phosphorylase 2* (*PHS2*; AT3G46970), *delta1-pyrroline-5-carboxylate synthase 1* (*P5CS1*; AT2G39800), and *isoamylase 3* (*ISA3*; AT4G09020) [−/+] displayed transcriptional drought memory behavior in Col-0 (−0.6 > log_2_FC > 0.6, *P* < 0.05) (Table S11; Fig. 4H). We tested the expression of the top hyper- and hypo-methylated genes in HIG and LIG by RT-qPCR and observed that, in most cases, differential methylation was reflected in higher relative D2/D1 gene expression in HIG (tolerant) ecotypes (Fig. 4I), indicating an upstream regulatory role of the differentially methylated genes in the superinduction of memory genes.

**Fig. 4.**
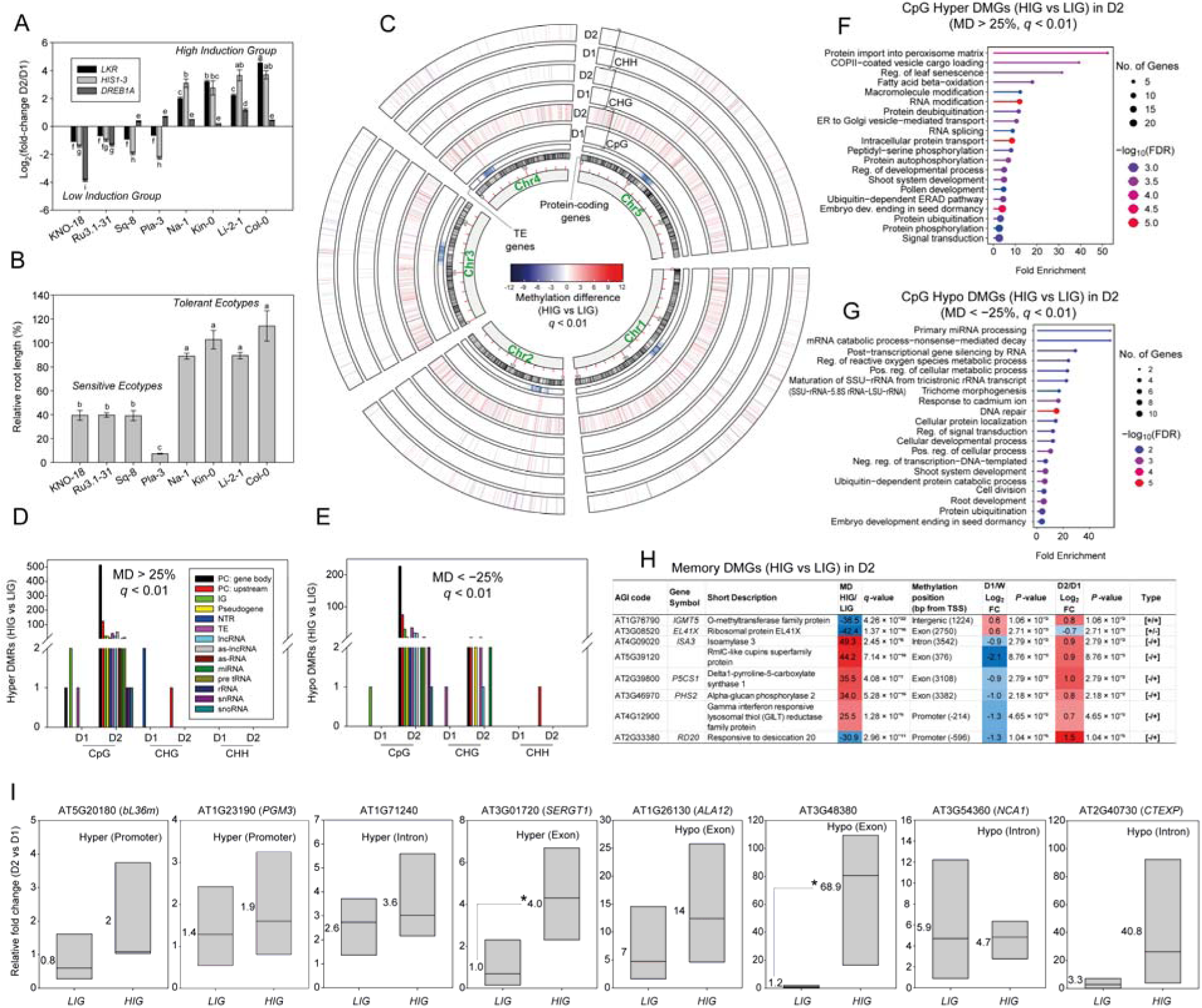
DNA methylation levels in high- and low-induction *Arabidopsis thaliana* ecotypes under recurrent droughts. (**A)** Four ecotypes having high fold-change of expression of the drought-memory genes: *LKR*, *HIS1-3*, and *DREB1A* in D2 vs. D1 (Na-1, Kin-0, Li-2-1, and Col-0), and four ecotypes with low D2/D1 expression (KNO-18, Ru3.1-31, Sq-8, and Pla-3) were selected and designated as the high induction group (HIG) and low induction group (LIG), respectively. Genome-wide methylation patterns were identified in these ecotypes at D1 and D2 by whole-genome bisulfite sequencing. (**B)** The drought tolerance phenotypes of the hydroponically grown *A. thaliana* ecotypes from the HIG and LIG are shown as the ratio of the root length in ¼-Hoagland’s solution with 2.5% PEG-6000 to the root length in the control ¼-Hoagland’s solution, i.e., relative root length (RRL; %) in a 14-day growth experiment (Marik et al. 2025). The average root lengths of 20 plants are displayed. (**C)** The Circos plot illustrates the methylation difference (MD; %) between HIG and LIG in both D1 and D2, across CpG, CHG, and CHH sequence contexts, with a *q*-value cutoff of < 0.01 (*N* = 4). The greatest difference in methylation is observed between HIG and LIG in D2. The locations of the protein-coding genes and transposable element (TE) genes on the five chromosomes of *A. thaliana* are indicated in the plot. (**D)** Regions having MD > 25 (*q* < 0.01) in HIG vs. LIG are designated as hyper-differentially methylated regions (hyper DMRs), and their numbers are presented separately for D1 and D2. The count of DMRs on gene bodies or upstream promoter regions of protein coding (PC) genes, intergenic sequences (IGs), pseudogenes, novel transcribed regions (NTRs), TEs, long non-coding RNAs (lncRNAs), antisense lncRNAs (as-lncRNAs), antisense RNAs (as-RNAs), miRNAs, pre tRNAs, rRNAs, small nuclear RNAs (snRNAs) and small nucleolar RNAs (snoRNAs) are shown separately (**E)** Regions having MD < 25 (*q* < 0.01) in HIG vs. LIG are designated as hypo-differentially methylated regions (hypo DMRs), and their numbers are presented similarly. Gene Ontology (GO) enrichment analysis of hyper-differentially methylated genes (hyper DMGs) between HIG vs. LIG (MD > 25; *q* < 0.01) in D2 **(F)** and hypo-differentially methylated genes (hypo DMGs) between HIG vs. LIG (MD < 25; *q* < 0.01) in D2 **(G)** are shown as lollipop plots. The circle size indicates the number of genes, and the circle color indicates –log_10_*FDR*. **(H)** The DMGs in D2 showing [+/+], [+/], and [/+] memory behavior (log_2_fold change > 0.6 or < 0.6; *P* < 0.05) in the Col-0 transcriptome data (Fig. 1) (memory DMGs) are presented along with gene symbols, short descriptions, MD (HIG vs. LIG), *q*-values, methylation positions, and expression levels in D1 vs. W, and D2 vs. D1. **(I)** Expression analysis of selected DMGs by RT-qPCR. The D2/D1 induction levels in the LIG and HIG ecotypes are shown as box plots. Each data point comprises the average of three biological replicates. *AtUBQ1* was used as an internal control. Hyper or hypomethylation status in HIG versus LIG within the gene body or promoter is indicated above the plots.

The genes hyper- or hypomethylated between the HIG and LIG ecotypes under D2 (regardless of genomic position) were subjected to functional enrichment analysis, revealing enrichment in stress signaling categories (Fig. 4F, 4G). Among the 873 hypermethylated genes (higher methylation levels in HIG than LIG), 23 were classified under signal transduction, which included *shaggy-like kinase 1* (*GSK1*; AT1G06390), a 14-3-3 protein *general regulatory factor 11* (*GRF11*; AT1G34760), *casein kinase 1-like protein 2* (*CKL2*; AT1G72710) that governs actin filament reorganization and stomatal closure, and *transmembrane kinase 1* (*TMK1*; AT1G66150) that controls apoplastic acidification and cell expansion through the phosphorylation of plasma membrane H^+^-ATPase (Zhao et al. 2016; Shang et al. 2024; Moscatiello, 2026). Another enriched category centered on protein phosphorylation, featuring genes AT1G04700, AT1G53430, and AT4G03230 classified as serine threonine kinases, *pre-mRNA processing factor 4 kinase A* (*PRP4KA*; AT3G25840) associated with alternative splicing, a leucine rich repeat receptor kinase, *BRI1-like 2* (*BRL2*; AT2G01950), linked with brassinosteroid signaling, and *MAP3K 2* (AT3G07980), a mitogen activated protein kinase kinase kinase that functions upstream in MAPK cascades and implicated in abiotic stress signaling, where it likely contributes to drought responses by modulating stress induced phosphorylation events that regulate stomatal closure, ROS homeostasis, and expression of drought responsive genes (Mitra et al. 2015; Kanno et al. 2018). DNA repair is a major outcome of drought stress and represents another enriched functional category including genes such as *DNA ligase 6* (*LIG6*; AT1G66730), that participates in single stranded break repair, *ataxia-telangiectasia mutated* (*ATM*; AT3G48190), related to telomere repair and maintenance, *BRCA1/BRCA2-containing complex 36 homolog A* (*BRCC36A*; AT1G80210), a deubiquitinating enzyme involved in homologous recombination and DNA repair, and *replication factor C4* (*RFC4*; AT1G21690), a crucial component of the DNA replication machinery (Aklilu et al. 2013; Sailer et al. 2018). The gene *cell division cycle 48A* (*CDC48A*; AT3G09840), which encodes an AAA-type ATPase, is involved in cell division and ERAD (Field et al. 2021). Other enriched processes include coat-protein complex II (COPII)-coated vesicle cargo loading, RNA splicing, and ubiquitin-dependent ERAD pathway. Among these, the Ubiquitin fusion degradation 1 (UFD1) family protein (AT2G29070) functions in ERAD, while *VPS25* (AT4G19003) and *VPS22* (AT4G27040) play roles in the ubiquitin-dependent sorting of plasma membrane proteins into multivesicular bodies (MVBs) for degradation in vacuoles (Richardson et al. 2011; Ucar et al. 2024). Additionally, *SEC23* (AT1G05520) and *SEC24* (AT4G32640), essential components of the COPII machinery, are involved in the transport of secretory proteins from the ER to the Golgi apparatus, and *SEC11* (AT1G12360) is related to vesicle trafficking (Karnik et al. 2015; Zeng et al. 2015).

Among the 364 hypomethylated genes (lower methylation in HIG than LIG), enriched categories include signal transduction, response to ABA and water deprivation, cell division, vesicle trafficking, gene silencing, DNA demethylation, etc. A total of twelve genes in signal transduction were identified including *casein kinase 1-like protein 2* (*CKL2*; AT1G72710), *RAF20* (AT1G79570), *CBL-interacting protein kinase* 3 (CIPK3; AT2G26980), and *G2484-1* (AT4G17330) which are involved in actin organization, SnRK2 activation, stomatal regulation, cell wall dynamics and root elongation under drought (Zhao et al. 2016; Chen et al. 2021a; Wang et al. 2021). Eleven hypomethylated genes were ABA-responsive, including *smaller with variable branches* (*SVB*; AT1G56580), *jumonji 21* (*JMJ21*; AT1G78280), *ABA hypersensitive 1* (*ABH1*; AT2G13540), and *RD20*, indicating the role of ABA signaling in upstream regulation of drought memory (Chen et al. 2015; Xiong et al. 2001). Among the nine genes categorized under DNA repair, *actin-related protein 5* (*ARP5*; AT3G12380) plays essential roles in DNA double-strand break repair, *zinc 4 finger DNA 3’-phosphoesterase* (*ZDP*; AT3G14890) participates in BER repair, *INO80* (AT3G57300) is a chromatin remodeling ATPase crucially involved in homologous recombination and genome stability maintenance, and AT5G63920 encodes a DNA topoisomerase TOP3α that relaxes supercoiled DNA (Roldán-Arjona et al. 2019; Fan et al. 2024). Out of the seven cell division category genes, *UBP16* (AT4G24560) acts as a ubiquitin-specific protease involved in salt tolerance, *cleavage stimulation factor 77* (*CSTF77*; AT1G17760) functions in pre-mRNA 3 -end polyadenylation machinery, and *ovate family protein 14* (*OFP14*; AT1G79960) is a transcription repressor (Feng et al. 2016; Zeng et al. 2018). In the gene silencing category, *silencing defective 3* (*SDE3*; AT1G05460) eliminates secondary structures in RNA-dependent RNA polymerase target transcripts, alongside *nuclear RNA polymerase D1B* (*NRDP1B*; AT2G40030), which functions in RdDM and siRNA biogenesis (Pontier et al. 2005; Linder and Owttrim 2009). The epigenetic regulators include the DNA demethylase *BRAT1-partner1* (*BRP1*; AT3G15120) and histone deacetylases *HD2C* (AT5G03740) and *HDA5* (AT5G61060) (Zhang et al. 2016; Chen et al. 2020).

## 4. Discussion

Our eGWAS study, accompanied by methylome profiling, dissected upstream signaling circuitry regulating drought memory genes, viz., *LKR*, *HIS1-3*, and *DREB1A*. The reverse genetic screen of eGWAS, on the other hand, showed clear attenuation/abolition of D2/D1 induction, while D1/C induction remained almost unchanged, providing evidence of regulatory activity at the second drought. Knockout/knockdown of LKR eGWAS-identified genes abolished D2/D1 induction of *LKR*, likely via the following mechanisms: disruption of AT2G19120 (a P-loop NTPase with a helicase domain) interferes with replication fork resolution and DNA damage checkpoints (Jia et al. 2016); disruption of AT3G23480 (cyclopropane-fatty-acyl-phospholipid synthase) rigidifies membranes, hindering ion homeostasis signals; disruption of AT4G16490 (an Armadillo repeat superfamily protein) may alter memory via chromatin-level control (Pieczynski et al. 2017); and disruption of mitochondrial *DEG3* impairs protein quality control, leading to accumulation of misfolded stress sensors that attenuate transcriptional memory (Ludwig et al. 2025). For *HIS1-3* eGWAS candidates, knocking out *CNGC10* blocks Ca² waves essential for *HIS1-3* activation; disruption of *RPP2A* and *GRF7* interferes with translation and 14 3 3 mediated coupling of kinase signalling to downstream transcriptional responses, respectively; and knocking out the splicing factor *STA1*/*EMB2770* prevents maturation of stress-responsive mRNAs, abolishing *HIS1-3* superinduction (Lee et al. 2006). Knockout of *DREB1A* eGWAS hits has analogous effects: loss of *LUP5* disrupts biosynthesis of triterpenoid alcohols required for membrane protection and reactive oxygen quenching (Shibuya et al. 2009); loss of the bromodomain transcription factor AT5G49430 disrupts acetyl histone–dependent recruitment or stabilisation of the transcriptional machinery at the *DREB1A* locus (Abiraami et al. 2023); loss of *JMJ12* causes retention of repressive H3K27me3 marks in *DREB1A* regulatory regions (Kim et al. 2025); loss of AT5G62110 likely disrupts imprinting linked chromatin control (Wolff et al. 2011); and knockdown of the cytokinin receptor *HK2* weakens cytokinin signalling, disturbing ABA–cytokinin balance (Bartrina et al. 2017) and thereby impairing *DREB1A* transcriptional priming during D2.

Bisulfite sequencing revealed the whole genome methylation status of Arabidopsis in D1 and D2 across ecotypes contrasting in drought tolerance, as well as D2/D1 induction, in relation to transcriptional memory behavior. It showed the largest methylation differences between the drought tolerant/HIG and drought sensitive/LIG ecotypes in D2 rather than D1, indicating stronger epigenetic regulatory activity during the second drought. Maximum D2/D1 differential methylation was also observed in sensitive ecotypes compared with tolerant ones, suggesting that sensitive lines undergo more extensive and unstable epigenome reprogramming under recurrent drought, a pattern also reported in rice varieties (Wang et al. 2016). As reported in the literature, exon methylation is generally associated with higher gene expression, whereas promoter or intron methylation often represses transcription. Gene body methylation epialleles have also been linked to environmental adaptation (Shahzad et al. 2025). Exon hypermethylation can either increase or decrease gene expression, depending on its precise location (e.g., first versus internal exons), gene context, and interactions with other regulatory features, such as promoters, transposable elements, and splicing signals. This is consistent with our RT qPCR validation of the methylome, in which both hypermethylation and hypomethylation were associated with higher expression in HIG versus LIG (Fig. 4I). In addition, we observed numerous methylation events in transposable elements, which typically induce silencing, confer genomic stability, and influence the expression of nearby genes (Shahzad et al. 2025). In D2, hypermethylation in HIG/tolerant exotypes of *GSK1* (intron) implies downregulation and enhanced BR signalling (Yan et al. 2009), whereas exon hypermethylation of *GRF11* and *BRL2* stabilizes BR receptors and likely increases BR signalling (Muyle et al. 2022; Li et al. 2024). Promoter hypermethylation of *CKL2* may delay stomatal closure by limiting ADF4 phosphorylation (Shi et al. 2022), while exon CpG hypermethylation of *RAF16* (AT1G04700) and *PRP4KA* (AT3G25840) suggests enhanced kinase and splicing activity, respectively, promoting drought tolerance (Fàbregas et al. 2020; Wang et al. 2022). DNA repair genes AT1G66730 and AT1G21690 show exon CpG hypermethylation consistent with upregulation, whereas hypomethylation of AT3G48190 and AT1G12360 points to increased expression in vesicle and SNARE mediated transport. Exon hypermethylation of *VPS25* contrasts with promoter silencing like effects at *VPS22*, indicating destabilization of the endosomal sorting complex (ESCRT II)–dependent sorting, while intron hypermethylation of Sec23/Sec24 transport family protein (AT4G14160) implies repression of COPII mediated vesicle coat assembly during drought in tolerant ecotypes.

In parallel, hypomethylation in HIG/tolerant exotypes at D2 highlights epigenetic activation of stress adaptive pathways: *RAF20* and *CIPK3* hypomethylation supports SnRK2 activation and Ca² –ABA signalling (Soma et al. 2020); and hypomethylation of *JMJ30* (AT1G78280), *RD20*, *ARP5*, *ZDP*, *INO80*, *TOP3*α, *UBP16*, *CstF77*, *BRP1*, and *HD2C* suggests coordinated modulation of histone demethylation, ABA responsive transcription, genome stability, mRNA processing, chromatin accessibility, and repression dynamics during stress recovery (Roldán Arjona et al. 2019; Fan et al. 2024; Vogel et al. 2024). Highly CpG hypermethylated genes in D2 include *bL36M* (AT5G20180, translational repression), *ATG1* (AT1G23190, autophagy; Cheng et al. 2025), *SMR5* (AT1G07500, cell cycle arrest; Braat et al. 2023), *CGLD11* (AT2G21385, photosynthesis; Grahl et al. 2016), *JMJ32* (AT3G45880, flowering; Gan et al. 2014), and *LIG6* (AT1G66730, error prone double-strand break repair; Waterworth et al. 2025), reflecting trade offs between growth, photosynthesis, and DNA repair. Conversely, CpG hypomethylation of *CTEXP* (AT2G40730, tRNA export; Johnstone et al. 2009) likely enhances translation, underscoring how epigenetic rewiring in D2 finetunes both stress protection and developmental trade offs.

Bisulfite sequencing revealed that sensitive ecotypes show extensive hypomethylation in 99 genes in D2 (Fig. S8; Table S12). The enriched categories in hypomethylated genes included regulation of ADP ribosylation factor (ARF) protein signal transduction, along with vesicle-mediated transport and epigenetic regulation (Fig. S8C). ARF GTPases relay signals inside the cell by switching between active (GTP bound) and inactive (GDP bound) states, thereby modulating downstream pathways. This suggests the potential to increase cellular responsiveness but also reflects a less stable, more labile epigenetic state under drought in sensitive ecotypes. Regulation of ARF protein signal transduction can influence drought responses by modulating membrane trafficking, hormone signaling, and cytoskeletal dynamics that underlie stomatal closure, root growth, and stress induced reorganization of cellular compartments. The 106 hypermethylated genes in the sensitive ecotypes were involved in the enriched processes of negative regulation of post-embryonic development, negative regulation of gene expression, proteasome-mediated ubiquitin-dependent protein catabolic process, PSII-associated light harvesting II catabolic process, ABA transport, and heterochromatin assembly (Fig. S8C). Hypermethylation of genes regulating development, gene expression, protein turnover, photosynthesis, ABA transport, and heterochromatin in sensitive ecotypes likely impairs their ability to balance growth and stress responses, reducing drought resilience and memory gene induction.

Methylome-eGWAS overlaps of candidate genes (Fig. 5A; Table S13) and enriched biological processes (Fig. 5B) provided epigenetic mechanistic depth. The enrichment of signal transduction among both hyper and hypomethylated genes, as well as DREB1A eGWAS hits, suggests that DNA methylation and natural genetic variation jointly target upstream signaling components to fine tune *DREB1A* induction under recurrent drought. *LKR* eGWAS detected hypermethylated ERAD and cell division genes (*UFD1*; AT2G29070 and *ABCC2*; AT2G34660), where hypermethylation or silencing retains ubiquitinated aquaporins (e.g., PIP2;1), elevating cellular water loss and eroding osmoprotectant memory (Chen et al. 2021b). On the other hand, hypomethylated translation factors (AT5G04090, AT5G16200) shared by *LKR* eGWAS hits may boost protein synthesis, thereby sustaining LKR priming. *LKR*/*HIS1-3* eGWAS common genes like Myb/SANT-like DNA-binding domain protein (AT2G24960; knockout validated; Fig. 3), a transcription elongation factor *Mediator 26A* (*MED26*; AT3G10820) bridging transcription factors and POL II (Guo et al. 2021), and an RNA-binding protein AT3G10845 (knockout validated; Fig. 3) together shape the transcription of both memory genes *LKR* and *HIS1-3*. AT3G01780 (TPLATE), detected by *HIS1-3* eGWAS as well as hypermethylated in HIG/tolerant ecotypes, regulates clathrin mediated endocytosis, and it indirectly modulates drought signaling by controlling membrane trafficking and endocytosis of ABA/other phytohormone receptors (Gadeyne et al. 2014). Another overlapping gene, *FATA* (AT3G25110), regulates oleic acid production, thereby influencing membrane integrity and signaling under drought. Hypomethylated *HIS1-3* hits (*STA1* and *PRP16*; AT5G13010) could enhance splicing, and miRNA processing of ABA-responsive transcripts (*HSFA3*, *COR15A*, *IDD14*), promoting superinduction of *HIS1-3* (Kim et al. 2017; Tsugeki et al. 2014). *DREB1A* eGWAS/hypermethylated overlaps (AT5G49430, AT5G23575, AT5G01950, and *ERF1,2*; AT1G12920) could suppress translation/release factors, curtailing *DREB1A* expression during recurrence. These overlaps align with the signaling roles of memory genes (Ding et al. 2013), revealing eGWAS/methylome convergence on DNA repair, epigenetics, and proteostasis as core memory hubs. This is also evident from co-expression and protein-protein interaction network connectivity between eGWAS and methylome-identified genes (Fig. 5C). Combining genes unravelled by both approaches yielded a big complex network (i) comprising genes involved in the enriched processes of RNA and DNA metabolism, post-transcriptional gene silencing by RNA, ubiquitin-dependent protein catabolic process, and chromatin organization, and a smaller network (ii) of genes involved in localization and vesicle-mediated transport (Fig. 5C).

**Fig. 5.**
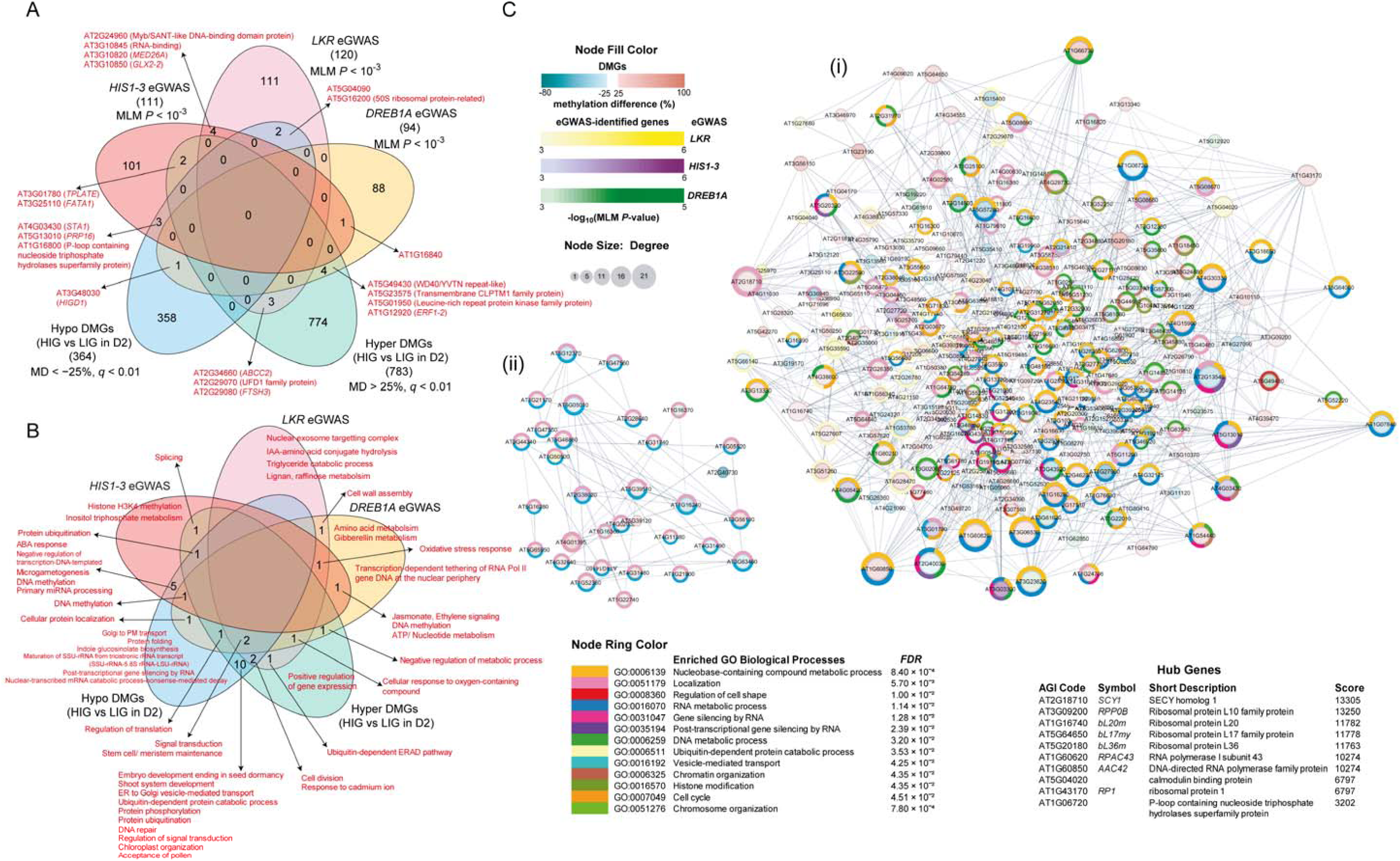
Network analysis of genes delineated by eGWAS and methylome analysis. (**A**) A Venn diagram illustrates the common genes between eGWAS and differential methylome analysis using whole-genome bisulfite sequencing. (**B**) A Venn diagram illustrates the common enriched GO biological processes identified between eGWAS and methylome analyses. (**C**) Network between genes in haploblocks of single-nucleotide polymorphisms (SNPs) with a mixed-linear model (MLM) *P*-value < 10^3^ identified by the eGWAS and the genes detected by bisulfite sequencing between HIG and LIG with MD cutoff > 25 and < 25 created through the STRING web server, with a high confidence score (0.7). Circles represent nodes, while lines represent the network’s edges. The size of the node indicates the degree of relatedness of the genes. The eGWAS-identified genes are color-coded (node fill color) in yellow, purple, and green, while the % methylation difference is shown in color shades of red and blue, indicating hyper- and hypo-methylation, respectively. Node ring colors indicate enriched Gene Ontology (GO) biological processes, as illustrated in the table with their corresponding *false discovery rate* (*FDR*) values.

In conclusion, by integrating eGWAS and differential methylome profiling, this study successfully identified a complex network of upstream signaling and epigenetic regulators that specifically modulate the transcriptional memory of recurrent drought in *A. thaliana*. Functional validation via T-DNA mutants confirmed that these identified loci primarily attenuate gene induction during repeated drought episodes rather than the initial stress, demonstrating their specialized role in memory maintenance and the rapid reactivation of poised stress-responsive genes. The results of the study could serve as a starting point for exploring the specific mechanisms by which the identified genes regulate drought memory. For example, how nuclear pore complexes, through components like NUP155, facilitate the “bookmarking” of drought genes at the nuclear periphery to maintain an environment permissive for rapid transcriptional re-initiation. Future work should dissect how candidate regulators directly engage the promoters of key memory genes and shape their chromatin environment during D1 versus D2. Promoter–reporter assays in WT and selected mutants (e.g., *at5g49430*, *jmj12*, *at5g62110*, *hk2*, *cngc10*, *emb2770*) under single and recurrent drought would clarify which factors control promoter responsiveness and D2/D1 superinduction. Complementarily, ChIP qPCR/ChIP seq for histone marks and for key regulators themselves (e.g., AT5G49430, *JMJ12*, *BRP1*, *HD2C*) at the memory genes across W/D1/D2 in tolerant versus sensitive ecotypes would test whether stable transcriptional memory correlates with a distinct primed promoter chromatin signature. Together, these approaches would directly link upstream chromatin readers, writers, and erasers to the establishment, maintenance, and erasure of transcriptional drought memory.

## Statements and declarations

### CRediT authorship contribution statement

DM: Investigation (all wet experiments), Visualization, Writing – Original Draft Preparation. RK: Investigation (network, ridge regression, and circos plots), Software, Visualization. AS: Conceptualization, Data Curation, Formal analysis, Visualization, Funding acquisition, Methodology, Resources (wet lab), Project Administration, Writing – Original Draft Preparation, Writing – Review and Editing.

### Data availability

All data are presented within the paper. The raw transcriptome data are submitted to the NCBI Sequence Read Archive with the accession number PRJNA1086208, and the whole-genome bisulfite sequencing raw data are submitted with the accession number PRJNA1147722.

