## Supplementary data for "Expression genome-wide association study and differential methylome profiling reveal upstream regulators of drought memory genes in *Arabidopsis thaliana*"

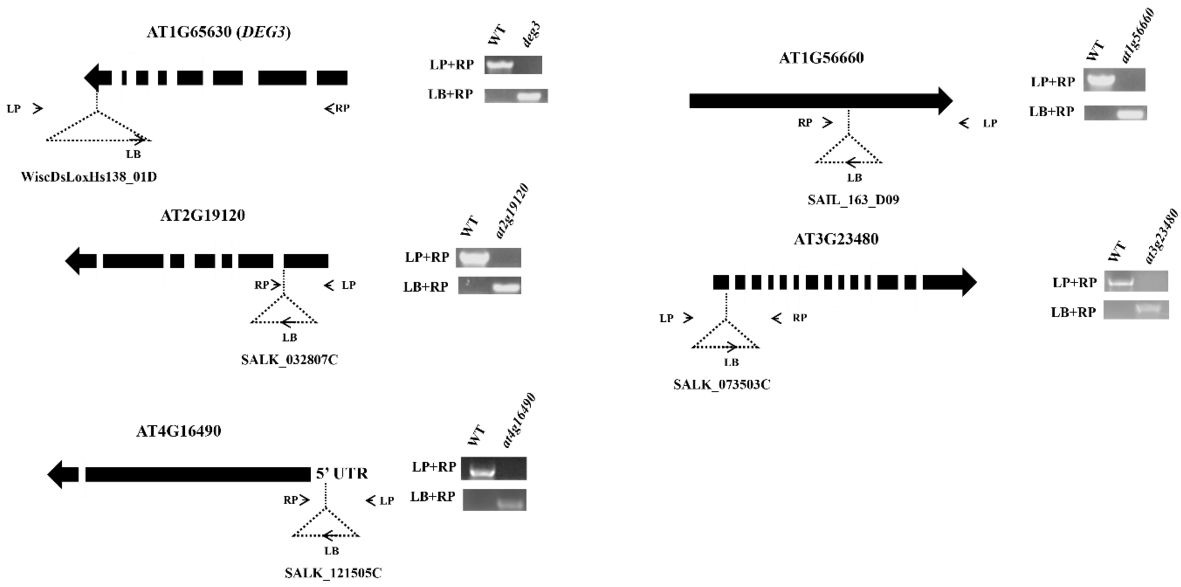


**Fig. S1 Validation of homozygosity of the TDNA insertion mutants used in *LKR* eGWAS**. TDNA insertion mutants used in the study, viz., *deg3* (WiscDsLoxHs138_01D), AT2G19120 (SALK_032807C), AT1G56660 (SAIL_163_D09), AT3G23480 (SALK_073503C), and AT4G16490 (SALK_121505C), were tested for homozygosity by a PCR-based method. The primer list is available in Table S10. The position of the inserted T-DNA is represented by dotted triangles, with the T-DNA left border (LB), forward genomic primer (LP), and reverse genomic primer (RP) labelled, along with gel images of the amplification. Gene coding sequences are denoted by thick black arrows in the 5' to 3' direction, where intermediate gaps indicate introns.

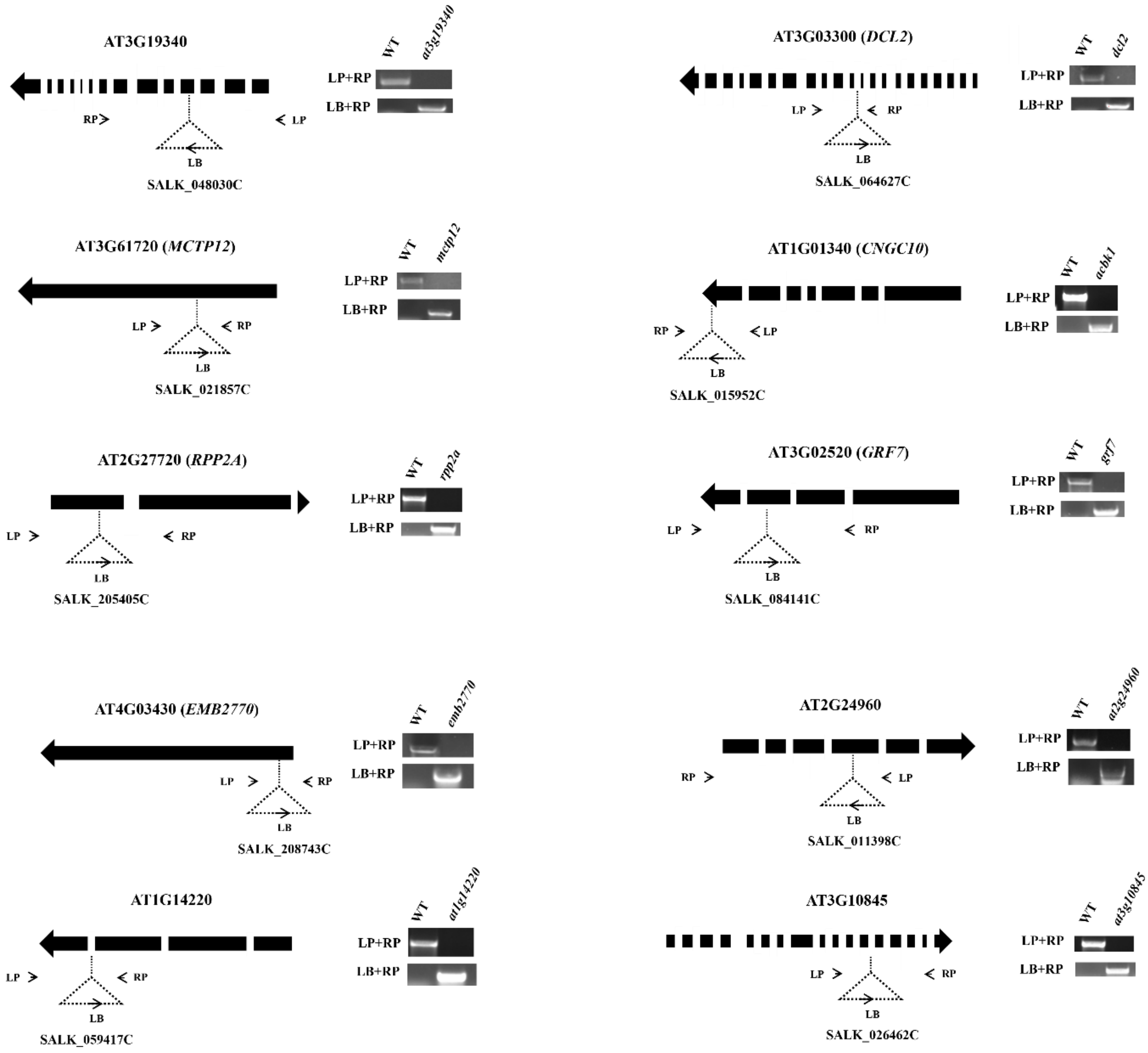


**Fig. S2 Validation of homozygosity of the TDNA insertion mutants used in *HIS1-3* eGWAS**. TDNA insertion mutants used in the study, viz., AT3G19340 (SALK_048030C), *dcl2* (SALK_064627C), *mctp12* (SALK_021857C), *cngc10* (SALK_015952C), *rpp2a* (SALK_205405C), *grf7* (SALK_084141C), *emb2770* (SALK_208743C), AT2G24960 (SALK_011398C), AT1G14220 (SALK_059417C), AT3G10845 (SALK_026462C), were tested for homozygosity by a PCR-based method. The primer list is available in Table S10. The position of the T-DNA insertion is represented by dotted triangles, with the T-DNA left border (LB), forward genomic primer (LP), and reverse genomic primer (RP) labelled, along with gel images of the amplification products. Gene coding sequences are denoted by thick black arrows in the 5' to 3' direction, where intermediate gaps indicate introns.

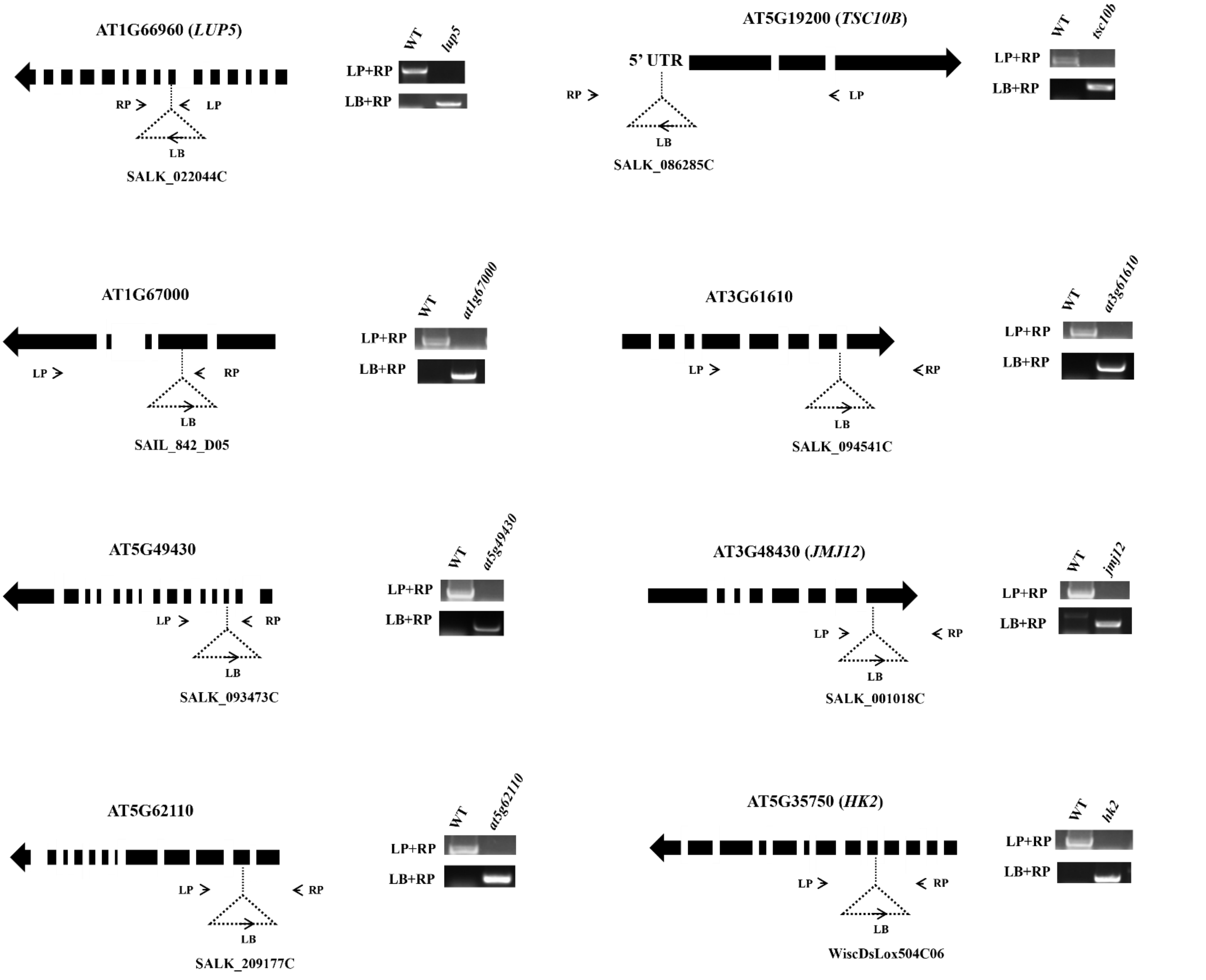


**Fig. S3 Validation of homozygosity of the TDNA insertion mutants used in *DREB1A* eGWAS**. TDNA insertion mutants used in the study, viz., *lup5* (SALK_022044C), *tsc10b* (SALK_086285C), AT1G67000 (SAIL_842_D05), AT3G61610 (SALK_094541C), AT5G49430 (SALK_093473C), *jmj12* (SALK_001018C), AT5G62110 (SALK_209177C), and *hk2* (WiscDsLox504C06) were tested for homozygosity by a PCR-based method. The primer list is available in Table S10. The position of the T-DNA insertion is represented by dotted triangles, with the T-DNA left border (LB), forward genomic primer (LP), and reverse genomic primer (RP) labelled, along with gel images of the amplification products. Gene coding sequences are denoted by thick black arrows in the 5' to 3' direction, where intermediate gaps indicate introns.

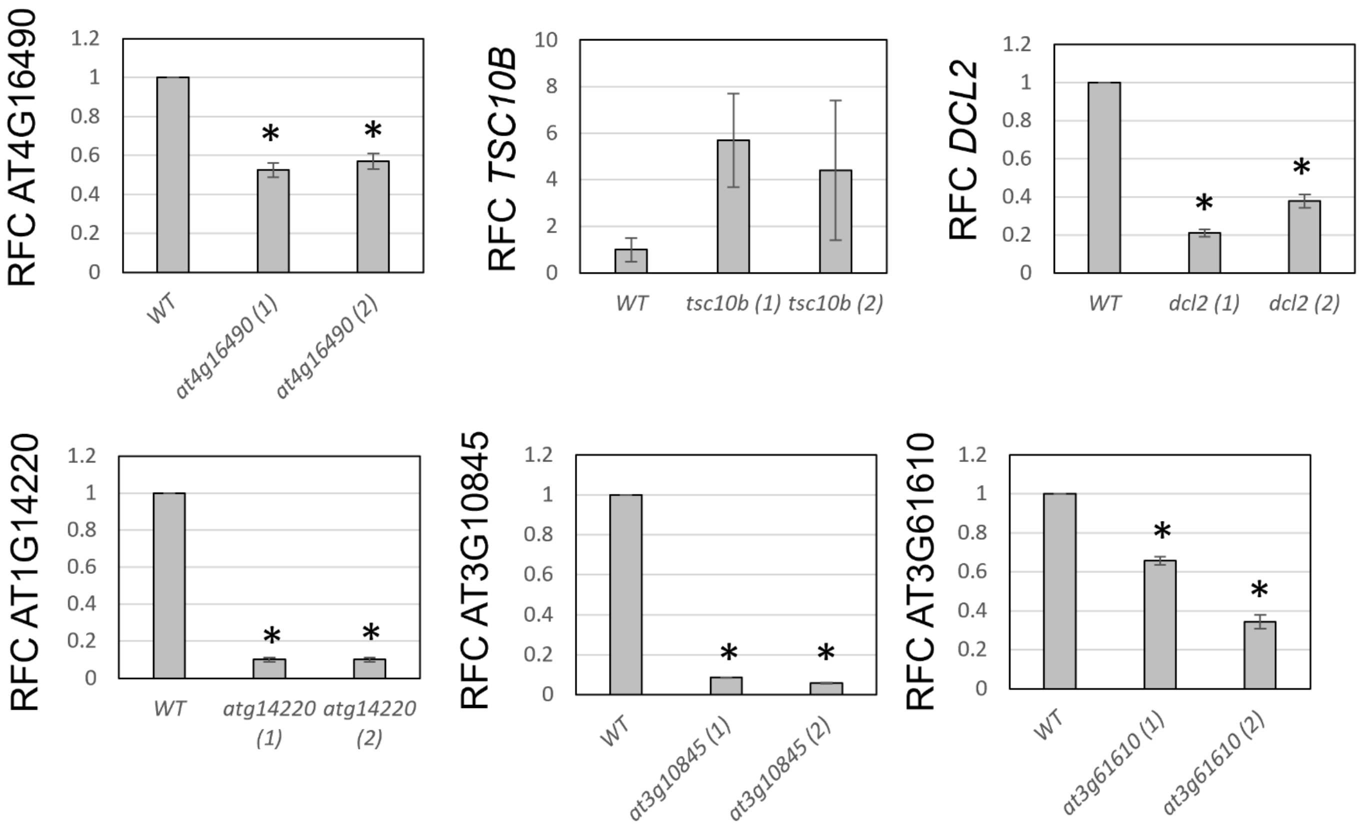


**Fig. S4 Gene expression levels of mutants with T-DNA insertion in the promoter, 5’-UTR and introns**. T-DNA insertion events in the promoter or 5’ UTR were tested for expression level reduction. The confirmed homozygous mutants: *at4g16490* (SALK_121505C), *tsc10b* (SALK_086285C), *dcl2* (SALK_064627C), *at1g14220* (SALK_059417C), *at3g10845* (SALK_026462C), and *at3g61610* (SALK_094541C) were grown in ¼-Hoagland’s media for 12 days from germination, followed by 3 days of drought stress in ¼-Hoagland’s media supplemented with 5% PEG-6000. The seedlings were harvested, followed by RNA isolation and qPCR analysis. Fold change in expression levels of two independent knockdown lines with respect to the wild type (WT) is represented. *AtUBQ1* was used as an internal control for normalization of expression levels, and fold change in expression was calculated using the $2^{-\Delta\Delta Ct}$ method. Asterisks indicate significant differences between WT and knockdown lines (Student’s *t*-test, *N* = 3 biological replicates).

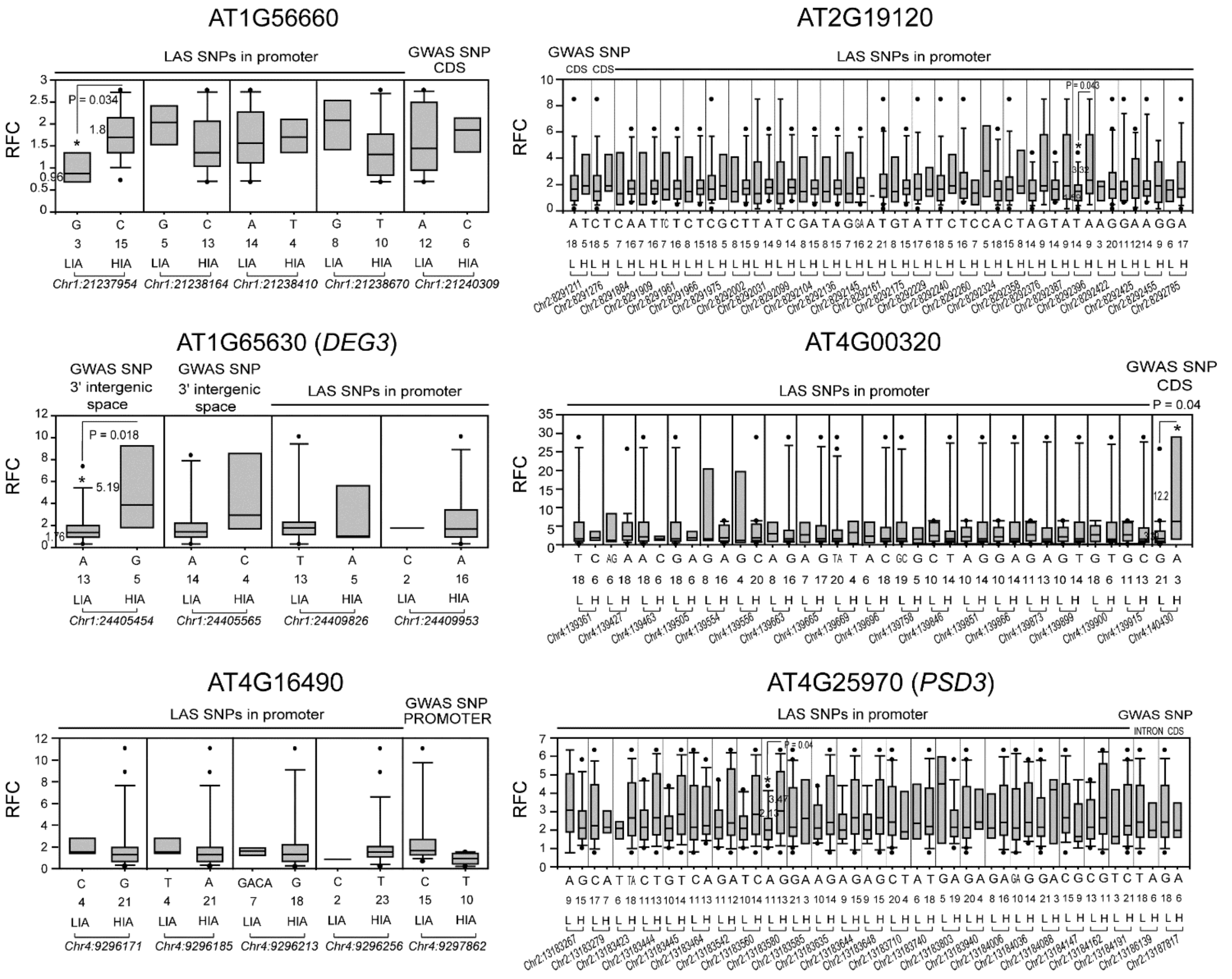


**Fig. S5. Expression level polymorphisms in *LKR* eGWAS-delineated genes.** The expression level variations of genes with -log$_{10}$ GWAS *p-value* > 4 in the genome-wide association study (eGWAS) were selected for ELP analysis. The genes with significantly associated promoter polymorphisms (GLM/MLM) in a local association study (LAS) conducted in randomly selected ecotypes among the 120 ecotypes used in the eGWAS are shown as box plots. Twenty plants of each ecotype were cultivated in ¼ Hoagland's solution for 12 days, then subjected to drought stress using ¼ Hoagland's media supplemented with 5% PEG-6000 (D1) for 3 days, followed by recovery period (R1) for 3 days in ¼ Hoagland's solution and another 3 days of drought stress in ¼ Hoagland's media supplemented with 5% PEG-6000 (D2). Expression levels in whole seedlings in D2 vs D1 were determined by quantitative real-time PCR and normalized to AtUBQ1 (AT3G52590). The expression levels were further normalized relative to the Columbia (Col-0) ecotype using the $2^{-\Delta\Delta Ct}$ method. An average of three technical replicates is presented. On the X-axis of each plot, ecotypes are grouped by alleles at the respective SNP loci, listed below as chromosome numbers and physical positions in base pairs. "N" indicates the number of ecotypes containing the SNP allele. The asterisks indicate significant differences (Student's *t*-test; $P < 0.05$). At significant positions, the numbers to the left of each box indicate the average relative fold change for ecotypes containing the SNP allele. LIA/L: low induction allele, HIA/H: high induction allele.

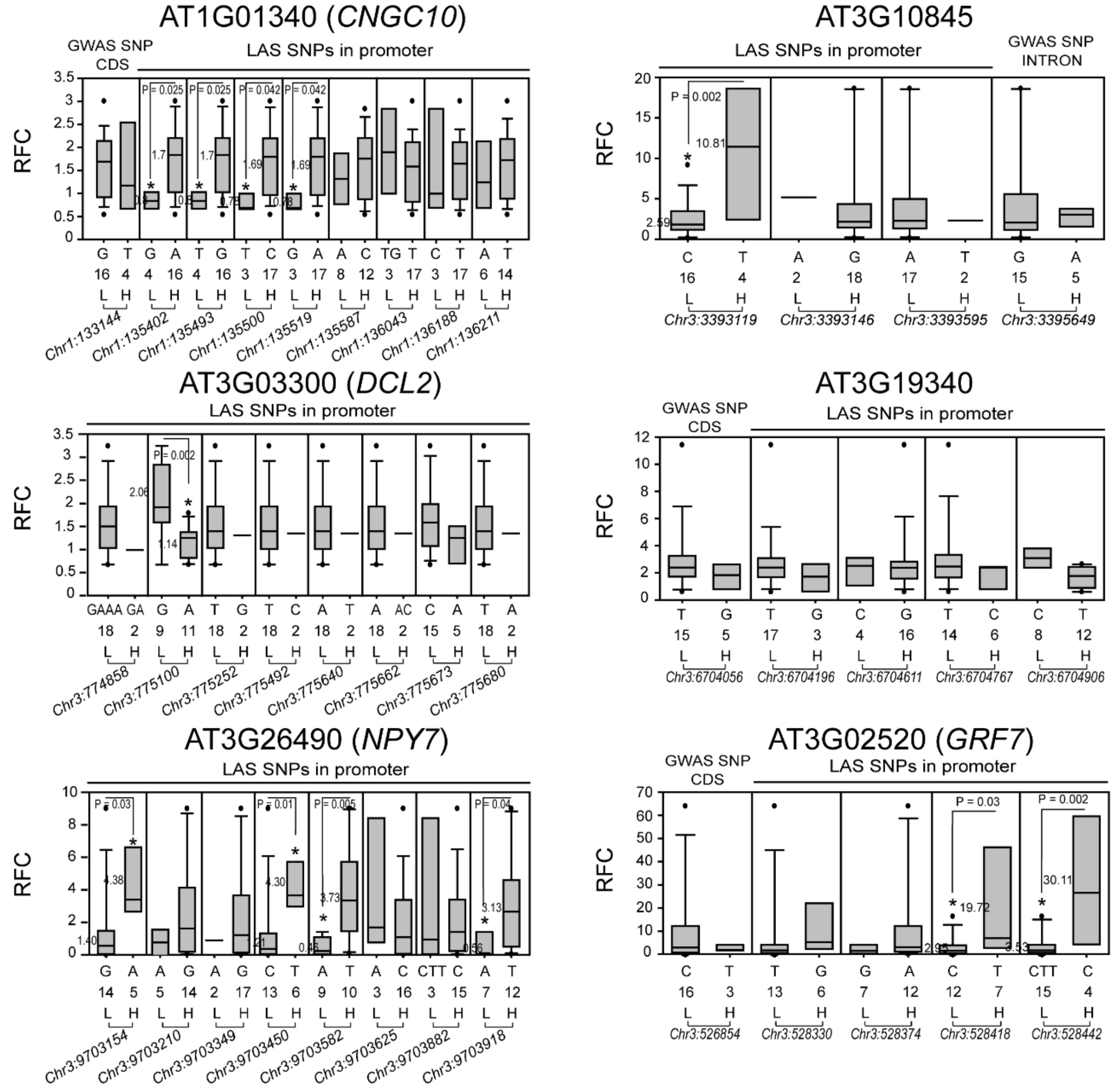


**Fig. S6. Expression level polymorphisms in *HIS1-3* eGWAS-delineated genes.** The expression level variations of genes with -$\log_{10}$ GWAS *p-value*>4 in the genome-wide association study (eGWAS) were selected for ELP analysis. The genes with significantly associated promoter polymorphisms (GLM/MLM) in a local association study (LAS) conducted in randomly selected ecotypes among the 120 ecotypes used in the eGWAS are shown as box plots. Twenty plants of each ecotype were cultivated in ¼ Hoagland's solution for 12 days, then subjected to drought stress using ¼ Hoagland's media supplemented with 5% PEG-6000 (D1) for 3 days, followed by recovery period (R1) for 3 days in ¼ Hoagland's solution and another 3 days of drought stress in ¼ Hoagland's media supplemented with 5% PEG-6000 (D2). Expression levels in whole seedlings in D2 vs D1 were determined by quantitative real-time PCR and normalized to AtUBQ1 (AT3G52590). The expression levels were further normalized relative to the Columbia (Col-0) ecotype using the $2^{-\Delta\Delta Ct}$ method. An average of three technical replicates is presented. On the X-axis of each plot, ecotypes are grouped by alleles at the respective SNP loci, listed below as chromosome numbers and physical positions in base pairs. "N" indicates the number of ecotypes containing the SNP allele. The asterisks indicate significant differences (Student's *t*-test; $P < 0.05$). At significant positions, the numbers to the left of each box indicate the average relative fold change for ecotypes containing the SNP allele. L: low induction allele, H: high induction allele

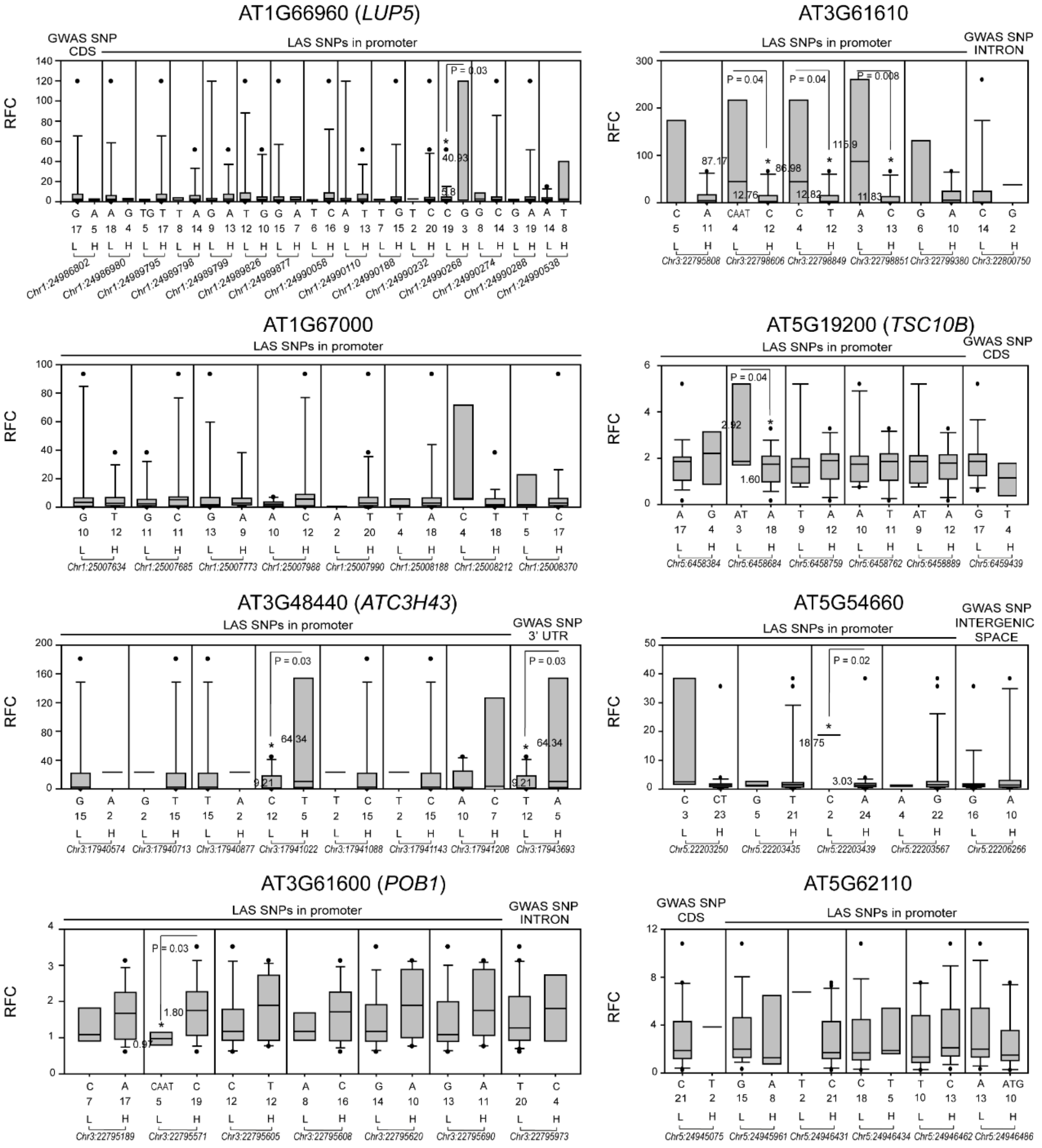


**Fig. S7. Expression level polymorphisms in *DREB1A* eGWAS-delineated genes.** The expression level variations of genes with -$\log_{10}$ GWAS *p-value*>4 in the genome-wide association study (eGWAS) were selected for ELP analysis The genes with significantly associated promoter polymorphisms (GLM/MLM) in a local association study (LAS) conducted in randomly selected ecotypes among the 120 ecotypes used in the eGWAS are shown as box plots. Twenty plants of each ecotype were cultivated in ¼ Hoagland's solution for 12 days, then subjected to drought stress using ¼ Hoagland's media supplemented with 5% PEG-6000 (D1) for 3 days, followed by recovery period (R1) for 3 days in ¼ Hoagland's solution and another 3 days of drought stress in ¼ Hoagland's media supplemented with 5% PEG-6000 (D2). Expression levels in whole seedlings for D1 vs D2 were determined by quantitative real-time PCR and normalized to *AtUBQ1* (AT3G52590). The expression levels were further normalized relative to the Columbia (Col-0) ecotype using the $2^{-\Delta\Delta Ct}$ method. An average of three technical replicates is presented. On the X-axis of each plot, ecotypes are grouped by alleles at the respective SNP loci, listed below as chromosome numbers and physical positions in base pairs. "N" indicates the number of ecotypes containing the SNP allele. The asterisks indicate significant differences (Student's *t*-test; $P < 0.05$). At significant positions, the numbers to the left of each box indicate the average relative fold change for ecotypes containing the SNP allele. L: low induction allele, H: high induction allele.

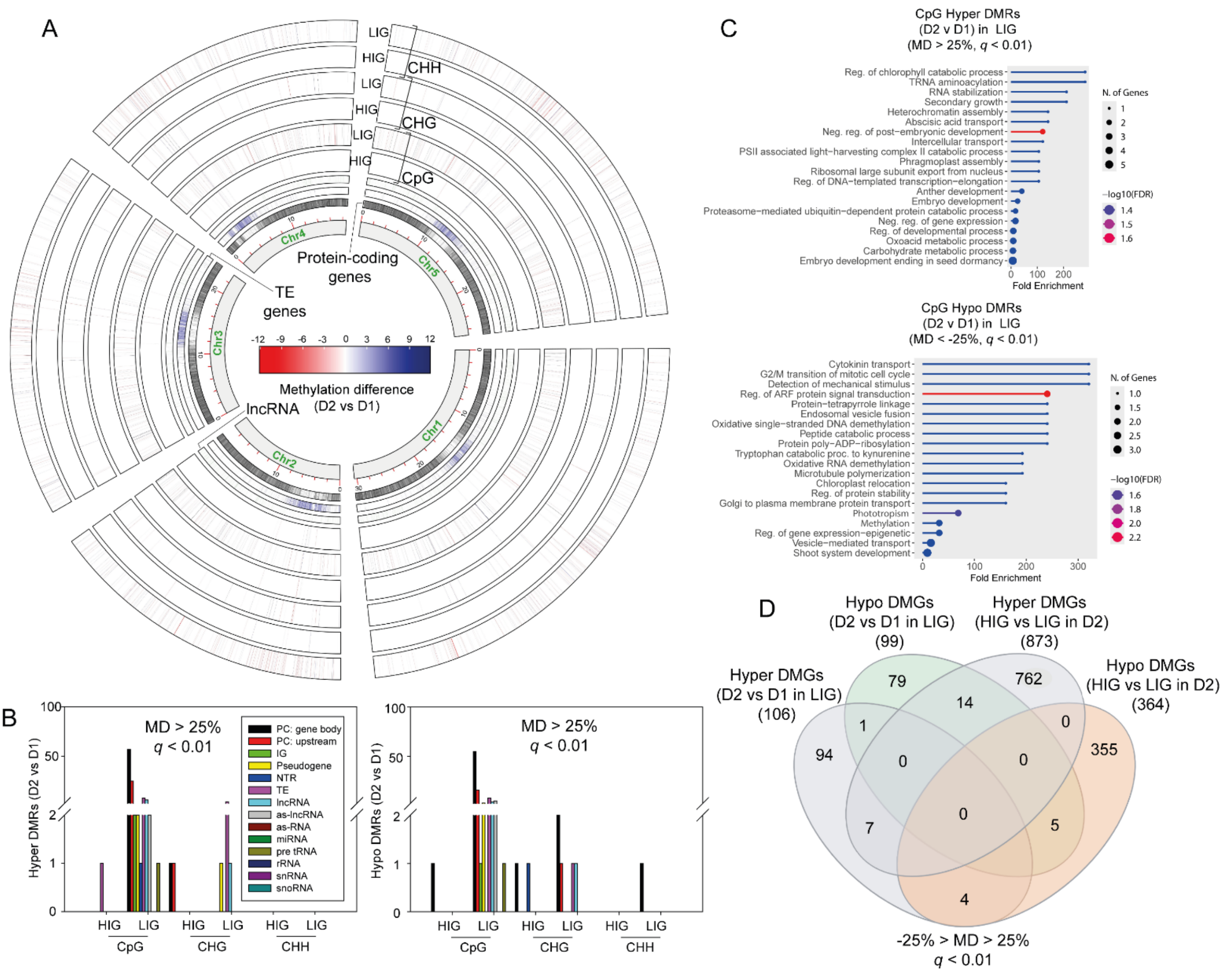


**Fig. S8. DNA methylation levels in high- and low-induction *Arabidopsis thaliana* ecotypes under recurrent droughts**. (**A**) The Circos plot illustrates the methylation difference (MD; %) between the D2 and D1 in both high induction group (HIG) and low induction group (LIG) (see Fig. 4), in the CpG, CHG, and CHH sequence contexts, at a $q$-value cutoff of < 0.01 ($N = 4$). The greatest difference in methylation is observed between D2 and D1 in LIG. The locations of the protein-coding genes, transposable element (TE) genes, and long non-coding RNAs on the five chromosomes of *A. thaliana* are indicated in the plot. (**B**) Regions having MD > 25 ($q < 0.01$) in D2 vs. D1 are designated as hyper-differentially methylated regions (hyper DMRs), and their numbers are presented separately for HIG and LIG. The count of DMRs on gene bodies or upstream promoter regions of protein coding (PC) genes, intergenic sequences (IGs), pseudogenes, novel transcribed regions (NTRs), TEs, long non-coding RNAs (lncRNAs), antisense lncRNAs (as-lncRNAs), antisense RNAs (as-RNAs), miRNAs, pre tRNAs, rRNAs, small nuclear RNAs (snRNAs) and small nucleolar RNAs (snoRNAs) are shown separately Regions having MD < –25 ($q < 0.01$) in HIG vs. LIG are designated as hypo-differentially methylated regions (hypo DMRs), and their numbers are presented similarly. (**C**) Gene Ontology (GO) biological process enrichment analysis of hyper-differentially methylated genes (hyper DMGs) between D2 vs. D1 (MD > 25; $q < 0.01$) in LIG and of hypo-differentially methylated genes (hypo DMGs) between D2 vs. D1 (MD < –25; $q < 0.01$) in LIG are shown as lollipop plots. The circle size indicates the number of genes, and the circle color indicates $-\log_{10}FDR$. (**D**) A Venn diagram shows the intersections between hyper- and hypo-DMGs in the two comparisons: D2 vs D1 in LIG and HIG vs LIG in D2.

### Table S1: Transcriptome of *Arabidopsis thaliana* under recurrent droughts in hydroponics

| AGI code | Drought 1 vs Watered | | | Recovery vs Watered | | | Drought 2 vs Watered | | | Drought 2 vs Drought 1 | | | Gene symbol | Short Description | CpG methylation difference (D2/D1) | | | CHG methylation difference (D2/D1) | | | CHH methylation difference (D2/D1) | | |
|---|---|---|---|---|---|---|---|---|---|---|---|---|---|---|---|---|---|---|---|---|---|---|---|
| | $Log_2$ FC | *P*-value | *FDR* | $Log_2$ FC | *P*-value | *FDR* | $Log_2$ FC | *P*-value | *FDR* | $Log_2$ FC | *P*-value | *FDR* | | | MD | *q*-value | Methylation position | MD | *q*-value | Methylation position | MD | *q*-value | Methylation position |
| AT2G39030 | -1.80 | **$4.61\times10^{-3}$** | **$3.06\times10^{-2}$** | -1.45 | $1.45\times10^{-1}$ | $2.90\times10^{-1}$ | 2.03 | **$3.85\times10^{-4}$** | **$2.31\times10^{-3}$** | 3.82 | **$2.68\times10^{-18}$** | **$5.19\times10^{-14}$** | *NATA1* | N-acetyltransferase activity 1 | | | | | | | 9.2 | **$3.26\times10^{-3}$** | Promoter-TSS (-459) |
| AT4G15210 | -2.42 | **$1.78\times10^{-3}$** | **$1.43\times10^{-2}$** | -1.23 | $1.23\times10^{-1}$ | $2.58\times10^{-1}$ | 1.09 | **$1.13\times10^{-3}$** | **$5.83\times10^{-3}$** | 3.51 | **$2.32\times10^{-6}$** | **$1.14\times10^{-3}$** | *BAM5* | Beta-amylase 5 | | | | | | | | | |
| AT1G58320 | -3.86 | **$1.56\times10^{-10}$** | **$9.32\times10^{-9}$** | -3.20 | **$1.98\times10^{-7}$** | **$3.87\times10^{-6}$** | -0.86 | $7.34\times10^{-2}$ | $1.77\times10^{-1}$ | 2.98 | **$5.52\times10^{-6}$** | **$1.98\times10^{-3}$** | | PLAC8 family protein | | | | | | | 7.3 | **$1.07\times10^{-4}$** | Promoter-TSS (-407) |
| AT5G60250 | -0.23 | $7.05\times10^{-1}$ | $8.67\times10^{-1}$ | 0.75 | $1.26\times10^{-1}$ | $2.63\times10^{-1}$ | 2.54 | **$6.27\times10^{-14}$** | **$2.13\times10^{-12}$** | 2.74 | **$3.49\times10^{-7}$** | **$2.82\times10^{-4}$** | | Zinc finger (C3HC4-type RING finger) family protein | | | | | | | | | |
| AT4G31970 | -2.93 | **$2.44\times10^{-5}$** | **$3.98\times10^{-4}$** | -3.30 | **$1.03\times10^{-5}$** | **$1.25\times10^{-4}$** | -0.35 | $5.50\times10^{-1}$ | $7.14\times10^{-1}$ | 2.58 | **$1.55\times10^{-3}$** | $1.22\times10^{-1}$ | *CYP82C2* | Cytochrome P45, family 82, subfamily C, polypeptide 2 | | | | | | | 7.9 | **$1.72\times10^{-3}$** | Exon (801) |
| AT1G30190 | -0.60 | $1.67\times10^{-1}$ | $3.95\times10^{-1}$ | -0.34 | $4.78\times10^{-1}$ | $6.50\times10^{-1}$ | 1.98 | **$2.20\times10^{-5}$** | **$1.82\times10^{-4}$** | 2.57 | **$7.57\times10^{-6}$** | **$2.57\times10^{-3}$** | | Cotton fiber protein | | | | | | | | | |
| AT1G16850 | -0.84 | $2.62\times10^{-1}$ | $5.20\times10^{-1}$ | -1.65 | **$3.81\times10^{-3}$** | **$1.77\times10^{-2}$** | 1.71 | **$3.76\times10^{-7}$** | **$4.49\times10^{-6}$** | 2.55 | **$9.67\times10^{-4}$** | $8.89\times10^{-2}$ | | Transmembrane protein | | | | | | | | | |
| AT3G02480 | 4.14 | **$1.43\times10^{-3}$** | **$1.20\times10^{-2}$** | 2.34 | **$4.10\times10^{-2}$** | $1.14\times10^{-1}$ | 6.63 | **$1.28\times10^{-42}$** | **$1.24\times10^{-28}$** | 2.49 | **$3.84\times10^{-2}$** | $7.02\times10^{-1}$ | *ABR* | Late embryogenesis abundant protein (LEA) family protein | | | | | | | | | |
| AT1G07985 | -0.54 | $3.06\times10^{-1}$ | $5.66\times10^{-1}$ | -1.13 | $1.20\times10^{-1}$ | $2.54\times10^{-1}$ | 1.90 | **$2.87\times10^{-7}$** | **$3.50\times10^{-6}$** | 2.44 | **$4.63\times10^{-10}$** | **$1.12\times10^{-6}$** | *LIPA* | Lipid droplet plasma membrane adaptor | | | | | | | | | |
| AT2G39330 | 0.39 | $5.31\times10^{-1}$ | $7.55\times10^{-1}$ | 0.78 | $1.74\times10^{-1}$ | $3.30\times10^{-1}$ | 2.82 | **$9.89\times10^{-11}$** | **$2.21\times10^{-9}$** | 2.41 | **$3.26\times10^{-4}$** | **$4.16\times10^{-2}$** | *JAL23* | Jacalin-related lectin 23 | | | | | | | 7.3 | **$2.77\times10^{-3}$** | Intron (1207) |
| AT1G51090 | -1.35 | $7.47\times10^{-2}$ | $2.35\times10^{-1}$ | -1.81 | **$9.45\times10^{-9}$** | **$2.41\times10^{-7}$** | 1.01 | **$4.59\times10^{-3}$** | **$1.93\times10^{-2}$** | 2.37 | **$4.10\times10^{-3}$** | $2.30\times10^{-1}$ | | Heavy metal transport/detoxification superfamily protein | | | | | | | | | |
| AT4G34510 | -0.44 | $2.12\times10^{-1}$ | $4.57\times10^{-1}$ | 0.16 | $6.90\times10^{-1}$ | $8.14\times10^{-1}$ | 1.89 | **$3.22\times10^{-14}$** | **$1.13\times10^{-12}$** | 2.32 | **$1.31\times10^{-12}$** | **$6.32\times10^{-9}$** | *KCS17* | 3-ketoacyl-CoA synthase 17 | | | | | | | | | |
| AT1G64110 | 3.05 | **$1.10\times10^{-3}$** | **$9.78\times10^{-3}$** | 2.30 | **$1.83\times10^{-2}$** | $6.16\times10^{-2}$ | 5.35 | **$4.34\times10^{-23}$** | **$3.95\times10^{-21}$** | 2.30 | **$1.06\times10^{-2}$** | $3.79\times10^{-1}$ | *DAA1* | DUO1-activated ATPase 1 | | | | | | | 8.8 | **$9.74\times10^{-4}$** | Intron (332) |
| AT2G18193 | -1.40 | **$7.10\times10^{-3}$** | **$4.29\times10^{-2}$** | -0.32 | $5.51\times10^{-1}$ | $7.11\times10^{-1}$ | 0.89 | **$1.38\times10^{-3}$** | **$6.95\times10^{-3}$** | 2.27 | **$4.89\times10^{-5}$** | **$1.10\times10^{-2}$** | | P-loop containing nucleoside triphosphate hydrolases superfamily protein | | | | | | | | | |
| AT2G45570 | -1.11 | **$9.46\times10^{-3}$** | $5.32\times10^{-2}$ | -1.83 | **$2.21\times10^{-5}$** | **$2.43\times10^{-4}$** | 1.16 | **$1.09\times10^{-2}$** | **$3.96\times10^{-2}$** | 2.26 | **$8.95\times10^{-5}$** | **$1.63\times10^{-2}$** | *CYP76C2* | Cytochrome P45, family 76, subfamily C, polypeptide 2 | | | | | | | 8.2 | **$3.18\times10^{-3}$** | Exon (322) |
| AT5G15960 | -0.24 | $7.24\times10^{-1}$ | $8.78\times10^{-1}$ | -0.57 | $5.91\times10^{-1}$ | $7.43\times10^{-1}$ | 1.94 | **$3.71\times10^{-12}$** | **$1.00\times10^{-10}$** | 2.18 | **$2.11\times10^{-3}$** | $1.55\times10^{-1}$ | *KIN1* | Stress-responsive protein (KIN1) / stress-induced protein (KIN1) | | | | | | | 9.5 | **$4.51\times10^{-6}$** | Exon (134) |
| AT5G66400 | 2.08 | **$1.35\times10^{-2}$** | $6.93\times10^{-2}$ | 0.91 | $2.81\times10^{-1}$ | $4.60\times10^{-1}$ | 4.25 | **$1.98\times10^{-47}$** | **$6.82\times10^{-45}$** | 2.17 | **$1.59\times10^{-2}$** | $4.70\times10^{-1}$ | *RAB18* | Responsive to ABA 18 | 26.8 | **$3.84\times10^{-4}$** | Promoter-TSS (-104) | | | | 6.1 | **$6.74\times10^{-4}$** | Promoter-TSS (-9) |
| AT2G34655 | 1.22 | **$1.61\times10^{-2}$** | $7.88\times10^{-2}$ | 1.95 | **$2.77\times10^{-3}$** | **$1.38\times10^{-2}$** | 3.37 | **$3.75\times10^{-25}$** | **$4.16\times10^{-23}$** | 2.13 | **$1.06\times10^{-6}$** | **$6.59\times10^{-4}$** | | Hypothetical protein | | | | | | | | | |
| AT5G52310 | -1.18 | **$2.95\times10^{-2}$** | $1.24\times10^{-1}$ | -1.96 | **$1.92\times10^{-4}$** | **$1.54\times10^{-3}$** | 0.94 | **$4.12\times10^{-3}$** | **$1.76\times10^{-2}$** | 2.12 | **$1.41\times10^{-3}$** | $1.16\times10^{-1}$ | *LTI78* | Low temperature induced 78 | | | | | | | | | |
| AT1G22990 | 0.69 | $1.60\times10^{-1}$ | $3.85\times10^{-1}$ | 0.34 | $2.80\times10^{-1}$ | $4.59\times10^{-1}$ | 2.79 | **$1.60\times10^{-19}$** | **$1.02\times10^{-17}$** | 2.09 | **$1.09\times10^{-5}$** | **$3.19\times10^{-3}$** | *HIPP22* | Heavy metal associated isoprenylated plant protein 22 | | | | | | | | | |
| AT5G01300 | -0.16 | $8.61\times10^{-1}$ | $9.44\times10^{-1}$ | -0.66 | $5.34\times10^{-1}$ | $6.97\times10^{-1}$ | 1.94 | **$4.47\times10^{-10}$** | **$9.16\times10^{-9}$** | 2.09 | **$1.77\times10^{-2}$** | $4.97\times10^{-1}$ | | Phosphatidylethanolamine-binding protein | | | | | | | | | |
| AT4G02360 | -0.05 | $9.33\times10^{-1}$ | $9.75\times10^{-1}$ | -0.46 | $1.10\times10^{-1}$ | $2.38\times10^{-1}$ | 2.04 | **$1.22\times10^{-10}$** | **$2.70\times10^{-9}$** | 2.09 | **$3.93\times10^{-5}$** | **$9.16\times10^{-3}$** | *ASD3* | ABA-inducible signal peptide- containing DUF538 protein 3 | | | | | | | 11.7 | **$2.45\times10^{-3}$** | 5' UTR-TSS (65) |
| AT3G57260 | -0.58 | $3.47\times10^{-1}$ | $6.07\times10^{-1}$ | 0.50 | **$9.23\times10^{-3}$** | **$3.60\times10^{-2}$** | 1.51 | **$1.25\times10^{-12}$** | **$3.61\times10^{-11}$** | 2.08 | **$1.55\times10^{-3}$** | $1.22\times10^{-1}$ | *PR2* | Beta-1,3-glucanase 2 | | | | | | | | | |
| AT1G11070 | -1.04 | **$4.00\times10^{-2}$** | $1.53\times10^{-1}$ | -1.06 | **$4.52\times10^{-3}$** | **$2.03\times10^{-2}$** | 1.04 | **$4.52\times10^{-4}$** | **$2.66\times10^{-3}$** | 2.07 | **$4.45\times10^{-6}$** | **$1.79\times10^{-3}$** | | Hydroxyproline-rich glycoprotein family protein | | | | | | | | | |
| AT2G16005 | 4.54 | **$9.27\times10^{-10}$** | **$4.62\times10^{-8}$** | 4.45 | **$3.29\times10^{-16}$** | **$3.06\times10^{-14}$** | 6.60 | **$9.42\times10^{-54}$** | **$4.66\times10^{-51}$** | 2.02 | **$5.44\times10^{-6}$** | **$1.98\times10^{-3}$** | *ROSY1* | MD-2-related lipid recognition domain-containing protein | | | | | | | | | |
| AT2G21820 | -0.32 | $6.75\times10^{-1}$ | $8.48\times10^{-1}$ | -0.67 | $5.73\times10^{-1}$ | $7.29\times10^{-1}$ | 1.70 | **$4.58\times10^{-4}$** | **$2.69\times10^{-3}$** | 2.01 | **$6.44\times10^{-4}$** | $7.00\times10^{-2}$ | | Seed maturation protein | | | | | | | | | |
| AT1G69140 | -2.14 | **$4.01\times10^{-7}$** | **$1.08\times10^{-5}$** | -2.37 | **$9.47\times10^{-17}$** | **$9.40\times10^{-15}$** | -0.15 | $6.31\times10^{-1}$ | $7.75\times10^{-1}$ | 2.00 | **$3.46\times10^{-5}$** | **$8.36\times10^{-3}$** | | Pseudogene of hypothetical protein | | | | | | | | | |
| AT4G28140 | 0.40 | $6.45\times10^{-1}$ | $8.30\times10^{-1}$ | 0.46 | $5.26\times10^{-1}$ | $6.90\times10^{-1}$ | 2.38 | **$4.07\times10^{-5}$** | **$3.16\times10^{-4}$** | 1.99 | **$6.02\times10^{-3}$** | $2.87\times10^{-1}$ | | Integrase-type DNA-binding superfamily protein | | | | | | | | | |
| AT1G52890 | -1.06 | **$1.72\times10^{-2}$** | $8.29\times10^{-2}$ | -1.50 | **$4.13\times10^{-4}$** | **$2.90\times10^{-3}$** | 0.88 | **$1.39\times10^{-2}$** | **$4.82\times10^{-2}$** | 1.95 | **$8.72\times10^{-6}$** | **$2.86\times10^{-3}$** | *NAC19* | NAC domain containing protein 19 | | | | | | | 9.1 | **$1.19\times10^{-3}$** | TTS-intergenic space (2131) |

| | | | | | | | | | | | | | | | | | | | | |
|---|---|---|---|---|---|---|---|---|---|---|---|---|---|---|---|---|---|---|---|---|
| AT4G25480 | -1.54 | **$4.87\times10^{-6}$** | **$9.88\times10^{-5}$** | -2.07 | **$7.63\times10^{-10}$** | **$2.42\times10^{-8}$** | 0.39 | $1.79\times10^{-1}$ | $3.42\times10^{-1}$ | 1.93 | **$3.91\times10^{-7}$** | **$2.91\times10^{-4}$** | *DREB1A* | Dehydration response element B1A | | | | | | |
| AT4G02280 | 0.58 | $3.58\times10^{-1}$ | $6.17\times10^{-1}$ | -0.13 | $8.16\times10^{-1}$ | $8.98\times10^{-1}$ | 2.47 | **$4.93\times10^{-9}$** | **$8.27\times10^{-8}$** | 1.89 | **$1.03\times10^{-2}$** | $3.73\times10^{-1}$ | *SUS3* | Sucrose synthase 3 | | | | | | |
| AT1G05680 | -3.22 | **$1.47\times10^{-15}$** | **$2.56\times10^{-13}$** | -2.06 | **$9.47\times10^{-4}$** | **$5.81\times10^{-3}$** | -1.32 | **$1.16\times10^{-6}$** | **$1.26\times10^{-5}$** | 1.88 | **$8.11\times10^{-7}$** | **$5.61\times10^{-4}$** | *UGT74E2* | Uridine diphosphate glycosyltransferase 74E2 | | | | | | |
| AT2G35070 | 1.42 | $8.61\times10^{-2}$ | $2.58\times10^{-1}$ | 0.60 | $1.90\times10^{-1}$ | $3.52\times10^{-1}$ | 3.25 | **$3.50\times10^{-17}$** | **$1.76\times10^{-15}$** | 1.84 | **$2.63\times10^{-2}$** | $5.97\times10^{-1}$ | | Transmembrane protein | | | | 5.9 | **$2.35\times10^{-3}$** | Intron (597) |
| AT5G24780 | 0.79 | $7.16\times10^{-2}$ | $2.29\times10^{-1}$ | 1.05 | $7.53\times10^{-2}$ | $1.80\times10^{-1}$ | 2.64 | **$2.61\times10^{-11}$** | **$6.31\times10^{-10}$** | 1.84 | **$1.04\times10^{-6}$** | **$6.59\times10^{-4}$** | *VSP1* | Vegetative storage protein 1 | | | | | | |
| AT3G03341 | -1.19 | $1.34\times10^{-1}$ | $3.44\times10^{-1}$ | -2.75 | **$2.14\times10^{-3}$** | **$1.12\times10^{-2}$** | 0.63 | **$1.89\times10^{-2}$** | $6.17\times10^{-2}$ | 1.82 | **$2.39\times10^{-2}$** | $5.70\times10^{-1}$ | | Cold-regulated protein | | | | 6.8 | **$4.34\times10^{-5}$** | TTS (1542) |
| AT1G53540 | -1.56 | **$7.15\times10^{-3}$** | **$4.31\times10^{-2}$** | -3.50 | **$1.36\times10^{-7}$** | **$2.75\times10^{-6}$** | 0.22 | $5.11\times10^{-1}$ | $6.84\times10^{-1}$ | 1.77 | **$2.74\times10^{-3}$** | $1.84\times10^{-1}$ | *HSP17.6C* | HSP2-like chaperones superfamily protein | | | | | | |
| AT2G17740 | -0.03 | $9.86\times10^{-1}$ | $9.98\times10^{-1}$ | -0.20 | $8.15\times10^{-1}$ | $8.98\times10^{-1}$ | 1.75 | **$2.05\times10^{-3}$** | **$9.70\times10^{-3}$** | 1.77 | **$4.85\times10^{-2}$** | $7.78\times10^{-1}$ | *VLG* | Cysteine/Histidine-rich C1 domain family protein | -56.5 | **$6.38\times10^{-3}$** | Exon (203) | 9.6 | **$3.65\times10^{-4}$** | Promoter-TSS (-597) |
| AT2G39920 | -1.30 | **$1.72\times10^{-2}$** | $8.30\times10^{-2}$ | -1.19 | **$2.75\times10^{-2}$** | $8.44\times10^{-2}$ | 0.47 | $2.46\times10^{-1}$ | $4.28\times10^{-1}$ | 1.77 | **$2.49\times10^{-4}$** | **$3.40\times10^{-2}$** | | HAD superfamily, subfamily IIIB acid phosphatase | 47.5 | **$1.62\times10^{-3}$** | TTS (1780) | | | |
| AT3G50970 | -0.59 | **$5.98\times10^{-3}$** | **$3.75\times10^{-2}$** | -0.74 | **$4.48\times10^{-3}$** | **$2.02\times10^{-2}$** | 1.18 | **$2.64\times10^{-9}$** | **$4.70\times10^{-8}$** | 1.77 | **$6.09\times10^{-16}$** | **$5.89\times10^{-12}$** | *LTI3* | Low temperature-induced 3 | | | | 5.7 | **$6.26\times10^{-4}$** | Promoter-TSS (-92) |
| AT2G04050 | -1.18 | $1.10\times10^{-1}$ | $3.02\times10^{-1}$ | -0.27 | $7.62\times10^{-1}$ | $8.63\times10^{-1}$ | 0.61 | $1.10\times10^{-1}$ | $2.41\times10^{-1}$ | 1.77 | **$1.12\times10^{-2}$** | $3.89\times10^{-1}$ | | MATE efflux family protein | | | | | | |
| AT4G25000 | -0.99 | $1.17\times10^{-1}$ | $3.15\times10^{-1}$ | -0.60 | $2.53\times10^{-1}$ | $4.29\times10^{-1}$ | 0.76 | $6.20\times10^{-2}$ | $1.56\times10^{-1}$ | 1.75 | **$3.22\times10^{-3}$** | $1.99\times10^{-1}$ | *AMY1* | Alpha-amylase-like | | | | | | |
| AT1G20120 | -0.68 | $2.00\times10^{-1}$ | $4.42\times10^{-1}$ | -0.08 | $8.58\times10^{-1}$ | $9.22\times10^{-1}$ | 1.06 | **$4.41\times10^{-3}$** | **$1.86\times10^{-2}$** | 1.73 | **$6.54\times10^{-5}$** | **$1.29\times10^{-2}$** | | GDSL-motif esterase/acyltransferase/lipase | -66.7 | **$5.29\times10^{-6}$** | Promoter (-1388) | | | |
| AT5G24770 | -1.02 | **$2.44\times10^{-2}$** | $1.08\times10^{-1}$ | -0.93 | $5.67\times10^{-2}$ | $1.46\times10^{-1}$ | 0.71 | **$3.67\times10^{-7}$** | **$4.40\times10^{-6}$** | 1.72 | **$3.27\times10^{-4}$** | **$4.16\times10^{-2}$** | *VSP2* | Vegetative storage protein 2 | | | | | | |
| AT5G53870 | -0.58 | **$1.84\times10^{-2}$** | $8.73\times10^{-2}$ | -0.53 | **$2.44\times10^{-2}$** | $7.70\times10^{-2}$ | 1.13 | **$2.93\times10^{-5}$** | **$2.35\times10^{-4}$** | 1.71 | **$1.12\times10^{-6}$** | **$6.75\times10^{-4}$** | *ENODL1* | Early nodulin-like protein 1 | | | | | | |
| AT4G16146 | -0.99 | **$7.26\times10^{-3}$** | **$4.36\times10^{-2}$** | -0.65 | $6.57\times10^{-2}$ | $1.63\times10^{-1}$ | 0.71 | $5.94\times10^{-2}$ | $1.51\times10^{-1}$ | 1.69 | **$3.08\times10^{-5}$** | **$7.69\times10^{-3}$** | | cAMP-regulated phosphoprotein 19-related protein | | | | | | |
| AT5G24280 | 0.11 | $7.55\times10^{-1}$ | $8.93\times10^{-1}$ | 0.58 | **$2.87\times10^{-2}$** | $8.73\times10^{-2}$ | 1.78 | **$4.78\times10^{-12}$** | **$1.27\times10^{-10}$** | 1.66 | **$5.73\times10^{-7}$** | **$4.11\times10^{-4}$** | *GMI1* | Gamma-irradiation and mitomycin c induced 1 | | | | | | |
| AT4G33905 | 0.13 | $8.58\times10^{-1}$ | $9.43\times10^{-1}$ | -0.09 | $9.09\times10^{-1}$ | $9.53\times10^{-1}$ | 1.78 | **$5.65\times10^{-7}$** | **$6.53\times10^{-6}$** | 1.66 | **$1.95\times10^{-2}$** | $5.21\times10^{-1}$ | | Peroxisomal membrane 22 kDa (Mpv17/PMP22) family protein | | | | | | |
| AT5G63350 | 0.58 | $4.01\times10^{-1}$ | $6.54\times10^{-1}$ | 0.23 | $6.81\times10^{-1}$ | $8.07\times10^{-1}$ | 2.21 | **$9.61\times10^{-12}$** | **$2.46\times10^{-10}$** | 1.64 | **$1.83\times10^{-2}$** | $5.06\times10^{-1}$ | *AITR6* | ABA-induced transcription repressor 6 | -8.4 | **$2.37\times10^{-3}$** | Exon (666) | 6.8 | **$6.86\times10^{-3}$** | 5' UTR-TSS (66) |
| AT5G60910 | -0.76 | $1.93\times10^{-1}$ | $4.32\times10^{-1}$ | -1.11 | **$4.50\times10^{-2}$** | $1.23\times10^{-1}$ | 0.87 | $5.30\times10^{-2}$ | $1.39\times10^{-1}$ | 1.63 | **$3.93\times10^{-5}$** | **$9.16\times10^{-3}$** | *AGL8* | Agamous-like 8 | | | | | | |
| AT5G17460 | -0.39 | $1.83\times10^{-1}$ | $4.16\times10^{-1}$ | -0.93 | **$3.95\times10^{-9}$** | **$1.07\times10^{-7}$** | 1.23 | **$9.61\times10^{-15}$** | **$3.57\times10^{-13}$** | 1.62 | **$3.88\times10^{-7}$** | **$2.91\times10^{-4}$** | *DFR1* | Glutamyl-tRNA (Gln) amidotransferase subunit C | 76.2 | **$9.37\times10^{-5}$** | Exon (1598) | 5.1 | **$3.36\times10^{-3}$** | 5' UTR-TSS (98) |
| AT2G47780 | -1.10 | **$7.81\times10^{-3}$** | **$4.60\times10^{-2}$** | -1.51 | **$1.07\times10^{-7}$** | **$2.22\times10^{-6}$** | 0.52 | $9.48\times10^{-2}$ | $2.16\times10^{-1}$ | 1.62 | **$1.07\times10^{-3}$** | $9.50\times10^{-2}$ | *LDAP2* | LD associated protein 2 | | | | 7.1 | **$9.93\times10^{-4}$** | Exon (511) |
| AT3G43715 | -1.94 | **$1.99\times10^{-3}$** | **$1.57\times10^{-2}$** | -1.21 | **$2.17\times10^{-2}$** | $7.05\times10^{-2}$ | -0.33 | $2.01\times10^{-1}$ | $3.72\times10^{-1}$ | 1.60 | **$9.24\times10^{-3}$** | $3.56\times10^{-1}$ | | Transposable_element_gene | | | | | | |
| AT5G49525 | -1.03 | **$2.47\times10^{-2}$** | $1.09\times10^{-1}$ | -1.13 | **$8.99\times10^{-4}$** | **$5.57\times10^{-3}$** | 0.55 | **$2.58\times10^{-2}$** | $7.92\times10^{-2}$ | 1.58 | **$3.18\times10^{-4}$** | **$4.13\times10^{-2}$** | | Transmembrane protein | | | | 6.2 | **$2.35\times10^{-4}$** | Intergenic(-1400) |
| AT4G37220 | -1.09 | $6.21\times10^{-2}$ | $2.07\times10^{-1}$ | -1.92 | **$2.59\times10^{-6}$** | **$3.74\times10^{-5}$** | 0.49 | $1.34\times10^{-1}$ | $2.78\times10^{-1}$ | 1.58 | **$6.49\times10^{-3}$** | $2.97\times10^{-1}$ | | Cold acclimation protein WCOR413 family | | | | | | |
| AT2G42530 | -1.77 | **$9.43\times10^{-10}$** | **$4.68\times10^{-8}$** | -1.78 | **$4.33\times10^{-15}$** | **$3.39\times10^{-13}$** | -0.20 | $4.61\times10^{-1}$ | $6.41\times10^{-1}$ | 1.57 | **$4.80\times10^{-5}$** | **$1.09\times10^{-2}$** | *COR15B* | Cold regulated 15b | | | | | | |
| AT3G05660 | -0.97 | **$9.67\times10^{-3}$** | $5.42\times10^{-2}$ | -0.67 | $1.79\times10^{-1}$ | $3.37\times10^{-1}$ | 0.59 | $1.58\times10^{-1}$ | $3.11\times10^{-1}$ | 1.56 | **$2.41\times10^{-3}$** | $1.69\times10^{-1}$ | *RLP33* | Receptor like protein 33 | | | | 6.8 | **$1.52\times10^{-4}$** | Exon (1298) |
| AT1G56300 | -0.72 | **$2.04\times10^{-3}$** | **$1.60\times10^{-2}$** | -0.66 | **$8.55\times10^{-4}$** | **$5.33\times10^{-3}$** | 0.85 | **$4.45\times10^{-6}$** | **$4.30\times10^{-5}$** | 1.56 | **$2.34\times10^{-9}$** | **$3.78\times10^{-6}$** | *DJC53* | Chaperone DnaJ-domain superfamily protein | 61.9 | **$1.19\times10^{-3}$** | TTS-3'UTR (2072) | | | |
| AT5G64120 | -0.94 | **$6.94\times10^{-4}$** | **$6.79\times10^{-3}$** | -0.92 | **$3.41\times10^{-2}$** | $9.94\times10^{-2}$ | 0.62 | $7.26\times10^{-2}$ | $1.76\times10^{-1}$ | 1.55 | **$2.69\times10^{-5}$** | **$6.93\times10^{-3}$** | *PRX71* | Peroxidase superfamily protein | | | | | | |
| AT1G68200 | -0.25 | $5.80\times10^{-1}$ | $7.89\times10^{-1}$ | 0.04 | $9.51\times10^{-1}$ | $9.76\times10^{-1}$ | 1.32 | **$2.53\times10^{-4}$** | **$1.61\times10^{-3}$** | 1.55 | **$1.95\times10^{-5}$** | **$5.24\times10^{-3}$** | *CDM1* | Zinc finger C-x8-C-x5-C-x3-H type family protein | | | | | | |
| AT2G47190 | -2.62 | **$6.13\times10^{-11}$** | **$4.02\times10^{-9}$** | -2.72 | **$4.37\times10^{-8}$** | **$9.87\times10^{-7}$** | -1.07 | **$5.06\times10^{-4}$** | **$2.93\times10^{-3}$** | 1.55 | **$9.01\times10^{-5}$** | **$1.63\times10^{-2}$** | *MYB2* | Myb domain protein 2 | | | | | | |
| AT5G15190 | 0.94 | **$3.18\times10^{-2}$** | $1.30\times10^{-1}$ | 0.53 | $3.37\times10^{-1}$ | $5.19\times10^{-1}$ | 2.48 | **$8.74\times10^{-14}$** | **$2.94\times10^{-12}$** | 1.54 | **$7.24\times10^{-4}$** | $7.38\times10^{-2}$ | | Hypothetical protein | | | | | | |
| AT1G05340 | 0.14 | $6.59\times10^{-1}$ | $8.39\times10^{-1}$ | -0.24 | $4.30\times10^{-1}$ | $6.10\times10^{-1}$ | 1.68 | **$4.77\times10^{-11}$** | **$1.11\times10^{-9}$** | 1.54 | **$5.56\times10^{-13}$** | **$3.59\times10^{-9}$** | | Cysteine-rich TM module stress tolerance protein | | | | | | |
| AT3G55970 | -0.57 | $3.49\times10^{-1}$ | $6.09\times10^{-1}$ | -0.08 | $8.64\times10^{-1}$ | $9.25\times10^{-1}$ | 0.97 | **$3.13\times10^{-2}$** | $9.23\times10^{-2}$ | 1.54 | **$1.50\times10^{-2}$** | $4.58\times10^{-1}$ | *JRG21* | Jasmonate-regulated gene 21 | | | | 8.0 | **$4.56\times10^{-4}$** | Intergenic(-3041) |

| | | | | | | | | | | | | | | | | | |
|---|---|---|---|---|---|---|---|---|---|---|---|---|---|---|---|---|---|
| AT3G28007 | 0.84 | $8.19\times10^{-2}$ | $2.50\times10^{-1}$ | 0.16 | $8.01\times10^{-1}$ | $8.89\times10^{-1}$ | 2.36 | **$2.08\times10^{-8}$** | **$3.11\times10^{-7}$** | 1.52 | **$3.91\times10^{-3}$** | $2.24\times10^{-1}$ | *SWEET4* | Nodulin MtN3 family protein | | | |
| AT5G22460 | 0.50 | $4.63\times10^{-1}$ | $7.06\times10^{-1}$ | 0.06 | $7.99\times10^{-1}$ | $8.87\times10^{-1}$ | 2.01 | **$1.02\times10^{-7}$** | **$1.34\times10^{-6}$** | 1.52 | **$4.62\times10^{-2}$** | $7.63\times10^{-1}$ | | Alpha/beta-Hydrolases superfamily protein | | | |
| AT1G73810 | -0.85 | $2.41\times10^{-1}$ | $4.94\times10^{-1}$ | 0.03 | $9.78\times10^{-1}$ | $9.91\times10^{-1}$ | 0.66 | $8.19\times10^{-2}$ | $1.93\times10^{-1}$ | 1.50 | **$2.41\times10^{-2}$** | $5.71\times10^{-1}$ | | Core-2/I-branching beta-1,6-N-acetylglucosaminyltransferase family protein | 7.9 | **$2.03\times10^{-3}$** | Intron (1746) |
| AT4G29610 | -1.58 | **$3.38\times10^{-3}$** | **$2.39\times10^{-2}$** | -0.88 | $1.15\times10^{-1}$ | $2.46\times10^{-1}$ | -0.06 | $8.94\times10^{-1}$ | $9.47\times10^{-1}$ | 1.50 | **$1.38\times10^{-4}$** | **$2.28\times10^{-2}$** | | Cytidine/deoxycytidylate deaminase family protein | 7.2 | **$4.11\times10^{-4}$** | Promoter-TSS (-242) |
| AT2G33380 | -1.30 | **$8.29\times10^{-6}$** | **$1.56\times10^{-4}$** | -1.68 | **$2.50\times10^{-14}$** | **$1.76\times10^{-12}$** | 0.21 | $4.08\times10^{-1}$ | $5.93\times10^{-1}$ | 1.50 | **$1.04\times10^{-5}$** | **$3.09\times10^{-3}$** | *RD2* | Responsive to dessication 2 | | | |
| AT2G21590 | 0.20 | $6.30\times10^{-1}$ | $8.21\times10^{-1}$ | -0.39 | $4.40\times10^{-1}$ | $6.18\times10^{-1}$ | 1.68 | **$1.02\times10^{-4}$** | **$7.22\times10^{-4}$** | 1.49 | **$1.13\times10^{-2}$** | $3.92\times10^{-1}$ | *APL4* | Glucose-1-phosphate adenylyltransferase family protein | | | |
| AT1G31750 | -0.23 | $5.98\times10^{-1}$ | $8.00\times10^{-1}$ | -0.75 | $1.85\times10^{-1}$ | $3.45\times10^{-1}$ | 1.23 | **$9.53\times10^{-6}$** | **$8.60\times10^{-5}$** | 1.45 | **$5.70\times10^{-4}$** | $6.34\times10^{-2}$ | *GPRP1* | Proline-rich family protein | | | |
| AT1G61800 | -1.68 | **$1.73\times10^{-4}$** | **$2.14\times10^{-3}$** | -1.87 | **$3.55\times10^{-8}$** | **$8.14\times10^{-7}$** | -0.23 | $6.16\times10^{-1}$ | $7.65\times10^{-1}$ | 1.45 | **$1.52\times10^{-2}$** | $4.61\times10^{-1}$ | *GPT2* | Glucose-6-phosphate/phosphate translocator 2 | | | |
| AT3G49120 | -0.36 | $1.56\times10^{-1}$ | $3.78\times10^{-1}$ | -0.18 | $5.54\times10^{-1}$ | $7.13\times10^{-1}$ | 1.09 | **$2.04\times10^{-6}$** | **$2.11\times10^{-5}$** | 1.44 | **$9.81\times10^{-6}$** | **$3.01\times10^{-3}$** | *PRXCB* | Peroxidase CB | | | |
| AT2G42540 | -0.69 | **$2.97\times10^{-2}$** | $1.24\times10^{-1}$ | -1.34 | **$1.45\times10^{-5}$** | **$1.69\times10^{-4}$** | 0.74 | **$1.55\times10^{-2}$** | $5.28\times10^{-2}$ | 1.43 | **$1.45\times10^{-3}$** | $1.18\times10^{-1}$ | *COR15A* | Cold-regulated 15a | | | |
| AT2G19190 | -0.25 | $6.61\times10^{-1}$ | $8.40\times10^{-1}$ | -0.22 | $6.93\times10^{-1}$ | $8.16\times10^{-1}$ | 1.19 | **$2.10\times10^{-4}$** | **$1.37\times10^{-3}$** | 1.43 | **$1.01\times10^{-2}$** | $3.70\times10^{-1}$ | *FRK1* | FLG22-induced receptor-like kinase 1 | 8.6 | **$1.58\times10^{-3}$** | Promoter-TSS (-7) |
| AT5G10760 | -1.01 | $8.72\times10^{-2}$ | $2.60\times10^{-1}$ | -0.05 | $8.71\times10^{-1}$ | $9.30\times10^{-1}$ | 0.43 | **$1.77\times10^{-3}$** | **$8.56\times10^{-3}$** | 1.43 | **$1.72\times10^{-2}$** | $4.90\times10^{-1}$ | *AED1* | Eukaryotic aspartyl protease family protein | | | |
| AT2G45550 | -0.88 | $7.10\times10^{-2}$ | $2.27\times10^{-1}$ | -0.24 | $6.10\times10^{-1}$ | $7.57\times10^{-1}$ | 0.53 | $1.47\times10^{-1}$ | $2.97\times10^{-1}$ | 1.39 | **$1.02\times10^{-2}$** | $3.72\times10^{-1}$ | *CYP76C4* | Cytochrome P45, family 76, subfamily C, polypeptide 4 | | | |
| AT1G05100 | -0.01 | $1.00\times10^{-0}$ | $1.00\times10^{-0}$ | -0.28 | $5.22\times10^{-1}$ | $6.88\times10^{-1}$ | 1.39 | **$3.08\times10^{-6}$** | **$3.09\times10^{-5}$** | 1.39 | **$9.07\times10^{-3}$** | $3.53\times10^{-1}$ | *MAPKKK18* | Mitogen-activated protein kinase kinase kinase 18 | 6.9 | **$2.26\times10^{-3}$** | Intergenic space/Promoter (1110) |
| AT1G52040 | 1.18 | **$1.11\times10^{-4}$** | **$1.46\times10^{-3}$** | 1.12 | **$3.01\times10^{-6}$** | **$4.26\times10^{-5}$** | 2.57 | **$3.02\times10^{-54}$** | **$1.57\times10^{-51}$** | 1.39 | **$5.07\times10^{-6}$** | **$1.92\times10^{-3}$** | *MBP1* | Myrosinase-binding protein 1 | | | |
| AT1G54160 | -0.37 | $3.23\times10^{-1}$ | $5.84\times10^{-1}$ | -0.84 | **$2.62\times10^{-2}$** | $8.13\times10^{-2}$ | 1.01 | **$9.58\times10^{-3}$** | **$3.55\times10^{-2}$** | 1.38 | **$4.77\times10^{-3}$** | $2.48\times10^{-1}$ | *NF-YA5* | Nuclear factor Y, subunit A5 | | | |
| AT3G03470 | -0.05 | $8.21\times10^{-1}$ | $9.24\times10^{-1}$ | -0.16 | $4.00\times10^{-1}$ | $5.81\times10^{-1}$ | 1.32 | **$6.73\times10^{-11}$** | **$1.54\times10^{-9}$** | 1.37 | **$2.68\times10^{-7}$** | **$2.26\times10^{-4}$** | *CYP89A9* | Cytochrome P45, family 87, subfamily A, polypeptide 9 | 21.0 | **$3.73\times10^{-3}$** | Promoter-TSS (-266) |
| AT1G14520 | -1.05 | $7.15\times10^{-2}$ | $2.28\times10^{-1}$ | -0.66 | $1.01\times10^{-1}$ | $2.24\times10^{-1}$ | 0.31 | $1.35\times10^{-1}$ | $2.80\times10^{-1}$ | 1.35 | **$2.42\times10^{-2}$** | $5.73\times10^{-1}$ | *MIOX1* | Myo-inositol oxygenase 1 | | | |
| AT5G48720 | -0.20 | $3.00\times10^{-1}$ | $5.60\times10^{-1}$ | -0.04 | $8.38\times10^{-1}$ | $9.11\times10^{-1}$ | 1.16 | **$3.26\times10^{-11}$** | **$7.77\times10^{-10}$** | 1.35 | **$5.86\times10^{-11}$** | **$1.89\times10^{-7}$** | *XRI1* | X-ray induced transcript 1 | | | |
| AT5G54190 | -1.54 | **$1.84\times10^{-2}$** | $8.73\times10^{-2}$ | -1.43 | **$3.88\times10^{-3}$** | **$1.80\times10^{-2}$** | -0.19 | $5.62\times10^{-1}$ | $7.23\times10^{-1}$ | 1.35 | **$4.13\times10^{-2}$** | $7.25\times10^{-1}$ | *PORA* | Protochlorophyllide oxidoreductase A | | | |
| AT4G12290 | -0.40 | $2.37\times10^{-1}$ | $4.89\times10^{-1}$ | -0.33 | $1.82\times10^{-1}$ | $3.41\times10^{-1}$ | 0.94 | **$1.29\times10^{-3}$** | **$6.54\times10^{-3}$** | 1.34 | **$2.19\times10^{-3}$** | $1.57\times10^{-1}$ | *CUAO* | Copper amine oxidase family protein | | | |
| AT4G33150 | 2.20 | **$8.82\times10^{-16}$** | **$1.60\times10^{-13}$** | 2.04 | **$1.11\times10^{-29}$** | **$4.95\times10^{-27}$** | 3.53 | **$1.53\times10^{-43}$** | **$3.73\times10^{-41}$** | 1.33 | **$3.50\times10^{-4}$** | **$4.40\times10^{-2}$** | *LKR* | Lysine-ketoglutarate reductase/saccharopine dehydrogenase bifunctional enzyme | 9.3 | **$7.71\times10^{-3}$** | Exon (690) |
| AT4G19390 | 0.13 | $6.53\times10^{-1}$ | $8.35\times10^{-1}$ | -0.22 | $3.66\times10^{-1}$ | $5.47\times10^{-1}$ | 1.46 | **$2.46\times10^{-11}$** | **$5.96\times10^{-10}$** | 1.32 | **$2.10\times10^{-6}$** | **$1.13\times10^{-3}$** | | Uncharacterized protein family (UPF0114) | | | |
| AT1G77450 | -2.36 | **$1.29\times10^{-8}$** | **$5.00\times10^{-7}$** | -2.55 | **$2.59\times10^{-13}$** | **$1.57\times10^{-11}$** | -1.04 | **$9.94\times10^{-6}$** | **$8.92\times10^{-5}$** | 1.32 | **$1.37\times10^{-3}$** | $1.13\times10^{-1}$ | *NAC32* | NAC domain containing protein 32 | | | |
| AT5G01600 | -0.44 | $1.32\times10^{-1}$ | $3.41\times10^{-1}$ | -0.43 | $1.15\times10^{-1}$ | $2.45\times10^{-1}$ | 0.89 | **$4.95\times10^{-5}$** | **$3.76\times10^{-4}$** | 1.32 | **$9.10\times10^{-6}$** | **$2.89\times10^{-3}$** | *FER1* | Ferretin 1 | | | |
| AT4G23600 | 0.08 | $7.30\times10^{-1}$ | $8.81\times10^{-1}$ | -0.07 | $7.82\times10^{-1}$ | $8.77\times10^{-1}$ | 1.35 | **$4.88\times10^{-7}$** | **$5.71\times10^{-6}$** | 1.27 | **$1.65\times10^{-4}$** | **$2.61\times10^{-2}$** | *CORI3* | Coronatine induced 1 | | | |
| AT1G11210 | 0.24 | $6.11\times10^{-1}$ | $8.09\times10^{-1}$ | -0.50 | $3.92\times10^{-1}$ | $5.73\times10^{-1}$ | 1.51 | **$6.98\times10^{-6}$** | **$6.48\times10^{-5}$** | 1.27 | **$9.21\times10^{-4}$** | $8.69\times10^{-2}$ | | Cotton fiber protein | | | |
| AT5G63580 | -0.40 | $3.56\times10^{-1}$ | $6.15\times10^{-1}$ | 0.06 | $8.86\times10^{-1}$ | $9.39\times10^{-1}$ | 0.85 | **$2.76\times10^{-2}$** | $8.36\times10^{-2}$ | 1.24 | **$3.50\times10^{-3}$** | $2.05\times10^{-1}$ | *FLS2* | Flavonol synthase 2 | | | |
| AT1G78990 | -1.24 | **$1.53\times10^{-3}$** | **$1.27\times10^{-2}$** | -0.77 | $2.87\times10^{-1}$ | $4.67\times10^{-1}$ | 0.01 | $9.90\times10^{-1}$ | $9.97\times10^{-1}$ | 1.24 | **$1.34\times10^{-3}$** | $1.11\times10^{-1}$ | | HXXXD-type acyl-transferase family protein | 5.1 | **$9.13\times10^{-3}$** | Exon (666) |
| AT1G20450 | -0.49 | $2.53\times10^{-1}$ | $5.09\times10^{-1}$ | -0.97 | **$1.60\times10^{-3}$** | **$8.88\times10^{-3}$** | 0.75 | **$2.04\times10^{-4}$** | **$1.33\times10^{-3}$** | 1.24 | **$9.83\times10^{-3}$** | $3.64\times10^{-1}$ | *ERD1* | Early responsive to dehydration 1 | | | |
| AT1G65890 | -1.33 | **$1.66\times10^{-5}$** | **$2.87\times10^{-4}$** | -0.77 | **$1.99\times10^{-2}$** | $6.58\times10^{-2}$ | -0.08 | $8.09\times10^{-1}$ | $8.94\times10^{-1}$ | 1.24 | **$3.19\times10^{-3}$** | $1.98\times10^{-1}$ | *AAE12* | Acyl activating enzyme 12 | | | |
| AT1G66700 | -3.80 | **$1.17\times10^{-18}$** | **$3.10\times10^{-16}$** | -3.52 | **$3.08\times10^{-26}$** | **$9.57\times10^{-24}$** | -2.56 | **$1.56\times10^{-28}$** | **$2.33\times10^{-26}$** | 1.24 | **$7.38\times10^{-3}$** | $3.14\times10^{-1}$ | *PXMT1* | S-adenosyl-L-methionine-dependent methyltransferases superfamily protein | 12.8 | **$9.25\times10^{-3}$** | Intergenic (-1385) |
| AT1G56600 | -0.29 | $5.83\times10^{-1}$ | $7.91\times10^{-1}$ | -0.85 | $9.16\times10^{-2}$ | $2.09\times10^{-1}$ | 0.94 | **$2.42\times10^{-5}$** | **$1.99\times10^{-4}$** | 1.23 | **$3.31\times10^{-2}$** | $6.64\times10^{-1}$ | *GolS2* | Galactinol synthase 2 | | | |

| | | | | | | | | | | | | | | | | | | | | |
|---|---|---|---|---|---|---|---|---|---|---|---|---|---|---|---|---|---|---|---|---|
| AT5G58310 | -1.03 | **$1.37\times10^{-3}$** | **$1.16\times10^{-2}$** | -0.71 | **$1.53\times10^{-2}$** | $5.32\times10^{-2}$ | 0.19 | $4.66\times10^{-1}$ | $6.45\times10^{-1}$ | 1.22 | **$5.14\times10^{-5}$** | **$1.12\times10^{-2}$** | *MES18* | Methyl esterase 18 | | | | 6.4 | **$6.43\times10^{-4}$** | Promoter-TSS (-167) |
| AT4G27410 | -0.03 | $9.40\times10^{-1}$ | $9.78\times10^{-1}$ | -0.55 | $8.98\times10^{-2}$ | $2.06\times10^{-1}$ | 1.19 | **$4.21\times10^{-12}$** | **$1.13\times10^{-10}$** | 1.22 | **$2.00\times10^{-3}$** | $1.50\times10^{-1}$ | *RD26* | Responsive to desiccation 26 | | | | | | |
| AT3G55880 | 0.74 | **$5.50\times10^{-3}$** | **$3.51\times10^{-2}$** | 0.67 | **$2.02\times10^{-3}$** | **$1.07\times10^{-2}$** | 1.96 | **$1.16\times10^{-13}$** | **$3.84\times10^{-12}$** | 1.21 | **$5.77\times10^{-5}$** | **$1.23\times10^{-2}$** | *SUE4* | Sulphate utilization efficiency 4 | | | | | | |
| AT1G76590 | 0.61 | $6.12\times10^{-2}$ | $2.04\times10^{-1}$ | 0.99 | **$3.28\times10^{-3}$** | **$1.57\times10^{-2}$** | 1.83 | **$2.57\times10^{-18}$** | **$1.41\times10^{-16}$** | 1.21 | **$3.21\times10^{-5}$** | **$7.86\times10^{-3}$** | *PLATZ2* | PLATZ transcription factor family protein | | | | | | |
| AT3G05640 | -0.23 | $5.54\times10^{-1}$ | $7.72\times10^{-1}$ | -0.54 | **$6.76\times10^{-3}$** | **$2.81\times10^{-2}$** | 0.97 | **$2.48\times10^{-4}$** | **$1.58\times10^{-3}$** | 1.20 | **$1.01\times10^{-2}$** | $3.70\times10^{-1}$ | *EGR1* | Protein phosphatase 2C family protein | 80.7 | **$3.63\times10^{-4}$** | Exon (3923) | | | |
| AT1G29395 | -1.38 | **$1.22\times10^{-8}$** | **$4.75\times10^{-7}$** | -1.82 | **$5.82\times10^{-23}$** | **$1.20\times10^{-20}$** | -0.19 | $4.77\times10^{-1}$ | $6.55\times10^{-1}$ | 1.19 | **$2.39\times10^{-4}$** | **$3.32\times10^{-2}$** | *COR413IM1* | COLD REGULATED 314 INNER MEMBRANE 1 | 69.4 | **$7.38\times10^{-3}$** | Promoter (-1885) | | | |
| AT3G13672 | 1.34 | **$1.08\times10^{-2}$** | $5.85\times10^{-2}$ | 0.22 | $6.94\times10^{-1}$ | $8.17\times10^{-1}$ | 2.52 | **$9.51\times10^{-19}$** | **$5.49\times10^{-17}$** | 1.18 | **$2.32\times10^{-2}$** | $5.64\times10^{-1}$ | | TRAF-like superfamily protein | | | | | | |
| AT1G69260 | -0.89 | $8.28\times10^{-2}$ | $2.51\times10^{-1}$ | -1.44 | **$5.15\times10^{-3}$** | **$2.27\times10^{-2}$** | 0.29 | $2.58\times10^{-1}$ | $4.41\times10^{-1}$ | 1.17 | **$3.99\times10^{-2}$** | $7.14\times10^{-1}$ | *AFP1* | ABI five binding protein | | | | | | |
| AT5G37690 | 0.64 | **$4.16\times10^{-2}$** | $1.57\times10^{-1}$ | 0.72 | **$1.99\times10^{-2}$** | $6.58\times10^{-2}$ | 1.82 | **$4.76\times10^{-11}$** | **$1.11\times10^{-9}$** | 1.17 | **$1.98\times10^{-4}$** | **$2.92\times10^{-2}$** | | SGNH hydrolase-type esterase superfamily protein | | | | | | |
| AT5G24470 | 0.16 | $6.82\times10^{-1}$ | $8.53\times10^{-1}$ | 0.34 | $3.50\times10^{-1}$ | $5.32\times10^{-1}$ | 1.33 | **$2.18\times10^{-5}$** | **$1.81\times10^{-4}$** | 1.17 | **$1.69\times10^{-7}$** | **$1.72\times10^{-4}$** | *PRR5* | Pseudo-response regulator 5 | | | | 5.8 | **$7.57\times10^{-4}$** | Exon (1295) |
| AT1G74460 | -0.36 | $1.37\times10^{-1}$ | $3.48\times10^{-1}$ | -0.17 | $5.69\times10^{-1}$ | $7.25\times10^{-1}$ | 0.81 | **$3.41\times10^{-4}$** | **$2.08\times10^{-3}$** | 1.17 | **$4.11\times10^{-6}$** | **$1.69\times10^{-3}$** | | GDSL-motif esterase/acyltransferase/lipase | | | | 9.0 | **$3.07\times10^{-6}$** | 5' UTR-TSS (14) |
| AT2G34850 | -0.65 | **$3.35\times10^{-2}$** | $1.35\times10^{-1}$ | -0.54 | **$4.68\times10^{-2}$** | $1.26\times10^{-1}$ | 0.51 | $8.42\times10^{-2}$ | $1.97\times10^{-1}$ | 1.16 | **$1.92\times10^{-3}$** | $1.47\times10^{-1}$ | *MEE25* | Maternal effect embryo arrest 25 | | | | | | |
| AT3G05650 | -0.82 | **$1.92\times10^{-2}$** | $9.00\times10^{-2}$ | -0.24 | $5.16\times10^{-1}$ | $6.82\times10^{-1}$ | 0.33 | $3.45\times10^{-1}$ | $5.32\times10^{-1}$ | 1.15 | **$2.71\times10^{-3}$** | $1.83\times10^{-1}$ | *RLP32* | Receptor like protein 32 | | | | | | |
| AT1G62570 | -0.07 | $7.58\times10^{-1}$ | $8.94\times10^{-1}$ | -0.14 | $4.01\times10^{-1}$ | $5.81\times10^{-1}$ | 1.07 | **$3.03\times10^{-14}$** | **$1.07\times10^{-12}$** | 1.14 | **$4.76\times10^{-6}$** | **$1.88\times10^{-3}$** | *FMO GS-OX4* | Flavin-monooxygenase glucosinolate S-oxygenase 4 | | | | | | |
| AT3G62550 | 0.47 | $7.37\times10^{-2}$ | $2.33\times10^{-1}$ | 0.26 | $2.66\times10^{-1}$ | $4.44\times10^{-1}$ | 1.62 | **$2.71\times10^{-21}$** | **$2.11\times10^{-19}$** | 1.14 | **$3.53\times10^{-6}$** | **$1.49\times10^{-3}$** | | Adenine nucleotide alpha hydrolases-like superfamily protein | | | | | | |
| AT3G61890 | -0.82 | $5.06\times10^{-2}$ | $1.79\times10^{-1}$ | -1.27 | **$4.99\times10^{-4}$** | **$3.40\times10^{-3}$** | 0.32 | $1.66\times10^{-1}$ | $3.24\times10^{-1}$ | 1.14 | **$1.33\times10^{-2}$** | $4.27\times10^{-1}$ | *HB-12* | Homeobox 12 | | | | | | |
| AT1G09500 | -0.31 | $3.93\times10^{-1}$ | $6.47\times10^{-1}$ | -0.69 | $1.10\times10^{-1}$ | $2.38\times10^{-1}$ | 0.82 | **$6.63\times10^{-3}$** | **$2.64\times10^{-2}$** | 1.14 | **$6.50\times10^{-4}$** | $7.02\times10^{-2}$ | | similar to Eucalyptus gunnii alcohol dehydrogenase | | | | | | |
| AT1G54100 | -0.41 | $7.40\times10^{-2}$ | $2.33\times10^{-1}$ | -0.72 | **$2.09\times10^{-4}$** | **$1.65\times10^{-3}$** | 0.72 | **$1.44\times10^{-3}$** | **$7.19\times10^{-3}$** | 1.13 | **$4.89\times10^{-4}$** | $5.71\times10^{-2}$ | *ALDH7B4* | Aldehyde dehydrogenase 7B4 | | | | | | |
| AT2G18050 | 2.67 | **$4.04\times10^{-15}$** | **$6.35\times10^{-13}$** | 2.68 | **$3.53\times10^{-6}$** | **$4.87\times10^{-5}$** | 3.80 | **$8.80\times10^{-77}$** | **$1.70\times10^{-73}$** | 1.13 | **$7.56\times10^{-6}$** | **$2.57\times10^{-3}$** | *HIS1-3* | Histone H1-3 | | | | | | |
| AT4G28940 | 1.51 | **$2.54\times10^{-3}$** | **$1.90\times10^{-2}$** | 1.29 | **$6.60\times10^{-3}$** | **$2.76\times10^{-2}$** | 2.64 | **$3.31\times10^{-8}$** | **$4.77\times10^{-7}$** | 1.12 | **$7.32\times10^{-3}$** | $3.14\times10^{-1}$ | | Phosphorylase superfamily protein | | | | | | |
| AT4G30650 | -1.50 | **$8.82\times10^{-6}$** | **$1.65\times10^{-4}$** | -1.87 | **$2.10\times10^{-23}$** | **$4.76\times10^{-21}$** | -0.39 | **$1.16\times10^{-2}$** | **$4.15\times10^{-2}$** | 1.10 | **$1.57\times10^{-3}$** | $1.23\times10^{-1}$ | | Low temperature and salt responsive protein family | | | | | | |
| AT1G30160 | -0.66 | $5.81\times10^{-2}$ | $1.97\times10^{-1}$ | -0.32 | $3.13\times10^{-1}$ | $4.94\times10^{-1}$ | 0.45 | $9.13\times10^{-2}$ | $2.10\times10^{-1}$ | 1.10 | **$2.30\times10^{-3}$** | $1.65\times10^{-1}$ | | Hypothetical protein (DUF295); | | | | | | |
| AT4G23680 | 1.31 | **$2.02\times10^{-11}$** | **$1.41\times10^{-9}$** | 1.39 | **$2.36\times10^{-14}$** | **$1.66\times10^{-12}$** | 2.41 | **$3.75\times10^{-46}$** | **$1.19\times10^{-43}$** | 1.10 | **$1.05\times10^{-7}$** | **$1.26\times10^{-4}$** | | Polyketide cyclase/dehydrase and lipid transport superfamily protein | | | | 15.3 | **$2.60\times10^{-4}$** | Exon (1348) |
| AT1G22370 | -0.67 | **$5.26\times10^{-4}$** | **$5.41\times10^{-3}$** | -0.76 | **$2.49\times10^{-5}$** | **$2.70\times10^{-4}$** | 0.43 | **$1.20\times10^{-2}$** | **$4.27\times10^{-2}$** | 1.10 | **$1.56\times10^{-6}$** | **$8.62\times10^{-4}$** | *UGT85A5* | UDP-glucosyl transferase 85A5 | | | | 13.1 | **$2.46\times10^{-3}$** | Promoter-TSS (-816) |
| AT1G10070 | 0.99 | $5.77\times10^{-2}$ | $1.96\times10^{-1}$ | 0.78 | $1.34\times10^{-1}$ | $2.74\times10^{-1}$ | 2.09 | **$1.29\times10^{-6}$** | **$1.40\times10^{-5}$** | 1.09 | **$9.60\times10^{-3}$** | $3.61\times10^{-1}$ | *BCAT-2* | Branched-chain amino acid transaminase 2 | | | | 7.7 | **$1.96\times10^{-5}$** | 5' UTR (-1536) |
| AT2G43570 | -0.58 | $1.34\times10^{-1}$ | $3.44\times10^{-1}$ | -1.09 | **$4.92\times10^{-4}$** | **$3.36\times10^{-3}$** | 0.50 | **$3.13\times10^{-2}$** | $9.21\times10^{-2}$ | 1.09 | **$7.60\times10^{-3}$** | $3.18\times10^{-1}$ | *CHI* | Chitinase, putative | | | | | | |
| AT3G26170 | 0.51 | $1.38\times10^{-1}$ | $3.50\times10^{-1}$ | 0.89 | **$8.86\times10^{-4}$** | **$5.50\times10^{-3}$** | 1.60 | **$7.07\times10^{-20}$** | **$4.76\times10^{-18}$** | 1.09 | **$5.14\times10^{-4}$** | $5.88\times10^{-2}$ | *CYP71B19* | Cytochrome P45, family 71, subfamily B, polypeptide 19 | | | | | | |
| AT2G29460 | -1.71 | **$9.51\times10^{-4}$** | **$8.71\times10^{-3}$** | -2.46 | **$1.10\times10^{-3}$** | **$6.54\times10^{-3}$** | -0.61 | $9.90\times10^{-2}$ | $2.23\times10^{-1}$ | 1.09 | **$1.45\times10^{-2}$** | $4.50\times10^{-1}$ | *GSTU4* | Glutathione S-transferase tau 4 | | | | | | |
| AT1G19250 | -1.95 | **$8.01\times10^{-5}$** | **$1.11\times10^{-3}$** | -2.07 | **$2.62\times10^{-6}$** | **$3.77\times10^{-5}$** | -0.86 | **$1.26\times10^{-2}$** | **$4.47\times10^{-2}$** | 1.08 | **$1.44\times10^{-2}$** | $4.50\times10^{-1}$ | *FMO1* | Flavin-dependent monooxygenase 1 | | | | | | |
| AT1G55990 | -2.12 | **$3.68\times10^{-8}$** | **$1.29\times10^{-6}$** | -1.80 | **$2.06\times10^{-6}$** | **$3.08\times10^{-5}$** | -1.03 | **$5.40\times10^{-4}$** | **$3.10\times10^{-3}$** | 1.08 | **$6.53\times10^{-3}$** | $2.97\times10^{-1}$ | | Glycine-rich protein | | | | | | |
| AT1G49720 | 0.04 | $8.79\times10^{-1}$ | $9.52\times10^{-1}$ | -0.09 | $7.06\times10^{-1}$ | $8.24\times10^{-1}$ | 1.12 | **$1.81\times10^{-16}$** | **$8.19\times10^{-15}$** | 1.07 | **$2.72\times10^{-5}$** | **$6.93\times10^{-3}$** | *ABF1* | Abscisic acid responsive element-binding factor 1 | | | | | | |
| AT2G26480 | -0.59 | $1.05\times10^{-1}$ | $2.93\times10^{-1}$ | -0.50 | $9.49\times10^{-2}$ | $2.14\times10^{-1}$ | 0.49 | $1.02\times10^{-1}$ | $2.26\times10^{-1}$ | 1.07 | **$5.75\times10^{-3}$** | $2.79\times10^{-1}$ | *UGT76D1* | UDP-glucosyl transferase 76D1 | | | | | | |
| AT3G57520 | 2.53 | **$9.95\times10^{-12}$** | **$7.65\times10^{-10}$** | 2.33 | **$1.73\times10^{-11}$** | **$7.67\times10^{-10}$** | 3.60 | **$1.96\times10^{-60}$** | **$1.45\times10^{-57}$** | 1.07 | **$2.88\times10^{-3}$** | $1.89\times10^{-1}$ | *SIP2* | Seed imbibition 2 | | | | | | |
| AT1G52400 | 0.38 | $4.50\times10^{-1}$ | $6.95\times10^{-1}$ | 0.51 | $1.83\times10^{-1}$ | $3.43\times10^{-1}$ | 1.46 | **$2.53\times10^{-11}$** | **$6.13\times10^{-10}$** | 1.06 | **$4.41\times10^{-2}$** | $7.46\times10^{-1}$ | *BGLU18* | Beta glucosidase 18 | | | | | | |
| AT5G47635 | -0.62 | $1.79\times10^{-1}$ | $4.11\times10^{-1}$ | -0.23 | $6.04\times10^{-1}$ | $7.52\times10^{-1}$ | 0.46 | $2.51\times10^{-1}$ | $4.34\times10^{-1}$ | 1.06 | **$4.39\times10^{-2}$** | $7.45\times10^{-1}$ | | Pollen Ole e 1 allergen and extensin family protein | | | | | | |

| | | | | | | | | | | | | | | | | | | | | |
|---|---|---|---|---|---|---|---|---|---|---|---|---|---|---|---|---|---|---|---|---|
| AT5G61820 | -0.67 | **$1.35\times10^{-2}$** | $6.92\times10^{-2}$ | -0.88 | **$1.83\times10^{-6}$** | **$2.77\times10^{-5}$** | 0.39 | $5.97\times10^{-2}$ | $1.52\times10^{-1}$ | 1.06 | **$1.47\times10^{-3}$** | $1.19\times10^{-1}$ | *NOD19* | Nodulin 19 | | | | | | |
| AT1G56650 | -0.97 | **$7.81\times10^{-4}$** | **$7.46\times10^{-3}$** | -1.09 | **$1.61\times10^{-4}$** | **$1.33\times10^{-3}$** | 0.08 | $7.66\times10^{-1}$ | $8.69\times10^{-1}$ | 1.05 | **$4.95\times10^{-3}$** | $2.53\times10^{-1}$ | *PAP1* | Production of anthocyanin pigment 1 | | | | | | |
| AT2G29500 | -2.55 | **$1.66\times10^{-7}$** | **$4.87\times10^{-6}$** | -3.09 | **$9.61\times10^{-18}$** | **$1.06\times10^{-15}$** | -1.50 | **$7.38\times10^{-9}$** | **$1.20\times10^{-7}$** | 1.05 | **$1.99\times10^{-2}$** | $5.25\times10^{-1}$ | | HSP2-like chaperones superfamily protein | | | | | | |
| AT4G12480 | 0.94 | **$1.41\times10^{-3}$** | **$1.19\times10^{-2}$** | 0.23 | $6.17\times10^{-1}$ | $7.61\times10^{-1}$ | 1.99 | **$5.25\times10^{-6}$** | **$5.00\times10^{-5}$** | 1.05 | **$2.16\times10^{-2}$** | $5.47\times10^{-1}$ | *EARLI1* | Early arabidopsis aluminum induced 1 | | | | | | |
| AT2G22510 | -0.70 | **$2.78\times10^{-3}$** | **$2.04\times10^{-2}$** | -0.34 | $2.14\times10^{-1}$ | $3.82\times10^{-1}$ | 0.35 | $1.78\times10^{-1}$ | $3.40\times10^{-1}$ | 1.04 | **$9.10\times10^{-4}$** | $8.63\times10^{-2}$ | | Hydroxyproline-rich glycoprotein family protein | | | | | | |
| AT3G26740 | -0.88 | **$1.27\times10^{-5}$** | **$2.27\times10^{-4}$** | -0.97 | $2.55\times10^{-8}$ | $6.00\times10^{-7}$ | 0.16 | $2.32\times10^{-1}$ | $4.10\times10^{-1}$ | 1.04 | **$1.79\times10^{-7}$** | **$1.74\times10^{-4}$** | *CCL* | CCR-like | | | | | | |
| AT5G03780 | 0.04 | $9.01\times10^{-1}$ | $9.61\times10^{-1}$ | 0.39 | $1.29\times10^{-1}$ | $2.67\times10^{-1}$ | 1.09 | **$1.02\times10^{-9}$** | **$1.97\times10^{-8}$** | 1.04 | **$7.25\times10^{-4}$** | $7.38\times10^{-2}$ | *TRFL1* | TRF-like 1 | | | | | | |
| AT5G45310 | 0.38 | $2.11\times10^{-1}$ | $4.57\times10^{-1}$ | 0.22 | $3.98\times10^{-1}$ | $5.79\times10^{-1}$ | 1.42 | **$1.12\times10^{-9}$** | **$2.15\times10^{-8}$** | 1.03 | **$1.26\times10^{-4}$** | **$2.17\times10^{-2}$** | | Coiled-coil protein | | | | 6.5 | **$4.28\times10^{-3}$** | Promoter-TSS (-238) |
| AT5G67480 | 0.44 | $8.17\times10^{-2}$ | $2.49\times10^{-1}$ | 0.09 | $6.58\times10^{-1}$ | $7.92\times10^{-1}$ | 1.48 | **$1.49\times10^{-12}$** | **$4.24\times10^{-11}$** | 1.03 | **$5.89\times10^{-5}$** | **$1.23\times10^{-2}$** | *BT4* | BTB and TAZ domain protein 4 | | | | | | |
| AT5G18130 | -0.26 | $1.61\times10^{-1}$ | $3.85\times10^{-1}$ | -0.37 | $5.63\times10^{-2}$ | $1.45\times10^{-1}$ | 0.77 | **$8.24\times10^{-7}$** | **$9.25\times10^{-6}$** | 1.02 | **$9.95\times10^{-7}$** | **$6.59\times10^{-4}$** | | Transmembrane protein | | | | | | |
| AT3G04040 | -1.41 | **$2.19\times10^{-4}$** | **$2.61\times10^{-3}$** | -1.20 | **$2.26\times10^{-3}$** | **$1.17\times10^{-2}$** | -0.40 | $2.74\times10^{-1}$ | $4.59\times10^{-1}$ | 1.02 | **$3.24\times10^{-2}$** | $6.55\times10^{-1}$ | | Transmembrane protein | | | | | | |
| AT2G39800 | -0.94 | **$1.48\times10^{-2}$** | $7.43\times10^{-2}$ | -1.25 | **$9.96\times10^{-4}$** | **$6.05\times10^{-3}$** | 0.08 | $7.51\times10^{-1}$ | $8.59\times10^{-1}$ | 1.02 | **$2.79\times10^{-2}$** | $6.13\times10^{-1}$ | *P5CS1* | Delta1-pyrroline-5-carboxylate synthase 1 | -18.4 | **$3.94\times10^{-3}$** | Promoter-TSS (-332) | 9.3 | **$7.00\times10^{-3}$** | Intron (1408) |
| AT3G55940 | -0.29 | $3.79\times10^{-1}$ | $6.35\times10^{-1}$ | -0.20 | $5.02\times10^{-1}$ | $6.71\times10^{-1}$ | 0.73 | **$8.02\times10^{-3}$** | **$3.09\times10^{-2}$** | 1.02 | **$2.98\times10^{-3}$** | $1.92\times10^{-1}$ | *PLC7* | Phosphoinositide-specific phospholipase C family protein | | | | | | |
| AT5G01520 | -0.21 | $5.11\times10^{-1}$ | $7.41\times10^{-1}$ | -0.46 | **$3.42\times10^{-2}$** | $9.97\times10^{-2}$ | 0.81 | **$3.39\times10^{-4}$** | **$2.07\times10^{-3}$** | 1.02 | **$9.58\times10^{-3}$** | $3.61\times10^{-1}$ | *AIRP2* | ABA Insensitive RING Protein 2 | | | | 7.0 | **$3.21\times10^{-3}$** | Intron (719) |
| AT5G63130 | -1.45 | **$2.25\times10^{-7}$** | **$6.43\times10^{-6}$** | -1.14 | **$1.17\times10^{-6}$** | **$1.87\times10^{-5}$** | -0.44 | $6.68\times10^{-2}$ | $1.65\times10^{-1}$ | 1.01 | **$3.07\times10^{-3}$** | $1.94\times10^{-1}$ | | Octicosapeptide/Phox/Bem1p family protein | | | | | | |
| AT1G52030 | 0.35 | $3.87\times10^{-1}$ | $6.43\times10^{-1}$ | 0.30 | $3.86\times10^{-1}$ | $5.68\times10^{-1}$ | 1.36 | **$3.32\times10^{-6}$** | **$3.30\times10^{-5}$** | 1.00 | **$3.30\times10^{-2}$** | $6.61\times10^{-1}$ | *MBP2* | Myrosinase-binding protein 2 | | | | | | |
| AT1G22570 | 0.32 | $1.87\times10^{-1}$ | $4.22\times10^{-1}$ | 0.49 | $5.44\times10^{-2}$ | $1.41\times10^{-1}$ | 1.32 | **$5.62\times10^{-15}$** | **$2.16\times10^{-13}$** | 1.00 | **$2.19\times10^{-6}$** | **$1.14\times10^{-3}$** | | Major facilitator superfamily protein | | | | 5.9 | **$1.49\times10^{-3}$** | TTS-intergenic space (3196) |
| AT1G04570 | -1.01 | **$3.63\times10^{-3}$** | **$2.53\times10^{-2}$** | -1.36 | **$8.06\times10^{-5}$** | **$7.42\times10^{-4}$** | -0.01 | $9.91\times10^{-1}$ | $9.97\times10^{-1}$ | 1.00 | **$2.38\times10^{-2}$** | $5.70\times10^{-1}$ | | Similar to plastid solute transporters | | | | | | |
| AT3G50400 | -0.60 | $1.21\times10^{-1}$ | $3.23\times10^{-1}$ | -0.70 | $7.02\times10^{-2}$ | $1.71\times10^{-1}$ | 0.41 | $3.17\times10^{-1}$ | $5.04\times10^{-1}$ | 1.00 | **$2.93\times10^{-2}$** | $6.26\times10^{-1}$ | | GDSL-motif esterase/acyltransferase/lipase | | | | 35.7 | **$1.53\times10^{-5}$** | Intron (587) |
| AT1G01570 | 0.92 | $7.47\times10^{-2}$ | $2.35\times10^{-1}$ | 0.29 | $6.51\times10^{-1}$ | $7.87\times10^{-1}$ | -0.08 | $7.54\times10^{-1}$ | $8.61\times10^{-1}$ | -1.00 | **$4.98\times10^{-2}$** | $7.86\times10^{-1}$ | | Transferring glycosyl group transferase | | | | | | |
| AT2G32030 | -0.41 | $3.61\times10^{-1}$ | $6.20\times10^{-1}$ | -0.14 | $7.68\times10^{-1}$ | $8.67\times10^{-1}$ | -1.40 | **$1.31\times10^{-5}$** | **$1.15\times10^{-4}$** | -1.01 | **$3.84\times10^{-2}$** | $7.02\times10^{-1}$ | | Acyl-CoA N-acyltransferases (NAT) superfamily protein | | | | 22.1 | **$2.91\times10^{-6}$** | Promoter-TSS (-500) |
| AT3G48700 | -0.43 | $2.28\times10^{-1}$ | $4.78\times10^{-1}$ | -0.89 | **$1.21\times10^{-2}$** | **$4.43\times10^{-2}$** | -1.44 | **$6.80\times10^{-13}$** | **$2.03\times10^{-11}$** | -1.01 | **$8.37\times10^{-3}$** | $3.39\times10^{-1}$ | *CXE13* | Carboxyesterase 13 | | | | | | |
| AT5G04080 | 0.31 | $1.46\times10^{-1}$ | $3.63\times10^{-1}$ | -0.05 | $8.93\times10^{-1}$ | $9.44\times10^{-1}$ | -0.69 | **$6.79\times10^{-4}$** | **$3.79\times10^{-3}$** | -1.02 | **$1.70\times10^{-5}$** | **$4.63\times10^{-3}$** | | Cysteine-rich TM module stress tolerance protein | | | | | | |
| AT3G53280 | 1.38 | **$1.80\times10^{-3}$** | **$1.45\times10^{-2}$** | 0.73 | $8.60\times10^{-2}$ | $1.99\times10^{-1}$ | 0.36 | **$2.08\times10^{-2}$** | $6.66\times10^{-2}$ | -1.03 | **$2.28\times10^{-2}$** | $5.62\times10^{-1}$ | *CYP71B5* | Cytochrome p45 71b5 | | | | | | |
| AT1G58590 | -0.15 | $7.87\times10^{-1}$ | $9.07\times10^{-1}$ | -0.11 | $8.18\times10^{-1}$ | $8.99\times10^{-1}$ | -1.18 | **$4.36\times10^{-3}$** | **$1.84\times10^{-2}$** | -1.04 | **$2.00\times10^{-2}$** | $5.27\times10^{-1}$ | | Non coding RNA | | | | | | |
| AT1G16000 | 0.62 | **$2.15\times10^{-3}$** | **$1.67\times10^{-2}$** | 0.32 | $3.56\times10^{-1}$ | $5.38\times10^{-1}$ | -0.43 | **$2.74\times10^{-2}$** | $8.31\times10^{-2}$ | -1.06 | **$3.31\times10^{-6}$** | **$1.42\times10^{-3}$** | *OEP9.1* | Outer envelope protein 9.1 | | | | 6.0 | **$4.80\times10^{-3}$** | TTS (1018) |
| AT1G44010 | 3.94 | **$2.11\times10^{-10}$** | **$1.21\times10^{-8}$** | 3.52 | **$2.56\times10^{-9}$** | **$7.19\times10^{-8}$** | 2.88 | **$1.94\times10^{-9}$** | **$3.55\times10^{-8}$** | -1.07 | **$3.80\times10^{-2}$** | $7.00\times10^{-1}$ | | Transmembrane protein | | | | 12.6 | **$1.18\times10^{-3}$** | Intergenic (-1971) |
| AT5G52547 | 0.14 | $7.39\times10^{-1}$ | $8.85\times10^{-1}$ | 0.17 | $6.49\times10^{-1}$ | $7.86\times10^{-1}$ | -0.92 | **$3.60\times10^{-3}$** | **$1.58\times10^{-2}$** | -1.07 | **$6.51\times10^{-3}$** | $2.97\times10^{-1}$ | | Hypothetical protein | | | | | | |
| AT1G60050 | -0.30 | $4.96\times10^{-1}$ | $7.30\times10^{-1}$ | 0.52 | $2.61\times10^{-1}$ | $4.37\times10^{-1}$ | -1.35 | **$4.01\times10^{-4}$** | **$2.40\times10^{-3}$** | -1.07 | **$4.27\times10^{-2}$** | $7.39\times10^{-1}$ | *UMAMIT35* | Usually multiple acids move in and out Transporters 35 | | | | | | |
| AT1G64065 | 0.06 | $9.10\times10^{-1}$ | $9.65\times10^{-1}$ | -0.13 | $7.37\times10^{-1}$ | $8.46\times10^{-1}$ | -1.01 | **$4.76\times10^{-4}$** | **$2.78\times10^{-3}$** | -1.08 | **$1.41\times10^{-2}$** | $4.46\times10^{-1}$ | | LEA hydroxyproline-rich glycoprotein family | 59.6 | **$3.32\times10^{-3}$** | Promoter (-3404) | 6.0 | **$3.37\times10^{-3}$** | Promoter (-4104) |
| AT1G68238 | 0.03 | $9.09\times10^{-1}$ | $9.65\times10^{-1}$ | -0.04 | $9.12\times10^{-1}$ | $9.55\times10^{-1}$ | -1.04 | **$4.26\times10^{-5}$** | **$3.29\times10^{-4}$** | -1.08 | **$6.40\times10^{-5}$** | **$1.28\times10^{-2}$** | | Transmembrane protein | | | | | | |
| AT1G66400 | 0.38 | $3.00\times10^{-1}$ | $5.60\times10^{-1}$ | 0.42 | $1.35\times10^{-1}$ | $2.75\times10^{-1}$ | -0.69 | $8.29\times10^{-2}$ | $1.95\times10^{-1}$ | -1.09 | **$2.13\times10^{-2}$** | $5.45\times10^{-1}$ | *CML23* | Calmodulin like 23 | | | | | | |
| AT4G25190 | -1.17 | **$1.95\times10^{-3}$** | **$1.54\times10^{-2}$** | -0.68 | $1.04\times10^{-1}$ | $2.28\times10^{-1}$ | -2.26 | **$7.17\times10^{-9}$** | **$1.17\times10^{-7}$** | -1.10 | **$2.02\times10^{-3}$** | $1.51\times10^{-1}$ | *QWRF7* | QWRF domain containing 7 | | | | | | |

| | | | | | | | | | | | | | | | | | | | | | | | |
|---|---|---|---|---|---|---|---|---|---|---|---|---|---|---|---|---|---|---|---|---|---|---|---|
| AT2G21195 | 0.51 | $1.49\times10^{-1}$ | $3.67\times10^{-1}$ | 0.14 | $6.63\times10^{-1}$ | $7.95\times10^{-1}$ | -0.59 | **$2.05\times10^{-2}$** | $6.57\times10^{-2}$ | -1.11 | **$2.93\times10^{-3}$** | $1.90\times10^{-1}$ | | Hypothetical protein | | | | | | | | | |
| AT5G59330 | -0.40 | $3.68\times10^{-1}$ | $6.26\times10^{-1}$ | -0.67 | $1.02\times10^{-1}$ | $2.25\times10^{-1}$ | -1.51 | **$1.70\times10^{-5}$** | **$1.45\times10^{-4}$** | -1.12 | **$7.11\times10^{-3}$** | $3.10\times10^{-1}$ | | Bifunctional inhibitor/lipid-transfer protein/seed storage 2S albumin superfamily protein | | | | | | | | | |
| AT2G45860 | 0.54 | **$4.14\times10^{-2}$** | $1.56\times10^{-1}$ | 0.34 | $3.68\times10^{-1}$ | $5.50\times10^{-1}$ | -0.57 | $5.50\times10^{-2}$ | $1.43\times10^{-1}$ | -1.12 | **$1.97\times10^{-4}$** | **$2.92\times10^{-2}$** | | Hypothetical protein | | | | | | | | | |
| AT5G37770 | 0.50 | $2.02\times10^{-1}$ | $4.44\times10^{-1}$ | 0.61 | **$1.15\times10^{-2}$** | **$4.26\times10^{-2}$** | -0.66 | **$7.34\times10^{-4}$** | **$4.06\times10^{-3}$** | -1.17 | **$5.84\times10^{-3}$** | $2.82\times10^{-1}$ | *TCH2* | TOUCH 2 | | | | | | | 15.6 | **$7.05\times10^{-4}$** | Promoter-TSS (-632) |
| AT2G29440 | 0.66 | $6.52\times10^{-2}$ | $2.14\times10^{-1}$ | 0.02 | $9.52\times10^{-1}$ | $9.77\times10^{-1}$ | -0.51 | $1.13\times10^{-1}$ | $2.45\times10^{-1}$ | -1.17 | **$2.00\times10^{-4}$** | **$2.92\times10^{-2}$** | *GSTU6* | Glutathione S-transferase tau 6 | | | | | | | | | |
| AT1G31310 | 0.20 | $5.62\times10^{-1}$ | $7.76\times10^{-1}$ | -0.06 | $8.66\times10^{-1}$ | $9.27\times10^{-1}$ | -0.98 | **$3.97\times10^{-3}$** | **$1.70\times10^{-2}$** | -1.18 | **$1.60\times10^{-3}$** | $1.25\times10^{-1}$ | | Hydroxyproline-rich glycoprotein family protein | | | | | | | | | |
| AT2G20835 | 0.85 | $6.03\times10^{-2}$ | $2.03\times10^{-1}$ | 1.03 | **$1.83\times10^{-2}$** | $6.14\times10^{-2}$ | -0.33 | $3.79\times10^{-1}$ | $5.66\times10^{-1}$ | -1.18 | **$2.52\times10^{-3}$** | $1.73\times10^{-1}$ | | Hypothetical protein | | | | | | | 9.0 | **$1.30\times10^{-3}$** | Promoter-TSS (-412) |
| AT1G80240 | -3.47 | **$1.76\times10^{-13}$** | **$2.03\times10^{-11}$** | -2.67 | **$9.00\times10^{-6}$** | **$1.12\times10^{-4}$** | -4.64 | **$7.98\times10^{-43}$** | **$1.83\times10^{-40}$** | -1.19 | **$4.66\times10^{-3}$** | $2.46\times10^{-1}$ | *DGR1* | DUF642 L-GalL responsive gene 1 | | | | | | | 5.6 | **$4.49\times10^{-3}$** | Intron (458) |
| AT5G57785 | -0.02 | $9.45\times10^{-1}$ | $9.80\times10^{-1}$ | -0.24 | $6.33\times10^{-1}$ | $7.73\times10^{-1}$ | -1.23 | **$7.71\times10^{-8}$** | **$1.04\times10^{-6}$** | -1.22 | **$9.43\times10^{-6}$** | **$2.94\times10^{-3}$** | | Hypothetical protein | | | | | | | | | |
| AT5G38200 | -4.13 | **$5.30\times10^{-12}$** | **$4.34\times10^{-10}$** | -3.54 | **$2.13\times10^{-9}$** | **$6.09\times10^{-8}$** | -5.33 | **$8.63\times10^{-90}$** | **$4.16\times10^{-86}$** | -1.22 | **$4.46\times10^{-2}$** | $7.49\times10^{-1}$ | | Class I glutamine amidotransferase-like superfamily protein | | | | | | | 9.0 | **$2.20\times10^{-4}$** | Intron (1275) |
| AT3G18010 | -0.14 | $6.21\times10^{-1}$ | $8.15\times10^{-1}$ | -0.12 | $6.10\times10^{-1}$ | $7.57\times10^{-1}$ | -1.36 | **$3.97\times10^{-9}$** | **$6.82\times10^{-8}$** | -1.23 | **$1.41\times10^{-5}$** | **$3.97\times10^{-3}$** | *WOX1* | WUSCHEL related homeobox 1 | | | | | | | | | |
| AT2G14580 | 0.28 | $4.50\times10^{-1}$ | $6.95\times10^{-1}$ | 0.46 | $3.53\times10^{-1}$ | $5.35\times10^{-1}$ | -1.02 | **$9.10\times10^{-3}$** | **$3.40\times10^{-2}$** | -1.31 | **$3.16\times10^{-4}$** | **$4.13\times10^{-2}$** | *PRB1* | Basic pathogenesis-related protein 1 | | | | | | | | | |
| AT5G47230 | -0.67 | $2.98\times10^{-1}$ | $5.58\times10^{-1}$ | -0.28 | $5.42\times10^{-1}$ | $7.02\times10^{-1}$ | -1.97 | **$7.44\times10^{-18}$** | **$3.95\times10^{-16}$** | -1.33 | **$4.89\times10^{-2}$** | $7.79\times10^{-1}$ | *ERF5* | Ethylene responsive element binding factor 5 | | | | | | | | | |
| AT5G60780 | 0.51 | $1.21\times10^{-1}$ | $3.22\times10^{-1}$ | 0.21 | $5.13\times10^{-1}$ | $6.80\times10^{-1}$ | -0.87 | **$1.37\times10^{-2}$** | **$4.75\times10^{-2}$** | -1.39 | **$1.61\times10^{-4}$** | **$2.57\times10^{-2}$** | *NRT2.3* | Nitrate transporter 2.3 | | | | | | | | | |
| AT3G22210 | 0.92 | $1.25\times10^{-1}$ | $3.29\times10^{-1}$ | 0.52 | $3.98\times10^{-1}$ | $5.79\times10^{-1}$ | -0.47 | $3.09\times10^{-1}$ | $4.97\times10^{-1}$ | -1.41 | **$9.03\times10^{-3}$** | $3.53\times10^{-1}$ | | Transmembrane protein | | | | | | | 7.9 | **$3.63\times10^{-4}$** | Exon (216) |
| AT3G59930 | -1.73 | **$4.63\times10^{-3}$** | **$3.07\times10^{-2}$** | -0.50 | $4.59\times10^{-1}$ | $6.35\times10^{-1}$ | -3.19 | **$1.31\times10^{-24}$** | **$1.41\times10^{-22}$** | -1.49 | **$2.13\times10^{-2}$** | $5.45\times10^{-1}$ | *DEFL206* | Defensin-like protein 206 | | | | | | | | | |
| AT2G30766 | -0.03 | $9.53\times10^{-1}$ | $9.83\times10^{-1}$ | -0.14 | $8.57\times10^{-1}$ | $9.21\times10^{-1}$ | -1.57 | **$3.32\times10^{-15}$** | **$1.32\times10^{-13}$** | -1.55 | **$3.33\times10^{-3}$** | $2.03\times10^{-1}$ | | Hypothetical protein | | | | | | | | | |
| AT1G47395 | 0.22 | $7.28\times10^{-1}$ | $8.80\times10^{-1}$ | -0.32 | $7.10\times10^{-1}$ | $8.27\times10^{-1}$ | -1.38 | **$8.78\times10^{-12}$** | **$2.26\times10^{-10}$** | -1.60 | **$1.48\times10^{-2}$** | $4.55\times10^{-1}$ | | Hypothetical protein | | | | -42.3 | **$6.48\times10^{-3}$** | Intergenic (-1047) | | | |
| AT4G21870 | 0.97 | $1.35\times10^{-1}$ | $3.46\times10^{-1}$ | 0.98 | **$1.16\times10^{-3}$** | **$6.80\times10^{-3}$** | -0.61 | $1.01\times10^{-1}$ | $2.25\times10^{-1}$ | -1.61 | **$2.48\times10^{-2}$** | $5.81\times10^{-1}$ | | HSP20-like chaperone | | | | | | | | | |
| AT2G22122 | 0.39 | $5.80\times10^{-1}$ | $7.89\times10^{-1}$ | 0.56 | $1.91\times10^{-1}$ | $3.53\times10^{-1}$ | -1.23 | **$6.07\times10^{-5}$** | **$4.52\times10^{-4}$** | -1.64 | **$3.37\times10^{-2}$** | $6.68\times10^{-1}$ | | Hypothetical protein; | | | | | | | | | |
| AT5G43580 | 1.94 | **$5.05\times10^{-5}$** | **$7.52\times10^{-4}$** | 2.01 | **$7.50\times10^{-13}$** | **$4.20\times10^{-11}$** | 0.28 | $5.16\times10^{-1}$ | $6.88\times10^{-1}$ | -1.68 | **$3.53\times10^{-3}$** | $2.06\times10^{-1}$ | *UPI* | Unusual serine protease inhibitor | | | | | | | | | |
| AT3G44260 | 0.59 | $4.80\times10^{-1}$ | $7.18\times10^{-1}$ | 1.18 | $5.10\times10^{-2}$ | $1.35\times10^{-1}$ | -1.09 | **$4.60\times10^{-7}$** | **$5.42\times10^{-6}$** | -1.69 | **$4.99\times10^{-2}$** | $7.87\times10^{-1}$ | *CAF1a* | CCR4- associated factor 1a | | | | | | | 9.9 | **$3.20\times10^{-3}$** | Exon (995) |
| AT1G47400 | 0.02 | $9.78\times10^{-1}$ | $9.95\times10^{-1}$ | -0.45 | $6.06\times10^{-1}$ | $7.53\times10^{-1}$ | -1.77 | **$4.40\times10^{-8}$** | **$6.21\times10^{-7}$** | -1.80 | **$1.46\times10^{-2}$** | $4.50\times10^{-1}$ | | Hypothetical protein | | | | | | | | | |
| AT1G20390 | -0.54 | $2.37\times10^{-1}$ | $4.90\times10^{-1}$ | -0.68 | **$1.18\times10^{-2}$** | **$4.34\times10^{-2}$** | -2.37 | **$1.32\times10^{-15}$** | **$5.43\times10^{-14}$** | -1.84 | **$1.94\times10^{-4}$** | **$2.92\times10^{-2}$** | | Transposable element gene | | | | | | | | | |
| AT1G64160 | 0.08 | $9.35\times10^{-1}$ | $9.76\times10^{-1}$ | 0.24 | $7.61\times10^{-1}$ | $8.62\times10^{-1}$ | -1.89 | **$8.34\times10^{-5}$** | **$6.04\times10^{-4}$** | -2.00 | **$4.82\times10^{-2}$** | $7.76\times10^{-1}$ | *DIR5* | Dirigent protein 5 | | | | | | | | | |
| AT4G24570 | 0.82 | $4.03\times10^{-1}$ | $6.56\times10^{-1}$ | 1.82 | **$3.15\times10^{-2}$** | $9.35\times10^{-2}$ | -1.45 | **$3.08\times10^{-8}$** | **$4.47\times10^{-7}$** | -2.29 | **$2.72\times10^{-2}$** | $6.07\times10^{-1}$ | *DIC2* | Dicarboxylate carrier 2 | | | | | | | | | |
| AT1G13609 | 1.46 | $8.82\times10^{-2}$ | $2.61\times10^{-1}$ | 1.48 | $2.57\times10^{-1}$ | $4.32\times10^{-1}$ | -0.91 | **$1.48\times10^{-4}$** | **$1.00\times10^{-3}$** | -2.38 | **$7.33\times10^{-3}$** | $3.14\times10^{-1}$ | | Defensin-like (DEFL) family protein | | | | | | | | | |
| AT5G33355 | -3.31 | **$3.46\times10^{-19}$** | **$1.03\times10^{-16}$** | -2.32 | **$1.54\times10^{-3}$** | **$8.59\times10^{-3}$** | -6.08 | **$5.70\times10^{-63}$** | **$5.50\times10^{-60}$** | -2.75 | **$2.28\times10^{-6}$** | **$1.14\times10^{-3}$** | *DEFL207* | Defensin like protein 207 | 14.0 | **$9.33\times10^{-3}$** | Promoter (-1199) | | | | 10.4 | **$4.24\times10^{-6}$** | Promoter (-1199) |

The results of a differential gene expression analysis of 12-d-old seedlings of *Arabidopsis thaliana*, ecotype: Columbia (Col-0), grown in 1/4-Hoagland's solution in hydroponics and then exposed for three days to a 5% PEG-induced first drought (D1) versus plants growing continuously for 15 days in hydroponics (watered), and plants exposed to another three-days second drought (D2) versus the first drought (D1), separated by a recovery interval (R1) of three days, are presented. Three biological replicates, each with 10 seedlings, were used for the transcriptome analysis, and Fisher's exact test *P*-values, as well as the Benjamini-Hochberg *false discovery rate* (*FDR*), are presented. Significantly up- or down-regulated genes in D2 versus D1 with $\log_2$(fold-change) >1 or <-1 are shown ($P < 0.05$) along with Arabidopsis Genome Initiative (AGI) code, gene symbols, and short description. The differentially methylated regions (DMRs) associated with the memory genes in the CpG, CHG, and CHH (H = A/C/T) sequence contexts are shown as the methylation difference (MD) between D2 versus D1-exposed Col-0 seedlings, *q*-values ($q < 0.01$), and relative position in base pairs (bp) with respect to the transcription start site (TSS) of each gene. Minus (-) denotes a DMR upstream to the TSS, and plus (+) denotes a DMR downstream to the TSS. TTS refers to the transcription termination site.

### Table S2. *Arabidopsis thaliana* genes showing transcriptional memory under recurrent drought stress in hydroponics

| Section | Drought 1 vs Watered | | | Recovery vs Watered | | | Drought 2 vs Watered | | | Drought 2 vs Drought 1 | | | | | CpG methylation difference (D2/D1) in Col-0 | | | CHH methylation difference (D2/D1) in Col-0 | | |
|---|---|---|---|---|---|---|---|---|---|---|---|---|---|---|---|---|---|---|---|---|
| AGI code | $Log_2FC$ | *P*-value | *FDR* | $Log_2FC$ | *P*-value | *FDR* | $Log_2FC$ | *P*-value | *FDR* | $Log_2FC$ | *P*-value | *FDR* | Gene symbol | Short Description | MD | *q*-value | Methylation Position (bp) relative to TSS | MD | *q*-value | Methylation Position (bp) relative to TSS |
| **Superinduced (+/+) genes** | | | | | | | | | | | | | | | | | | | | |
| AT3G02480 | 4.140 | **1.43 × $10^{-3}$** | **1.20 × $10^{-2}$** | 2.34 | **4.10 × $10^{-2}$** | 1.14 × $10^{-1}$ | 6.63 | **1.28 × $10^{-32}$** | **1.24 × $10^{-28}$** | 2.49 | **3.84 × $10^{-2}$** | 7.02 × $10^{-1}$ | | Late embryogenesis abundant protein (LEA) family protein | | | | | | |
| AT1G64110 | 3.052 | **1.10 × $10^{-3}$** | **9.78 × $10^{-3}$** | 2.30 | **1.83 × $10^{-2}$** | 6.16 × $10^{-2}$ | 5.35 | **4.34 × $10^{-23}$** | **3.95 × $10^{-21}$** | 2.30 | **1.06 × $10^{-2}$** | 3.79 × $10^{-1}$ | *DAA1* | DUO1-activated ATPase 1 | | | | 8.76 | **9.74× $10^{-4}$** | Intron (332) |
| AT5G66400 | 2.081 | **1.35 × $10^{-2}$** | 6.93 × $10^{-2}$ | 0.91 | 2.81 × $10^{-1}$ | 4.60 × $10^{-1}$ | 4.25 | **1.98 × $10^{-37}$** | **6.82 × $10^{-35}$** | 2.17 | **1.59 × $10^{-2}$** | 4.70 × $10^{-1}$ | *RAB18* | Responsive to ABA 18 (dehydrin) | 26.79 | **3.84× $10^{-4}$** | Promoter (-104) | 6.07 | **6.74× $10^{-4}$** | Promoter (-9) |
| AT2G34655 | 1.222 | **1.61 × $10^{-2}$** | 7.88 × $10^{-2}$ | 1.95 | **2.77 × $10^{-3}$** | **1.38 × $10^{-2}$** | 3.37 | **3.75 × $10^{-25}$** | **4.16 × $10^{-23}$** | 2.13 | **1.06 × $10^{-6}$** | **6.59 × $10^{-4}$** | | Hypothetical protein | | | | | | |
| AT2G16005 | 4.535 | **9.27 × $10^{-10}$** | **4.62 × $10^{-8}$** | 4.45 | **3.29 × $10^{-16}$** | **3.06 × $10^{-14}$** | 6.60 | **9.42 × $10^{-44}$** | **4.66 × $10^{-41}$** | 2.02 | **5.44 × $10^{-6}$** | **1.98 × $10^{-3}$** | | MD-2-related lipid recognition domain-containing protein | | | | | | |
| AT1G52040 | 1.175 | **1.11 × $10^{-4}$** | **1.46 × $10^{-3}$** | 1.12 | **3.01 × $10^{-6}$** | **4.26 × $10^{-5}$** | 2.57 | **3.02 × $10^{-44}$** | **1.57 × $10^{-41}$** | 1.39 | **5.07 × $10^{-6}$** | **1.92 × $10^{-3}$** | *MBP1* | Myrosinase-binding protein 1 | | | | | | |
| AT4G33150 | 2.200 | **8.82 × $10^{-16}$** | **1.60 × $10^{-13}$** | 2.04 | **1.11 × $10^{-29}$** | **4.95 × $10^{-27}$** | 3.53 | **1.53 × $10^{-33}$** | **3.73 × $10^{-31}$** | 1.33 | **3.50 × $10^{-4}$** | **4.40 × $10^{-2}$** | ***LKR*** | Lysine-ketoglutarate reductase/ saccharopine dehydrogenase bifunctional enzyme | | | | 9.30 | **7.71× $10^{-3}$** | Exon (690) |
| AT3G13672 | 1.337 | **1.08 × $10^{-2}$** | 5.85 × $10^{-2}$ | 0.22 | 6.94 × $10^{-1}$ | 8.17 × $10^{-1}$ | 2.52 | **9.51 × $10^{-19}$** | **5.49 × $10^{-17}$** | 1.18 | **2.32 × $10^{-2}$** | 5.64 × $10^{-1}$ | *SINAT6* | Traf-like superfamily protein | | | | | | |
| AT2G18050 | 2.670 | **4.04 × $10^{-15}$** | **6.35 × $10^{-13}$** | 2.68 | **3.53 × $10^{-6}$** | **4.87 × $10^{-5}$** | 3.80 | **8.80 × $10^{-67}$** | **1.70 × $10^{-63}$** | 1.13 | **7.56 × $10^{-6}$** | **2.57 × $10^{-3}$** | ***HIS1-3*** | Histone H1-3 | | | | | | |
| AT4G28940 | 1.507 | **2.54 × $10^{-3}$** | **1.90 × $10^{-2}$** | 1.29 | **6.60 × $10^{-3}$** | **2.76 × $10^{-2}$** | 2.64 | **3.31 × $10^{-8}$** | **4.77 × $10^{-7}$** | 1.12 | **7.32 × $10^{-3}$** | 3.14 × $10^{-1}$ | | Phosphorylase superfamily protein | | | | | | |
| AT4G23680 | 1.306 | **2.02 × $10^{-11}$** | **1.41 × $10^{-9}$** | 1.39 | **2.36 × $10^{-14}$** | **1.66 × $10^{-12}$** | 2.41 | **3.75 × $10^{-36}$** | **1.19 × $10^{-33}$** | 1.10 | **1.05 × $10^{-7}$** | **1.26 × $10^{-4}$** | | Polyketide cyclase/dehydrase and lipid transport superfamily protein | | | | 15.34 | **2.60× $10^{-4}$** | Exon (1348) |
| AT3G57520 | 2.531 | **9.95 × $10^{-12}$** | **7.65 × $10^{-10}$** | 2.33 | **1.73 × $10^{-11}$** | **7.67 × $10^{-10}$** | 3.60 | **1.96 × $10^{-50}$** | **1.45 × $10^{-47}$** | 1.07 | **2.88 × $10^{-3}$** | 1.89 × $10^{-1}$ | *SIP2* | Seed imbibition 2 | | | | | | |
| **Down-regulated in Drought 1 and up-regulated in drought 2 (-/+) genes** | | | | | | | | | | | | | | | | | | | | |
| AT2G39030 | -1.801 | **4.61 × $10^{-3}$** | **3.06 × $10^{-2}$** | -1.45 | 1.45 × $10^{-1}$ | 2.90 × $10^{-1}$ | 2.03 | **3.85 × $10^{-4}$** | **2.31 × $10^{-3}$** | 3.82 | **2.68 × $10^{-18}$** | **5.19 × $10^{-14}$** | *NATA1* | N-acetyltransferase activity 1 | | | | 9.18 | **3.26× $10^{-3}$** | Promoter (-459) |
| AT4G15210 | -2.424 | **1.78 × $10^{-3}$** | **1.43 × $10^{-2}$** | -1.23 | 1.13 × $10^{-1}$ | 2.58 × $10^{-1}$ | 1.09 | **1.13 × $10^{-3}$** | **5.83 × $10^{-3}$** | 3.51 | **2.32 × $10^{-6}$** | **1.14 × $10^{-3}$** | *BAM5* | Beta-amylase 5 | | | | | | |
| AT1G58320 | -3.856 | **1.56 × $10^{-10}$** | **9.32 × $10^{-9}$** | -3.20 | **1.98 × $10^{-7}$** | **3.87 × $10^{-6}$** | -0.86 | 7.34 × $10^{-2}$ | 1.77 × $10^{-1}$ | 2.98 | **5.52 × $10^{-6}$** | **1.98 × $10^{-3}$** | | PLAC8 family protein | | | | 7.33 | **1.07× $10^{-4}$** | Promoter (-407) |
| AT4G31970 | -2.935 | **2.44 × $10^{-5}$** | **3.98 × $10^{-4}$** | -3.30 | **1.03 × $10^{-5}$** | **1.25 × $10^{-4}$** | -0.35 | 5.50 × $10^{-1}$ | 7.14 × $10^{-1}$ | 2.58 | **1.55 × $10^{-3}$** | 1.22 × $10^{-1}$ | *CYP82C2* | Cytochrome p450, family 82, subfamily c, polypeptide 2 | | | | 7.88 | **1.72× $10^{-3}$** | Exon (801) |
| AT2G18193 | -1.396 | **7.10 × $10^{-3}$** | **4.29 × $10^{-2}$** | -0.32 | 5.51 × $10^{-1}$ | 7.11 × $10^{-1}$ | 0.89 | **1.38 × $10^{-3}$** | **6.95 × $10^{-3}$** | 2.27 | **4.89 × $10^{-5}$** | **1.10 × $10^{-2}$** | | P-loop containing nucleoside triphosphate hydrolases superfamily protein | | | | | | |
| AT2G45570 | -1.107 | **9.46 × $10^{-3}$** | 5.32 × $10^{-2}$ | -1.83 | **2.21 × $10^{-5}$** | **2.43 × $10^{-4}$** | 1.16 | **1.09 × $10^{-2}$** | **3.96 × $10^{-2}$** | 2.26 | **8.95 × $10^{-5}$** | **1.63 × $10^{-2}$** | *CYP76C2* | Cytochrome p450, family 76, subfamily c, polypeptide 2 | | | | 8.18 | **3.18× $10^{-3}$** | Exon (322) |
| AT5G52310 | -1.182 | **2.95 × $10^{-2}$** | 1.24 × $10^{-1}$ | -1.96 | **1.92 × $10^{-4}$** | **1.54 × $10^{-3}$** | 0.94 | **4.12 × $10^{-3}$** | **1.76 × $10^{-2}$** | 2.12 | **1.41 × $10^{-3}$** | 1.16 × $10^{-1}$ | *LTI78* | Low-temperature-induced 78 | | | | | | |
| AT1G11070 | -1.043 | **4.00 × $10^{-2}$** | 1.53 × $10^{-1}$ | -1.06 | **4.52 × $10^{-3}$** | **2.03 × $10^{-2}$** | 1.04 | **4.52 × $10^{-4}$** | **2.66 × $10^{-3}$** | 2.07 | **4.45 × $10^{-6}$** | **1.79 × $10^{-3}$** | | Hydroxyproline-rich glycoprotein family protein | | | | | | |

| | | | | | | | | | | | | | | | | | | | | |
|---|---|---|---|---|---|---|---|---|---|---|---|---|---|---|---|---|---|---|---|---|
| AT1G69140 | -2.145 | **4.01 ×** **$10^{-7}$** | **1.08 ×** **$10^{-5}$** | -2.37 | **9.47 ×** **$10^{-17}$** | **9.40 ×** **$10^{-15}$** | -0.15 | 6.31 × $10^{-1}$ | 7.75 × $10^{-1}$ | 2.00 | **3.46 ×** **$10^{-5}$** | **8.36 ×** **$10^{-3}$** | | Pseudogene of hypothetical protein | | | | | | |
| AT1G52890 | -1.064 | **1.72 ×** **$10^{-2}$** | 8.29 × $10^{-2}$ | -1.50 | **4.13 ×** **$10^{-4}$** | **2.90 ×** **$10^{-3}$** | 0.88 | **1.39 ×** **$10^{-2}$** | **4.82 ×** **$10^{-2}$** | 1.95 | **8.72 ×** **$10^{-6}$** | **2.86 ×** **$10^{-3}$** | *NAC019* | NAC domain containing protein 19 | | | | 9.09 | **1.19×** **$10^{-3}$** | TTS-intergenic space (2131) |
| AT4G25480 | -1.543 | **4.87 ×** **$10^{-6}$** | **9.88 ×** **$10^{-5}$** | -2.07 | **7.63 ×** **$10^{-10}$** | **2.42 ×** **$10^{-8}$** | 0.39 | 1.79 × $10^{-1}$ | 3.42 × $10^{-1}$ | 1.93 | **3.91 ×** **$10^{-7}$** | **2.91 ×** **$10^{-4}$** | ***DREB1A*** | Dehydration response element binding 1A | | | | | | |
| AT1G05680 | -3.218 | **1.47 ×** **$10^{-15}$** | **2.56 ×** **$10^{-13}$** | -2.06 | **9.47 ×** **$10^{-4}$** | **5.81 ×** **$10^{-3}$** | -1.32 | **1.16 ×** **$10^{-6}$** | **1.26 ×** **$10^{-5}$** | 1.88 | **8.11 ×** **$10^{-7}$** | **5.61 ×** **$10^{-4}$** | *UGT74E2* | Uridine diphosphate glycosyltransferase 74E2 | | | | | | |
| AT1G53540 | -1.564 | **7.15 ×** **$10^{-3}$** | **4.31 ×** **$10^{-2}$** | -3.50 | **1.36 ×** **$10^{-7}$** | **2.75 ×** **$10^{-6}$** | 0.22 | 5.11 × $10^{-1}$ | 6.84 × $10^{-1}$ | 1.77 | **2.74 ×** **$10^{-3}$** | 1.84 × $10^{-1}$ | | HSP20-like chaperones superfamily protein | | | | | | |
| AT2G39920 | -1.300 | **1.72 ×** **$10^{-2}$** | 8.30 × $10^{-2}$ | -1.19 | **2.75 ×** **$10^{-2}$** | 8.44 × $10^{-2}$ | 0.47 | 2.46 × $10^{-1}$ | 4.28 × $10^{-1}$ | 1.77 | **2.49 ×** **$10^{-4}$** | **3.40 ×** **$10^{-2}$** | | HAD superfamily, subfamily IIIB acid phosphatase | 47.51 | **1.62×** **$10^{-3}$** | TTS (1780) | | | |
| AT5G24770 | -1.023 | **2.44 ×** **$10^{-2}$** | 1.08 × $10^{-1}$ | -0.93 | 5.67 × $10^{-2}$ | 1.46 × $10^{-1}$ | 0.71 | **3.67 ×** **$10^{-7}$** | **4.40 ×** **$10^{-6}$** | 1.72 | **3.27 ×** **$10^{-4}$** | **4.16 ×** **$10^{-2}$** | *VSP2* | Vegetative storage protein 2 | | | | | | |
| AT2G47780 | -1.096 | **7.81 ×** **$10^{-3}$** | **4.60 ×** **$10^{-2}$** | -1.51 | **1.07 ×** **$10^{-7}$** | **2.22 ×** **$10^{-6}$** | 0.52 | 9.48 × $10^{-2}$ | 2.16 × $10^{-1}$ | 1.62 | **1.07 ×** **$10^{-3}$** | 9.50 × $10^{-2}$ | | Rubber elongation factor protein (REF) | | | | 7.08 | **9.93×** **$10^{-4}$** | Exon (511) |
| AT3G43715 | -1.943 | **1.99 ×** **$10^{-3}$** | **1.57 ×** **$10^{-2}$** | -1.21 | **2.17 ×** **$10^{-2}$** | 7.05 × $10^{-2}$ | -0.33 | 2.01 × $10^{-1}$ | 3.72 × $10^{-1}$ | 1.60 | **9.24 ×** **$10^{-3}$** | 3.56 × $10^{-1}$ | | Transposable element gene | | | | | | |
| AT5G49525 | -1.029 | **2.47 ×** **$10^{-2}$** | 1.09 × $10^{-1}$ | -1.13 | **8.99 ×** **$10^{-4}$** | **5.57 ×** **$10^{-3}$** | 0.55 | **2.58 ×** **$10^{-2}$** | 7.92 × $10^{-2}$ | 1.58 | **3.18 ×** **$10^{-4}$** | **4.13 ×** **$10^{-2}$** | | Transmembrane protein | | | | 6.21 | **2.35×** **$10^{-4}$** | Intergenic (-1400) |
| AT2G42530 | -1.767 | **9.43 ×** **$10^{-10}$** | **4.68 ×** **$10^{-8}$** | -1.78 | **4.33 ×** **$10^{-15}$** | **3.39 ×** **$10^{-13}$** | -0.20 | 4.61 × $10^{-1}$ | 6.41 × $10^{-1}$ | 1.57 | **4.80 ×** **$10^{-5}$** | **1.09 ×** **$10^{-2}$** | *COR15B* | Cold regulated 15b | | | | | | |
| AT2G47190 | -2.618 | **6.13 ×** **$10^{-11}$** | **4.02 ×** **$10^{-9}$** | -2.72 | **4.37 ×** **$10^{-8}$** | **9.87 ×** **$10^{-7}$** | -1.07 | **5.06 ×** **$10^{-4}$** | **2.93 ×** **$10^{-3}$** | 1.55 | **9.01 ×** **$10^{-5}$** | **1.63 ×** **$10^{-2}$** | *MYB2* | Myb domain protein 2 | | | | | | |
| AT4G29610 | -1.581 | **3.38 ×** **$10^{-3}$** | **2.39 ×** **$10^{-2}$** | -0.88 | 1.15 × $10^{-1}$ | 2.46 × $10^{-1}$ | -0.06 | 8.94 × $10^{-1}$ | 9.47 × $10^{-1}$ | 1.50 | **1.38 ×** **$10^{-4}$** | **2.28 ×** **$10^{-2}$** | | Cytidine/deoxycytidylate deaminase family protein | | | | 7.21 | **4.11×** **$10^{-4}$** | Promoter (-242) |
| AT2G33380 | -1.298 | **8.29 ×** **$10^{-6}$** | **1.56 ×** **$10^{-4}$** | -1.68 | **2.50 ×** **$10^{-14}$** | **1.76 ×** **$10^{-12}$** | 0.21 | 4.08 × $10^{-1}$ | 5.93 × $10^{-1}$ | 1.50 | **1.04 ×** **$10^{-5}$** | **3.09 ×** **$10^{-3}$** | *RD20* | Responsive to dessication 20 | | | | | | |
| AT1G61800 | -1.676 | **1.73 ×** **$10^{-4}$** | **2.14 ×** **$10^{-3}$** | -1.87 | **3.55 ×** **$10^{-8}$** | **8.14 ×** **$10^{-7}$** | -0.23 | 6.16 × $10^{-1}$ | 7.65 × $10^{-1}$ | 1.45 | **1.52 ×** **$10^{-2}$** | 4.61 × $10^{-1}$ | | Glucose-6-phosphate/phosphate translocator 2 | | | | | | |
| AT5G54190 | -1.544 | **1.84 ×** **$10^{-2}$** | 8.73 × $10^{-2}$ | -1.43 | **3.88 ×** **$10^{-3}$** | **1.80 ×** **$10^{-2}$** | -0.19 | 5.62 × $10^{-1}$ | 7.23 × $10^{-1}$ | 1.35 | **4.13 ×** **$10^{-2}$** | 7.25 × $10^{-1}$ | *PORA* | Protochlorophyllide oxidoreductase a | | | | | | |
| AT1G77450 | -2.358 | **1.29 ×** **$10^{-8}$** | **5.00 ×** **$10^{-7}$** | -2.55 | **2.59 ×** **$10^{-13}$** | **1.57 ×** **$10^{-11}$** | -1.04 | **9.94 ×** **$10^{-6}$** | **8.92 ×** **$10^{-5}$** | 1.32 | **1.37 ×** **$10^{-3}$** | 1.13 × $10^{-1}$ | *NAC032* | NAC domain containing protein 32 | | | | | | |
| AT1G78990 | -1.245 | **1.53 ×** **$10^{-3}$** | **1.27 ×** **$10^{-2}$** | -0.77 | 2.87 × $10^{-1}$ | 4.67 × $10^{-1}$ | 0.01 | 9.90 × $10^{-1}$ | 9.97 × $10^{-1}$ | 1.24 | **1.34 ×** **$10^{-3}$** | 1.11 × $10^{-1}$ | | HXXXD-type acyl-transferase family protein | | | | 5.14 | **9.13×** **$10^{-3}$** | Exon (666) |
| AT1G65890 | -1.327 | **1.66 ×** **$10^{-5}$** | **2.87 ×** **$10^{-4}$** | -0.77 | **1.99 ×** **$10^{-2}$** | 6.58 × $10^{-2}$ | -0.08 | 8.09 × $10^{-1}$ | 8.94 × $10^{-1}$ | 1.24 | **3.19 ×** **$10^{-3}$** | 1.98 × $10^{-1}$ | *AAE12* | Acyl activating enzyme 12 | | | | | | |
| AT1G66700 | -3.804 | **1.17 ×** **$10^{-18}$** | **3.10 ×** **$10^{-16}$** | -3.52 | 3.08 × $10^{-26}$ | **9.57 ×** **$10^{-24}$** | -2.56 | **1.56 ×** **$10^{-28}$** | **2.33 ×** **$10^{-26}$** | 1.24 | **7.38 ×** **$10^{-3}$** | 3.14 × $10^{-1}$ | *PXMT1* | A member of the Arabidopsis SABATH methyltransferase gene family | | | | 12.85 | **9.25×** **$10^{-3}$** | Intergenic (-1385) |
| AT5G58310 | -1.029 | **1.37 ×** **$10^{-3}$** | **1.16 ×** **$10^{-2}$** | -0.71 | **1.53 ×** **$10^{-2}$** | **5.32 ×** **$10^{-2}$** | 0.19 | 4.66 × $10^{-1}$ | 6.45 × $10^{-1}$ | 1.22 | **5.14 ×** **$10^{-5}$** | **1.12 ×** **$10^{-2}$** | *MES18* | Methyl esterase 18 | | | | 6.38 | **6.43×** **$10^{-4}$** | Promoter (-167) |
| AT1G29395 | -1.379 | **1.22 ×** **$10^{-8}$** | **4.75 ×** **$10^{-7}$** | -1.82 | **5.82 ×** **$10^{-23}$** | **1.20 ×** **$10^{-20}$** | -0.19 | 4.77 × $10^{-1}$ | 6.55 × $10^{-1}$ | 1.19 | **2.39 ×** **$10^{-4}$** | **3.32 ×** **$10^{-2}$** | *COR413IM1* | Cold regulated 314 inner membrane 1 | 69.44 | **7.38×** **$10^{-3}$** | Promoter (-1885) | | | |
| AT4G30650 | -1.498 | **8.82 ×** **$10^{-6}$** | **1.65 ×** **$10^{-4}$** | -1.87 | **2.10 ×** **$10^{-23}$** | **4.76 ×** **$10^{-21}$** | -0.39 | **1.16 ×** **$10^{-2}$** | **4.15 ×** **$10^{-2}$** | 1.10 | **1.57 ×** **$10^{-3}$** | 1.23 × $10^{-1}$ | | Low temperature and salt responsive protein family | | | | | | |
| AT2G29460 | -1.706 | **9.51 ×** **$10^{-4}$** | **8.71 ×** **$10^{-3}$** | -2.46 | **1.10 ×** **$10^{-3}$** | **6.54 ×** **$10^{-3}$** | -0.61 | 9.90 × $10^{-2}$ | 2.23 × $10^{-1}$ | 1.09 | **1.45 ×** **$10^{-2}$** | 4.50 × $10^{-1}$ | *GSTU4* | Glutathione s-transferase tau 4 | | | | | | |
| AT1G19250 | -1.949 | **8.01 ×** **$10^{-5}$** | **1.11 ×** **$10^{-3}$** | -2.07 | **2.62 ×** **$10^{-6}$** | **3.77 ×** **$10^{-5}$** | -0.86 | **1.26 ×** **$10^{-2}$** | **4.47 ×** **$10^{-2}$** | 1.08 | **1.44 ×** **$10^{-2}$** | 4.50 × $10^{-1}$ | *FMO1* | Flavin-dependent monooxygenase 1 | | | | | | |
| AT1G55990 | -2.122 | **3.68 ×** **$10^{-8}$** | **1.29 ×** **$10^{-6}$** | -1.80 | **2.06 ×** **$10^{-6}$** | **3.08 ×** **$10^{-5}$** | -1.03 | **5.40 ×** **$10^{-4}$** | **3.10 ×** **$10^{-3}$** | 1.08 | **6.53 ×** **$10^{-3}$** | 2.97 × $10^{-1}$ | | Glycine-rich protein | | | | | | |
| AT2G29500 | -2.550 | **1.66 ×** **$10^{-7}$** | **4.87 ×** **$10^{-6}$** | -3.09 | **9.61 ×** **$10^{-18}$** | **1.06 ×** **$10^{-15}$** | -1.50 | **7.38 ×** **$10^{-9}$** | **1.20 ×** **$10^{-7}$** | 1.05 | **1.99 ×** **$10^{-2}$** | 5.25 × $10^{-1}$ | | HSP20-like chaperones superfamily protein | | | | | | |
| AT3G04040 | -1.412 | **2.19 ×** **$10^{-4}$** | **2.61 ×** **$10^{-3}$** | -1.20 | **2.26 ×** **$10^{-3}$** | **1.17 ×** **$10^{-2}$** | -0.40 | 2.74 × $10^{-1}$ | 4.59 × $10^{-1}$ | 1.02 | **3.24 ×** **$10^{-2}$** | 6.55 × $10^{-1}$ | | Transmembrane protein | | | | | | |
| AT5G63130 | -1.451 | **2.25 ×** **$10^{-7}$** | **6.43 ×** **$10^{-6}$** | -1.14 | **1.17 ×** **$10^{-6}$** | **1.87 ×** **$10^{-5}$** | -0.44 | 6.68 × $10^{-2}$ | 1.65 × $10^{-1}$ | 1.01 | **3.07 ×** **$10^{-3}$** | 1.94 × $10^{-1}$ | | Octicosapeptide/Phox/Bem1p family protein | | | | | | |
| AT1G04570 | -1.014 | **3.63 ×** **$10^{-3}$** | **2.53 ×** **$10^{-2}$** | -1.36 | **8.06 ×** **$10^{-5}$** | **7.42 ×** **$10^{-4}$** | -0.01 | 9.91 × $10^{-1}$ | 9.97 × $10^{-1}$ | 1.00 | **2.38 ×** **$10^{-2}$** | 5.70 × $10^{-1}$ | | Similar to plastid solute transporters | | | | | | |

**Up-regulated in Drought 1 and down-regulated in drought 2 (+/-) genes**

| | | | | | | | | | | | | | | | | | | | | |
|---|---|---|---|---|---|---|---|---|---|---|---|---|---|---|---|---|---|---|---|---|
| AT3G53280 | 1.377 | **1.80 ×** **$10^{-3}$** | **1.45 ×** **$10^{-2}$** | 0.73 | 8.60 × $10^{-2}$ | 1.99 × $10^{-1}$ | 0.36 | **2.08 ×** **$10^{-2}$** | 6.66 × $10^{-2}$ | -1.03 | **2.28 ×** **$10^{-2}$** | 5.62 × $10^{-1}$ | *CYP71B5* | Cytochrome P450 71B5 | | | | | | |
| AT1G44010 | 3.940 | **2.11 ×** **$10^{-10}$** | **1.21 ×** **$10^{-8}$** | 3.52 | **2.56 ×** **$10^{-9}$** | **7.19 ×** **$10^{-8}$** | 2.88 | **1.94 ×** **$10^{-9}$** | **3.55 ×** **$10^{-8}$** | -1.07 | **3.80 ×** **$10^{-2}$** | 7.00 × $10^{-1}$ | | Transmembrane protein | | | | 12.59 | **1.18×** **$10^{-3}$** | Intergenic (-1971) |
| ATCG00180 | 1.594 | **4.39 ×** **$10^{-3}$** | **2.94 ×** **$10^{-2}$** | 0.66 | 8.75 × $10^{-2}$ | 2.02 × $10^{-1}$ | 0.31 | 2.86 × $10^{-1}$ | 4.71 × $10^{-1}$ | -1.29 | **1.50 ×** **$10^{-2}$** | 4.58 × $10^{-1}$ | *rpoC1* | RNA polymerase beta subunit | | | | | | |

| | | | | | | | | | | | | | | | | | | | |
|---|---|---|---|---|---|---|---|---|---|---|---|---|---|---|---|---|---|---|---|
| ATCG00350 | 1.285 | **1.76 × $10^{-2}$** | 8.43 × $10^{-2}$ | -0.26 | 4.62 × $10^{-1}$ | 6.37 × $10^{-1}$ | -0.24 | 3.28 × $10^{-1}$ | 5.16 × $10^{-1}$ | -1.54 | **6.49 × $10^{-3}$** | 2.97 × $10^{-1}$ | *psaA* | Photosystem I P700 chlorophyll a apoprotein A1 | | | | | |
| AT5G43580 | 1.945 | **5.05 × $10^{-5}$** | **7.52 × $10^{-4}$** | 2.01 | **7.50 × $10^{-13}$** | **4.20 × $10^{-11}$** | 0.28 | 5.16 × $10^{-1}$ | 6.88 × $10^{-1}$ | -1.68 | **3.53 × $10^{-3}$** | 2.06 × $10^{-1}$ | *UPI* | Serine protease inhibitor, potato inhibitor I-type family protein | | | | | |
| ATCG00340 | 1.505 | **5.04 × $10^{-3}$** | **3.27 × $10^{-2}$** | -0.18 | 6.07 × $10^{-1}$ | 7.54 × $10^{-1}$ | -0.17 | 4.59 × $10^{-1}$ | 6.39 × $10^{-1}$ | -1.69 | **2.53 × $10^{-3}$** | 1.73 × $10^{-1}$ | *psaB* | Photosystem I P700 chlorophyll a apoprotein A2 | | | | | |
| **Progressively down-regulated (-/-) genes** | | | | | | | | | | | | | | | | | | | |
| AT4G25190 | -1.171 | **1.95 × $10^{-3}$** | **1.54 × $10^{-2}$** | -0.68 | 1.04 × $10^{-1}$ | 2.28 × $10^{-1}$ | -2.26 | **7.17 × $10^{-9}$** | **1.17 × $10^{-7}$** | -1.10 | **2.02 × $10^{-3}$** | 1.51 × $10^{-1}$ | *QWRF7* | QWRF motif protein | | | | | |
| AT1G80240 | -3.469 | **1.80× $10^{-13}$** | **2.03 × $10^{-11}$** | -2.67 | **9.00 × $10^{-6}$** | **1.12 × $10^{-4}$** | -4.64 | **7.98 × $10^{-33}$** | **1.83 × $10^{-30}$** | -1.19 | **4.66 × $10^{-3}$** | 2.46 × $10^{-1}$ | *DGR1* | choice-of-anchor C domain protein | | | | 5.59 | **4.49× $10^{-3}$** | Intron (458) |
| AT5G38200 | -4.133 | **5.30× $10^{-12}$** | **4.34 × $10^{-10}$** | -3.54 | **2.13 × $10^{-9}$** | **6.09 × $10^{-8}$** | -5.33 | **8.63 × $10^{-80}$** | **4.16 × $10^{-76}$** | -1.22 | **4.46 × $10^{-2}$** | 7.49 × $10^{-1}$ | | Class I glutamine amidotransferase-like superfamily protein | | | | 9.00 | **2.20× $10^{-4}$** | Intron (1275) |
| AT3G59930 | -1.725 | **4.60× $10^{-3}$** | **3.07 × $10^{-2}$** | -0.50 | 4.59 × $10^{-1}$ | 6.35 × $10^{-1}$ | -3.19 | **1.31 × $10^{-24}$** | **1.41 × $10^{-22}$** | -1.49 | **2.13 × $10^{-2}$** | 5.45 × $10^{-1}$ | | Defensin-like protein 206 | | | | | | |
| AT5G33355 | -3.305 | **3.50× $10^{-19}$** | **1.03 × $10^{-16}$** | -2.32 | **1.54 × $10^{-3}$** | **8.59 × $10^{-3}$** | -6.08 | **5.70 × $10^{-53}$** | **5.50 × $10^{-50}$** | -2.75 | **2.28 × $10^{-6}$** | **1.14 × $10^{-3}$** | | Defensin-like protein 207 | 13.95 | **9.33× $10^{-3}$** | Promoter (-1199) | 10.44 | **4.24× $10^{-6}$** | Promoter (-1199) |

The results of a differential gene expression analysis of 12-d-old seedlings of *Arabidopsis thaliana*, ecotype: Columbia (Col-0), grown in 1/4-Hoagland's solution in hydroponics and then exposed for three days to a 5% PEG-induced first drought (D1) versus plants growing continuously for 15 days in hydroponics (watered), and plants exposed to another three-days second drought (D2) versus the first drought (D1), separated by a recovery interval of three days, are presented. Three biological replicates, each with 10 seedlings, were used for the transcriptome analysis, and Fisher's exact test *P*-values, as well as the Benjamini-Hochberg *false discovery rate* (*FDR*), are presented. Significantly up- or down-regulated genes in D2 versus D1 with log2(fold-change) >1 or <-1 are shown ($P < 0.05$) along with Arabidopsis Genome Initiative (AGI) code, gene symbols, and short description. Differentially methylated regions (DMRs) associated with the memory genes in the CpG, CHG, and CHH (H = A/C/T) sequence contexts are shown as the methylation difference (MD) between D2 versus D1-exposed Col-0 seedlings, *q*-values ($q < 0.01$), and relative position in base pairs (bp) with respect to the transcription start site (TSS) of each gene. Minus (-) denotes a DMR upstream to the TSS, and plus (+) denotes a DMR downstream to the TSS. TTS refers to the transcription termination site.

**Table S3 Genes associated with the most significant SNPs in an expression genome-wide association study of *LKR* superinduction in *Arabidopsis thaliana* under recurrent droughts**

| Chromosome | Physical position (bp) | GWAS *P*-value | Frequency of low expression allele | RFC *LKR* (D2/D1) of low induction allele | Frequency of high induction allele | RFC *LKR* (D2/D1) of high induction allele | *t*-test *P*-value (segregation of alleles) | Nearest Gene | SNP position with respect to the gene | Gene Symbol | Short Description |
|---|---|---|---|---|---|---|---|---|---|---|---|
| 1 | 593706 | **$3.70\times 10^{-4}$** | 101/T | 3.2 ± 0.6 | 14/A | 5.4 ± 1.9 | $1.90\times 10^{-1}$ | AT1G02720 | Intron | *GATL5* | Galacturonosyltransferase-like 5 |
| 1 | 3916947 | **$5.40\times 10^{-4}$** | 81/G | 2.4 ± 0.3 | 34/A | 5.9 ± 1.8 | **$3.60\times 10^{-3}$** | AT1G11650 | Intron | *RBP45B* | RNA-binding (RRM/RBD/RNP motifs) family protein |
| 1<br>Chr1:3911676 - Chr1:3921016 | 3918247 | **$6.60\times 10^{-4}$** | 87/A | 2.5 ± 0.3 | 28/G | 6.6 ± 2.1 | **$1.10\times 10^{-3}$** | AT1G11650 | 306 | *RBP45B* | RNA-binding (RRM/RBD/RNP motifs) family protein |
| | | | | | | | | AT1G11620 | 6323 | | F-box and associated interaction domains-containing protein |
| | | | | | | | | AT1G11660 | -2809 | *HSP70-16* | Heat shock protein 70-16 |
| 1 | 9632376 | **$4.00\times 10^{-4}$** | 89/C | 2.5 ± 0.2 | 26/T | 6.8 ± 2.3 | **$7.50\times 10^{-4}$** | AT1G27680 | Exon | *APL2* | ADPGLC-PPASE large subunit |
| 1 | 11182719 | **$7.80\times 10^{-4}$** | 93/T | 2.9 ± 0.7 | 22/C | 5.6 ± 1.2 | $5.70\times 10^{-2}$ | AT1G31280 | Exon | *AGO2* | Argonaute 2 |
| 1 | 11463473 | **$9.60\times 10^{-4}$** | 102/G | 3.2 ± 0.6 | 13/T | 5 ± 2 | $3.00\times 10^{-1}$ | AT1G31920 | Exon | | Tetratricopeptide repeat (TPR)-like superfamily protein |
| 1 | 17080042 | **$6.50\times 10^{-4}$** | 101/T | 3.2 ± 0.6 | 14/C | 5 ± 1.8 | $2.90\times 10^{-1}$ | AT1G45150 | Intron | | Alpha-1,6-mannosyl-glycoprotein 2-beta-N-acetylglucosaminyltransferase |
| 1<br>Chr1:17518622 - Chr1:17525577 | 17518622 | **$7.50\times 10^{-4}$** | 77/T | 3 ± 0.8 | 38/C | 4.4 ± 0.8 | $2.50\times 10^{-1}$ | AT1G47640 | Exon | | Seven transmembrane domain protein |
| | | | | | | | | AT1G47655 | -6815 | | Dof-type zinc finger DNA-binding family protein |
| 1<br>Chr1:18711140 - Chr1:18717401 | 18711140 | **$1.80\times 10^{-4}$** | 103/T | 3.1 ± 0.6 | 12/G | 6.8 ± 2.1 | **$3.80\times 10^{-2}$** | AT1G50500 | Intron | *VPS53* | Arabidopsis thaliana VPS53 homolog |
| | | | | | | | | AT1G50510 | -5438 | *PUMY* | Pseudouridine monophosphate glycosylase |
| 1 | 18726721 | **$1.10\times 10^{-4}$** | 101/A | 3 ± 0.6 | 14/G | 6.8 ± 1.8 | **$2.00\times 10^{-2}$** | AT1G50570 | Exon | | Calcium-dependent lipid-binding (CaLB domain) family protein |
| 1<br>Chr1:18728221 - Chr1:18731586 | 18730274 | **$6.20\times 10^{-4}$** | 102/G | 3 ± 0.6 | 13/A | 6.6 ± 2 | **$3.90\times 10^{-2}$** | AT1G50575 | Exon | | Putative lysine decarboxylase family protein |
| | | | | | | | | AT1G50570 | -2708 | | Calcium-dependent lipid-binding (CaLB domain) family protein |
| | | | | | | | | AT1G50580 | -557 | | UDP-Glycosyltransferase superfamily protein |
| 1 | 21240309 | **$2.30\times 10^{-5}$** | 103/A | 3.1 ± 0.6 | 12/C | 6.8 ± 2 | **$3.80\times 10^{-2}$** | AT1G56660 | Exon | | MAEBL domain protein |
| 1 | 21242893 | **$6.00\times 10^{-4}$** | 103/A | 3.2 ± 0.6 | 12/T | 5.3 ± 2.1 | $2.50\times 10^{-1}$ | AT1G56670 | Exon | | GDSL-motif esterase/acyltransferase/lipase |
| 1 | 21243257 | **$6.00\times 10^{-4}$** | 103/T | 3.2 ± 0.6 | 12/A | 5.3 ± 2.1 | $2.50\times 10^{-1}$ | AT1G56670 | Exon | | GDSL-motif esterase/acyltransferase/lipase |
| 1 | 22227483 | **$1.70\times 10^{-4}$** | 98/T | 3.2 ± 0.6 | 17/G | 5.1 ± 1.5 | $2.00\times 10^{-1}$ | AT1G60280 | Exon | *NAC023* | NAC domain containing protein 23 |
| 1 | 22595123 | **$8.30\times 10^{-4}$** | 72/C | 2.2 ± 0.3 | 43/T | 5.5 ± 1.4 | **$3.20\times 10^{-3}$** | AT1G61260 | -137 | | Cotton fiber (DUF761) |
| 1 | 23043312 | **$2.40\times 10^{-4}$** | 103/A | 3.3 ± 0.6 | 12/G | 5 ± 2 | $3.40\times 10^{-1}$ | AT1G62320 | Exon | *OSCA1.4* | ERD (early-responsive to dehydration stress) family protein |
| 1 | 24405454 | **$1.90\times 10^{-6}$** | 104/A | 3 ± 0.6 | 11/G | 7.7 ± 2.2 | **$1.00\times 10^{-2}$** | AT1G65630 | 1088 | *DEG3* | Degp protease 3 |
| 1 | 24405565 | **$2.80\times 10^{-5}$** | 104/A | 3 ± 0.6 | 11/C | 7.2 ± 2.2 | **$2.40\times 10^{-2}$** | AT1G65630 | 977 | *DEG3* | Degp protease 3 |
| 1<br>Chr1:25133155 - Chr1:25142069 | 25142069 | **$4.90\times 10^{-4}$** | 106/A | 2.7 ± 0.3 | 9/C | 12.1 ± 6.1 | **$1.60\times 10^{-6}$** | AT1G67210 | Intron | *ZCCHC8B* | Proline-rich spliceosome-associated (PSP) family protein |
| | | | | | | | | AT1G67190 | 7756 | | F-box/RNI-like superfamily protein |
| 1 | 25143543 | **$6.00\times 10^{-5}$** | 107/A | 2.7 ± 0.3 | 8/T | 13.5 ± 6.7 | **$1.30\times 10^{-7}$** | AT1G67210 | Exon | *ZCCHC8B* | Proline-rich spliceosome-associated (PSP) family protein |

| | | | | | | | | | | | |
|---|---|---|---|---|---|---|---|---|---|---|---|
| | *Chr1:25143543 - Chr1:25145105* | | | | | | | AT1G08913 | -1501 | | Long non-coding RNA |
| 1 | 25603964 | **$9.80\times 10^{-4}$** | 99/G | 3.2 ± 0.6 | 16/A | 4.8 ± 1.6 | $3.30\times 10^{-1}$ | AT1G68320 | Exon | *MYB62* | Myb domain protein 62 |
| 1 | 26442483 | **$2.80\times 10^{-4}$** | 111/C | 3.3 ± 0.6 | 4/T | 7.8 ± 5.7 | $1.30\times 10^{-1}$ | AT1G70210 | Exon | *CYCD1;1* | Cyclin D1 |
| | *Chr1:26442483 - Chr1:26447108* | | | | | | | AT1G70220 | 3847 | | RNA-processing, Lsm domain-containing protein |
| 1 | 26800918 | **$6.40\times 10^{-4}$** | 68/G | 3.1 ± 0.9 | 47/C | 3.9 ± 0.7 | $4.80\times 10^{-1}$ | AT1G71040 | -694 | *LPR2* | Low phosphate root2 |
| 1 | 27861354 | **$3.90\times 10^{-4}$** | 107/C | 3.3 ± 0.6 | 8/A | 5.5 ± 2.9 | $3.10\times 10^{-1}$ | AT1G74090 | -1649 | *SOT18* | Arabidopsis sulfotransferase 5b |
| 2 | 862457 | **$6.80\times 10^{-4}$** | 105/A | 3.1 ± 0.6 | 10/G | 6.6 ± 2.4 | **$7.40\times 10^{-2}$** | AT2G02960 | Exon | | RING/FYVE/PHD zinc finger superfamily protein |
| 2 | 5016037 | **$5.30\times 10^{-4}$** | 100/T | 2.7 ± 0.3 | 15/A | 8.4 ± 3.8 | **$3.10\times 10^{-4}$** | AT2TE20600 | 699 | | ATHILA4A transposon family,LTR/Gypsy transposon super family |
| 2 | 5040311 | **$8.00\times 10^{-4}$** | 106/G | 2.7 ± 0.3 | 9/A | 11.9 ± 6.2 | **$3.10\times 10^{-6}$** | AT2TE20730 | -637 | | ATLANTYS1 transposon family,LTR/Gypsy transposon super family |
| 2 | 6037239 | **$4.60\times 10^{-4}$** | 94/G | 3.2 ± 0.7 | 21/A | 4.8 ± 1.3 | $2.60\times 10^{-1}$ | AT2G14255 | Intron | | Ankyrin repeat family protein with DHHC zinc finger domain-containing protein |
| 2 | 7629760 | **$1.30\times 10^{-4}$** | 87/G | 3.1 ± 0.7 | 28/C | 4.7 ± 1 | $2.00\times 10^{-1}$ | AT2G17540 | 902 | | Hypothetical protein |
| 2 | 7699748 | **$7.10\times 10^{-4}$** | 80/T | 2.4 ± 0.3 | 35/G | 5.8 ± 1.7 | **$4.30\times 10^{-3}$** | AT2G17723 | -386 | | Defensin-like (DEFL) family protein |
| 2 | 8285903 | **$1.80\times 10^{-4}$** | 102/A | 2.6 ± 0.3 | 13/T | 10.3 ± 4.3 | **$3.30\times 10^{-6}$** | AT2G19110 | Exon | *HMA4* | Arabidopsis heavy metal ATPase 4 |
| | *Chr2:8285903 - Chr2:8290712* | | | | | | | AT2G19120 | 1133 | | P-loop containing nucleoside triphosphate hydrolases superfamily protein |
| 2 | 8291211 | **$8.10\times 10^{-5}$** | 100/A | 2.5 ± 0.2 | 15/T | 9.5 ± 3.8 | **$9.10\times 10^{-6}$** | AT2G19120 | Exon | | P-loop containing nucleoside triphosphate hydrolases superfamily protein |
| 2 | 8291276 | **$2.00\times 10^{-5}$** | 98/C | 2.4 ± 0.2 | 17/T | 9.5 ± 3.3 | **$1.40\times 10^{-6}$** | AT2G19120 | Exon | | P-loop containing nucleoside triphosphate hydrolases superfamily protein |
| 2 | 8757608 | **$4.50\times 10^{-4}$** | 93/G | 2.5 ± 0.2 | 22/A | 7.4 ± 2.7 | **$2.80\times 10^{-4}$** | AT2G20300 | Exon | *ALE2* | Abnormal leaf shape 2 |
| 2 | 9050750 | **$5.20\times 10^{-4}$** | 103/T | 2.6 ± 0.3 | 12/A | 10.4 ± 4.6 | $7.80\times 10^{-6}$ | AT2G21110 | Exon | *DIR4* | Dirigent protein 4 |
| | | | | | | | | AT2G21080 | 5637 | | Ras guanine nucleotide exchange factor K |
| | | | | | | | | AT2G21090 | -3262 | | Pentatricopeptide repeat (PPR-like) superfamily protein |
| | *Chr2:9044450 - Chr2:9052185* | | | | | | | AT2G21120 | 1353 | *AVI2, ENOR3* | Magnesium transporter, putative (DUF803) |
| 2 | 10619291 | **$2.00\times 10^{-4}$** | 103/C | 3.2 ± 0.6 | 12/T | 5.6 ± 2 | $1.70\times 10^{-1}$ | AT2G24960 | Exon | | Myb/SANT-like DNA-binding domain protein |
| 2 | 10619519 | **$2.00\times 10^{-4}$** | 103/A | 3.2 ± 0.6 | 12/T | 5.6 ± 2 | $1.70\times 10^{-1}$ | AT2G24960 | Intron | | Myb/SANT-like DNA-binding domain protein |
| 2 | 10620788 | **$3.30\times 10^{-4}$** | 97/A | 3.1 ± 0.6 | 18/T | 5.2 ± 1.5 | $1.70\times 10^{-1}$ | AT2G24970 | Intron | | Spindle/kinetochore-associated protein |
| | *Chr2:10620788 - Chr2:10631922* | | | | | | | AT2G25010 | -10903 | *MAIL1* | Aminotransferase-like, plant mobile domain family protein |
| 2 | 10860253 | **$6.20\times 10^{-4}$** | 105/C | 2.7 ± 0.3 | 10/A | 11 ± 5.6 | **$1.10\times 10^{-5}$** | AT2G25520 | -697 | | Drug/metabolite transporter superfamily protein |
| 2 | 12471135 | **$5.20\times 10^{-4}$** | 103/C | 3.2 ± 0.6 | 12/T | 5.4 ± 2 | $2.40\times 10^{-1}$ | AT2G29020 | Intron | | Rab5-interacting family protein |
| 2 | 12485512 | **$4.00\times 10^{-4}$** | 104/G | 3.2 ± 0.6 | 11/A | 5.7 ± 2.2 | $1.90\times 10^{-1}$ | AT2G29060 | Exon | | GRAS family transcription factor |
| 2 | 12488101 | **$5.20\times 10^{-4}$** | 102/T | 3.3 ± 0.6 | 13/A | 4.9 ± 2 | $3.40\times 10^{-1}$ | AT2G29070 | Intron | | Ubiquitin fusion degradation UFD1 family protein |
| | *Chr2:12488101 - Chr2:12490396* | | | | | | | AT2G29080 | 1810 | *FTSH3* | FTSH protease 3 |
| 2 | 14603435 | **$1.70\times 10^{-4}$** | 102/A | 2.7 ± 0.3 | 13/G | 9 ± 4.4 | $2.30\times 10^{-4}$ | AT2G34660 | Exon | *ABCC2* | Multidrug resistance-associated protein 2 |
| 3 | 1114598 | **$7.40\times 10^{-4}$** | 104/T | 3.1 ± 0.6 | 11/G | 6.3 ± 2.2 | $9.10\times 10^{-2}$ | AT3G04240 | Exon | *SEC* | Tetratricopeptide repeat (TPR)-like superfamily protein |
| 3 | 1143847 | **$2.80\times 10^{-4}$** | 105/C | 3.1 ± 0.6 | 10/T | 6.7 ± 2.4 | $7.00\times 10^{-2}$ | AT3G04320 | Exon | | Kunitz family trypsin and protease inhibitor protein |
| | | | | | | | | AT3G04290 | -7624 | *LTL1* | Li-tolerant lipase 1 |

| | | | | | | | | | | | |
|---|---|---|---|---|---|---|---|---|---|---|---|
| | | | | | | | | AT3G04300 | 3124 | | RmlC-like cupins superfamily protein |
| | | | | | | | | AT3G04310 | -1319 | | Transmembrane protein |
| | | | | | | | | AT3G04340 | 3096 | *TSL2* | FtsH extracellular protease family |
| *Chr3:1135444 - Chr3:1154899* | | | | | | | | AT3G04350 | 10125 | | Vacuolar sorting-associated protein (DUF946) |
| 3 | 1153108 | **$1.60\times 10^{-4}$** | 107/A | 3.2 ± 0.6 | 8/G | 7.5 ± 2.9 | **$4.50\times 10^{-2}$** | AT3G04340 | Exon | *TSL2* | FtsH extracellular protease family |
| 3 | 1155166 | **$1.60\times 10^{-4}$** | 107/C | 3.2 ± 0.6 | 8/G | 7.5 ± 2.9 | **$4.50\times 10^{-2}$** | AT3G04350 | Exon | | Vacuolar sorting-associated protein (DUF946) |
| | | | | | | | | AT3G04340 | -1825 | *TSL2* | FtsH extracellular protease family |
| *Chr3:1151538 - Chr3:1157673* | | | | | | | | AT3G04360 | 1859 | | Calcium-dependent lipid-binding (CaLB domain) family protein |
| 3 | 1157673 | **$2.80\times 10^{-4}$** | 106/G | 3.1 ± 0.6 | 9/A | 7.1 ± 2.6 | $5.30\times 10^{-2}$ | AT3G04360 | Exon | | Calcium-dependent lipid-binding (CaLB domain) family protein |
| | | | | | | | | AT3G04340 | -4332 | *TSL2* | FtsH extracellular protease family |
| *Chr3:1151538 - Chr3:1157673* | | | | | | | | AT3G04350 | -1204 | | Vacuolar sorting-associated protein (DUF946) |
| 3 | 1157955 | **$8.40\times 10^{-4}$** | 106/G | 3.2 ± 0.6 | 9/T | 6.8 ± 2.6 | $7.10\times 10^{-2}$ | AT3G04360 | Exon | | Calcium-dependent lipid-binding (CaLB domain) family protein |
| *Chr3:1155283 - Chr3:1157955* | | | | | | | | AT3G04350 | -1486 | | Vacuolar sorting-associated protein (DUF946) |
| 3 | 3395510 | **$2.60\times 10^{-4}$** | 107/G | 2.7 ± 0.3 | 8/A | 12.9 ± 6.8 | **$8.50\times 10^{-7}$** | AT3G10845 | Exon | | RNA-binding (RRM/RBD/RNP motifs) family protein |
| 3 | 3395649 | **$3.60\times 10^{-4}$** | 108/G | 2.8 ± 0.3 | 7/A | 13.9 ± 7.8 | **$3.10\times 10^{-7}$** | AT3G10845 | Intron | | RNA-binding (RRM/RBD/RNP motifs) family protein |
| | | | | | | | | AT3G10820 | 6623 | *MED26A* | Transcription elongation factor (TFIIS) family protein |
| *Chr3:3389313 - Chr3:3399271* | | | | | | | | AT3G10850 | 2107 | *GLX2-2* | Metallo-hydrolase/oxidoreductase superfamily protein |
| 3 | 5210413 | **$6.20\times 10^{-4}$** | 100/G | 3.1 ± 0.6 | 15/A | 5.5 ± 1.7 | $1.40\times 10^{-1}$ | AT3G15430 | Exon | | Regulator of chromosome condensation (RCC1) family protein |
| 3 | 5657132 | **$4.60\times 10^{-4}$** | 96/A | 2.6 ± 0.3 | 19/T | 7.9 ± 3.1 | **$2.30\times 10^{-4}$** | AT3G16610 | Exon | | Pentatricopeptide (PPR) repeat-containing protein |
| 3 | 5672604 | **$7.80\times 10^{-4}$** | 93/T | 3.1 ± 0.7 | 22/G | 5 ± 1.3 | $1.80\times 10^{-1}$ | AT3G16650 | Intron | *PRL2* | Transducin/WD40 repeat-like superfamily protein |
| *Chr3:5670598 - Chr3:5674751* | | | | | | | | AT3G16640 | -1875 | *TCTP1* | Translationally controlled tumor protein |
| 3 | 6469820 | **$9.30\times 10^{-4}$** | 99/T | 3.2 ± 0.6 | 16/G | 5 ± 1.6 | $2.60\times 10^{-1}$ | AT3G18773 | 2854 | *ATL77* | RING/U-box superfamily protein |
| 3 | 8422878 | **$7.70\times 10^{-5}$** | 106/C | 3.2 ± 0.6 | 9/G | 6.7 ± 2.7 | $7.90\times 10^{-2}$ | AT3G23480 | Exon | | Cyclopropane-fatty-acyl-phospholipid synthase |
| 3 | 19447028 | **$7.70\times 10^{-4}$** | 101/C | 2.7 ± 0.3 | 14/A | 9.2 ± 4 | **$5.50\times 10^{-5}$** | AT3G52460 | Exon | | Hydroxyproline-rich glycoprotein family protein |
| 3 | 19481922 | **$1.70\times 10^{-4}$** | 103/T | 3.2 ± 0.6 | 12/A | 5.2 ± 2.1 | $2.70\times 10^{-1}$ | AT3G52530 | 2945 | *ZRK11* | Protein kinase superfamily protein |
| 3 | 23191968 | **$3.60\times 10^{-4}$** | 110/G | 3.3 ± 0.6 | 5/A | 6.7 ± 4.5 | $2.10\times 10^{-1}$ | AT3G62700 | Exon | *MRP10* | Multidrug resistance-associated protein 10 |
| 3 | 23193409 | **$2.00\times 10^{-4}$** | 109/T | 3.3 ± 0.6 | 6/C | 6.9 ± 3.7 | $1.30\times 10^{-1}$ | AT3G62700 | Exon | *MRP10* | Multidrug resistance-associated protein 10 |
| 3 | 23194429 | **$5.10\times 10^{-4}$** | 109/A | 3.3 ± 0.6 | 6/G | 6.5 ± 3.7 | $1.90\times 10^{-1}$ | AT3G62700 | Exon | *MRP10* | Multidrug resistance-associated protein 10 |
| 4 | 140430 | **$5.80\times 10^{-6}$** | 111/G | 3.2 ± 0.6 | 4/A | 9.2 ± 5.6 | **$4.70\times 10^{-2}$** | AT4G00320 | Exon | | F-box/RNI-like superfamily protein |
| 4 | 2662015 | **$6.10\times 10^{-4}$** | 102/C | 3.2 ± 0.6 | 13/A | 5.4 ± 1.9 | $2.10\times 10^{-1}$ | AT4G05150 | Exon | | Octicosapeptide/Phox/Bem1p family protein |
| 4 | 7070654 | **$4.50\times 10^{-4}$** | 96/A | 3.2 ± 0.7 | 19/G | 4.9 ± 1.3 | $2.30\times 10^{-1}$ | AT4G11730 | Exon | | Cation transporter/ E1-E2 ATPase family protein |
| *Chr4:7069534 - Chr4:7073549* | | | | | | | | AT4G11740 | -1301 | *SAY1* | Ubiquitin-like superfamily protein |
| 4 | 7199963 | **$3.60\times 10^{-4}$** | 92/C | 2.4 ± 0.3 | 23/T | 7.5 ± 2.5 | **$1.60\times 10^{-4}$** | AT4G12010 | Exon | *DSC1* | Disease resistance protein (TIR-NBS-LRR class) family |
| 4 | 7948195 | **$8.70\times 10^{-4}$** | 89/G | 2.5 ± 0.3 | 26/A | 6.6 ± 2.3 | **$1.70\times 10^{-3}$** | AT4G13660 | Exon | *PRR2* | Pinoresinol reductase 2 |
| 4 | 9044548 | **$5.90\times 10^{-4}$** | 99/A | 3.2 ± 0.6 | 16/G | 5.3 ± 1.6 | $1.90\times 10^{-1}$ | AT4G15955 | Intron | | Alpha/beta-Hydrolases superfamily protein |

| | | | | | | | | | | | |
|---|---|---|---|---|---|---|---|---|---|---|---|
| 4 | 9297862 | **$2.20\times 10^{-5}$** | 86/C | 2.5 ± 0.3 | 29/T | 6.3 ± 2.1 | **$2.20\times 10^{-3}$** | AT4G16490 | -2332 | | ARM repeat superfamily protein |
| 4 | 10893131 | **$6.50\times 10^{-4}$** | 106/T | 3.2 ± 0.6 | 9/G | 6.7 ± 2.6 | $8.80\times 10^{-2}$ | AT4G20160 | Exon | | Golgin family A protein |
| 4 | 11279865 | **$7.10\times 10^{-4}$** | 102/T | 3.1 ± 0.6 | 13/G | 6.1 ± 2 | $8.50\times 10^{-2}$ | AT4G21150 | Exon | *HAP6* | Ribophorin II (RPN2) family protein |
| 4 | 12374490 | **$7.40\times 10^{-4}$** | 100/A | 3.1 ± 0.6 | 15/C | 5.5 ± 1.7 | $1.40\times 10^{-1}$ | AT4G23750 | -2261 | *CRF2* | Cytokinin response factor 2 |
| 4 | 13186139 | **$4.50\times 10^{-5}$** | 104/T | 2.7 ± 0.3 | 11/A | 10.9 ± 5.1 | **$3.70\times 10^{-6}$** | AT4G25970 | Intron | *PSD3* | Phosphatidylserine decarboxylase 3 |
| 4 | 13187817 | **$4.50\times 10^{-5}$** | 104/G | 2.7 ± 0.3 | 11/A | 10.9 ± 5.1 | **$3.70\times 10^{-6}$** | AT4G25970 | Exon | *PSD3* | Phosphatidylserine decarboxylase 3 |
| 4 | 13226704 | **$2.70\times 10^{-4}$** | 109/T | 2.8 ± 0.3 | 6/A | 14.8 ± 8.6 | **$2.90\times 10^{-7}$** | AT4G26090 | Exon | *RPS2* | Resistant to P. Syringae 2 |
| 4 | 13229245 | **$2.70\times 10^{-4}$** | 109/G | 2.8 ± 0.3 | 6/A | 14.8 ± 8.6 | **$2.90\times 10^{-7}$** | AT4G26100 | Intron | *CKL1* | Casein kinase 1 |
| 4 | 14014214<br>*Chr4:14006703 - Chr4:14014214* | **$3.60\times 10^{-4}$** | 94/T | 3.1 ± 0.7 | 21/C | 5.2 ± 1.3 | $1.30\times 10^{-1}$ | AT4G28300<br>AT4G28260 | -646<br>7452 | *FLOE1* | Formin-like protein (DUF1421)<br>Acyl-UDP-N-acetylglucosamine O-acyltransferase |
| 4 | 14512202 | **$3.70\times 10^{-4}$** | 101/A | 3.1 ± 0.6 | 14/G | 5.7 ± 1.8 | $1.20\times 10^{-1}$ | AT4G29580 | 16 | | Cytidine/deoxycytidylate deaminase family protein |
| 4 | 14513116 | **$8.70\times 10^{-4}$** | 101/T | 3.1 ± 0.6 | 14/A | 5.6 ± 1.8 | $1.40\times 10^{-1}$ | AT4G29590 | Intron | | Putative methyltransferase |
| 4 | 17850934 | **$9.30\times 10^{-4}$** | 96/G | 2.6 ± 0.3 | 19/A | 7.5 ± 3 | **$7.30\times 10^{-4}$** | AT4G37970 | Exon | *CAD6* | Cinnamyl alcohol dehydrogenase 6 |
| 4 | 17877877<br>*Chr4:17875134 - Chr4:17877877* | **$8.20\times 10^{-4}$** | 97/T | 2.7 ± 0.3 | 18/C | 7.6 ± 3.2 | **$9.00\times 10^{-4}$** | AT4G38070<br>AT4G38062 | Exon<br>1810 | | Transcription factor bHLH131-like protein<br>Basic helix loop helix (bHLH) DNA-binding superfamily protein |
| 5 | 1083021<br>*Chr5:1080404 - Chr5:1083021* | **$2.80\times 10^{-4}$** | 103/G | 3.2 ± 0.6 | 12/A | 5.4 ± 2.1 | $2.30\times 10^{-1}$ | AT5G04020<br>AT5G04010 | Exon<br>-1953 | | Calmodulin binding protein<br>F-box family protein |
| 5 | 1084636 | **$9.20\times 10^{-4}$** | 103/A | 3.3 ± 0.6 | 12/C | 5.1 ± 2.1 | $3.10\times 10^{-1}$ | AT5G04020 | Exon | | Calmodulin binding protein |
| 5 | 1105815<br>*Chr5:1093060 - Chr5:1105815* | **$3.30\times 10^{-4}$** | 102/T | 3.3 ± 0.6 | 13/G | 5 ± 2 | $3.30\times 10^{-1}$ | AT5G04090<br>AT5G04060<br>AT5G04040 | Intron<br>4005<br>12812 | <br><br>*SDP1* | Histidine-tRNA ligase<br>S-adenosyl-L-methionine-dependent methyltransferases superfamily protein<br>Patatin-like phospholipase family protein |
| 5 | 3796248<br>*Chr5:3796248 - Chr5:3806411* | **$7.40\times 10^{-5}$** | 88/T | 2.5 ± 0.3 | 27/A | 6.5 ± 2.2 | **$1.90\times 10^{-3}$** | AT5G11780<br>AT5G11800 | 265<br>7387 | <br>*KEA6* | SNF2 domain protein<br>K+ efflux antiporter 6 |
| 5 | 5290037<br>*Chr5:5288006 - Chr5:5292164* | **$6.30\times 10^{-4}$** | 100/G | 3.1 ± 0.6 | 15/A | 5.6 ± 1.7 | $1.30\times 10^{-1}$ | AT5G16200<br>AT5G16210 | Exon<br>962 | | 50S ribosomal protein-like protein<br>HEAT repeat-containing protein |
| 5 | 5466004<br>*Chr5:5461864 - Chr5:5466004* | **$3.00\times 10^{-4}$** | 100/C | 3.1 ± 0.6 | 15/T | 6.1 ± 1.6 | $6.30\times 10^{-2}$ | AT5G16660<br>AT5G16640 | Intron<br>3459 | | Low-density receptor-like protein<br>Pentatricopeptide repeat (PPR) superfamily protein |
| 5 | 7128699 | **$9.80\times 10^{-4}$** | 92/G | 2.5 ± 0.3 | 23/C | 7.3 ± 2.5 | $3.30\times 10^{-4}$ | AT5G20990 | 38 | *B73* | Molybdopterin biosynthesis CNX1 protein |
| 5 | 7241108<br>*Chr5:7241108 - Chr5:7242531* | **$6.20\times 10^{-4}$** | 99/T | 3.1 ± 0.6 | 16/C | 5.9 ± 1.6 | $7.00\times 10^{-2}$ | AT5G21910<br>AT5G21920 | Exon<br>242 | <br>*YLMG2* | Transmembrane protein<br>YGGT family protein |
| 5 | 13768447 | **$1.10\times 10^{-4}$** | 104/G | 3.2 ± 0.6 | 11/T | 5.9 ± 2.2 | $1.60\times 10^{-1}$ | AT5G35590 | -679 | *PAA1* | Proteasome alpha subunit A1 |
| 5 | 13958580 | **$7.40\times 10^{-4}$** | 84/A | 3.3 ± 0.7 | 31/G | 3.8 ± 0.9 | $7.20\times 10^{-1}$ | AT5G35790 | Intron | *G6PD1* | Glucose-6-phosphate dehydrogenase 1 |
| 5 | 14012117 | **$6.20\times 10^{-4}$** | 95/G | 3.2 ± 0.7 | 20/A | 4.7 ± 1.3 | $3.00\times 10^{-1}$ | AT5TE49950 | Exon | | ATHILA7 transposon family,LTR/Gypsy transposon super family |
| 5 | 16162612 | **$1.90\times 10^{-4}$** | 104/C | 2.7 ± 0.3 | 11/A | 10.3 ± 5.1 | **$3.00\times 10^{-5}$** | AT5G40390 | Intron | *RS5, SIP1* | Raffinose synthase family protein |
| 5 | 18375331 | **$6.00\times 10^{-4}$** | 73/G | 2.3 ± 0.3 | 42/T | 5.4 ± 1.5 | $5.50\times 10^{-3}$ | AT5TE66150 | -2701 | | HELITRONY1D transposon family,RC/Helitron transposon super family |

| | | | | | | | | | | | |
|---|---|---|---|---|---|---|---|---|---|---|---|
| 5 | 21148701 | **$1.50\times 10^{-4}$** | 99/T | 3.1 ± 0.6 | 16/C | 5.5 ± 1.6 | $1.40\times 10^{-1}$ | AT5G52060 | 3748 | *BAG1* | BCL-2-associated athanogene 1 |
| 5 | 21380524<br>*Chr5:21380524 - Chr5:21391426* | **$6.70\times 10^{-4}$** | 104/G | 3.3 ± 0.6 | 11/A | 5.3 ± 2.3 | $2.60\times 10^{-1}$ | AT5G52730 | Exon | | Copper transport protein family |
| | | | | | | | | AT5G52780 | 10419 | *PAM68L* | Transmembrane protein, putative (DUF3464) |
| 5 | 22314362 | **$3.50\times 10^{-4}$** | 57/A | 2 ± 0.2 | 58/T | 4.8 ± 1.1 | **$1.00\times 10^{-2}$** | AT5G54980 | -1437 | *CASPL2D1* | Uncharacterized protein family (UPF0497) |
| 5 | 22314362 | **$3.50\times 10^{-4}$** | 57/A | 2 ± 0.2 | 58/T | 4.8 ± 1.1 | **$1.00\times 10^{-2}$** | AT5G54970 | 714 | | Hypothetical protein |
| 5 | 22937262<br>*Chr5:22934538 - Chr5:22941977* | **$3.20\times 10^{-4}$** | 105/T | 3.3 ± 0.6 | 10/C | 5.1 ± 2.5 | $3.60\times 10^{-1}$ | AT5G56680 | Exon | *EMB2755* | Embryo defective 2755 |
| | | | | | | | | AT5G56660 | 2251 | *ILL2* | IAA-leucine resistant (ILR)-like 2 |
| | | | | | | | | AT5G56700 | -4729 | | FBD / Leucine Rich Repeat domains containing protein |
| 5 | 22937474 | **$1.40\times 10^{-4}$** | 105/T | 3.2 ± 0.6 | 10/C | 5.7 ± 2.5 | $2.10\times 10^{-1}$ | AT5G56680 | Exon | *EMB2755* | Embryo defective 2755 |
| 5 | 22937694<br>*Chr5:22934538 - Chr5:22938254* | **$3.20\times 10^{-4}$** | 105/A | 3.3 ± 0.6 | 10/G | 5.1 ± 2.5 | $3.60\times 10^{-1}$ | AT5G56680 | Exon | *EMB2755* | Embryo defective 2755 |
| | | | | | | | | AT5G56660 | 2683 | *ILL2* | IAA-leucine resistant (ILR)-like 2 |
| 5 | 22938254<br>*Chr5:22934538 - Chr5:22938254* | **$3.20\times 10^{-4}$** | 105/A | 3.3 ± 0.6 | 10/G | 5.1 ± 2.5 | $3.60\times 10^{-1}$ | AT5G56680 | Intron | *EMB2755* | Embryo defective 2755 |
| | | | | | | | | AT5G56660 | 3243 | *ILL2* | IAA-leucine resistant (ILR)-like 2 |
| 5 | 22939011 | **$2.50\times 10^{-4}$** | 104/T | 3.3 ± 0.6 | 11/A | 5 ± 2.3 | $3.60\times 10^{-1}$ | AT5G56680 | 170 | *EMB2755* | Embryo defective 2755 |
| 5 | 26542207 | **$1.40\times 10^{-4}$** | 112/G | 3.3 ± 0.6 | 3/A | 9.2 ± 7.7 | $8.30\times 10^{-2}$ | AT5G66470 | Exon | *ERA1* | GTP-binding protein Era-like protein |

Genes within haploblocks of the most significantly associated single-nucleotide polymorphisms (SNPs) in an expression genome-wide association study (eGWAS) of LKR superinduction (D2/D1) under recurrent droughts (D1 and D2 separated by a nonstressful recovery interval R) in 115 *Arabidopsis thaliana* ecotypes. Top-ranking SNPs having a *P*-value $< 10^{-3}$ are represented in the table. The haploblocks are indicated by bold rectangles. The locations of the SNPs with respect to the nearest genes are shown. Minus and plus signs denote upstream and downstream locations of the SNP with respect to the gene. The relative fold change (RFC) of expression levels of the *A. thaliana* ecotypes having SNP alleles associated with high and low D2/D1 induction are shown as average values with standard error, along with the allele frequencies and a *t*-test *P*-value between these groups.

#### Table S4 Genes associated with the most significant SNPs in an expression genome-wide association study of *HIS1-3* superinduction in *Arabidopsis thaliana* under recurrent droughts

| Chromosome | Physical position (bp) | GWAS *P*-value | Frequency of low induction allele | RFC *HIS1-3* (D2/D1) of low induction allele | Frequency of high induction allele | RFC *HIS1-3* (D2/D1) of high induction allele | *t*-test *P*-value (segregation of alleles) | Nearest Gene | SNP position with respect to the gene | Gene Symbol | Short Description |
|---|---|---|---|---|---|---|---|---|---|---|---|
| 1 | 133144 | **$3.28\times 10^{-5}$** | 108/G | 2.6 ± 0.3 | 6/T | 9.7 ± 3.7 | **$9.70\times 10^{-7}$** | AT1G01340 | Exon | *ACBK1, CNGC10* | Cyclic nucleotide gated channel 10 |
| 1 | 3682074 | **$4.18\times 10^{-4}$** | 108/T | 2.7 ± 0.3 | 6/C | 8.5 ± 3.6 | **$8.69\times 10^{-5}$** | AT1G11050 | Exon | | Protein kinase superfamily protein |
| 1 | 4516808 | **$4.31\times 10^{-4}$** | 95/A | 2.5 ± 0.3 | 19/G | 5.3 ± 1.4 | **$1.80\times 10^{-3}$** | AT1G13220 | Exon | *CRWN2* | Crowded nuclei 2 |
| 1 | 4859326 | **$3.38\times 10^{-6}$** | 105/A | 2.5 ± 0.3 | 9/C | 9 ± 2.6 | **$3.71\times 10^{-8}$** | AT1G14220 | Exon | | Ribonuclease T2 family protein |
| | | | | | | | | AT1G14230 | -2171 | | GDA1/CD39 nucleoside phosphatase family protein |
| *Chr1:4859052 - Chr1:4865227* | | | | | | | | AT1G14240 | -5833 | *APY3* | GDA1/CD39 nucleoside phosphatase family protein |
| 1 | 5052910 | **$1.33\times 10^{-4}$** | 104/A | 2.6 ± 0.3 | 10/T | 7.1 ± 2.5 | **$9.57\times 10^{-5}$** | AT1G14690 | Intron | *MAP65-7* | Microtubule-associated protein 65-7 |
| 1 | 5237652 | **$4.66\times 10^{-4}$** | 63/A | 2 ± 0.3 | 51/G | 4.2 ± 0.7 | **$1.10\times 10^{-3}$** | AT1G15215 | -444 | *DTF1* | DNA-binding transcription factor 1 |
| 1 | 5755331 | **$7.95\times 10^{-4}$** | 106/G | 2.7 ± 0.3 | 8/A | 6.9 ± 3.1 | **$1.13\times 10^{-3}$** | AT1G16800 | -68 | | P-loop containing nucleoside triphosphate hydrolases superfamily protein |
| 1 | 5763685 | **$7.27\times 10^{-4}$** | 16/G | 1.3 ± 0.3 | 98/A | 3.3 ± 0.4 | **$4.20\times 10^{-2}$** | AT1G16840 | Exon | | Hypothetical protein |
| 1 | 8436256 | **$1.55\times 10^{-4}$** | 85/C | 2.4 ± 0.3 | 29/A | 4.7 ± 1 | **$2.94\times 10^{-3}$** | AT1G23880 | Exon | | NHL domain-containing protein |
| *Chr1:8436256 - Chr1:8439351* | | | | | | | | AT1G23890 | 2644 | | NHL domain-containing protein |
| 1 | 9984544 | **$8.26\times 10^{-4}$** | 79/A | 2.5 ± 0.4 | 35/C | 4 ± 0.8 | $5.30\times 10^{-2}$ | AT1G28420 | Exon | *HB1* | Homeobox-1 |
| 1 | 21410324 | **$7.94\times 10^{-4}$** | 57/G | 2.2 ± 0.4 | 57/C | 3.8 ± 0.6 | **$2.05\times 10^{-2}$** | AT1G57800 | Intron | *VIM5* | Variant in methylation 5 |
| 1 | 26014202 | **$8.83\times 10^{-4}$** | 94/G | 2.8 ± 0.4 | 20/T | 3.9 ± 1.2 | $2.04\times 10^{-1}$ | AT1G69190 | Exon | *DHPS;HPPK* | Dihydropterin pyrophosphokinase / Dihydropteroate synthase |
| 2 | 7707454 | **$8.71\times 10^{-5}$** | 100/G | 2.5 ± 0.3 | 14/A | 6.6 ± 1.6 | **$5.07\times 10^{-5}$** | AT2G17740 | Exon | *VLG* | Vacuoleless gametophytes |
| 2 | 8936574 | **$7.03\times 10^{-4}$** | 106/C | 2.7 ± 0.3 | 8/A | 6.9 ± 3.1 | **$1.22\times 10^{-3}$** | AT2G20740 | Intron | | Tetraspanin family protein |
| 2 | 10262887 | **$6.40\times 10^{-4}$** | 73/T | 2.1 ± 0.3 | 41/C | 4.5 ± 0.8 | **$8.56\times 10^{-4}$** | AT2G24140 | Exon | *MRF7* | Myosin-J heavy chain-like protein |
| 2 | 10263271 | **$6.72\times 10^{-4}$** | 71/A | 2.1 ± 0.3 | 43/T | 4.5 ± 0.8 | **$3.74\times 10^{-4}$** | AT2G24140 | Exon | *MRF7* | Myosin-J heavy chain-like protein |
| 2 | 10263420 | **$6.69\times 10^{-4}$** | 69/C | 2 ± 0.3 | 45/A | 4.4 ± 0.7 | **$4.73\times 10^{-4}$** | AT2G24140 | Exon | *MRF7* | Myosin-J heavy chain-like protein |
| 2 | 10263564 | **$8.00\times 10^{-4}$** | 53/T | 1.9 ± 0.4 | 61/A | 3.9 ± 0.6 | **$2.57\times 10^{-3}$** | AT2G24140 | Exon | *MRF7* | Myosin-J heavy chain-like protein |
| 2 | 10619291 | **$9.28\times 10^{-5}$** | 102/C | 2.5 ± 0.3 | 12/T | 7 ± 2.1 | **$3.43\times 10^{-5}$** | AT2G24960 | Exon | | Myb/SANT-like DNA-binding domain protein |
| 2 | 10619519 | **$9.28\times 10^{-5}$** | 102/A | 2.5 ± 0.3 | 12/T | 7 ± 2.1 | **$3.43\times 10^{-5}$** | AT2G24960 | Intron | | Myb/SANT-like DNA-binding domain protein |
| 2 | 10978006 | **$5.11\times 10^{-4}$** | 101/T | 2.8 ± 0.3 | 13/C | 4.5 ± 2.1 | $1.04\times 10^{-1}$ | AT2G25737 | Intron | | Sulfite exporter TauE/SafE family protein |
| 2 | 11818914 | **$5.02\times 10^{-5}$** | 104/A | 2.5 ± 0.3 | 10/C | 7.8 ± 2.5 | **$5.47\times 10^{-6}$** | AT2G27720 | Intron | *P2Z, RPP2A* | Ribosomal protein |
| 2 | 15915329 | **$2.10\times 10^{-4}$** | 15/C | 1.6 ± 0.7 | 99/T | 3.2 ± 0.4 | $1.24\times 10^{-1}$ | AT2G38025 | Exon | *OTU3* | Cysteine proteinases superfamily protein |
| | | | | | | | | AT2G38010 | 5462 | *ATNCER2* | Neutral ceramidase 2 |
| | | | | | | | | AT2G38020 | -1139 | *VCL1* | MVB-to-vacuole trafficking protein |
| | | | | | | | | AT2G38040 | -2283 | *CAC3* | Acetyl co-enzyme a carboxylase carboxyltransferase alpha subunit |

| | | | | | | | | | | | |
|---|---|---|---|---|---|---|---|---|---|---|---|
| *Chr2:15907320 - Chr2:15924114* | | | | | | | | AT2G38060 | 7398 | *PHT4;2* | Phosphate transporter 4;2 |
| 2 | 16148964 | **$1.46\times 10^{-4}$** | 109/A | 2.6 ± 0.3 | 5/T | 10.3 ± 3 | **$1.61\times 10^{-6}$** | AT2G38610 | Exon | *KH9* | RNA-binding KH domain-containing protein |
| | | | | | | | | AT2G38630 | 5319 | *FBIP2* | D FBS interacting protein 2 |
| *Chr2:16148964 - Chr2:16158095* | | | | | | | | AT2G38640 | 8761 | | LURP-one-like protein (DUF567) |
| 2 | 16959062 | **$6.51\times 10^{-4}$** | 98/T | 2.4 ± 0.3 | 16/C | 6.4 ± 1.7 | **$3.10\times 10^{-5}$** | AT2G40630 | Exon | *BTE1* | Uncharacterized conserved protein (UCP030365) |
| 2 | 17870970 | **$9.96\times 10^{-4}$** | 110/G | 2.8 ± 0.4 | 4/A | 7.9 ± 3.7 | **$5.02\times 10^{-3}$** | AT2G42960 | -340 | | Protein kinase superfamily protein |
| *Chr2:17870970 - Chr2:17872673* | | | | | | | | AT2G42975 | -3028 | | Myosin-G heavy chain-like protein |
| 2 | 17908108 | **$6.41\times 10^{-4}$** | 96/T | 2.4 ± 0.3 | 18/G | 6 ± 1.5 | **$6.50\times 10^{-5}$** | AT2G43060 | -1139 | *IBH1* | ILI1 binding BHLH 1 |
| *Chr2:17908108 - Chr2:17913697* | | | | | | | | AT2G43070 | 3125 | *SPPL3* | SIGNAL PEPTIDE PEPTIDASE-LIKE 3 |
| 2 | 18210302 | **$6.61\times 10^{-4}$** | 110/C | 2.8 ± 0.3 | 4/G | 8.3 ± 5.1 | **$2.71\times 10^{-3}$** | AT2G43980 | Exon | *ITPK4* | Inositol 1,3,4-trisphosphate 5/6-kinase 4 |
| *Chr2:18210302 - Chr2:18217701* | | | | | | | | AT2G44020 | 7435 | | Mitochondrial transcription termination factor family protein |
| 2 | 18440051 | **$4.72\times 10^{-4}$** | 98/C | 2.6 ± 0.3 | 16/A | 5.2 ± 1.6 | **$6.79\times 10^{-3}$** | AT2G44735 | Intron | | Transmembrane protein |
| | | | | | | | | AT2G44710 | 3422 | *HRLP* | RNA-binding (RRM/RBD/RNP motifs) family protein |
| *Chr2:18436357 - Chr2:18458125* | | | | | | | | AT2G44770 | -17766 | *MCR* | Engulfment and cell motility (elmo) domain B |
| 3 | 94424 | **$3.01\times 10^{-4}$** | 100/G | 2.6 ± 0.4 | 14/A | 5.4 ± 1.7 | **$8.20\times 10^{-3}$** | AT3G01310 | Exon | *VIP1* | Arabidopsis homolog of yeast VIP1 1 |
| | | | | | | | | AT3G01260 | -12419 | | Galactose mutarotase-like superfamily protein |
| *Chr3:81887 - Chr3:99914* | | | | | | | | AT3G01300 | -1089 | *PBL35* | PBS 1 like 35 |
| 3 | 274655 | **$8.21\times 10^{-5}$** | 107/A | 2.6 ± 0.3 | 7/T | 8.1 ± 3.5 | **$6.87\times 10^{-5}$** | AT3G01760 | Exon | *LHT6* | Lysine-histidine-like transporter 6 |
| | | | | | | | | AT3G01770 | 927 | *BET10, GTE11* | Bromodomain and extraterminal domain protein 10 |
| *Chr3:274655 - Chr3:280492* | | | | | | | | AT3G01780 | -4516 | *TPLATE* | ARM repeat superfamily protein |
| 3 | 352736 | **$6.22\times 10^{-4}$** | 94/A | 2.6 ± 0.4 | 20/G | 4.8 ± 1.3 | **$1.42\times 10^{-2}$** | AT3G02050 | Exon | *KUP3* | Potassium transporter KUP3p (KUP3) |
| *Chr3:352736 - Chr3:356927* | | | | | | | | AT3G02060 | -1673 | | DEAD/DEAH box helicase |
| 3 | 526854 | **$8.07\times 10^{-5}$** | 105/C | 2.6 ± 0.3 | 9/T | 7.3 ± 2.8 | **$1.19\times 10^{-4}$** | AT3G02520 | Exon | *GRF7* | General regulatory factor 7 |
| | | | | | | | | AT3G02500 | 5226 | | Mental retardation GTPase activating protein |
| | | | | | | | | AT3G02510 | -2318 | | Regulator of chromosome condensation (RCC1) family protein |
| *Chr3:519454 - Chr3:530468* | | | | | | | | AT3G02530 | 1952 | *CCT6-2* | TCP-1/cpn60 chaperonin family protein |
| 3 | 650206 | **$7.65\times 10^{-4}$** | 71/C | 2.8 ± 0.5 | 43/A | 3.2 ± 0.5 | $5.43\times 10^{-1}$ | AT3G02910 | Exon | | AIG2-like (avirulence induced gene) family protein |
| 3 | 773844 | **$1.04\times 10^{-6}$** | 112/C | 2.7 ± 0.3 | 2/T | 15.8 ± 7.5 | **$9.41\times 10^{-8}$** | AT3G03300 | Exon | *DCL2* | Dicer like 2 |
| 3 | 3133260 | **$1.24\times 10^{-4}$** | 101/T | 2.5 ± 0.3 | 13/A | 6.8 ± 2 | **$2.90\times 10^{-5}$** | AT3G10130 | -102 | *SOUL4* | SOUL heme-binding family protein |
| 3 | 3395649 | **$6.65\times 10^{-6}$** | 106/G | 2.5 ± 0.3 | 8/A | 9.2 ± 2.5 | **$9.80\times 10^{-8}$** | AT3G10845 | Intron | | RNA-binding (RRM/RBD/RNP motifs) family protein |
| | | | | | | | | AT3G10820 | 6623 | *MED26A* | Transcription elongation factor (TFIIS) family protein |
| *Chr3:3389313 - Chr3:3399271* | | | | | | | | AT3G10850 | 2107 | *GLX2-2* | Metallo-hydrolase/oxidoreductase superfamily protein |
| 3 | 3464013 | **$8.12\times 10^{-4}$** | 94/T | 2.5 ± 0.3 | 20/C | 5 ± 1.5 | **$5.20\times 10^{-3}$** | AT3G11050 | Intron | *FER2* | Ferritin 2 |
| 3 | 6122556 | **$4.19\times 10^{-4}$** | 109/G | 2.7 ± 0.3 | 5/A | 9.4 ± 3.7 | **$2.98\times 10^{-5}$** | AT3G17860 | Intron | *JAI3* | Jasmonate-zim-domain protein 3 |
| *Chr3:6121042 - Chr3:6123547* | | | | | | | | AT3G17880 | -978 | *ATHIP2* | Tetraticopeptide domain-containing thioredoxin |
| 3 | 6123206 | **$6.25\times 10^{-4}$** | 108/T | 2.7 ± 0.3 | 6/A | 8.9 ± 3.1 | **$2.66\times 10^{-5}$** | AT3G17880 | -328 | *ATHIP2* | Tetraticopeptide domain-containing thioredoxin |
| *Chr3:6122288 - Chr3:6123864* | | | | | | | | AT3G17860 | 515 | *JAI3* | Jasmonate-zim-domain protein 3 |

| | | | | | | | | | | | |
|---|---|---|---|---|---|---|---|---|---|---|---|
| 3 | 6704056 | **$5.42\times 10^{-7}$** | 108/T | 2.5 ± 0.3 | 6/G | 11 ± 3.5 | **$2.92\times 10^{-9}$** | AT3G19340 | Exon | | Aminopeptidase (DUF3754) |
| 3 | 8323524 | **$1.40\times 10^{-4}$** | 104/T | 2.6 ± 0.3 | 10/G | 7.3 ± 2.3 | **$4.42\times 10^{-5}$** | AT3G23280 | Intron | *XBAT35* | XB3 ortholog 5 in *Arabidopsis thaliana* |
| 3 | 8930393 | **$8.14\times 10^{-4}$** | 91/A | 2.5 ± 0.3 | 23/T | 4.9 ± 1.3 | **$5.17\times 10^{-3}$** | AT3G24508 | Exon | | Encodes a defensin-like (DEFL) family protein |
| 3 | 9028198 | **$4.56\times 10^{-4}$** | 96/T | 2.7 ± 0.4 | 18/G | 4.3 ± 1.4 | $1.03\times 10^{-1}$ | AT3G24715 | Exon | *HCR1* | Kinase superfamily with octicosapeptide/Phox/Bem1p domain-containing protein |
| 3 | 9146019 | **$6.45\times 10^{-4}$** | 107/T | 2.8 ± 0.4 | 7/A | 5.2 ± 3.2 | $9.15\times 10^{-2}$ | AT3G25100 | Exon | *CDC45* | Cell division cycle 45 |
| *Chr3:9145272 - Chr3:9147480* | | | | | | | | AT3G25110 | 570 | *FATA1* | Fata acyl-acp thioesterase |
| 3 | 9146043 | **$5.78\times 10^{-4}$** | 104/A | 2.8 ± 0.3 | 10/G | 5.2 ± 2.4 | **$4.54\times 10^{-2}$** | AT3G25100 | Exon | *CDC45* | Cell division cycle 45 |
| *Chr3:9146043 - Chr3:9147766* | | | | | | | | AT3G25110 | 546 | *FATA1* | Fata acyl-acp thioesterase |
| 3 | 9703450 | **$1.10\times 10^{-4}$** | 86/C | 2.1 ± 0.3 | 28/T | 5.5 ± 1.1 | **$8.74\times 10^{-6}$** | AT3G26490 | -692 | *NPY7* | Phototropic-responsive NPH3 family protein |
| 3 | 9703450 | **$1.10\times 10^{-4}$** | 86/C | 2.1 ± 0.3 | 28/T | 5.5 ± 1.1 | **$8.74\times 10^{-6}$** | AT3TE40445 | Exon | | ATLINEIII transposon family,LINE/L1 transposon super family |
| 3 | 10653911 | **$6.49\times 10^{-4}$** | 101/C | 2.7 ± 0.4 | 13/T | 5.5 ± 1.8 | **$7.68\times 10^{-3}$** | AT3G28415 | -2371 | *ABCB22* | ABC transporter family protein |
| 3 | 10677398 | **$3.81\times 10^{-4}$** | 98/C | 2.5 ± 0.3 | 16/G | 5.9 ± 1.6 | **$3.84\times 10^{-4}$** | AT3G28480 | Intron | | Oxoglutarate/iron-dependent oxygenase |
| 3 | 20799517 | **$7.04\times 10^{-4}$** | 51/T | 2.1 ± 0.4 | 63/A | 3.7 ± 0.6 | **$2.54\times 10^{-2}$** | AT3G56050 | Intron | | Protein kinase family protein |
| 3 | 22843017 | **$9.37\times 10^{-6}$** | 107/C | 2.5 ± 0.3 | 7/G | 9.6 ± 3 | **$1.35\times 10^{-7}$** | AT3G61720 | Exon | *MCTP12* | Multiple c2 domain and transmembrane region protein 12 |
| | | | | | | | | AT3G61700 | 4214 | | Helicase with zinc finger protein |
| | | | | | | | | AT3G61710 | -764 | *ATG6* | Autophagy protein 6 |
| | | | | | | | | AT3G61723 | 3451 | | PHD finger protein |
| | | | | | | | | AT3G61730 | 5410 | *RMF1* | Reduced male fertility |
| *Chr3:22837659 - Chr3:22853608* | | | | | | | | AT3G61740 | 8116 | *ATX3* | Arabidopsis trithorax 3 |
| 4 | 1516897 | **$8.51\times 10^{-5}$** | 111/G | 2.7 ± 0.3 | 3/T | 12.2 ± 5.7 | **$3.64\times 10^{-6}$** | AT4G03430 | 514 | *EMB2770* | Embryo defective 2770 |
| *Chr4:1513501 - Chr4:1516897* | | | | | | | | AT4G03420 | 3303 | | Hypothetical protein (DUF789) |
| 4 | 2413073 | **$9.47\times 10^{-4}$** | 104/T | 2.6 ± 0.3 | 10/G | 7.2 ± 2.4 | **$8.29\times 10^{-5}$** | AT4G04745 | Exon | | Hypothetical protein |
| *Chr4:2406215 - Chr4:2416340* | | | | | | | | AT4G04740 | -4580 | *CPK23* | Calcium-dependent protein kinase 23 |
| 4 | 2740856 | **$8.92\times 10^{-5}$** | 110/T | 2.7 ± 0.3 | 4/A | 10.3 ± 4.6 | **$2.09\times 10^{-5}$** | AT4G05390 | -373 | *RFNR1* | Root FNR1 |
| *Chr4:2740856 - Chr4:2748870* | | | | | | | | AT4G05420 | -5432 | *DDB1A* | Damaged dna binding protein 1a |
| 4 | 5724843 | **$8.48\times 10^{-4}$** | 96/T | 2.5 ± 0.3 | 18/C | 5.6 ± 1.6 | **$7.10\times 10^{-4}$** | AT4G08920 | Exon | *CRY1* | Cryptochrome 1 |
| 4 | 6088747 | **$6.35\times 10^{-4}$** | 110/T | 2.8 ± 0.4 | 4/G | 9 ± 3.9 | **$5.63\times 10^{-4}$** | AT4G09640 | Exon | *ENOR3L6* | Magnesium transporter, putative (DUF803) |
| *Chr4:6084522 - Chr4:6088803* | | | | | | | | AT4G09630 | 945 | | Transmembrane protein (DUF616) |
| 4 | 6745253 | **$2.74\times 10^{-4}$** | 19/T | 1.4 ± 0.4 | 95/C | 3.3 ± 0.4 | **$4.11\times 10^{-2}$** | AT4G11040 | Exon | *DOG18* | Delay of germination 18 |
| *Chr4:6741269 - Chr4:6745253* | | | | | | | | AT4G11030 | 3024 | *LACS5* | AMP-dependent synthetase and ligase family protein |
| 4 | 7247349 | **$9.80\times 10^{-4}$** | 103/A | 2.6 ± 0.4 | 11/T | 6.1 ± 1.7 | **$2.67\times 10^{-3}$** | AT4G12100 | Intron | | Cullin family protein |
| 4 | 7499646 | **$3.50\times 10^{-4}$** | 97/T | 2.6 ± 0.4 | 17/C | 4.9 ± 1.3 | **$1.66\times 10^{-2}$** | AT4G12750 | Exon | | Homeodomain-like transcriptional regulator |
| 4 | 8912575 | **$6.37\times 10^{-4}$** | 103/G | 2.7 ± 0.3 | 11/T | 5.8 ± 2.1 | **$6.53\times 10^{-3}$** | AT4G15620 | -1360 | *CASPL1E2* | Uncharacterized protein family (UPF0497) |
| 4 | 8914066 | **$6.37\times 10^{-4}$** | 103/A | 2.7 ± 0.3 | 11/G | 5.8 ± 2.1 | **$6.53\times 10^{-3}$** | AT4G15620 | Exon | *CASPL1E2* | Uncharacterized protein family (UPF0497) |
| 4 | 10148636 | **$7.42\times 10^{-4}$** | 110/A | 2.8 ± 0.3 | 4/G | 8.3 ± 5.1 | **$2.45\times 10^{-3}$** | AT4G18360 | Intron | *GOX3* | Glycolate oxidase 3 |
| *Chr4:10148636 - Chr4:10152022* | | | | | | | | AT4G18372 | 2713 | | Small nuclear ribonucleoprotein family protein |
| 4 | 13362035 | **$1.72\times 10^{-4}$** | 103/C | 2.5 ± 0.3 | 11/A | 7.2 ± 2 | **$3.34\times 10^{-5}$** | AT4G26450 | -521 | | Hypothetical protein |

| | | | | | | | | | | | |
|---|---|---|---|---|---|---|---|---|---|---|---|
| 5 | 612228 | **$8.09\times 10^{-4}$** | 84/A | 2.4 ± 0.4 | 30/G | 4.5 ± 1 | **$6.61\times 10^{-3}$** | AT5G02710 | -400 | | Zinc/iron-chelating domain protein |
| 5 | 2767295 | **$1.73\times 10^{-4}$** | 109/C | 2.7 ± 0.3 | 5/G | 8 ± 4.1 | **$1.33\times 10^{-3}$** | AT5G08550 | Intron | *ILP1* | Increased level of polyploidy1-1d |
| | *Chr5:2766133 - Chr5:2772413* | | | | | | | AT5G08560 | 3809 | *WDR26* | Transducin family protein / WD-40 repeat family protein |
| 5 | 2774512 | **$6.12\times 10^{-4}$** | 110/G | 2.8 ± 0.3 | 4/T | 8.7 ± 5.2 | **$1.08\times 10^{-3}$** | AT5G08560 | -685 | *WDR26* | Transducin family protein / WD-40 repeat family protein |
| 5 | 2807059 | **$8.19\times 10^{-4}$** | 109/T | 2.8 ± 0.3 | 5/A | 7.3 ± 4.3 | **$5.92\times 10^{-3}$** | AT5G08650 | Exon | *CPLEPA* | Small GTP-binding protein |
| | *Chr5:2807059 - Chr5:2814169* | | | | | | | AT5G08660 | -7467 | *PSI3* | D-lactate dehydrogenase (DUF668) |
| 5 | 2822494 | **$4.46\times 10^{-5}$** | 106/T | 2.6 ± 0.3 | 8/A | 8.6 ± 3.1 | **$2.57\times 10^{-6}$** | AT5G08680 | Intron | | ATP synthase alpha/beta family protein |
| | *Chr5:2821212 - Chr5:2823797* | | | | | | | AT5G08670 | -1345 | | ATP synthase alpha/beta family protein |
| 5 | 2822703 | **$6.93\times 10^{-4}$** | 109/C | 2.8 ± 0.3 | 5/T | 7.3 ± 4.3 | **$6.40\times 10^{-3}$** | AT5G08680 | Exon | | ATP synthase alpha/beta family protein |
| | *Chr5:2821212 - Chr5:2823797* | | | | | | | AT5G08670 | -1554 | | ATP synthase alpha/beta family protein |
| 5 | 2822975 | **$6.12\times 10^{-4}$** | 110/T | 2.8 ± 0.3 | 4/A | 8.7 ± 5.2 | **$1.08\times 10^{-3}$** | AT5G08680 | Intron | | ATP synthase alpha/beta family protein |
| | *Chr5:2821212 - Chr5:2823797* | | | | | | | AT5G08670 | -1826 | | ATP synthase alpha/beta family protein |
| 5 | 2823332 | **$2.59\times 10^{-4}$** | 108/A | 2.7 ± 0.3 | 6/G | 8.2 ± 3.6 | **$2.11\times 10^{-4}$** | AT5G08680 | Exon | | ATP synthase alpha/beta family protein |
| | | | | | | | | AT5G08660 | 5487 | *PSI3* | D-lactate dehydrogenase (DUF668) |
| | *Chr5:2815853 - Chr5:2828537* | | | | | | | AT5G08690 | -2407 | | ATP synthase alpha/beta family protein |
| 5 | 2824717 | **$6.93\times 10^{-4}$** | 109/C | 2.8 ± 0.3 | 5/T | 7.3 ± 4.3 | **$6.40\times 10^{-3}$** | AT5G08680 | Exon | | ATP synthase alpha/beta family protein |
| | | | | | | | | AT5G08660 | 6872 | *PSI3* | D-lactate dehydrogenase (DUF668) |
| | *Chr5:2817107 - Chr5:2828537* | | | | | | | AT5G08690 | -1022 | | ATP synthase alpha/beta family protein |
| 5 | 3571025 | **$3.86\times 10^{-4}$** | 83/A | 2.5 ± 0.4 | 31/G | 4.2 ± 0.9 | **$2.72\times 10^{-2}$** | AT5G11210 | 189 | *GLR2.5* | Glutamate receptor 2.5 |
| | *Chr5:3562004 - Chr5:3571025* | | | | | | | AT5G11200 | 339 | *UAP56B* | DEAD/DEAH box RNA helicase family protein |
| 5 | 4123962 | **$2.00\times 10^{-4}$** | 92/A | 2.5 ± 0.4 | 22/T | 4.9 ± 1.2 | **$5.21\times 10^{-3}$** | AT5G13010 | Intron | *EMB3011* | RNA helicase family protein |
| | *Chr5:4117470 - Chr5:4123962* | | | | | | | AT5G13000 | -2760 | *GSL12* | Glucan synthase-like 12 |
| 5 | 6358402 | **$8.20\times 10^{-4}$** | 87/C | 2.6 ± 0.4 | 27/G | 4.1 ± 1.1 | $7.26\times 10^{-2}$ | AT5G19030 | Intron | | Cloroplast RRM-containing protein.RNA-binding (RRM/RBD/RNP motifs) family protein |
| 5 | 6794840 | **$1.90\times 10^{-4}$** | 100/C | 2.6 ± 0.3 | 14/G | 5.5 ± 1.7 | **$4.63\times 10^{-3}$** | AT5G20120 | -1049 | | Testis- and ovary-specific PAZ domain protein |
| 5 | 6797770 | **$3.43\times 10^{-4}$** | 98/A | 2.5 ± 0.3 | 16/T | 6 ± 1.7 | **$2.22\times 10^{-4}$** | AT5G20130 | Intron | | Sulfate adenylyltransferase subunit |
| 5 | 9096895 | **$6.30\times 10^{-4}$** | 58/A | 2.2 ± 0.4 | 56/T | 3.8 ± 0.6 | **$1.58\times 10^{-2}$** | AT5G26030 | Exon | *FC1* | Ferrochelatase 1 |
| | *Chr5:9079839 - Chr5:9096895* | | | | | | | AT5G26000 | -14548 | *BGLU38, TGG1* | Thioglucoside glucohydrolase 1 |

Genes within haploblocks of the most significantly associated single-nucleotide polymorphisms (SNPs) in an expression genome-wide association study (eGWAS) of *HIS1-3* superinduction (D2/D1) under recurrent droughts (D1 and D2 separated by a nonstressful recovery interval) in 115 *Arabidopsis thaliana* ecotypes. Top-ranking SNPs having a *P*-value $< 10^{-3}$ are represented in the table. The haploblocks are indicated by bold rectangles. The locations of the SNPs with respect to the nearest genes are shown. Minus and plus signs denote upstream and downstream locations of the SNP with respect to the gene. The relative fold change (RFC) of expression levels of the *A. thaliana* ecotypes having SNP alleles associated with high and low D2/D1 induction are shown as average values with standard error, along with the allele frequencies and a *t*-test *P*-value between these groups.

**Table S5 Genes associated with the most significant SNPs in an expression genome-wide association study of delayed induction of *DREB1A* in *Arabidopsis thaliana* under recurrent droughts**

| Chromosome | Physical position (bp) | GWAS *P*-value | Frequency of low induction allele | RFC *DREB1A* (D2/D1) of low induction allele | Frequency of high induction allele | RFC *DREB1A* (D2/D1) of high induction allele | *t*-test *P*-value (segregation of alleles) | AGI | Region | Gene Symbol | Short Description |
|---|---|---|---|---|---|---|---|---|---|---|---|
| 1 | 4394035 | **$9.85\times 10^{-4}$** | 99/G | 1.7 ± 0.4 | 16/A | 3 ± 0.9 | $1.30\times 10^{-1}$ | AT1G12900 | Intron | *GAPA-2* | Glyceraldehyde 3-phosphate dehydrogenase A subunit 2 |
| | | | | | | | | AT1G12870 | -4925 | | F-box and associated interaction domains-containing protein |
| *Chr1:4388837 - Chr1:4396705* | | | | | | | | AT1G12920 | 2520 | *ERF1-2* | Eukaryotic release factor 1-2 |
| 1 | 4397797 | **$2.37\times 10^{-4}$** | 103/A | 1.8 ± 0.4 | 12/T | 3.2 ± 1.2 | $1.40\times 10^{-1}$ | AT1G12920 | Exon | *ERF1-2* | Eukaryotic release factor 1-2 |
| 1 | 5097674 | **$7.25\times 10^{-4}$** | 102/A | 1.7 ± 0.4 | 13/G | 3.6 ± 1.1 | **$3.60\times 10^{-2}$** | AT1G14790 | Exon | *RDR1* | Rna-dependent rna polymerase 1 |
| | | | | | | | | AT1G14780 | 3801 | *MACP1* | Membrane attack complex/perforin-like 1 |
| | | | | | | | | AT1G14800 | -1898 | | Nucleic acid-binding, OB-fold-like protein |
| | | | | | | | | AT1G14810 | 5010 | | Encodes an aspartate semialdehyde dehydrogenase |
| *Chr1:5093024 - Chr1:5111585* | | | | | | | | AT1G14830 | 10025 | *ADL5* | Arabidopsis dynamin-like protein 5 |
| 1 | 5097963 | **$7.25\times 10^{-4}$** | 102/A | 1.7 ± 0.4 | 13/G | 3.6 ± 1.1 | **$3.60\times 10^{-2}$** | AT1G14790 | Exon | *RDR1* | Rna-dependent rna polymerase 1 |
| | | | | | | | | AT1G14800 | -1609 | | Nucleic acid-binding, OB-fold-like protein |
| | | | | | | | | AT1G14810 | 4721 | | Encodes an aspartate semialdehyde dehydrogenase |
| | | | | | | | | AT1G14820 | 7274 | | Sec14p-like phosphatidylinositol transfer family protein |
| | | | | | | | | AT1G14830 | 9736 | *ADL5* | Arabidopsis dynamin-like protein 5 |
| *Chr1:5094641 - Chr1:5113941* | | | | | | | | AT1G14840 | 14452 | *MAP70-4* | Microtubule-associated proteins 70-4 |
| 1 | 5102442 | **$7.25\times 10^{-4}$** | 102/G | 1.7 ± 0.4 | 13/A | 3.6 ± 1.1 | **$3.60\times 10^{-2}$** | AT1G14800 | Exon | | Nucleic acid-binding, OB-fold-like protein |
| | | | | | | | | AT1G14790 | -4625 | *RDR1* | Rna-dependent rna polymerase 1 |
| | | | | | | | | AT1G14810 | 242 | | Encodes an aspartate semialdehyde dehydrogenase |
| | | | | | | | | AT1G14820 | 2795 | | Sec14p-like phosphatidylinositol transfer family protein |
| | | | | | | | | AT1G14830 | 5257 | *ADL5* | Arabidopsis dynamin-like protein 5 |
| | | | | | | | | AT1G14840 | 9973 | *MAP70-4* | Microtubule-associated proteins 70-4 |
| *Chr1:5094641 - Chr1:5118790* | | | | | | | | AT1G14850 | 14479 | *NUP155* | Nucleoporin 155 |
| 1 | 5102442 | **$7.25\times 10^{-4}$** | 102/G | 1.7 ± 0.4 | 13/A | 3.6 ± 1.1 | **$3.60\times 10^{-2}$** | AT1G14810 | Exon | | Encodes an aspartate semialdehyde dehydrogenase |
| | | | | | | | | AT1G14800 | Exon | | Nucleic acid-binding, OB-fold-like protein |
| | | | | | | | | AT1G14790 | -4625 | *RDR1* | Rna-dependent rna polymerase 1 |
| | | | | | | | | AT1G14820 | 2795 | | Sec14p-like phosphatidylinositol transfer family protein |
| | | | | | | | | AT1G14830 | 5257 | *ADL5* | Arabidopsis dynamin-like protein 5 |
| | | | | | | | | AT1G14840 | 9973 | *MAP70-4* | Microtubule-associated proteins 70-4 |
| *Chr1:5094641 - Chr1:5118790* | | | | | | | | AT1G14850 | 14479 | *NUP155* | Nucleoporin 155 |

| | | | | | | | | | | | |
|---|---|---|---|---|---|---|---|---|---|---|---|
| 1 | 5102675 | **$6.72\times 10^{-4}$** | 100/G | 1.7 ± 0.4 | 15/T | 3.5 ± 1 | **$3.80\times 10^{-2}$** | AT1G14810 | Exon | | Semialdehyde dehydrogenase family protein |
| | | | | | | | | AT1G14790 | -4858 | *RDR1* | Rna-dependent rna polymerase 1 |
| | | | | | | | | AT1G14800 | 117 | | Nucleic acid-binding, OB-fold-like protein |
| | | | | | | | | AT1G14820 | 2562 | | Sec14p-like phosphatidylinositol transfer family protein |
| | | | | | | | | AT1G14830 | 5024 | *ADL5* | Arabidopsis dynamin-like protein 5 |
| | | | | | | | | AT1G14840 | 9740 | *MAP70-4* | Microtubule-associated proteins 70-4 |
| *Chr1:5094806 - Chr1:5120711* | | | | | | | | AT1G14850 | 14246 | *NUP155* | Nucleoporin 155 |
| 1 | 5103585 | **$7.25\times 10^{-4}$** | 102/A | 1.7 ± 0.4 | 13/G | 3.6 ± 1.1 | **$3.60\times 10^{-2}$** | AT1G14810 | Exon | | Encodes an aspartate semialdehyde dehydrogenase |
| 1 | 5597232 | **$5.84\times 10^{-4}$** | 109/C | 1.8 ± 0.3 | 6/T | 3.9 ± 1.8 | $1.20\times 10^{-1}$ | AT1G16370 | Exon | *(OCT6)* | Organic cation/carnitine transporter 6 |
| | | | | | | | | AT1G16340 | -6763 | *ATKDSA2* | Aldolase superfamily protein |
| | | | | | | | | AT1G16360 | -2053 | | LEM3 (ligand-effect modulator 3) family protein / CDC50 family protein |
| *Chr1:5587472 - Chr1:5599872* | | | | | | | | AT1G16380 | 1221 | *CAX1* | Cation exchanger 1 |
| 1 | 5756321 | **$4.39\times 10^{-4}$** | 89/C | 1.7 ± 0.4 | 26/T | 2.7 ± 0.7 | $1.50\times 10^{-1}$ | AT1G16810 | Intron | | 7-dehydrocholesterol reductase-like protein |
| *Chr1:5756321 - Chr1:5757864* | | | | | | | | AT1G16820 | 878 | | Vacuolar ATP synthase catalytic subunit-related / vacuolar proton pump-like protein |
| 1 | 5762904 | **$4.56\times 10^{-4}$** | 84/C | 1.6 ± 0.4 | 31/T | 2.7 ± 0.6 | $1.10\times 10^{-1}$ | AT1G16840 | 207 | | Hypothetical protein |
| 1 | 5763685 | **$4.64\times 10^{-4}$** | 99/A | 1.7 ± 0.4 | 16/G | 3.3 ± 0.9 | $6.10\times 10^{-2}$ | AT1G16840 | Exon | | Hypothetical protein |
| 1 | 21905349 | **$1.18\times 10^{-4}$** | 97/A | 1.6 ± 0.4 | 18/G | 3.3 ± 0.9 | **$3.80\times 10^{-2}$** | AT1G59620 | Exon | *CW9* | Encodes CW9 |
| 1 | 23274203 | **$7.26\times 10^{-4}$** | 93/T | 1.7 ± 0.4 | 22/A | 3 ± 0.6 | $7.80\times 10^{-2}$ | AT1G62850 | Exon | | Class I peptide chain release factor |
| *Chr1:23274203 - Chr1:23275214* | | | | | | | | AT1G62860 | | | Pseudogene of pentatricopeptide (PPR) repeat-containing protein |
| 1 | 24986802 | **$3.31\times 10^{-5}$** | 80/G | 1.2 ± 0.2 | 35/A | 3.6 ± 0.9 | **$1.30\times 10^{-4}$** | AT1G66960 | Exon | *LUP5* | Terpenoid cyclases family protein |
| 1 | 25004491 | **$7.89\times 10^{-5}$** | 100/C | 1.4 ± 0.2 | 15/G | 5.1 ± 1.9 | **$1.00\times 10^{-5}$** | AT1G67000 | Exon | | Protein kinase superfamily protein |
| 2 | 16326247 | **$3.48\times 10^{-4}$** | 80/A | 1.6 ± 0.4 | 35/G | 2.6 ± 0.5 | $1.10\times 10^{-1}$ | AT2G39130 | Exon | | Transmembrane amino acid transporter family protein |
| 2 | 16841888 | **$9.54\times 10^{-4}$** | 94/A | 1.7 ± 0.4 | 21/C | 2.7 ± 0.7 | $1.90\times 10^{-1}$ | AT2G40320 | Exon | *TBL33* | Trichome birefringence-like 33 |
| 3 | 1594337 | **$5.04\times 10^{-4}$** | 94/G | 1.6 ± 0.4 | 21/T | 3.2 ± 0.8 | **$4.50\times 10^{-2}$** | AT3G05500 | Exon | *LDAP3* | Ld-associated protein 3 |
| 3 | 2174061 | **$5.12\times 10^{-4}$** | 112/A | 1.8 ± 0.3 | 3/T | 5.9 ± 2.7 | **$2.60\times 10^{-2}$** | AT3G06880 | Exon | | Transducin/WD40 repeat-like superfamily protein |
| 3 | 3967255 | **$8.70\times 10^{-4}$** | 33/A | 0.9 ± 0.2 | 82/G | 2.3 ± 0.5 | **$3.30\times 10^{-2}$** | AT3G12510 | Exon | | MADS-box family protein |
| | | | | | | | | AT3G12490 | -5379 | *CYS6* | Arabidopsis thaliana phytocystatin 6 |
| | | | | | | | | AT3G12500 | -3310 | *CHI-B, PR3* | Basic chitinase |
| *Chr3:3961815 - Chr3:3970620* | | | | | | | | AT3G12520 | 721 | *SULTR4;2* | Sulfate transporter 4;2 |
| 3 | 8481106 | **$8.31\times 10^{-4}$** | 98/G | 1.7 ± 0.4 | 17/C | 2.9 ± 0.9 | $1.80\times 10^{-1}$ | AT3G23620 | Exon | *ARPF2* | Arabidopsis homolog of yeast rpf2 |
| *Chr3:8479372 - Chr3:8481259* | | | | | | | | AT3G23610 | 1465 | *DSPTP1* | Dual specificity protein phosphatase 1 |
| 3 | 17312716 | **$5.90\times 10^{-4}$** | 108/G | 1.7 ± 0.3 | 7/A | 4.7 ± 1.5 | **$1.60\times 10^{-2}$** | AT3G46990 | Exon | *OPL3* | DUF740 family protein, putative (DUF740) |
| *Chr3:17309901 - Chr3:17314792* | | | | | | | | AT3G47000 | 1095 | | Glycosyl hydrolase family protein |
| 3 | 17723914 | **$4.01\times 10^{-4}$** | 104/T | 1.5 ± 0.2 | 11/C | 5.8 ± 2.5 | **$9.30\times 10^{-6}$** | AT3G48010 | Exon | *CNGC16* | Cyclic nucleotide-gated channel 16 |
| *Chr3:17722899 - Chr3:17725635* | | | | | | | | AT3G48030 | 1496 | *ATL48* | Arabidopsis toxicos en levadura 48 |

| | | | | | | | | | | | |
|---|---|---|---|---|---|---|---|---|---|---|---|
| 3 | 17725048 | **$4.42\times 10^{-4}$** | 102/A | 1.5 ± 0.2 | 13/T | 5.4 ± 2.2 | **$1.40\times 10^{-5}$** | AT3G48020 | Exon | | Hypothetical protein |
| *Chr3:17725048 - Chr3:17726259* | | | | | | | | AT3G48030 | 362 | *ATL48* | Arabidopsis toxicos en levadura 48 |
| 3 | 17726259 | **$2.31\times 10^{-4}$** | 101/A | 1.4 ± 0.2 | 14/T | 5.2 ± 2 | **$1.30\times 10^{-5}$** | AT3G48030 | Intron | *ATL48* | Arabidopsis toxicos en levadura 48 |
| | | | | | | | | AT3G48020 | 1259 | | Hypothetical protein |
| *Chr3:17725048 - Chr3:17737234* | | | | | | | | AT3G48050 | 8011 | *SUO* | Shuttle' in chinese |
| 3 | 17936675 | **$2.40\times 10^{-4}$** | 95/G | 1.6 ± 0.4 | 20/C | 3.2 ± 0.8 | **$4.80\times 10^{-2}$** | AT3G48430 | Intron | *JMJ12* | Jumonji domain-containing protein 12 |
| *Chr3:17934266 - Chr3:17940650* | | | | | | | | AT3G48425 | 2182 | *APE1L* | DNAse I-like superfamily protein |
| 3 | 17943693 | **$2.40\times 10^{-4}$** | 95/T | 1.6 ± 0.4 | 20/A | 3.2 ± 0.8 | **$4.80\times 10^{-2}$** | AT3G48440 | 117 | *ATC3H43* | Zinc finger C-x8-C-x5-C-x3-H type family protein |
| 3 | 18581486 | **$5.37\times 10^{-4}$** | 90/C | 1.4 ± 0.2 | 25/A | 3.8 ± 1.2 | **$4.00\times 10^{-4}$** | AT3G50110 | Exon | *PTEN3* | *Arabidopsis thaliana* phosphatase and tensin homolog deleted on chromosome ten 3 |
| 3 | 18934040 | **$8.37\times 10^{-4}$** | 105/A | 1.6 ± 0.2 | 10/T | 5.3 ± 2.7 | **$2.40\times 10^{-4}$** | AT3G50940 | Intron | | P-loop containing nucleoside triphosphate hydrolases superfamily protein |
| 3 | 22795973 | **$7.12\times 10^{-5}$** | 106/T | 1.7 ± 0.3 | 9/C | 4.5 ± 1.3 | **$1.10\times 10^{-2}$** | AT3G61600 | Intron | *POB1* | Poz/btb containin g-protein 1 |
| *Chr3:22793523 - Chr3:22796077* | | | | | | | | AT3G61590 | 1824 | *HWS* | Hawaiian skirt |
| 3 | 22800750 | **$1.15\times 10^{-4}$** | 105/G | 1.7 ± 0.3 | 10/A | 4.2 ± 1.2 | **$1.40\times 10^{-2}$** | AT3G61610 | Intron | | Galactose mutarotase-like superfamily protein |
| *Chr3:22795835 - Chr3:22800750* | | | | | | | | AT3G61600 | 2681 | *POB1* | Poz/btb containin g-protein 1 |
| 3 | 22801488 | **$4.50\times 10^{-4}$** | 104/A | 1.7 ± 0.3 | 11/C | 3.9 ± 1.1 | **$2.60\times 10^{-2}$** | AT3G61620 | Intron | *RRP41* | Exonuclease RRP41 |
| 4 | 309005 | **$7.39\times 10^{-4}$** | 112/C | 1.8 ± 0.3 | 3/G | 5.9 ± 2.5 | **$2.70\times 10^{-2}$** | AT4G00740 | Exon | *QUA3* | Quasimodo 3 |
| 4 | 9229870 | **$8.59\times 10^{-4}$** | 94/A | 1.7 ± 0.4 | 21/T | 2.8 ± 0.8 | $1.40\times 10^{-1}$ | AT4G16340 | Exon | *SPK1* | Spike 1 |
| *Chr4:9222034 - Chr4:9229870* | | | | | | | | AT4G16310 | 5106 | *LDL3* | LSD-1 like 3 |
| 4 | 10422057 | **$6.28\times 10^{-4}$** | 93/T | 1.6 ± 0.4 | 22/G | 3 ± 0.7 | $6.10\times 10^{-2}$ | AT4G19030 | Exon | *NLM1* | Nod26-like major intrinsic protein 1 |
| 4 | 11287349 | **$1.68\times 10^{-4}$** | 106/G | 1.7 ± 0.3 | 9/A | 4.5 ± 1.4 | **$1.10\times 10^{-2}$** | AT4G21170 | Exon | | Tetratricopeptide repeat (TPR)-like superfamily protein |
| 4 | 11308042 | **$3.95\times 10^{-4}$** | 102/C | 1.7 ± 0.4 | 13/T | 3.6 ± 1.1 | **$4.10\times 10^{-2}$** | AT4G21210 | Exon | *RP1* | PPDK regulatory protein |
| *Chr4:11306012 - Chr4:11308889* | | | | | | | | AT4G21200 | 1441 | *GA2OX8* | *Arabidopsis thaliana* gibberellin 2-oxidase 8 |
| 4 | 11338818 | **$7.84\times 10^{-4}$** | 101/T | 1.7 ± 0.4 | 14/G | 3.4 ± 1 | $6.00\times 10^{-2}$ | AT4G21300 | Exon | *ATDEK36* | Tetratricopeptide repeat (TPR)-like superfamily protein |
| | | | | | | | | AT4G21310 | 316 | *DEAL2* | DUF1218 family member |
| *Chr4:11338818 - Chr4:11340653* | | | | | | | | AT4G21320 | -1674 | *HSA32* | Heat-stress-associated 32 |
| 4 | 13950736 | **$6.19\times 10^{-4}$** | 99/T | 1.7 ± 0.4 | 16/G | 3.2 ± 0.9 | $7.00\times 10^{-2}$ | AT4G28080 | Exon | *REC2* | Reduced chloroplast coverage 2 |
| 4 | 16057037 | **$7.92\times 10^{-4}$** | 60/A | 1 ± 0.2 | 55/G | 2.9 ± 0.6 | **$1.30\times 10^{-3}$** | AT4G33320 | Exon | *CHIQL2* | DUF641 domain protein, probable psuedogene |
| 4 | 16386814 | **$4.78\times 10^{-4}$** | 104/A | 1.5 ± 0.2 | 11/G | 6 ± 2.5 | **$2.30\times 10^{-6}$** | AT4G34230 | Exon | *CAD5* | Cinnamyl alcohol dehydrogenase 5 |
| 5 | 368967 | **$7.13\times 10^{-4}$** | 108/T | 1.5 ± 0.2 | 7/A | 7.4 ± 3.8 | **$5.40\times 10^{-7}$** | AT5G01950 | Intron | | Leucine-rich repeat protein kinase family protein |
| 5 | 4073312 | **$8.57\times 10^{-4}$** | 100/G | 1.4 ± 0.2 | 15/C | 5 ± 1.9 | **$3.20\times 10^{-5}$** | AT5G12900 | Exon | | DNA double-strand break repair RAD50 ATPase |
| | | | | | | | | AT5G12920 | 4605 | | Transducin/WD40 repeat-like superfamily protein |
| *Chr5:4073312 - Chr5:4084108* | | | | | | | | AT5G12930 | 9843 | | Inactive rhomboid protein |
| 5 | 6166559 | **$6.47\times 10^{-4}$** | 106/T | 1.7 ± 0.3 | 9/C | 4.3 ± 1.2 | **$1.70\times 10^{-2}$** | AT5G18560 | -654 | *PUCHI* | Encodes PUCHI, a member of the ERF subfamily B-1 |
| *Chr5:6160520 - Chr5:6166559* | | | | | | | | AT5G18550 | 3830 | *ATC3H58* | Zinc finger C-x8-C-x5-C-x3-H type family protein |
| 5 | 6459439 | **$3.33\times 10^{-5}$** | 103/G | 1.8 ± 0.4 | 12/T | 3.2 ± 1.2 | $1.30\times 10^{-1}$ | AT5G19200 | Exon | *TSC10B* | NAD(P)-binding Rossmann-fold superfamily protein |

| | | | | | | | | | | | |
|---|---|---|---|---|---|---|---|---|---|---|---|
| | | | | | | | | AT5G19190 | 1521 | | Hypothetical protein |
| | | | | | | | | AT5G19210 | -2005 | | P-loop containing nucleoside triphosphate hydrolases superfamily protein |
| *Chr5:6457786 - Chr5:6466043* | | | | | | | | AT5G19220 | 4492 | *ADG2* | ADP glucose pyrophosphorylase 2 |
| 5 | 7953822 | **$3.14\times 10^{-4}$** | 67/T | $1.2 \pm 0.2$ | 48/C | $2.8 \pm 0.7$ | **$6.90\times 10^{-3}$** | AT5G23590 | 48 | | DNAJ heat shock N-terminal domain-containing protein |
| *Chr5:7948442 - Chr5:7953822* | | | | | | | | AT5G23575 | 3781 | | Transmembrane CLPTM1 family protein |
| 5 | 7960433 | **$3.79\times 10^{-4}$** | 86/G | $1.4 \pm 0.2$ | 29/T | $3.5 \pm 1.1$ | **$1.70\times 10^{-3}$** | AT5G23610 | -465 | | DYAD protein |
| | | | | | | | | AT5G23580 | -8000 | *CPK12* | *Arabidopsis thaliana* calmodulin-like domain protein kinase 9 |
| *Chr5:7950453 - Chr5:7962579* | | | | | | | | AT5G23630 | 323 | *MIA* | Male gametogenesis impaired anthers |
| 5 | 7960433 | **$3.79\times 10^{-4}$** | 86/G | $1.4 \pm 0.2$ | 29/T | $3.5 \pm 1.1$ | **$1.70\times 10^{-3}$** | AT5G23630 | 323 | *MIA* | Male gametogenesis impaired anthers |
| | | | | | | | | AT5G23610 | -465 | | DYAD protein |
| *Chr5:7950453 - Chr5:7962579* | | | | | | | | AT5G23580 | -8000 | *CPK12* | *Arabidopsis thaliana* calmodulin-like domain protein kinase 9 |
| 5 | 9209620 | **$2.67\times 10^{-4}$** | 111/G | $1.8 \pm 0.3$ | 4/A | $5 \pm 1.5$ | **$4.30\times 10^{-2}$** | AT5G26280 | Exon | | TRAF-like family protein |
| 5 | 13908934 | **$9.88\times 10^{-4}$** | 95/G | $1.7 \pm 0.4$ | 20/C | $2.9 \pm 0.7$ | $1.00\times 10^{-1}$ | AT5TE49575 | 614 | | ATREP9 transposon family,RC/Helitron transposon super family |
| 5 | 13910038 | **$5.30\times 10^{-4}$** | 96/T | $1.7 \pm 0.4$ | 19/G | $3.1 \pm 0.7$ | $7.50\times 10^{-2}$ | AT5G35740 | 534 | | Carbohydrate-binding X8 domain superfamily protein |
| *Chr5:13908380 - Chr5:13913339* | | | | | | | | AT5G35750 | 1705 | *HK2* | Histidine kinase 2 |
| 5 | 13910408 | **$5.30\times 10^{-4}$** | 96/G | $1.7 \pm 0.4$ | 19/T | $3.1 \pm 0.7$ | $7.50\times 10^{-2}$ | AT5G35740 | 164 | | Carbohydrate-binding X8 domain superfamily protein |
| *Chr5:13908380 - Chr5:13913339* | | | | | | | | AT5G35750 | 1335 | *HK2* | Histidine kinase 2 |
| 5 | 13910590 | **$5.28\times 10^{-4}$** | 95/C | $1.7 \pm 0.4$ | 95/C | $1.7 \pm 0.4$ | $1.00\times 10^{-0}$ | AT5G35740 | Exon | | Carbohydrate-binding X8 domain superfamily protein |
| 5 | 13912805 | **$4.86\times 10^{-4}$** | 98/G | $1.7 \pm 0.4$ | 17/A | $3.1 \pm 0.8$ | $8.00\times 10^{-2}$ | AT5G35750 | Exon | *HK2* | Histidine kinase 2 |
| 5 | 13913597 | **$1.50\times 10^{-4}$** | 97/G | $1.7 \pm 0.4$ | 18/T | $3.2 \pm 0.7$ | $5.20\times 10^{-2}$ | AT5G35750 | Exon | *HK2* | Histidine kinase 2 |
| *Chr5:13909662 - Chr5:13924157* | | | | | | | | AT5G35753 | 8972 | | DnaJ heat shock amino-terminal domain protein (DUF3444) |
| 5 | 13926798 | **$1.50\times 10^{-4}$** | 97/G | $1.7 \pm 0.4$ | 18/A | $3.2 \pm 0.7$ | $5.20\times 10^{-2}$ | AT5G35753 | -1878 | | DnaJ heat shock amino-terminal domain protein (DUF3444) |
| 5 | 13930156 | **$1.50\times 10^{-4}$** | 97/T | $1.7 \pm 0.4$ | 18/G | $3.2 \pm 0.7$ | $5.20\times 10^{-2}$ | AT5TE49650 | Exon | | ATLINE1_2 transposon family,LINE/L1 transposon super family |
| 5 | 13930909 | **$1.50\times 10^{-4}$** | 96/G | $1.7 \pm 0.4$ | 19/T | $3.2 \pm 0.7$ | $5.10\times 10^{-2}$ | AT5TE49655 | Exon | | ATCOPIA8A transposon family,LTR/Copia transposon super family |
| 5 | 14954326 | **$4.58\times 10^{-4}$** | 90/T | $1.4 \pm 0.2$ | 25/A | $3.8 \pm 1.2$ | **$7.10\times 10^{-4}$** | AT5G37650 | Exon | | Hypothetical protein (DUF577) |
| 5 | 14955338 | **$8.39\times 10^{-4}$** | 83/T | $1.4 \pm 0.2$ | 32/C | $3.2 \pm 1$ | **$4.40\times 10^{-3}$** | AT5G37650 | Exon | | Hypothetical protein (DUF577) |
| 5 | 16111268 | **$3.45\times 10^{-4}$** | 82/T | $1.6 \pm 0.4$ | 33/C | $2.7 \pm 0.5$ | $9.10\times 10^{-2}$ | AT5G40300 | Intron | *CASPL4A1* | Uncharacterized protein family (UPF0497) |
| 5 | 17130737 | **$7.41\times 10^{-4}$** | 93/G | $1.6 \pm 0.4$ | 22/A | $3.2 \pm 0.8$ | **$3.30\times 10^{-2}$** | AT5G42720 | Exon | | Glycosyl hydrolase family 17 protein |
| 5 | 20040327 | **$1.66\times 10^{-4}$** | 107/A | $1.7 \pm 0.3$ | 8/G | $4.5 \pm 1.1$ | **$1.30\times 10^{-2}$** | AT5G49430 | Exon | | WD40/YVTN repeat and Bromo-WDR9-I-like domain-containing protein |
| 5 | 21112829 | **$6.81\times 10^{-4}$** | 97/G | $1.7 \pm 0.4$ | 18/A | $3.3 \pm 0.9$ | **$4.80\times 10^{-2}$** | AT5G51970 | Exon | *ATSDH* | Sorbitol dehydrogenase |
| | | | | | | | | AT5G51960 | 1713 | | Complex 1 protein, LYR family protein |
| | | | | | | | | AT5G51980 | 821 | *ATC3H63* | Transducin/WD40 repeat-like superfamily protein |
| *Chr5:21111077 -Chr5:21117488* | | | | | | | | AT5G51990 | 4284 | *DREB1D* | Dehydration-responsive element-binding protein 1D |
| 5 | 22206266 | **$8.08\times 10^{-5}$** | 97/G | $1.7 \pm 0.4$ | 18/A | $3 \pm 0.8$ | $1.10\times 10^{-1}$ | AT5G54660 | 1280 | | HSP20-like chaperones superfamily protein |
| 5 | 24945075 | **$1.60\times 10^{-5}$** | 112/C | $1.8 \pm 0.3$ | 3/T | $7.4 \pm 1.5$ | **$2.10\times 10^{-3}$** | AT5G62110 | Exon | | Homeodomain-like superfamily protein |

| | | | | |
|---|---|---|---|---|
| Chr5:24938349 - Chr5:24945075 | AT5G62090 | -6535 | *SLK2* | Seuss-like 2 |

Genes within haploblocks of the most significantly associated single-nucleotide polymorphisms (SNPs) in an expression genome-wide association study (eGWAS) of delayed induction (D2/D1) of *DREB1A* under recurrent droughts (D1 and D2 separated by a nonstressful recovery interval) in 115 *Arabidopsis thaliana* ecotypes. Top-ranking SNPs having a *P*-value < $10^{-3}$ are represented in the table. The haploblocks are indicated by bold rectangles. The locations of the SNPs with respect to the nearest genes are shown. Minus and plus signs denote upstream and downstream locations of the SNP with respect to the gene. The relative fold change (RFC) of expression levels of the *A. thaliana* ecotypes having SNP alleles associated with high and low D2/D1 induction are shown as average values with standard error, along with the allele frequencies and a *t*-test *P*-value between these groups.

#### Table S6 Memory behavior of eGWAS-delineated genes

| AGI code | Gene Symbol | Short Description | D1 vs W | | | R vs W | | | D2 vs W | | | D2 vs D1 | | | Category of memory behavior |
|---|---|---|---|---|---|---|---|---|---|---|---|---|---|---|---|
| | | | $log_2FC$ | *P*-value | *FDR* | $log_2FC$ | *P*-value | *FDR* | $log_2FC$ | *P*-value | *FDR* | $log_2FC$ | *P*-value | *FDR* | |
| ***LKR* eGWAS** | | | | | | | | | | | | | | | |
| AT1G02720 | *GATL5* | Galacturonosyltransferase-like 5 | | | | | | | -0.3 | **4.94× $10^{-2}$** | 1.31× $10^{-1}$ | | | | |
| AT1G11650 | *RBP45B* | RNA-binding (RRM/RBD/RNP motifs) family protein | | | | -0.2 | **5.43× $10^{-2}$** | 1.41× $10^{-1}$ | | | | | | | |
| AT1G11650 | *RBP45B* | RNA-binding (RRM/RBD/RNP motifs) family protein | | | | -0.2 | **5.43× $10^{-2}$** | 1.41× $10^{-1}$ | | | | | | | |
| AT1G27680 | *APL2* | ADPGLC-PPASE large subunit | | | | -0.4 | **1.52× $10^{-2}$** | **5.29× $10^{-2}$** | | | | | | | |
| AT1G50580 | | UDP-Glycosyltransferase superfamily protein | 1.0 | **2.01× $10^{-2}$** | 9.31× $10^{-2}$ | 1.1 | **1.49× $10^{-3}$** | **8.38× $10^{-3}$** | 0.9 | **1.98× $10^{-2}$** | 6.39× $10^{-2}$ | | | | |
| AT1G56660 | | MAEBL domain protein | | | | | | | | | | | | | |
| AT1G56670 | | GDSL-motif esterase/acyltransferase/lipase | -0.5 | **4.35× $10^{-2}$** | 1.62× $10^{-1}$ | -0.5 | **9.91× $10^{-3}$** | **3.80× $10^{-2}$** | -0.7 | **1.27× $10^{-4}$** | **8.75× $10^{-4}$** | | | | |
| AT1G56670 | | GDSL-motif esterase/acyltransferase/lipase | -0.5 | **4.35× $10^{-2}$** | 1.62× $10^{-1}$ | -0.5 | **9.91× $10^{-3}$** | **3.80× $10^{-2}$** | -0.7 | **1.27× $10^{-4}$** | **8.75× $10^{-4}$** | | | | |
| AT1G60280 | *NAC023* | NAC domain containing protein 23 | | | | | | | | | | | | | |
| AT1G61260 | | Cotton fiber (DUF761) | | | | | | | | | | | | | |
| AT1G62320 | *OSCA1.4* | ERD (early-responsive to dehydration stress) family protein | 1.6 | **3.34× $10^{-4}$** | **3.76× $10^{-3}$** | 1.2 | **6.19× $10^{-4}$** | **4.07× $10^{-3}$** | 1.4 | **4.09× $10^{-5}$** | **3.17× $10^{-4}$** | | | | |
| AT1G65630 | *DEG3* | Degp protease 3 | | | | | | | | | | | | | |
| AT1G65630 | *DEG3* | Degp protease 3 | | | | | | | | | | | | | |
| AT1G67210 | *ZCCHC8B* | Proline-rich spliceosome-associated (PSP) family protein | | | | | | | | | | | | | |
| AT1G67190 | | F-box/RNI-like superfamily protein | | | | | | | | | | | | | |
| AT1G67210 | *ZCCHC8B* | Proline-rich spliceosome-associated (PSP) family protein | | | | | | | | | | | | | |
| AT1G08913 | | Long non-coding RNA | | | | | | | | | | | | | |
| AT1G68320 | *MYB62* | Myb domain protein 62 | | | | | | | | | | | | | |
| AT1G70210 | *CYCD1;1* | Cyclin D1 | | | | 0.4 | **1.91× $10^{-2}$** | 6.36× $10^{-2}$ | | | | | | | |
| AT1G70220 | | RNA-processing, Lsm domain-containing protein | | | | | | | | | | | | | |
| AT1G71040 | *LPR2* | Low phosphate root2 | | | | | | | 0.3 | **1.78× $10^{-2}$** | **5.88× $10^{-2}$** | 0.5 | **2.14× $10^{-2}$** | 5.45× $10^{-1}$ | |
| AT1G74090 | *SOT18* | Arabidopsis sulfotransferase 5b | | | | -0.4 | **8.48× $10^{-3}$** | **3.37× $10^{-2}$** | | | | | | | |
| AT2G02960 | | RING/FYVE/PHD zinc finger superfamily protein | 0.7 | **4.89× $10^{-6}$** | **9.91× $10^{-5}$** | 0.8 | **5.19× $10^{-6}$** | **6.94× $10^{-5}$** | 0.4 | **6.49× $10^{-4}$** | **3.64× $10^{-3}$** | | | | |
| AT2TE20600 | | ATHILA4A transposon family,LTR/Gypsy transposon super family | | | | | | | | | | | | | |

| | | | | | | | | | | | | | | |
|---|---|---|---|---|---|---|---|---|---|---|---|---|---|---|
| AT2TE20730 | | ATLANTYS1 transposon family,LTR/Gypsy transposon super family | | | | | | | | | | | | |
| AT2G14255 | | Ankyrin repeat family protein with DHHC zinc finger domain-containing protein | | | | | | | | | | | | |
| AT2G17540 | | Hypothetical protein | | | | | | | -0.3 | **$2.69\times10^{-2}$** | **$8.18\times10^{-2}$** | | | |
| AT2G17723 | | Defensin-like (DEFL) family protein | | | | | | | | | | | | |
| AT2G19110 | *HMA4* | Arabidopsis heavy metal ATPase 4 | 0.9 | **$3.67\times10^{-4}$** | **$4.07\times10^{-3}$** | | | | 1.0 | **$2.88\times10^{-7}$** | **$3.51\times10^{-6}$** | | | |
| AT2G19120 | | P-loop containing nucleoside triphosphate hydrolases superfamily protein | | | | | | | | | | | | |
| AT2G19120 | | P-loop containing nucleoside triphosphate hydrolases superfamily protein | | | | | | | | | | | | |
| AT2G19120 | | P-loop containing nucleoside triphosphate hydrolases superfamily protein | | | | | | | | | | | | |
| AT2G20300 | *ALE2* | Abnormal leaf shape 2 | | | | | | | | | | | | |
| AT2G21110 | *DIR4* | Dirigent protein 4 | | | | | | | | | | | | |
| AT2G21080 | | Ras guanine nucleotide exchange factor K | | | | | | | -0.6 | **$3.50\times10^{-3}$** | **$1.54\times10^{-2}$** | | | |
| AT2G21090 | | Pentatricopeptide repeat (PPR-like) superfamily protein | | | | | | | | | | | | |
| AT2G21120 | *AVI2, ENOR3* | Magnesium transporter, putative (DUF803) | | | | | | | | | | | | |
| AT2G24960 | | Myb/SANT-like DNA-binding domain protein | | | | 1.1 | **$1.71\times10^{-4}$** | **$1.40\times10^{-3}$** | | | | | | |
| AT2G24960 | | Myb/SANT-like DNA-binding domain protein | | | | 1.1 | **$1.71\times10^{-4}$** | **$1.40\times10^{-3}$** | | | | | | |
| AT2G24970 | | Spindle/kinetochore-associated protein | | | | | | | | | | | | |
| AT2G25010 | *MAIL1* | Aminotransferase-like, plant mobile domain family protein | | | | | | | | | | | | |
| AT2G25520 | | Drug/metabolite transporter superfamily protein | | | | -0.3 | **$5.35\times10^{-2}$** | $1.40\times10^{-1}$ | | | | | | |
| AT2G29020 | | Rab5-interacting family protein | | | | -0.3 | **$5.81\times10^{-3}$** | **$2.50\times10^{-2}$** | | | | | | |
| AT2G29060 | | GRAS family transcription factor | 1.2 | **$6.29\times10^{-4}$** | **$6.28\times10^{-3}$** | 1.8 | **$1.11\times10^{-6}$** | **$1.79\times10^{-5}$** | 1.3 | **$5.82\times10^{-4}$** | **$3.32\times10^{-3}$** | | | |
| AT2G29070 | | Ubiquitin fusion degradation UFD1 family protein | | | | | | | 0.4 | **$4.88\times10^{-2}$** | $1.30\times10^{-1}$ | | | |
| AT2G29080 | *FTSH3* | FTSH protease 3 | | | | | | | | | | | | |
| AT2G34660 | *ABCC2* | Multidrug resistance-associated protein 2 | -0.4 | **$4.04\times10^{-2}$** | $1.53\times10^{-1}$ | | | | -0.4 | **$1.62\times10^{-2}$** | $5.44\times10^{-2}$ | | | |
| AT3G04240 | *SEC* | Tetratricopeptide repeat (TPR)-like superfamily protein | | | | | | | | | | | | |
| AT3G04320 | | Kunitz family trypsin and protease inhibitor protein | 3.0 | **$2.00\times10^{-6}$** | **$4.53\times10^{-5}$** | 2.6 | **$7.88\times10^{-11}$** | **$3.07\times10^{-9}$** | 3.2 | **$2.00\times10^{-17}$** | **$1.03\times10^{-15}$** | | | |
| AT3G04290 | *LTL1* | Li-tolerant lipase 1 | | | | | | | -0.5 | **$4.35\times10^{-4}$** | **$2.57\times10^{-3}$** | -0.4 | **$1.04\times10^{-2}$** | $3.76\times10^{-1}$ |
| AT3G04300 | | RmlC-like cupins superfamily protein | | | | | | | | | | | | |
| AT3G04310 | | Transmembrane protein | | | | | | | | | | | | |
| AT3G04340 | *TSL2* | FtsH extracellular protease family | | | | 0.3 | **$3.32\times10^{-2}$** | $9.73\times10^{-2}$ | | | | | | |
| AT3G04350 | | Vacuolar sorting-associated protein (DUF946) | | | | 0.3 | **$3.10\times10^{-2}$** | $9.24\times10^{-2}$ | | | | | | |
| AT3G04340 | *TSL2* | FtsH extracellular protease family | | | | 0.3 | **$3.32\times10^{-2}$** | $9.73\times10^{-2}$ | | | | | | |
| AT3G04350 | | Vacuolar sorting-associated protein (DUF946) | | | | 0.3 | **$3.10\times10^{-2}$** | $9.24\times10^{-2}$ | | | | | | |
| AT3G04340 | *TSL2* | FtsH extracellular protease family | | | | 0.3 | **$3.32\times10^{-2}$** | $9.73\times10^{-2}$ | | | | | | |
| AT3G04360 | | Calcium-dependent lipid-binding (CaLB domain) family protein | | | | | | | | | | | | |
| AT3G04360 | | Calcium-dependent lipid-binding (CaLB domain) family protein | | | | | | | | | | | | |
| AT3G04340 | *TSL2* | FtsH extracellular protease family | | | | 0.3 | **$3.32\times10^{-2}$** | $9.73\times10^{-2}$ | | | | | | |
| AT3G04350 | | Vacuolar sorting-associated protein (DUF946) | | | | 0.3 | **$3.10\times10^{-2}$** | $9.24\times10^{-2}$ | | | | | | |
| AT3G04360 | | Calcium-dependent lipid-binding (CaLB domain) family protein | | | | | | | | | | | | |
| AT3G04350 | | Vacuolar sorting-associated protein (DUF946) | | | | 0.3 | **$3.10\times10^{-2}$** | $9.24\times10^{-2}$ | | | | | | |

| | | | | | | | | | | | | | | |
|---|---|---|---|---|---|---|---|---|---|---|---|---|---|---|
| AT3G10845 | | RNA-binding (RRM/RBD/RNP motifs) family protein | | | | | | | | | | | | |
| AT3G10845 | | RNA-binding (RRM/RBD/RNP motifs) family protein | | | | | | | | | | | | |
| AT3G10820 | *MED26A* | Transcription elongation factor (TFIIS) family protein | | | | | | | | | | | | |
| AT3G10850 | *GLX2-2* | Metallo-hydrolase/oxidoreductase superfamily protein | -0.4 | **$4.83\times 10^{-2}$** | $1.74\times 10^{-1}$ | -0.5 | **$1.40\times 10^{-4}$** | **$1.19\times 10^{-3}$** | | | | | | |
| AT3G15430 | | Regulator of chromosome condensation (RCC1) family protein | | | | | | | | | | | | |
| AT3G16610 | | Pentatricopeptide (PPR) repeat-containing protein | | | | | | | | | | | | |
| AT3G16650 | *PRL2* | Transducin/WD40 repeat-like superfamily protein | | | | | | | | | | | | |
| AT3G16640 | *TCTP1* | Translationally controlled tumor protein | | | | | | | -0.3 | **$4.09\times 10^{-2}$** | $1.13\times 10^{-1}$ | | | |
| AT3G18773 | *ATL77* | RING/U-box superfamily protein | | | | | | | | | | | | |
| AT3G23480 | | Cyclopropane-fatty-acyl-phospholipid synthase | | | | 0.7 | **$1.79\times 10^{-2}$** | $6.04\times 10^{-2}$ | | | | | | |
| AT3G52460 | | Hydroxyproline-rich glycoprotein family protein | 0.7 | **$1.73\times 10^{-2}$** | $8.35\times 10^{-2}$ | 0.4 | $9.04\times 10^{-2}$ | $2.07\times 10^{-1}$ | 0.7 | **$3.47\times 10^{-2}$** | $9.96\times 10^{-2}$ | | | |
| AT3G52530 | *ZRK11* | Protein kinase superfamily protein | | | | | | | | | | | | |
| AT3G62700 | *MRP10* | Multidrug resistance-associated protein 10 | | | | | | | | | | | | |
| AT3G62700 | *MRP10* | Multidrug resistance-associated protein 10 | | | | | | | | | | | | |
| AT3G62700 | *MRP10* | Multidrug resistance-associated protein 10 | | | | | | | | | | | | |
| AT4G00320 | | F-box/RNI-like superfamily protein | | | | | | | | | | | | |
| AT4G05150 | | Octicosapeptide/Phox/Bem1p family protein | -0.6 | **$2.16\times 10^{-3}$** | **$1.68\times 10^{-2}$** | -0.4 | **$2.76\times 10^{-2}$** | $8.46\times 10^{-2}$ | | | | 0.5 | **$1.19\times 10^{-2}$** | $4.05\times 10^{-1}$ |
| AT4G11730 | | Cation transporter/ E1-E2 ATPase family protein | | | | | | | | | | | | |
| AT4G11740 | *SAY1* | Ubiquitin-like superfamily protein | | | | | | | | | | | | |
| AT4G12010 | *DSC1* | Disease resistance protein (TIR-NBS-LRR class) family | | | | | | | | | | | | |
| AT4G13660 | *PRR2* | Pinoresinol reductase 2 | 1.1 | **$3.46\times 10^{-3}$** | **$2.43\times 10^{-2}$** | 1.0 | **$1.43\times 10^{-3}$** | **$8.08\times 10^{-3}$** | 1.1 | **$6.73\times 10^{-4}$** | **$3.76\times 10^{-3}$** | | | |
| AT4G15955 | | Alpha/beta-Hydrolases superfamily protein | | | | -0.4 | **$2.54\times 10^{-2}$** | **$7.94\times 10^{-2}$** | | | | | | |
| AT4G16490 | | ARM repeat superfamily protein | | | | | | | | | | | | |
| AT4G20160 | | Golgin family A protein | 2.8 | **$2.94\times 10^{-5}$** | **$4.68\times 10^{-4}$** | 2.9 | **$1.42\times 10^{-9}$** | **$4.21\times 10^{-8}$** | 2.8 | **$1.46\times 10^{-10}$** | **$3.18\times 10^{-9}$** | | | |
| AT4G21150 | *HAP6* | Ribophorin II (RPN2) family protein | 0.4 | **$2.27\times 10^{-2}$** | $1.02\times 10^{-1}$ | 0.4 | **$1.13\times 10^{-2}$** | **$4.22\times 10^{-2}$** | 0.3 | **$1.17\times 10^{-2}$** | **$4.18\times 10^{-2}$** | | | |
| AT4G23750 | *CRF2* | Cytokinin response factor 2 | 0.6 | **$3.29\times 10^{-2}$** | $1.33\times 10^{-1}$ | | | | 0.4 | **$1.90\times 10^{-3}$** | **$9.08\times 10^{-3}$** | | | |
| AT4G25970 | *PSD3* | Phosphatidylserine decarboxylase 3 | | | | | | | | | | | | |
| AT4G25970 | *PSD3* | Phosphatidylserine decarboxylase 3 | | | | | | | | | | | | |
| AT4G26090 | *RPS2* | Resistant to P. Syringae 2 | | | | | | | | | | | | |
| AT4G26100 | *CKL1* | Casein kinase 1 | | | | | | | | | | | | |
| AT4G28300 | *FLOE1* | Formin-like protein (DUF1421) | | | | | | | | | | | | |
| AT4G28260 | | Acyl-UDP-N-acetylglucosamine O-acyltransferase | | | | | | | | | | | | |
| AT4G29580 | | Cytidine/deoxycytidylate deaminase family protein | | | | | | | | | | | | |
| AT4G29590 | | Putative methyltransferase | -0.5 | **$5.18\times 10^{-3}$** | **$3.35\times 10^{-2}$** | -0.5 | **$2.56\times 10^{-4}$** | **$1.96\times 10^{-3}$** | -0.5 | **$7.22\times 10^{-4}$** | **$4.00\times 10^{-3}$** | | | |
| AT4G37970 | *CAD6* | Cinnamyl alcohol dehydrogenase 6 | | | | -0.6 | **$4.09\times 10^{-3}$** | **$1.88\times 10^{-2}$** | | | | | | |
| AT4G38070 | | Transcription factor bHLH131-like protein | | | | | | | | | | | | |
| AT4G38062 | | Basic helix loop helix (bHLH) DNA-binding superfamily protein | | | | | | | | | | | | |
| AT5G04020 | | Calmodulin binding protein | | | | 0.7 | **$6.09\times 10^{-4}$** | **$4.02\times 10^{-3}$** | 0.3 | **$2.69\times 10^{-2}$** | $8.18\times 10^{-2}$ | | | |
| AT5G04010 | | F-box family protein | | | | | | | | | | | | |

| | | | | | | | | | | | |
|---|---|---|---|---|---|---|---|---|---|---|---|
| AT5G04020 | | Calmodulin binding protein | | | | 0.7 | **$6.09\times 10^{-4}$** | **$4.02\times 10^{-3}$** | 0.3 | **$2.69\times 10^{-2}$** | $8.18\times 10^{-2}$ |
| AT5G04090 | | Histidine-tRNA ligase | | | | | | | | | |
| AT5G04060 | | S-adenosyl-L-methionine-dependent methyltransferases superfamily protein | | | | | | | | | |
| AT5G04040 | *SDP1* | Patatin-like phospholipase family protein | | | | | | | | | |
| AT5G11780 | | SNF2 domain protein | | | | | | | | | |
| AT5G11800 | *KEA6* | K+ efflux antiporter 6 | | | | 0.4 | **$1.04\times 10^{-2}$** | **$3.94\times 10^{-2}$** | | | |
| AT5G16200 | | 50S ribosomal protein-like protein | -0.7 | **$5.49\times 10^{-3}$** | **$3.51\times 10^{-2}$** | -0.8 | **$6.18\times 10^{-3}$** | **$2.62\times 10^{-2}$** | -1.1 | **$1.58\times 10^{-6}$** | **$1.68\times 10^{-5}$** |
| AT5G16210 | | HEAT repeat-containing protein | | | | | | | | | |
| AT5G16660 | | Low-density receptor-like protein | | | | -0.4 | **$1.56\times 10^{-2}$** | **$5.42\times 10^{-2}$** | -0.3 | **$3.63\times 10^{-2}$** | $1.03\times 10^{-1}$ |
| AT5G16640 | | Pentatricopeptide repeat (PPR) superfamily protein | | | | | | | | | |
| AT5G20990 | *B73* | Molybdopterin biosynthesis CNX1 protein | | | | 0.3 | **$1.76\times 10^{-2}$** | **$5.96\times 10^{-2}$** | 0.3 | **$3.19\times 10^{-2}$** | $9.33\times 10^{-2}$ |
| AT5G21910 | | Transmembrane protein | | | | | | | | | |
| AT5G21920 | *YLMG2* | YGGT family protein | | | | -0.3 | **$1.56\times 10^{-2}$** | **$5.42\times 10^{-2}$** | | | |
| AT5G35590 | *PAA1* | Proteasome alpha subunit A1 | | | | -0.3 | **$1.53\times 10^{-2}$** | **$5.33\times 10^{-2}$** | | | |
| AT5G35790 | *G6PD1* | Glucose-6-phosphate dehydrogenase 1 | | | | | | | | | |
| AT5TE49950 | | ATHILA7 transposon family,LTR/Gypsy transposon super family | | | | | | | | | |
| AT5G40390 | *RS5, SIP1* | Raffinose synthase family protein | | | | | | | 0.4 | **$2.65\times 10^{-3}$** | **$1.21\times 10^{-2}$** |
| AT5TE66150 | | HELITRONY1D transposon family,RC/Helitron transposon super family | | | | | | | | | |
| AT5G52060 | *BAG1* | BCL-2-associated athanogene 1 | | | | | | | -0.4 | **$1.73\times 10^{-3}$** | **$8.39\times 10^{-3}$** |
| AT5G52730 | | Copper transport protein family | | | | | | | | | |
| AT5G52780 | *PAM68L* | Transmembrane protein, putative (DUF3464) | | | | -0.5 | **$1.65\times 10^{-3}$** | **$9.09\times 10^{-3}$** | | | |
| AT5G54980 | *CASPL2D1* | Uncharacterized protein family (UPF0497) | | | | | | | | | |
| AT5G54970 | | Hypothetical protein | | | | | | | | | |
| AT5G56680 | *EMB2755* | Embryo defective 2755 | | | | | | | 0.3 | **$8.61\times 10^{-3}$** | **$3.26\times 10^{-2}$** |
| AT5G56660 | *ILL2* | IAA-leucine resistant (ILR)-like 2 | | | | | | | | | |
| AT5G56700 | | FBD / Leucine Rich Repeat domains containing protein | | | | | | | | | |
| AT5G56680 | *EMB2755* | Embryo defective 2755 | | | | | | | 0.3 | **$8.61\times 10^{-3}$** | **$3.26\times 10^{-2}$** |
| AT5G56680 | *EMB2755* | Embryo defective 2755 | | | | | | | 0.3 | **$8.61\times 10^{-3}$** | **$3.26\times 10^{-2}$** |
| AT5G56660 | *ILL2* | IAA-leucine resistant (ILR)-like 2 | | | | | | | | | |
| AT5G56680 | *EMB2755* | Embryo defective 2755 | | | | | | | 0.3 | **$8.61\times 10^{-3}$** | **$3.26\times 10^{-2}$** |
| AT5G56660 | *ILL2* | IAA-leucine resistant (ILR)-like 2 | | | | | | | | | |
| AT5G56680 | *EMB2755* | Embryo defective 2755 | | | | | | | 0.3 | **$8.61\times 10^{-3}$** | **$3.26\times 10^{-2}$** |
| AT5G66470 | *ERA1* | GTP-binding protein Era-like protein | | | | | | | 0.3 | **$4.47\times 10^{-2}$** | $1.21\times 10^{-1}$ |
| ***HIS1-3*** **eGWAS** | | | | | | | | | | | |
| AT1G01340 | *ACBK1, CNGC10* | Cyclic nucleotide gated channel 10 | | | | | | | | | |
| AT1G11050 | | Protein kinase superfamily protein | | | | | | | | | |
| AT1G13220 | *CRWN2* | Crowded nuclei 2 | | | | 0.4 | **$1.56\times 10^{-2}$** | **$5.40\times 10^{-2}$** | | | |
| AT1G14220 | | Ribonuclease T2 family protein | 4.3 | **$6.69\times 10^{-9}$** | **$2.75\times 10^{-7}$** | 3.8 | **$3.15\times 10^{-11}$** | **$1.34\times 10^{-9}$** | 4.1 | **$3.00\times 10^{-12}$** | **$8.23\times 10^{-11}$** |

| | | | | | | | | | | | | | | | |
|---|---|---|---|---|---|---|---|---|---|---|---|---|---|---|---|
| AT1G14230 | | GDA1/CD39 nucleoside phosphatase family protein | | | | | | | | | | | | | |
| AT1G14240 | *APY3* | GDA1/CD39 nucleoside phosphatase family protein | 2.3 | **$1.16\times10^{-12}$** | **$1.11\times10^{-10}$** | 2.1 | **$5.43\times10^{-10}$** | **$1.79\times10^{-8}$** | 2.9 | **$6.79\times10^{-22}$** | **$5.55\times10^{-20}$** | 0.6 | **$3.14\times10^{-2}$** | $6.45\times10^{-1}$ | (+/+) |
| AT1G14690 | *MAP65-7* | Microtubule-associated protein 65-7 | | | | | | | | | | | | | |
| AT1G15215 | *DTF1* | DNA-binding transcription factor 1 | | | | -0.5 | **$2.21\times10^{-2}$** | $7.15\times10^{-2}$ | | | | | | | |
| AT1G16800 | | P-loop containing nucleoside triphosphate hydrolases superfamily protein | | | | 0.6 | **$1.48\times10^{-2}$** | **$5.19\times10^{-2}$** | | | | | | | |
| AT1G16840 | | Hypothetical protein | -0.9 | **$6.64\times10^{-7}$** | **$1.70\times10^{-5}$** | -0.9 | **$1.08\times10^{-9}$** | **$3.28\times10^{-8}$** | -0.6 | **$1.06\times10^{-7}$** | **$1.40\times10^{-6}$** | | | | |
| AT1G23880 | | NHL domain-containing protein | | | | | | | | | | | | | |
| AT1G23890 | | NHL domain-containing protein | | | | | | | -0.4 | **$3.71\times10^{-2}$** | $1.05\times10^{-1}$ | | | | |
| AT1G28420 | *HB1* | Homeobox-1 | | | | 0.3 | **$3.44\times10^{-2}$** | $1.00\times10^{-1}$ | 0.3 | **$2.70\times10^{-2}$** | $8.20\times10^{-2}$ | | | | |
| AT1G57800 | *VIM5* | Variant in methylation 5 | | | | | | | | | | | | | |
| AT1G69190 | *DHPS;HPPK* | Dihydropterin pyrophosphokinase / Dihydropteroate synthase | | | | | | | | | | | | | |
| AT2G17740 | *VLG* | Vacuoleless gametophytes | | | | | | | 1.8 | **$2.05\times10^{-3}$** | **$9.70\times10^{-3}$** | 1.8 | **$4.85\times10^{-2}$** | $7.78\times10^{-1}$ | |
| AT2G20740 | | Tetraspanin family protein | | | | | | | | | | | | | |
| AT2G24140 | *MRF7* | Myosin-J heavy chain-like protein | | | | | | | | | | | | | |
| AT2G24140 | *MRF7* | Myosin-J heavy chain-like protein | | | | | | | | | | | | | |
| AT2G24140 | *MRF7* | Myosin-J heavy chain-like protein | | | | | | | | | | | | | |
| AT2G24140 | *MRF7* | Myosin-J heavy chain-like protein | | | | | | | | | | | | | |
| AT2G24960 | | Myb/SANT-like DNA-binding domain protein | | | | 1.1 | **$1.71\times10^{-4}$** | **$1.40\times10^{-3}$** | | | | | | | |
| AT2G24960 | | Myb/SANT-like DNA-binding domain protein | | | | 1.1 | **$1.71\times10^{-4}$** | **$1.40\times10^{-3}$** | | | | | | | |
| AT2G25737 | | Sulfite exporter TauE/SafE family protein | | | | | | | | | | | | | |
| AT2G27720 | *P2Z, RPP2A* | Ribosomal protein | | | | -0.4 | **$2.54\times10^{-2}$** | $7.94\times10^{-2}$ | | | | | | | |
| AT2G38025 | *OTU3* | Cysteine proteinases superfamily protein | | | | | | | | | | | | | |
| AT2G38010 | *ATNCER2* | Neutral ceramidase 2 | -0.5 | **$1.15\times10^{-2}$** | $6.14\times10^{-2}$ | | | | | | | | | | |
| AT2G38020 | *VCL1* | MVB-to-vacuole trafficking protein | | | | | | | | | | | | | |
| AT2G38040 | *CAC3* | Acetyl co-enzyme a carboxylase carboxyltransferase alpha subunit | | | | | | | | | | | | | |
| AT2G38060 | *PHT4;2* | Phosphate transporter 4;2 | 1.0 | **$1.66\times10^{-2}$** | $8.09\times10^{-2}$ | | | | 1.0 | **$1.28\times10^{-3}$** | **$6.52\times10^{-3}$** | | | | |
| AT2G38610 | *KH9* | RNA-binding KH domain-containing protein | | | | | | | | | | | | | |
| AT2G38630 | *FBIP2* | D FBS interacting protein 2 | | | | | | | | | | | | | |
| AT2G38640 | | LURP-one-like protein (DUF567) | 0.4 | **$1.35\times10^{-2}$** | $6.93\times10^{-2}$ | 0.6 | **$5.29\times10^{-3}$** | **$2.32\times10^{-2}$** | 0.5 | **$1.08\times10^{-4}$** | **$7.56\times10^{-4}$** | | | | |
| AT2G40630 | *BTE1* | Uncharacterized conserved protein (UCP030365) | 0.5 | **$3.98\times10^{-2}$** | $1.52\times10^{-1}$ | 0.8 | **$1.43\times10^{-5}$** | **$1.68\times10^{-4}$** | 0.6 | **$1.55\times10^{-3}$** | **$7.63\times10^{-3}$** | | | | |
| AT2G42960 | | Protein kinase superfamily protein | | | | | | | | | | | | | |
| AT2G42975 | | Myosin-G heavy chain-like protein | -1.2 | **$3.73\times10^{-12}$** | **$3.21\times10^{-10}$** | -1.1 | **$1.98\times10^{-8}$** | **$4.77\times10^{-7}$** | -1.0 | **$3.04\times10^{-14}$** | **$1.07\times10^{-12}$** | | | | |
| AT2G43060 | *IBH1* | ILI1 binding BHLH 1 | | | | | | | | | | | | | |
| AT2G43070 | *SPPL3* | SIGNAL PEPTIDE PEPTIDASE-LIKE 3 | | | | | | | | | | | | | |

| | | | | | | | | | | | | | | |
|---|---|---|---|---|---|---|---|---|---|---|---|---|---|---|
| AT2G43980 | *ITPK4* | Inositol 1,3,4-trisphosphate 5/6-kinase 4 | | | | | | | | | | | | |
| AT2G44020 | | Mitochondrial transcription termination factor family protein | | | | | | | | | | | | |
| AT2G44735 | | Transmembrane protein | | | | | | | | | | | | |
| AT2G44710 | *HRLP* | RNA-binding (RRM/RBD/RNP motifs) family protein | | | | | | | | | | | | |
| AT2G44770 | *MCR* | Engulfment and cell motility (elmo) domain B | | | | | | | | | | | | |
| AT3G01310 | *VIP1* | Arabidopsis homolog of yeast VIP1 1 | | | | | | | 0.3 | **$7.66\times 10^{-3}$** | **$2.97\times 10^{-2}$** | 0.5 | **$1.54\times 10^{-2}$** | $4.61\times 10^{-1}$ |
| AT3G01260 | | Galactose mutarotase-like superfamily protein | 5.0 | **$1.14\times 10^{-58}$** | **$7.34\times 10^{-55}$** | 4.8 | **$3.19\times 10^{-80}$** | **$3.07\times 10^{-76}$** | 5.3 | **$5.11\times 10^{-80}$** | **$1.10\times 10^{-76}$** | | | |
| AT3G01300 | *PBL35* | PBS 1 like 35 | | | | | | | -0.4 | **$8.64\times 10^{-3}$** | **$3.27\times 10^{-2}$** | | | |
| AT3G01760 | *LHT6* | Lysine-histidine-like transporter 6 | | | | | | | | | | | | |
| AT3G01770 | *BET10, GTE11* | Bromodomain and extraterminal domain protein 10 | | | | | | | | | | | | |
| AT3G01780 | *TPLATE* | ARM repeat superfamily protein | | | | 0.4 | **$2.24\times 10^{-2}$** | $7.21\times 10^{-2}$ | | | | | | |
| AT3G02050 | *KUP3* | Potassium transporter KUP3p (KUP3) | | | | | | | -0.3 | **$5.09\times 10^{-2}$** | $1.34\times 10^{-1}$ | | | |
| AT3G02060 | | DEAD/DEAH box helicase | 0.5 | **$2.03\times 10^{-3}$** | **$1.59\times 10^{-2}$** | 0.6 | **$8.72\times 10^{-7}$** | **$1.44\times 10^{-5}$** | 0.5 | **$4.42\times 10^{-4}$** | **$2.61\times 10^{-3}$** | | | |
| AT3G02520 | *GRF7* | General regulatory factor 7 | | | | | | | | | | | | |
| AT3G02500 | | Mental retardation GTPase activating protein | | | | | | | | | | | | |
| AT3G02510 | | Regulator of chromosome condensation (RCC1) family protein | | | | | | | | | | | | |
| AT3G02530 | *CCT6-2* | TCP-1/cpn60 chaperonin family protein | | | | | | | | | | | | |
| AT3G02910 | | AIG2-like (avirulence induced gene) family protein | | | | | | | | | | | | |
| AT3G03300 | *DCL2* | Dicer like 2 | | | | | | | | | | | | |
| AT3G10130 | *SOUL4* | SOUL heme-binding family protein | -0.6 | **$8.92\times 10^{-4}$** | **$8.26\times 10^{-3}$** | -0.6 | **$5.37\times 10^{-4}$** | **$3.61\times 10^{-3}$** | -0.4 | **$1.31\times 10^{-4}$** | **$9.01\times 10^{-4}$** | | | |
| AT3G10845 | | RNA-binding (RRM/RBD/RNP motifs) family protein | | | | | | | | | | | | |
| AT3G10820 | *MED26A* | Transcription elongation factor (TFIIS) family protein | | | | | | | | | | | | |
| AT3G10850 | *GLX2-2* | Metallo-hydrolase/oxidoreductase superfamily protein | -0.4 | **$4.83\times 10^{-2}$** | $1.74\times 10^{-1}$ | -0.5 | **$1.40\times 10^{-4}$** | **$1.19\times 10^{-3}$** | | | | | | |
| AT3G11050 | *FER2* | Ferritin 2 | | | | | | | | | | | | |
| AT3G17860 | *JAI3* | Jasmonate-zim-domain protein 3 | -0.8 | **$1.43\times 10^{-3}$** | **$1.20\times 10^{-2}$** | -0.7 | **$1.07\times 10^{-2}$** | **$4.04\times 10^{-2}$** | -0.5 | **$9.17\times 10^{-4}$** | **$4.90\times 10^{-3}$** | | | |
| AT3G17880 | *ATHIP2* | Tetraticopeptide domain-containing thioredoxin | | | | | | | | | | | | |
| AT3G17880 | *ATHIP2* | Tetraticopeptide domain-containing thioredoxin | | | | | | | | | | | | |
| AT3G17860 | *JAI3* | Jasmonate-zim-domain protein 3 | -0.8 | **$1.43\times 10^{-3}$** | **$1.20\times 10^{-2}$** | -0.7 | **$1.07\times 10^{-2}$** | **$4.04\times 10^{-2}$** | -0.5 | **$9.17\times 10^{-4}$** | **$4.90\times 10^{-3}$** | | | |
| AT3G19340 | | Aminopeptidase (DUF3754) | -0.5 | **$1.72\times 10^{-3}$** | **$1.39\times 10^{-2}$** | -0.5 | **$2.61\times 10^{-5}$** | **$2.80\times 10^{-4}$** | -0.3 | **$4.16\times 10^{-3}$** | **$1.77\times 10^{-2}$** | | | |
| AT3G23280 | *XBAT35* | XB3 ortholog 5 in *Arabidopsis thaliana* | | | | | | | | | | 0.4 | **$2.96\times 10^{-2}$** | $6.27\times 10^{-1}$ |
| AT3G24508 | | Encodes a defensin-like (DEFL) family protein | | | | | | | | | | | | |
| AT3G24715 | *HCR1* | Kinase superfamily with octicosapeptide/Phox/Bem1p domain-containing protein | | | | 1.1 | **$6.64\times 10^{-5}$** | **$6.31\times 10^{-4}$** | 0.5 | **$3.23\times 10^{-2}$** | $9.43\times 10^{-2}$ | | | |
| AT3G25100 | *CDC45* | Cell division cycle 45 | | | | | | | | | | | | |
| AT3G25110 | *FATA1* | Fata acyl-acp thioesterase | | | | | | | | | | | | |

| | | | | | | | | | | | | | | |
|---|---|---|---|---|---|---|---|---|---|---|---|---|---|---|
| AT3G25100 | *CDC45* | Cell division cycle 45 | | | | | | | | | | | | |
| AT3G25110 | *FATA1* | Fata acyl-acp thioesterase | | | | | | | | | | | | |
| AT3G26490 | *NPY7* | Phototropic-responsive NPH3 family protein | 0.6 | **$4.68\times 10^{-2}$** | $1.71\times 10^{-1}$ | 0.5 | **$4.12\times 10^{-2}$** | $1.15\times 10^{-1}$ | 0.5 | **$3.26\times 10^{-2}$** | $9.48\times 10^{-2}$ | | | |
| AT3TE40445 | | ATLINEIII transposon family,LINE/L1 transposon super family | | | | | | | | | | | | |
| AT3G28415 | *ABCB22* | ABC transporter family protein | | | | | | | | | | | | |
| AT3G28480 | | Oxoglutarate/iron-dependent oxygenase | | | | -0.4 | **$3.22\times 10^{-2}$** | $9.53\times 10^{-2}$ | -0.7 | **$2.79\times 10^{-5}$** | **$2.25\times 10^{-4}$** | -0.3 | **$4.82\times 10^{-2}$** | $7.76\times 10^{-1}$ |
| AT3G56050 | | Protein kinase family protein | | | | | | | | | | | | |
| AT3G61720 | *MCTP12* | Multiple c2 domain and transmembrane region protein 12 | | | | | | | | | | | | |
| AT3G61700 | | Helicase with zinc finger protein | | | | | | | | | | | | |
| AT3G61710 | *ATG6* | Autophagy protein 6 | | | | | | | | | | | | |
| AT3G61723 | | PHD finger protein | | | | | | | | | | | | |
| AT3G61730 | *RMF1* | Reduced male fertility | | | | | | | | | | | | |
| AT3G61740 | *ATX3* | Arabidopsis trithorax 3 | | | | 0.4 | **$4.18\times 10^{-2}$** | $1.16\times 10^{-1}$ | | | | | | |
| AT4G03430 | *EMB2770* | Embryo defective 2770 | | | | 0.3 | **$4.13\times 10^{-2}$** | $1.15\times 10^{-1}$ | | | | | | |
| AT4G03420 | | Hypothetical protein (DUF789) | | | | 0.4 | **$3.15\times 10^{-2}$** | $9.36\times 10^{-2}$ | 1.0 | **$1.70\times 10^{-14}$** | **$6.11\times 10^{-13}$** | 0.7 | **$1.20\times 10^{-5}$** | **$3.42\times 10^{-3}$** |
| AT4G04745 | | Hypothetical protein | | | | | | | | | | | | |
| AT4G04740 | *CPK23* | Calcium-dependent protein kinase 23 | | | | 0.8 | **$2.01\times 10^{-5}$** | **$2.23\times 10^{-4}$** | 0.4 | **$1.11\times 10^{-2}$** | **$4.00\times 10^{-2}$** | | | |
| AT4G05390 | *RFNR1* | Root FNR1 | -0.9 | **$3.57\times 10^{-4}$** | **$3.97\times 10^{-3}$** | -1.1 | **$1.47\times 10^{-8}$** | **$3.65\times 10^{-7}$** | -1.0 | **$2.89\times 10^{-9}$** | **$5.11\times 10^{-8}$** | | | |
| AT4G05420 | *DDB1A* | Damaged dna binding protein 1a | | | | 0.4 | **$1.47\times 10^{-2}$** | **$5.18\times 10^{-2}$** | | | | | | |
| AT4G08920 | *CRY1* | Cryptochrome 1 | | | | | | | | | | | | |
| AT4G09640 | *ENOR3L6* | Magnesium transporter, putative (DUF803) | | | | | | | | | | | | |
| AT4G09630 | | Transmembrane protein (DUF616) | | | | 0.4 | **$1.76\times 10^{-3}$** | **$9.53\times 10^{-3}$** | | | | | | |
| AT4G11040 | *DOG18* | Delay of germination 18 | | | | | | | | | | | | |
| AT4G11030 | *LACS5* | AMP-dependent synthetase and ligase family protein | | | | | | | | | | | | |
| AT4G12100 | | Cullin family protein | | | | | | | | | | | | |
| AT4G12750 | | Homeodomain-like transcriptional regulator | 0.6 | **$1.98\times 10^{-2}$** | $9.21\times 10^{-2}$ | 0.7 | **$4.95\times 10^{-4}$** | **$3.38\times 10^{-3}$** | 0.7 | **$1.37\times 10^{-4}$** | **$9.35\times 10^{-4}$** | | | |
| AT4G15620 | *CASPL1E2* | Uncharacterized protein family (UPF0497) | -1.2 | **$2.47\times 10^{-8}$** | **$8.98\times 10^{-7}$** | -1.6 | **$9.31\times 10^{-13}$** | **$5.13\times 10^{-11}$** | -1.4 | **$3.00\times 10^{-48}$** | **$1.14\times 10^{-45}$** | | | |
| AT4G15620 | *CASPL1E2* | Uncharacterized protein family (UPF0497) | -1.2 | **$2.47\times 10^{-8}$** | **$8.98\times 10^{-7}$** | -1.6 | **$9.31\times 10^{-13}$** | **$5.13\times 10^{-11}$** | -1.4 | **$3.00\times 10^{-48}$** | **$1.14\times 10^{-45}$** | | | |
| AT4G18360 | *GOX3* | Glycolate oxidase 3 | -0.7 | **$1.64\times 10^{-2}$** | **$7.99\times 10^{-2}$** | -1.0 | **$1.58\times 10^{-3}$** | **$8.78\times 10^{-3}$** | -0.6 | **$2.48\times 10^{-2}$** | $7.65\times 10^{-2}$ | | | |
| AT4G18372 | | Small nuclear ribonucleoprotein family protein | | | | | | | | | | | | |
| AT4G26450 | | Hypothetical protein | | | | | | | | | | | | |
| AT5G02710 | | Zinc/iron-chelating domain protein | | | | | | | | | | | | |
| AT5G08550 | *ILP1* | Increased level of polyploidy1-1d | | | | 0.4 | **$2.32\times 10^{-2}$** | $7.42\times 10^{-2}$ | | | | | | |
| AT5G08560 | *WDR26* | Transducin family protein / WD-40 repeat family protein | | | | | | | | | | | | |

| | | | | | | | | | | | |
|---|---|---|---|---|---|---|---|---|---|---|---|
| AT5G08560 | *WDR26* | Transducin family protein / WD-40 repeat family protein | | | | | | | | | |
| AT5G08650 | *CPLEPA* | Small GTP-binding protein | | | | | | | | | |
| AT5G08660 | *PSI3* | D-lactate dehydrogenase (DUF668) | | | | | | | | | |
| AT5G08680 | | ATP synthase alpha/beta family protein | | | | | | | | | |
| AT5G08670 | | ATP synthase alpha/beta family protein | | | | | | | | | |
| AT5G08680 | | ATP synthase alpha/beta family protein | | | | | | | | | |
| AT5G08670 | | ATP synthase alpha/beta family protein | | | | | | | | | |
| AT5G08680 | | ATP synthase alpha/beta family protein | | | | | | | | | |
| AT5G08670 | | ATP synthase alpha/beta family protein | | | | | | | | | |
| AT5G08680 | | ATP synthase alpha/beta family protein | | | | | | | | | |
| AT5G08660 | *PSI3* | D-lactate dehydrogenase (DUF668) | | | | | | | | | |
| AT5G08690 | | ATP synthase alpha/beta family protein | | | | | | | 0.2 | **$3.26\times10^{-2}$** | $9.49\times10^{-2}$ |
| AT5G08680 | | ATP synthase alpha/beta family protein | | | | | | | | | |
| AT5G08660 | *PSI3* | D-lactate dehydrogenase (DUF668) | | | | | | | | | |
| AT5G08690 | | ATP synthase alpha/beta family protein | | | | | | | 0.2 | **$3.26\times10^{-2}$** | $9.49\times10^{-2}$ |
| AT5G11210 | *GLR2.5* | Glutamate receptor 2.5 | | | | | | | | | |
| AT5G11200 | *UAP56B* | DEAD/DEAH box RNA helicase family protein | 0.4 | **$1.09\times10^{-2}$** | **$5.93\times10^{-2}$** | 0.4 | **$1.07\times10^{-3}$** | **$6.41\times10^{-3}$** | 0.3 | **$9.79\times10^{-3}$** | **$3.61\times10^{-2}$** |
| AT5G13010 | *EMB3011* | RNA helicase family protein | | | | | | | | | |
| AT5G13000 | *GSL12* | Glucan synthase-like 12 | | | | | | | | | |
| AT5G19030 | | Cloroplast RRM-containing protein.RNA-binding (RRM/RBD/RNP motifs) family protein | | | | -0.6 | **$5.85\times10^{-3}$** | **$2.51\times10^{-2}$** | | | |
| AT5G20120 | | Testis- and ovary-specific PAZ domain protein | | | | | | | | | |
| AT5G20130 | | Sulfate adenylyltransferase subunit | | | | | | | | | |
| AT5G26030 | *FC1* | Ferrochelatase 1 | | | | | | | -0.3 | **$3.33\times10^{-2}$** | $9.64\times10^{-2}$ |
| AT5G26000 | *BGLU38, TGG1* | Thioglucoside glucohydrolase 1 | | | | | | | -0.3 | **$5.95\times10^{-2}$** | $1.51\times10^{-1}$ |
| ***DREB1A* eGWAS** | | | | | | | | | | | |
| AT1G12900 | *GAPA-2* | Glyceraldehyde 3-phosphate dehydrogenase A subunit 2 | -0.5 | **$1.13\times10^{-2}$** | $6.06\times10^{-2}$ | | | | -0.5 | **$4.58\times10^{-5}$** | **$3.52\times10^{-4}$** |
| AT1G12870 | | F-box and associated interaction domains-containing protein | | | | | | | | | |
| AT1G12920 | *ERF1-2* | Eukaryotic release factor 1-2 | 0.4 | **$1.74\times10^{-2}$** | $8.39\times10^{-2}$ | 0.3 | **$1.41\times10^{-2}$** | **$5.03\times10^{-2}$** | 0.4 | **$2.53\times10^{-4}$** | **$1.61\times10^{-3}$** |
| AT1G12920 | *ERF1-2* | Eukaryotic release factor 1-2 | 0.4 | **$1.74\times10^{-2}$** | $8.39\times10^{-2}$ | 0.3 | **$1.41\times10^{-2}$** | **$5.03\times10^{-2}$** | 0.4 | **$2.53\times10^{-4}$** | **$1.61\times10^{-3}$** |
| AT1G14790 | *RDR1* | Rna-dependent rna polymerase 1 | | | | | | | | | |
| AT1G14780 | *MACP1* | Membrane attack complex/perforin-like 1 | | | | | | | 0.7 | **$3.85\times10^{-2}$** | $1.08\times10^{-1}$ |
| AT1G14800 | | Nucleic acid-binding, OB-fold-like protein | | | | | | | | | |
| AT1G14810 | | Encodes an aspartate semialdehyde dehydrogenase | | | | | | | 0.3 | **$3.67\times10^{-2}$** | $1.04\times10^{-1}$ |

| | | | | | | | | | | | |
|---|---|---|---|---|---|---|---|---|---|---|---|
| AT1G14830 | *ADL5* | Arabidopsis dynamin-like protein 5 | | | | | | | | | |
| AT1G14790 | *RDR1* | Rna-dependent rna polymerase 1 | | | | | | | | | |
| AT1G14800 | | Nucleic acid-binding, OB-fold-like protein | | | | | | | | | |
| AT1G14810 | | Encodes an aspartate semialdehyde dehydrogenase | | | | | | | 0.3 | **$3.67\times 10^{-2}$** | $1.04\times 10^{-1}$ |
| AT1G14820 | | Sec14p-like phosphatidylinositol transfer family protein | | | | | | | | | |
| AT1G14830 | *ADL5* | Arabidopsis dynamin-like protein 5 | | | | | | | | | |
| AT1G14840 | *MAP70-4* | Microtubule-associated proteins 70-4 | | | | | | | -0.4 | **$9.78\times 10^{-4}$** | **$5.16\times 10^{-3}$** |
| AT1G14800 | | Nucleic acid-binding, OB-fold-like protein | | | | | | | | | |
| AT1G14790 | *RDR1* | Rna-dependent rna polymerase 1 | | | | | | | | | |
| AT1G14810 | | Encodes an aspartate semialdehyde dehydrogenase | | | | | | | 0.3 | **$3.67\times 10^{-2}$** | $1.04\times 10^{-1}$ |
| AT1G14820 | | Sec14p-like phosphatidylinositol transfer family protein | | | | | | | | | |
| AT1G14830 | *ADL5* | Arabidopsis dynamin-like protein 5 | | | | | | | | | |
| AT1G14840 | *MAP70-4* | Microtubule-associated proteins 70-4 | | | | | | | -0.4 | **$9.78\times 10^{-4}$** | **$5.16\times 10^{-3}$** |
| AT1G14850 | *NUP155* | Nucleoporin 155 | 0.5 | **$1.14\times 10^{-3}$** | **$1.00\times 10^{-2}$** | 0.6 | **$1.26\times 10^{-4}$** | **$1.09\times 10^{-3}$** | 0.4 | **$2.09\times 10^{-3}$** | **$9.85\times 10^{-3}$** |
| AT1G14810 | | Encodes an aspartate semialdehyde dehydrogenase | | | | | | | 0.3 | **$3.67\times 10^{-2}$** | $1.04\times 10^{-1}$ |
| AT1G14800 | | Nucleic acid-binding, OB-fold-like protein | | | | | | | | | |
| AT1G14790 | *RDR1* | Rna-dependent rna polymerase 1 | | | | | | | | | |
| AT1G14820 | | Sec14p-like phosphatidylinositol transfer family protein | | | | | | | | | |
| AT1G14830 | *ADL5* | Arabidopsis dynamin-like protein 5 | | | | | | | | | |
| AT1G14840 | *MAP70-4* | Microtubule-associated proteins 70-4 | | | | | | | -0.4 | **$9.78\times 10^{-4}$** | **$5.16\times 10^{-3}$** |
| AT1G14850 | *NUP155* | Nucleoporin 155 | 0.5 | **$1.14\times 10^{-3}$** | **$1.00\times 10^{-2}$** | 0.6 | **$1.26\times 10^{-4}$** | **$1.09\times 10^{-3}$** | 0.4 | **$2.09\times 10^{-3}$** | **$9.85\times 10^{-3}$** |
| AT1G14810 | | Semialdehyde dehydrogenase family protein | | | | | | | 0.3 | **$3.67\times 10^{-2}$** | $1.04\times 10^{-1}$ |
| AT1G14790 | *RDR1* | Rna-dependent rna polymerase 1 | | | | | | | | | |
| AT1G14800 | | Nucleic acid-binding, OB-fold-like protein | | | | | | | | | |
| AT1G14820 | | Sec14p-like phosphatidylinositol transfer family protein | | | | | | | | | |
| AT1G14830 | *ADL5* | Arabidopsis dynamin-like protein 5 | | | | | | | | | |
| AT1G14840 | *MAP70-4* | Microtubule-associated proteins 70-4 | | | | | | | -0.4 | **$9.78\times 10^{-4}$** | **$5.16\times 10^{-3}$** |
| AT1G14850 | *NUP155* | Nucleoporin 155 | 0.5 | **$1.14\times 10^{-3}$** | **$1.00\times 10^{-2}$** | 0.6 | **$1.26\times 10^{-4}$** | **$1.09\times 10^{-3}$** | 0.4 | **$2.09\times 10^{-3}$** | **$9.85\times 10^{-3}$** |
| AT1G14810 | | Encodes an aspartate semialdehyde dehydrogenase | | | | | | | 0.3 | **$3.67\times 10^{-2}$** | $1.04\times 10^{-1}$ |
| AT1G16370 | *(OCT6)* | Organic cation/carnitine transporter 6 | 2.3 | **$1.79\times 10^{-8}$** | **$6.69\times 10^{-7}$** | 2.3 | **$1.96\times 10^{-8}$** | **$4.74\times 10^{-7}$** | 2.1 | **$1.29\times 10^{-10}$** | **$2.84\times 10^{-9}$** |
| AT1G16340 | *ATKDSA2* | Aldolase superfamily protein | | | | -0.4 | **$4.28\times 10^{-2}$** | $1.18\times 10^{-1}$ | | | |
| AT1G16360 | | LEM3 (ligand-effect modulator 3) family protein / CDC50 family protein | | | | | | | | | |
| AT1G16380 | *CAX1* | Cation exchanger 1 | | | | | | | | | |
| AT1G16810 | | 7-dehydrocholesterol reductase-like protein | | | | -0.3 | **$5.19\times 10^{-3}$** | **$2.28\times 10^{-2}$** | | | |

| | | | | | | | | | | | | | | | |
|---|---|---|---|---|---|---|---|---|---|---|---|---|---|---|---|
| AT1G16820 | | Vacuolar ATP synthase catalytic subunit-related / vacuolar proton pump-like protein | | | | | | | | | | | | | |
| AT1G16840 | | Hypothetical protein | -0.9 | $6.64 \times 10^{-7}$ | $1.70 \times 10^{-5}$ | -0.9 | $1.08 \times 10^{-9}$ | $3.28 \times 10^{-8}$ | -0.6 | $1.06 \times 10^{-7}$ | $1.40 \times 10^{-6}$ | | | | |
| AT1G16840 | | Hypothetical protein | -0.9 | $6.64 \times 10^{-7}$ | $1.70 \times 10^{-5}$ | -0.9 | $1.08 \times 10^{-9}$ | $3.28 \times 10^{-8}$ | -0.6 | $1.06 \times 10^{-7}$ | $1.40 \times 10^{-6}$ | | | | |
| AT1G59620 | *CW9* | Encodes CW9 | | | | | | | | | | | | | |
| AT1G62850 | | Class I peptide chain release factor | -0.4 | $2.14 \times 10^{-2}$ | $9.76 \times 10^{-2}$ | -0.5 | $1.01 \times 10^{-3}$ | $6.12 \times 10^{-3}$ | -0.6 | $1.89 \times 10^{-3}$ | $9.06 \times 10^{-3}$ | | | | |
| AT1G62860 | | Pseudogene of pentatricopeptide (PPR) repeat-containing protein | | | | | | | | | | | | | |
| AT1G66960 | *LUP5* | Terpenoid cyclases family protein | | | | | | | | | | | | | |
| AT1G67000 | | Protein kinase superfamily protein | | | | | | | | | | | | | |
| AT2G39130 | | Transmembrane amino acid transporter family protein | 0.6 | $2.47 \times 10^{-3}$ | $1.86 \times 10^{-2}$ | 0.6 | $4.13 \times 10^{-3}$ | $1.90 \times 10^{-2}$ | 0.6 | $1.38 \times 10^{-5}$ | $1.20 \times 10^{-4}$ | | | | |
| AT2G40320 | *TBL33* | Trichome birefringence-like 33 | 0.6 | $4.82 \times 10^{-3}$ | $3.17 \times 10^{-2}$ | 0.7 | $3.47 \times 10^{-4}$ | $2.51 \times 10^{-3}$ | 0.6 | $8.13 \times 10^{-4}$ | $4.42 \times 10^{-3}$ | | | | |
| AT3G05500 | *LDAP3* | Ld-associated protein 3 | -0.7 | $1.26 \times 10^{-3}$ | $1.09 \times 10^{-2}$ | -0.7 | $8.91 \times 10^{-5}$ | $8.08 \times 10^{-4}$ | -0.8 | $3.52 \times 10^{-10}$ | $7.30 \times 10^{-9}$ | | | | |
| AT3G06880 | | Transducin/WD40 repeat-like superfamily protein | | | | | | | | | | | | | |
| AT3G12510 | | MADS-box family protein | | | | | | | | | | | | | |
| AT3G12490 | *CYS6* | Arabidopsis thaliana phytocystatin 6 | -0.5 | $2.15 \times 10^{-2}$ | $9.80 \times 10^{-2}$ | -0.6 | $3.23 \times 10^{-3}$ | $1.55 \times 10^{-2}$ | | | | | | | |
| AT3G12500 | *CHI-B, PR3* | Basic chitinase | -2.2 | $6.74 \times 10^{-9}$ | $2.76 \times 10^{-7}$ | -2.5 | $3.10 \times 10^{-10}$ | $1.07 \times 10^{-8}$ | -1.9 | $9.68 \times 10^{-10}$ | $1.87 \times 10^{-8}$ | | | | |
| AT3G12520 | *SULTR4;2* | Sulfate transporter 4;2 | -0.9 | $4.46 \times 10^{-4}$ | $4.74 \times 10^{-3}$ | -0.6 | $8.16 \times 10^{-3}$ | $3.26 \times 10^{-2}$ | -0.7 | $2.86 \times 10^{-4}$ | $1.78 \times 10^{-3}$ | | | | |
| AT3G23620 | *ARPF2* | Arabidopsis homolog of yeast rpf2 | 0.8 | $5.83 \times 10^{-5}$ | $8.47 \times 10^{-4}$ | 0.5 | $4.66 \times 10^{-3}$ | $2.08 \times 10^{-2}$ | 0.4 | $2.15 \times 10^{-3}$ | $1.01 \times 10^{-2}$ | | | | |
| AT3G23610 | *DSPTP1* | Dual specificity protein phosphatase 1 | | | | | | | | | | | | | |
| AT3G46990 | *OPL3* | DUF740 family protein, putative (DUF740) | | | | | | | | | | | | | |
| AT3G47000 | | Glycosyl hydrolase family protein | | | | | | | -0.3 | $8.76 \times 10^{-3}$ | $3.30 \times 10^{-2}$ | | | | |
| AT3G48010 | *CNGC16* | Cyclic nucleotide-gated channel 16 | | | | | | | | | | | | | |
| AT3G48030 | *ATL48* | Arabidopsis toxicos en levadura 48 | | | | -0.8 | $1.12 \times 10^{-8}$ | $2.82 \times 10^{-7}$ | -0.3 | $1.57 \times 10^{-2}$ | $5.32 \times 10^{-2}$ | | | | |
| AT3G48020 | | Hypothetical protein | -1.1 | $1.71 \times 10^{-3}$ | $1.38 \times 10^{-2}$ | -1.2 | $1.34 \times 10^{-3}$ | $7.68 \times 10^{-3}$ | | | | 0.9 | $2.01 \times 10^{-4}$ | $2.92 \times 10^{-2}$ | (-/+) |
| AT3G48030 | *ATL48* | Arabidopsis toxicos en levadura 48 | | | | -0.8 | $1.12 \times 10^{-8}$ | $2.82 \times 10^{-7}$ | -0.3 | $1.57 \times 10^{-2}$ | $5.32 \times 10^{-2}$ | | | | |
| AT3G48030 | *ATL48* | Arabidopsis toxicos en levadura 48 | | | | -0.8 | $1.12 \times 10^{-8}$ | $2.82 \times 10^{-7}$ | -0.3 | $1.57 \times 10^{-2}$ | $5.32 \times 10^{-2}$ | | | | |
| AT3G48020 | | Hypothetical protein | -1.1 | $1.71 \times 10^{-3}$ | $1.38 \times 10^{-2}$ | -1.2 | $1.34 \times 10^{-3}$ | $7.68 \times 10^{-3}$ | | | | 0.9 | $2.01 \times 10^{-4}$ | $2.92 \times 10^{-2}$ | (-/+) |
| AT3G48050 | *SUO* | Shuttle' in chinese | | | | | | | | | | | | | |
| AT3G48430 | *JMJ12* | Jumonji domain-containing protein 12 | | | | 0.3 | $2.54 \times 10^{-2}$ | $7.92 \times 10^{-2}$ | | | | | | | |
| AT3G48425 | *APE1L* | DNAse I-like superfamily protein | | | | | | | | | | | | | |
| AT3G48440 | *ATC3H43* | Zinc finger C-x8-C-x5-C-x3-H type family protein | -0.4 | $4.10 \times 10^{-2}$ | $1.55 \times 10^{-1}$ | -0.4 | $2.75 \times 10^{-2}$ | $8.42 \times 10^{-2}$ | -0.5 | $1.34 \times 10^{-2}$ | $4.69 \times 10^{-2}$ | | | | |
| AT3G50110 | *PTEN3* | *Arabidopsis thaliana* phosphatase and tensin homolog deleted on chromosome ten 3 | | | | | | | | | | | | | |
| AT3G50940 | | P-loop containing nucleoside triphosphate hydrolases superfamily protein | | | | | | | | | | | | | |
| AT3G61600 | *POB1* | Poz/btb containin g-protein 1 | | | | | | | | | | | | | |
| AT3G61590 | *HWS* | Hawaiian skirt | | | | | | | | | | | | | |

| | | | | | | | | | | | | | | | |
|---|---|---|---|---|---|---|---|---|---|---|---|---|---|---|---|
| AT3G61610 | | Galactose mutarotase-like superfamily protein | | | | | | | | | | | | | |
| AT3G61600 | *POB1* | Poz/btb containin g-protein 1 | | | | | | | | | | | | | |
| AT3G61620 | *RRP41* | Exonuclease RRP41 | | | | 0.4 | **$1.29\times 10^{-2}$** | **$4.68\times 10^{-2}$** | | | | | | | |
| AT4G00740 | *QUA3* | Quasimodo 3 | | | | | | | | | | | | | |
| AT4G16340 | *SPK1* | Spike 1 | | | | 0.4 | **$3.29\times 10^{-3}$** | **$1.57\times 10^{-2}$** | | | | | | | |
| AT4G16310 | *LDL3* | LSD-1 like 3 | | | | 0.3 | **$4.55\times 10^{-2}$** | $1.24\times 10^{-1}$ | | | | | | | |
| AT4G19030 | *NLM1* | Nod26-like major intrinsic protein 1 | 2.8 | **$3.77\times 10^{-7}$** | **$1.02\times 10^{-5}$** | 2.2 | **$5.70\times 10^{-6}$** | **$7.51\times 10^{-5}$** | 2.6 | **$7.47\times 10^{-17}$** | **$3.58\times 10^{-15}$** | | | | |
| AT4G21170 | | Tetratricopeptide repeat (TPR)-like superfamily protein | | | | | | | | | | | | | |
| AT4G21210 | *RP1* | PPDK regulatory protein | | | | | | | | | | | | | |
| AT4G21200 | *GA2OX8* | *Arabidopsis thaliana* gibberellin 2-oxidase 8 | | | | | | | | | | | | | |
| AT4G21300 | *ATDEK36* | Tetratricopeptide repeat (TPR)-like superfamily protein | | | | | | | | | | | | | |
| AT4G21310 | *DEAL2* | DUF1218 family member | | | | | | | -0.8 | **$1.87\times 10^{-2}$** | $6.11\times 10^{-2}$ | | | | |
| AT4G21320 | *HSA32* | Heat-stress-associated 32 | | | | -0.9 | **$1.08\times 10^{-3}$** | **$6.44\times 10^{-3}$** | | | | | | | |
| AT4G28080 | *REC2* | Reduced chloroplast coverage 2 | | | | | | | | | | | | | |
| AT4G33320 | *CHIQL2* | DUF641 domain protein, probable psuedogene | | | | | | | | | | | | | |
| AT4G34230 | *CAD5* | Cinnamyl alcohol dehydrogenase 5 | | | | -0.4 | **$4.53\times 10^{-2}$** | $1.23\times 10^{-1}$ | | | | | | | |
| AT5G01950 | | Leucine-rich repeat protein kinase family protein | -0.5 | **$4.69\times 10^{-2}$** | $1.71\times 10^{-1}$ | | | | -0.6 | **$2.69\times 10^{-4}$** | **$1.69\times 10^{-3}$** | | | | |
| AT5G12900 | | DNA double-strand break repair RAD50 ATPase | 0.5 | **$6.60\times 10^{-3}$** | **$4.04\times 10^{-2}$** | 0.4 | **$2.71\times 10^{-2}$** | $8.35\times 10^{-2}$ | | | | | | | |
| AT5G12920 | | Transducin/WD40 repeat-like superfamily protein | | | | | | | | | | | | | |
| AT5G12930 | | Inactive rhomboid protein | -1.2 | **$7.83\times 10^{-6}$** | **$1.49\times 10^{-4}$** | -0.8 | **$1.42\times 10^{-2}$** | **$5.05\times 10^{-2}$** | -0.7 | **$1.45\times 10^{-5}$** | **$1.26\times 10^{-4}$** | | | | |
| AT5G18560 | *PUCHI* | Encodes PUCHI, a member of the ERF subfamily B-1 | | | | | | | | | | | | | |
| AT5G18550 | *ATC3H58* | Zinc finger C-x8-C-x5-C-x3-H type family protein | -0.4 | **$2.07\times 10^{-2}$** | $9.51\times 10^{-2}$ | | | | -0.4 | **$1.10\times 10^{-2}$** | **$3.97\times 10^{-2}$** | | | | |
| AT5G19200 | *TSC10B* | NAD(P)-binding Rossmann-fold superfamily protein | | | | | | | | | | | | | |
| AT5G19190 | | Hypothetical protein | 1.1 | **$6.14\times 10^{-6}$** | **$1.22\times 10^{-4}$** | 1.0 | **$2.19\times 10^{-8}$** | **$5.22\times 10^{-7}$** | 0.6 | **$7.29\times 10^{-5}$** | **$5.34\times 10^{-4}$** | -0.5 | **$2.80\times 10^{-2}$** | $6.14\times 10^{-1}$ | (+/-) |
| AT5G19210 | | P-loop containing nucleoside triphosphate hydrolases superfamily protein | | | | | | | | | | | | | |
| AT5G19220 | *ADG2* | ADP glucose pyrophosphorylase 2 | | | | 0.5 | **$4.13\times 10^{-3}$** | **$1.89\times 10^{-2}$** | | | | | | | |
| AT5G23590 | | DNAJ heat shock N-terminal domain-containing protein | | | | | | | | | | | | | |
| AT5G23575 | | Transmembrane CLPTM1 family protein | 0.6 | **$1.60\times 10^{-3}$** | **$1.31\times 10^{-2}$** | 0.6 | **$2.65\times 10^{-5}$** | **$2.83\times 10^{-4}$** | 1.0 | **$2.06\times 10^{-21}$** | **$1.63\times 10^{-19}$** | 0.5 | **$1.51\times 10^{-2}$** | $4.58\times 10^{-1}$ | (+/+) |
| AT5G23610 | | DYAD protein | | | | | | | | | | | | | |
| AT5G23580 | *CPK12* | *Arabidopsis thaliana* calmodulin-like domain protein kinase 9 | | | | | | | | | | | | | |
| AT5G23630 | *MIA* | Male gametogenesis impaired anthers | 0.3 | **$4.36\times 10^{-2}$** | $1.62\times 10^{-1}$ | 0.5 | **$1.85\times 10^{-5}$** | **$2.08\times 10^{-4}$** | 0.3 | **$4.40\times 10^{-2}$** | $1.20\times 10^{-1}$ | | | | |
| AT5G23630 | *MIA* | Male gametogenesis impaired anthers | 0.3 | **$4.36\times 10^{-2}$** | $1.62\times 10^{-1}$ | 0.5 | **$1.85\times 10^{-5}$** | **$2.08\times 10^{-4}$** | 0.3 | **$4.40\times 10^{-2}$** | $1.20\times 10^{-1}$ | | | | |
| AT5G23610 | | DYAD protein | | | | | | | | | | | | | |
| AT5G23580 | *CPK12* | *Arabidopsis thaliana* calmodulin-like domain protein kinase 9 | | | | | | | | | | | | | |

| | | | | | | | | | | | |
|---|---|---|---|---|---|---|---|---|---|---|---|
| AT5G26280 | | TRAF-like family protein | 1.2 | **$1.06 \times 10^{-5}$** | **$1.94 \times 10^{-4}$** | 1.3 | **$1.76 \times 10^{-5}$** | **$1.99 \times 10^{-4}$** | 1.4 | **$2.53 \times 10^{-5}$** | **$2.07 \times 10^{-4}$** |
| AT5TE49575 | | ATREP9 transposon family,RC/Helitron transposon super family | | | | | | | | | |
| AT5G35740 | | Carbohydrate-binding X8 domain superfamily protein | 0.6 | **$2.30 \times 10^{-2}$** | $1.03 \times 10^{-1}$ | | | | 0.4 | **$1.47 \times 10^{-2}$** | **$5.05 \times 10^{-2}$** |
| AT5G35750 | *HK2* | Histidine kinase 2 | 0.6 | **$1.20 \times 10^{-3}$** | **$1.05 \times 10^{-2}$** | 0.6 | **$1.61 \times 10^{-5}$** | **$1.85 \times 10^{-4}$** | 0.4 | **$2.72 \times 10^{-3}$** | **$1.24 \times 10^{-2}$** |
| AT5G35740 | | Carbohydrate-binding X8 domain superfamily protein | 0.6 | **$2.30 \times 10^{-2}$** | $1.03 \times 10^{-1}$ | | | | 0.4 | **$1.47 \times 10^{-2}$** | **$5.05 \times 10^{-2}$** |
| AT5G35750 | *HK2* | Histidine kinase 2 | 0.6 | **$1.20 \times 10^{-3}$** | **$1.05 \times 10^{-2}$** | 0.6 | **$1.61 \times 10^{-5}$** | **$1.85 \times 10^{-4}$** | 0.4 | **$2.72 \times 10^{-3}$** | **$1.24 \times 10^{-2}$** |
| AT5G35740 | | Carbohydrate-binding X8 domain superfamily protein | 0.6 | **$2.30 \times 10^{-2}$** | $1.03 \times 10^{-1}$ | | | | 0.4 | **$1.47 \times 10^{-2}$** | **$5.05 \times 10^{-2}$** |
| AT5G35750 | *HK2* | Histidine kinase 2 | 0.6 | **$1.20 \times 10^{-3}$** | **$1.05 \times 10^{-2}$** | 0.6 | **$1.61 \times 10^{-5}$** | **$1.85 \times 10^{-4}$** | 0.4 | **$2.72 \times 10^{-3}$** | **$1.24 \times 10^{-2}$** |
| AT5G35750 | *HK2* | Histidine kinase 2 | 0.6 | **$1.20 \times 10^{-3}$** | **$1.05 \times 10^{-2}$** | 0.6 | **$1.61 \times 10^{-5}$** | **$1.85 \times 10^{-4}$** | 0.4 | **$2.72 \times 10^{-3}$** | **$1.24 \times 10^{-2}$** |
| AT5G35753 | | DnaJ heat shock amino-terminal domain protein (DUF3444) | | | | | | | | | |
| AT5G35753 | | DnaJ heat shock amino-terminal domain protein (DUF3444) | | | | | | | | | |
| AT5TE49650 | | ATLINE1_2 transposon family,LINE/L1 transposon super family | | | | | | | | | |
| AT5TE49655 | | ATCOPIA8A transposon family,LTR/Copia transposon super family | | | | | | | | | |
| AT5G37650 | | Hypothetical protein (DUF577) | | | | | | | | | |
| AT5G37650 | | Hypothetical protein (DUF577) | | | | | | | | | |
| AT5G40300 | *CASPL4A1* | Uncharacterized protein family (UPF0497) | | | | | | | | | |
| AT5G42720 | | Glycosyl hydrolase family 17 protein | -0.5 | **$1.13 \times 10^{-2}$** | $6.09 \times 10^{-2}$ | -0.9 | **$7.54 \times 10^{-5}$** | **$7.02 \times 10^{-4}$** | -0.9 | **$1.14 \times 10^{-10}$** | **$2.54 \times 10^{-9}$** |
| AT5G49430 | | WD40/YVTN repeat and Bromo-WDR9-I-like domain-containing protein | 0.4 | **$4.59 \times 10^{-2}$** | $1.68 \times 10^{-1}$ | 0.5 | **$1.28 \times 10^{-3}$** | **$7.40 \times 10^{-3}$** | | | |
| AT5G51970 | *ATSDH* | Sorbitol dehydrogenase | | | | | | | | | |
| AT5G51960 | | Complex 1 protein, LYR family protein | | | | | | | | | |
| AT5G51980 | *ATC3H63* | Transducin/WD40 repeat-like superfamily protein | | | | | | | | | |
| AT5G51990 | *DREB1D* | Dehydration-responsive element-binding protein 1D | | | | | | | | | |
| AT5G54660 | | HSP20-like chaperones superfamily protein | | | | | | | | | |
| AT5G62110 | | Homeodomain-like superfamily protein | | | | | | | | | |
| AT5G62090 | *SLK2* | Seuss-like 2 | | | | | | | | | |

The transcriptional memory behaviour of the genes delineated by eGWAS are shown. The results of a differential gene expression analysis of 12-d-old seedlings of *Arabidopsis thaliana*, ecotype: Columbia (Col-0), grown in 1/4-Hoagland's solution in hydroponics and then exposed for three days to a 5% PEG-induced first drought (D1) versus plants growing continuously for 15 days in hydroponics (watered), and plants exposed to another three-days second drought (D2) versus the first drought (D1), separated by a recovery interval (R) of three days, are presented. Three biological replicates, each comprising 10 seedlings, were used for the transcriptome analysis. Fisher's exact test *P*-values, as well as the Benjamini-Hochberg *false discovery rate* (*FDR*), are presented.

### Table S7 Local association study of superinduction of *LKR* in 80 *Arabidopsis thaliana* ecotypes under recurrent drought

| Chromosome | Physical position | Frequency of high expression allele | Average RFC high expression allele | Frequency of low-expression allele | Average RFC low-expression allele | Student's *t*-test *P*-value | LAS GLM *F*-statistic | LAS GLM *P*-value | LAS MLM *F*-statistic | LAS MLM *P*-value | GWAS-detected SNP | Polymorphism effects | Impact | Gene ID | Gene symbol | Short description |
|---|---|---|---|---|---|---|---|---|---|---|---|---|---|---|---|---|
| 1 | 21237954 | 70/C | 3.9 ± 0.9 | 10/G | 2.8 ± 0.8 | $6.27 \times 10^{-1}$ | 2.0 | $1.64 \times 10^{-1}$ | 1.27 | $2.63 \times 10^{-1}$ | | 5'-UTR intron variant | Moderate | AT1G56660 | | MAEBL domain protein |
| 1 | 21238164 | 58/C | 4.1 ± 1.1 | 22/G | 3 ± 0.8 | $5.50 \times 10^{-1}$ | 0.0 | $9.92 \times 10^{-1}$ | 0.01 | $9.33 \times 10^{-1}$ | | 5'-UTR intron variant | Moderate | | | |
| 1 | 21238410 | 9/T | 6.5 ± 2.7 | 71/A | 3.4 ± 0.8 | $2.10 \times 10^{-1}$ | **5.0** | **$2.91 \times 10^{-2}$** | **3.27** | **$7.49 \times 10^{-2}$** | | 5'-UTR intron variant | Moderate | | | |
| 1 | 21238670 | 58/T | 4 ± 1.1 | 20/G | 3.2 ± 0.8 | $6.49 \times 10^{-1}$ | 0.2 | $8.01 \times 10^{-1}$ | 0.17 | $8.40 \times 10^{-1}$ | | 5'-UTR intron variant | Moderate | | | |
| 1 | 21239069 | 75/G | 3.9 ± 0.9 | 5/T | 2.1 ± 0.6 | $5.74 \times 10^{-1}$ | 0.0 | $9.82 \times 10^{-1}$ | 0.01 | $9.33 \times 10^{-1}$ | | amino acid substitution (Met87Ile) | Moderate | | | |
| 1 | 21239103 | 75/C | 3.9 ± 0.9 | 5/G | 2.1 ± 0.6 | $5.71 \times 10^{-1}$ | 0.3 | $5.84 \times 10^{-1}$ | 0.48 | $4.90 \times 10^{-1}$ | | amino acid substitution (Leu99Val) | Moderate | | | |
| 1 | 21240160 | 75/A | 3.9 ± 0.9 | 5/G | 1.7 ± 0.4 | $4.94 \times 10^{-1}$ | 0.1 | $7.44 \times 10^{-1}$ | 0.15 | $6.99 \times 10^{-1}$ | | amino acid substitution (Asp451Gly) | Moderate | | | |
| 1 | 21240309 | 10/C | 6.9 ± 2.4 | 70/A | 3.3 ± 0.9 | $1.23 \times 10^{-1}$ | **4.3** | **$4.06 \times 10^{-2}$** | 2.67 | $1.06 \times 10^{-1}$ | ✓ | amino acid substitution (Ile501Leu) | Moderate | | | |
| 1 | 24405454 | 9/G | 8.4 ± 2.7 | 71/A | 3.2 ± 0.8 | **$3.10 \times 10^{-2}$** | **4.2** | **$4.49 \times 10^{-2}$** | 2.25 | $1.38 \times 10^{-1}$ | ✓ | intergenic space variant | Moderate | AT1G65630 | *DEG3* | Degp protease 3 |
| 1 | 24405565 | 8/C | 7.8 ± 2.9 | 72/A | 3.3 ± 0.8 | **$7.87 \times 10^{-2}$** | **3.0** | **$9.01 \times 10^{-2}$** | 1.82 | $1.81 \times 10^{-1}$ | ✓ | intergenic space variant | Moderate | | | |
| 1 | 24406598 | 79/G | 3.8 ± 0.8 | 1/A | 3.4 | | 0.00 | $9.67 \times 10^{-1}$ | 0.01 | $9.08 \times 10^{-1}$ | | stop codon gain (Gln542*) | **High** | | | |
| 1 | 24407370 | 79/T | 3.8 ± 0.8 | 1/A | 1.4 | | 0.07 | $7.99 \times 10^{-1}$ | 0.07 | $7.89 \times 10^{-1}$ | | stop codon gain (Lys385*) | **High** | | | |
| 1 | 24407561 | 12/T | 7.9 ± 4.8 | 68/C | 3 ± 0.4 | **$2.25 \times 10^{-2}$** | **3.7** | **$5.96 \times 10^{-2}$** | **3.29** | **$7.41 \times 10^{-2}$** | | amino acid substitution (Ile344Val) | Moderate | | | |
| 1 | 24407670 | 37/G | 4.3 ± 1.6 | 43/C | 3.3 ± 0.7 | $5.24 \times 10^{-1}$ | 0.0 | $9.84 \times 10^{-1}$ | 0.05 | $8.17 \times 10^{-1}$ | | amino acid substitution (Asp307Glu) | Moderate | | | |
| 1 | 24407717 | 67/C | 4 ± 1 | 13/G | 2.8 ± 0.7 | $5.91 \times 10^{-1}$ | 0.4 | $5.09 \times 10^{-1}$ | 0.31 | $5.79 \times 10^{-1}$ | | amino acid substitution (Glu292Gln) | Moderate | | | |
| 1 | 24407909 | 79/A | 3.8 ± 0.8 | 1/AT | 1.9 | | 0.00 | $9.65 \times 10^{-1}$ | 0.07 | $7.85 \times 10^{-1}$ | | splice donor variant | Moderate | | | |
| 1 | 24408977 | 38/A | 4.1 ± 1.5 | 42/G | 3.5 ± 0.7 | $6.71 \times 10^{-1}$ | 0.2 | $6.47 \times 10^{-1}$ | 0.39 | $5.32 \times 10^{-1}$ | | amino acid substitution (Val31Ala) | Moderate | | | |
| 1 | 24409269 | 72/A | 3.9 ± 0.9 | 8/G | 2.7 ± 0.4 | $6.29 \times 10^{-1}$ | 1.0 | $3.17 \times 10^{-1}$ | 1.74 | $1.91 \times 10^{-1}$ | | Promoter variant | Moderate | | | |
| 1 | 24409811 | 7/A | 4.2 ± 1 | 73/G | 3.7 ± 0.9 | $8.78 \times 10^{-1}$ | 0.0 | $9.93 \times 10^{-1}$ | 0.10 | $7.58 \times 10^{-1}$ | | Promoter variant | Moderate | | | |
| 1 | 24409826 | 21/A | 4.6 ± 1.2 | 59/T | 3.5 ± 1 | $5.17 \times 10^{-1}$ | 1.2 | $2.82 \times 10^{-1}$ | 0.85 | $3.60 \times 10^{-1}$ | | Promoter variant | Moderate | | | |
| 1 | 24409953 | 72/A | 3.9 ± 0.9 | 8/C | 2.5 ± 0.4 | $5.97 \times 10^{-1}$ | 0.5 | $4.95 \times 10^{-1}$ | 1.18 | $2.80 \times 10^{-1}$ | | Promoter variant | Moderate | | | |
| 1 | 25140573 | 52/C | 3.9 ± 1.1 | 28/T | 3.6 ± 1 | $8.89 \times 10^{-1}$ | 0.1 | $7.31 \times 10^{-1}$ | 0.14 | $7.09 \times 10^{-1}$ | | Promoter variant | Moderate | AT1G67210 | *ZCCHC8B* | Proline-rich spliceosome-associated (PSP) family protein |
| 1 | 25140936 | 76/A | 3.9 ± 0.9 | 4/T | 2.2 ± 1 | $6.35 \times 10^{-1}$ | 0.2 | $6.76 \times 10^{-1}$ | 0.08 | $7.84 \times 10^{-1}$ | | 5'-UTR intron variant | Moderate | | | |
| 1 | 25141110 | 57/A | 4.3 ± 1.1 | 23/G | 2.5 ± 0.4 | $3.04 \times 10^{-1}$ | 1.1 | $2.97 \times 10^{-1}$ | 0.61 | $4.37 \times 10^{-1}$ | | 5'-UTR intron variant | Moderate | | | |
| 1 | 25141267 | 3/C | 5.4 ± 3.9 | 77/G | 3.7 ± 0.8 | $6.79 \times 10^{-1}$ | 0.0 | $9.90 \times 10^{-1}$ | 0.05 | $8.26 \times 10^{-1}$ | | splice region variant | **High** | | | |
| 1 | 25141315 | 57/A | 4.3 ± 1.1 | 23/T | 2.6 ± 0.4 | $3.34 \times 10^{-1}$ | 1.5 | $2.27 \times 10^{-1}$ | 1.10 | $2.97 \times 10^{-1}$ | | 5'-UTR variant | Moderate | | | |
| 1 | 25143543 | 8/T | 13.5 ± 6.7 | 72/A | 2.7 ± 0.3 | **$9.77 \times 10^{-6}$** | **24.5** | **$5.00 \times 10^{-6}$** | **16.48** | **$1.23 \times 10^{-4}$** | ✓ | synonymous variant | Moderate | | | |
| 1 | 25143688 | 63/T | 4.1 ± 1 | 17/G | 2.7 ± 0.8 | $4.55 \times 10^{-1}$ | **3.5** | **$6.37 \times 10^{-2}$** | **3.45** | **$6.75 \times 10^{-2}$** | | amino acid substitution (Tyr378Asp) | Moderate | | | |
| 1 | 25143712 | 75/C | 3.9 ± 0.9 | 5/G | 2.1 ± 0.5 | $5.86 \times 10^{-1}$ | 0.0 | $8.95 \times 10^{-1}$ | 0.03 | $8.66 \times 10^{-1}$ | | amino acid substitution (Gln386Glu) | Moderate | | | |
| 1 | 25144630 | 69/A | 3.9 ± 0.9 | 11/C | 3 ± 1.1 | $6.81 \times 10^{-1}$ | **3.1** | **$8.33 \times 10^{-2}$** | 2.27 | $1.36 \times 10^{-1}$ | | Promoter variant | Moderate | AT1G08913 | | Long non-coding RNA |
| 1 | 25144639 | 25/G | 4.6 ± 2.2 | 55/A | 3.4 ± 0.6 | $4.73 \times 10^{-1}$ | 2.8 | **$9.93 \times 10^{-2}$** | 2.75 | $1.02 \times 10^{-1}$ | | Promoter variant | Moderate | | | |

| | | | | | | | | | | | | | | | |
|---|---|---|---|---|---|---|---|---|---|---|---|---|---|---|---|
| 1 | 25144663 | 52/C | 4.2 ± 1.2 | 28/T | 3 ± 0.5 | $4.36 \times 10^{-1}$ | 0.2 | $6.98 \times 10^{-1}$ | 0.01 | $9.33 \times 10^{-1}$ | | Promoter variant | Moderate | | |
| 1 | 25144748 | 61/C | 4 ± 1 | 19/T | 2.9 ± 0.8 | $5.43 \times 10^{-1}$ | 0.7 | $4.07 \times 10^{-1}$ | 0.31 | $5.81 \times 10^{-1}$ | | Promoter variant | Moderate | | |
| 1 | 25144752 | 70/C | 3.9 ± 0.9 | 10/T | 3.1 ± 1.2 | $7.56 \times 10^{-1}$ | 0.8 | $3.82 \times 10^{-1}$ | 0.30 | $5.83 \times 10^{-1}$ | | Promoter variant | Moderate | | |
| 1 | 25144780 | 33/A | 4.9 ± 1.8 | 47/G | 3 ± 0.5 | $2.16 \times 10^{-1}$ | 2.2 | $1.38 \times 10^{-1}$ | 1.30 | $2.57 \times 10^{-1}$ | | Promoter variant | Moderate | | |
| 1 | 25144800 | 29/T | 5.3 ± 2.1 | 51/C | 2.9 ± 0.4 | $1.24 \times 10^{-1}$ | **3.4** | **$6.87 \times 10^{-2}$** | 2.30 | $1.34 \times 10^{-1}$ | | Promoter variant | Moderate | | |
| 1 | 25145105 | 55/G | 4.2 ± 1.1 | 25/A | 2.8 ± 0.6 | $3.76 \times 10^{-1}$ | 2.1 | $1.51 \times 10^{-1}$ | 1.76 | $1.89 \times 10^{-1}$ | | Promoter variant | Moderate | | |
| 1 | 25145183 | 65/G | 4 ± 1 | 15/A | 2.7 ± 0.9 | $5.04 \times 10^{-1}$ | 2.8 | **$9.89 \times 10^{-2}$** | 2.18 | $1.44 \times 10^{-1}$ | | Promoter variant | Moderate | | |
| 1 | 25145374 | 13/T | 10.1 ± 4.3 | 67/A | 2.5 ± 0.3 | **$1.75 \times 10^{-4}$** | **15.5** | **$1.89 \times 10^{-4}$** | **9.38** | **$3.09 \times 10^{-3}$** | | Promoter variant | Moderate | | |
| 2 | 8287360 | 54/C | 4 ± 1.2 | 26/T | 3.4 ± 0.6 | $7.49 \times 10^{-1}$ | 0.3 | $5.92 \times 10^{-1}$ | 0.00 | $9.80 \times 10^{-1}$ | | amino acid substitution (Arg1013Lys) | Moderate | AT2G19120 | P-loop containing nucleoside triphosphate hydrolases superfamily protein |
| 2 | 8287513 | 55/T | 4 ± 1.1 | 25/G | 3.3 ± 0.6 | $6.48 \times 10^{-1}$ | 0.5 | $4.79 \times 10^{-1}$ | 0.19 | $6.66 \times 10^{-1}$ | | amino acid substitution (Asp962Ala) | Moderate | | |
| 2 | 8287529 | 57/T | 4.2 ± 1.1 | 23/C | 2.6 ± 0.6 | $3.41 \times 10^{-1}$ | 0.7 | $4.05 \times 10^{-1}$ | 0.80 | $3.74 \times 10^{-1}$ | | amino acid substitution (Asn957Asp) | Moderate | | |
| 2 | 8287649 | 53/A | 4 ± 1.2 | 27/C | 3.3 ± 0.6 | $6.94 \times 10^{-1}$ | 0.5 | $4.93 \times 10^{-1}$ | 0.10 | $7.56 \times 10^{-1}$ | | amino acid substitution (Cys917Gly) | Moderate | | |
| 2 | 8287776 | 52/G | 4 ± 1.2 | 28/A | 3.3 ± 0.5 | $6.56 \times 10^{-1}$ | 0.6 | $4.23 \times 10^{-1}$ | 0.30 | $5.86 \times 10^{-1}$ | | amino acid substitution (Ser874Arg) | Moderate | | |
| 2 | 8288763 | 52/C | 4 ± 1.2 | 28/A | 3.3 ± 0.5 | $6.56 \times 10^{-1}$ | 0.6 | $4.23 \times 10^{-1}$ | 0.30 | $5.86 \times 10^{-1}$ | | amino acid substitution (Glu545Asp) | Moderate | | |
| 2 | 8291131 | 52/A | 4 ± 1.2 | 28/T | 3.3 ± 0.5 | $6.56 \times 10^{-1}$ | 0.6 | $4.23 \times 10^{-1}$ | 0.30 | $5.86 \times 10^{-1}$ | | amino acid substitution (Ser224Arg) | Moderate | | |
| 2 | 8291211 | 11/T | 11.3 ± 5.1 | 69/A | 2.6 ± 0.3 | **$4.41 \times 10^{-5}$** | **13.6** | **$4.35 \times 10^{-4}$** | **9.19** | **$3.38 \times 10^{-3}$** | ✓ | amino acid substitution (Ile198Leu) | Moderate | | |
| 2 | 8291276 | 12/T | 10.6 ± 4.7 | 68/C | 2.6 ± 0.3 | **$9.99 \times 10^{-5}$** | **9.0** | **$3.71 \times 10^{-3}$** | **5.22** | **$2.52 \times 10^{-2}$** | ✓ | amino acid substitution (Glu176Gly) | Moderate | | |
| 2 | 8291447 | 70/G | 4 ± 0.9 | 10/A | 2.5 ± 0.4 | $5.45 \times 10^{-1}$ | 0.3 | $5.60 \times 10^{-1}$ | 0.28 | $5.99 \times 10^{-1}$ | | amino acid substitution (Pro119Leu) | Moderate | | |
| 2 | 8291696 | 53/G | 4 ± 1.2 | 27/T | 3.4 ± 0.6 | $7.12 \times 10^{-1}$ | 0.6 | $4.44 \times 10^{-1}$ | 0.20 | $6.55 \times 10^{-1}$ | | amino acid substitution (Pro36Gln) | Moderate | | |
| 2 | 8291771 | 53/G | 4 ± 1.2 | 27/T | 3.3 ± 0.6 | $6.93 \times 10^{-1}$ | 0.4 | $5.27 \times 10^{-1}$ | 0.05 | $8.24 \times 10^{-1}$ | | amino acid substitution (Thr11Lys) | Moderate | | |
| 2 | 8291790 | 56/C | 4 ± 1.1 | 24/T | 3.4 ± 0.6 | $7.29 \times 10^{-1}$ | 0.5 | $4.80 \times 10^{-1}$ | 0.15 | $6.97 \times 10^{-1}$ | | amino acid substitution (Ala5Thr) | Moderate | | |
| 2 | 8291884 | 52/A | 4 ± 1.2 | 28/C | 3.3 ± 0.5 | $6.56 \times 10^{-1}$ | 0.6 | $4.23 \times 10^{-1}$ | 0.30 | $5.86 \times 10^{-1}$ | | 5'-UTR intron variant | Moderate | | |
| 2 | 8291909 | 52/T | 4 ± 1.2 | 28/A | 3.3 ± 0.5 | $6.56 \times 10^{-1}$ | 0.6 | $4.23 \times 10^{-1}$ | 0.30 | $5.86 \times 10^{-1}$ | | 5'-UTR intron variant | Moderate | | |
| 2 | 8291961 | 48/T | 4 ± 1.3 | 32/TC | 3.4 ± 0.5 | $6.92 \times 10^{-1}$ | 0.4 | $5.25 \times 10^{-1}$ | 0.00 | $9.57 \times 10^{-1}$ | | 5'-UTR intron variant | Moderate | | |
| 2 | 8291966 | 47/T | 4.1 ± 1.3 | 33/C | 3.4 ± 0.5 | $6.51 \times 10^{-1}$ | 0.2 | $6.54 \times 10^{-1}$ | 0.02 | $8.88 \times 10^{-1}$ | | 5'-UTR intron variant | Moderate | | |
| 2 | 8291975 | 15/G | 8.5 ± 3.9 | 65/C | 2.7 ± 0.3 | **$2.95 \times 10^{-3}$** | **7.2** | **$9.10 \times 10^{-3}$** | **3.95** | **$5.08 \times 10^{-2}$** | | 5'-UTR intron variant | Moderate | | |
| 2 | 8292002 | 45/T | 4.2 ± 1.4 | 35/C | 3.2 ± 0.5 | $5.31 \times 10^{-1}$ | 0.3 | $5.77 \times 10^{-1}$ | 0.00 | $9.81 \times 10^{-1}$ | | 5'-UTR intron variant | Moderate | | |
| 2 | 8292031 | 43/A | 4.3 ± 1.4 | 37/T | 3.1 ± 0.4 | $4.38 \times 10^{-1}$ | 0.9 | $3.39 \times 10^{-1}$ | 0.41 | $5.23 \times 10^{-1}$ | | 5'-UTR intron variant | Moderate | | |
| 2 | 8292099 | 43/C | 4.3 ± 1.4 | 37/T | 3.1 ± 0.4 | $4.50 \times 10^{-1}$ | 1.0 | $3.31 \times 10^{-1}$ | 0.45 | $5.04 \times 10^{-1}$ | | 5'-UTR intron variant | Moderate | | |
| 2 | 8292104 | 45/A | 4.2 ± 1.4 | 35/G | 3.2 ± 0.5 | $5.07 \times 10^{-1}$ | 0.3 | $5.68 \times 10^{-1}$ | 0.00 | $9.59 \times 10^{-1}$ | | 5'-UTR intron variant | Moderate | | |
| 2 | 8292136 | 51/A | 4.1 ± 1.2 | 29/T | 3.2 ± 0.5 | $5.83 \times 10^{-1}$ | 0.2 | $6.39 \times 10^{-1}$ | 0.00 | $9.51 \times 10^{-1}$ | | 5'-UTR variant | Moderate | | |
| 2 | 8292145 | 60/GA | 4.1 ± 1.1 | 20/G | 2.7 ± 0.5 | $4.27 \times 10^{-1}$ | 0.1 | $7.01 \times 10^{-1}$ | 0.02 | $8.98 \times 10^{-1}$ | | 5'-UTR variant | Moderate | | |
| 2 | 8292161 | 70/T | 4 ± 0.9 | 10/A | 2.5 ± 0.4 | $5.45 \times 10^{-1}$ | 0.3 | $5.60 \times 10^{-1}$ | 0.28 | $5.99 \times 10^{-1}$ | | 5'-UTR variant | Moderate | | |
| 2 | 8292175 | 48/T | 4.2 ± 1.3 | 32/G | 3.2 ± 0.5 | $5.25 \times 10^{-1}$ | 0.3 | $6.00 \times 10^{-1}$ | 0.00 | $9.60 \times 10^{-1}$ | | 5'-UTR variant | Moderate | | |
| 2 | 8292229 | 19/T | 6.9 ± 3.1 | 61/A | 2.8 ± 0.3 | **$2.06 \times 10^{-2}$** | **3.8** | **$5.44 \times 10^{-2}$** | 1.39 | $2.43 \times 10^{-1}$ | | 5'-UTR variant | Moderate | | |
| 2 | 8292240 | 16/C | 8 ± 3.7 | 64/T | 2.7 ± 0.3 | **$4.81 \times 10^{-3}$** | **7.9** | **$6.26 \times 10^{-3}$** | **4.31** | **$4.15 \times 10^{-2}$** | | 5'-UTR variant | Moderate | | |
| 2 | 8292260 | 19/C | 7.4 ± 3.1 | 61/T | 2.7 ± 0.3 | **$8.42 \times 10^{-3}$** | **7.0** | **$1.00 \times 10^{-2}$** | **4.55** | **$3.63 \times 10^{-2}$** | | 5'-UTR variant | Moderate | | |

| | | | | | | | | | | | | | | |
|---|---|---|---|---|---|---|---|---|---|---|---|---|---|---|
| 2 | 8292324 | 66/A | 3.8 ± 1 | 14/C | 3.4 ± 0.8 | 8.40 × $10^{-1}$ | 0.9 | 3.39 × $10^{-1}$ | 0.64 | 4.26 × $10^{-1}$ | 5'-UTR premature start codon gain variant | Moderate | | |
| 2 | 8292358 | 25/T | 6.2 ± 2.4 | 55/C | 2.7 ± 0.3 | **3.01 × $10^{-2}$** | 1.7 | 1.99 × $10^{-1}$ | 0.42 | 5.19 × $10^{-1}$ | 5'-UTR premature start codon gain variant | Moderate | | |
| 2 | 8292376 | 28/G | 5.7 ± 2.2 | 52/A | 2.7 ± 0.4 | **6.01 × $10^{-2}$** | 1.2 | 2.77 × $10^{-1}$ | 0.06 | 8.11 × $10^{-1}$ | 5'-UTR variant | Moderate | | |
| 2 | 8292387 | 28/A | 5.8 ± 2.1 | 52/T | 2.7 ± 0.3 | **4.90 × $10^{-2}$** | 1.6 | 2.10 × $10^{-1}$ | 0.36 | 5.50 × $10^{-1}$ | 5'-UTR variant | Moderate | | |
| 2 | 8292396 | 31/A | 5.4 ± 2 | 49/T | 2.7 ± 0.4 | **9.31 × $10^{-2}$** | 0.7 | 3.91 × $10^{-1}$ | 0.01 | 9.19 × $10^{-1}$ | 5'-UTR variant | Moderate | | |
| 2 | 8292422 | 73/G | 4 ± 0.9 | 7/A | 1.9 ± 0.7 | 4.46 × $10^{-1}$ | 0.9 | 3.37 × $10^{-1}$ | 0.65 | 4.24 × $10^{-1}$ | 5'-UTR premature start codon gain variant | Moderate | | |
| 2 | 8292425 | 33/A | 5.6 ± 1.8 | 47/G | 2.5 ± 0.3 | **5.08 × $10^{-2}$** | 2.3 | 1.36 × $10^{-1}$ | 1.63 | 2.06 × $10^{-1}$ | 5'-UTR variant | Moderate | | |
| 2 | 8292455 | 30/G | 4.4 ± 1.9 | 50/A | 3.4 ± 0.6 | 5.20 × $10^{-1}$ | 0.3 | 6.08 × $10^{-1}$ | 0.63 | 4.31 × $10^{-1}$ | 5'-UTR variant | Moderate | | |
| 2 | 8292785 | 64/A | 4.2 ± 1 | 16/G | 1.9 ± 0.3 | 2.23 × $10^{-1}$ | 0.5 | 4.88 × $10^{-1}$ | 0.17 | 6.78 × $10^{-1}$ | Promoter variant | Moderate | | |
| 3 | 8419320 | 74/A | 3.8 ± 0.9 | 6/G | 3.4 ± 1 | 9.01 × $10^{-1}$ | 1.0 | 3.16 × $10^{-1}$ | 1.16 | 2.85 × $10^{-1}$ | 5'-UTR intron variant | Moderate | AT3G23480 | Cyclopropane-fatty-acyl-phospholipid synthase |
| 3 | 8419377 | 73/T | 4 ± 0.9 | 7/C | 1.7 ± 0.4 | 4.14 × $10^{-1}$ | 0.5 | 4.95 × $10^{-1}$ | 0.19 | 6.61 × $10^{-1}$ | 5'-UTR intron variant | Moderate | | |
| 3 | 8419402 | 73/C | 4 ± 0.9 | 7/A | 1.5 ± 0.3 | 3.69 × $10^{-1}$ | 0.4 | 5.41 × $10^{-1}$ | 0.16 | 6.90 × $10^{-1}$ | 5'-UTR intron variant | Moderate | | |
| 3 | 8419445 | 74/G | 3.8 ± 0.9 | 6/T | 3.4 ± 1 | 9.01 × $10^{-1}$ | 1.0 | 3.16 × $10^{-1}$ | 1.16 | 2.85 × $10^{-1}$ | 5'-UTR intron variant | Moderate | | |
| 3 | 8419452 | 75/A | 3.8 ± 0.9 | 5/G | 3.7 ± 1.2 | 9.71 × $10^{-1}$ | 0.1 | 7.61 × $10^{-1}$ | 0.12 | 7.29 × $10^{-1}$ | 5'-UTR intron variant | Moderate | | |
| 3 | 8419522 | 74/C | 3.8 ± 0.9 | 6/A | 3.4 ± 1.4 | 8.97 × $10^{-1}$ | 0.0 | 9.97 × $10^{-1}$ | 0.00 | 9.80 × $10^{-1}$ | 5'-UTR intron variant | Moderate | | |
| 3 | 8419547 | 49/T | 4.5 ± 1.3 | 31/C | 2.7 ± 0.5 | 2.60 × $10^{-1}$ | 1.5 | 2.17 × $10^{-1}$ | 0.83 | 3.67 × $10^{-1}$ | 5'-UTR intron variant | Moderate | | |
| 3 | 8419553 | 74/T | 3.8 ± 0.9 | 6/C | 3.5 ± 1.4 | 9.25 × $10^{-1}$ | 0.4 | 5.19 × $10^{-1}$ | 0.65 | 4.21 × $10^{-1}$ | 5'-UTR intron variant | Moderate | | |
| 3 | 8419669 | 73/A | 3.9 ± 0.9 | 7/G | 2.9 ± 1.3 | 7.12 × $10^{-1}$ | 0.5 | 4.96 × $10^{-1}$ | 0.30 | 5.87 × $10^{-1}$ | 5'-UTR intron variant | Moderate | | |
| 3 | 8419677 | 21/G | 5.3 ± 2.6 | 59/A | 3.2 ± 0.6 | 2.44 × $10^{-1}$ | 2.5 | 1.18 × $10^{-1}$ | 1.86 | 1.77 × $10^{-1}$ | 5'-UTR intron variant | Moderate | | |
| 3 | 8419984 | 75/G | 3.8 ± 0.9 | 5/A | 2.9 ± 0.6 | 7.83 × $10^{-1}$ | 0.1 | 7.36 × $10^{-1}$ | 0.41 | 5.23 × $10^{-1}$ | 5'-UTR intron variant | Moderate | | |
| 3 | 8420110 | 30/A | 4.3 ± 1.9 | 50/T | 3.5 ± 0.6 | 6.10 × $10^{-1}$ | 0.3 | 5.79 × $10^{-1}$ | 0.06 | 8.14 × $10^{-1}$ | 5'-UTR intron variant | Moderate | | |
| 3 | 8420140 | 73/C | 3.8 ± 0.9 | 7/T | 3.1 ± 1.3 | 7.93 × $10^{-1}$ | 0.8 | 3.66 × $10^{-1}$ | 0.67 | 4.15 × $10^{-1}$ | 5'-UTR intron variant | Moderate | | |
| 3 | 8420143 | 73/A | 3.8 ± 0.9 | 7/T | 3.1 ± 1.3 | 7.93 × $10^{-1}$ | 0.8 | 3.66 × $10^{-1}$ | 0.67 | 4.15 × $10^{-1}$ | 5'-UTR intron variant | Moderate | | |
| 3 | 8420207 | 19/T | 4.1 ± 1.3 | 61/C | 3.7 ± 1 | 8.36 × $10^{-1}$ | 1.1 | 3.05 × $10^{-1}$ | 0.84 | 3.63 × $10^{-1}$ | 5'-UTR intron variant | Moderate | | |
| 3 | 8420218 | 72/G | 3.8 ± 0.9 | 8/A | 3.2 ± 0.8 | 8.05 × $10^{-1}$ | 1.8 | 1.80 × $10^{-1}$ | 2.02 | 1.60 × $10^{-1}$ | 5'-UTR intron variant | Moderate | | |
| 3 | 8420239 | 62/A | 4 ± 1 | 18/G | 2.9 ± 0.6 | 5.64 × $10^{-1}$ | 0.0 | 8.78 × $10^{-1}$ | 0.06 | 8.02 × $10^{-1}$ | 5'-UTR intron variant | Moderate | | |
| 3 | 8420258 | 72/T | 3.9 ± 0.9 | 8/C | 3 ± 1.1 | 7.24 × $10^{-1}$ | 1.1 | 3.06 × $10^{-1}$ | 1.12 | 2.93 × $10^{-1}$ | 5'-UTR intron variant | Moderate | | |
| 3 | 8420277 | 72/G | 3.9 ± 0.9 | 8/A | 3.1 ± 0.8 | 7.65 × $10^{-1}$ | 0.4 | 5.41 × $10^{-1}$ | 0.33 | 5.69 × $10^{-1}$ | splice acceptor variant | **High** | | |
| 3 | 8420287 | 67/G | 3.9 ± 1 | 13/A | 3 ± 0.5 | 6.76 × $10^{-1}$ | 0.9 | 3.44 × $10^{-1}$ | 1.53 | 2.21 × $10^{-1}$ | amino acid substitution (Val2Ile) | Moderate | | |
| 3 | 8420290 | 67/A | 3.9 ± 1 | 13/G | 3.1 ± 0.8 | 7.02 × $10^{-1}$ | 0.1 | 7.36 × $10^{-1}$ | 0.17 | 6.83 × $10^{-1}$ | amino acid substitution (Ile3Val) | Moderate | | |
| 3 | 8420326 | 71/G | 3.9 ± 0.9 | 9/C | 3 ± 0.7 | 7.38 × $10^{-1}$ | 1.1 | 2.94 × $10^{-1}$ | 1.11 | 2.96 × $10^{-1}$ | amino acid substitution (Val15Met) | Moderate | | |
| 3 | 8420332 | 71/A | 3.9 ± 0.9 | 9/G | 2.7 ± 0.8 | 6.27 × $10^{-1}$ | 1.0 | 3.24 × $10^{-1}$ | 1.12 | 2.93 × $10^{-1}$ | amino acid substitution (Asn17Asp) | Moderate | | |
| 3 | 8420334 | 71/T | 3.9 ± 0.9 | 9/A | 3 ± 0.7 | 7.38 × $10^{-1}$ | 1.1 | 2.94 × $10^{-1}$ | 1.11 | 2.96 × $10^{-1}$ | amino acid substitution (Asn17Lys) | Moderate | | |
| 3 | 8420411 | 77/C | 3.8 ± 0.8 | 3/A | 2.1 ± 0.6 | 6.75 × $10^{-1}$ | 0.00 | 9.67 × $10^{-1}$ | 0.19 | 6.61 × $10^{-1}$ | stop codon gain (Ser43*) | **High** | | |
| 3 | 8420752 | 73/G | 3.9 ± 0.9 | 7/T | 2.8 ± 0.6 | 6.83 × $10^{-1}$ | 0.09 | 7.67 × $10^{-1}$ | 0.02 | 8.90 × $10^{-1}$ | stop codon gain (Gly107*) | **High** | | |
| 3 | 8420999 | 79/AG | 3.8 ± 0.8 | 1/A | 0.4 | | 0.01 | 9.16 × $10^{-1}$ | 0.08 | 7.78 × $10^{-1}$ | splice acceptor variant | **High** | | |
| 3 | 8421018 | 76/TG | 3.8 ± 0.9 | 4/T | 3.1 ± 0.7 | 8.47 × $10^{-1}$ | 0.01 | 9.34 × $10^{-1}$ | 0.03 | 8.73 × $10^{-1}$ | frameshift variant (Val138) | **High** | | |

| | | | | | | | | | | | | | | | |
|---|---|---|---|---|---|---|---|---|---|---|---|---|---|---|---|
| 3 | 8421494 | 70/A | 3.8 ± 0.9 | 10/G | 3.7 ± 1.5 | $9.87 \times 10^{-1}$ | 0.02 | $9.02 \times 10^{-1}$ | 0.37 | $5.43 \times 10^{-1}$ | | splice acceptor variant | **High** | | |
| 3 | 8421509 | 59/CAT | 4 ± 1.1 | 21/C | 3.2 ± 0.7 | $6.47 \times 10^{-1}$ | 0.20 | $6.54 \times 10^{-1}$ | 0.01 | $9,27 \times 10^{-1}$ | | frameshift variant (Pro219) | **High** | | |
| 3 | 8421607 | 50/G | 3.9 ± 1.2 | 30/A | 3.6 ± 0.9 | $8.60 \times 10^{-1}$ | 0.0 | $9.34 \times 10^{-1}$ | 0.03 | $8.55 \times 10^{-1}$ | | amino acid substitution (Glu252Lys) | Moderate | | |
| 3 | 8421646 | 73/G | 3.9 ± 0.9 | 7/T | 2.8 ± 0.6 | $6.83 \times 10^{-1}$ | 0.09 | $7.67 \times 10^{-1}$ | 0.02 | $8.90 \times 10^{-1}$ | | stop codon gain (Gly265*) | **High** | | |
| 3 | 8421655 | 5/T | 4.4 ± 1.8 | 75/G | 3.7 ± 0.9 | $8.37 \times 10^{-1}$ | 0.11 | $7.43 \times 10^{-1}$ | 0.03 | $8.74 \times 10^{-1}$ | | splice donor variant | **High** | | |
| 3 | 8421725 | 37/C | 3.8 ± 0.8 | 43/G | 3.7 ± 1.3 | $9.43 \times 10^{-1}$ | 0.1 | $7.85 \times 10^{-1}$ | 0.13 | $7.15 \times 10^{-1}$ | | amino acid substitution (Glu268Asp) | Moderate | | |
| 3 | 8421746 | 64/C | 3.9 ± 1 | 16/A | 3.2 ± 0.7 | $6.93 \times 10^{-1}$ | 0.08 | $7.84 \times 10^{-1}$ | 0.10 | $7.54 \times 10^{-1}$ | | stop codon gain (Cys275*) | **High** | | |
| 3 | 8421921 | 79/CTG | 3.8 ± 0.8 | 1/C | 0.5 | | 0.00 | $9.79 \times 10^{-1}$ | 0.02 | $8.77 \times 10^{-1}$ | | frameshift variant (Cys310) | **High** | | |
| 3 | 8422269 | 77/G | 3.9 ± 0.8 | 3/A | 1.7 ± 1.2 | $5.95 \times 10^{-1}$ | 0.41 | $5.24 \times 10^{-1}$ | 0.24 | $6.26 \times 10^{-1}$ | | splice acceptor variant | **High** | | |
| 3 | 8422555 | 76/AAC | 3.8 ± 0.9 | 4/A | 3 ± 0.6 | $8.14 \times 10^{-1}$ | 0.07 | $7.93 \times 10^{-1}$ | 0.19 | $6.64 \times 10^{-1}$ | | frameshift variant (Thr433) | **High** | | |
| 3 | 8422878 | 9/G | 6.7 ± 2.7 | 71/C | 3.4 ± 0.8 | $1.71 \times 10^{-1}$ | **3.8** | $\mathbf{5.38 \times 10^{-2}}$ | 1.51 | $2.23 \times 10^{-1}$ | ✓ | 3' UTR variant | Moderate | | |
| 4 | 139361 | 51/G | 4.7 ± 1.2 | 29/A | 2.2 ± 0.3 | $1.22 \times 10^{-1}$ | 0.0 | $9.27 \times 10^{-1}$ | 0.13 | $7.18 \times 10^{-1}$ | | Promoter variant | Moderate | AT4G00320 | F-box/RNI-like superfamily protein |
| 4 | 139427 | 19/A | 5.8 ± 2.9 | 61/C | 3.2 ± 0.5 | $1.46 \times 10^{-1}$ | 0.1 | $7.23 \times 10^{-1}$ | 0.03 | $8.59 \times 10^{-1}$ | | Promoter variant | Moderate | | |
| 4 | 139463 | 61/TA | 4.1 ± 1 | 19/T | 2.8 ± 0.6 | $4.66 \times 10^{-1}$ | 0.0 | $9.49 \times 10^{-1}$ | 0.14 | $7.10 \times 10^{-1}$ | | Promoter variant | Moderate | | |
| 4 | 139505 | 48/T | 3.8 ± 1.2 | 32/C | 3.8 ± 0.9 | $9.97 \times 10^{-1}$ | 0.0 | $9.45 \times 10^{-1}$ | 0.22 | $6.43 \times 10^{-1}$ | | Promoter variant | Moderate | | |
| 4 | 139554 | 50/T | 4.1 ± 1.2 | 30/G | 3.2 ± 0.6 | $5.58 \times 10^{-1}$ | 0.2 | $6.27 \times 10^{-1}$ | 0.04 | $8.35 \times 10^{-1}$ | | Promoter variant | Moderate | | |
| 4 | 139556 | 43/A | 4.4 ± 1.4 | 37/C | 3.1 ± 0.5 | $3.94 \times 10^{-1}$ | 0.0 | $8.40 \times 10^{-1}$ | 0.02 | $8.88 \times 10^{-1}$ | | Promoter variant | Moderate | | |
| 4 | 139663 | 41/A | 4 ± 1.4 | 32/G | 3.8 ± 0.9 | $8.99 \times 10^{-1}$ | 0.0 | $8.73 \times 10^{-1}$ | 0.00 | $9.76 \times 10^{-1}$ | | Promoter variant | Moderate | | |
| 4 | 139665 | 43/C | 4.4 ± 1.4 | 37/T | 3 ± 0.5 | $3.58 \times 10^{-1}$ | 0.0 | $8.64 \times 10^{-1}$ | 0.09 | $7.67 \times 10^{-1}$ | | Promoter variant | Moderate | | |
| 4 | 139669 | 43/G | 4.5 ± 1.4 | 37/A | 3 ± 0.5 | $3.38 \times 10^{-1}$ | 0.7 | $4.15 \times 10^{-1}$ | 0.93 | $3.37 \times 10^{-1}$ | | Promoter variant | Moderate | | |
| 4 | 139696 | 13/A | 5.4 ± 1.9 | 67/G | 3.5 ± 0.9 | $3.47 \times 10^{-1}$ | 0.1 | $7.19 \times 10^{-1}$ | 0.22 | $6.40 \times 10^{-1}$ | | Promoter variant | Moderate | | |
| 4 | 139758 | 48/G | 4.2 ± 1.3 | 32/A | 3.1 ± 0.6 | $4.95 \times 10^{-1}$ | 0.0 | $9.53 \times 10^{-1}$ | 0.02 | $9.00 \times 10^{-1}$ | | Promoter variant | Moderate | | |
| 4 | 139846 | 48/G | 4.2 ± 1.3 | 32/A | 3.2 ± 0.6 | $5.49 \times 10^{-1}$ | 0.0 | $8.33 \times 10^{-1}$ | 0.00 | $9.66 \times 10^{-1}$ | | Promoter variant | Moderate | | |
| 4 | 139851 | 47/G | 4.2 ± 1.3 | 33/A | 3.2 ± 0.6 | $5.07 \times 10^{-1}$ | 0.2 | $6.22 \times 10^{-1}$ | 0.11 | $7.46 \times 10^{-1}$ | | Promoter variant | Moderate | | |
| 4 | 139866 | 15/T | 4.6 ± 1.6 | 65/C | 3.6 ± 0.9 | $6.31 \times 10^{-1}$ | 0.2 | $6.86 \times 10^{-1}$ | 0.01 | $9.29 \times 10^{-1}$ | | Promoter variant | Moderate | | |
| 4 | 139873 | 61/T | 3.8 ± 1 | 19/A | 3.8 ± 1.3 | $9.87 \times 10^{-1}$ | 0.3 | $5.80 \times 10^{-1}$ | 0.16 | $6.91 \times 10^{-1}$ | | Promoter variant | Moderate | | |
| 4 | 139899 | 61/A | 4.2 ± 1 | 19/G | 2.5 ± 0.4 | $3.49 \times 10^{-1}$ | 0.0 | $8.31 \times 10^{-1}$ | 0.03 | $8.73 \times 10^{-1}$ | | Promoter variant | Moderate | | |
| 4 | 139900 | 12/A | 5.2 ± 2.1 | 68/G | 3.5 ± 0.9 | $4.41 \times 10^{-1}$ | 0.0 | $8.31 \times 10^{-1}$ | 0.02 | $9.03 \times 10^{-1}$ | | Promoter variant | Moderate | | |
| 4 | 139915 | 53/A | 3.8 ± 1.1 | 27/G | 3.7 ± 1 | $9.65 \times 10^{-1}$ | 0.3 | $6.10 \times 10^{-1}$ | 0.04 | $8.51 \times 10^{-1}$ | | Promoter variant | Moderate | | |
| 4 | 139946 | 49/G | 4.5 ± 1.3 | 31/GA | 2.6 ± 0.5 | $2.21 \times 10^{-1}$ | 0.3 | $6.02 \times 10^{-1}$ | 0.12 | $7.26 \times 10^{-1}$ | | amino acid substitution (Ser3Leu) | Moderate | | |
| 4 | 139994 | 11/A | 5.2 ± 2.1 | 69/G | 3.6 ± 0.9 | $4.72 \times 10^{-1}$ | 0.3 | $5.65 \times 10^{-1}$ | 0.82 | $3.69 \times 10^{-1}$ | | amino acid substitution (Pro19Gln) | Moderate | | |
| 4 | 139999 | 30/G | 4.1 ± 0.9 | 50/C | 3.6 ± 1.2 | $7.30 \times 10^{-1}$ | 0.0 | $9.93 \times 10^{-1}$ | 0.10 | $7.55 \times 10^{-1}$ | | amino acid substitution (Ala21Thr) | Moderate | | |
| 4 | 140135 | 39/G | 4.1 ± 1.5 | 41/C | 3.4 ± 0.7 | $6.46 \times 10^{-1}$ | 0.0 | $8.43 \times 10^{-1}$ | 0.00 | $9.83 \times 10^{-1}$ | | amino acid substitution (Arg66Gln) | Moderate | | |
| 4 | 140235 | 1/AT | 13.1 ± 0 | 79/A | 3.7 ± 0.8 | | 1.26 | $2.65 \times 10^{-1}$ | 0.14 | $7.06 \times 10^{-1}$ | | frameshift variant (Phe100_Tyr101) | **High** | | |
| 4 | 140430 | 62/C | 3.9 ± 1 | 18/T | 3.4 ± 0.7 | $7.92 \times 10^{-1}$ | **3.8** | $\mathbf{5.44 \times 10^{-2}}$ | 2.42 | $1.24 \times 10^{-1}$ | ✓ | synonymous variant | Moderate | | |
| 4 | 140885 | 7/A | 4.3 ± 1.6 | 73/C | 3.7 ± 0.9 | $8.37 \times 10^{-1}$ | 1.3 | $2.61 \times 10^{-1}$ | 0.78 | $3.79 \times 10^{-1}$ | | amino acid substitution (Thr316Lys) | Moderate | | |
| 4 | 141073 | 62/A | 4.2 ± 1 | 18/T | 2.5 ± 0.5 | $3.60 \times 10^{-1}$ | 1.7 | $1.96 \times 10^{-1}$ | 1.56 | $2.16 \times 10^{-1}$ | | amino acid substitution (Thr348Ser) | Moderate | | |

| | | | | | | | | | | | | | | | | |
|---|---|---|---|---|---|---|---|---|---|---|---|---|---|---|---|---|
| 4 | 141340 | 72/G | 3.9 ± 0.9 | 8/T | 2.4 ± 0.6 | $5.50 \times 10^{-1}$ | 0.0 | $9.22 \times 10^{-1}$ | 0.07 | $7.99 \times 10^{-1}$ | | amino acid substitution (Asp407Glu) | Moderate | | | |
| 4 | 9295636 | 8/A | 14.2 ± 6.7 | 72/T | 2.6 ± 0.3 | **$1.41 \times 10^{-6}$** | 2.1 | $1.54 \times 10^{-1}$ | 2.02 | $1.60 \times 10^{-1}$ | | 5' UTR variant | Moderate | AT4G16490 | | ARM repeat superfamily protein |
| 4 | 9296072 | 8/A | 14.2 ± 6.7 | 72/G | 2.6 ± 0.3 | **$1.41 \times 10^{-6}$** | 0.0 | $9.99 \times 10^{-1}$ | 0.01 | $9.20 \times 10^{-1}$ | | 5' UTR variant | Moderate | | | |
| 4 | 9296171 | 70/C | 4 ± 0.9 | 10/T | 2.4 ± 0.8 | $4.87 \times 10^{-1}$ | 0.0 | $8.38 \times 10^{-1}$ | 0.01 | $9.04 \times 10^{-1}$ | | Promoter variant | Moderate | | | |
| 4 | 9296185 | 19/C | 5.6 ± 2.9 | 61/T | 3.2 ± 0.5 | $1.98 \times 10^{-1}$ | 0.1 | $7.05 \times 10^{-1}$ | 0.00 | $9.93 \times 10^{-1}$ | | Promoter variant | Moderate | | | |
| 4 | 9296213 | 57/A | 4.3 ± 1.1 | 23/AG | 2.4 ± 0.4 | $2.57 \times 10^{-1}$ | 2.3 | $1.37 \times 10^{-1}$ | 1.58 | $2.12 \times 10^{-1}$ | | Promoter variant | Moderate | | | |
| 4 | 9296256 | 22/C | 5.1 ± 2.5 | 58/A | 3.3 ± 0.6 | $3.05 \times 10^{-1}$ | 2.9 | **$9.24 \times 10^{-2}$** | **3.00** | **$8.75 \times 10^{-2}$** | | Promoter variant | Moderate | | | |
| 4 | 9297862 | 21/A | 5.2 ± 2.6 | 59/G | 3.3 ± 0.6 | $2.90 \times 10^{-1}$ | **11.0** | **$1.45 \times 10^{-3}$** | **10.49** | **$1.82 \times 10^{-3}$** | ✓ | Promoter variant | Moderate | | | |
| 4 | 13183267 | 52/A | 4.4 ± 1.2 | 28/G | 2.6 ± 0.5 | $2.74 \times 10^{-1}$ | 1.3 | $2.50 \times 10^{-1}$ | 1.38 | $2.45 \times 10^{-1}$ | | Promoter variant | Moderate | AT4G25970 | *PSD3* | Phosphatidylserine decarboxylase 3 |
| 4 | 13183279 | 64/C | 4.1 ± 1 | 16/G | 2.3 ± 0.4 | $3.47 \times 10^{-1}$ | 1.3 | $2.55 \times 10^{-1}$ | 0.88 | $3.50 \times 10^{-1}$ | | Promoter variant | Moderate | | | |
| 4 | 13183423 | 55/G | 4 ± 1.1 | 25/A | 3.3 ± 0.8 | $6.60 \times 10^{-1}$ | 1.5 | $2.31 \times 10^{-1}$ | 0.81 | $3.72 \times 10^{-1}$ | | Promoter variant | Moderate | | | |
| 4 | 13183444 | 61/G | 3.8 ± 1 | 19/A | 3.7 ± 0.9 | $9.78 \times 10^{-1}$ | 0.3 | $5.98 \times 10^{-1}$ | 1.14 | $2.90 \times 10^{-1}$ | | Promoter variant | Moderate | | | |
| 4 | 13183445 | 13/T | 4.7 ± 1.3 | 67/TA | 3.6 ± 0.9 | $5.93 \times 10^{-1}$ | 0.0 | $8.50 \times 10^{-1}$ | 0.11 | $7.38 \times 10^{-1}$ | | Promoter variant | Moderate | | | |
| 4 | 13183464 | 56/C | 4 ± 1.1 | 24/A | 3.3 ± 0.7 | $6.67 \times 10^{-1}$ | 0.3 | $6.06 \times 10^{-1}$ | 0.00 | $9.58 \times 10^{-1}$ | | Promoter variant | Moderate | | | |
| 4 | 13183542 | 12/G | 3.8 ± 1.2 | 68/GC | 3.8 ± 0.9 | $9.91 \times 10^{-1}$ | 0.8 | $4.46 \times 10^{-1}$ | 0.65 | $5.24 \times 10^{-1}$ | | Promoter variant | Moderate | | | |
| 4 | 13183560 | 42/T | 4.4 ± 1.4 | 38/C | 3.1 ± 0.6 | $4.11 \times 10^{-1}$ | 0.7 | $4.13 \times 10^{-1}$ | 0.32 | $5.73 \times 10^{-1}$ | | Promoter variant | Moderate | | | |
| 4 | 13183580 | 41/G | 4.4 ± 1.5 | 39/A | 3.1 ± 0.5 | $3.76 \times 10^{-1}$ | 1.9 | $1.76 \times 10^{-1}$ | 1.80 | $1.84 \times 10^{-1}$ | | Promoter variant | Moderate | | | |
| 4 | 13183585 | 47/A | 4.2 ± 1.3 | 33/G | 3.2 ± 0.6 | $5.14 \times 10^{-1}$ | 0.4 | $5.18 \times 10^{-1}$ | 0.41 | $5.25 \times 10^{-1}$ | | Promoter variant | Moderate | | | |
| 4 | 13183635 | 44/A | 4.3 ± 1.4 | 36/G | 3.1 ± 0.6 | $4.39 \times 10^{-1}$ | 0.3 | $5.94 \times 10^{-1}$ | 0.07 | $7.93 \times 10^{-1}$ | | Promoter variant | Moderate | | | |
| 4 | 13183644 | 47/T | 4.1 ± 1.3 | 33/G | 3.3 ± 0.6 | $5.85 \times 10^{-1}$ | 0.2 | $6.47 \times 10^{-1}$ | 0.03 | $8.68 \times 10^{-1}$ | | Promoter variant | Moderate | | | |
| 4 | 13183648 | 49/T | 4 ± 1.3 | 31/G | 3.4 ± 0.7 | $7.08 \times 10^{-1}$ | 0.5 | $4.83 \times 10^{-1}$ | 0.26 | $6.09 \times 10^{-1}$ | | Promoter variant | Moderate | | | |
| 4 | 13183710 | 43/C | 4.3 ± 1.4 | 37/G | 3.1 ± 0.6 | $4.40 \times 10^{-1}$ | 0.0 | $8.61 \times 10^{-1}$ | 0.00 | $9.93 \times 10^{-1}$ | | Promoter variant | Moderate | | | |
| 4 | 13183740 | 62/C | 4.2 ± 1 | 18/T | 2.2 ± 0.3 | $2.83 \times 10^{-1}$ | 0.2 | $6.60 \times 10^{-1}$ | 0.00 | $9.45 \times 10^{-1}$ | | Promoter variant | Moderate | | | |
| 4 | 13183803 | 65/C | 4 ± 1 | 15/A | 2.7 ± 0.6 | $4.99 \times 10^{-1}$ | 0.3 | $5.70 \times 10^{-1}$ | 0.25 | $6.21 \times 10^{-1}$ | | Promoter variant | Moderate | | | |
| 4 | 13183940 | 62/G | 4.1 ± 1 | 18/A | 2.7 ± 0.6 | $4.59 \times 10^{-1}$ | 0.0 | $8.69 \times 10^{-1}$ | 0.17 | $6.83 \times 10^{-1}$ | | Promoter variant | Moderate | | | |
| 4 | 13184006 | 61/G | 4.1 ± 1 | 19/A | 2.6 ± 0.5 | $4.04 \times 10^{-1}$ | 0.1 | $7.51 \times 10^{-1}$ | 0.01 | $9.11 \times 10^{-1}$ | | Promoter variant | Moderate | | | |
| 4 | 13184036 | 4/A | 9.2 ± 5.6 | 76/G | 3.5 ± 0.8 | $1.08 \times 10^{-1}$ | 2.2 | $1.44 \times 10^{-1}$ | 0.97 | $3.28 \times 10^{-1}$ | | Promoter variant | Moderate | | | |
| 4 | 13184088 | 30/C | 5.3 ± 2 | 50/A | 2.9 ± 0.5 | $1.23 \times 10^{-1}$ | 0.1 | $8.10 \times 10^{-1}$ | 0.01 | $9.06 \times 10^{-1}$ | | Promoter variant | Moderate | | | |
| 4 | 13184147 | 55/A | 4.3 ± 1.1 | 25/T | 2.7 ± 0.6 | $3.57 \times 10^{-1}$ | 0.1 | $7.38 \times 10^{-1}$ | 0.24 | $6.26 \times 10^{-1}$ | | 5'-UTR variant | Moderate | | | |
| 4 | 13184162 | 57/T | 4.1 ± 1.1 | 23/G | 2.9 ± 0.6 | $4.66 \times 10^{-1}$ | 0.1 | $8.16 \times 10^{-1}$ | 0.00 | $9.61 \times 10^{-1}$ | | 5'-UTR variant | Moderate | | | |
| 4 | 13184191 | 71/A | 4 ± 0.9 | 9/C | 2 ± 0.3 | $4.04 \times 10^{-1}$ | 0.0 | $8.32 \times 10^{-1}$ | 0.26 | $6.15 \times 10^{-1}$ | | 5'-UTR variant | Moderate | | | |
| 4 | 13184321 | 72/C | 3.9 ± 0.9 | 8/T | 2.9 ± 0.4 | $7.10 \times 10^{-1}$ | 0.0 | $8.82 \times 10^{-1}$ | 0.02 | $8.98 \times 10^{-1}$ | | amino acid substitution (Arg28Ser) | Moderate | | | |
| 4 | 13185020 | 72/G | 3.8 ± 0.9 | 8/C | 3.6 ± 1.4 | $9.45 \times 10^{-1}$ | 0.5 | $4.64 \times 10^{-1}$ | 0.58 | $4.47 \times 10^{-1}$ | | amino acid substitution (Gln140Leu) | Moderate | | | |
| 4 | 13185332 | 71/A | 3.8 ± 0.9 | 9/T | 3.4 ± 1.3 | $8.45 \times 10^{-1}$ | 1.5 | $2.31 \times 10^{-1}$ | 1.45 | $2.32 \times 10^{-1}$ | | amino acid substitution (Ser162Ile) | Moderate | | | |
| 4 | 13186139 | 49/G | 4.7 ± 1.3 | 31/GACA | 2.3 ± 0.3 | $1.25 \times 10^{-1}$ | **15.6** | **$1.81 \times 10^{-4}$** | **11.02** | **$1.41 \times 10^{-3}$** | ✓ | intron | Moderate | | | |
| 4 | 13187817 | 71/T | 3.9 ± 0.9 | 9/C | 2.5 ± 0.5 | $5.65 \times 10^{-1}$ | **15.6** | **$1.81 \times 10^{-4}$** | **11.02** | **$1.41 \times 10^{-3}$** | ✓ | synonymous variant | Moderate | | | |
| 4 | 13187825 | 18/T | 8.2 ± 3.2 | 62/C | 2.5 ± 0.3 | **$1.57 \times 10^{-3}$** | 0.1 | $7.43 \times 10^{-1}$ | 0.13 | $7.19 \times 10^{-1}$ | | amino acid substitution (Ser450Leu) | Moderate | | | |

| | | | | | | | | | | | | | | | | |
|---|---|---|---|---|---|---|---|---|---|---|---|---|---|---|---|---|
| 5 | 3793170 | 57/C | 4.2 ± 1.1 | 23/A | 2.7 ± 0.6 | $3.94 \times 10^{-1}$ | 1.7 | $1.96 \times 10^{-1}$ | 0.88 | $3.52 \times 10^{-1}$ | | Promoter variant | Moderate | AT5G11780 | | SNF2 domain protein |
| 5 | 3793239 | 41/T | 4.9 ± 1.5 | 39/C | 2.5 ± 0.4 | $1.20 \times 10^{-1}$ | 1.8 | $1.80 \times 10^{-1}$ | 0.70 | $4.04 \times 10^{-1}$ | | Promoter variant | Moderate | | | |
| 5 | 3793331 | 36/A | 4.7 ± 1.6 | 44/G | 3 ± 0.7 | $2.82 \times 10^{-1}$ | 0.2 | $6.40 \times 10^{-1}$ | 0.06 | $8.08 \times 10^{-1}$ | | Promoter variant | Moderate | | | |
| 5 | 3793369 | 59/A | 4.1 ± 1.1 | 21/C | 3 ± 0.7 | $5.48 \times 10^{-1}$ | 0.6 | $4.58 \times 10^{-1}$ | 0.09 | $7.67 \times 10^{-1}$ | | Promoter variant | Moderate | | | |
| 5 | 3793370 | 47/C | 4.1 ± 1.2 | 33/T | 3.4 ± 0.8 | $6.48 \times 10^{-1}$ | 0.1 | $7.46 \times 10^{-1}$ | 0.01 | $9.13 \times 10^{-1}$ | | Promoter variant | Moderate | | | |
| 5 | 3793388 | 54/G | 4 ± 1.1 | 26/T | 3.3 ± 1 | $6.79 \times 10^{-1}$ | 0.0 | $9.37 \times 10^{-1}$ | 0.03 | $8.53 \times 10^{-1}$ | | Promoter variant | Moderate | | | |
| 5 | 3793434 | 37/A | 4.7 ± 1.5 | 43/G | 3 ± 0.7 | $2.76 \times 10^{-1}$ | 0.6 | $4.59 \times 10^{-1}$ | 0.15 | $7.03 \times 10^{-1}$ | | Promoter variant | Moderate | | | |
| 5 | 3793499 | 60/A | 4.1 ± 1.1 | 20/AC | 2.9 ± 0.7 | $4.93 \times 10^{-1}$ | 0.3 | $5.87 \times 10^{-1}$ | 0.11 | $7.44 \times 10^{-1}$ | | Promoter variant | Moderate | | | |
| 5 | 3793595 | 66/T | 3.9 ± 1 | 14/A | 3.2 ± 1 | $7.42 \times 10^{-1}$ | 0.2 | $6.37 \times 10^{-1}$ | 0.50 | $4.82 \times 10^{-1}$ | | Promoter variant | Moderate | | | |
| 5 | 3794673 | 63/A | 4.2 ± 1 | 17/T | 2.3 ± 0.5 | $3.09 \times 10^{-1}$ | 1.7 | $2.03 \times 10^{-1}$ | 1.74 | $1.91 \times 10^{-1}$ | | amino acid substitution (His249Leu) | Moderate | | | |
| 5 | 3794682 | 60/T | 4 ± 1 | 20/A | 3.2 ± 0.8 | $6.84 \times 10^{-1}$ | 0.6 | $4.40 \times 10^{-1}$ | 0.43 | $5.16 \times 10^{-1}$ | | amino acid substitution (Phe252Tyr) | Moderate | | | |
| 5 | 3795628 | 49/T | 4.5 ± 1.3 | 31/A | 2.7 ± 0.5 | $2.60 \times 10^{-1}$ | 0.9 | $3.48 \times 10^{-1}$ | 0.71 | $4.01 \times 10^{-1}$ | | amino acid substitution (Leu387Gln) | Moderate | | | |
| 5 | 3795643 | 48/C | 4.5 ± 1.3 | 32/T | 2.7 ± 0.5 | $2.38 \times 10^{-1}$ | 1.1 | $3.06 \times 10^{-1}$ | 1.01 | $3.17 \times 10^{-1}$ | | amino acid substitution (Pro392Leu) | Moderate | | | |
| 5 | 3795741 | 48/G | 4.5 ± 1.3 | 32/C | 2.7 ± 0.5 | $2.38 \times 10^{-1}$ | 1.1 | $3.06 \times 10^{-1}$ | 1.01 | $3.17 \times 10^{-1}$ | | amino acid substitution (Glu425Gln) | Moderate | | | |
| 5 | 3796248 | 19/A | 7.7 ± 3 | 61/T | 2.6 ± 0.3 | **$3.87 \times 10^{-3}$** | **7.9** | **$6.44 \times 10^{-3}$** | **5.99** | **$1.68 \times 10^{-2}$** | ✓ | intergenic space variant | Moderate | | | |
| 5 | 3808018 | 65/C | 4.1 ± 1 | 15/G | 2.3 ± 0.5 | $3.70 \times 10^{-1}$ | 0.7 | $4.05 \times 10^{-1}$ | 0.39 | $5.35 \times 10^{-1}$ | | amino acid substitution (Ala18Pro) | Moderate | AT5G11800 | *KEA6* | $K^+$ efflux antiporter 6 |
| 5 | 3808248 | 73/C | 3.8 ± 0.9 | 7/CT | 3.3 ± 1.7 | $8.37 \times 10^{-1}$ | 0.1 | $7.46 \times 10^{-1}$ | 0.00 | $9.71 \times 10^{-1}$ | | Promoter variant | Moderate | | | |
| 5 | 3808963 | 73/C | 3.8 ± 0.9 | 7/G | 3.2 ± 1.1 | $8.25 \times 10^{-1}$ | 0.4 | $5.40 \times 10^{-1}$ | 0.48 | $4.89 \times 10^{-1}$ | | Promoter variant | Moderate | | | |

The results of a local association study (LAS) of the variation in relative fold-change (RFC) in expression levels of *LKR* in 80 *A. thaliana* ecotypes in a second drought (D2) versus the a drought (D1) separated by a recovery interval, calculated in Tassel separately for the individual genomic regions are presented. The polymorphisms in coding sequences and upstream regulatory regions (5'-UTR and promoters within 1 kb from the start codon) of genes directly associated with the top most significant ($P < 10^{-4}$) single nucleotide polymorphisms (SNPs) detected by a genome-wide association study (GWAS) (see Table 1) were downloaded from the 1001 genomes database (https://tools.1001genomes.org/polymorph/). Data of only 80 out of the 115 ecotypes used for the expression GWAS (see Figure 1, Table 1) were available in the 1001 genomes database. The frequency of the tolerant and sensitive alleles among the 80 ecotypes is shown with the average RFC. The *F*-statistic and *P*-values of the generalized linear model (GLM) and the mixed linear model (MLM) methods of LAS calculations are shown. The effects of the GWAS-detected SNPs and additional polymorphisms in each gene mined from the 1001 genomes database are shown in terms of amino acid substitutions, frameshift, promoter, or 5'-UTR variations.

### Table S8 Local association study of superinduction of *HIS1-3* in 81 *Arabidopsis thaliana* ecotypes under recurrent drought

| Chromosome | Physical position | Frequency of high expression allele | Average RFC high expression allele | Frequency of low-expression allele | Average RFC low-expressionallele | Student's *t*-test *P*-value | LAS GLM *F*-statistic | LAS GLM *P*-value | LAS MLM *F*-statistic | LAS MLM *P*-value | GWAS-detected SNP | Polymorphism effects | Impact | Gene ID | Gene symbol | Short description |
|---|---|---|---|---|---|---|---|---|---|---|---|---|---|---|---|---|
| 1 | 132430 | 74/T | 3 ± 0.5 | 7/C | 3 ± 1.1 | $9.99 \times 10^{-1}$ | 0.1 | $7.89 \times 10^{-1}$ | 0.9 | $3.37 \times 10^{-1}$ | | amino acid substitution (His702Arg) | Moderate | AT1G01340 | *ACBK1, CNGC10* | Cyclic nucleotide gated channel 10 |
| 1 | 133144 | 4/T | 9.7 ± 4.8 | 77/G | 2.7 ± 0.4 | **$1.51 \times 10^{-4}$** | **15.9** | **$1.55 \times 10^{-4}$** | 2 | $1.60 \times 10^{-1}$ | ✓ | synonymous variant | Moderate | | | |
| 1 | 135402 | 63/A | 3.2 ± 0.6 | 18/G | 2.6 ± 0.6 | $6.05 \times 10^{-1}$ | 0 | $9.33 \times 10^{-1}$ | 0.2 | $6.86 \times 10^{-1}$ | | 5'-UTR intron variant | Moderate | | | |
| 1 | 135493 | 66/G | 3.1 ± 0.5 | 15/T | 2.9 ± 0.7 | $8.49 \times 10^{-1}$ | 0 | $8.71 \times 10^{-1}$ | 0.5 | $4.72 \times 10^{-1}$ | | 5'-UTR intron variant | Moderate | | | |
| 1 | 135500 | 66/C | 3.2 ± 0.5 | 15/T | 2.5 ± 0.6 | $5.67 \times 10^{-1}$ | 0 | $8.45 \times 10^{-1}$ | 0 | $8.33 \times 10^{-1}$ | | 5'-UTR intron variant | Moderate | | | |
| 1 | 135519 | 66/A | 3.2 ± 0.5 | 15/G | 2.5 ± 0.6 | $5.67 \times 10^{-1}$ | 0 | $8.45 \times 10^{-1}$ | 0 | $8.33 \times 10^{-1}$ | | 5'-UTR intron variant | Moderate | | | |
| 1 | 135587 | 42/C | 3.7 ± 0.7 | 39/A | 2.3 ± 0.5 | $1.08 \times 10^{-1}$ | 1.9 | $1.75 \times 10^{-1}$ | 0.2 | $6.36 \times 10^{-1}$ | | 5'-UTR intron variant | Moderate | | | |
| 1 | 136043 | 68/T | 3.1 ± 0.5 | 13/TG | 2.8 ± 0.8 | $7.88 \times 10^{-1}$ | 0.1 | $7.91 \times 10^{-1}$ | 0.2 | $6.90 \times 10^{-1}$ | | Promoter variant | Moderate | | | |
| 1 | 136188 | 62/T | 3.1 ± 0.6 | 19/C | 3 ± 0.7 | $9.24 \times 10^{-1}$ | 0 | $9.54 \times 10^{-1}$ | 0.8 | $3.70 \times 10^{-1}$ | | 5'-UTR variant | Moderate | | | |
| 1 | 136211 | 54/T | 3.1 ± 0.6 | 27/A | 2.9 ± 0.5 | $7.62 \times 10^{-1}$ | 0 | $8.77 \times 10^{-1}$ | 0.3 | $5.74 \times 10^{-1}$ | | 5'-UTR variant | Moderate | | | |
| 1 | 4859326 | 7/C | 7.8 ± 3.1 | 74/A | 2.6 ± 0.4 | **$2.53 \times 10^{-4}$** | **14.7** | **$2.66 \times 10^{-4}$** | **3.5** | $6.68 \times 10^{-2}$ | ✓ | synonymous variant | Moderate | AT1G14220 | | Ribonuclease T2 family protein |
| 1 | 4859930 | 66/A | 3.2 ± 0.5 | 15/G | 2.4 ± 0.9 | $4.39 \times 10^{-1}$ | 0.7 | $3.93 \times 10^{-1}$ | 0.7 | $3.99 \times 10^{-1}$ | | Promoter variant | Moderate | | | |
| 1 | 4860000 | 67/G | 3.2 ± 0.5 | 14/C | 2.5 ± 0.9 | $5.28 \times 10^{-1}$ | 0.5 | $4.96 \times 10^{-1}$ | 0.4 | $5.12 \times 10^{-1}$ | | Promoter variant | Moderate | | | |
| 1 | 4860332 | 70/G | 3.1 ± 0.5 | 11/A | 2.5 ± 0.8 | $5.76 \times 10^{-1}$ | 0.9 | $3.39 \times 10^{-1}$ | 0.6 | $4.40 \times 10^{-1}$ | | Promoter variant | Moderate | | | |
| 1 | 4860367 | 67/C | 3.2 ± 0.5 | 14/T | 2.4 ± 0.9 | $5.08 \times 10^{-1}$ | 0.7 | $4.10 \times 10^{-1}$ | 0.7 | $4.21 \times 10^{-1}$ | | Promoter variant | Moderate | | | |
| 1 | 4860388 | 66/A | 3.2 ± 0.5 | 14/G | 2.3 ± 0.7 | $3.92 \times 10^{-1}$ | 1.1 | $3.47 \times 10^{-1}$ | 0.7 | $5.05 \times 10^{-1}$ | | Promoter variant | Moderate | | | |
| 1 | 4860488 | 29/A | 3.3 ± 0.9 | 52/G | 2.9 ± 0.5 | $6.93 \times 10^{-1}$ | 0.1 | $7.05 \times 10^{-1}$ | 0.1 | $7.34 \times 10^{-1}$ | | Promoter variant | Moderate | | | |
| 1 | 4860493 | 68/C | 3.1 ± 0.5 | 13/T | 2.5 ± 1 | $5.70 \times 10^{-1}$ | 0.3 | $5.90 \times 10^{-1}$ | 0.2 | $6.88 \times 10^{-1}$ | | Promoter variant | Moderate | | | |
| 1 | 4860528 | 66/G | 3.3 ± 0.5 | 15/A | 2 ± 0.6 | $2.25 \times 10^{-1}$ | 2.5 | $1.16 \times 10^{-1}$ | 1.7 | $1.92 \times 10^{-1}$ | | missense variant | Moderate | AT1G14220, | | Ribonuclease T2 family protein; GDA1/CD39 nucleoside phosphatase family protein |
| 1 | 4860539 | 65/G | 3.2 ± 0.5 | 16/A | 2.2 ± 0.8 | $3.32 \times 10^{-1}$ | 1.1 | $3.09 \times 10^{-1}$ | 1 | $3.25 \times 10^{-1}$ | | synonymous variant | Moderate | AT1G14230 | | |
| 1 | 4860618 | 9/A | 3.9 ± 0.9 | 72/G | 2.9 ± 0.5 | $4.86 \times 10^{-1}$ | 0.3 | $6.16 \times 10^{-1}$ | 1.3 | $2.59 \times 10^{-1}$ | | missense variant | Moderate | AT1G14230 | | GDA1/CD39 nucleoside phosphatase family protein |
| 1 | 4860778 | 65/A | 3.2 ± 0.5 | 16/T | 2.6 ± 0.8 | $5.69 \times 10^{-1}$ | 0.5 | $4.67 \times 10^{-1}$ | 0.5 | $4.84 \times 10^{-1}$ | | Promoter variant | Moderate | | | |
| 1 | 4860883 | 29/C | 3.3 ± 0.6 | 52/G | 2.9 ± 0.6 | $6.27 \times 10^{-1}$ | 0 | $9.44 \times 10^{-1}$ | 0.5 | $4.86 \times 10^{-1}$ | | Promoter variant | Moderate | | | |
| 1 | 4861125 | 9/T | 3.9 ± 0.9 | 72/A | 2.9 ± 0.5 | $4.86 \times 10^{-1}$ | 0.3 | $6.16 \times 10^{-1}$ | 1.3 | $2.59 \times 10^{-1}$ | | Promoter variant | Moderate | | | |
| 1 | 4861444 | 66/G | 3.3 ± 0.5 | 15/A | 2 ± 0.6 | $2.25 \times 10^{-1}$ | 2.5 | $1.16 \times 10^{-1}$ | 1.7 | $1.92 \times 10^{-1}$ | | 5'-UTR variant | Moderate | | | |
| 1 | 4861445 | 67/A | 3.3 ± 0.5 | 14/G | 2 ± 0.7 | $2.73 \times 10^{-1}$ | 2.3 | $1.35 \times 10^{-1}$ | 1.2 | $2.85 \times 10^{-1}$ | | 5'-UTR variant | Moderate | | | |
| 1 | 4861689 | 24/G | 3.1 ± 0.8 | 57/T | 3 ± 0.6 | $8.93 \times 10^{-1}$ | 0.1 | $7.03 \times 10^{-1}$ | 0.7 | $4.08 \times 10^{-1}$ | | amino acid substitution (Gly65Cys) | Moderate | | | |
| 1 | 4862355 | 66/G | 3.1 ± 0.5 | 15/C | 2.7 ± 0.9 | $6.70 \times 10^{-1}$ | 0.3 | $5.79 \times 10^{-1}$ | 0.3 | $6.10 \times 10^{-1}$ | | amino acid substitution (Gly207Ala) | Moderate | | | |
| 1 | 4864204 | 66/G | 3.2 ± 0.5 | 15/C | 2.3 ± 0.8 | $4.03 \times 10^{-1}$ | 1 | $3.22 \times 10^{-1}$ | 1.2 | $2.85 \times 10^{-1}$ | | Promoter variant | Moderate | AT1G14240 | *APY3* | GDA1/CD39 nucleoside phosphatase family protein |
| 1 | 4864241 | 31/TGA | 4 ± 0.9 | 50/T | 2.5 ± 0.5 | $7.62 \times 10^{-2}$ | 1.6 | $2.06 \times 10^{-1}$ | 0 | $8.50 \times 10^{-1}$ | | Promoter variant | Moderate | | | |

| | | | | | | | | | | | | | | | | |
|---|---|---|---|---|---|---|---|---|---|---|---|---|---|---|---|---|
| 1 | 4864345 | 9/G | 3.9 ± 0.9 | 72/A | 2.9 ± 0.5 | $4.86 \times 10^{-1}$ | 0.3 | $6.16 \times 10^{-1}$ | 1.3 | $2.59 \times 10^{-1}$ | | Promoter variant | Moderate | | | |
| 1 | 4864468 | 13/G | 3.4 ± 1.8 | 68/GT | 3 ± 0.4 | $7.35 \times 10^{-1}$ | 0.1 | $8.09 \times 10^{-1}$ | 1.5 | $2.22 \times 10^{-1}$ | | Promoter variant | Moderate | | | |
| 1 | 4864474 | 12/A | 3.1 ± 1.1 | 69/T | 3 ± 0.5 | $9.76 \times 10^{-1}$ | 0 | $9.92 \times 10^{-1}$ | 0 | $8.68 \times 10^{-1}$ | | Promoter variant | Moderate | | | |
| 1 | 4864475 | 13/A | 4 ± 1.8 | 68/T | 2.9 ± 0.4 | $3.25 \times 10^{-1}$ | 1.2 | $2.74 \times 10^{-1}$ | 0 | $9.06 \times 10^{-1}$ | | Promoter variant | Moderate | | | |
| 1 | 4864719 | 29/A | 3.2 ± 0.9 | 52/G | 2.9 ± 0.5 | $7.65 \times 10^{-1}$ | 0.1 | $7.48 \times 10^{-1}$ | 0.2 | $6.31 \times 10^{-1}$ | | Promoter variant | Moderate | | | |
| 1 | 4864724 | 27/G | 3.3 ± 1 | 54/C | 2.9 ± 0.5 | $6.57 \times 10^{-1}$ | 0.2 | $6.98 \times 10^{-1}$ | 0.1 | $7.51 \times 10^{-1}$ | | Promoter variant | Moderate | | | |
| 1 | 4864729 | 72/GT | 3.1 ± 0.5 | 9/G | 2.6 ± 1 | $6.94 \times 10^{-1}$ | 0.7 | $4.03 \times 10^{-1}$ | 0.4 | $5.34 \times 10^{-1}$ | | Promoter variant | Moderate | | | |
| 1 | 4864806 | 30/T | 3.1 ± 0.9 | 51/A | 3 ± 0.5 | $8.83 \times 10^{-1}$ | 0 | $8.51 \times 10^{-1}$ | 0.4 | $5.37 \times 10^{-1}$ | | Promoter variant | Moderate | | | |
| 1 | 4864840 | 26/C | 3.4 ± 1 | 55/A | 2.9 ± 0.5 | $5.43 \times 10^{-1}$ | 0.3 | $5.65 \times 10^{-1}$ | 0.1 | $7.88 \times 10^{-1}$ | | Promoter variant | Moderate | | | |
| 1 | 4864935 | 71/A | 3.1 ± 0.5 | 10/T | 2.5 ± 1.3 | $6.06 \times 10^{-1}$ | 0 | $9.04 \times 10^{-1}$ | 0.1 | $7.98 \times 10^{-1}$ | | 5'-UTR variant | Moderate | | | |
| 1 | 4864981 | 57/G | 3.2 ± 0.6 | 24/T | 2.8 ± 0.7 | $6.72 \times 10^{-1}$ | 0.3 | $5.59 \times 10^{-1}$ | 0 | $8.64 \times 10^{-1}$ | | 5'-UTR intron variant | Moderate | | | |
| 1 | 4865091 | 3/C | 3.1 ± 1.8 | 78/T | 3 ± 0.5 | $9.94 \times 10^{-1}$ | 0.1 | $8.10 \times 10^{-1}$ | 0.1 | $7.10 \times 10^{-1}$ | | 5'-UTR intron variant | Moderate | | | |
| 1 | 4865117 | 55/C | 3.2 ± 0.6 | 26/T | 2.7 ± 0.6 | $6.16 \times 10^{-1}$ | 0.7 | $4.17 \times 10^{-1}$ | 0.2 | $6.54 \times 10^{-1}$ | | 5'-UTR intron variant | Moderate | | | |
| 1 | 4865177 | 67/G | 3.1 ± 0.5 | 14/T | 2.6 ± 0.9 | $6.46 \times 10^{-1}$ | 0.3 | $5.94 \times 10^{-1}$ | 0.3 | $6.18 \times 10^{-1}$ | | amino acid substitution (Ala7Ser) | Moderate | | | |
| 1 | 4865501 | 48/G | 3.1 ± 0.6 | 33/T | 3 ± 0.6 | $8.82 \times 10^{-1}$ | 0 | $9.84 \times 10^{-1}$ | 0.2 | $6.39 \times 10^{-1}$ | | amino acid substitution (Ser115Ala) | Moderate | | | |
| 1 | 4865721 | 22/A | 3.2 ± 0.7 | 59/C | 3 ± 0.6 | $7.85 \times 10^{-1}$ | 0.2 | $6.85 \times 10^{-1}$ | 0.6 | $4.38 \times 10^{-1}$ | | amino acid substitution (Asn188Thr) | Moderate | | | |
| 2 | 7707454 | 12/A | 6.9 ± 1.8 | 69/G | 2.4 ± 0.4 | **$5.31 \times 10^{-5}$** | **24.4** | **$4.80 \times 10^{-6}$** | **12.8** | **$6.27 \times 10^{-4}$** | ✓ | synonymous variant | Moderate | AT2G17740 | *VLG* | Vacuoleless gametophytes |
| 2 | 7707930 | 11/C | 4.2 ± 1.2 | 70/A | 2.9 ± 0.5 | $2.71 \times 10^{-1}$ | 1.4 | $2.46 \times 10^{-1}$ | 2.7 | $1.07 \times 10^{-1}$ | | Promoter variant | Moderate | | | |
| 2 | 7708164 | 8/GA | 4.2 ± 2.8 | 3/GAAA | 0.7 ± 0.3 | $4.78 \times 10^{-1}$ | 1.1 | $3.55 \times 10^{-1}$ | 0.9 | $4.28 \times 10^{-1}$ | | Promoter variant | Moderate | | | |
| 2 | 7708290 | 25/G | 3.4 ± 1 | 56/A | 2.9 ± 0.5 | $5.35 \times 10^{-1}$ | 0.2 | $6.52 \times 10^{-1}$ | 0.6 | $4.59 \times 10^{-1}$ | | Promoter variant | Moderate | | | |
| 2 | 7708360 | 47/T | 3.1 ± 0.7 | 34/A | 3 ± 0.6 | $9.32 \times 10^{-1}$ | 0 | $9.30 \times 10^{-1}$ | 0.4 | $5.43 \times 10^{-1}$ | | Promoter variant | Moderate | | | |
| 2 | 7708513 | 42/T | 3.3 ± 0.7 | 39/C | 2.7 ± 0.5 | $5.00 \times 10^{-1}$ | 0.6 | $4.53 \times 10^{-1}$ | 0.2 | $6.21 \times 10^{-1}$ | | Promoter variant | Moderate | | | |
| 2 | 7708584 | 33/T | 3.3 ± 0.8 | 48/C | 2.9 ± 0.5 | $6.52 \times 10^{-1}$ | 0.1 | $7.91 \times 10^{-1}$ | 0.4 | $5.20 \times 10^{-1}$ | | Promoter variant | Moderate | | | |
| 2 | 7708597 | 10/T | 4 ± 1.4 | 71/C | 2.9 ± 0.5 | $3.79 \times 10^{-1}$ | 0.4 | $5.20 \times 10^{-1}$ | 1.2 | $2.73 \times 10^{-1}$ | | Promoter variant | Moderate | | | |
| 2 | 10616314 | 57/C | 3.3 ± 0.6 | 24/G | 2.3 ± 0.5 | $2.72 \times 10^{-1}$ | 1.4 | $2.43 \times 10^{-1}$ | 0.2 | $6.81 \times 10^{-1}$ | | Promoter variant | Moderate | AT2G24960 | | Myb/SANT-like DNA-binding domain protein |
| 2 | 10616329 | 25/C | 3.6 ± 1 | 56/G | 2.8 ± 0.5 | $3.52 \times 10^{-1}$ | 0.8 | $3.82 \times 10^{-1}$ | 0.7 | $4.05 \times 10^{-1}$ | | Promoter variant | Moderate | | | |
| 2 | 10616370 | 22/G | 3.8 ± 1.1 | 59/T | 2.7 ± 0.5 | $2.45 \times 10^{-1}$ | 1.6 | $2.08 \times 10^{-1}$ | 1.3 | $2.66 \times 10^{-1}$ | | Promoter variant | Moderate | | | |
| 2 | 10616371 | 20/T | 4.1 ± 1.2 | 61/A | 2.7 ± 0.4 | $1.52 \times 10^{-1}$ | 2.6 | $1.08 \times 10^{-1}$ | 2.5 | $1.17 \times 10^{-1}$ | | Promoter variant | Moderate | | | |
| 2 | 10616399 | 25/T | 3.9 ± 1 | 56/A | 2.6 ± 0.5 | $1.53 \times 10^{-1}$ | 2.3 | $1.31 \times 10^{-1}$ | 2.3 | $1.34 \times 10^{-1}$ | | Promoter variant | Moderate | | | |
| 2 | 10616406 | 27/G | 3.7 ± 1 | 54/T | 2.7 ± 0.5 | $2.58 \times 10^{-1}$ | 1.5 | $2.26 \times 10^{-1}$ | 1.3 | $2.56 \times 10^{-1}$ | | Promoter variant | Moderate | | | |
| 2 | 10616427 | 16/CT | 4 ± 0.8 | 65/C | 2.8 ± 0.5 | $2.53 \times 10^{-1}$ | 1 | $3.26 \times 10^{-1}$ | **4.7** | **$3.29 \times 10^{-2}$** | | Promoter variant | Moderate | | | |
| 2 | 10616495 | 29/C | 3.5 ± 0.9 | 52/T | 2.8 ± 0.5 | $4.22 \times 10^{-1}$ | 0.7 | $4.20 \times 10^{-1}$ | 0.6 | $4.44 \times 10^{-1}$ | | Promoter variant | Moderate | | | |
| 2 | 10616514 | 57/T | 3 ± 0.5 | 24/A | 3 ± 1 | $9.99 \times 10^{-1}$ | 0 | $9.63 \times 10^{-1}$ | 0.2 | $6.77 \times 10^{-1}$ | | Promoter variant | Moderate | | | |
| 2 | 10616948 | 68/T | 3.4 ± 0.5 | 13/G | 1.2 ± 0.4 | **$4.98 \times 10^{-2}$** | **4.3** | **$4.23 \times 10^{-2}$** | **5.3** | **$2.38 \times 10^{-2}$** | | Promoter variant | Moderate | | | |
| 2 | 10617051 | 71/A | 3.1 ± 0.5 | 10/C | 2.8 ± 1 | $8.13 \times 10^{-1}$ | 0.2 | $6.26 \times 10^{-1}$ | 0.2 | $6.81 \times 10^{-1}$ | | Promoter variant | Moderate | | | |
| 2 | 10617053 | 14/G | 4.2 ± 1.7 | 67/C | 2.8 ± 0.4 | $2.16 \times 10^{-1}$ | 1.3 | $2.51 \times 10^{-1}$ | 0 | $9.40 \times 10^{-1}$ | | Promoter variant | Moderate | | | |
| 2 | 10618462 | 10/T | 3.9 ± 1.3 | 71/C | 2.9 ± 0.5 | $4.49 \times 10^{-1}$ | 0.3 | $5.99 \times 10^{-1}$ | 0.7 | $4.20 \times 10^{-1}$ | | amino acid substitution (Ile325Thr) | Moderate | | | |

| | | | | | | | | | | | | | | | | |
|---|---|---|---|---|---|---|---|---|---|---|---|---|---|---|---|---|
| 2 | 10618783 | 55/T | 3.3 ± 0.6 | 26/C | 2.6 ± 0.6 | $4.60 \times 10^{-1}$ | 1.2 | $2.80 \times 10^{-1}$ | 0.7 | $4.15 \times 10^{-1}$ | | amino acid substitution (Ile432Thr) | Moderate | | | |
| 2 | 10619291 | 7/T | 6.6 ± 3 | 74/C | 2.7 ± 0.4 | **$8.14 \times 10^{-3}$** | **7** | **$1.00 \times 10^{-2}$** | 1 | $3.16 \times 10^{-1}$ | ✓ | synonymous variant | Moderate | | | |
| 2 | 10619519 | 7/T | 6.6 ± 3 | 74/A | 2.7 ± 0.4 | **$8.14 \times 10^{-3}$** | **7** | **$1.00 \times 10^{-2}$** | 1 | $3.16 \times 10^{-1}$ | ✓ | intron variant | Moderate | | | |
| 2 | 11817721 | 54/A | 3.1 ± 0.6 | 27/G | 3 ± 0.5 | $9.05 \times 10^{-1}$ | 0.1 | $8.17 \times 10^{-1}$ | 0.2 | $6.83 \times 10^{-1}$ | | Promoter variant | Moderate | AT2G27720 | *P2Z, RPP2A* | Ribosomal protein |
| 2 | 11817899 | 5/T | 3.9 ± 2.4 | 75/A | 2.9 ± 0.5 | $5.87 \times 10^{-1}$ | 1.2 | $3.00 \times 10^{-1}$ | 2.4 | $9.52 \times 10^{-2}$ | | Promoter variant | Moderate | | | |
| 2 | 11818261 | 39/T | 3.3 ± 0.6 | 42/G | 2.8 ± 0.7 | $5.41 \times 10^{-1}$ | 0.2 | $6.32 \times 10^{-1}$ | **3** | $8.51 \times 10^{-2}$ | | Promoter variant | Moderate | | | |
| 2 | 11818599 | 69/C | 3.1 ± 0.5 | 12/A | 2.5 ± 0.8 | $5.61 \times 10^{-1}$ | 1.2 | $2.77 \times 10^{-1}$ | 1.8 | $1.83 \times 10^{-1}$ | | 5'-UTR variant | Moderate | | | |
| 2 | 11818636 | 54/C | 3.1 ± 0.6 | 27/A | 2.9 ± 0.5 | $8.34 \times 10^{-1}$ | 0.1 | $7.12 \times 10^{-1}$ | 0 | $9.12 \times 10^{-1}$ | | Promoter variant | Moderate | | | |
| 2 | 11818914 | 7/C | 7.2 ± 3.1 | 74/A | 2.6 ± 0.4 | **$1.56 \times 10^{-3}$** | **10.5** | **$1.77 \times 10^{-3}$** | 2.6 | $1.13 \times 10^{-1}$ | ✓ | intron variant | Moderate | | | |
| 3 | 273034 | 72/G | 3.2 ± 0.5 | 9/C | 1.9 ± 0.8 | $3.47 \times 10^{-1}$ | 0.3 | $6.01 \times 10^{-1}$ | 0.2 | $6.20 \times 10^{-1}$ | | Promoter variant | Moderate | AT3G01760 | *LHT6* | Lysine-histidine-like transporter 6 |
| 3 | 273099 | 71/C | 3.2 ± 0.5 | 10/T | 1.8 ± 0.7 | $2.55 \times 10^{-1}$ | 0.5 | $4.83 \times 10^{-1}$ | 0.3 | $6.15 \times 10^{-1}$ | | Promoter variant | Moderate | | | |
| 3 | 274655 | 5/T | 6.1 ± 4.4 | 76/A | 2.8 ± 0.4 | $5.59 \times 10^{-2}$ | **4.9** | **$2.95 \times 10^{-2}$** | 0.2 | $6.40 \times 10^{-1}$ | ✓ | synonymous variant | Moderate | | | |
| 3 | 274712 | 78/T | 3.1 ± 0.5 | 3/C | 0.9 ± 0.2 | $3.15 \times 10^{-1}$ | 0.9 | $3.40 \times 10^{-1}$ | 0.6 | $4.58 \times 10^{-1}$ | | splice donor variant | **High** | | | |
| 3 | 275226 | 72/A | 3.2 ± 0.5 | 9/G | 1.8 ± 0.8 | $3.12 \times 10^{-1}$ | 0.3 | $5.71 \times 10^{-1}$ | 0 | $8.73 \times 10^{-1}$ | | amino acid substitution (Lys442Glu) | Moderate | | | |
| 3 | 275682 | 72/T | 3.2 ± 0.5 | 9/C | 1.8 ± 0.8 | $3.12 \times 10^{-1}$ | 0.3 | $5.71 \times 10^{-1}$ | 0 | $8.73 \times 10^{-1}$ | | amino acid substitution (Gln588Arg) | Moderate | AT3G01770 | *BET10, GTE11* | Bromodomain and extraterminal domain protein 10 |
| 3 | 276546 | 11/T | 3.9 ± 2.1 | 70/C | 2.9 ± 0.4 | $3.94 \times 10^{-1}$ | 1.6 | $2.12 \times 10^{-1}$ | 0 | $8.69 \times 10^{-1}$ | | amino acid substitution (Arg418Lys) | Moderate | | | |
| 3 | 278097 | 13/T | 3.5 ± 1.8 | 68/C | 3 ± 0.4 | $6.33 \times 10^{-1}$ | 0.7 | $4.05 \times 10^{-1}$ | 0.4 | $5.08 \times 10^{-1}$ | | amino acid substitution (Ser97Asn) | Moderate | | | |
| 3 | 278139 | 13/G | 3.5 ± 1.8 | 68/A | 3 ± 0.4 | $6.33 \times 10^{-1}$ | 0.7 | $4.05 \times 10^{-1}$ | 0.4 | $5.08 \times 10^{-1}$ | | amino acid substitution (Ile83Thr) | Moderate | | | |
| 3 | 278163 | 13/A | 3.5 ± 1.8 | 68/C | 3 ± 0.4 | $6.33 \times 10^{-1}$ | 0.7 | $4.05 \times 10^{-1}$ | 0.4 | $5.08 \times 10^{-1}$ | | amino acid substitution (Gly75Val) | Moderate | | | |
| 3 | 278857 | 10/T | 3.6 ± 2.3 | 71/A | 3 ± 0.4 | $6.41 \times 10^{-1}$ | 0.6 | $4.26 \times 10^{-1}$ | 0.6 | $4.41 \times 10^{-1}$ | | Promoter variant | Moderate | | | |
| 3 | 279329 | 12/A | 3.2 ± 1.9 | 69/T | 3 ± 0.4 | $8.55 \times 10^{-1}$ | 0.2 | $6.76 \times 10^{-1}$ | 1.5 | $2.18 \times 10^{-1}$ | | synonymous variant | Moderate | | | |
| 3 | 278188 | 13/A | 3.5 ± 1.8 | 68/G | 3 ± 0.4 | $6.33 \times 10^{-1}$ | 0.7 | $4.05 \times 10^{-1}$ | 0.4 | $5.08 \times 10^{-1}$ | | synonymous variant | Moderate | AT3G01780 | *TPLATE* | ARM repeat superfamily protein |
| 3 | 278234 | 69/C | 3 ± 0.5 | 12/T | 3 ± 1.1 | $9.84 \times 10^{-1}$ | 0 | $9.38 \times 10^{-1}$ | 0 | $9.46 \times 10^{-1}$ | | synonymous variant | Moderate | | | |
| 3 | 283148 | 33/T | 3.9 ± 0.6 | 48/C | 2.5 ± 0.6 | $1.08 \times 10^{-1}$ | 1.7 | $2.00 \times 10^{-1}$ | **5.4** | **$2.31 \times 10^{-2}$** | | amino acid substitution (Ser1094Pro) | Moderate | | | |
| 3 | 278495 | 37/G | 3.7 ± 0.6 | 44/A | 2.5 ± 0.7 | $1.46 \times 10^{-1}$ | 1.3 | $2.50 \times 10^{-1}$ | **5.4** | **$2.27 \times 10^{-2}$** | | 5'-UTR intron variant | Moderate | AT3G01770, AT3G01780 | *BET10, GTE11; TPLATE* | Bromodomain and extraterminal domain protein 10; ARM repeat superfamily protein |
| 3 | 278504 | 12/A | 3.7 ± 1.9 | 69/G | 2.9 ± 0.4 | $5.08 \times 10^{-1}$ | 1.1 | $3.04 \times 10^{-1}$ | 0.2 | $6.25 \times 10^{-1}$ | | 5'-UTR intron variant | Moderate | | | |
| 3 | 278510 | 11/C | 3.9 ± 2.1 | 70/A | 2.9 ± 0.4 | $3.94 \times 10^{-1}$ | 1.6 | $2.12 \times 10^{-1}$ | 0 | $8.69 \times 10^{-1}$ | | 5'-UTR intron variant | Moderate | | | |
| 3 | 278566 | 39/C | 3.4 ± 0.6 | 42/T | 2.7 ± 0.7 | $3.59 \times 10^{-1}$ | 0.5 | $5.00 \times 10^{-1}$ | 2.6 | $1.09 \times 10^{-1}$ | | 5'-UTR variant | Moderate | | | |
| 3 | 278593 | 64/G | 3.2 ± 0.5 | 17/A | 2.3 ± 0.8 | $3.84 \times 10^{-1}$ | 0.8 | $3.89 \times 10^{-1}$ | 0.6 | $4.59 \times 10^{-1}$ | | 5'-UTR premature start codon gain variant | Moderate | | | |
| 3 | 278597 | 74/T | 3.2 ± 0.5 | 7/C | 1.8 ± 0.9 | $3.65 \times 10^{-1}$ | 0.2 | $6.68 \times 10^{-1}$ | 0 | $9.46 \times 10^{-1}$ | | 5'-UTR premature start codon gain variant | Moderate | | | |
| 3 | 278615 | 7/T | 5.4 ± 1 | 74/A | 2.8 ± 0.5 | $8.47 \times 10^{-2}$ | 1.4 | $2.47 \times 10^{-1}$ | 2.5 | $1.20 \times 10^{-1}$ | | 5'-UTR premature start codon gain variant | Moderate | | | |
| 3 | 278626 | 71/T | 3.2 ± 0.5 | 10/A | 1.9 ± 0.7 | $2.88 \times 10^{-1}$ | 0.5 | $4.81 \times 10^{-1}$ | 0.2 | $6.93 \times 10^{-1}$ | | 5'-UTR variant | Moderate | | | |
| 3 | 278637 | 71/A | 3.2 ± 0.5 | 10/G | 1.9 ± 0.7 | $2.88 \times 10^{-1}$ | 0.5 | $4.81 \times 10^{-1}$ | 0.2 | $6.93 \times 10^{-1}$ | | 5'-UTR variant | Moderate | | | |
| 3 | 278672 | 65/T | 3.2 ± 0.5 | 16/C | 2.4 ± 0.9 | $4.17 \times 10^{-1}$ | 0.8 | $3.77 \times 10^{-1}$ | 0.6 | $4.42 \times 10^{-1}$ | | 5'-UTR variant | Moderate | | | |
| 3 | 278731 | 11/C | 3.3 ± 2.1 | 70/T | 3 ± 0.4 | $7.77 \times 10^{-1}$ | 0.4 | $5.53 \times 10^{-1}$ | 1.1 | $2.88 \times 10^{-1}$ | | 5'-UTR premature start codon gain variant | Moderate | | | |
| 3 | 279009 | 55/C | 3.5 ± 0.6 | 26/G | 2 ± 0.6 | $9.43 \times 10^{-2}$ | 2.3 | $1.33 \times 10^{-1}$ | 2.4 | $1.26 \times 10^{-1}$ | | Promoter variant | Moderate | | | |

| | | | | | | | | | | | | | | | | |
|---|---|---|---|---|---|---|---|---|---|---|---|---|---|---|---|---|
| 3 | 279049 | 32/A | 3.7 ± 0.9 | 49/G | 2.6 ± 0.4 | $2.17 \times 10^{-1}$ | 1.6 | $2.06 \times 10^{-1}$ | 0.4 | $5.21 \times 10^{-1}$ | | Promoter variant | Moderate | | | |
| 3 | 279139 | 13/T | 4.2 ± 1.1 | 68/G | 2.8 ± 0.5 | $2.16 \times 10^{-1}$ | 0.5 | $4.69 \times 10^{-1}$ | 1.3 | $2.66 \times 10^{-1}$ | | 5'-UTR premature start codon gain variant | Moderate | | | |
| 3 | 518477 | 19/C | 5.5 ± 1.3 | 62/T | 2.3 ± 0.4 | **$9.93 \times 10^{-4}$** | **10.2** | **$2.09 \times 10^{-3}$** | **4.7** | **$3.29 \times 10^{-2}$** | | Promoter variant | Moderate | AT3G02500 | | Mental retardation GTPase activating protein |
| 3 | 518585 | 65/A | 3.4 ± 0.5 | 16/T | 1.4 ± 0.4 | $5.05 \times 10^{-2}$ | **4.4** | **$3.95 \times 10^{-2}$** | 2.5 | $1.17 \times 10^{-1}$ | | Promoter variant | Moderate | | | |
| 3 | 520068 | 62/A | 3.4 ± 0.6 | 19/C | 1.9 ± 0.4 | $1.20 \times 10^{-1}$ | **3.4** | $7.07 \times 10^{-2}$ | 1.9 | $1.74 \times 10^{-1}$ | | amino acid substitution (Tyr87Ser) | Moderate | | | |
| 3 | 520275 | 72/T | 3.1 ± 0.5 | 9/A | 2.6 ± 1 | $7.33 \times 10^{-1}$ | 0 | $9.40 \times 10^{-1}$ | 0.2 | $6.54 \times 10^{-1}$ | | amino acid substitution (Ile156Asn) | Moderate | | | |
| 3 | 520449 | 60/G | 3.5 ± 0.6 | 21/C | 1.8 ± 0.4 | $7.46 \times 10^{-2}$ | **4.8** | **$3.23 \times 10^{-2}$** | **3.1** | $8.34 \times 10^{-2}$ | | amino acid substitution (Gly186Arg) | Moderate | | | |
| 3 | 520501 | 72/C | 3.1 ± 0.5 | 9/A | 2.6 ± 1 | $7.33 \times 10^{-1}$ | 0 | $9.40 \times 10^{-1}$ | 0.2 | $6.54 \times 10^{-1}$ | | amino acid substitution (Thr203Lys) | Moderate | | | |
| 3 | 522102 | 23/C | 4 ± 1.1 | 58/T | 2.7 ± 0.4 | $1.36 \times 10^{-1}$ | 1.7 | $1.98 \times 10^{-1}$ | 0 | $8.80 \times 10^{-1}$ | | amino acid substitution (Lys389Arg) | Moderate | AT3G02510 | | Regulator of chromosome condensation (RCC1) family protein |
| 3 | 522216 | 73/T | 3.1 ± 0.5 | 8/A | 2.9 ± 1.1 | $9.05 \times 10^{-1}$ | 0.1 | $8.08 \times 10^{-1}$ | 0.6 | $4.45 \times 10^{-1}$ | | amino acid substitution (Lys351Met) | Moderate | | | |
| 3 | 523584 | 73/G | 3.1 ± 0.5 | 8/C | 2.9 ± 1.1 | $9.05 \times 10^{-1}$ | 0.1 | $8.08 \times 10^{-1}$ | 0.6 | $4.45 \times 10^{-1}$ | | amino acid substitution (Ala114Gly) | Moderate | | | |
| 3 | 524543 | 75/C | 3.1 ± 0.5 | 6/G | 2.6 ± 1.3 | $7.88 \times 10^{-1}$ | 0 | $8.40 \times 10^{-1}$ | 0.4 | $5.43 \times 10^{-1}$ | | 5'-UTR premature start codon gain variant | Moderate | | | |
| 3 | 524549 | 6/G | 3.6 ± 1.3 | 75/A | 3 ± 0.5 | $7.17 \times 10^{-1}$ | 0.4 | $5.35 \times 10^{-1}$ | 1.7 | $1.94 \times 10^{-1}$ | | 5'-UTR variant | Moderate | | | |
| 3 | 524601 | 74/G | 3.1 ± 0.5 | 7/A | 2.4 ± 1.1 | $6.40 \times 10^{-1}$ | 0 | $9.14 \times 10^{-1}$ | 0.1 | $7.40 \times 10^{-1}$ | | 5'-UTR variant | Moderate | | | |
| 3 | 524631 | 7/G | 3.2 ± 1.2 | 74/A | 3 ± 0.5 | $8.90 \times 10^{-1}$ | 0.3 | $5.60 \times 10^{-1}$ | 1.3 | $2.66 \times 10^{-1}$ | | 5'-UTR variant | Moderate | | | |
| 3 | 524678 | 55/T | 3.6 ± 0.6 | 26/A | 1.8 ± 0.4 | **$3.53 \times 10^{-2}$** | **6** | **$1.66 \times 10^{-2}$** | **3.5** | $6.55 \times 10^{-2}$ | | Promoter variant | Moderate | | | |
| 3 | 524766 | 16/A | 5 ± 0.9 | 65/T | 2.6 ± 0.5 | **$2.01 \times 10^{-2}$** | **3.5** | $6.52 \times 10^{-2}$ | **4.8** | **$3.17 \times 10^{-2}$** | | Promoter variant | Moderate | | | |
| 3 | 524845 | 61/G | 3.4 ± 0.6 | 20/A | 2 ± 0.4 | $1.36 \times 10^{-1}$ | 2.6 | $1.12 \times 10^{-1}$ | 1.2 | $2.76 \times 10^{-1}$ | | Promoter variant | Moderate | | | |
| 3 | 524905 | 39/G | 3.7 ± 0.6 | 42/A | 2.4 ± 0.7 | $1.29 \times 10^{-1}$ | 2.3 | $1.33 \times 10^{-1}$ | **4.7** | **$3.31 \times 10^{-2}$** | | Promoter variant | Moderate | | | |
| 3 | 524954 | 38/G | 4 ± 0.6 | 43/C | 2.2 ± 0.6 | **$3.31 \times 10^{-2}$** | **5.8** | **$1.85 \times 10^{-2}$** | **9.7** | **$2.60 \times 10^{-3}$** | | Promoter variant | Moderate | | | |
| 3 | 524990 | 52/G | 3.5 ± 0.6 | 29/T | 2.1 ± 0.5 | $1.09 \times 10^{-1}$ | 2.3 | $1.36 \times 10^{-1}$ | 1.2 | $2.86 \times 10^{-1}$ | | Promoter variant | Moderate | | | |
| 3 | 525013 | 43/C | 3.4 ± 0.6 | 37/T | 2.6 ± 0.7 | $3.40 \times 10^{-1}$ | 0.8 | $4.63 \times 10^{-1}$ | 1.8 | $1.74 \times 10^{-1}$ | | Promoter variant | Moderate | | | |
| 3 | 525286 | 63/C | 3.5 ± 0.6 | 18/T | 1.5 ± 0.4 | **$4.84 \times 10^{-2}$** | **4.9** | $2.93 \times 10^{-2}$ | 3.3 | $7.14 \times 10^{-2}$ | | Promoter variant | Moderate | | | |
| 3 | 525352 | 54/G | 3.7 ± 0.6 | 27/A | 1.7 ± 0.3 | **$1.80 \times 10^{-2}$** | **7.5** | $7.94 \times 10^{-3}$ | 5 | **$2.87 \times 10^{-2}$** | | Promoter variant | Moderate | | | |
| 3 | 525380 | 11/T | 3.4 ± 1.3 | 70/A | 3 ± 0.5 | $7.28 \times 10^{-1}$ | 0.3 | $5.99 \times 10^{-1}$ | 1.4 | $2.36 \times 10^{-1}$ | | Promoter variant | Moderate | | | |
| 3 | 525469 | 36/A | 3.9 ± 0.6 | 45/T | 2.4 ± 0.6 | $6.48 \times 10^{-2}$ | **3.2** | $7.62 \times 10^{-2}$ | 5.6 | **$2.03 \times 10^{-2}$** | | Promoter variant | Moderate | | | |
| 3 | 525475 | 48/C | 3.7 ± 0.7 | 33/T | 2.2 ± 0.5 | $7.65 \times 10^{-2}$ | **4.7** | $3.29 \times 10^{-2}$ | 1.9 | $1.78 \times 10^{-1}$ | | Promoter variant | Moderate | | | |
| 3 | 526854 | 7/T | 5.7 ± 3.2 | 74/C | 2.8 ± 0.4 | **$4.99 \times 10^{-2}$** | **7.4** | $7.97 \times 10^{-3}$ | 0.8 | $3.71 \times 10^{-1}$ | ✓ | synonymous variant | Moderate | AT3G02520 | *GRF7* | General regulatory factor 7 |
| 3 | 526998 | 73/A | 3.1 ± 0.5 | 8/G | 2.9 ± 1.1 | $9.05 \times 10^{-1}$ | 0.1 | $8.08 \times 10^{-1}$ | 0.6 | $4.45 \times 10^{-1}$ | | amino acid substitution (Ile238Thr) | Moderate | | | |
| 3 | 528330 | 21/G | 4.4 ± 0.8 | 60/T | 2.6 ± 0.5 | $6.06 \times 10^{-2}$ | 2.8 | $9.65 \times 10^{-2}$ | 3.9 | $5.30 \times 10^{-2}$ | | Promoter variant | Moderate | | | |
| 3 | 528374 | 49/A | 3.6 ± 0.7 | 32/G | 2.2 ± 0.5 | $1.01 \times 10^{-1}$ | **4.4** | $3.85 \times 10^{-2}$ | 1.8 | $1.80 \times 10^{-1}$ | | Promoter variant | Moderate | | | |
| 3 | 528418 | 23/T | 4.2 ± 0.7 | 58/C | 2.6 ± 0.6 | $8.31 \times 10^{-2}$ | 2.2 | $1.44 \times 10^{-1}$ | 2.7 | $1.03 \times 10^{-1}$ | | Promoter variant | Moderate | | | |
| 3 | 528442 | 12/C | 5.3 ± 1.1 | 69/CTT | 2.6 ± 0.5 | **$2.02 \times 10^{-2}$** | **4.1** | $4.75 \times 10^{-2}$ | 5.7 | **$1.93 \times 10^{-2}$** | | Promoter variant | Moderate | | | |
| 3 | 532603 | 7/C | 3.2 ± 1.2 | 74/A | 3 ± 0.5 | $8.90 \times 10^{-1}$ | 0.3 | $5.60 \times 10^{-1}$ | 1.3 | $2.66 \times 10^{-1}$ | | Promoter variant | Moderate | AT3G02530 | *CCT6-2* | TCP-1/cpn60 chaperonin family protein |
| 3 | 532818 | 77/A | 3.2 ± 0.5 | 4/T | 0.8 ± 0.3 | $2.22 \times 10^{-1}$ | 1.2 | $2.71 \times 10^{-1}$ | 0.8 | $3.84 \times 10^{-1}$ | | Promoter variant | Moderate | | | |
| 3 | 768931 | 74/T | 3.2 ± 0.5 | 7/C | 0.9 ± 0.2 | $1.12 \times 10^{-1}$ | **3.5** | $6.72 \times 10^{-2}$ | 3.5 | $6.70 \times 10^{-2}$ | | amino acid substitution (Lys1145Glu) | Moderate | AT3G03300 | *DCL2* | Dicer like 2 |
| 3 | 773843 | 5/T | 6.8 ± 4.3 | 76/C | 2.8 ± 0.4 | **$1.96 \times 10^{-2}$** | **7.5** | $7.57 \times 10^{-3}$ | 0.4 | $5.08 \times 10^{-1}$ | | amino acid substitution (Val148Ile) | Moderate | | | |

| | | | | | | | | | | | | | | | | |
|---|---|---|---|---|---|---|---|---|---|---|---|---|---|---|---|---|
| 3 | 773844 | 1/T | 23.3 | 80/C | 2.8 ± 0.4 | | **55.2** | $1.65 \times 10^{-10}$ | 2 | $1.62 \times 10^{-1}$ | ✓ | synonymous variant | Moderate | | | |
| 3 | 774858 | 10/GA | 4.6 ± 1 | 71/GAAA | 2.8 ± 0.5 | $1.53 \times 10^{-1}$ | 1.8 | $1.82 \times 10^{-1}$ | 3 | $8.60 \times 10^{-2}$ | | 5'-UTR variant | Moderate | | | |
| 3 | 775100 | 41/A | 3.5 ± 0.7 | 40/G | 2.6 ± 0.5 | $2.65 \times 10^{-1}$ | 0.8 | $3.86 \times 10^{-1}$ | 0 | $9.56 \times 10^{-1}$ | | 5'-UTR premature start codon gain variant | Moderate | | | |
| 3 | 775252 | 10/G | 5.2 ± 2.2 | 71/T | 2.7 ± 0.4 | $4.61 \times 10^{-2}$ | 4.5 | $3.73 \times 10^{-2}$ | 0.4 | $5.10 \times 10^{-1}$ | | 5'-UTR intron variant | Moderate | | | |
| 3 | 775492 | 5/C | 6.8 ± 4.3 | 76/T | 2.8 ± 0.4 | $1.96 \times 10^{-2}$ | 7.5 | $7.57 \times 10^{-3}$ | 0.4 | $5.08 \times 10^{-1}$ | | 5'-UTR intron variant | Moderate | | | |
| 3 | 775640 | 6/T | 5.7 ± 3.7 | 75/A | 2.8 ± 0.4 | $6.88 \times 10^{-2}$ | 6.9 | $1.03 \times 10^{-2}$ | 0.5 | $5.03 \times 10^{-1}$ | | 5'-UTR intron variant | Moderate | | | |
| 3 | 775662 | 4/AC | 8 ± 5.3 | 77/A | 2.8 ± 0.4 | **$5.84 \times 10^{-3}$** | 9.3 | $3.24 \times 10^{-3}$ | 0.6 | $4.56 \times 10^{-1}$ | | Promoter variant | Moderate | | | |
| 3 | 775673 | 18/A | 3.1 ± 0.6 | 63/C | 3 ± 0.6 | $9.26 \times 10^{-1}$ | 0.4 | $5.25 \times 10^{-1}$ | 0.9 | $3.59 \times 10^{-1}$ | | Promoter variant | Moderate | | | |
| 3 | 775680 | 5/A | 6.8 ± 4.3 | 76/T | 2.8 ± 0.4 | **$1.96 \times 10^{-2}$** | 7.5 | $7.57 \times 10^{-3}$ | 0.4 | $5.08 \times 10^{-1}$ | | Promoter variant | Moderate | | | |
| 3 | 3386436 | 26/C | 4.1 ± 1 | 55/CA | 2.6 ± 0.4 | $9.15 \times 10^{-2}$ | 2.4 | $1.22 \times 10^{-1}$ | 0.4 | $5.08 \times 10^{-1}$ | | Promoter variant | Moderate | AT3G10820 | *MED26A* | Transcription elongation factor (TFIIS) family protein |
| 3 | 3386763 | 12/C | 3.5 ± 1.1 | 69/T | 3 ± 0.5 | $6.28 \times 10^{-1}$ | 0 | $9.52 \times 10^{-1}$ | 0 | $8.73 \times 10^{-1}$ | | Promoter variant | Moderate | | | |
| 3 | 3387178 | 46/A | 3.2 ± 0.7 | 35/C | 2.9 ± 0.5 | $6.94 \times 10^{-1}$ | 0.4 | $5.08 \times 10^{-1}$ | 0 | $8.79 \times 10^{-1}$ | | Promoter variant | Moderate | | | |
| 3 | 3387229 | 76/A | 3.1 ± 0.5 | 5/T | 1.7 ± 0.6 | $4.28 \times 10^{-1}$ | 0.2 | $6.20 \times 10^{-1}$ | 0 | $9.15 \times 10^{-1}$ | | 5'-UTR variant | Moderate | | | |
| 3 | 3387271 | 76/A | 3.1 ± 0.5 | 5/G | 2.6 ± 1.5 | $7.96 \times 10^{-1}$ | 0.2 | $6.91 \times 10^{-1}$ | 0.2 | $6.53 \times 10^{-1}$ | | 5'-UTR variant | Moderate | | | |
| 3 | 3387498 | 49/A | 3.3 ± 0.7 | 32/T | 2.7 ± 0.6 | $4.93 \times 10^{-1}$ | 1.5 | $2.22 \times 10^{-1}$ | 1.3 | $2.64 \times 10^{-1}$ | | amino acid substitution (Ser47Arg) | Moderate | | | |
| 3 | 3388426 | 79/G | 3.1 ± 0.5 | 2/A | 2.6 ± 2.2 | $8.68 \times 10^{-1}$ | 0.0 | $9.14 \times 10^{-1}$ | 0.0 | $9.56 \times 10^{-1}$ | | stop codon gain (Trp183*) | **High** | | | |
| 3 | 3388954 | 17/A | 3.5 ± 0.9 | 64/G | 2.9 ± 0.5 | $6.04 \times 10^{-1}$ | 0.4 | $5.47 \times 10^{-1}$ | 0.8 | $3.71 \times 10^{-1}$ | | amino acid substitution (Met265Ile) | Moderate | | | |
| 3 | 3393119 | 9/T | 4.2 ± 2.5 | 72/C | 2.9 ± 0.4 | $3.24 \times 10^{-1}$ | 1.7 | $1.95 \times 10^{-1}$ | 0.4 | $5.23 \times 10^{-1}$ | | Promoter variant | Moderate | AT3G10845 | | RNA-binding (RRM/RBD/RNP motifs) family protein |
| 3 | 3393146 | 70/G | 3.3 ± 0.5 | 11/A | 1.3 ± 0.5 | $9.98 \times 10^{-2}$ | **3.4** | $6.96 \times 10^{-2}$ | 4.5 | **$3.72 \times 10^{-2}$** | | Promoter variant | Moderate | | | |
| 3 | 3393595 | 7/T | 3.7 ± 1.8 | 73/A | 3 ± 0.5 | $6.26 \times 10^{-1}$ | 0.2 | $8.25 \times 10^{-1}$ | 0.4 | $6.42 \times 10^{-1}$ | | Promoter variant | Moderate | | | |
| 3 | 3394337 | 69/T | 3.3 ± 0.5 | 12/G | 1.4 ± 0.5 | $9.98 \times 10^{-2}$ | **3.6** | $6.23 \times 10^{-2}$ | 4.9 | **$2.94 \times 10^{-2}$** | | amino acid substitution (Ile57Ser) | Moderate | | | |
| 3 | 3394877 | 12/G | 3.8 ± 1.9 | 69/T | 2.9 ± 0.4 | $4.44 \times 10^{-1}$ | 1.4 | $2.36 \times 10^{-1}$ | 0.1 | $7.58 \times 10^{-1}$ | | amino acid substitution (Leu88Arg) | Moderate | | | |
| 3 | 3394900 | 54/G | 3.1 ± 0.6 | 27/C | 2.9 ± 0.6 | $7.61 \times 10^{-1}$ | 0 | $8.44 \times 10^{-1}$ | 0.1 | $7.31 \times 10^{-1}$ | | amino acid substitution (Val96Leu) | Moderate | | | |
| 3 | 3395184 | 68/C | 3.4 ± 0.5 | 13/T | 1.4 ± 0.4 | $7.87 \times 10^{-2}$ | **3.6** | $6.04 \times 10^{-2}$ | 5.1 | **$2.71 \times 10^{-2}$** | | amino acid substitution (Ala125Val) | Moderate | | | |
| 3 | 3395453 | 1/G | 12.3 ± 0 | 80/GT | 2.9 ± 0.5 | | 5.6 | $2.08 \times 10^{-2}$ | 9.6 | $2.73 \times 10^{-3}$ | | frameshift variant (Gly145) | **High** | | | |
| 3 | 3395510 | 17/A | 3.3 ± 1 | 64/G | 3 ± 0.5 | $7.25 \times 10^{-1}$ | 0 | $9.88 \times 10^{-1}$ | 0.5 | $4.94 \times 10^{-1}$ | | amino acid substitution (Asp164Gly) | Moderate | | | |
| 3 | 3395649 | 6/A | 11.2 ± 2.8 | 75/G | 2.4 ± 0.3 | **$8.93 \times 10^{-10}$** | **44.4** | **$4.42 \times 10^{-9}$** | **19.2** | **$3.95 \times 10^{-5}$** | ✓ | intron variant | Moderate | | | |
| 3 | 3395733 | 11/A | 3.7 ± 1.1 | 70/T | 2.9 ± 0.5 | $5.29 \times 10^{-1}$ | 0.7 | $4.02 \times 10^{-1}$ | 1.5 | $2.23 \times 10^{-1}$ | | amino acid substitution (Glu190Asp) | Moderate | | | |
| 3 | 3395739 | 11/T | 3.4 ± 1.2 | 70/G | 3 ± 0.5 | $7.37 \times 10^{-1}$ | 0.7 | $4.08 \times 10^{-1}$ | 2 | $1.63 \times 10^{-1}$ | | amino acid substitution (Phe192Leu) | Moderate | | | |
| 3 | 3395785 | 8/G | 5.2 ± 1.8 | 73/A | 2.8 ± 0.5 | $8.77 \times 10^{-2}$ | 2.7 | $1.07 \times 10^{-1}$ | **4.8** | **$3.20 \times 10^{-2}$** | | amino acid substitution (Asp208Asn) | Moderate | | | |
| 3 | 3396135 | 80/AGG | 3.1 ± 0.5 | 1/A | 1.1 ± 0 | | 0.2 | $6.24 \times 10^{-1}$ | 1.0 | $3.22 \times 10^{-1}$ | | frameshift variant (Lys225) | **High** | | | |
| 3 | 3396350 | 80/G | 3.1 ± 0.5 | 1/T | 1 ± 0 | | 0.4 | $5.24 \times 10^{-1}$ | 0.8 | $3.64 \times 10^{-1}$ | | splice donor variant | **High** | | | |
| 3 | 3396911 | 15/A | 3.4 ± 0.9 | 66/T | 3 ± 0.5 | $6.75 \times 10^{-1}$ | 0.1 | $7.40 \times 10^{-1}$ | 1.1 | $2.99 \times 10^{-1}$ | | amino acid substitution (Leu314Met) | Moderate | | | |
| 3 | 3397142 | 71/C | 3.1 ± 0.5 | 10/T | 2.3 ± 0.6 | $5.17 \times 10^{-1}$ | 0.6 | $4.31 \times 10^{-1}$ | 0 | $9.13 \times 10^{-1}$ | | amino acid substitution (Arg362Cys) | Moderate | | | |
| 3 | 3397147 | 1/G | 4.4 ± 0 | 80/T | 3 ± 0.5 | | 0.3 | $5.95 \times 10^{-1}$ | 0.7 | $4.09 \times 10^{-1}$ | | stop codon gain (Tyr363*) | **High** | | | |
| 3 | 3397200 | 71/T | 3.1 ± 0.5 | 10/A | 2.3 ± 0.6 | $5.17 \times 10^{-1}$ | 0.6 | $4.31 \times 10^{-1}$ | 0 | $9.13 \times 10^{-1}$ | | amino acid substitution (Met381Lys) | Moderate | | | |
| 3 | 3397443 | 10/T | 4 ± 1.2 | 71/G | 2.9 ± 0.5 | $4.03 \times 10^{-1}$ | 0.5 | $4.80 \times 10^{-1}$ | 3.5 | $6.71 \times 10^{-2}$ | | stop codon gain (Glu397*) | **High** | | | |

| | | | | | | | | | | | | | | | | |
|---|---|---|---|---|---|---|---|---|---|---|---|---|---|---|---|---|
| 3 | 3397444 | 9/T | 4 ± 1.3 | 72/A | 2.9 ± 0.5 | $4.40 \times 10^{-1}$ | 0.5 | $4.69 \times 10^{-1}$ | **3.7** | $5.97 \times 10^{-2}$ | | amino acid substitution (Glu397Val) | Moderate | | | |
| 3 | 3397446 | 49/T | 3.4 ± 0.7 | 32/C | 2.6 ± 0.6 | $3.48 \times 10^{-1}$ | 0.5 | $4.88 \times 10^{-1}$ | 0 | $9.12 \times 10^{-1}$ | | amino acid substitution (Ser398Pro) | Moderate | | | |
| 3 | 3397461 | 73/C | 3.1 ± 0.5 | 8/G | 2.9 ± 0.8 | $9.24 \times 10^{-1}$ | 0 | $9.34 \times 10^{-1}$ | 0.6 | $4.36 \times 10^{-1}$ | | amino acid substitution (Gln403Glu) | Moderate | | | |
| 3 | 3397468 | 9/A | 4 ± 1.3 | 72/C | 2.9 ± 0.5 | $4.40 \times 10^{-1}$ | 0.5 | $4.69 \times 10^{-1}$ | **3.7** | $5.97 \times 10^{-2}$ | | amino acid substitution (Ala405Glu) | Moderate | | | |
| 3 | 3398537 | 40/T | 3.7 ± 0.8 | 41/G | 2.4 ± 0.5 | $1.11 \times 10^{-1}$ | 2.1 | $1.48 \times 10^{-1}$ | 0.7 | $3.94 \times 10^{-1}$ | | amino acid substitution (Asn122Thr) | Moderate | AT3G10850 | *GLX2-2* | Metallo-hydrolase/oxidoreductase superfamily protein |
| 3 | 3399303 | 73/T | 3.1 ± 0.5 | 8/G | 2.1 ± 0.6 | $4.80 \times 10^{-1}$ | 0.9 | $3.59 \times 10^{-1}$ | 0.1 | $7.35 \times 10^{-1}$ | | amino acid substitution (Lys42Gln) | Moderate | | | |
| 3 | 3399314 | 73/G | 3.1 ± 0.5 | 8/T | 2.1 ± 0.6 | $4.80 \times 10^{-1}$ | 0.9 | $3.59 \times 10^{-1}$ | 0.1 | $7.35 \times 10^{-1}$ | | amino acid substitution (Ala38Glu) | Moderate | | | |
| 3 | 3399553 | 37/C | 3.4 ± 0.8 | 44/A | 2.7 ± 0.5 | $4.14 \times 10^{-1}$ | 0.9 | $3.43 \times 10^{-1}$ | 0.2 | $6.36 \times 10^{-1}$ | | 5'-UTR premature start codon gain variant | Moderate | | | |
| 3 | 3399554 | 51/A | 3.1 ± 0.5 | 30/G | 3 ± 0.9 | $9.09 \times 10^{-1}$ | 0 | $9.04 \times 10^{-1}$ | 1.5 | $2.26 \times 10^{-1}$ | | 5'-UTR premature start codon gain variant | Moderate | | | |
| 3 | 3399660 | 25/A | 3.2 ± 1 | 56/C | 3 ± 0.5 | $8.10 \times 10^{-1}$ | 0.1 | $7.91 \times 10^{-1}$ | 0.4 | $5.05 \times 10^{-1}$ | | Promoter variant | Moderate | | | |
| 3 | 3399808 | 30/A | 3.4 ± 0.9 | 51/G | 2.8 ± 0.5 | $4.91 \times 10^{-1}$ | 0.2 | $6.19 \times 10^{-1}$ | 0 | $8.66 \times 10^{-1}$ | | 5'-UTR variant | Moderate | | | |
| 3 | 3399880 | 50/T | 3.1 ± 0.5 | 31/C | 3 ± 0.9 | $9.67 \times 10^{-1}$ | 0 | $8.78 \times 10^{-1}$ | 1 | $3.14 \times 10^{-1}$ | | synonymous variant | Moderate | | | |
| 3 | 3399919 | 53/T | 3.1 ± 0.5 | 28/C | 2.9 ± 0.9 | $8.29 \times 10^{-1}$ | 0 | $9.03 \times 10^{-1}$ | 1.1 | $3.00 \times 10^{-1}$ | | synonymous variant | Moderate | | | |
| 3 | 3399992 | 26/A | 3.5 ± 1.1 | 55/T | 2.8 ± 0.4 | $4.24 \times 10^{-1}$ | 0.4 | $5.23 \times 10^{-1}$ | 0 | $9.06 \times 10^{-1}$ | | Promoter variant | Moderate | | | |
| 3 | 3400012 | 53/A | 3.1 ± 0.5 | 28/C | 3 ± 0.9 | $9.01 \times 10^{-1}$ | 0.1 | $7.59 \times 10^{-1}$ | 2 | $1.66 \times 10^{-1}$ | | Promoter variant | Moderate | | | |
| 3 | 3400015 | 29/C | 3.1 ± 0.9 | 52/G | 3 ± 0.5 | $9.67 \times 10^{-1}$ | 0 | $9.94 \times 10^{-1}$ | 0.8 | $3.62 \times 10^{-1}$ | | Promoter variant | Moderate | | | |
| 3 | 3400116 | 61/T | 3.2 ± 0.5 | 20/G | 2.7 ± 0.8 | $6.09 \times 10^{-1}$ | 0.6 | $4.55 \times 10^{-1}$ | 0.5 | $4.66 \times 10^{-1}$ | | Promoter variant | Moderate | | | |
| 3 | 6702806 | 12/G | 3.4 ± 1.9 | 69/T | 3 ± 0.4 | $7.08 \times 10^{-1}$ | 0 | $9.23 \times 10^{-1}$ | 2 | $1.58 \times 10^{-1}$ | | amino acid substitution (Lys218Gln) | Moderate | AT3G19340 | | Aminopeptidase (DUF3754) |
| 3 | 6704056 | 5/G | 9.6 ± 3.9 | 76/T | 2.6 ± 0.4 | **$2.33 \times 10^{-5}$** | **23.2** | **$7.84 \times 10^{-6}$** | **5.1** | **$2.67 \times 10^{-2}$** | ✓ | synonymous variant | Moderate | | | |
| 3 | 6704196 | 16/G | 3.3 ± 1.5 | 65/T | 3 ± 0.4 | $7.40 \times 10^{-1}$ | 0.2 | $6.82 \times 10^{-1}$ | 0.5 | $5.02 \times 10^{-1}$ | | 5'-UTR variant | Moderate | | | |
| 3 | 6704611 | 46/G | 3.1 ± 0.5 | 35/C | 3 ± 0.8 | $8.89 \times 10^{-1}$ | 0.2 | $6.24 \times 10^{-1}$ | 2 | $1.57 \times 10^{-1}$ | | Promoter variant | Moderate | | | |
| 3 | 6704767 | 16/C | 3.2 ± 1.1 | 65/T | 3 ± 0.5 | $8.54 \times 10^{-1}$ | 0.1 | $7.63 \times 10^{-1}$ | 0.4 | $5.20 \times 10^{-1}$ | | Promoter variant | Moderate | | | |
| 3 | 6704906 | 48/T | 3.1 ± 0.7 | 33/C | 3 ± 0.6 | $8.58 \times 10^{-1}$ | 0.2 | $6.86 \times 10^{-1}$ | 2 | $1.64 \times 10^{-1}$ | | Promoter variant | Moderate | | | |
| 3 | 9703154 | 18/A | 5.8 ± 1.4 | 63/G | 2.3 ± 0.4 | **$2.41 \times 10^{-4}$** | **14.5** | **$2.86 \times 10^{-4}$** | **11.9** | **$9.19 \times 10^{-4}$** | | Promoter variant | Moderate | AT3G26490 | *NPY7* | Phototropic-responsive NPH3 family protein |
| 3 | 9703210 | 54/G | 3.5 ± 0.6 | 27/A | 2.1 ± 0.6 | $1.02 \times 10^{-1}$ | 2.6 | $1.13 \times 10^{-1}$ | 1.7 | $1.96 \times 10^{-1}$ | | Promoter variant | Moderate | | | |
| 3 | 9703302 | 4/C | 4.4 ± 3 | 77/G | 3 ± 0.5 | $4.63 \times 10^{-1}$ | 0.2 | $6.55 \times 10^{-1}$ | 0.1 | $7.88 \times 10^{-1}$ | | Promoter variant | Moderate | | | |
| 3 | 9703304 | 60/A | 3.3 ± 0.6 | 21/G | 2.2 ± 0.7 | $2.32 \times 10^{-1}$ | 1.2 | $2.81 \times 10^{-1}$ | 0.9 | $3.33 \times 10^{-1}$ | | Promoter variant | Moderate | | | |
| 3 | 9703349 | 59/G | 3.3 ± 0.6 | 22/A | 2.3 ± 0.7 | $3.11 \times 10^{-1}$ | 0.9 | $3.54 \times 10^{-1}$ | 0.7 | $4.05 \times 10^{-1}$ | | Promoter variant | Moderate | | | |
| 3 | 9703402 | 67/C | 3.1 ± 0.5 | 14/T | 2.8 ± 1 | $8.03 \times 10^{-1}$ | 0.3 | $5.91 \times 10^{-1}$ | 0.1 | $7.98 \times 10^{-1}$ | | Promoter variant | Moderate | | | |
| 3 | 9703450 | 19/T | 5.9 ± 1.3 | 62/C | 2.2 ± 0.4 | **$5.79 \times 10^{-5}$** | **15.6** | **$1.79 \times 10^{-4}$** | **12.8** | **$6.18 \times 10^{-4}$** | | Promoter variant | Moderate | | | |
| 3 | 9703582 | 49/T | 3.8 ± 0.7 | 32/A | 1.9 ± 0.4 | **$2.94 \times 10^{-2}$** | **3.9** | $5.30 \times 10^{-2}$ | 3 | $8.73 \times 10^{-2}$ | | Promoter variant | Moderate | | | |
| 3 | 9703625 | 55/C | 3.2 ± 0.6 | 26/A | 2.6 ± 0.7 | $5.19 \times 10^{-1}$ | 0.3 | $5.82 \times 10^{-1}$ | 0 | $8.31 \times 10^{-1}$ | | Promoter variant | Moderate | | | |
| 3 | 9703882 | 66/C | 3.1 ± 0.5 | 3/CT | 0.4 ± 0.2 | $2.41 \times 10^{-1}$ | 1 | $4.19 \times 10^{-1}$ | 1.6 | $1.93 \times 10^{-1}$ | | Promoter variant | Moderate | | | |
| 3 | 9703918 | 47/T | 3.7 ± 0.7 | 34/A | 2.1 ± 0.5 | **$4.82 \times 10^{-2}$** | **3.2** | $7.69 \times 10^{-2}$ | 2.5 | $1.17 \times 10^{-1}$ | | Promoter variant | Moderate | | | |
| 3 | 9704368 | 77/GT | 3.1 ± 0.5 | 4/G | 1.8 ± 1.2 | $5.14 \times 10^{-1}$ | 0.8 | $3.63 \times 10^{-1}$ | 1.2 | $2.72 \times 10^{-1}$ | | splice donor variant | **High** | | | |
| 3 | 9704481 | 12/A | 6 ± 2 | 69/G | 2.5 ± 0.4 | **$2.91 \times 10^{-3}$** | **8.5** | **$4.72 \times 10^{-3}$** | **4.9** | **$3.05 \times 10^{-2}$** | | amino acid substitution (Arg52His) | Moderate | | | |
| 3 | 9704562 | 12/G | 6 ± 2 | 69/A | 2.5 ± 0.4 | **$2.91 \times 10^{-3}$** | **8.5** | **$4.72 \times 10^{-3}$** | **4.9** | **$3.05 \times 10^{-2}$** | | amino acid substitution (Glu79Gly) | Moderate | | | |

| | | | | | | | | | | | | | | | | |
|---|---|---|---|---|---|---|---|---|---|---|---|---|---|---|---|---|
| 3 | 9704694 | 59/C | 3.4 ± 0.6 | 22/G | 2.1 ± 0.4 | $1.90 \times 10^{-1}$ | 1.7 | $1.93 \times 10^{-1}$ | 1.5 | $2.28 \times 10^{-1}$ | | amino acid substitution (Thr123Arg) | Moderate | | | |
| 3 | 9704856 | 11/A | 5.7 ± 2.1 | 70/C | 2.6 ± 0.4 | **$9.85 \times 10^{-3}$** | **6.4** | **$1.34 \times 10^{-2}$** | 2.6 | $1.12 \times 10^{-1}$ | | amino acid substitution (Thr177Asn) | Moderate | | | |
| 3 | 9704937 | 12/A | 6 ± 2 | 69/T | 2.5 ± 0.4 | **$2.91 \times 10^{-3}$** | **8.5** | **$4.72 \times 10^{-3}$** | 4.9 | **$3.05 \times 10^{-2}$** | | amino acid substitution (Ile204Asn) | Moderate | | | |
| 3 | 9704938 | 11/G | 4.4 ± 1.2 | 70/C | 2.8 ± 0.5 | $2.03 \times 10^{-1}$ | 1 | $3.30 \times 10^{-1}$ | 4.5 | **$3.78 \times 10^{-2}$** | | amino acid substitution (Ile204Met) | Moderate | | | |
| 3 | 9705045 | 12/T | 6 ± 2 | 69/A | 2.5 ± 0.4 | **$2.91 \times 10^{-3}$** | **8.5** | **$4.72 \times 10^{-3}$** | **4.9** | **$3.05 \times 10^{-2}$** | | amino acid substitution (Lys240Ile) | Moderate | | | |
| 3 | 9705053 | 12/A | 6 ± 2 | 69/C | 2.5 ± 0.4 | **$2.91 \times 10^{-3}$** | **8.5** | **$4.72 \times 10^{-3}$** | **4.9** | **$3.05 \times 10^{-2}$** | | amino acid substitution (His243Asn) | Moderate | | | |
| 3 | 9705125 | 12/G | 6 ± 2 | 69/C | 2.5 ± 0.4 | **$2.91 \times 10^{-3}$** | **8.5** | **$4.72 \times 10^{-3}$** | **4.9** | **$3.05 \times 10^{-2}$** | | amino acid substitution (Gln267Glu) | Moderate | | | |
| 3 | 9705267 | 12/C | 6 ± 2 | 69/T | 2.5 ± 0.4 | **$2.91 \times 10^{-3}$** | **8.5** | **$4.72 \times 10^{-3}$** | **4.9** | **$3.05 \times 10^{-2}$** | | amino acid substitution (Met314Thr) | Moderate | | | |
| 3 | 9705641 | 15/T | 5.3 ± 1.6 | 66/A | 2.5 ± 0.4 | **$9.34 \times 10^{-3}$** | **5.5** | **$2.12 \times 10^{-2}$** | 2.9 | $9.15 \times 10^{-2}$ | | amino acid substitution (Thr439Ser) | Moderate | | | |
| 3 | 9705935 | 15/C | 5.3 ± 1.6 | 66/G | 2.5 ± 0.4 | **$9.34 \times 10^{-3}$** | **5.5** | **$2.12 \times 10^{-2}$** | 2.9 | $9.15 \times 10^{-2}$ | | amino acid substitution (Arg514Thr) | Moderate | | | |
| 3 | 9706111 | 18/A | 5.1 ± 1.4 | 63/C | 2.5 ± 0.4 | **$8.56 \times 10^{-3}$** | **5.2** | **$2.49 \times 10^{-2}$** | 2.6 | $1.15 \times 10^{-1}$ | | amino acid substitution (Leu573Met) | Moderate | | | |
| 3 | 22835428 | 11/T | 5.1 ± 2 | 70/C | 2.7 ± 0.4 | **$4.66 \times 10^{-2}$** | **4.2** | **$4.45 \times 10^{-2}$** | 0.5 | $4.68 \times 10^{-1}$ | | 5'-UTR variant | Moderate | AT3G61700 | | Helicase with zinc finger protein |
| 3 | 22835433 | 11/A | 5.1 ± 2 | 70/C | 2.7 ± 0.4 | **$4.66 \times 10^{-2}$** | **4.2** | **$4.45 \times 10^{-2}$** | 0.5 | $4.68 \times 10^{-1}$ | | 5'-UTR variant | Moderate | | | |
| 3 | 22835558 | 58/T | 3.3 ± 0.6 | 23/G | 2.5 ± 0.7 | $3.75 \times 10^{-1}$ | 0.5 | $4.75 \times 10^{-1}$ | 0.4 | $5.29 \times 10^{-1}$ | | 5'-UTR variant | Moderate | | | |
| 3 | 22835716 | 76/GT | 3.1 ± 0.5 | 5/G | 2.9 ± 0.9 | $9.38 \times 10^{-1}$ | 0.1 | $7.82 \times 10^{-1}$ | 0 | $9.17 \times 10^{-1}$ | | 5'-UTR intron variant | Moderate | | | |
| 3 | 22835809 | 10/C | 4.9 ± 2.2 | 71/A | 2.8 ± 0.4 | $9.76 \times 10^{-2}$ | 2 | $1.65 \times 10^{-1}$ | 0.1 | $7.24 \times 10^{-1}$ | | 5'-UTR intron variant | Moderate | | | |
| 3 | 22836223 | 13/C | 4.7 ± 1.8 | 68/T | 2.7 ± 0.4 | $8.85 \times 10^{-2}$ | 2.2 | $1.43 \times 10^{-1}$ | 0.1 | $7.68 \times 10^{-1}$ | | 5'-UTR intron variant | Moderate | | | |
| 3 | 22837659 | 19/A | 4.6 ± 1.4 | 62/G | 2.6 ± 0.4 | **$3.95 \times 10^{-2}$** | **7.6** | **$7.39 \times 10^{-3}$** | **5.1** | **$2.70 \times 10^{-2}$** | | amino acid substitution (Val141Ile) | Moderate | | | |
| 3 | 22842900 | 18/T | 4.6 ± 1.4 | 63/G | 2.6 ± 0.4 | **$4.50 \times 10^{-2}$** | **3.9** | $5.31 \times 10^{-2}$ | 1.3 | $2.54 \times 10^{-1}$ | | Promoter variant | Moderate | AT3G61710 | *ATG6* | Autophagy protein 6 |
| 3 | 22843017 | 5/G | 11.4 ± 3.8 | 76/C | 2.5 ± 0.3 | **$1.90 \times 10^{-8}$** | **37.2** | **$4.55 \times 10^{-8}$** | **14.9** | **$2.40 \times 10^{-4}$** | ✓ | Promoter variant | Moderate | | | |
| 3 | 22845358 | 18/G | 5.4 ± 1.5 | 63/A | 2.4 ± 0.4 | **$2.32 \times 10^{-3}$** | **7.6** | **$7.38 \times 10^{-3}$** | **4.7** | **$3.36 \times 10^{-2}$** | | amino acid substitution (Phe14Ser) | Moderate | AT3G61720 | *MCTP12* | Multiple c2 domain and transmembrane region protein 12 |
| 3 | 22845423 | 16/G | 5.8 ± 1.6 | 65/A | 2.4 ± 0.4 | **$8.02 \times 10^{-4}$** | **9.6** | **$2.74 \times 10^{-3}$** | **5.1** | **$2.71 \times 10^{-2}$** | | Promoter variant | Moderate | | | |
| 3 | 22846770 | 3/C | 3.2 ± 2.3 | 78/CA | 3 ± 0.5 | $9.54 \times 10^{-1}$ | 0.0 | $9.29 \times 10^{-1}$ | 0.0 | $8.62 \times 10^{-1}$ | | frameshift variant (Cys6) | **High** | AT3G61723 | | PHD finger protein |
| 3 | 22847503 | 9/T | 4.8 ± 1.8 | 72/C | 2.8 ± 0.5 | $1.46 \times 10^{-1}$ | **5.8** | **$1.89 \times 10^{-2}$** | **13** | **$5.57 \times 10^{-4}$** | | Promoter variant | Moderate | | | |
| 3 | 22849190 | 1/G | 12.7 ± 0 | 80/A | 2.9 ± 0.5 | | 7.7 | $6.94 \times 10^{-3}$ | 11.4 | $1.15 \times 10^{-3}$ | | splice donor variant | **High** | AT3G61730 | *RMF1* | Reduced male fertility |
| 3 | 22850995 | 77/A | 3 ± 0.5 | 4/C | 3 ± 1.3 | $9.66 \times 10^{-1}$ | 0 | $8.63 \times 10^{-1}$ | 0.1 | $7.68 \times 10^{-1}$ | | Promoter variant | Moderate | | | |
| 3 | 22856400 | 10/C | 4.2 ± 1.6 | 71/T | 2.9 ± 0.5 | $2.88 \times 10^{-1}$ | 3 | $8.75 \times 10^{-2}$ | **8.3** | **$5.14 \times 10^{-3}$** | | amino acid substitution (Lys50Arg) | Moderate | AT3G61740 | *ATX3* | Arabidopsis trithorax 3 |
| 3 | 22856657 | 5/G | 5.6 ± 2.9 | 76/A | 2.9 ± 0.5 | $1.21 \times 10^{-1}$ | **4.4** | **$3.91 \times 10^{-2}$** | **7** | **$9.77 \times 10^{-3}$** | | 5'-UTR variant | Moderate | | | |
| 3 | 22856986 | 9/G | 3.3 ± 1.5 | 72/C | 3 ± 0.5 | $8.06 \times 10^{-1}$ | 0.8 | $3.62 \times 10^{-1}$ | **3.4** | $6.85 \times 10^{-2}$ | | Promoter variant | Moderate | | | |
| 3 | 22857378 | 16/C | 3.8 ± 0.7 | 65/G | 2.9 ± 0.5 | $3.92 \times 10^{-1}$ | 0 | $8.27 \times 10^{-1}$ | 0.3 | $5.85 \times 10^{-1}$ | | Promoter variant | Moderate | | | |
| 4 | 1511512 | 52/C | 3.4 ± 0.6 | 29/T | 2.4 ± 0.6 | $2.17 \times 10^{-1}$ | 2 | $1.60 \times 10^{-1}$ | **3.2** | $7.67 \times 10^{-2}$ | | Promoter variant | Moderate | AT4G03420 | | Hypothetical protein (DUF789) |
| 4 | 1511544 | 41/C | 3.1 ± 0.7 | 40/T | 3 ± 0.6 | $9.62 \times 10^{-1}$ | 0 | $9.63 \times 10^{-1}$ | 0.9 | $3.38 \times 10^{-1}$ | | Promoter variant | Moderate | | | |
| 4 | 1511929 | 70/T | 3.2 ± 0.5 | 11/A | 2.2 ± 0.8 | $4.51 \times 10^{-1}$ | 0.6 | $4.27 \times 10^{-1}$ | 0.2 | $6.34 \times 10^{-1}$ | | 5'-UTR variant | Moderate | | | |
| 4 | 1512143 | 15/C | 3.3 ± 1 | 66/CT | 3 ± 0.5 | $7.55 \times 10^{-1}$ | 0 | $9.21 \times 10^{-1}$ | 0.5 | $4.74 \times 10^{-1}$ | | 5'-UTR variant | Moderate | | | |
| 4 | 1516897 | 2/T | 14.1 ± 9.3 | 79/G | 2.8 ± 0.4 | **$8.71 \times 10^{-6}$** | **21.6** | **$1.47 \times 10^{-5}$** | 0.2 | $6.41 \times 10^{-1}$ | ✓ | intron variant | Moderate | AT4G03430 | *EMB2770* | Embryo defective 2770 |
| 4 | 1520259 | 61/G | 3.2 ± 0.6 | 20/A | 2.4 ± 0.7 | $3.98 \times 10^{-1}$ | 1.9 | $1.74 \times 10^{-1}$ | **4.5** | **$3.76 \times 10^{-2}$** | | amino acid substitution (Leu81Pro) | Moderate | | | |

| | | | | | | | | | | | | | | | | |
|---|---|---|---|---|---|---|---|---|---|---|---|---|---|---|---|---|
| 4 | 1520884 | 62/G | 3.1 ± 0.5 | 17/A | 3 ± 0.9 | $8.85 \times 10^{-1}$ | 0.3 | $8.02 \times 10^{-1}$ | 0.1 | $9.36 \times 10^{-1}$ | | Promoter variant | Moderate | | | |
| 4 | 1520985 | 12/G | 3.2 ± 1.2 | 69/C | 3 ± 0.5 | $8.69 \times 10^{-1}$ | 0 | $9.72 \times 10^{-1}$ | 0 | $8.59 \times 10^{-1}$ | | Promoter variant | Moderate | | | |
| 4 | 1521119 | 11/T | 3.3 ± 1.3 | 70/C | 3 ± 0.5 | $8.04 \times 10^{-1}$ | 0 | $9.59 \times 10^{-1}$ | 0 | $8.56 \times 10^{-1}$ | | Promoter variant | Moderate | | | |
| 4 | 2739972 | 14/A | 4 ± 1.3 | 67/C | 2.9 ± 0.5 | $3.20 \times 10^{-1}$ | 0.2 | $6.57 \times 10^{-1}$ | 0.7 | $3.90 \times 10^{-1}$ | | amino acid substitution (Ser67Ala) | Moderate | AT4G05390 | *RFNR1* | Root FNR1 |
| 4 | 2740784 | 8/A | 4 ± 1.6 | 73/T | 2.9 ± 0.5 | $4.39 \times 10^{-1}$ | 0.4 | $5.44 \times 10^{-1}$ | 1.6 | $2.08 \times 10^{-1}$ | | Promoter variant | Moderate | | | |
| 4 | 2740856 | 3/A | 11 ± 6.4 | 78/T | 2.7 ± 0.4 | **$1.05 \times 10^{-4}$** | **19** | **$4.18 \times 10^{-5}$** | 1.8 | $1.86 \times 10^{-1}$ | ✓ | 5'-UTR variant | Moderate | | | |
| 4 | 2741285 | 4/A | 6.8 ± 5.6 | 77/C | 2.8 ± 0.4 | **$3.69 \times 10^{-2}$** | **7.4** | **$8.04 \times 10^{-3}$** | 0.2 | $6.76 \times 10^{-1}$ | | synonymous variant | Moderate | | | |
| 4 | 2746234 | 75/G | 3.1 ± 0.5 | 6/A | 2.1 ± 1 | $5.10 \times 10^{-1}$ | 0 | $8.93 \times 10^{-1}$ | 0 | $8.39 \times 10^{-1}$ | | 5'-UTR variant | Moderate | AT4G05420 | *DDB1A* | Damaged DNA binding protein 1a |
| 4 | 2746265 | 7/T | 4.6 ± 1.7 | 74/C | 2.9 ± 0.5 | $2.67 \times 10^{-1}$ | 0.4 | $5.08 \times 10^{-1}$ | 1.8 | $1.84 \times 10^{-1}$ | | 5'-UTR variant | Moderate | | | |
| 4 | 2748127 | 9/G | 3.2 ± 1.4 | 72/A | 3 ± 0.5 | $8.72 \times 10^{-1}$ | 0 | $9.03 \times 10^{-1}$ | 0.2 | $6.21 \times 10^{-1}$ | | amino acid substitution (Asn146Ser) | Moderate | | | |
| 5 | 2821040 | 8/G | 3.7 ± 1.6 | 73/T | 3 ± 0.5 | $6.21 \times 10^{-1}$ | 0 | $9.84 \times 10^{-1}$ | 0 | $8.57 \times 10^{-1}$ | | missense variant | Moderate | AT5G08680, | | ATP synthase alpha/beta family protein |
| 5 | 2821247 | 63/C | 3.4 ± 0.6 | 18/T | 1.9 ± 0.5 | $1.30 \times 10^{-1}$ | **3.2** | $7.87 \times 10^{-2}$ | **3.6** | $6.10 \times 10^{-2}$ | | Promoter variant | Moderate | AT5G08670 | | |
| 5 | 2821378 | 72/GTTA | 3.2 ± 0.5 | 9/G | 2.1 ± 0.9 | $4.07 \times 10^{-1}$ | 1.3 | $2.53 \times 10^{-1}$ | 1 | $3.19 \times 10^{-1}$ | | Promoter variant | Moderate | | | |
| 5 | 2821433 | 10/C | 4.4 ± 2.3 | 71/T | 2.9 ± 0.4 | $2.25 \times 10^{-1}$ | 1.1 | $2.91 \times 10^{-1}$ | 0.1 | $7.62 \times 10^{-1}$ | | Promoter variant | Moderate | | | |
| 5 | 2821724 | 27/T | 3.1 ± 1 | 54/C | 3 ± 0.5 | $9.85 \times 10^{-1}$ | 0 | $8.78 \times 10^{-1}$ | 0.7 | $4.13 \times 10^{-1}$ | | Promoter variant | Moderate | | | |
| 5 | 2821726 | 55/A | 3.1 ± 0.5 | 26/T | 3 ± 1 | $8.98 \times 10^{-1}$ | 0 | $9.66 \times 10^{-1}$ | 1.3 | $2.50 \times 10^{-1}$ | | Promoter variant | Moderate | | | |
| 5 | 2821904 | 62/A | 3.4 ± 0.6 | 19/G | 1.8 ± 0.5 | $1.08 \times 10^{-1}$ | **3.4** | $6.82 \times 10^{-2}$ | **4** | $5.06 \times 10^{-2}$ | | Promoter variant | Moderate | | | |
| 5 | 2822109 | 29/G | 3.5 ± 0.7 | 52/C | 2.8 ± 0.6 | $4.45 \times 10^{-1}$ | 0.3 | $6.16 \times 10^{-1}$ | 0.8 | $3.68 \times 10^{-1}$ | | amino acid substitution (Gly40Arg) | Moderate | AT5G08680 | | ATP synthase alpha/beta family protein |
| 5 | 2822109 | 29/G | 3.5 ± 0.7 | 52/C | 2.8 ± 0.6 | $4.45 \times 10^{-1}$ | 0.3 | $6.16 \times 10^{-1}$ | 0.8 | $3.68 \times 10^{-1}$ | | Promoter variant | Moderate | AT5G08670 | | ATP synthase alpha/beta family protein |
| 5 | 2822494 | 5/A | 8.1 ± 4.4 | 76/T | 2.7 ± 0.4 | **$1.35 \times 10^{-3}$** | **14.4** | **$3.05 \times 10^{-4}$** | **5.4** | **$2.29 \times 10^{-2}$** | ✓ | Promoter variant | Moderate | | | |

The results of a local association study (LAS) of the variation in relative fold-change (RFC) in expression levels of *HIS1-3* in 81 *A. thaliana ecotypes* in the second drought (D2) versus the first drought (D1) separated by a recovery interval, calculated in Tassel separately for the individual genomic regions, are presented. The polymorphisms in coding sequences and upstream regulatory regions (5'-UTR and promoters within 1 kb from the start codon) of genes directly associated with the top most significant ($P < 10^{-4}$) single nucleotide polymorphisms (SNPs) detected by a genome-wide association study (GWAS) (see Table 1) were downloaded from the 1001 genomes database (https://tools.1001genomes.org/polymorph/). Data of only 80 out of the 114 ecotypes used for the expression GWAS (see Figure 1, Table 1) were available in the 1001 genomes database. The frequency of the tolerant and sensitive alleles among the 81 ecotypes is shown with the average REL. The *F*-statistic and *P*-values of the generalized linear model (GLM) and the mixed linear model (MLM) methods of LAS calculations are shown. The effects of the GWAS-detected SNPs and additional polymorphisms in each gene mined from the 1001 genomes database are shown in terms of amino acid substitutions, frameshift, promoter, or 5'-UTR variations.

#### Table S9 Local association study of superinduction of *DREB1A* in 80 *Arabidopsis thaliana* ecotypes under recurrent drought

| Chromosome | Physical position | Frequency of high expression allele | Average RFC high expression allele | Frequency of low-expression allele | Average RFC low-expression allele | Student's *t*-test *P*-value | LAS GLM *F*-statistic | LAS GLM *P*-value | LAS MLM *F*-statistic | LAS MLM *P*-value | GWAS-detected SNP | Polymorphism effects | Impact | Gene ID | Gene symbol | Short description |
|---|---|---|---|---|---|---|---|---|---|---|---|---|---|---|---|---|
| 1 | 4397797 | 7/T | 3.8 ± 1.7 | 73/A | 1.5 ± 0.3 | **$6.40 \times 10^{-3}$** | **10.96** | **$1.46 \times 10^{-3}$** | **6.12** | **$1.57 \times 10^{-2}$** | ✓ | synonymous variant | Moderate | AT1G12920 | *ERF1-2* | Eukaryotic release factor 1-2 |
| 1 | 4397978 | 13/T | 1.9 ± 0.7 | 67/G | 1.6 ± 0.3 | $6.75 \times 10^{-1}$ | 0.37 | $5.45 \times 10^{-1}$ | 0.00 | $9.93 \times 10^{-1}$ | | 5'-UTR intron variant | Moderate | | | |
| 1 | 4398370 | 49/G | 1.7 ± 0.4 | 31/T | 1.6 ± 0.5 | $8.04 \times 10^{-1}$ | 0.03 | $8.56 \times 10^{-1}$ | 0.43 | $5.14 \times 10^{-1}$ | | Promoter variant | Moderate | | | |
| 1 | 4398567 | 12/G | 2 ± 0.8 | 68/A | 1.6 ± 0.3 | $5.34 \times 10^{-1}$ | 0.47 | $4.97 \times 10^{-1}$ | 0.26 | $6.12 \times 10^{-1}$ | | Promoter variant | Moderate | | | |
| 1 | 4398585 | 53/G | 1.7 ± 0.3 | 27/T | 1.7 ± 0.5 | $9.57 \times 10^{-1}$ | 0.00 | $9.59 \times 10^{-1}$ | 0.12 | $7.33 \times 10^{-1}$ | | Promoter variant | Moderate | | | |
| 1 | 21901686 | 18/A | 2 ± 0.7 | 62/G | 1.6 ± 0.3 | $4.39 \times 10^{-1}$ | 0.51 | $4.78 \times 10^{-1}$ | 0.72 | $3.99 \times 10^{-1}$ | | Promoter variant | Moderate | AT1G59620 | *CW9* | Disease resistance protein (CC-NBS-LRR class) family |
| 1 | 21901697 | 68/T | 1.8 ± 0.3 | 12/C | 1.2 ± 0.4 | $4.03 \times 10^{-1}$ | 0.83 | $3.66 \times 10^{-1}$ | 0.26 | $6.09 \times 10^{-1}$ | | Promoter variant | Moderate | | | |
| 1 | 21901702 | 69/C | 1.7 ± 0.3 | 11/G | 1.3 ± 0.5 | $4.88 \times 10^{-1}$ | 0.59 | $4.44 \times 10^{-1}$ | 0.16 | $6.94 \times 10^{-1}$ | | Promoter variant | Moderate | | | |
| 1 | 21901742 | 20/A | 2 ± 0.5 | 60/C | 1.6 ± 0.3 | $4.47 \times 10^{-1}$ | 0.51 | $4.79 \times 10^{-1}$ | 0.53 | $4.70 \times 10^{-1}$ | | Promoter variant | Moderate | | | |
| 1 | 21901766 | 13/G | 1.8 ± 0.8 | 67/A | 1.7 ± 0.3 | $8.84 \times 10^{-1}$ | 0.01 | $9.25 \times 10^{-1}$ | 0.01 | $9.33 \times 10^{-1}$ | | Promoter variant | Moderate | | | |
| 1 | 21901860 | 68/C | 1.8 ± 0.3 | 12/T | 1.1 ± 0.4 | $3.52 \times 10^{-1}$ | 1.03 | $3.13 \times 10^{-1}$ | 0.00 | $9.91 \times 10^{-1}$ | | Promoter variant | Moderate | | | |
| 1 | 21901866 | 47/G | 1.7 ± 0.4 | 33/A | 1.6 ± 0.4 | $8.31 \times 10^{-1}$ | 0.11 | $7.42 \times 10^{-1}$ | 0.92 | $3.41 \times 10^{-1}$ | | Promoter variant | Moderate | | | |
| 1 | 21901892 | 17/C | 1.9 ± 0.6 | 63/T | 1.6 ± 0.3 | $6.60 \times 10^{-1}$ | 0.12 | $7.30 \times 10^{-1}$ | 0.07 | $7.91 \times 10^{-1}$ | | Promoter variant | Moderate | | | |
| 1 | 21901949 | 14/A | 1.9 ± 0.7 | 66/T | 1.6 ± 0.3 | $6.28 \times 10^{-1}$ | 0.17 | $6.86 \times 10^{-1}$ | 0.00 | $9.96 \times 10^{-1}$ | | Promoter variant | Moderate | | | |
| 1 | 21902006 | 67/G | 1.8 ± 0.3 | 13/T | 0.9 ± 0.2 | $1.49 \times 10^{-1}$ | 1.72 | $1.94 \times 10^{-1}$ | 0.72 | $3.99 \times 10^{-1}$ | | Promoter variant | Moderate | | | |
| 1 | 21902008 | 67/T | 1.8 ± 0.3 | 13/G | 0.9 ± 0.2 | $1.49 \times 10^{-1}$ | 1.72 | $1.94 \times 10^{-1}$ | 0.72 | $3.99 \times 10^{-1}$ | | Promoter variant | Moderate | | | |
| 1 | 21902187 | 62/G | 1.8 ± 0.3 | 18/C | 1.1 ± 0.3 | $1.86 \times 10^{-1}$ | 1.25 | $2.68 \times 10^{-1}$ | 1.27 | $2.63 \times 10^{-1}$ | | Promoter variant | Moderate | | | |
| 1 | 21902207 | 64/C | 1.8 ± 0.3 | 16/T | 1.2 ± 0.3 | $2.86 \times 10^{-1}$ | 1.05 | $3.09 \times 10^{-1}$ | 2.01 | $1.61 \times 10^{-1}$ | | Promoter variant | Moderate | | | |
| 1 | 21902244 | 8/G | 2.4 ± 1.3 | 72/T | 1.6 ± 0.3 | $3.18 \times 10^{-1}$ | 0.93 | $3.39 \times 10^{-1}$ | 0.63 | $4.30 \times 10^{-1}$ | | Promoter variant | Moderate | | | |
| 1 | 21902326 | 36/C | 1.7 ± 0.4 | 44/T | 1.6 ± 0.4 | $7.79 \times 10^{-1}$ | 0.02 | $8.78 \times 10^{-1}$ | 1.58 | $2.13 \times 10^{-1}$ | | 5'-UTR variant | Moderate | | | |
| 1 | 21902394 | 6/G | 2.1 ± 0.8 | 74/GA | 1.6 ± 0.3 | $6.13 \times 10^{-1}$ | 0.19 | $6.62 \times 10^{-1}$ | 0.00 | $9.69 \times 10^{-1}$ | | 5'-UTR variant | Moderate | | | |
| 1 | 21902572 | 65/C | 1.8 ± 0.3 | 15/T | 0.9 ± 0.2 | $1.32 \times 10^{-1}$ | 2.70 | $1.05 \times 10^{-1}$ | **4.77** | **$3.21 \times 10^{-2}$** | | 5'-UTR intron variant | Moderate | | | |
| 1 | 21902738 | 13/T | 1.9 ± 0.7 | 67/C | 1.6 ± 0.3 | $6.62 \times 10^{-1}$ | 0.16 | $6.88 \times 10^{-1}$ | 1.04 | $3.11 \times 10^{-1}$ | | amino acid substitution (Leu38Ile) | Moderate | | | |
| 1 | 21902868 | 78/G | 1.7 ± 0.3 | 2/GA | 0.8 ± 0.3 | $5.71 \times 10^{-1}$ | 0.31 | $5.80 \times 10^{-1}$ | 0.21 | $6.47 \times 10^{-1}$ | | frameshift variant (Arg81_Lys82) | **High** | | | |
| 1 | 21902891 | 79/A | 1.7 ± 0.3 | 1/T | 0.5 | | 0.27 | $6.02 \times 10^{-1}$ | 0.10 | $7.56 \times 10^{-1}$ | | stop codon gain (Arg89*) | **High** | | | |
| 1 | 21902915 | 56/G | 1.9 ± 0.4 | 24/C | 1.1 ± 0.3 | $1.15 \times 10^{-1}$ | 2.20 | $1.43 \times 10^{-1}$ | 0.30 | $5.86 \times 10^{-1}$ | | amino acid substitution (Glu97Gln) | Moderate | | | |
| 1 | 21902915 | 56/G | 1.9 ± 0.4 | 24/C | 1.1 ± 0.3 | $1.15 \times 10^{-1}$ | 2.20 | $1.43 \times 10^{-1}$ | 1.41 | $2.40 \times 10^{-1}$ | | stop codon gain (Glu97*) | **High** | | | |
| 1 | 21902981 | 77/A | 1.7 ± 0.3 | 3/AT | 1.5 ± 0.6 | $8.72 \times 10^{-1}$ | 0.02 | $8.82 \times 10^{-1}$ | 0.30 | $5.83 \times 10^{-1}$ | | frameshift variant (Ile119_Ala120) | **High** | | | |

| | | | | | | | | | | | | | | | | |
|---|---|---|---|---|---|---|---|---|---|---|---|---|---|---|---|---|
| 1 | 21903649 | 8/C | 3.6 ± 1.4 | 72/T | 1.5 ± 0.3 | **$8.11 \times 10^{-3}$** | **7.06** | **$9.70 \times 10^{-3}$** | 2.66 | $1.07 \times 10^{-1}$ | | amino acid substitution (Val269Ala) | Moderate | | | |
| 1 | 21903687 | 70/C | 1.7 ± 0.3 | 10/G | 1.5 ± 0.6 | $8.36 \times 10^{-1}$ | 0.03 | $8.54 \times 10^{-1}$ | 0.01 | $9.18 \times 10^{-1}$ | | amino acid substitution (Pro282Ala) | Moderate | | | |
| 1 | 21903716 | 1/A | 9.2 ± 0 | 79/G | 1.6 ± 0.3 | | **14.22** | **$3.31 \times 10^{-4}$** | 2.40 | $1.26 \times 10^{-1}$ | | stop codon gain (Trp291*) | **High** | | | |
| 1 | 21903726 | 1/T | 6.2 ± 0 | 79/C | 1.6 ± 0.3 | | **5.58** | **$2.09 \times 10^{-2}$** | **6.21** | **$1.50 \times 10^{-2}$** | | stop codon gain (Arg295*) | **High** | | | |
| 1 | 21903944 | 1/G | 4.4 ± 0 | 79/GC | 1.6 ± 0.3 | | 1.65 | $2.03 \times 10^{-1}$ | **3.92** | **$5.17 \times 10^{-2}$** | | frameshift variant (Gly335) | **High** | | | |
| 1 | 21904065 | 72/T | 1.7 ± 0.3 | 8/G | 1.3 ± 0.5 | $6.06 \times 10^{-1}$ | 0.36 | $5.49 \times 10^{-1}$ | 2.58 | $1.12 \times 10^{-1}$ | | amino acid substitution (Ile375Met) | Moderate | | | |
| 1 | 21904087 | 11/A | 2.2 ± 0.8 | 69/C | 1.6 ± 0.3 | $3.65 \times 10^{-1}$ | 0.63 | $4.30 \times 10^{-1}$ | 0.47 | $4.96 \times 10^{-1}$ | | amino acid substitution (Leu383Met) | Moderate | | | |
| 1 | 21904156 | 79/G | 1.7 ± 0.3 | 1/T | 1.1 | | 0.08 | $7.76 \times 10^{-1}$ | 0.07 | $7.99 \times 10^{-1}$ | | stop codon gain (Glu406*) | **High** | | | |
| 1 | 21904225 | 1/T | 3.8 | 79/C | 1.6 ± 0.3 | | 0.88 | $3.52 \times 10^{-1}$ | 0.06 | $8.03 \times 10^{-1}$ | | stop codon gain (Arg429*) | **High** | | | |
| 1 | 21904247 | 78/TAG | 1.7 ± 0.3 | 2/T | 0.8 ± 0.4 | $5.50 \times 10^{-1}$ | 0.46 | $4.98 \times 10^{-1}$ | **4.36** | **$4.03 \times 10^{-2}$** | | frameshift variant (Ile436) | **High** | | | |
| 1 | 21904281 | 1/T | 2.3 ± 0 | 79/TG | 1.7 ± 0.3 | | 0.08 | $7.73 \times 10^{-1}$ | 0.20 | $6.52 \times 10^{-1}$ | | frameshift variant (Glu448) | **High** | | | |
| 1 | 21904429 | 10/A | 1.7 ± 0.8 | 70/G | 1.7 ± 0.3 | $9.37 \times 10^{-1}$ | 0.00 | $9.62 \times 10^{-1}$ | 0.00 | $1.00 \times 10^{-0}$ | | amino acid substitution (Gly497Arg) | Moderate | | | |
| 1 | 21904433 | 9/T | 1.8 ± 0.9 | 71/G | 1.7 ± 0.3 | $8.25 \times 10^{-1}$ | 0.04 | $8.44 \times 10^{-1}$ | 0.01 | $9.40 \times 10^{-1}$ | | amino acid substitution (Gly498Val) | Moderate | | | |
| 1 | 21904543 | 6/T | 2 ± 1.1 | 74/C | 1.6 ± 0.3 | $6.87 \times 10^{-1}$ | 1.42 | $2.37 \times 10^{-1}$ | 1.88 | $1.75 \times 10^{-1}$ | | stop codon gain (Gln535*) | **High** | | | |
| 1 | 21904676 | 18/G | 1.8 ± 0.6 | 62/A | 1.6 ± 0.3 | $7.09 \times 10^{-1}$ | 0.49 | $4.87 \times 10^{-1}$ | 0.26 | $6.13 \times 10^{-1}$ | | amino acid substitution (Gln579Arg) | Moderate | | | |
| 1 | 21904705 | 1/C | 4.4 | 79/CTA | 1.6 ± 0.3 | | 1.50 | $2.25 \times 10^{-1}$ | **4.87** | **$3.05 \times 10^{-2}$** | | frameshift variant (Leu589) | **High** | | | |
| 1 | 21904797 | 70/T | 1.7 ± 0.3 | 10/A | 1.5 ± 0.5 | $8.21 \times 10^{-1}$ | 0.12 | $7.33 \times 10^{-1}$ | 0.13 | $7.24 \times 10^{-1}$ | | amino acid substitution (Asp619Glu) | Moderate | | | |
| 1 | 21904874 | 32/T | 1.9 ± 0.4 | 48/TG | 1.6 ± 0.4 | $5.45 \times 10^{-1}$ | 0.27 | $6.04 \times 10^{-1}$ | 0.30 | $5.85 \times 10^{-1}$ | | frameshift variant (Trp626) | **High** | | | |
| 1 | 21904938 | 33/A | 2.2 ± 0.6 | 47/C | 1.3 ± 0.2 | **$6.51 \times 10^{-2}$** | **3.72** | **$5.78 \times 10^{-2}$** | **5.03** | **$2.80 \times 10^{-2}$** | | amino acid substitution (Asn647Thr) | Moderate | | | |
| 1 | 21905080 | 79/A | 1.7 ± 0.3 | 1/AAT | 0.8 | | 0.18 | $6.74 \times 10^{-1}$ | 1.34 | $2.50 \times 10^{-1}$ | | frameshift variant (Ile695_Tyr696) | **High** | | | |
| 1 | 21905286 | 77/A | 1.7 ± 0.3 | 3/AT | 0.8 ± 0.3 | $4.91 \times 10^{-1}$ | 0.54 | $4.66 \times 10^{-1}$ | 0.82 | $3.67 \times 10^{-1}$ | | frameshift variant (Asp763_Leu764) | **High** | | | |
| 1 | 21905349 | 13/G | 3.9 ± 1.1 | 67/A | 1.2 ± 0.2 | **$2.65 \times 10^{-5}$** | **18.61** | **$5.03 \times 10^{-5}$** | **6.71** | **$1.16 \times 10^{-2}$** | ✓ | synonymous variant | Moderate | | | |
| 1 | 21905361 | 75/T | 1.7 ± 0.3 | 5/TC | 0.9 ± 0.3 | $4.07 \times 10^{-1}$ | 0.73 | $3.95 \times 10^{-1}$ | **5.11** | **$2.68 \times 10^{-2}$** | | frameshift variant (Ile788_Arg789) | **High** | | | |
| 1 | 24985508 | 66/A | 1.8 ± 0.3 | 14/G | 1.2 ± 0.4 | $3.71 \times 10^{-1}$ | 0.82 | $3.70 \times 10^{-1}$ | 0.51 | $4.77 \times 10^{-1}$ | | amino acid substitution (Ile701Thr) | Moderate | AT1G66960 | *LUP5* | Terpenoid cyclases family protein |
| 1 | 24985672 | 42/C | 2 ± 0.4 | 38/G | 1.3 ± 0.3 | $1.90 \times 10^{-1}$ | 1.36 | $2.47 \times 10^{-1}$ | 1.01 | $3.18 \times 10^{-1}$ | | amino acid substitution (Ala689Gly) | Moderate | | | |
| 1 | 24986119 | 66/A | 1.8 ± 0.3 | 14/T | 1.1 ± 0.4 | $2.78 \times 10^{-1}$ | 1.34 | $2.50 \times 10^{-1}$ | 0.01 | $9.41 \times 10^{-1}$ | | amino acid substitution (Leu604Met) | Moderate | | | |
| 1 | 24986125 | 66/T | 1.8 ± 0.3 | 14/C | 1.1 ± 0.4 | $2.78 \times 10^{-1}$ | 1.34 | $2.50 \times 10^{-1}$ | 0.01 | $9.41 \times 10^{-1}$ | | amino acid substitution (Lys602Glu) | Moderate | | | |
| 1 | 24986185 | 68/C | 1.8 ± 0.3 | 12/T | 1.1 ± 0.4 | $3.34 \times 10^{-1}$ | 1.02 | $3.17 \times 10^{-1}$ | 0.00 | $9.73 \times 10^{-1}$ | | amino acid substitution (Asp582Asn) | Moderate | | | |
| 1 | 24986197 | 70/G | 1.8 ± 0.3 | 10/T | 1.1 ± 0.3 | $3.54 \times 10^{-1}$ | 1.02 | $3.16 \times 10^{-1}$ | 1.33 | $2.52 \times 10^{-1}$ | | amino acid substitution (Gln578Lys) | Moderate | | | |
| 1 | 24986198 | 44/T | 1.9 ± 0.4 | 36/A | 1.4 ± 0.3 | $2.35 \times 10^{-1}$ | 1.78 | $1.87 \times 10^{-1}$ | 0.74 | $3.91 \times 10^{-1}$ | | amino acid substitution (Asn577Lys) | Moderate | | | |
| 1 | 24986592 | 68/C | 1.8 ± 0.3 | 12/T | 1 ± 0.4 | $2.50 \times 10^{-1}$ | 1.60 | $2.10 \times 10^{-1}$ | 0.13 | $7.23 \times 10^{-1}$ | | amino acid substitution (Glu508Lys) | Moderate | | | |
| 1 | 24986597 | 68/G | 1.8 ± 0.3 | 12/A | 1 ± 0.4 | $2.50 \times 10^{-1}$ | 1.60 | $2.10 \times 10^{-1}$ | 0.13 | $7.23 \times 10^{-1}$ | | amino acid substitution (Thr506Ile) | Moderate | | | |
| 1 | 24986603 | 11/A | 1.9 ± 0.7 | 69/T | 1.6 ± 0.3 | $6.82 \times 10^{-1}$ | 0.12 | $7.25 \times 10^{-1}$ | 0.03 | $8.70 \times 10^{-1}$ | | amino acid substitution (Asp504Val) | Moderate | | | |
| 1 | 24986802 | 24/A | 2.9 ± 0.7 | 56/G | 1.1 ± 0.2 | **$6.03 \times 10^{-4}$** | **15.40** | **$1.97 \times 10^{-4}$** | **6.13** | **$1.56 \times 10^{-2}$** | ✓ | synonymous variant | Moderate | | | |
| 1 | 24986835 | 23/T | 1.7 ± 0.5 | 57/G | 1.7 ± 0.3 | $9.25 \times 10^{-1}$ | 0.09 | $7.61 \times 10^{-1}$ | 0.70 | $4.07 \times 10^{-1}$ | | amino acid substitution (Asp455Glu) | Moderate | | | |

| | | | | | | | | | | | | | | | |
|---|---|---|---|---|---|---|---|---|---|---|---|---|---|---|---|
| 1 | 24987392 | 51/C | 1.7 ± 0.3 | 29/T | 1.6 ± 0.5 | $8.18 \times 10^{-1}$ | 0.11 | $7.40 \times 10^{-1}$ | 0.18 | $6.69 \times 10^{-1}$ | | amino acid substitution (Asn389Asp) | Moderate | | |
| 1 | 24987635 | 23/G | 1.8 ± 0.5 | 57/C | 1.6 ± 0.3 | $7.09 \times 10^{-1}$ | 0.10 | $7.49 \times 10^{-1}$ | 0.12 | $7.32 \times 10^{-1}$ | | amino acid substitution (Leu335Phe) | Moderate | | |
| 1 | 24988110 | 10/T | 2.3 ± 0.9 | 70/A | 1.6 ± 0.3 | $3.14 \times 10^{-1}$ | 0.98 | $3.26 \times 10^{-1}$ | 0.53 | $4.67 \times 10^{-1}$ | | amino acid substitution (Asp232Glu) | Moderate | | |
| 1 | 24988250 | 42/G | 1.8 ± 0.4 | 38/A | 1.5 ± 0.4 | $4.81 \times 10^{-1}$ | 0.34 | $5.64 \times 10^{-1}$ | 0.37 | $5.46 \times 10^{-1}$ | | amino acid substitution (Ile215Thr) | Moderate | | |
| 1 | 24989501 | 70/G | 1.7 ± 0.3 | 10/A | 1.5 ± 0.8 | $8.11 \times 10^{-1}$ | 0.09 | $7.66 \times 10^{-1}$ | 0.14 | $7.14 \times 10^{-1}$ | | amino acid substitution (Ser55Phe) | Moderate | | |
| 1 | 24989680 | 17/G | 1.9 ± 0.6 | 63/A | 1.6 ± 0.3 | $6.11 \times 10^{-1}$ | 0.70 | $4.04 \times 10^{-1}$ | 0.20 | $6.53 \times 10^{-1}$ | | 5'-UTR variant | Moderate | | |
| 1 | 24989795 | 56/T | 1.9 ± 0.4 | 24/TG | 1.2 ± 0.3 | $2.09 \times 10^{-1}$ | 1.63 | $2.06 \times 10^{-1}$ | 0.60 | $4.41 \times 10^{-1}$ | | 5'-UTR variant | Moderate | | |
| 1 | 24989798 | 52/A | 2 ± 0.4 | 28/T | 1.1 ± 0.2 | **$6.16 \times 10^{-2}$** | **3.04** | **$8.56 \times 10^{-2}$** | 0.15 | $7.00 \times 10^{-1}$ | | Promoter variant | Moderate | | |
| 1 | 24989799 | 49/A | 2 ± 0.4 | 30/G | 1.1 ± 0.2 | **$7.87 \times 10^{-2}$** | 2.23 | $1.15 \times 10^{-1}$ | **3.06** | **$5.32 \times 10^{-2}$** | | Promoter variant | Moderate | | |
| 1 | 24989826 | 35/G | 1.8 ± 0.4 | 45/T | 1.6 ± 0.4 | $6.26 \times 10^{-1}$ | 0.44 | $5.09 \times 10^{-1}$ | 0.07 | $7.91 \times 10^{-1}$ | | Promoter variant | Moderate | | |
| 1 | 24989877 | 24/A | 1.8 ± 0.5 | 56/G | 1.6 ± 0.3 | $7.27 \times 10^{-1}$ | 0.32 | $5.74 \times 10^{-1}$ | 0.07 | $7.97 \times 10^{-1}$ | | Promoter variant | Moderate | | |
| 1 | 24990058 | 49/C | 1.9 ± 0.4 | 31/T | 1.3 ± 0.3 | $2.55 \times 10^{-1}$ | 1.05 | $3.08 \times 10^{-1}$ | 0.39 | $5.37 \times 10^{-1}$ | | Promoter variant | Moderate | | |
| 1 | 24990110 | 44/T | 1.9 ± 0.4 | 36/A | 1.3 ± 0.3 | $2.10 \times 10^{-1}$ | 1.27 | $2.64 \times 10^{-1}$ | 1.38 | $2.45 \times 10^{-1}$ | | Promoter variant | Moderate | | |
| 1 | 24990188 | 47/G | 1.9 ± 0.4 | 33/T | 1.3 ± 0.3 | $2.08 \times 10^{-1}$ | 1.23 | $2.71 \times 10^{-1}$ | 1.05 | $3.09 \times 10^{-1}$ | | Promoter variant | Moderate | | |
| 1 | 24990232 | 70/C | 1.7 ± 0.3 | 10/T | 1.5 ± 0.8 | $8.11 \times 10^{-1}$ | 0.09 | $7.66 \times 10^{-1}$ | 0.14 | $7.14 \times 10^{-1}$ | | Promoter variant | Moderate | | |
| 1 | 24990268 | 15/G | 1.9 ± 0.7 | 65/C | 1.6 ± 0.3 | $6.18 \times 10^{-1}$ | 0.18 | $6.73 \times 10^{-1}$ | 0.60 | $4.43 \times 10^{-1}$ | | Promoter variant | Moderate | | |
| 1 | 24990274 | 55/C | 1.8 ± 0.4 | 25/G | 1.4 ± 0.3 | $4.57 \times 10^{-1}$ | 0.70 | $4.05 \times 10^{-1}$ | 0.08 | $7.72 \times 10^{-1}$ | | Promoter variant | Moderate | | |
| 1 | 24990288 | 67/A | 1.8 ± 0.3 | 13/G | 0.9 ± 0.2 | $1.79 \times 10^{-1}$ | 1.16 | $2.85 \times 10^{-1}$ | 1.15 | $2.88 \times 10^{-1}$ | | Promoter variant | Moderate | | |
| 1 | 24990538 | 29/T | 1.8 ± 0.5 | 51/A | 1.6 ± 0.3 | $6.05 \times 10^{-1}$ | 0.48 | $4.91 \times 10^{-1}$ | 0.52 | $4.71 \times 10^{-1}$ | | Promoter variant | Moderate | | |
| 1 | 25004218 | 12/G | 2.5 ± 1 | 68/C | 1.5 ± 0.3 | $1.35 \times 10^{-1}$ | 2.00 | $1.61 \times 10^{-1}$ | 0.03 | $8.52 \times 10^{-1}$ | | stop codon lost (Ter893Serext*) | **High** | AT1G67000 | Protein kinase superfamily protein |
| 1 | 25004491 | 6/G | 4.6 ± 1.8 | 74/C | 1.4 ± 0.3 | **$3.93 \times 10^{-4}$** | **14.18** | **$3.37 \times 10^{-4}$** | 1.23 | $2.70 \times 10^{-1}$ | ✓ | amino acid substitution (Gly802Ala) | Moderate | | |
| 1 | 25004494 | 21/A | 2 ± 0.6 | 59/T | 1.6 ± 0.3 | $4.72 \times 10^{-1}$ | 0.39 | $5.35 \times 10^{-1}$ | 0.02 | $8.75 \times 10^{-1}$ | | amino acid substitution (Asn801Ile) | Moderate | | |
| 1 | 25004536 | 45/A | 1.8 ± 0.4 | 35/T | 1.5 ± 0.4 | $6.13 \times 10^{-1}$ | 0.14 | $7.09 \times 10^{-1}$ | 0.18 | $6.75 \times 10^{-1}$ | | amino acid substitution (Ile787Asn) | Moderate | | |
| 1 | 25004540 | 78/TGGCC | 1.7 ± 0.3 | 2/T | 1.3 ± 1.1 | $7.98 \times 10^{-1}$ | 0.09 | $7.69 \times 10^{-1}$ | 0.12 | $7.34 \times 10^{-1}$ | | frameshift variant (Lys784) | **High** | | |
| 1 | 25004545 | 1/T | 2.3 | 79/TTCTCAAG | 1.7 ± 0.3 | | 0.07 | $7.88 \times 10^{-1}$ | 0.74 | $3.93 \times 10^{-1}$ | | frameshift variant (Leu782) | **High** | | |
| 1 | 25004599 | 60/G | 1.7 ± 0.3 | 20/C | 1.7 ± 0.6 | $9.74 \times 10^{-1}$ | 0.00 | $9.47 \times 10^{-1}$ | 1.16 | $2.86 \times 10^{-1}$ | | amino acid substitution (Arg766Pro) | Moderate | | |
| 1 | 25004615 | 58/A | 1.7 ± 0.3 | 22/C | 1.6 ± 0.5 | $7.91 \times 10^{-1}$ | 0.13 | $7.21 \times 10^{-1}$ | 0.00 | $9.82 \times 10^{-1}$ | | amino acid substitution (Phe761Val) | Moderate | | |
| 1 | 25004962 | 54/A | 1.7 ± 0.4 | 5/T | 1.3 ± 0.5 | $6.55 \times 10^{-1}$ | 0.15 | $8.63 \times 10^{-1}$ | 0.27 | $7.65 \times 10^{-1}$ | | amino acid substitution (Ser645Ile) | Moderate | | |
| 1 | 25004980 | 55/C | 1.9 ± 0.4 | 25/T | 1.2 ± 0.3 | $2.25 \times 10^{-1}$ | 1.71 | $1.95 \times 10^{-1}$ | 0.01 | $9.15 \times 10^{-1}$ | | amino acid substitution (Lys639Arg) | Moderate | | |
| 1 | 25005234 | 69/C | 1.8 ± 0.3 | 11/A | 1.1 ± 0.3 | $3.76 \times 10^{-1}$ | 0.94 | $3.36 \times 10^{-1}$ | 0.29 | $5.93 \times 10^{-1}$ | | amino acid substitution (Met554Ile) | Moderate | | |
| 1 | 25006072 | 49/C | 1.8 ± 0.4 | 31/A | 1.5 ± 0.3 | $4.89 \times 10^{-1}$ | 0.73 | $3.94 \times 10^{-1}$ | 1.22 | $2.74 \times 10^{-1}$ | | amino acid substitution (Arg423Ser) | Moderate | | |
| 1 | 25006124 | 55/C | 1.7 ± 0.4 | 25/G | 1.6 ± 0.4 | $7.52 \times 10^{-1}$ | 0.22 | $6.44 \times 10^{-1}$ | 1.01 | $3.17 \times 10^{-1}$ | | amino acid substitution (Arg406Thr) | Moderate | | |
| 1 | 25006141 | 10/A | 1.8 ± 1.1 | 70/C | 1.6 ± 0.3 | $8.08 \times 10^{-1}$ | 0.04 | $8.47 \times 10^{-1}$ | 0.01 | $9.42 \times 10^{-1}$ | | amino acid substitution (Leu400Phe) | Moderate | | |
| 1 | 25006262 | 76/GATGAA | 1.7 ± 0.3 | 4/G | 1.1 ± 0.5 | $5.87 \times 10^{-1}$ | 0.13 | $7.15 \times 10^{-1}$ | 0.12 | $7.28 \times 10^{-1}$ | | frameshift variant (Asn358) | **High** | | |
| 1 | 25006272 | 74/AACGG | 1.7 ± 0.3 | 6/A | 1.1 ± 0.6 | $5.21 \times 10^{-1}$ | 0.05 | $8.30 \times 10^{-1}$ | 0.09 | $7.65 \times 10^{-1}$ | | frameshift variant (Cys355) | **High** | | |

| | | | | | | | | | | | | | | | | |
|---|---|---|---|---|---|---|---|---|---|---|---|---|---|---|---|---|
| 1 | 25006285 | 2/C | 2.3 ± 0.1 | 78/G | 1.7 ± 0.3 | $6.81 \times 10^{-1}$ | 0.17 | $6.86 \times 10^{-1}$ | 1.27 | $2.64 \times 10^{-1}$ | | stop codon gain (Tyr352*) | **High** | | | |
| 1 | 25006355 | 46/T | 1.9 ± 0.4 | 34/G | 1.3 ± 0.3 | $2.32 \times 10^{-1}$ | 1.83 | $1.81 \times 10^{-1}$ | 1.65 | $2.03 \times 10^{-1}$ | | amino acid substitution (Thr329Asn) | Moderate | | | |
| 1 | 25006541 | 65/T | 1.7 ± 0.3 | 14/C | 1.4 ± 0.4 | $5.96 \times 10^{-1}$ | 0.37 | $6.90 \times 10^{-1}$ | 1.78 | $1.77 \times 10^{-1}$ | | amino acid substitution (Asn267Ser) | Moderate | | | |
| 1 | 25006593 | 55/C | 1.7 ± 0.4 | 25/G | 1.6 ± 0.4 | $8.91 \times 10^{-1}$ | 0.00 | $9.49 \times 10^{-1}$ | 0.02 | $9.02 \times 10^{-1}$ | | amino acid substitution (Val250Leu) | Moderate | | | |
| 1 | 25006601 | 17/C | 1.9 ± 0.6 | 63/G | 1.6 ± 0.3 | $6.97 \times 10^{-1}$ | 0.47 | $4.97 \times 10^{-1}$ | 2.74 | $1.02 \times 10^{-1}$ | | amino acid substitution (Ala247Gly) | Moderate | | | |
| 1 | 25006871 | 34/G | 1.9 ± 0.4 | 46/T | 1.5 ± 0.4 | $4.61 \times 10^{-1}$ | 0.77 | $3.82 \times 10^{-1}$ | 0.90 | $3.45 \times 10^{-1}$ | | amino acid substitution (His245Pro) | Moderate | | | |
| 1 | 25006877 | 34/G | 1.9 ± 0.4 | 46/T | 1.5 ± 0.4 | $4.61 \times 10^{-1}$ | 0.77 | $3.82 \times 10^{-1}$ | 0.90 | $3.45 \times 10^{-1}$ | | amino acid substitution (Tyr243Ser) | Moderate | | | |
| 1 | 25007070 | 57/T | 1.8 ± 0.3 | 23/G | 1.4 ± 0.5 | $5.36 \times 10^{-1}$ | 0.50 | $4.82 \times 10^{-1}$ | 0.06 | $8.02 \times 10^{-1}$ | | amino acid substitution (His179Asn) | Moderate | | | |
| 1 | 25007127 | 44/G | 1.9 ± 0.4 | 36/C | 1.5 ± 0.3 | $4.09 \times 10^{-1}$ | 0.92 | $3.41 \times 10^{-1}$ | 0.08 | $7.82 \times 10^{-1}$ | | amino acid substitution (Val160Leu) | Moderate | | | |
| 1 | 25007202 | 63/G | 1.8 ± 0.3 | 17/A | 1.4 ± 0.4 | $4.96 \times 10^{-1}$ | 0.63 | $4.30 \times 10^{-1}$ | 1.35 | $2.49 \times 10^{-1}$ | | amino acid substitution (His135Tyr) | Moderate | | | |
| 1 | 25007207 | 26/C | 1.7 ± 0.4 | 54/A | 1.7 ± 0.4 | $9.82 \times 10^{-1}$ | 0.01 | $9.16 \times 10^{-1}$ | 0.00 | $9.74 \times 10^{-1}$ | | amino acid substitution (Leu133Trp) | Moderate | | | |
| 1 | 25007232 | 26/A | 1.7 ± 0.4 | 54/G | 1.7 ± 0.4 | $9.82 \times 10^{-1}$ | 0.01 | $9.16 \times 10^{-1}$ | 0.00 | $9.74 \times 10^{-1}$ | | amino acid substitution (Pro125Ser) | Moderate | | | |
| 1 | 25007249 | 53/T | 1.7 ± 0.3 | 27/C | 1.6 ± 0.5 | $8.73 \times 10^{-1}$ | 0.06 | $8.05 \times 10^{-1}$ | 0.51 | $4.79 \times 10^{-1}$ | | amino acid substitution (Glu119Gly) | Moderate | | | |
| 1 | 25007257 | 18/A | 2.1 ± 0.8 | 62/C | 1.5 ± 0.3 | $3.35 \times 10^{-1}$ | 1.80 | $1.84 \times 10^{-1}$ | 0.08 | $7.83 \times 10^{-1}$ | | amino acid substitution (Leu116Phe) | Moderate | | | |
| 1 | 25007561 | 67/A | 1.8 ± 0.3 | 13/T | 1.2 ± 0.4 | $3.87 \times 10^{-1}$ | 0.99 | $3.24 \times 10^{-1}$ | 1.30 | $2.57 \times 10^{-1}$ | | amino acid substitution (Leu15His) | Moderate | | | |
| 1 | 25007592 | 10/G | 2.7 ± 1.1 | 70/A | 1.5 ± 0.3 | $\mathbf{9.93 \times 10^{-2}}$ | **4.75** | $\mathbf{3.25 \times 10^{-2}}$ | 0.08 | $7.78 \times 10^{-1}$ | | amino acid substitution (Ser5Pro) | Moderate | | | |
| 1 | 25007598 | 24/T | 1.8 ± 0.6 | 56/A | 1.6 ± 0.3 | $7.81 \times 10^{-1}$ | 0.05 | $8.29 \times 10^{-1}$ | 0.15 | $7.04 \times 10^{-1}$ | | amino acid substitution (Tyr3Asn) | Moderate | | | |
| 1 | 25007634 | 29/T | 1.7 ± 0.5 | 51/G | 1.6 ± 0.3 | $8.23 \times 10^{-1}$ | 0.18 | $6.70 \times 10^{-1}$ | 0.03 | $8.67 \times 10^{-1}$ | | 5'-UTR variant | Moderate | | | |
| 1 | 25007685 | 43/C | 1.8 ± 0.4 | 37/G | 1.6 ± 0.4 | $7.02 \times 10^{-1}$ | 0.25 | $6.21 \times 10^{-1}$ | 0.24 | $6.22 \times 10^{-1}$ | | Promoter variant | Moderate | | | |
| 1 | 25007773 | 19/A | 2 ± 0.7 | 61/G | 1.6 ± 0.3 | $4.72 \times 10^{-1}$ | 0.42 | $5.17 \times 10^{-1}$ | 1.26 | $2.66 \times 10^{-1}$ | | Promoter variant | Moderate | | | |
| 1 | 25007988 | 43/C | 1.9 ± 0.4 | 37/A | 1.4 ± 0.3 | $3.61 \times 10^{-1}$ | 1.06 | $3.07 \times 10^{-1}$ | 0.02 | $8.83 \times 10^{-1}$ | | Promoter variant | Moderate | | | |
| 1 | 25007990 | 65/T | 1.8 ± 0.3 | 15/A | 1.1 ± 0.4 | $2.64 \times 10^{-1}$ | 1.56 | $2.15 \times 10^{-1}$ | 0.67 | $4.16 \times 10^{-1}$ | | Promoter variant | Moderate | | | |
| 1 | 25008188 | 66/A | 1.8 ± 0.3 | 14/T | 1 ± 0.2 | $1.69 \times 10^{-1}$ | 2.37 | $1.28 \times 10^{-1}$ | 1.80 | $1.83 \times 10^{-1}$ | | Promoter variant | Moderate | | | |
| 1 | 25008212 | 61/T | 1.7 ± 0.4 | 19/C | 1.6 ± 0.3 | $9.03 \times 10^{-1}$ | 0.05 | $8.24 \times 10^{-1}$ | 0.01 | $9.12 \times 10^{-1}$ | | Promoter variant | Moderate | | | |
| 1 | 25008370 | 69/C | 1.7 ± 0.3 | 11/T | 1.6 ± 0.8 | $9.40 \times 10^{-1}$ | 0.01 | $9.10 \times 10^{-1}$ | 0.00 | $9.44 \times 10^{-1}$ | | Promoter variant | Moderate | | | |
| 3 | 17723736 | 25/C | 1.7 ± 0.4 | 55/A | 1.6 ± 0.4 | $8.80 \times 10^{-1}$ | 0.00 | $9.67 \times 10^{-1}$ | 0.69 | $4.10 \times 10^{-1}$ | | missense variant | Moderate | AT3G48020 | | Hypothetical protein |
| 3 | 17723993 | 55/G | 1.8 ± 0.3 | 25/A | 1.5 ± 0.5 | $6.29 \times 10^{-1}$ | 0.41 | $5.23 \times 10^{-1}$ | 0.23 | $6.34 \times 10^{-1}$ | | synonymous variant | Moderate | | | |
| 3 | 17724131 | 14/A | 2.1 ± 0.7 | 66/T | 1.6 ± 0.3 | $4.49 \times 10^{-1}$ | 0.51 | $4.78 \times 10^{-1}$ | 0.30 | $5.83 \times 10^{-1}$ | | Promoter variant | Moderate | | | |
| 3 | 17724364 | 69/T | 1.8 ± 0.3 | 11/TA | 1 ± 0.4 | $2.40 \times 10^{-1}$ | 1.27 | $2.64 \times 10^{-1}$ | 0.00 | $9.57 \times 10^{-1}$ | | Promoter variant | Moderate | | | |
| 3 | 17724516 | 19/G | 2.1 ± 0.6 | 61/A | 1.5 ± 0.3 | $2.72 \times 10^{-1}$ | 1.39 | $2.42 \times 10^{-1}$ | 0.21 | $6.46 \times 10^{-1}$ | | Promoter variant | Moderate | | | |
| 3 | 17726083 | 54/A | 2 ± 0.4 | 26/C | 1 ± 0.3 | $\mathbf{6.08 \times 10^{-2}}$ | **4.53** | $\mathbf{3.68 \times 10^{-2}}$ | 0.22 | $6.42 \times 10^{-1}$ | | amino acid substitution (Gly126Val) | Moderate | AT3G48030 | *ATL48* | Arabidopsis toxicos en levadura 48 |
| 3 | 17726259 | 8/T | 3.5 ± 1.5 | 72/A | 1.5 ± 0.3 | $\mathbf{1.08 \times 10^{-2}}$ | **6.75** | $\mathbf{1.13 \times 10^{-2}}$ | 1.96 | $1.66 \times 10^{-1}$ | ✓ | intron variant | Moderate | | | |
| 3 | 17728032 | 18/A | 2.3 ± 0.8 | 62/G | 1.5 ± 0.3 | $1.37 \times 10^{-1}$ | 2.20 | $1.42 \times 10^{-1}$ | 0.75 | $3.89 \times 10^{-1}$ | | Promoter variant | Moderate | | | |
| 3 | 17728680 | 25/C | 1.8 ± 0.4 | 55/T | 1.6 ± 0.4 | $7.93 \times 10^{-1}$ | 0.04 | $8.45 \times 10^{-1}$ | 0.02 | $8.84 \times 10^{-1}$ | | Promoter variant | Moderate | | | |
| 3 | 17728685 | 41/C | 1.8 ± 0.4 | 39/T | 1.5 ± 0.4 | $4.77 \times 10^{-1}$ | 0.44 | $5.10 \times 10^{-1}$ | 0.08 | $7.81 \times 10^{-1}$ | | Promoter variant | Moderate | | | |

| | | | | | | | | | | | | | | | | |
|---|---|---|---|---|---|---|---|---|---|---|---|---|---|---|---|---|
| 3 | 17728812 | 47/A | 1.7 ± 0.4 | 33/G | 1.7 ± 0.4 | $9.88 \times 10^{-1}$ | 0.03 | $8.62 \times 10^{-1}$ | 0.02 | $8.99 \times 10^{-1}$ | | Promoter variant | Moderate | | | |
| 3 | 17728854 | 10/C | 2.2 ± 0.9 | 70/G | 1.6 ± 0.3 | $4.08 \times 10^{-1}$ | 0.74 | $3.94 \times 10^{-1}$ | 0.18 | $6.75 \times 10^{-1}$ | | Promoter variant | Moderate | | | |
| 3 | 17728922 | 64/A | 1.7 ± 0.3 | 16/G | 1.4 ± 0.4 | $5.19 \times 10^{-1}$ | 0.62 | $4.34 \times 10^{-1}$ | 0.11 | $7.38 \times 10^{-1}$ | | Promoter variant | Moderate | AT3G48050 | *SUO* | Shuttle' in chinese |
| 3 | 17735286 | 38/C | 2 ± 0.5 | 42/G | 1.4 ± 0.3 | $2.68 \times 10^{-1}$ | 1.12 | $2.92 \times 10^{-1}$ | 0.10 | $7.57 \times 10^{-1}$ | | amino acid substitution (Pro1276Ala) | Moderate | | | |
| 3 | 17735436 | 16/C | 2.1 ± 0.8 | 64/G | 1.6 ± 0.3 | $3.89 \times 10^{-1}$ | 0.75 | $3.89 \times 10^{-1}$ | 0.02 | $8.99 \times 10^{-1}$ | | amino acid substitution (Pro1226Ala) | Moderate | | | |
| 3 | 17736098 | 19/C | 2.3 ± 0.7 | 61/T | 1.5 ± 0.3 | $1.56 \times 10^{-1}$ | 1.90 | $1.72 \times 10^{-1}$ | 1.17 | $2.83 \times 10^{-1}$ | | amino acid substitution (Gly1005Asp) | Moderate | | | |
| 3 | 17736962 | 53/A | 1.8 ± 0.4 | 27/G | 1.5 ± 0.4 | $5.92 \times 10^{-1}$ | 0.14 | $7.06 \times 10^{-1}$ | 0.02 | $8.84 \times 10^{-1}$ | | amino acid substitution (Leu717Pro) | Moderate | | | |
| 3 | 17738487 | 49/G | 1.9 ± 0.4 | 31/C | 1.3 ± 0.3 | $1.78 \times 10^{-1}$ | 1.56 | $2.16 \times 10^{-1}$ | 0.58 | $4.48 \times 10^{-1}$ | | amino acid substitution (Pro209Ala) | Moderate | | | |
| 3 | 17739711 | 35/A | 1.8 ± 0.4 | 45/C | 1.5 ± 0.4 | $5.16 \times 10^{-1}$ | 0.36 | $5.51 \times 10^{-1}$ | 0.03 | $8.68 \times 10^{-1}$ | | Promoter variant | Moderate | | | |
| 3 | 17740073 | 23/G | 2.3 ± 0.6 | 57/T | 1.4 ± 0.3 | $9.38 \times 10^{-2}$ | 2.73 | $1.03 \times 10^{-1}$ | 2.87 | **$9.46 \times 10^{-2}$** | | Promoter variant | Moderate | | | |
| 3 | 17931230 | 6/G | 3.3 ± 1.3 | 74/A | 1.5 ± 0.3 | **$4.68 \times 10^{-2}$** | **3.70** | **$5.84 \times 10^{-2}$** | 0.71 | $4.02 \times 10^{-1}$ | | Promoter variant | Moderate | AT3G48425 | *APE1L* | DNAse I-like superfamily protein |
| 3 | 17931532 | 8/G | 1.8 ± 0.6 | 72/A | 1.7 ± 0.3 | $8.78 \times 10^{-1}$ | 0.01 | $9.41 \times 10^{-1}$ | 2.63 | $1.09 \times 10^{-1}$ | | missense variant | Moderate | | | |
| 3 | 17931556 | 7/T | 4.7 ± 1.5 | 73/C | 1.4 ± 0.2 | **$5.12 \times 10^{-5}$** | **17.09** | **$9.50 \times 10^{-5}$** | 2.84 | **$9.66 \times 10^{-2}$** | | 5'-UTR | Moderate | | | |
| 3 | 17931707 | 11/C | 1.9 ± 0.6 | 69/T | 1.6 ± 0.3 | $7.54 \times 10^{-1}$ | 0.12 | $7.32 \times 10^{-1}$ | 0.50 | $4.83 \times 10^{-1}$ | | Promoter variant | Moderate | | | |
| 3 | 17931992 | 29/T | 2.2 ± 0.6 | 51/G | 1.4 ± 0.3 | $1.31 \times 10^{-1}$ | 1.95 | $1.66 \times 10^{-1}$ | 2.13 | $1.49 \times 10^{-1}$ | | amino acid substitution (Gly28Val) | Moderate | | | |
| 3 | 17934815 | 65/T | 1.8 ± 0.3 | 15/G | 1.2 ± 0.4 | $3.80 \times 10^{-1}$ | 0.77 | $3.82 \times 10^{-1}$ | 0.74 | $3.94 \times 10^{-1}$ | | Promoter variant | Moderate | AT3G48430 | *JMJ12* | Jumonji domain-containing protein 12 |
| 3 | 17934988 | 65/C | 1.7 ± 0.3 | 15/T | 1.6 ± 0.5 | $9.65 \times 10^{-1}$ | 0.00 | $9.58 \times 10^{-1}$ | 0.35 | $5.57 \times 10^{-1}$ | | Promoter variant | Moderate | | | |
| 3 | 17935042 | 64/A | 1.7 ± 0.3 | 16/T | 1.6 ± 0.4 | $8.15 \times 10^{-1}$ | 0.08 | $7.83 \times 10^{-1}$ | 1.24 | $2.70 \times 10^{-1}$ | | Promoter variant | Moderate | | | |
| 3 | 17935050 | 70/C | 1.7 ± 0.3 | 10/T | 1.2 ± 0.5 | $4.85 \times 10^{-1}$ | 0.50 | $4.82 \times 10^{-1}$ | 0.82 | $3.68 \times 10^{-1}$ | | Promoter variant | Moderate | | | |
| 3 | 17935120 | 69/T | 1.7 ± 0.3 | 11/A | 1.4 ± 0.4 | $6.33 \times 10^{-1}$ | 0.23 | $6.31 \times 10^{-1}$ | 0.56 | $4.55 \times 10^{-1}$ | | Promoter variant | Moderate | | | |
| 3 | 17936675 | 11/C | 3.8 ± 1.2 | 69/G | 1.3 ± 0.2 | **$2.19 \times 10^{-4}$** | **14.45** | **$2.99 \times 10^{-4}$** | 2.43 | $1.23 \times 10^{-1}$ | ✓ | intron variant | Moderate | | | |
| 3 | 17937841 | 62/G | 1.7 ± 0.3 | 18/A | 1.4 ± 0.4 | $6.01 \times 10^{-1}$ | 0.29 | $5.93 \times 10^{-1}$ | 0.06 | $8.02 \times 10^{-1}$ | | amino acid substitution (Met572Ile) | Moderate | | | |
| 3 | 17938290 | 61/T | 1.8 ± 0.3 | 19/A | 1.4 ± 0.4 | $5.21 \times 10^{-1}$ | 0.43 | $5.16 \times 10^{-1}$ | 0.01 | $9.25 \times 10^{-1}$ | | amino acid substitution (Cys699Ser) | Moderate | | | |
| 3 | 17940034 | 62/A | 1.7 ± 0.3 | 18/G | 1.4 ± 0.4 | $5.57 \times 10^{-1}$ | 0.35 | $5.56 \times 10^{-1}$ | 0.04 | $8.47 \times 10^{-1}$ | | amino acid substitution (Lys1221Glu) | Moderate | | | |
| 3 | 17940574 | 10/A | 2 ± 0.7 | 70/G | 1.6 ± 0.3 | $6.29 \times 10^{-1}$ | 0.30 | $5.86 \times 10^{-1}$ | 0.82 | $3.68 \times 10^{-1}$ | | Promoter variant | Moderate | AT3G48440 | *ATC3H43* | Zinc finger C-x8-C-x5-C-x3-H type family protein |
| 3 | 17940713 | 62/T | 1.7 ± 0.3 | 18/G | 1.5 ± 0.4 | $6.42 \times 10^{-1}$ | 0.22 | $6.42 \times 10^{-1}$ | 0.01 | $9.04 \times 10^{-1}$ | | Promoter variant | Moderate | | | |
| 3 | 17940877 | 10/A | 2 ± 0.7 | 70/T | 1.6 ± 0.3 | $6.29 \times 10^{-1}$ | 0.30 | $5.86 \times 10^{-1}$ | 0.82 | $3.68 \times 10^{-1}$ | | Promoter variant | Moderate | | | |
| 3 | 17941022 | 13/T | 3.4 ± 1.1 | 67/C | 1.3 ± 0.2 | **$1.33 \times 10^{-3}$** | **10.28** | **$2.01 \times 10^{-3}$** | 1.93 | $1.69 \times 10^{-1}$ | | Promoter variant | Moderate | | | |
| 3 | 17941088 | 62/C | 1.8 ± 0.3 | 18/T | 1.1 ± 0.3 | $2.24 \times 10^{-1}$ | 1.48 | $2.27 \times 10^{-1}$ | 1.39 | $2.42 \times 10^{-1}$ | | Promoter variant | Moderate | | | |
| 3 | 17941143 | 61/C | 1.8 ± 0.3 | 19/T | 1.4 ± 0.4 | $5.21 \times 10^{-1}$ | 0.43 | $5.16 \times 10^{-1}$ | 0.01 | $9.25 \times 10^{-1}$ | | Promoter variant | Moderate | | | |
| 3 | 17941208 | 29/C | 2.4 ± 0.6 | 51/A | 1.3 ± 0.2 | **$2.51 \times 10^{-2}$** | **4.57** | **$3.59 \times 10^{-2}$** | 0.15 | $6.97 \times 10^{-1}$ | | Promoter variant | Moderate | | | |
| 3 | 17941684 | 10/A | 2 ± 0.7 | 70/G | 1.6 ± 0.3 | $6.29 \times 10^{-1}$ | 0.30 | $5.86 \times 10^{-1}$ | 0.82 | $3.68 \times 10^{-1}$ | | amino acid substitution (Gly95Ser) | Moderate | | | |
| 3 | 17943535 | 64/C | 1.9 ± 0.3 | 16/G | 0.6 ± 0.2 | **$2.50 \times 10^{-2}$** | **4.24** | **$4.32 \times 10^{-2}$** | 0.52 | $4.72 \times 10^{-1}$ | | amino acid substitution (Gln436Glu) | Moderate | | | |
| 3 | 17943693 | 11/A | 3.8 ± 1.2 | 69/T | 1.3 ± 0.2 | **$2.19 \times 10^{-4}$** | **14.45** | **$2.99 \times 10^{-4}$** | 2.43 | $1.23 \times 10^{-1}$ | ✓ | 3' - UTR variant | Moderate | | | |
| 3 | 22792092 | 67/A | 1.8 ± 0.3 | 13/C | 0.8 ± 0.2 | $9.30 \times 10^{-2}$ | 2.45 | $1.22 \times 10^{-1}$ | 0.12 | $7.29 \times 10^{-1}$ | | 5'-UTR variant | Moderate | AT3G61590 | *HWS* | Hawaiian skirt |

| | | | | | | | | | | | | | | | | |
|---|---|---|---|---|---|---|---|---|---|---|---|---|---|---|---|---|
| 3 | 22792297 | 75/C | 1.7 ± 0.3 | 5/G | 1.1 ± 0.5 | $5.39 \times 10^{-1}$ | 0.48 | $4.93 \times 10^{-1}$ | 0.00 | $9.61 \times 10^{-1}$ | | 5'-UTR variant | Moderate | | | |
| 3 | 22792412 | 11/T | 1.8 ± 0.6 | 69/G | 1.6 ± 0.3 | $8.18 \times 10^{-1}$ | 0.04 | $8.48 \times 10^{-1}$ | 0.02 | $8.85 \times 10^{-1}$ | | 5'-UTR variant | Moderate | | | |
| 3 | 22792573 | 54/TG | 2 ± 0.4 | 26/T | 1.1 ± 0.3 | **$7.62 \times 10^{-2}$** | 2.71 | $1.04 \times 10^{-1}$ | 0.02 | $8.78 \times 10^{-1}$ | | 5'-UTR intron variant | Moderate | | | |
| 3 | 22792583 | 44/A | 2 ± 0.4 | 36/C | 1.2 ± 0.3 | $1.12 \times 10^{-1}$ | 2.23 | $1.40 \times 10^{-1}$ | 0.64 | $4.28 \times 10^{-1}$ | | 5'-UTR intron variant | Moderate | | | |
| 3 | 22792586 | 42/G | 2.1 ± 0.5 | 38/C | 1.2 ± 0.3 | **$6.58 \times 10^{-2}$** | **3.09** | **$8.30 \times 10^{-2}$** | 0.26 | $6.10 \times 10^{-1}$ | | 5'-UTR intron variant | Moderate | | | |
| 3 | 22792816 | 66/T | 1.7 ± 0.3 | 14/C | 1.4 ± 0.7 | $5.51 \times 10^{-1}$ | 0.10 | $7.48 \times 10^{-1}$ | 0.01 | $9.07 \times 10^{-1}$ | | 5'-UTR intron variant | Moderate | | | |
| 3 | 22792904 | 3/A | 2.5 ± 1.9 | 77/G | 1.6 ± 0.3 | $4.85 \times 10^{-1}$ | 1.03 | $3.14 \times 10^{-1}$ | 2.27 | $1.36 \times 10^{-1}$ | | 5'-UTR variant | Moderate | | | |
| 3 | 22793392 | 68/C | 1.8 ± 0.3 | 12/G | 1.1 ± 0.3 | $3.44 \times 10^{-1}$ | 1.20 | $2.76 \times 10^{-1}$ | 0.03 | $8.73 \times 10^{-1}$ | | amino acid substitution (Thr160Arg) | Moderate | | | |
| 3 | 22795189 | 49/A | 1.9 ± 0.4 | 31/C | 1.3 ± 0.3 | $1.82 \times 10^{-1}$ | 1.57 | $2.14 \times 10^{-1}$ | 0.39 | $5.35 \times 10^{-1}$ | | Promoter variant | Moderate | AT3G61600 | *POB1* | Poz/btb containing-protein 1 |
| 3 | 22795571 | 62/C | 1.8 ± 0.3 | 18/CAAT | 1.1 ± 0.3 | $2.43 \times 10^{-1}$ | 0.90 | $3.47 \times 10^{-1}$ | 0.07 | $7.92 \times 10^{-1}$ | | Promoter variant | Moderate | | | |
| 3 | 22795605 | 38/T | 2.2 ± 0.5 | 42/C | 1.2 ± 0.2 | **$3.69 \times 10^{-2}$** | **3.85** | **$5.37 \times 10^{-2}$** | 1.52 | $2.22 \times 10^{-1}$ | | 5'-UTR variant | Moderate | | | |
| 3 | 22795608 | 46/C | 2.1 ± 0.4 | 34/A | 1.1 ± 0.2 | **$3.12 \times 10^{-2}$** | **4.61** | **$3.51 \times 10^{-2}$** | **3.05** | **$8.51 \times 10^{-2}$** | | 5'-UTR variant | Moderate | | | |
| 3 | 22795620 | 35/A | 1.9 ± 0.5 | 45/G | 1.5 ± 0.3 | $4.50 \times 10^{-1}$ | 0.39 | $5.35 \times 10^{-1}$ | 0.01 | $9.40 \times 10^{-1}$ | | 5'-UTR variant | Moderate | | | |
| 3 | 22795690 | 36/A | 2 ± 0.5 | 44/G | 1.4 ± 0.3 | $2.49 \times 10^{-1}$ | 1.09 | $3.00 \times 10^{-1}$ | 0.98 | $3.26 \times 10^{-1}$ | | 5'-UTR variant | Moderate | | | |
| 3 | 22795973 | 7/C | 5 ± 1.6 | 73/T | 1.4 ± 0.2 | **$6.38 \times 10^{-6}$** | **23.78** | **$6.26 \times 10^{-6}$** | **9.57** | **$2.82 \times 10^{-3}$** | ✓ | intron variant | Moderate | | | |
| 3 | 22798508 | 45/C | 2.2 ± 0.4 | 35/T | 1 ± 0.2 | **$1.62 \times 10^{-2}$** | **5.78** | **$1.88 \times 10^{-2}$** | **5.15** | **$2.62 \times 10^{-2}$** | | Promoter variant | Moderate | AT3G61610 | | Galactose mutarotase-like superfamily protein |
| 3 | 22798606 | 46/C | 1.7 ± 0.4 | 34/G | 1.6 ± 0.4 | $7.47 \times 10^{-1}$ | 0.11 | $7.39 \times 10^{-1}$ | 0.01 | $9.28 \times 10^{-1}$ | | Promoter variant | Moderate | | | |
| 3 | 22798735 | 70/T | 1.7 ± 0.3 | 10/TTA | 1.3 ± 0.5 | $5.68 \times 10^{-1}$ | 0.25 | $6.16 \times 10^{-1}$ | 0.65 | $4.22 \times 10^{-1}$ | | Promoter variant | Moderate | | | |
| 3 | 22798849 | 50/T | 1.9 ± 0.4 | 30/C | 1.2 ± 0.3 | $1.52 \times 10^{-1}$ | 2.09 | $1.53 \times 10^{-1}$ | 1.06 | $3.08 \times 10^{-1}$ | | Promoter variant | Moderate | | | |
| 3 | 22798851 | 52/C | 1.9 ± 0.4 | 28/A | 1.2 ± 0.3 | $1.98 \times 10^{-1}$ | 1.66 | $2.01 \times 10^{-1}$ | 1.09 | $3.00 \times 10^{-1}$ | | Promoter variant | Moderate | | | |
| 3 | 22799380 | 43/C | 1.8 ± 0.4 | 37/G | 1.5 ± 0.4 | $5.29 \times 10^{-1}$ | 0.24 | $6.23 \times 10^{-1}$ | 0.02 | $8.77 \times 10^{-1}$ | | 5'-UTR variant | Moderate | | | |
| 3 | 22800750 | 8/A | 4.6 ± 1.4 | 72/G | 1.3 ± 0.2 | **$1.36 \times 10^{-5}$** | **21.43** | **$1.59 \times 10^{-5}$** | **8.25** | **$5.34 \times 10^{-3}$** | ✓ | intron variant | Moderate | | | |
| 4 | 11286937 | 7/A | 1.9 ± 1.2 | 73/C | 1.6 ± 0.3 | $7.69 \times 10^{-1}$ | 0.04 | $8.37 \times 10^{-1}$ | 0.01 | $9.26 \times 10^{-1}$ | | amino acid substitution (Val492Gly) | Moderate | AT4G21170 | | Tetratricopeptide repeat (TPR)-like superfamily protein |
| 4 | 11287036 | 51/C | 2 ± 0.4 | 29/T | 1.1 ± 0.3 | **$5.93 \times 10^{-2}$** | **3.67** | **$5.94 \times 10^{-2}$** | 0.01 | $9.26 \times 10^{-1}$ | | amino acid substitution (Lys459Arg) | Moderate | | | |
| 4 | 11287349 | 7/A | 4.3 ± 1.6 | 73/G | 1.4 ± 0.2 | **$4.79 \times 10^{-4}$** | **13.45** | **$4.66 \times 10^{-4}$** | **5.14** | **$2.64 \times 10^{-2}$** | ✓ | synonymous variant | Moderate | | | |
| 4 | 11287503 | 67/A | 1.8 ± 0.3 | 13/C | 0.9 ± 0.2 | $1.72 \times 10^{-1}$ | 1.62 | $2.07 \times 10^{-1}$ | **4.12** | **$4.62 \times 10^{-2}$** | | amino acid substitution (Asp337Glu) | Moderate | | | |
| 4 | 11288356 | 66/G | 1.7 ± 0.3 | 14/A | 1.6 ± 0.3 | $9.11 \times 10^{-1}$ | 0.03 | $8.61 \times 10^{-1}$ | 0.15 | $7.00 \times 10^{-1}$ | | amino acid substitution (Thr53Ile) | Moderate | | | |
| 4 | 11288658 | 44/A | 2.2 ± 0.4 | 36/C | 1.1 ± 0.2 | **$1.95 \times 10^{-2}$** | **5.92** | **$1.75 \times 10^{-2}$** | 0.48 | $4.92 \times 10^{-1}$ | | Promoter variant | Moderate | | | |
| 4 | 11288745 | 68/C | 1.7 ± 0.3 | 12/G | 1.3 ± 0.5 | $5.54 \times 10^{-1}$ | 0.51 | $4.78 \times 10^{-1}$ | 0.03 | $8.61 \times 10^{-1}$ | | Promoter variant | Moderate | | | |
| 4 | 11289138 | 13/C | 1.7 ± 0.8 | 67/T | 1.7 ± 0.3 | $9.48 \times 10^{-1}$ | 0.00 | $9.79 \times 10^{-1}$ | 0.21 | $6.50 \times 10^{-1}$ | | Promoter variant | Moderate | | | |
| 5 | 6456413 | 32/A | 1.9 ± 0.4 | 48/C | 1.5 ± 0.4 | $4.54 \times 10^{-1}$ | 0.48 | $4.90 \times 10^{-1}$ | 0.22 | $6.38 \times 10^{-1}$ | | Promoter variant | Moderate | AT5G19190 | | Hypothetical protein |
| 5 | 6456455 | 29/A | 1.7 ± 0.3 | 51/G | 1.7 ± 0.4 | $9.09 \times 10^{-1}$ | 0.00 | $9.70 \times 10^{-1}$ | 0.07 | $7.97 \times 10^{-1}$ | | Promoter variant | Moderate | | | |
| 5 | 6456500 | 58/CA | 1.9 ± 0.3 | 22/C | 1.1 ± 0.4 | $1.69 \times 10^{-1}$ | 1.83 | $1.80 \times 10^{-1}$ | 0.85 | $3.59 \times 10^{-1}$ | | Promoter variant | Moderate | | | |
| 5 | 6456521 | 45/A | 1.7 ± 0.3 | 35/T | 1.7 ± 0.5 | $9.46 \times 10^{-1}$ | 0.00 | $9.45 \times 10^{-1}$ | 0.27 | $6.02 \times 10^{-1}$ | | Promoter variant | Moderate | | | |
| 5 | 6456527 | 57/T | 1.8 ± 0.4 | 23/A | 1.3 ± 0.4 | $3.00 \times 10^{-1}$ | 1.21 | $2.75 \times 10^{-1}$ | 0.06 | $8.12 \times 10^{-1}$ | | Promoter variant | Moderate | | | |

| | | | | | | | | | | | | | | | | |
|---|---|---|---|---|---|---|---|---|---|---|---|---|---|---|---|---|
| 5 | 6456628 | 41/A | 2 ± 0.4 | 39/AG | 1.4 ± 0.4 | $2.08 \times 10^{-1}$ | 1.34 | $2.51 \times 10^{-1}$ | 2.15 | $1.47 \times 10^{-1}$ | | Promoter variant | Moderate | | | |
| 5 | 6456708 | 59/C | 2 ± 0.4 | 21/A | 0.7 ± 0.1 | **$1.59 \times 10^{-2}$** | **6.53** | **$1.27 \times 10^{-2}$** | 0.46 | $5.00 \times 10^{-1}$ | | Promoter variant | Moderate | | | |
| 5 | 6456730 | 36/A | 2.1 ± 0.4 | 44/G | 1.3 ± 0.4 | $1.33 \times 10^{-1}$ | 2.46 | $1.21 \times 10^{-1}$ | 0.03 | $8.65 \times 10^{-1}$ | | Promoter variant | Moderate | | | |
| 5 | 6456731 | 38/A | 2 ± 0.4 | 42/T | 1.4 ± 0.4 | $2.11 \times 10^{-1}$ | 2.09 | $1.53 \times 10^{-1}$ | 0.11 | $7.43 \times 10^{-1}$ | | Promoter variant | Moderate | | | |
| 5 | 6456739 | 35/T | 2.3 ± 0.5 | 45/C | 1.2 ± 0.3 | **$1.55 \times 10^{-2}$** | **6.48** | **$1.31 \times 10^{-2}$** | 0.67 | $4.14 \times 10^{-1}$ | | Promoter variant | Moderate | | | |
| 5 | 6456821 | 39/T | 1.7 ± 0.5 | 41/C | 1.7 ± 0.3 | $9.56 \times 10^{-1}$ | 0.06 | $8.08 \times 10^{-1}$ | 0.32 | $5.75 \times 10^{-1}$ | | Promoter variant | Moderate | | | |
| 5 | 6456845 | 56/AT | 2 ± 0.4 | 24/A | 1 ± 0.4 | **$6.06 \times 10^{-2}$** | **3.61** | **$6.16 \times 10^{-2}$** | 0.43 | $5.15 \times 10^{-1}$ | | Promoter variant | Moderate | | | |
| 5 | 6456872 | 34/G | 1.9 ± 0.4 | 46/C | 1.5 ± 0.4 | $3.57 \times 10^{-1}$ | 0.75 | $3.90 \times 10^{-1}$ | 0.14 | $7.10 \times 10^{-1}$ | | Promoter variant | Moderate | | | |
| 5 | 6456877 | 32/T | 2 ± 0.4 | 48/C | 1.4 ± 0.4 | $2.51 \times 10^{-1}$ | 1.34 | $2.50 \times 10^{-1}$ | 0.08 | $7.76 \times 10^{-1}$ | | Promoter variant | Moderate | | | |
| 5 | 6456897 | 30/G | 1.9 ± 0.4 | 50/A | 1.5 ± 0.4 | $3.78 \times 10^{-1}$ | 0.74 | $3.92 \times 10^{-1}$ | 0.01 | $9.25 \times 10^{-1}$ | | Promoter variant | Moderate | | | |
| 5 | 6456901 | 13/C | 2 ± 0.4 | 67/T | 1.6 ± 0.3 | $5.46 \times 10^{-1}$ | 0.29 | $5.93 \times 10^{-1}$ | 0.67 | $4.15 \times 10^{-1}$ | | Promoter variant | Moderate | | | |
| 5 | 6456995 | 50/G | 1.9 ± 0.4 | 30/GT | 1.3 ± 0.4 | $1.84 \times 10^{-1}$ | 1.37 | $2.46 \times 10^{-1}$ | 0.27 | $6.06 \times 10^{-1}$ | | Promoter variant | Moderate | | | |
| 5 | 6457210 | 46/TG | 1.9 ± 0.4 | 34/T | 1.4 ± 0.3 | $3.56 \times 10^{-1}$ | 0.97 | $3.28 \times 10^{-1}$ | 0.14 | $7.13 \times 10^{-1}$ | | Promoter variant | Moderate | | | |
| 5 | 6457212 | 34/C | 1.9 ± 0.4 | 46/A | 1.5 ± 0.4 | $4.69 \times 10^{-1}$ | 0.38 | $5.40 \times 10^{-1}$ | 0.09 | $7.62 \times 10^{-1}$ | | Promoter variant | Moderate | | | |
| 5 | 6458384 | 13/G | 3.2 ± 1 | 67/A | 1.4 ± 0.3 | **$4.19 \times 10^{-3}$** | **9.10** | **$3.53 \times 10^{-3}$** | **7.14** | **$9.33 \times 10^{-3}$** | | Promoter variant | Moderate | AT5G19200 | *TSC10B* | NAD(P)-binding Rossmann-fold superfamily protein |
| 5 | 6458684 | 70/A | 1.8 ± 0.3 | 10/AT | 0.9 ± 0.3 | $2.13 \times 10^{-1}$ | 1.73 | $1.92 \times 10^{-1}$ | 1.63 | $2.05 \times 10^{-1}$ | | Promoter variant | Moderate | | | |
| 5 | 6458759 | 36/A | 2.2 ± 0.5 | 44/T | 1.2 ± 0.3 | **$3.51 \times 10^{-2}$** | **6.06** | **$1.62 \times 10^{-2}$** | 1.09 | $3.00 \times 10^{-1}$ | | Promoter variant | Moderate | | | |
| 5 | 6458762 | 36/T | 2.1 ± 0.5 | 44/A | 1.3 ± 0.3 | **$8.05 \times 10^{-2}$** | **4.24** | **$4.31 \times 10^{-2}$** | 1.84 | $1.79 \times 10^{-1}$ | | Promoter variant | Moderate | | | |
| 5 | 6458889 | 56/A | 2.1 ± 0.4 | 24/AT | 0.7 ± 0.2 | **$9.20 \times 10^{-3}$** | **7.96** | **$6.17 \times 10^{-3}$** | 2.49 | $1.19 \times 10^{-1}$ | | Promoter variant | Moderate | | | |
| 5 | 6458972 | 76/G | 1.7 ± 0.3 | 4/A | 0.7 ± 0.2 | $3.66 \times 10^{-1}$ | 0.97 | $3.27 \times 10^{-1}$ | 0.38 | $5.42 \times 10^{-1}$ | | 5'-UTR premature start codon gain variant | Moderate | | | |
| 5 | 6459296 | 57/C | 1.9 ± 0.4 | 23/G | 1 ± 0.3 | **$9.90 \times 10^{-2}$** | **4.44** | **$3.85 \times 10^{-2}$** | 1.17 | $2.83 \times 10^{-1}$ | | amino acid substitution (Ala101Gly) | Moderate | | | |
| 5 | 6459439 | 7/T | 3.7 ± 1.7 | 73/G | 1.5 ± 0.3 | **$7.18 \times 10^{-3}$** | **16.63** | **$1.16 \times 10^{-4}$** | **13.83** | **$3.93 \times 10^{-4}$** | ✓ | synonymous variant | Moderate | | | |
| 5 | 6459783 | 10/G | 1.8 ± 0.5 | 70/A | 1.6 ± 0.3 | $8.05 \times 10^{-1}$ | 0.04 | $8.41 \times 10^{-1}$ | 0.56 | $4.58 \times 10^{-1}$ | | amino acid substitution (Gly208Asp) | Moderate | | | |
| 5 | 6460438 | 33/G | 1.7 ± 0.3 | 47/T | 1.7 ± 0.4 | $9.86 \times 10^{-1}$ | 0.00 | $9.99 \times 10^{-1}$ | 0.02 | $9.02 \times 10^{-1}$ | | amino acid substitution (Asn248Lys) | Moderate | | | |
| 5 | 6460598 | 71/C | 1.7 ± 0.3 | 9/T | 1.5 ± 0.7 | $8.02 \times 10^{-1}$ | 0.09 | $7.59 \times 10^{-1}$ | 0.14 | $7.11 \times 10^{-1}$ | | amino acid substitution (Leu302Phe) | Moderate | | | |
| 5 | 6460602 | 70/C | 1.7 ± 0.3 | 10/T | 1.4 ± 0.6 | $6.58 \times 10^{-1}$ | 0.25 | $6.15 \times 10^{-1}$ | 0.04 | $8.39 \times 10^{-1}$ | | amino acid substitution (Ala303Val) | Moderate | | | |
| 5 | 6460444 | 37/G | 1.8 ± 0.4 | 43/A | 1.6 ± 0.4 | $7.58 \times 10^{-1}$ | 0.07 | $7.97 \times 10^{-1}$ | 0.01 | $9.37 \times 10^{-1}$ | | Promoter variant | Moderate | AT5G19210 | | P-loop containing nucleoside triphosphate hydrolases superfamily protein |
| 5 | 6460450 | 36/A | 1.8 ± 0.5 | 44/G | 1.6 ± 0.3 | $6.82 \times 10^{-1}$ | 0.08 | $7.74 \times 10^{-1}$ | 0.01 | $9.19 \times 10^{-1}$ | | Promoter variant | Moderate | | | |
| 5 | 6460483 | 23/C | 1.9 ± 0.5 | 57/T | 1.6 ± 0.3 | $6.01 \times 10^{-1}$ | 0.19 | $6.60 \times 10^{-1}$ | 0.32 | $5.76 \times 10^{-1}$ | | Promoter variant | Moderate | | | |
| 5 | 6460519 | 38/T | 1.7 ± 0.4 | 42/A | 1.6 ± 0.4 | $9.07 \times 10^{-1}$ | 0.02 | $8.81 \times 10^{-1}$ | 0.03 | $8.53 \times 10^{-1}$ | | Promoter variant | Moderate | | | |
| 5 | 6460552 | 13/G | 2.6 ± 1 | 67/T | 1.5 ± 0.3 | $1.06 \times 10^{-1}$ | 2.63 | $1.09 \times 10^{-1}$ | 0.19 | $6.61 \times 10^{-1}$ | | Promoter variant | Moderate | | | |
| 5 | 6463258 | 9/A | 1.7 ± 0.5 | 71/G | 1.7 ± 0.3 | $9.64 \times 10^{-1}$ | 0.00 | $9.65 \times 10^{-1}$ | 0.03 | $8.59 \times 10^{-1}$ | | amino acid substitution (Glu199Lys) | Moderate | | | |
| 5 | 6463601 | 9/A | 1.7 ± 0.5 | 71/G | 1.7 ± 0.3 | $9.64 \times 10^{-1}$ | 0.00 | $9.65 \times 10^{-1}$ | 0.03 | $8.59 \times 10^{-1}$ | | amino acid substitution (Ala288Thr) | Moderate | | | |
| 5 | 6464350 | 10/A | 1.8 ± 0.5 | 70/T | 1.6 ± 0.3 | $7.94 \times 10^{-1}$ | 0.04 | $8.33 \times 10^{-1}$ | 1.02 | $3.16 \times 10^{-1}$ | | amino acid substitution (Leu452Gln) | Moderate | AT5G19220 | *ADG2* | ADP glucose pyrophosphorylase 2 |
| 5 | 6466552 | 27/C | 1.8 ± 0.3 | 53/T | 1.6 ± 0.4 | $6.86 \times 10^{-1}$ | 0.10 | $7.52 \times 10^{-1}$ | 0.01 | $9.12 \times 10^{-1}$ | | amino acid substitution (Ser75Asn) | Moderate | | | |

| | | | | | | | | | | | | | | | |
|---|---|---|---|---|---|---|---|---|---|---|---|---|---|---|---|
| 5 | 6466909 | 68/C | 1.8 ± 0.3 | 12/T | 0.7 ± 0.2 | **$7.71 \times 10^{-2}$** | **3.79** | **$5.56 \times 10^{-2}$** | 1.09 | $3.01 \times 10^{-1}$ | | Promoter variant | Moderate | | |
| 5 | 6467109 | 17/T | 2.1 ± 0.6 | 63/A | 1.6 ± 0.3 | $3.69 \times 10^{-1}$ | 0.73 | $3.97 \times 10^{-1}$ | 0.05 | $8.23 \times 10^{-1}$ | | Promoter variant | Moderate | | |
| 5 | 6467203 | 17/G | 2.1 ± 0.5 | 63/A | 1.6 ± 0.3 | $4.06 \times 10^{-1}$ | 0.68 | $4.12 \times 10^{-1}$ | 0.04 | $8.51 \times 10^{-1}$ | | Promoter variant | Moderate | | |
| 5 | 6467254 | 66/A | 1.9 ± 0.3 | 14/C | 0.7 ± 0.2 | **$6.17 \times 10^{-2}$** | **4.62** | **$3.50 \times 10^{-2}$** | 0.25 | $6.22 \times 10^{-1}$ | | Promoter variant | Moderate | | |
| 5 | 6467325 | 16/C | 2.1 ± 0.6 | 64/T | 1.6 ± 0.3 | $3.74 \times 10^{-1}$ | 0.71 | $4.03 \times 10^{-1}$ | 0.22 | $6.38 \times 10^{-1}$ | | Promoter variant | Moderate | | |
| 5 | 6467371 | 16/A | 2.3 ± 0.6 | 64/C | 1.5 ± 0.3 | $2.24 \times 10^{-1}$ | 1.34 | $2.51 \times 10^{-1}$ | 1.58 | $2.13 \times 10^{-1}$ | | Promoter variant | Moderate | | |
| 5 | 6467374 | 48/T | 1.7 ± 0.4 | 32/A | 1.7 ± 0.4 | $9.95 \times 10^{-1}$ | 0.00 | $9.49 \times 10^{-1}$ | 0.01 | $9.43 \times 10^{-1}$ | | Promoter variant | Moderate | | |
| 5 | 6467380 | 30/A | 2.1 ± 0.5 | 50/T | 1.4 ± 0.3 | $1.49 \times 10^{-1}$ | 1.90 | $1.73 \times 10^{-1}$ | 0.33 | $5.68 \times 10^{-1}$ | | Promoter variant | Moderate | | |
| 5 | 6467558 | 43/C | 2.2 ± 0.4 | 37/T | 1.1 ± 0.4 | **$2.29 \times 10^{-2}$** | **5.36** | **$2.34 \times 10^{-2}$** | 2.61 | $1.10 \times 10^{-1}$ | | 5'-UTR variant | Moderate | | |
| 5 | 6467562 | 54/C | 2.2 ± 0.4 | 26/CT | 0.7 ± 0.1 | **$3.09 \times 10^{-3}$** | **8.78** | **$4.13 \times 10^{-3}$** | 2.11 | $1.51 \times 10^{-1}$ | | 5'-UTR variant | Moderate | | |
| 5 | 7953822 | 32/C | 2.6 ± 0.6 | 48/T | 1.1 ± 0.2 | **$1.28 \times 10^{-3}$** | **13.10** | **$5.47 \times 10^{-4}$** | **3.06** | **$8.47 \times 10^{-2}$** | ✓ | | Moderate | AT5G23590 | DNAJ heat shock N-terminal domain-containing protein |
| 5 | 7954097 | 60/G | 1.7 ± 0.3 | 20/A | 1.6 ± 0.6 | $7.91 \times 10^{-1}$ | 0.06 | $8.09 \times 10^{-1}$ | 0.14 | $7.14 \times 10^{-1}$ | | Promoter variant | Moderate | | |
| 5 | 7954208 | 54/C | 2 ± 0.4 | 26/G | 1 ± 0.2 | **$4.82 \times 10^{-2}$** | **5.04** | **$2.78 \times 10^{-2}$** | 1.66 | $2.02 \times 10^{-1}$ | | Promoter variant | Moderate | | |
| 5 | 7954224 | 22/T | 2.4 ± 0.7 | 58/G | 1.4 ± 0.3 | **$7.17 \times 10^{-2}$** | **4.18** | **$4.45 \times 10^{-2}$** | 0.32 | $5.74 \times 10^{-1}$ | | Promoter variant | Moderate | | |
| 5 | 7954727 | 63/T | 1.8 ± 0.3 | 17/C | 1.1 ± 0.4 | $2.28 \times 10^{-1}$ | 1.30 | $2.58 \times 10^{-1}$ | 0.13 | $7.22 \times 10^{-1}$ | | Promoter variant | Moderate | | |
| 5 | 7946585 | 63/C | 1.7 ± 0.3 | 17/G | 1.6 ± 0.4 | $8.64 \times 10^{-1}$ | 0.00 | $9.75 \times 10^{-1}$ | 0.04 | $8.34 \times 10^{-1}$ | | amino acid substitution (Thr8Arg) | Moderate | AT5G23575 | Transmembrane CLPTM1 family protein |
| 5 | 7946143 | 32/C | 1.8 ± 0.4 | 48/T | 1.6 ± 0.4 | $5.59 \times 10^{-1}$ | 0.47 | $4.95 \times 10^{-1}$ | 0.03 | $8.65 \times 10^{-1}$ | | Promoter variant | Moderate | | |
| 5 | 7946222 | 58/C | 1.8 ± 0.4 | 22/A | 1.4 ± 0.4 | $5.32 \times 10^{-1}$ | 0.47 | $4.95 \times 10^{-1}$ | 0.58 | $4.50 \times 10^{-1}$ | | Promoter variant | Moderate | | |
| 5 | 7946251 | 26/T | 2.2 ± 0.6 | 54/G | 1.4 ± 0.3 | $1.37 \times 10^{-1}$ | 2.23 | $1.40 \times 10^{-1}$ | 1.13 | $2.90 \times 10^{-1}$ | | Promoter variant | Moderate | | |
| 5 | 7946361 | 52/G | 1.9 ± 0.4 | 28/C | 1.3 ± 0.3 | $2.01 \times 10^{-1}$ | 1.98 | $1.64 \times 10^{-1}$ | 2.33 | $1.31 \times 10^{-1}$ | | 5'-- UTR_variant | Moderate | | |
| 5 | 9207766 | 69/A | 1.8 ± 0.3 | 1/T | 1.2 ± 0.5 | $4.43 \times 10^{-1}$ | 0.82 | $4.45 \times 10^{-1}$ | 0.61 | $5.44 \times 10^{-1}$ | | Promoter variant | Moderate | AT5G26280 | TRAF-like family protein |
| 5 | 9208140 | 28/G | 2 ± 0.5 | 52/A | 1.5 ± 0.3 | $2.56 \times 10^{-1}$ | 1.09 | $3.00 \times 10^{-1}$ | 0.57 | $4.52 \times 10^{-1}$ | | Promoter variant | Moderate | | |
| 5 | 9208174 | 69/A | 1.8 ± 0.3 | 11/T | 1 ± 0.3 | $2.70 \times 10^{-1}$ | 1.49 | $2.26 \times 10^{-1}$ | 0.20 | $6.58 \times 10^{-1}$ | | Promoter variant | Moderate | | |
| 5 | 9208198 | 15/G | 1.9 ± 0.4 | 65/C | 1.6 ± 0.3 | $6.40 \times 10^{-1}$ | 0.17 | $6.79 \times 10^{-1}$ | 0.17 | $6.84 \times 10^{-1}$ | | Promoter variant | Moderate | | |
| 5 | 9208206 | 66/T | 1.7 ± 0.3 | 14/A | 1.5 ± 0.6 | $7.20 \times 10^{-1}$ | 0.18 | $6.73 \times 10^{-1}$ | 0.05 | $8.23 \times 10^{-1}$ | | Promoter variant | Moderate | | |
| 5 | 9208291 | 35/C | 1.9 ± 0.4 | 45/T | 1.5 ± 0.4 | $4.91 \times 10^{-1}$ | 0.35 | $5.55 \times 10^{-1}$ | 0.13 | $7.15 \times 10^{-1}$ | | Promoter variant | Moderate | | |
| 5 | 9208335 | 44/C | 1.8 ± 0.4 | 36/A | 1.5 ± 0.3 | $6.47 \times 10^{-1}$ | 0.38 | $5.42 \times 10^{-1}$ | 0.14 | $7.05 \times 10^{-1}$ | | Promoter variant | Moderate | | |
| 5 | 9208400 | 45/A | 1.8 ± 0.4 | 35/G | 1.6 ± 0.3 | $7.11 \times 10^{-1}$ | 0.27 | $6.08 \times 10^{-1}$ | 0.47 | $4.96 \times 10^{-1}$ | | Promoter variant | Moderate | | |
| 5 | 9208426 | 36/C | 1.8 ± 0.4 | 44/T | 1.6 ± 0.4 | $7.32 \times 10^{-1}$ | 0.05 | $8.21 \times 10^{-1}$ | 0.02 | $8.87 \times 10^{-1}$ | | Promoter variant | Moderate | | |
| 5 | 9208452 | 47/C | 1.8 ± 0.4 | 33/G | 1.5 ± 0.3 | $6.28 \times 10^{-1}$ | 0.37 | $5.46 \times 10^{-1}$ | 0.10 | $7.58 \times 10^{-1}$ | | Promoter variant | Moderate | | |
| 5 | 9208521 | 12/A | 2.1 ± 0.5 | 68/G | 1.6 ± 0.3 | $4.56 \times 10^{-1}$ | 0.48 | $4.91 \times 10^{-1}$ | 0.58 | $4.48 \times 10^{-1}$ | | Promoter variant | Moderate | | |
| 5 | 9208553 | 69/A | 1.7 ± 0.3 | 11/C | 1.5 ± 0.8 | $7.83 \times 10^{-1}$ | 0.11 | $7.41 \times 10^{-1}$ | 0.05 | $8.23 \times 10^{-1}$ | | Promoter variant | Moderate | | |
| 5 | 9208664 | 65/T | 1.7 ± 0.3 | 15/TA | 1.5 ± 0.4 | $7.82 \times 10^{-1}$ | 0.10 | $7.53 \times 10^{-1}$ | 0.01 | $9.43 \times 10^{-1}$ | | Promoter variant | Moderate | | |
| 5 | 9209364 | 12/G | 2.1 ± 0.8 | 68/T | 1.6 ± 0.3 | $4.52 \times 10^{-1}$ | 0.51 | $4.76 \times 10^{-1}$ | 0.47 | $4.97 \times 10^{-1}$ | | amino acid substitution (Leu123Val) | Moderate | | |
| 5 | 9209620 | 3/A | 6.2 ± 1.4 | 77/G | 1.5 ± 0.3 | **$1.20 \times 10^{-4}$** | **16.85** | **$1.05 \times 10^{-4}$** | **7.46** | **$7.94 \times 10^{-3}$** | ✓ | synonymous variant | Moderate | | |

| | | | | | | | | | | | | | | | | |
|---|---|---|---|---|---|---|---|---|---|---|---|---|---|---|---|---|
| 5 | 9209651 | 16/T | 1.8 ± 0.6 | 64/C | 1.6 ± 0.3 | $8.03 \times 10^{-1}$ | 0.03 | $8.67 \times 10^{-1}$ | 0.12 | $7.33 \times 10^{-1}$ | | amino acid substitution (Pro168Ser) | Moderate | | | |
| 5 | 9209666 | 14/G | 2 ± 0.4 | 66/T | 1.6 ± 0.3 | $5.76 \times 10^{-1}$ | 0.25 | $6.20 \times 10^{-1}$ | 0.52 | $4.72 \times 10^{-1}$ | | amino acid substitution (Tyr173Asp) | Moderate | | | |
| 5 | 9210060 | 68/C | 1.7 ± 0.3 | 12/T | 1.6 ± 0.5 | $8.98 \times 10^{-1}$ | 0.05 | $8.25 \times 10^{-1}$ | 0.02 | $8.97 \times 10^{-1}$ | | amino acid substitution (Ala273Val) | Moderate | | | |
| 5 | 13913080 | 39/G | 1.7 ± 0.4 | 41/T | 1.7 ± 0.4 | $9.46 \times 10^{-1}$ | 0.01 | $9.11 \times 10^{-1}$ | 0.99 | $3.23 \times 10^{-1}$ | | amino acid substitution (Gln789Lys) | Moderate | AT5G35750 | *HK2* | Histidine kinase 2 |
| 5 | 13913597 | 14/T | 2.9 ± 0.8 | 66/G | 1.4 ± 0.3 | **$2.09 \times 10^{-2}$** | **5.98** | **$1.70 \times 10^{-2}$** | 2.42 | $1.24 \times 10^{-1}$ | ✓ | synonymous variant | Moderate | | | |
| 5 | 13915231 | 43/G | 1.8 ± 0.4 | 37/A | 1.5 ± 0.4 | $5.72 \times 10^{-1}$ | 0.63 | $4.29 \times 10^{-1}$ | 0.41 | $5.26 \times 10^{-1}$ | | amino acid substitution (Ile302Thr) | Moderate | | | |
| 5 | 13916422 | 10/T | 2.8 ± 0.9 | 70/G | 1.5 ± 0.3 | **$7.26 \times 10^{-2}$** | **3.52** | **$6.46 \times 10^{-2}$** | 1.96 | $1.65 \times 10^{-1}$ | | Promoter variant | Moderate | | | |
| 5 | 13916440 | 32/C | 1.7 ± 0.4 | 48/T | 1.7 ± 0.4 | $9.24 \times 10^{-1}$ | 0.11 | $7.46 \times 10^{-1}$ | 0.09 | $7.69 \times 10^{-1}$ | | Promoter variant | Moderate | | | |
| 5 | 13916644 | 15/G | 2.7 ± 0.8 | 65/T | 1.4 ± 0.3 | **$4.15 \times 10^{-2}$** | **4.48** | **$3.77 \times 10^{-2}$** | 1.15 | $2.87 \times 10^{-1}$ | | Promoter variant | Moderate | | | |
| 5 | 13916664 | 68/T | 1.8 ± 0.3 | 12/TC | 0.9 ± 0.2 | $1.51 \times 10^{-1}$ | 1.78 | $1.86 \times 10^{-1}$ | 2.03 | $1.59 \times 10^{-1}$ | | Promoter variant | Moderate | | | |
| 5 | 13916737 | 11/C | 2.4 ± 0.8 | 69/G | 1.6 ± 0.3 | $2.06 \times 10^{-1}$ | 1.74 | $1.91 \times 10^{-1}$ | 0.52 | $4.75 \times 10^{-1}$ | | Promoter variant | Moderate | | | |
| 5 | 13916811 | 55/T | 1.7 ± 0.3 | 25/A | 1.5 ± 0.4 | $6.42 \times 10^{-1}$ | 0.06 | $8.06 \times 10^{-1}$ | 0.02 | $8.92 \times 10^{-1}$ | | Promoter variant | Moderate | | | |
| 5 | 13916833 | 55/G | 1.8 ± 0.4 | 25/T | 1.5 ± 0.4 | $6.05 \times 10^{-1}$ | 0.10 | $7.49 \times 10^{-1}$ | 1.50 | $2.24 \times 10^{-1}$ | | Promoter variant | Moderate | | | |
| 5 | 13916835 | 56/T | 1.7 ± 0.3 | 24/C | 1.5 ± 0.5 | $6.63 \times 10^{-1}$ | 0.05 | $8.22 \times 10^{-1}$ | 2.06 | $1.56 \times 10^{-1}$ | | Promoter variant | Moderate | | | |
| 5 | 13916868 | 58/C | 1.7 ± 0.3 | 22/T | 1.5 ± 0.5 | $6.56 \times 10^{-1}$ | 0.04 | $8.38 \times 10^{-1}$ | 0.51 | $4.78 \times 10^{-1}$ | | Promoter variant | Moderate | | | |
| 5 | 13916940 | 60/G | 1.8 ± 0.4 | 20/T | 1.2 ± 0.3 | $2.59 \times 10^{-1}$ | 0.84 | $3.61 \times 10^{-1}$ | 0.03 | $8.70 \times 10^{-1}$ | | Promoter variant | Moderate | | | |
| 5 | 13916943 | 66/A | 1.9 ± 0.3 | 14/G | 0.8 ± 0.2 | **$9.80 \times 10^{-2}$** | 2.05 | $1.57 \times 10^{-1}$ | 0.00 | $9.80 \times 10^{-1}$ | | Promoter variant | Moderate | | | |
| 5 | 13916964 | 68/T | 1.8 ± 0.3 | 12/C | 0.8 ± 0.2 | $1.16 \times 10^{-1}$ | 2.20 | $1.43 \times 10^{-1}$ | 1.20 | $2.77 \times 10^{-1}$ | | Promoter variant | Moderate | | | |
| 5 | 13922702 | 79/ATC | 1.7 ± 0.3 | 1/A | 0.9 | | 0.17 | $6.80 \times 10^{-1}$ | 0.00 | $9.63 \times 10^{-1}$ | | frameshift variant (Glu546) | **High** | AT5G35753 | | DnaJ heat shock amino-terminal domain protein (DUF3444) |
| 5 | 13923351 | 79/T | 1.7 ± 0.3 | 1/A | 0.7 | | 0.23 | $6.34 \times 10^{-1}$ | 0.00 | $9.70 \times 10^{-1}$ | | stop codon gain (Lys331*) | **High** | | | |
| 5 | 13923474 | 39/T | 1.7 ± 0.4 | 41/G | 1.7 ± 0.4 | $9.41 \times 10^{-1}$ | 0.01 | $9.30 \times 10^{-1}$ | 1.38 | $2.43 \times 10^{-1}$ | | amino acid substitution (Ile290Leu) | Moderate | | | |
| 5 | 13923564 | 66/C | 1.8 ± 0.3 | 14/G | 0.9 ± 0.2 | $1.23 \times 10^{-1}$ | 1.70 | $1.97 \times 10^{-1}$ | 0.07 | $7.87 \times 10^{-1}$ | | amino acid substitution (Ala260Pro) | Moderate | | | |
| 5 | 13924040 | 15/C | 2.7 ± 0.8 | 65/A | 1.4 ± 0.3 | **$4.01 \times 10^{-2}$** | **4.60** | **$3.54 \times 10^{-2}$** | 1.91 | $1.71 \times 10^{-1}$ | | amino acid substitution (Leu138Arg) | Moderate | | | |
| 5 | 13924157 | 40/G | 1.7 ± 0.4 | 40/T | 1.6 ± 0.4 | $9.28 \times 10^{-1}$ | 0.01 | $9.20 \times 10^{-1}$ | 1.43 | $2.36 \times 10^{-1}$ | | amino acid substitution (Tyr99Ser) | Moderate | | | |
| 5 | 13924265 | 50/A | 1.8 ± 0.4 | 30/G | 1.5 ± 0.4 | $5.48 \times 10^{-1}$ | 0.18 | $6.72 \times 10^{-1}$ | 0.18 | $6.69 \times 10^{-1}$ | | 5'-UTR intron variant | Moderate | | | |
| 5 | 13924444 | 16/T | 2.6 ± 0.8 | 64/C | 1.4 ± 0.3 | **$5.54 \times 10^{-2}$** | **4.02** | **$4.88 \times 10^{-2}$** | 0.26 | $6.15 \times 10^{-1}$ | | 5'-UTR intron variant | Moderate | | | |
| 5 | 13924575 | 70/G | 1.8 ± 0.3 | 10/A | 0.9 ± 0.3 | $2.03 \times 10^{-1}$ | 1.00 | $3.21 \times 10^{-1}$ | 0.05 | $8.31 \times 10^{-1}$ | | amino acid substitution (Ala25Val) | Moderate | | | |
| 5 | 13924658 | 12/A | 2.3 ± 0.8 | 68/G | 1.6 ± 0.3 | $3.10 \times 10^{-1}$ | 1.09 | $3.01 \times 10^{-1}$ | 0.33 | $5.66 \times 10^{-1}$ | | 5'-UTR premature start codon gain variant | Moderate | | | |
| 5 | 13924698 | 13/T | 2.4 ± 0.7 | 67/C | 1.5 ± 0.3 | $1.96 \times 10^{-1}$ | 1.71 | $1.95 \times 10^{-1}$ | 0.56 | $4.56 \times 10^{-1}$ | | 5'-UTR variant | Moderate | | | |
| 5 | 13924704 | 42/G | 1.8 ± 0.4 | 38/C | 1.5 ± 0.4 | $4.69 \times 10^{-1}$ | 0.90 | $3.45 \times 10^{-1}$ | 0.20 | $6.60 \times 10^{-1}$ | | 5'-UTR variant | Moderate | | | |
| 5 | 13924822 | 70/T | 1.8 ± 0.3 | 10/C | 1.1 ± 0.3 | $3.49 \times 10^{-1}$ | 0.37 | $5.47 \times 10^{-1}$ | 0.06 | $8.08 \times 10^{-1}$ | | 5'-UTR intron variant | Moderate | | | |
| 5 | 13924872 | 32/C | 1.8 ± 0.4 | 48/T | 1.6 ± 0.4 | $7.76 \times 10^{-1}$ | 0.28 | $6.00 \times 10^{-1}$ | 0.21 | $6.50 \times 10^{-1}$ | | 5'-UTR intron variant | Moderate | | | |
| 5 | 13926798 | 14/A | 2.9 ± 0.8 | 66/G | 1.4 ± 0.3 | **$2.09 \times 10^{-2}$** | **5.98** | **$1.70 \times 10^{-2}$** | 2.42 | $1.24 \times 10^{-1}$ | ✓ | synonymous variant | Moderate | AT5G35756 | | Copia-like retrotransposon family |
| 5 | 13930156 | 14/G | 2.9 ± 0.8 | 66/T | 1.4 ± 0.3 | **$2.09 \times 10^{-2}$** | **5.98** | **$1.70 \times 10^{-2}$** | 2.42 | $1.24 \times 10^{-1}$ | ✓ | synonymous variant | Moderate | | | |
| 5 | 13930909 | 15/T | 2.8 ± 0.8 | 65/G | 1.4 ± 0.3 | **$1.82 \times 10^{-2}$** | **6.26** | **$1.46 \times 10^{-2}$** | 2.85 | **$9.55 \times 10^{-2}$** | ✓ | synonymous variant | Moderate | | | |

| | | | | | | | | | | | | | | | | |
|---|---|---|---|---|---|---|---|---|---|---|---|---|---|---|---|---|
| 5 | 20039699 | 10/C | 1.7 ± 0.5 | 70/T | 1.7 ± 0.3 | $9.86 \times 10^{-1}$ | 0.00 | $9.93 \times 10^{-1}$ | 0.00 | $9.53 \times 10^{-1}$ | | amino acid substitution (Asn1017Asp) | Moderate | AT5G49430 | | WD40/YVTN repeat and Bromo-WDR9-I-like domain-containing protein |
| 5 | 20039717 | 16/A | 2.4 ± 0.7 | 64/G | 1.5 ± 0.3 | $1.43 \times 10^{-1}$ | **3.54** | **$6.41 \times 10^{-2}$** | 0.28 | $5.98 \times 10^{-1}$ | | amino acid substitution (Pro1011Ser) | Moderate | | | |
| 5 | 20040327 | 6/G | 4.5 ± 1.5 | 74/A | 1.4 ± 0.3 | **$5.60 \times 10^{-4}$** | **14.28** | **$3.22 \times 10^{-4}$** | **4.70** | **$3.34 \times 10^{-2}$** | ✓ | synonymous variant | Moderate | | | |
| 5 | 20045743 | 13/C | 2.3 ± 0.9 | 67/A | 1.6 ± 0.3 | $2.65 \times 10^{-1}$ | 1.26 | $2.66 \times 10^{-1}$ | 1.83 | $1.80 \times 10^{-1}$ | | Promoter variant | Moderate | | | |
| 5 | 20045797 | 69/A | 1.8 ± 0.3 | 11/G | 0.7 ± 0.2 | **$8.90 \times 10^{-2}$** | **4.19** | **$4.44 \times 10^{-2}$** | 0.08 | $7.80 \times 10^{-1}$ | | Promoter variant | Moderate | | | |
| 5 | 20045854 | 16/A | 1.9 ± 0.7 | 64/G | 1.6 ± 0.3 | $6.14 \times 10^{-1}$ | 0.23 | $6.32 \times 10^{-1}$ | 0.10 | $7.49 \times 10^{-1}$ | | Promoter variant | Moderate | | | |
| 5 | 22203250 | 57/CT | 1.8 ± 0.4 | 23/C | 1.4 ± 0.3 | $4.93 \times 10^{-1}$ | 0.62 | $4.34 \times 10^{-1}$ | 0.03 | $8.71 \times 10^{-1}$ | | Promoter variant | Moderate | AT5G54660 | | HSP20-like chaperones superfamily protein |
| 5 | 22203435 | 61/T | 1.8 ± 0.3 | 19/G | 1.4 ± 0.4 | $4.76 \times 10^{-1}$ | 0.59 | $4.44 \times 10^{-1}$ | 0.78 | $3.81 \times 10^{-1}$ | | Promoter variant | Moderate | | | |
| 5 | 22203439 | 70/A | 1.7 ± 0.3 | 10/C | 1.3 ± 0.5 | $5.69 \times 10^{-1}$ | 0.45 | $5.05 \times 10^{-1}$ | 0.15 | $7.00 \times 10^{-1}$ | | Promoter variant | Moderate | | | |
| 5 | 22203567 | 62/G | 1.8 ± 0.4 | 18/A | 1.1 ± 0.2 | $2.08 \times 10^{-1}$ | 1.65 | $2.03 \times 10^{-1}$ | 2.65 | $1.08 \times 10^{-1}$ | | Promoter variant | Moderate | | | |
| 5 | 22204077 | 35/T | 1.9 ± 0.4 | 45/A | 1.5 ± 0.4 | $4.50 \times 10^{-1}$ | 0.83 | $3.65 \times 10^{-1}$ | 1.17 | $2.83 \times 10^{-1}$ | | amino acid substitution (Lys43Met) | Moderate | | | |
| 5 | 22206266 | 13/A | 2.8 ± 0.9 | 67/G | 1.5 ± 0.3 | **$4.36 \times 10^{-2}$** | **7.17** | **$9.19 \times 10^{-3}$** | 2.09 | $1.53 \times 10^{-1}$ | ✓ | intergenic region variant | Moderate | | | |
| 5 | 24935391 | 73/C | 1.7 ± 0.3 | 7/T | 1.3 ± 0.3 | $6.27 \times 10^{-1}$ | 0.28 | $5.96 \times 10^{-1}$ | 0.58 | $4.49 \times 10^{-1}$ | | amino acid substitution (Ala761Thr) | Moderate | AT5G62090 | *SLK2* | Seuss-like 2 |
| 5 | 24938566 | 11/C | 1.9 ± 0.8 | 69/G | 1.6 ± 0.3 | $6.99 \times 10^{-1}$ | 0.14 | $7.12 \times 10^{-1}$ | 0.03 | $8.66 \times 10^{-1}$ | | 5'-UTR premature start codon gain variant | Moderate | | | |
| 5 | 24938570 | 11/G | 1.9 ± 0.8 | 69/T | 1.6 ± 0.3 | $6.99 \times 10^{-1}$ | 0.14 | $7.12 \times 10^{-1}$ | 0.03 | $8.66 \times 10^{-1}$ | | 5'-UTR variant | Moderate | | | |
| 5 | 24938629 | 6/T | 1.8 ± 0.7 | 74/C | 1.7 ± 0.3 | $8.88 \times 10^{-1}$ | 0.02 | $9.01 \times 10^{-1}$ | 0.00 | $9.87 \times 10^{-1}$ | | splice region variant | **High** | | | |
| 5 | 24938639 | 70/G | 1.7 ± 0.3 | 10/T | 1.2 ± 0.5 | $4.99 \times 10^{-1}$ | 0.49 | $4.88 \times 10^{-1}$ | 0.84 | $3.62 \times 10^{-1}$ | | 5'-UTR variant | Moderate | | | |
| 5 | 24938702 | 73/A | 1.7 ± 0.3 | 7/G | 1.4 ± 0.6 | $6.83 \times 10^{-1}$ | 0.19 | $6.68 \times 10^{-1}$ | 0.96 | $3.29 \times 10^{-1}$ | | 5'-UTR variant | Moderate | | | |
| 5 | 24938811 | 21/T | 1.7 ± 0.6 | 59/A | 1.7 ± 0.3 | $9.96 \times 10^{-1}$ | 0.00 | $9.81 \times 10^{-1}$ | 0.00 | $9.81 \times 10^{-1}$ | | 5'-UTR variant | Moderate | | | |
| 5 | 24938813 | 74/A | 1.8 ± 0.3 | 6/T | 0.6 ± 0.2 | $2.25 \times 10^{-1}$ | 1.53 | $2.20 \times 10^{-1}$ | **3.09** | **$8.30 \times 10^{-2}$** | | 5'-UTR variant | Moderate | | | |
| 5 | 24938850 | 9/A | 2.1 ± 1 | 71/G | 1.6 ± 0.3 | $5.55 \times 10^{-1}$ | 0.34 | $5.63 \times 10^{-1}$ | 0.01 | $9.05 \times 10^{-1}$ | | 5'-UTR variant | Moderate | | | |
| 5 | 24938958 | 74/A | 1.8 ± 0.3 | 6/C | 0.6 ± 0.2 | $1.97 \times 10^{-1}$ | 2.00 | $1.62 \times 10^{-1}$ | 0.04 | $8.36 \times 10^{-1}$ | | 5'-UTR premature start codon gain variant | Moderate | | | |
| 5 | 24938984 | 8/C | 2.3 ± 1.1 | 72/T | 1.6 ± 0.3 | $4.21 \times 10^{-1}$ | 0.66 | $4.18 \times 10^{-1}$ | 0.01 | $9.25 \times 10^{-1}$ | | 5'-UTR premature start codon gain variant | Moderate | | | |
| 5 | 24939005 | 9/C | 2.1 ± 1 | 71/T | 1.6 ± 0.3 | $5.55 \times 10^{-1}$ | 0.34 | $5.63 \times 10^{-1}$ | 0.01 | $9.05 \times 10^{-1}$ | | 5'-UTR variant | Moderate | | | |
| 5 | 24939398 | 68/T | 1.8 ± 0.3 | 12/C | 1.1 ± 0.4 | $3.43 \times 10^{-1}$ | 0.98 | $3.25 \times 10^{-1}$ | 0.62 | $4.33 \times 10^{-1}$ | | Promoter variant | Moderate | | | |
| 5 | 24942983 | 79/T | 1.7 ± 0.3 | 1/TG | 0.4 | | 0.39 | $5.32 \times 10^{-1}$ | 0.05 | $8.19 \times 10^{-1}$ | | frameshift variant (Glu633_Ile634) | **High** | AT5G62110 | | Homeodomain-like superfamily protein |
| 5 | 24942989 | 43/T | 2.2 ± 0.5 | 37/C | 1.1 ± 0.2 | **$2.92 \times 10^{-2}$** | **5.44** | **$2.25 \times 10^{-2}$** | 0.52 | $4.73 \times 10^{-1}$ | | amino acid substitution (Asp631Gly) | Moderate | | | |
| 5 | 24942996 | 2/A | 2.4 ± 2.1 | 78/C | 1.7 ± 0.3 | $6.49 \times 10^{-1}$ | 0.16 | $6.88 \times 10^{-1}$ | **3.23** | **$7.67 \times 10^{-2}$** | | stop codon gain (Glu629*) | **High** | | | |
| 5 | 24943410 | 70/C | 1.8 ± 0.3 | 10/T | 1 ± 0.3 | $3.06 \times 10^{-1}$ | 1.41 | $2.39 \times 10^{-1}$ | 0.02 | $9.00 \times 10^{-1}$ | | amino acid substitution (Met587Ile) | Moderate | | | |
| 5 | 24943453 | 39/A | 2 ± 0.5 | 41/G | 1.4 ± 0.3 | $1.95 \times 10^{-1}$ | 1.39 | $2.43 \times 10^{-1}$ | 0.13 | $7.16 \times 10^{-1}$ | | amino acid substitution (Phe573Ser) | Moderate | | | |
| 5 | 24943460 | 28/T | 2.3 ± 0.6 | 52/C | 1.3 ± 0.3 | **$6.40 \times 10^{-2}$** | **3.45** | **$6.74 \times 10^{-2}$** | 0.24 | $6.23 \times 10^{-1}$ | | amino acid substitution (Asn571Asp) | Moderate | | | |
| 5 | 24943573 | 66/G | 1.7 ± 0.3 | 6/A | 0.6 ± 0.2 | $2.21 \times 10^{-1}$ | 1.06 | $3.52 \times 10^{-1}$ | 0.20 | $8.20 \times 10^{-1}$ | | amino acid substitution (Thr565Ser) | Moderate | | | |
| 5 | 24943599 | 30/T | 2.1 ± 0.6 | 50/C | 1.4 ± 0.3 | $1.38 \times 10^{-1}$ | 2.21 | $1.41 \times 10^{-1}$ | 0.00 | $9.88 \times 10^{-1}$ | | amino acid substitution (Ile556Met) | Moderate | | | |
| 5 | 24944039 | 36/A | 1.9 ± 0.5 | 44/G | 1.5 ± 0.3 | $3.16 \times 10^{-1}$ | 1.10 | $2.97 \times 10^{-1}$ | 0.32 | $5.71 \times 10^{-1}$ | | amino acid substitution (Phe477Leu) | Moderate | | | |
| 5 | 24944044 | 66/A | 1.8 ± 0.3 | 14/C | 1.1 ± 0.3 | $2.65 \times 10^{-1}$ | 1.28 | $2.62 \times 10^{-1}$ | 1.67 | $2.01 \times 10^{-1}$ | | amino acid substitution (Val475Gly) | Moderate | | | |

| | | | | | | | | | | | | | |
|---|---|---|---|---|---|---|---|---|---|---|---|---|---|
| 5 | 24944387 | 42/A | 2.2 ± 0.5 | 38/G | 1.1 ± 0.2 | **$2.59 \times 10^{-2}$** | **6.22** | **$1.50 \times 10^{-2}$** | 1.70 | $1.96 \times 10^{-1}$ | | amino acid substitution (Leu419Pro) | Moderate |
| 5 | 24944973 | 16/T | 2 ± 0.8 | 64/C | 1.6 ± 0.3 | $5.16 \times 10^{-1}$ | 0.96 | $3.30 \times 10^{-1}$ | 0.01 | $9.05 \times 10^{-1}$ | | amino acid substitution (Glu289Lys) | Moderate |
| 5 | 24945075 | 2/T | 8.8 ± 0.5 | 78/C | 1.5 ± 0.3 | **$3.52 \times 10^{-7}$** | **32.16** | **$2.76 \times 10^{-7}$** | **12.76** | **$6.36 \times 10^{-4}$** | ✓ | synonymous variant | Moderate |
| 5 | 24945204 | 7/A | 2.2 ± 0.7 | 73/G | 1.6 ± 0.3 | $4.86 \times 10^{-1}$ | 0.38 | $5.39 \times 10^{-1}$ | 0.14 | $7.09 \times 10^{-1}$ | | amino acid substitution (Ser212Pro) | Moderate |
| 5 | 24945961 | 13/A | 1.7 ± 0.8 | 67/G | 1.7 ± 0.3 | $9.42 \times 10^{-1}$ | 0.02 | $8.92 \times 10^{-1}$ | 0.41 | $5.24 \times 10^{-1}$ | | Promoter variant | Moderate |
| 5 | 24946218 | 69/G | 1.8 ± 0.3 | 11/A | 0.9 ± 0.3 | $2.04 \times 10^{-1}$ | 1.89 | $1.73 \times 10^{-1}$ | 0.16 | $6.91 \times 10^{-1}$ | | Promoter variant | Moderate |
| 5 | 24946431 | 69/C | 1.8 ± 0.3 | 11/T | 0.9 ± 0.3 | $2.20 \times 10^{-1}$ | 1.79 | $1.85 \times 10^{-1}$ | 0.09 | $7.65 \times 10^{-1}$ | | Promoter variant | Moderate |
| 5 | 24946434 | 20/T | 2.9 ± 0.8 | 60/C | 1.3 ± 0.2 | **$2.94 \times 10^{-3}$** | **10.85** | **$1.54 \times 10^{-3}$** | 2.92 | **$9.19 \times 10^{-2}$** | | Promoter variant | Moderate |
| 5 | 24946462 | 66/C | 1.8 ± 0.3 | 14/T | 0.9 ± 0.2 | $1.27 \times 10^{-1}$ | 1.78 | $1.86 \times 10^{-1}$ | 2.27 | $1.36 \times 10^{-1}$ | | Promoter variant | Moderate |
| 5 | 24946486 | 26/ATG | 1.7 ± 0.5 | 54/A | 1.7 ± 0.3 | $9.08 \times 10^{-1}$ | 0.02 | $8.91 \times 10^{-1}$ | 0.55 | $4.59 \times 10^{-1}$ | | Promoter variant | Moderate |

The results of a local association study (LAS) of the variation in relative fold-change (RFC) in expression levels of *DREB1A* in 80 *A. thaliana ecotypes* in the second drought (D2) versus the first drought (D1) separated by a recovery interval, calculated in Tassel separately for the individual genomic regions are presented. The polymorphisms in coding sequences and upstream regulatory regions (5'-UTR and promoters within 1 kb from the start codon) of genes directly associated with the top most significant ($P < 10^{-4}$) single nucleotide polymorphisms (SNPs) detected by a genome-wide association study (GWAS) (see Table 1) were downloaded from the 1001 genomes database (https://tools.1001genomes.org/polymorph/). Data of only 80 out of the 115 ecotypes used for the expression GWAS (see Figure 1, Table 1) were available in the 1001 genomes database. The frequency of the tolerant and sensitive alleles among the 80 ecotypes is shown with the average REL. The *F*-statistic and *P*-values of the generalized linear model (GLM) and the mixed linear model (MLM) methods of LAS calculations are shown. The effects of the GWAS-detected SNPs and additional polymorphisms in each gene mined from the 1001 genomes database are shown in terms of amino acid substitutions, frameshift, promoter, or 5'-UTR variations.

**Table S10 List of oligonucleotides used in this study**

| AGI code | Primer name | Sequence (5'--> 3') |
|---|---|---|
| **Real time q-PCR** | | |
| AT4G25480 | *DREB1A*_RT_Fw | GCTTCCGTCCGTACAGTGGA |
| | *DREB1A*_RT_Rv | AGCCCACACTTATACTGAAACTGAA |
| AT4G33150 | *LKR*_RT_Fw | CAGCTTTGTCTCCAACTCCAAGT |
| | *LKR*_RT_Rv | AAAGCTGATGGAGAAGGCAGAAT |
| AT2G18050 | *HIS1-3*_RT_Fw | CTCAGTGATCAAGCAGCGGA |
| | *HIS1-3*_RT_Rv | AGTGAAGAAGGCGAGACAGC |
| AT3G52590 | *UBQ1* _RT_Fw | TCGTAAGTACAATCAGGATAAGATG |
| | *UBQ1* _RT_Rv | CACTGAAACAAGAAAAACAAACCCT |
| AT1G01340 | AT1G01340_RT_Fw | ACGCGCTTCACAATCTGAGG |
| | AT1G01340_RT_Rv | AGTCAGGCTCGGTGGGTTTT |
| AT1G14220 | AT1G14220_RT_Fw | CGTAATGCCCAGCTTCATCAGA |
| | AT1G14220_RT_Rv | CTCTAGGCAAAACTGGACATTCGA |
| AT3G03300 | AT3G03300_RT_Fw | GCATCCTGTGAGAGAGTTGACA |
| | AT3G03300_RT_Rv | CCTTAGCTTTCACCTCAACCGT |
| AT3G10845 | AT3G10845_RT_Fw | ACAGAGGAATGGTAAAAGTTTGCGC |
| | AT3G10845_RT_Rv | TGTGTTGGAGAGGGTATGGAGC |
| AT3G19340 | AT3G19340_RT_Fw | GGGGATAGTTGCTCGGGACA |
| | AT3G19340_RT_Rv | TGAATTCAAGAAGGTGTGACGCC |
| AT3G61720 | AT3G61720_RT_Fw | GTGCAACTACCGTGGTTCCG |
| | AT3G61720_RT_Rv | CAGAACATGAGGTCTTCGTTGCT |
| AT2G24960 | AT2G24960_RT_Fw | AGAGCAGGTGAGTAGAGGGAAT |
| | AT2G24960_RT_Rv | CATGTCAGTCTGCAACCCGA |
| AT3G26490 | AT3G26490_RT_Fw | CGAGAAGAGAGGAAGGAGGACAAG |
| | AT3G26490_RT_Rv | GGATCGCACACAACGCCATA |
| AT3G02520 | AT3G02520_RT_Fw | CCACTGCATCTTTGGCCGAG |
| | AT3G02520_RT_Rv | CAGCAGCTTCTTTCCTCTCAGC |
| AT1G14220 | AT1G14220_RT_Fw | CGTAATGCCCAGCTTCATCAGA |
| | AT1G14220_RT_Rv | CTCTAGGCAAAACTGGACATTCGA |
| AT1G56660 | AT1G56660_RT_Fw | TGGAGGAGAAGGCAGAAATCGA |
| | AT1G56660_RT_Rv | ACATAACATTAAGCTGCTCCCATCA |
| AT1G65630 | AT1G65630_RT_Fw | GCTTGTAGAGGAATGCTGCACA |
| | AT1G65630_RT_Rv | CTCAGACACGGCCGATGGTA |
| AT2G19120 | AT2G19120_RT_Fw | ACCGATCATAGGCCTGCACA |
| | AT2G19120_RT_Rv | CCGTCTTGAGAGTCTTCACATCG |
| AT4G00320 | AT4G00320_RT_Fw | GCGGGTCAGATAAAGCAGGT |
| | AT4G00320_RT_Rv | CATCACATCTTCATCCTCTGGTGT |
| AT4G16490 | AT4G16490_RT_Fw | TCTGGGGTATCTTCGGGAGC |
| | AT4G16490_RT_Rv | AGACTGATGAAACCTCCACATGG |
| AT4G25970 | AT4G25970_RT_Fw | ACAGTGCTAGGTCCTTGGAGA |
| | AT4G25970_RT_Rv | GCAGTTCAGGGCTCAAGTACA |
| AT1G66960 | AT1G66960_RT_Fw | TGCAGAGTACCGGAAAGCTG |
| | AT1G66960_RT_Rv | GCCTGGCAAGATGGAAGACC |
| AT1G67000 | AT1G67000_RT_Fw | GCCACCTAGACCTGTTTTGCA |
| | AT1G67000_RT_Rv | GGCACTTGAACTCTCCTCTGAAT |
| AT3G61600 | AT3G61600_RT_Fw | TGGCATTCTCCATCTCAGAGCA |
| | AT3G61600_RT_Rv | AGCGAGAGATTTAGCAGCAGGT |
| AT5G19200 | AT5G19200_RT_Fw | TCAATCATAGCCGCATCATCTGG |
| | AT5G19200_RT_Rv | GCATGTCACTGTGAATTTTCCTGC |
| AT5G54660 | AT5G54660_RT_Fw | AGCAACAAGAAAGCGGCGAC |

| | | |
|---|---|---|
| | AT5G54660_RT_Rv | AGCTCGAGACGGCGGACATA |
| AT5G62110 | AT5G62110_RT_Fw | AAGTCTAGGCGCCATGAACTG |
| | AT5G62110_RT_Rv | AGCAGACCATCCGTTTGTGAA |
| AT3G48440 | AT3G48440_RT_Fw | TGCAAATTCGGTCCAGCTTGT |
| | AT3G48440_RT_Rv | TCTACAATGGCCTGGGAGCT |
| AT3G61610 | AT3G61610_RT_Fw | CAGGAGAAGAATGGACCGGC |
| | AT3G61610_RT_Rv | GGTGGCTCTGCTCTTCCAAT |
| AT5G49430 | AT5G49430_RT_Fw | ACCGAAGTGTGGAAGCATTACG |
| | AT5G49430_RT_Rv | GAAGGTAGCGTCCGGTCGAA |
| **Homozygosity assessment** | | |
| SALK T-DNA left border primer | SALK-LB-1.3 | ATTTTGCCGATTTCGGAAC |
| SAIL T-DNA left border primer | SAIL-LB-1 | TAGCATCTGAATTTCATAACCAATCTCGATACAC |
| Wisc T-DNA left border primer | WiscDsLoxHs LB | TGATCCATGTAGATTTCCCGGACATGAAG |
| AT1G65630 | SALK_063352C_KO_LP | TTCAAGCTTAAGAGTGTGGCC |
| | SALK_063352C_KO_RP | AAGAAGGAAACCAGGAGCAAG |
| AT1G65630 | WiscDsLoxHs138_01D_KO_LP | ACACATACGTGATACGCCCAG |
| | WiscDsLoxHs138_01D_KO_RP | GCTTGTAGAGGAATGCTGCAC |
| AT4G00320 | SALK_031180C_KO_LP | AAAACCCTGAATTTGGTTTGG |
| | SALK_031180C_KO_RP | TTGCATATTCCCATGAAAAGC |
| AT2G19120 | SALK_032807C_KO_LP | CAAGAACCAGGTTTGATGGTG |
| | SALK_032807C_KO_RP | AGTTTGGACCAGAGTGTCAGC |
| AT4G16490 | SALK_121505C_KO_LP | GAATTTAGCTCGAAAAATCAGAATC |
| | SALK_121505C_KO_RP | GAAACTCCGACGCATACTCTG |
| AT1G56660 | SAIL_163_D09_KO_LP | TCTAATCGGTTGAGCCAACTG |
| | SAIL_163_D09_KO_RP | AGGGAAAGAAAGGAAAGGGAG |
| AT4G25970 | SALK_055283C_KO_LP | CTTCCTGTGAGTTGATCGAGC |
| | SALK_055283C_KO_RP | CCCTAATCCAACCCAAAGATC |
| AT5G11780 | SALK_070669C_KO_LP | GATGGGAACAAATCACTCACG |
| | SALK_070669C_KO_RP | GCTGATTTGAGTCGAGTTTGG |
| AT3G23480 | SALK_073503C_KO_LP | TCCATCACATCCGCAAATATC |
| | SALK_073503C_KO_RP | GCAAGGTTTGATTTTCGTGAG |
| AT3G19340 | SALK_048030C_KO_LP | GTTTTGTTCCTCCTCAGGGTC |
| | SALK_048030C_KO_RP | TGGGAAACGTGTTCTAACCTG |
| AT3G03300 | SALK_064627C_KO_LP | TGAATCATCTGGAAGAGGTGG |
| | SALK_064627C_KO_RP | CTTCACAGGAGTTTTTGGCTG |
| AT3G61720 | SALK_021857C_KO_LP | TGACGTTCCGATAAAACAACC |
| | SALK_021857C_KO_RP | ACAACACGAAGGATGCATAGG |
| AT1G01340 | SALK_015952C_KO_LP | CCTGTGCTCTACACCGAGAAC |
| | SALK_015952C_KO_RP | AGCATTCTCCCTCAGGTTTTC |
| AT2G27720 | SALK_205405C_KO_LP | ACGTTGTCGTTCATTTGGAAC |
| | SALK_205405C_KO_RP | CCAGATGTAGCAGAAGCAACC |
| AT3G02520 | SALK_084141C_KO_LP | GTCTTAAACCAAAAGCGGACC |
| | SALK_084141C_KO_RP | AGGAGCTGTCATGATGAATGC |
| AT4G03430 | SALK_208743C_KO_LP | TTTTCTTTCTTCGCCTTCTCC |
| | SALK_208743C_KO_RP | CAAAAAGCCCATTAAAAAGCC |
| AT2G24960 | SALK_011398C_KO_LP | TTGCCATAATTTTCCAACTCG |
| | SALK_011398C_KO_RP | ATCACAGAGATGATGGTTCGG |
| AT1G14220 | SALK_059417C_KO_LP | TTGAACTGGGATCCGATAATG |
| | SALK_059417C_KO_RP | TGGAGCCTATTGCGATACATC |

| | | |
|---|---|---|
| AT3G10845 | SALK_026462C_KO_LP | ACTCACTTGGCTGTTAAACGC |
| | SALK_026462C_KO_RP | AGCGCAAACTTTTACCATTCC |
| AT1G66960 | SALK_022044C_KO_LP | TTGATATGGGATGCTCTCCAC |
| | SALK_022044C_KO_RP | ATCAGTTGGCATTGTGGAAAG |
| AT5G19200 | SALK_086285C_KO_LP | CTGACCAGCTTGAGAGGACAC |
| | SALK_086285C_KO_RP | GCTCACACACCTTCTGTCCTC |
| AT3G61600 | SALK_044446C_KO_LP | CCAAAGCCAAAAGAGTAAGGG |
| | SALK_044446C_KO_RP | TCAAGAGCACATGAGATGGTG |
| AT1G67000 | SAIL_842_D05_KO_LP | AACACCTGTTAATGCACCCAG |
| | SAIL_842_D05_KO_RP | CTTCAAGCTCAACTGTAGCGG |
| AT1G67000 | SALK_093030C_KO_LP | CCTCGAACTCACTGGTGAGAC |
| | SALK_093030C_KO_RP | TGTCATTTTCTTCACTTCCGC |
| AT3G61610 | SALK_094541C_KO_LP | TTTGCAAGGAACAAAATTTGG |
| | SALK_094541C_KO_RP | CTGGTCTGAGAAATCACCAGG |
| AT1G59620 | SALK_024312C_KO_LP | TGTTCATCAGGTAACCTTGGC |
| | SALK_024312C_KO_RP | GGTTACCGTTGGCTCTTAAGG |
| AT5G49430 | SALK_093473C_KO_LP | TCCAGTATGTCCACGGAGAAC |
| | SALK_093473C_KO_RP | ACCTGGTGGAGCTTCTAAAGC |
| AT3G48430 | SALK_001018C_KO_LP | TCATATACAAGGCGTTCGGTC |
| | SALK_001018C_KO_RP | CAGTTGCAACTCTGGAGAAGG |
| AT5G62110 | SALK_209177C_KO_LP | TTCCTTTCCAGTTGATCATCG |
| | SALK_209177C_KO_RP | CACTCCATGTACGATGGATCC |
| AT5G35750 | WiscDsLox504C06_KO_LP | TAAGATCGACGGTGTTATCGC |
| | WiscDsLox504C06_KO_RP | TCATGGTAAAATCCCATCTGC |

### Table S11 Differential methylation between tolerant (high induction) and sensitive (low induction) ecotypes under recurrent droughts

| AGI code | Type of gene | Gene Symbol | HIG vs LIG CpG methylation difference | | | | | HIG vs LIG CHG methylation difference | | | | | HIG vs LIG CHH methylation difference | | | | |
|---|---|---|---|---|---|---|---|---|---|---|---|---|---|---|---|---|---|
| | | | MD | *p*-value | *q*-value | Methylation position | Distance to TSS | MD | *p*-value | *q*-value | Methylation position | Distance to TSS | MD | *p*-value | *q*-value | Methylation position | Distance to TSS |
| **Drought-1** | | | | | | | | | | | | | | | | | |
| AT3G55650 | protein coding gene | | 52.86 | $3.84 \times 10^{-37}$ | $3.63 \times 10^{-36}$ | mRNA-TTS | 766 | | | | | | | | | | |
| AT5G31804 | transposable element gene | | 28.69 | $4.90 \times 10^{-11}$ | $8.21 \times 10^{-11}$ | Intergenic | -5287 | | | | | | | | | | |
| AT5G31804 | transposable element gene | | 27.42 | $2.96 \times 10^{-19}$ | $9.21 \times 10^{-19}$ | Intergenic | -5387 | | | | | | | | | | |
| AT2G07700 | transposable element gene | | -28.71 | $1.40 \times 10^{-12}$ | $2.63 \times 10^{-12}$ | Intergenic | -1350 | | | | | | | | | | |
| AT3G06365 | novel transcribed region | | | | | | | 31.9 | $2.49 \times 10^{-34}$ | $6.94 \times 10^{-33}$ | lnc_RNA-TTS | 836 | | | | | |
| AT3G06365 | novel transcribed region | | | | | | | 25.2 | | | transcript-exon | 2736 | | | | | |
| AT4G06700 | transposable element gene | | | | | | | -36.6 | $1.47 \times 10^{-43}$ | $2.33 \times 10^{-41}$ | mRNA-promoter-TSS | -849 | | | | | |
| **Drought-2** | | | | | | | | | | | | | | | | | |
| AT5G20180 | protein coding gene | *bL36m* | 93.2 | $5.05 \times 10^{-34}$ | $5.27 \times 10^{-32}$ | | -1157 | | | | | | | | | | |
| AT1G23190 | protein coding gene | *PGM3* | 87.1 | $1.21 \times 10^{-28}$ | $8.00 \times 10^{-27}$ | | -1357 | | | | | | | | | | |
| AT5G06285 | lncRNA | | 86.0 | $1.19 \times 10^{-28}$ | $7.90 \times 10^{-27}$ | | -2479 | | | | | | | | | | |
| AT5G39140 | protein coding gene | | 79.7 | $2.95 \times 10^{-37}$ | $3.06 \times 10^{-34}$ | mRNA-promoter-TSS | 310 | | | | | | | | | | |
| AT2G09975 | anti sense lncRNA | | 76.5 | $4.64 \times 10^{-46}$ | $5.51 \times 10^{-43}$ | lnc RNA-promoter-TSS | -649 | | | | | | | | | | |
| AT1G07500 | protein coding gene | | 75.7 | $4.15 \times 10^{-60}$ | $1.60 \times 10^{-57}$ | | -1420 | | | | | | | | | | |
| AT1G71240 | protein coding gene | | 75.6 | $1.52 \times 10^{-65}$ | $7.01 \times 10^{-63}$ | mRNA-intron | 1038 | | | | | | | | | | |
| AT3G01720 | protein coding gene | *AtSERGT1* | 73.1 | $2.08 \times 10^{-43}$ | $3.68 \times 10^{-41}$ | mRNA-exon | 1911 | | | | | | | | | | |
| AT2G01960 | protein coding gene | *TET14* | 73.1 | $5.56 \times 10^{-23}$ | $2.20 \times 10^{-21}$ | mRNA-TTS | 2072 | | | | | | | | | | |
| AT5G14580 | protein coding gene | | 72.6 | $1.53 \times 10^{-18}$ | $4.01 \times 10^{-17}$ | mRNA-intron | 2440 | | | | | | | | | | |
| AT2G44540 | protein coding gene | *GH9B9* | 72.4 | $2.24 \times 10^{-75}$ | $1.33 \times 10^{-72}$ | mRNA-promoter-TSS | 59 | | | | | | | | | | |
| AT2G21385 | protein coding gene | | 71.7 | $6.91 \times 10^{-32}$ | $6.01 \times 10^{-30}$ | | -1326 | | | | | | | | | | |
| AT2G40770 | protein coding gene | | 71.3 | $4.80 \times 10^{-23}$ | $1.90 \times 10^{-21}$ | mRNA-intron | 2464 | | | | | | | | | | |
| AT3G45880 | protein coding gene | | 71.2 | $5.76 \times 10^{-16}$ | $1.17 \times 10^{-14}$ | | -1942 | | | | | | | | | | |
| AT2G34680 | protein coding gene | | 70.7 | $6.04 \times 10^{-32}$ | $5.28 \times 10^{-30}$ | mRNA-intron | 3732 | | | | | | | | | | |
| AT1G66730 | protein coding gene | *LIG6* | 70.0 | $3.82 \times 10^{-17}$ | $8.73 \times 10^{-16}$ | mRNA-exon | 2070 | | | | | | | | | | |
| AT1G05570 | protein coding gene | *CALS1* | 69.2 | $1.55 \times 10^{-31}$ | $1.28 \times 10^{-29}$ | mRNA-intron | 755 | | | | | | | | | | |
| AT1G31740 | protein coding gene | *BGAL15* | 68.4 | $5.40 \times 10^{-29}$ | $3.71 \times 10^{-27}$ | mRNA-exon | 1688 | | | | | | | | | | |

| | | | | | | | | | | | | |
|---|---|---|---|---|---|---|---|---|---|---|---|---|
| AT3G54360 | protein coding gene | | 67.2 | $1.09 \times 10^{-16}$ | $2.38 \times 10^{-15}$ | mRNA-intron | 1462 | | | | | |
| AT5G42270 | protein coding gene | *VAR1* | 66.9 | $1.02 \times 10^{-36}$ | $1.30 \times 10^{-34}$ | mRNA-exon | 1376 | | | | | |
| AT2G27610 | protein coding gene | | 66.7 | $1.30 \times 10^{-21}$ | $4.62 \times 10^{-20}$ | mRNA-TTS | 1982 | | | | | |
| AT4G14760 | protein coding gene | *NET1B* | 66.2 | $3.96 \times 10^{-22}$ | $1.48 \times 10^{-20}$ | mRNA-exon | 2995 | | | | | |
| AT1G16280 | protein coding gene | *RH36* | 66.1 | $3.03 \times 10^{-26}$ | $1.62 \times 10^{-24}$ | mRNA-TTS | 1628 | | | | | |
| AT5G48910 | protein coding gene | *LPA66* | 65.6 | $1.83 \times 10^{-34}$ | $1.94 \times 10^{-32}$ | mRNA-exon | 1497 | | | | | |
| AT5G33234 | transposable element gene | | 65.0 | $6.94 \times 10^{-40}$ | $1.04 \times 10^{-37}$ | mRNA-promoter-TSS | -132 | | | | | |
| AT4G39370 | protein coding gene | *UBP27* | 64.9 | $9.79 \times 10^{-21}$ | $3.21 \times 10^{-19}$ | mRNA-TTS | 1713 | | | | | |
| AT3G56150 | protein coding gene | *EIF3C* | 64.7 | $1.54 \times 10^{-23}$ | $6.36 \times 10^{-22}$ | | -1476 | | | | | |
| AT5G50990 | protein coding gene | | 64.6 | $2.44 \times 10^{-40}$ | $3.71 \times 10^{-38}$ | | -1268 | | | | | |
| AT3G25460 | protein coding gene | | 64.5 | $3.20 \times 10^{-30}$ | $2.41 \times 10^{-28}$ | mRNA-promoter-TSS | 115 | | | | | |
| AT4G19003 | protein coding gene | *VPS25* | 64.3 | $1.25 \times 10^{-25}$ | $6.35 \times 10^{-24}$ | mRNA-TTS | 1296 | | | | | |
| AT4G02580 | protein coding gene | | 63.9 | $9.11 \times 10^{-41}$ | $1.43 \times 10^{-38}$ | | -1878 | | | | | |
| AT3G17205 | protein coding gene | *UPL6* | 63.7 | $3.15 \times 10^{-13}$ | $4.59 \times 10^{-12}$ | mRNA-exon | 2557 | | | | | |
| AT5G25760 | protein coding gene | *PEX4* | 63.2 | $2.03 \times 10^{-39}$ | $2.98 \times 10^{-37}$ | mRNA-TTS | 2056 | | | | | |
| AT2G15029 | protein coding gene | | 63.2 | $5.58 \times 10^{-37}$ | $7.25 \times 10^{-35}$ | mRNA-promoter-TSS | -180 | | | | | |
| AT2G15029 | protein coding gene | | 63.2 | $5.58 \times 10^{-37}$ | $7.25 \times 10^{-35}$ | mRNA-promoter-TSS | -180 | 25.7 | $1.37 \times 10^{-7}$ | $5.74 \times 10^{-6}$ | mRNA-promoter-TSS | -180 |
| AT1G47470 | protein coding gene | | 62.9 | $1.11 \times 10^{-14}$ | $1.93 \times 10^{-13}$ | mRNA-TTS | 1399 | | | | | |
| AT1G23190 | protein coding gene | *PGM3* | 62.8 | $2.18 \times 10^{-30}$ | $1.66 \times 10^{-28}$ | | -1257 | | | | | |
| AT4G05740 | lncRNA | | 60.2 | $6.70 \times 10^{-45}$ | $1.28 \times 10^{-42}$ | Intergenic | -1854 | | | | | |
| AT3G01810 | protein coding gene | | 60.1 | $3.00 \times 10^{-19}$ | $8.43 \times 10^{-18}$ | mRNA-exon | 3210 | | | | | |
| AT5G45250 | protein coding gene | *RPS4* | 59.9 | $3.71 \times 10^{-18}$ | $9.31 \times 10^{-17}$ | mRNA-exon | 3224 | | | | | |
| AT3G46610 | protein coding gene | | 59.8 | $2.59 \times 10^{-27}$ | $1.52 \times 10^{-25}$ | mRNA-exon | 1324 | | | | | |
| AT3G54280 | protein coding gene | *RGD3* | 59.7 | $8.74 \times 10^{-28}$ | $5.34 \times 10^{-26}$ | mRNA-intron | 2075 | | | | | |
| AT4G16630 | protein coding gene | | 59.5 | $3.34 \times 10^{-20}$ | $1.03 \times 10^{-18}$ | mRNA-intron | 1997 | | | | | |
| AT5G18633 | transposable element gene | | 59.4 | $4.94 \times 10^{-38}$ | $7.07 \times 10^{-36}$ | | -1362 | | | | | |
| AT2G03380 | protein coding gene | | 59.4 | $6.05 \times 10^{-47}$ | $1.31 \times 10^{-44}$ | mRNA-TTS | 1923 | | | | | |
| AT3G58820 | protein coding gene | | 59.3 | $3.54 \times 10^{-30}$ | $2.65 \times 10^{-28}$ | mRNA-TTS | 1518 | | | | | |
| AT1G03475 | protein coding gene | *LIN2* | 59.1 | $5.44 \times 10^{-18}$ | $1.34 \times 10^{-16}$ | mRNA-exon | 1357 | | | | | |
| AT5G49720 | protein coding gene | *GH9A1* | 59.0 | $1.24 \times 10^{-28}$ | $8.18 \times 10^{-27}$ | mRNA-exon | 1717 | | | | | |
| AT1G31740 | protein coding gene | *BGAL15* | 58.8 | $1.05 \times 10^{-32}$ | $9.77 \times 10^{-31}$ | mRNA-intron | 1588 | | | | | |
| AT3G57620 | protein coding gene | | 58.8 | $1.61 \times 10^{-42}$ | $2.70 \times 10^{-40}$ | mRNA-TTS | 2018 | | | | | |
| AT1G49990 | protein coding gene | | 58.8 | $1.10 \times 10^{-34}$ | $1.19 \times 10^{-32}$ | mRNA-TTS | 1023 | | | | | |
| AT1G76140 | protein coding gene | | 58.8 | $2.46 \times 10^{-24}$ | $1.12 \times 10^{-22}$ | mRNA-exon | 2021 | | | | | |
| AT2G21720 | protein coding gene | | 58.6 | $4.86 \times 10^{-40}$ | $7.34 \times 10^{-38}$ | mRNA-exon | 1255 | | | | | |
| AT2G17510 | protein coding gene | *EMB2763* | 58.6 | $2.21 \times 10^{-21}$ | $7.71 \times 10^{-20}$ | mRNA-exon | 4249 | | | | | |
| AT2G41616 | miRNA | | 58.4 | $3.00 \times 10^{-17}$ | $6.94 \times 10^{-16}$ | | -1477 | | | | | |

| | | | | | | | |
|---|---|---|---|---|---|---|---|
| AT2G33260 | protein coding gene | | 58.2 | $1.53 \times 10^{-37}$ | $2.10 \times 10^{-35}$ | mRNA-TTS | 877 |
| AT5G01950 | protein coding gene | | 57.8 | $1.58 \times 10^{-8}$ | $1.25 \times 10^{-7}$ | mRNA-exon | 1749 |
| AT5G22830 | protein coding gene | *MGT1* | 57.4 | $4.20 \times 10^{-17}$ | $9.55 \times 10^{-16}$ | mRNA-intron | 1654 |
| AT4G16430 | protein coding gene | | 56.9 | $2.09 \times 10^{-31}$ | $1.71 \times 10^{-29}$ | mRNA-promoter-TSS | -309 |
| AT1G65190 | protein coding gene | | 56.9 | $7.82 \times 10^{-41}$ | $1.24 \times 10^{-38}$ | | -2215 |
| AT5G64640 | protein coding gene | | 56.6 | $1.08 \times 10^{-18}$ | $2.85 \times 10^{-17}$ | | -1563 |
| AT4G31150 | protein coding gene | | 56.5 | $9.69 \times 10^{-15}$ | $1.70 \times 10^{-13}$ | | -4511 |
| AT3G20790 | protein coding gene | | 56.3 | $8.53 \times 10^{-22}$ | $3.09 \times 10^{-20}$ | | -2063 |
| AT5G18650 | protein coding gene | | 56.3 | $3.78 \times 10^{-17}$ | $8.65 \times 10^{-16}$ | mRNA-TTS | 2261 |
| AT1G60200 | protein coding gene | | 56.3 | $3.98 \times 10^{-14}$ | $6.49 \times 10^{-13}$ | mRNA-exon | 2885 |
| AT4G13410 | protein coding gene | *ATCSLA15* | 56.2 | $7.02 \times 10^{-61}$ | $2.78 \times 10^{-58}$ | | -2210 |
| AT4G06632 | transposable element gene | | 56.2 | $1.12 \times 10^{-21}$ | $3.99 \times 10^{-20}$ | mRNA-exon | 890 |
| AT3G51260 | protein coding gene | *PAD1* | 55.9 | $1.02 \times 10^{-26}$ | $5.67 \times 10^{-25}$ | mRNA-promoter-TSS | -675 |
| AT4G01395 | protein coding gene | | 55.9 | $2.28 \times 10^{-33}$ | $2.26 \times 10^{-31}$ | mRNA-exon | 1422 |
| AT3G27730 | protein coding gene | | 55.9 | $2.88 \times 10^{-23}$ | $1.16 \times 10^{-21}$ | mRNA-exon | 3611 |
| AT1G63690 | protein coding gene | *SPPL2* | 55.6 | $8.20 \times 10^{-29}$ | $5.54 \times 10^{-27}$ | mRNA-intron | 2773 |
| AT4G30860 | protein coding gene | *SDG4* | 55.5 | $1.82 \times 10^{-11}$ | $2.13 \times 10^{-10}$ | mRNA-exon | 1207 |
| AT2G15240 | protein coding gene | | 55.3 | $3.76 \times 10^{-28}$ | $2.37 \times 10^{-26}$ | mRNA-intron | 1723 |
| AT2G22410 | protein coding gene | *SLO1* | 54.3 | $3.44 \times 10^{-21}$ | $1.18 \times 10^{-19}$ | mRNA-promoter-TSS | 735 |
| AT1G64780 | protein coding gene | *AMT1222* | 54.2 | $1.24 \times 10^{-21}$ | $4.43 \times 10^{-20}$ | | -4247 |
| AT1G56340 | protein coding gene | *CRT1a* | 54.0 | $2.49 \times 10^{-47}$ | $5.45 \times 10^{-45}$ | mRNA-exon | 2008 |
| AT4G28600 | protein coding gene | | 53.8 | $9.30 \times 10^{-22}$ | $3.36 \times 10^{-20}$ | mRNA-intron | 1166 |
| AT5G51340 | protein coding gene | | 53.7 | $1.24 \times 10^{-20}$ | $4.02 \times 10^{-19}$ | mRNA-exon | 1349 |
| AT5G48410 | protein coding gene | *GLR1.3* | 53.6 | $8.80 \times 10^{-29}$ | $5.92 \times 10^{-27}$ | mRNA-exon | 1084 |
| AT4G27700 | protein coding gene | | 53.3 | $1.10 \times 10^{-34}$ | $1.19 \times 10^{-32}$ | mRNA-TTS | 1177 |
| AT2G29590 | protein coding gene | | 53.2 | $1.02 \times 10^{-27}$ | $6.20 \times 10^{-26}$ | | -1394 |
| AT3G51260 | protein coding gene | *PAD1* | 53.0 | $7.72 \times 10^{-14}$ | $1.21 \times 10^{-12}$ | mRNA-exon | 1156 |
| AT3G01720 | protein coding gene | | 53.0 | $1.90 \times 10^{-31}$ | $1.56 \times 10^{-29}$ | mRNA-exon | 1811 |
| AT1G18840 | protein coding gene | *IQD3* | 52.3 | $3.45 \times 10^{-11}$ | $3.93 \times 10^{-10}$ | mRNA-exon | 2059 |
| AT3G27450 | pseudogene | | 52.0 | $2.59 \times 10^{-26}$ | $1.38 \times 10^{-24}$ | transcript-promoter-TSS | -90 |
| AT3G06930 | protein coding gene | *PRMT4B* | 51.8 | $2.16 \times 10^{-17}$ | $5.08 \times 10^{-16}$ | mRNA-intron | 1336 |
| AT2G34090 | protein coding gene | | 51.8 | $2.77 \times 10^{-12}$ | $3.61 \times 10^{-11}$ | mRNA-intron | 1219 |
| AT2G41550 | protein coding gene | | 51.7 | $4.41 \times 10^{-12}$ | $5.61 \times 10^{-11}$ | | -1878 |
| AT2G07495 | anti sense lncRNA | | 51.7 | $6.13 \times 10^{-17}$ | $1.38 \times 10^{-15}$ | mRNA-intron | 2534 |
| AT4G25130 | protein coding gene | *PMSR4* | 51.6 | $5.28 \times 10^{-14}$ | $8.51 \times 10^{-13}$ | mRNA-intron | 4139 |
| AT5G33280 | protein coding gene | | 51.6 | $6.34 \times 10^{-20}$ | $1.91 \times 10^{-18}$ | mRNA-exon | 2508 |
| AT4G23300 | protein coding gene | *CRK22* | 51.6 | $1.59 \times 10^{-47}$ | $3.51 \times 10^{-45}$ | mRNA-exon | 1626 |

| | | | | | | | |
|---|---|---|---|---|---|---|---|
| AT5G51340 | protein coding gene | | 51.2 | $3.46 \times 10^{-18}$ | $8.75 \times 10^{-17}$ | mRNA-intron | 1249 |
| AT2G25020 | pseudogene | | 51.0 | $1.56 \times 10^{-26}$ | $8.51 \times 10^{-25}$ | transcript-exon | 1062 |
| AT5G35170 | protein coding gene | | 50.9 | $7.94 \times 10^{-12}$ | $9.76 \times 10^{-11}$ | mRNA-exon | 1885 |
| AT1G55490 | protein coding gene | *CPN6B* | 50.7 | $4.10 \times 10^{-8}$ | $3.03 \times 10^{-7}$ | mRNA-exon | 1314 |
| AT5G56590 | protein coding gene | | 50.7 | $4.32 \times 10^{-19}$ | $1.19 \times 10^{-17}$ | mRNA-exon | 1487 |
| AT2G26070 | protein coding gene | | 50.2 | $5.21 \times 10^{-9}$ | $4.36 \times 10^{-8}$ | | -2963 |
| AT1G24706 | protein coding gene | *THO2* | 50.2 | $3.39 \times 10^{-13}$ | $4.92 \times 10^{-12}$ | mRNA-exon | 5313 |
| AT4G00110 | protein coding gene | *GAE3* | 50.0 | $1.57 \times 10^{-32}$ | $1.42 \times 10^{-30}$ | mRNA-TTS | 1547 |
| AT5G37560 | protein coding gene | | 49.9 | $2.12 \times 10^{-25}$ | $1.05 \times 10^{-23}$ | mRNA-promoter-TSS | -35 |
| AT1G07500 | protein coding gene | | 49.9 | $2.12 \times 10^{-20}$ | $6.67 \times 10^{-19}$ | | -1320 |
| AT2G33720 | protein coding gene | | 49.9 | $1.02 \times 10^{-24}$ | $4.80 \times 10^{-23}$ | mRNA-TTS | 2119 |
| AT5G33253 | transposable element gene | | 49.9 | $8.21 \times 10^{-36}$ | $9.99 \times 10^{-34}$ | mRNA-promoter-TSS | -73 |
| AT3G21130 | protein coding gene | | 49.7 | $2.90 \times 10^{-13}$ | $4.25 \times 10^{-12}$ | mRNA-TTS | 505 |
| AT5G11730 | protein coding gene | | 49.6 | $6.00 \times 10^{-18}$ | $1.47 \times 10^{-16}$ | mRNA-TTS | 849 |
| AT5G04720 | protein coding gene | *ADR1-L2* | 49.6 | $4.39 \times 10^{-22}$ | $1.63 \times 10^{-20}$ | mRNA-exon | 2062 |
| AT4G28880 | protein coding gene | *ckl3* | 49.4 | $2.29 \times 10^{-27}$ | $1.35 \times 10^{-25}$ | mRNA-intron | 1512 |
| AT4G09020 | protein coding gene | *ISA3* | 49.3 | $2.83 \times 10^{-9}$ | $2.45 \times 10^{-8}$ | mRNA-intron | 3542 |
| AT2G14690 | protein coding gene | | 49.3 | $2.03 \times 10^{-15}$ | $3.86 \times 10^{-14}$ | mRNA-TTS | 2904 |
| AT1G34760 | protein coding gene | *GRF11* | 49.3 | $2.84 \times 10^{-32}$ | $2.53 \times 10^{-30}$ | mRNA-exon | 1197 |
| AT5G43090 | protein coding gene | *PUM13* | 49.2 | $2.17 \times 10^{-21}$ | $7.61 \times 10^{-20}$ | mRNA-exon | 718 |
| AT1G50250 | protein coding gene | *FTSH1* | 49.2 | $6.57 \times 10^{-16}$ | $1.32 \times 10^{-14}$ | | -1896 |
| AT3G56180 | protein coding gene | | 49.2 | $4.65 \times 10^{-12}$ | $5.90 \times 10^{-11}$ | mRNA-TTS | 1784 |
| AT2G34450 | protein coding gene | | 49.1 | $1.93 \times 10^{-10}$ | $1.97 \times 10^{-9}$ | mRNA-TTS | 1141 |
| AT3G22590 | protein coding gene | *PHP* | 49.0 | $4.15 \times 10^{-24}$ | $1.83 \times 10^{-22}$ | mRNA-exon | 1053 |
| AT5G52710 | protein coding gene | | 48.7 | $4.57 \times 10^{-28}$ | $2.85 \times 10^{-26}$ | | -1253 |
| AT2G34880 | protein coding gene | *MEE27* | 48.7 | $4.69 \times 10^{-8}$ | $3.45 \times 10^{-7}$ | | -1817 |
| AT5G01935 | anti sense lncRNA | | 48.5 | $4.81 \times 10^{-19}$ | $1.32 \times 10^{-17}$ | mRNA-TTS | 462 |
| AT1G13630 | protein coding gene | | 48.4 | $9.77 \times 10^{-25}$ | $4.62 \times 10^{-23}$ | mRNA-exon | 626 |
| AT4G31480 | protein coding gene | | 48.4 | $1.93 \times 10^{-20}$ | $6.13 \times 10^{-19}$ | mRNA-exon | 797 |
| AT3G48990 | protein coding gene | *AAE3* | 48.3 | $8.21 \times 10^{-22}$ | $2.98 \times 10^{-20}$ | mRNA-exon | 1535 |
| AT4G27040 | protein coding gene | *VPS22* | 48.3 | $1.00 \times 10^{-55}$ | $3.33 \times 10^{-53}$ | mRNA-promoter-TSS | -512 |
| AT5G32726 | transposable element gene | | 48.3 | $2.07 \times 10^{-9}$ | $1.82 \times 10^{-8}$ | | -2374 |
| AT4G38510 | protein coding gene | *VAB2* | 48.3 | $9.29 \times 10^{-13}$ | $1.28 \times 10^{-11}$ | mRNA-intron | 2750 |
| AT4G27080 | protein coding gene | *PDIL5-4* | 48.2 | $2.28 \times 10^{-21}$ | $7.91 \times 10^{-20}$ | mRNA-intron | 2079 |
| AT3G23540 | protein coding gene | | 48.2 | $4.19 \times 10^{-10}$ | $4.05 \times 10^{-9}$ | mRNA-intron | 1213 |
| AT3G59470 | protein coding gene | | 47.9 | $2.85 \times 10^{-17}$ | $6.63 \times 10^{-16}$ | mRNA-exon | 904 |
| AT2G29080 | protein coding gene | *FTSH3* | 47.7 | $8.69 \times 10^{-16}$ | $1.73 \times 10^{-14}$ | mRNA-intron | 1178 |

| | | | | | | | |
|---|---|---|---|---|---|---|---|
| AT5G18510 | protein coding gene | | 47.7 | $1.55 \times 10^{-18}$ | $4.03 \times 10^{-17}$ | mRNA-TTS | 1435 |
| AT5G64650 | protein coding gene | | 47.4 | $4.66 \times 10^{-22}$ | $1.72 \times 10^{-20}$ | mRNA-TTS | 2172 |
| AT5G03835 | anti sense lncRNA | | 47.3 | $3.12 \times 10^{-15}$ | $5.80 \times 10^{-14}$ | lnc RNA-promoter-TSS | -623 |
| AT4G24560 | protein coding gene | *UBP16* | 47.3 | $7.36 \times 10^{-13}$ | $1.03 \times 10^{-11}$ | mRNA-exon | 2825 |
| AT2G13865 | transposable element gene | | 47.3 | $2.26 \times 10^{-22}$ | $8.60 \times 10^{-21}$ | mRNA-promoter-TSS | -579 |
| AT2G09975 | anti sense lncRNA | | 47.2 | $4.26 \times 10^{-15}$ | $7.82 \times 10^{-14}$ | | -3049 |
| AT5G28910 | protein coding gene | | 47.1 | $4.97 \times 10^{-13}$ | $7.06 \times 10^{-12}$ | mRNA-TTS | 2810 |
| AT5G18633 | transposable element gene | | 47.1 | $6.22 \times 10^{-17}$ | $1.39 \times 10^{-15}$ | | -1462 |
| AT5G46070 | protein coding gene | | 47.0 | $7.37 \times 10^{-13}$ | $1.03 \times 10^{-11}$ | mRNA-exon | 1832 |
| AT3G22210 | protein coding gene | | 47.0 | $1.50 \times 10^{-27}$ | $8.91 \times 10^{-26}$ | | -1284 |
| AT3G54290 | protein coding gene | | 46.8 | $2.37 \times 10^{-11}$ | $2.75 \times 10^{-10}$ | mRNA-intron | 4064 |
| AT4G27040 | protein coding gene | *VPS22* | 46.8 | $5.86 \times 10^{-51}$ | $1.47 \times 10^{-48}$ | | -1012 |
| AT1G17720 | protein coding gene | *ATB BETA* | 46.7 | $5.28 \times 10^{-15}$ | $9.60 \times 10^{-14}$ | mRNA-intron | 3088 |
| AT2G03670 | protein coding gene | *CDC48B* | 46.7 | $3.55 \times 10^{-38}$ | $5.12 \times 10^{-36}$ | mRNA-TTS | 765 |
| AT1G18450 | protein coding gene | | 46.7 | $9.92 \times 10^{-12}$ | $1.20 \times 10^{-10}$ | mRNA-intron | 1603 |
| AT1G29962 | protein coding gene | *AGL64* | 46.6 | $2.56 \times 10^{-7}$ | $1.71 \times 10^{-6}$ | mRNA-promoter-TSS | -179 |
| AT3G53930 | protein coding gene | | 46.6 | $7.86 \times 10^{-12}$ | $9.67 \times 10^{-11}$ | mRNA-TTS | 2525 |
| AT3G04830 | protein coding gene | | 46.5 | $8.75 \times 10^{-18}$ | $2.12 \times 10^{-16}$ | mRNA-intron | 1602 |
| AT1G20870 | protein coding gene | | 46.5 | $2.55 \times 10^{-51}$ | $6.51 \times 10^{-49}$ | mRNA-promoter-TSS | 24 |
| AT5G23575 | protein coding gene | | 46.5 | $7.99 \times 10^{-13}$ | $1.11 \times 10^{-11}$ | mRNA-TTS | 3166 |
| AT2G17477 | pseudogene | | 46.4 | $3.25 \times 10^{-22}$ | $1.22 \times 10^{-20}$ | mRNA-intron | 1943 |
| AT2G35100 | protein coding gene | *ARAD1* | 46.3 | $1.14 \times 10^{-18}$ | $3.02 \times 10^{-17}$ | mRNA-promoter-TSS | -850 |
| AT1G60850 | protein coding gene | *ATRPAC42* | 46.2 | $3.55 \times 10^{-16}$ | $7.31 \times 10^{-15}$ | | -1538 |
| AT3G21220 | protein coding gene | *MKK5* | 46.0 | $7.89 \times 10^{-45}$ | $1.49 \times 10^{-42}$ | mRNA-TTS | 2720 |
| AT4G09915 | lncRNA | | 45.9 | $1.88 \times 10^{-24}$ | $8.63 \times 10^{-23}$ | mRNA-TTS | 920 |
| AT2G26790 | protein coding gene | | 45.9 | $5.39 \times 10^{-55}$ | $1.66 \times 10^{-52}$ | mRNA-exon | 623 |
| AT5G00665 | novel transcribed region | | 45.9 | $1.78 \times 10^{-18}$ | $4.62 \times 10^{-17}$ | transcript-TTS | 1849 |
| AT1G42490 | pseudogene | | 45.9 | $5.21 \times 10^{-8}$ | $3.80 \times 10^{-7}$ | transcript-exon | 334 |
| AT5G65950 | protein coding gene | | 45.9 | $1.51 \times 10^{-9}$ | $1.36 \times 10^{-8}$ | mRNA-exon | 1728 |
| AT4G32920 | protein coding gene | | 45.8 | $1.34 \times 10^{-24}$ | $6.27 \times 10^{-23}$ | mRNA-intron | 4655 |
| AT3G17830 | protein coding gene | | 45.8 | $3.40 \times 10^{-17}$ | $7.81 \times 10^{-16}$ | mRNA-exon | 1898 |
| AT3G18290 | protein coding gene | *BTS* | 45.8 | $2.93 \times 10^{-20}$ | $9.09 \times 10^{-19}$ | mRNA-exon | 2485 |
| AT3G04460 | protein coding gene | *PEX12* | 45.7 | $2.84 \times 10^{-12}$ | $3.71 \times 10^{-11}$ | mRNA-intron | 1167 |
| AT3G03895 | anti sense lncRNA | | 45.6 | $3.37 \times 10^{-10}$ | $3.32 \times 10^{-9}$ | lnc RNA-promoter-TSS | -149 |
| AT4G25660 | protein coding gene | | 45.6 | $2.63 \times 10^{-16}$ | $5.51 \times 10^{-15}$ | mRNA-TTS | 1344 |
| AT3G46180 | protein coding gene | *UTR5* | 45.6 | $1.13 \times 10^{-14}$ | $1.97 \times 10^{-13}$ | mRNA-intron | 1384 |
| AT3G48810 | protein coding gene | | 45.4 | $1.27 \times 10^{-13}$ | $1.95 \times 10^{-12}$ | mRNA-promoter-TSS | 841 |

| | | | | | | | |
|---|---|---|---|---|---|---|---|
| AT4G15900 | protein coding gene | *PRL1* | 45.3 | $2.59 \times 10^{-18}$ | $6.59 \times 10^{-17}$ | mRNA-intron | 985 |
| AT5G64390 | protein coding gene | *HEN4* | 45.3 | $2.07 \times 10^{-10}$ | $2.10 \times 10^{-9}$ | mRNA-exon | 1905 |
| AT1G07707 | lncRNA | | 45.3 | $4.36 \times 10^{-7}$ | $2.82 \times 10^{-6}$ | lnc RNA-promoter-TSS | -942 |
| AT5G04890 | protein coding gene | *RTM2* | 45.3 | $8.69 \times 10^{-14}$ | $1.36 \times 10^{-12}$ | mRNA-promoter-TSS | -866 |
| AT5G04230 | protein coding gene | *PAL3* | 45.3 | $3.50 \times 10^{-45}$ | $6.92 \times 10^{-43}$ | mRNA-exon | 2017 |
| AT2G32000 | protein coding gene | | 45.2 | $3.68 \times 10^{-21}$ | $1.26 \times 10^{-19}$ | mRNA-intron | 2190 |
| AT4G20725 | transposable element gene | | 45.2 | $4.06 \times 10^{-55}$ | $1.32 \times 10^{-52}$ | transcript-TTS | 2115 |
| AT1G19100 | protein coding gene | *DMS11* | 45.1 | $1.00 \times 10^{-11}$ | $1.21 \times 10^{-10}$ | | -2132 |
| AT2G35030 | protein coding gene | | 45.1 | $1.38 \times 10^{-27}$ | $8.25 \times 10^{-26}$ | mRNA-TTS | 2183 |
| AT1G66150 | protein coding gene | *TMK1* | 45.1 | $8.47 \times 10^{-32}$ | $7.25 \times 10^{-30}$ | mRNA-exon | 2613 |
| AT2G08050 | lncRNA | | 44.9 | $5.67 \times 10^{-14}$ | $9.08 \times 10^{-13}$ | lnc RNA-TTS | 1271 |
| AT3G57170 | protein coding gene | | 44.8 | $4.03 \times 10^{-13}$ | $5.77 \times 10^{-12}$ | mRNA-intron | 1145 |
| AT1G32950 | protein coding gene | | 44.4 | $1.15 \times 10^{-50}$ | $2.85 \times 10^{-48}$ | mRNA-TTS | 655 |
| AT3G04670 | protein coding gene | *WRKY39* | 44.3 | $4.02 \times 10^{-15}$ | $7.41 \times 10^{-14}$ | mRNA-TTS | 1799 |
| AT1G12920 | protein coding gene | *ERF1-2* | 44.3 | $2.15 \times 10^{-11}$ | $2.50 \times 10^{-10}$ | | -2558 |
| AT5G39120 | protein coding gene | | 44.2 | $2.28 \times 10^{-20}$ | $7.14 \times 10^{-19}$ | mRNA-exon | 376 |
| AT2G16630 | protein coding gene | | 44.1 | $1.17 \times 10^{-15}$ | $2.28 \times 10^{-14}$ | mRNA-TTS | 1743 |
| AT4G02570 | protein coding gene | *CUL1* | 44.0 | $5.49 \times 10^{-7}$ | $3.50 \times 10^{-6}$ | mRNA-intron | 1832 |
| AT4G21326 | protein coding gene | *SBT3.12* | 43.9 | $1.03 \times 10^{-19}$ | $3.02 \times 10^{-18}$ | | -1921 |
| AT2G14570 | transposable element gene | | 43.9 | $5.46 \times 10^{-6}$ | $2.98 \times 10^{-5}$ | mRNA-promoter-TSS | -934 |
| AT5G35370 | protein coding gene | | 43.8 | $1.34 \times 10^{-19}$ | $3.90 \times 10^{-18}$ | mRNA-TTS | 2120 |
| AT1G54440 | protein coding gene | | 43.8 | $2.64 \times 10^{-17}$ | $6.16 \times 10^{-16}$ | mRNA-intron | 2398 |
| AT1G53430 | protein coding gene | | 43.7 | $2.55 \times 10^{-15}$ | $4.79 \times 10^{-14}$ | mRNA-exon | 5188 |
| AT2G33260 | protein coding gene | | 43.7 | $2.43 \times 10^{-25}$ | $1.21 \times 10^{-23}$ | mRNA-exon | 777 |
| AT1G63240 | protein coding gene | | 43.7 | $1.22 \times 10^{-20}$ | $3.96 \times 10^{-19}$ | mRNA-exon | 1577 |
| AT4G05495 | pseudogene | | 43.6 | $2.23 \times 10^{-17}$ | $5.25 \times 10^{-16}$ | transcript-promoter-TSS | -801 |
| AT5G03145 | miRNA | | 43.6 | $4.06 \times 10^{-21}$ | $1.38 \times 10^{-19}$ | mRNA-intron | 3779 |
| AT4G15025 | protein coding gene | | 43.6 | $2.10 \times 10^{-14}$ | $3.53 \times 10^{-13}$ | mRNA-promoter-TSS | -664 |
| AT5G53010 | protein coding gene | | 43.4 | $9.69 \times 10^{-20}$ | $2.86 \times 10^{-18}$ | mRNA-exon | 2186 |
| AT5G51250 | protein coding gene | | 43.3 | $3.15 \times 10^{-24}$ | $1.40 \times 10^{-22}$ | | -2092 |
| AT4G38230 | protein coding gene | *CPK26* | 43.3 | $7.19 \times 10^{-19}$ | $1.93 \times 10^{-17}$ | mRNA-exon | 1231 |
| AT1G06423 | anti sense lncRNA | | 43.3 | $4.37 \times 10^{-53}$ | $1.19 \times 10^{-50}$ | mRNA-TTS | 244 |
| AT3G01045 | anti sense lncRNA | | 43.2 | $3.37 \times 10^{-26}$ | $1.79 \times 10^{-24}$ | mRNA-TTS | 132 |
| AT3G11830 | protein coding gene | | 43.2 | $2.50 \times 10^{-24}$ | $1.13 \times 10^{-22}$ | mRNA-intron | 1794 |
| AT5G18830 | protein coding gene | *SPL7* | 43.2 | $4.69 \times 10^{-15}$ | $8.59 \times 10^{-14}$ | mRNA-exon | 2629 |
| AT1G50250 | protein coding gene | *FTSH1* | 43.2 | $1.28 \times 10^{-9}$ | $1.16 \times 10^{-8}$ | | -1796 |
| AT4G23040 | protein coding gene | | 43.1 | $5.94 \times 10^{-13}$ | $8.40 \times 10^{-12}$ | mRNA-exon | 1526 |

| | | | | | | | | | | | | |
|---|---|---|---|---|---|---|---|---|---|---|---|---|
| AT2G23348 | protein coding gene | | 43.0 | $9.05 \times 10^{-6}$ | $4.73 \times 10^{-5}$ | primary transcript-TTS | 124 | | | | | |
| AT2G38550 | protein coding gene | | 42.9 | $1.53 \times 10^{-9}$ | $1.37 \times 10^{-8}$ | mRNA-exon | 1321 | | | | | |
| AT1G21910 | protein coding gene | *DREB26* | 42.9 | $2.10 \times 10^{-31}$ | $1.71 \times 10^{-29}$ | Intergenic | -1576 | | | | | |
| AT2G34560 | protein coding gene | *CCP1* | 42.9 | $2.17 \times 10^{-13}$ | $3.23 \times 10^{-12}$ | mRNA-exon | 1263 | | | | | |
| AT3G20790 | protein coding gene | | 42.7 | $2.50 \times 10^{-19}$ | $7.07 \times 10^{-18}$ | | -2163 | | | | | |
| AT5G04895 | protein coding gene | *ABO6* | 42.7 | $3.70 \times 10^{-13}$ | $5.34 \times 10^{-12}$ | mRNA-intron | 3054 | | | | | |
| AT4G10130 | protein coding gene | | 42.7 | $2.28 \times 10^{-10}$ | $2.30 \times 10^{-9}$ | mRNA-TTS | 2592 | | | | | |
| AT5G08630 | protein coding gene | | 42.7 | $9.37 \times 10^{-10}$ | $8.64 \times 10^{-9}$ | mRNA-exon | 1120 | | | | | |
| AT2G05725 | lncRNA | | 42.6 | $4.20 \times 10^{-8}$ | $3.10 \times 10^{-7}$ | lnc RNA-promoter-TSS | -21 | | | | | |
| AT3G14690 | protein coding gene | | 42.6 | $1.09 \times 10^{-32}$ | $1.00 \times 10^{-30}$ | mRNA-exon | 1334 | | | | | |
| AT3G19360 | protein coding gene | | 42.6 | $4.26 \times 10^{-11}$ | $4.78 \times 10^{-10}$ | mRNA-intron | 1300 | | | | | |
| AT2G04700 | protein coding gene | | 42.6 | $2.35 \times 10^{-36}$ | $2.94 \times 10^{-34}$ | mRNA-promoter-TSS | -748 | | | | | |
| AT3G56190 | protein coding gene | *ALPHA-SNAP2* | 42.1 | $4.16 \times 10^{-12}$ | $5.31 \times 10^{-11}$ | mRNA-intron | 1283 | | | | | |
| AT5G11470 | protein coding gene | | 42.1 | $9.75 \times 10^{-12}$ | $1.18 \times 10^{-10}$ | mRNA-intron | 1672 | | | | | |
| AT3G60310 | protein coding gene | | 42.1 | $7.14 \times 10^{-9}$ | $5.86 \times 10^{-8}$ | mRNA-intron | 1269 | | | | | |
| AT4G08136 | pseudogene | | 42.1 | $4.86 \times 10^{-21}$ | $1.64 \times 10^{-19}$ | transcript-exon | 1550 | 26.6 | $6.62 \times 10^{-8}$ | $2.95 \times 10^{-6}$ | transcript-exon | 1550 |
| AT2G12325 | transposable element gene | | 41.9 | $1.60 \times 10^{-11}$ | $1.89 \times 10^{-10}$ | mRNA-promoter-TSS | 16 | | | | | |
| AT4G19220 | protein coding gene | | 41.8 | $7.82 \times 10^{-28}$ | $4.81 \times 10^{-26}$ | mRNA-exon | 1370 | | | | | |
| AT2G25040 | pseudogene | | 41.7 | $6.72 \times 10^{-19}$ | $1.81 \times 10^{-17}$ | mRNA-intron | 3352 | | | | | |
| AT1G09720 | protein coding gene | *NET2B* | 41.7 | $6.32 \times 10^{-27}$ | $3.63 \times 10^{-25}$ | | -2448 | | | | | |
| AT5G22785 | protein coding gene | | 41.7 | $6.21 \times 10^{-17}$ | $1.39 \times 10^{-15}$ | mRNA-TTS | 708 | | | | | |
| AT2G30105 | protein coding gene | | 41.7 | $1.60 \times 10^{-9}$ | $1.43 \times 10^{-8}$ | mRNA-exon | 3433 | | | | | |
| AT4G32450 | protein coding gene | *MEF8S* | 41.6 | $2.00 \times 10^{-21}$ | $7.04 \times 10^{-20}$ | mRNA-TTS | 1224 | | | | | |
| AT5G25260 | protein coding gene | | 41.6 | $6.65 \times 10^{-20}$ | $1.99 \times 10^{-18}$ | mRNA-TTS | 1067 | | | | | |
| AT1G50890 | protein coding gene | | 41.5 | $1.84 \times 10^{-20}$ | $5.84 \times 10^{-19}$ | mRNA-exon | 2362 | | | | | |
| AT5G05235 | lncRNA | | 41.5 | $5.87 \times 10^{-12}$ | $7.33 \times 10^{-11}$ | mRNA-exon | 1705 | | | | | |
| AT4G10110 | protein coding gene | | 41.5 | $1.92 \times 10^{-17}$ | $4.53 \times 10^{-16}$ | mRNA-TTS | 759 | | | | | |
| AT5G51230 | protein coding gene | *EMF2* | 41.5 | $1.95 \times 10^{-5}$ | $9.64 \times 10^{-5}$ | mRNA-intron | 1617 | | | | | |
| AT1G79080 | protein coding gene | | 41.4 | $1.37 \times 10^{-29}$ | $9.83 \times 10^{-28}$ | mRNA-promoter-TSS | -507 | | | | | |
| AT4G27080 | protein coding gene | *PDIL5-4* | 41.3 | $1.98 \times 10^{-12}$ | $2.62 \times 10^{-11}$ | mRNA-exon | 1779 | | | | | |
| AT1G56510 | protein coding gene | *WRR4* | 41.2 | $7.54 \times 10^{-20}$ | $2.25 \times 10^{-18}$ | | -3714 | | | | | |
| AT3G48150 | protein coding gene | *APC8* | 41.2 | $5.43 \times 10^{-12}$ | $6.82 \times 10^{-11}$ | mRNA-exon | 1018 | | | | | |
| AT5G64480 | protein coding gene | | 41.1 | $8.40 \times 10^{-14}$ | $1.32 \times 10^{-12}$ | mRNA-promoter-TSS | -479 | | | | | |
| AT3G44060 | protein coding gene | | 41.1 | $7.89 \times 10^{-21}$ | $2.61 \times 10^{-19}$ | mRNA-exon | 2940 | | | | | |
| AT1G65750 | transposable element gene | | 41.0 | $9.14 \times 10^{-25}$ | $4.35 \times 10^{-23}$ | mRNA-promoter-TSS | 79 | | | | | |
| AT1G05520 | protein coding gene | | 41.0 | $1.72 \times 10^{-21}$ | $6.05 \times 10^{-20}$ | mRNA-intron | 1667 | | | | | |
| AT4G03035 | novel transcribed region | | 41.0 | $5.42 \times 10^{-52}$ | $1.43 \times 10^{-49}$ | transcript-TTS | 2115 | | | | | |

| | | | | | | | |
|---|---|---|---|---|---|---|---|
| AT2G47070 | protein coding gene | *SPL1* | 41.0 | $1.39 \times 10^{-16}$ | $3.00 \times 10^{-15}$ | mRNA-TTS | 3756 |
| AT1G08073 | anti sense lncRNA | | 40.8 | $4.20 \times 10^{-15}$ | $7.72 \times 10^{-14}$ | | -4007 |
| AT1G11310 | protein coding gene | *MLO2* | 40.8 | $1.66 \times 10^{-13}$ | $2.51 \times 10^{-12}$ | mRNA-exon | 1344 |
| AT3G20570 | protein coding gene | *ENODL9* | 40.8 | $4.22 \times 10^{-19}$ | $1.17 \times 10^{-17}$ | mRNA-intron | 2670 |
| AT4G30320 | protein coding gene | | 40.8 | $1.56 \times 10^{-13}$ | $2.36 \times 10^{-12}$ | | -1141 |
| AT2G35900 | protein coding gene | | 40.7 | $4.22 \times 10^{-18}$ | $1.05 \times 10^{-16}$ | mRNA-intron | 1130 |
| AT1G72960 | protein coding gene | | 40.6 | $1.88 \times 10^{-33}$ | $1.89 \times 10^{-31}$ | mRNA-TTS | 415 |
| AT5G04605 | lncRNA | | 40.6 | $7.49 \times 10^{-25}$ | $3.60 \times 10^{-23}$ | Intergenic | 2353 |
| AT2G35730 | protein coding gene | | 40.6 | $4.82 \times 10^{-14}$ | $7.79 \times 10^{-13}$ | mRNA-exon | 2245 |
| AT3G59210 | protein coding gene | | 40.5 | $1.00 \times 10^{-12}$ | $1.37 \times 10^{-11}$ | mRNA-exon | 913 |
| AT1G35467 | protein coding gene | *RALFL5* | 40.5 | $2.73 \times 10^{-29}$ | $1.91 \times 10^{-27}$ | mRNA-TTS | 737 |
| AT3G04135 | anti sense lncRNA | | 40.5 | $4.52 \times 10^{-12}$ | $5.75 \times 10^{-11}$ | lnc RNA-promoter-TSS | -148 |
| AT3G03400 | protein coding gene | | 40.5 | $1.16 \times 10^{-10}$ | $1.22 \times 10^{-9}$ | mRNA-exon | 4007 |
| AT3G54350 | protein coding gene | *emb1967* | 40.5 | $4.99 \times 10^{-35}$ | $5.60 \times 10^{-33}$ | | -1070 |
| AT1G06710 | protein coding gene | | 40.4 | $2.09 \times 10^{-10}$ | $2.11 \times 10^{-9}$ | | -1422 |
| AT1G64750 | protein coding gene | *DSS1(I)* | 40.4 | $3.00 \times 10^{-28}$ | $1.91 \times 10^{-26}$ | | -1128 |
| AT3G08840 | protein coding gene | | 40.4 | $9.44 \times 10^{-12}$ | $1.15 \times 10^{-10}$ | mRNA-exon | 1036 |
| AT5G22794 | protein coding gene | | 40.3 | $4.05 \times 10^{-19}$ | $1.12 \times 10^{-17}$ | mRNA-intron | 1109 |
| AT5G51390 | protein coding gene | | 40.2 | $8.49 \times 10^{-37}$ | $1.09 \times 10^{-34}$ | mRNA-promoter-TSS | -487 |
| AT4G21323 | protein coding gene | | 40.2 | $4.59 \times 10^{-26}$ | $2.41 \times 10^{-24}$ | mRNA-exon | 1888 |
| AT4G01230 | protein coding gene | | 40.1 | $4.53 \times 10^{-55}$ | $1.45 \times 10^{-52}$ | mRNA-promoter-TSS | -835 |
| AT2G47015 | miRNA | | 40.1 | $9.56 \times 10^{-16}$ | $1.89 \times 10^{-14}$ | mRNA-TTS | 818 |
| AT5G57590 | protein coding gene | *BIO1* | 40.0 | $6.11 \times 10^{-10}$ | $5.78 \times 10^{-9}$ | mRNA-intron | 2119 |
| AT4G09955 | anti sense lncRNA | | 39.9 | $7.15 \times 10^{-24}$ | $3.08 \times 10^{-22}$ | | -1843 |
| AT1G77460 | protein coding gene | | 39.9 | $4.32 \times 10^{-16}$ | $8.81 \times 10^{-15}$ | mRNA-exon | 2209 |
| AT4G28710 | protein coding gene | *XIH* | 39.8 | $1.17 \times 10^{-16}$ | $2.54 \times 10^{-15}$ | mRNA-intron | 2153 |
| AT5G37990 | protein coding gene | | 39.8 | $9.60 \times 10^{-35}$ | $1.06 \times 10^{-32}$ | mRNA-exon | 318 |
| AT1G64570 | protein coding gene | *DUO3* | 39.7 | $4.45 \times 10^{-6}$ | $2.45 \times 10^{-5}$ | mRNA-intron | 2238 |
| AT5G13050 | protein coding gene | *5-FCL* | 39.7 | $1.40 \times 10^{-9}$ | $1.26 \times 10^{-8}$ | Intergenic | -1359 |
| AT4G12115 | pre tRNA | | 39.7 | $3.62 \times 10^{-14}$ | $5.93 \times 10^{-13}$ | mRNA-intron | 1236 |
| AT2G34090 | protein coding gene | | 39.7 | $8.77 \times 10^{-9}$ | $7.11 \times 10^{-8}$ | mRNA-exon | 919 |
| AT1G51540 | protein coding gene | | 39.7 | $1.54 \times 10^{-27}$ | $9.15 \times 10^{-26}$ | | -1106 |
| AT4G10865 | transposable element gene | | 39.7 | $1.01 \times 10^{-15}$ | $1.99 \times 10^{-14}$ | mRNA-exon | 211 |
| AT1G54530 | protein coding gene | | 39.4 | $9.30 \times 10^{-11}$ | $9.92 \times 10^{-10}$ | | -1262 |
| AT1G64050 | protein coding gene | | 39.4 | $1.51 \times 10^{-10}$ | $1.57 \times 10^{-9}$ | mRNA-exon | 3098 |
| AT4G02570 | protein coding gene | *CUL1* | 39.4 | $7.33 \times 10^{-8}$ | $5.22 \times 10^{-7}$ | mRNA-exon | 2832 |
| AT5G22340 | protein coding gene | | 39.3 | $1.27 \times 10^{-8}$ | $1.01 \times 10^{-7}$ | mRNA-intron | 1278 |

| | | | | | | | |
|---|---|---|---|---|---|---|---|
| AT3G49140 | protein coding gene | | 39.3 | $1.99 \times 10^{-13}$ | $2.97 \times 10^{-12}$ | | -1008 |
| AT1G13448 | anti sense lncRNA | | 39.3 | $1.86 \times 10^{-13}$ | $2.79 \times 10^{-12}$ | mRNA-TTS | 1108 |
| AT5G37990 | protein coding gene | | 39.3 | $1.41 \times 10^{-16}$ | $3.05 \times 10^{-15}$ | mRNA-exon | 518 |
| AT3G09710 | protein coding gene | *IQD1* | 39.3 | $6.49 \times 10^{-22}$ | $2.38 \times 10^{-20}$ | | -1067 |
| AT1G04720 | pre tRNA | | 39.2 | $7.57 \times 10^{-11}$ | $8.16 \times 10^{-10}$ | mRNA-exon | 1128 |
| AT1G72710 | protein coding gene | *CKL2* | 39.2 | $2.75 \times 10^{-35}$ | $3.11 \times 10^{-33}$ | mRNA-promoter-TSS | -846 |
| AT1G50380 | protein coding gene | | 39.2 | $8.80 \times 10^{-10}$ | $8.16 \times 10^{-9}$ | | -1815 |
| AT1G11755 | protein coding gene | *LEW1* | 39.2 | $1.03 \times 10^{-17}$ | $2.48 \times 10^{-16}$ | mRNA-TTS | 1160 |
| AT5G21070 | protein coding gene | | 39.2 | $5.98 \times 10^{-11}$ | $6.57 \times 10^{-10}$ | mRNA-TTS | 1088 |
| AT1G10450 | protein coding gene | | 39.1 | $4.17 \times 10^{-12}$ | $5.32 \times 10^{-11}$ | mRNA-exon | 1791 |
| AT3G28370 | protein coding gene | | 39.0 | $7.48 \times 10^{-29}$ | $5.07 \times 10^{-27}$ | mRNA-TTS | 3624 |
| AT3G13340 | protein coding gene | | 39.0 | $1.74 \times 10^{-13}$ | $2.62 \times 10^{-12}$ | mRNA-exon | 1621 |
| AT5G37560 | protein coding gene | | 39.0 | $6.14 \times 10^{-21}$ | $2.05 \times 10^{-19}$ | mRNA-exon | 265 |
| AT3G53340 | protein coding gene | *NF-YB1* | 39.0 | $5.42 \times 10^{-22}$ | $2.00 \times 10^{-20}$ | mRNA-TTS | 1516 |
| AT2G04940 | protein coding gene | | 38.9 | $5.00 \times 10^{-18}$ | $1.24 \times 10^{-16}$ | mRNA-intron | 3017 |
| AT1G10810 | protein coding gene | | 38.9 | $1.16 \times 10^{-10}$ | $1.23 \times 10^{-9}$ | mRNA-TTS | 1785 |
| AT5G06060 | protein coding gene | | 38.9 | $5.16 \times 10^{-11}$ | $5.71 \times 10^{-10}$ | mRNA-intron | 975 |
| AT1G26390 | protein coding gene | | 38.9 | $3.80 \times 10^{-11}$ | $4.29 \times 10^{-10}$ | mRNA-exon | 1328 |
| AT3G13330 | protein coding gene | *PA2* | 38.8 | $2.87 \times 10^{-11}$ | $3.30 \times 10^{-10}$ | mRNA-intron | 7167 |
| AT4G28470 | protein coding gene | *RPN1B* | 38.8 | $1.72 \times 10^{-17}$ | $4.10 \times 10^{-16}$ | mRNA-intron | 1531 |
| AT5G14770 | protein coding gene | | 38.7 | $1.62 \times 10^{-12}$ | $2.16 \times 10^{-11}$ | mRNA-exon | 1082 |
| AT1G68020 | protein coding gene | *ATTPS6* | 38.6 | $2.79 \times 10^{-10}$ | $2.78 \times 10^{-9}$ | mRNA-exon | 1961 |
| AT4G16190 | protein coding gene | | 38.5 | $5.55 \times 10^{-17}$ | $1.25 \times 10^{-15}$ | mRNA-TTS | 1171 |
| AT2G17115 | transposable element gene | | 38.5 | $1.38 \times 10^{-12}$ | $1.86 \times 10^{-11}$ | Intergenic | -2260 |
| AT5G08020 | protein coding gene | *RPA7B* | 38.5 | $3.62 \times 10^{-20}$ | $1.11 \times 10^{-18}$ | | -1006 |
| AT1G04247 | anti sense lncRNA | | 38.5 | $2.52 \times 10^{-9}$ | $2.19 \times 10^{-8}$ | | -1874 |
| AT5G25360 | protein coding gene | | 38.5 | $1.61 \times 10^{-17}$ | $3.85 \times 10^{-16}$ | mRNA-exon | 593 |
| AT4G39470 | protein coding gene | | 38.5 | $1.45 \times 10^{-7}$ | $9.98 \times 10^{-7}$ | mRNA-exon | 931 |
| AT3G07522 | protein coding gene | | 38.5 | $9.67 \times 10^{-9}$ | $7.80 \times 10^{-8}$ | mRNA-promoter-TSS | -577 |
| AT5G06600 | protein coding gene | *UBP12* | 38.5 | $3.41 \times 10^{-16}$ | $7.03 \times 10^{-15}$ | mRNA-exon | 1892 |
| AT4G03515 | novel transcribed region | | 38.4 | $1.64 \times 10^{-15}$ | $3.13 \times 10^{-14}$ | | -3940 |
| AT5G39150 | protein coding gene | | 38.3 | $1.90 \times 10^{-19}$ | $5.45 \times 10^{-18}$ | mRNA-exon | 853 |
| AT5G01225 | protein coding gene | | 38.3 | $1.04 \times 10^{-14}$ | $1.82 \times 10^{-13}$ | mRNA-exon | 1765 |
| AT1G61350 | protein coding gene | | 38.2 | $1.23 \times 10^{-19}$ | $3.62 \times 10^{-18}$ | mRNA-exon | 1877 |
| AT1G45180 | protein coding gene | | 38.1 | $4.19 \times 10^{-16}$ | $8.56 \times 10^{-15}$ | mRNA-exon | 2647 |
| AT4G06990 | anti sense lncRNA | | 38.1 | $1.13 \times 10^{-12}$ | $1.53 \times 10^{-11}$ | lnc RNA-promoter-TSS | -274 |
| AT5G65687 | protein coding gene | | 38.0 | $1.88 \times 10^{-10}$ | $1.92 \times 10^{-9}$ | mRNA-TTS | 2449 |

| | | | | | | | | | | | | |
|---|---|---|---|---|---|---|---|---|---|---|---|---|
| AT4G17880 | protein coding gene | *MYC4* | 38.0 | $1.96 \times 10^{-20}$ | $6.22 \times 10^{-19}$ | mRNA-exon | 2034 | | | | | |
| AT1G43170 | protein coding gene | *RP1* | 38.0 | $5.40 \times 10^{-5}$ | $2.45 \times 10^{-4}$ | mRNA-promoter-TSS | -702 | | | | | |
| AT4G14240 | protein coding gene | | 37.9 | $4.59 \times 10^{-20}$ | $1.40 \times 10^{-18}$ | mRNA-exon | 1412 | | | | | |
| AT4G02945 | novel transcribed region | | 37.9 | $2.85 \times 10^{-14}$ | $4.73 \times 10^{-13}$ | | -2877 | | | | | |
| AT3G52250 | protein coding gene | | 37.8 | $1.22 \times 10^{-9}$ | $1.11 \times 10^{-8}$ | mRNA-exon | 2413 | | | | | |
| AT1G61370 | protein coding gene | | 37.8 | $3.37 \times 10^{-19}$ | $9.44 \times 10^{-18}$ | mRNA-promoter-TSS | -529 | | | | | |
| AT5G05995 | anti sense lncRNA | | 37.7 | $5.74 \times 10^{-9}$ | $4.77 \times 10^{-8}$ | lnc RNA-promoter-TSS | -321 | | | | | |
| AT1G61850 | protein coding gene | | 37.7 | $7.18 \times 10^{-24}$ | $3.08 \times 10^{-22}$ | mRNA-exon | 2200 | | | | | |
| AT2G27820 | protein coding gene | *PD1* | 37.6 | $1.68 \times 10^{-11}$ | $1.98 \times 10^{-10}$ | | -1620 | | | | | |
| AT5G23910 | protein coding gene | | 37.6 | $5.91 \times 10^{-23}$ | $2.32 \times 10^{-21}$ | mRNA-promoter-TSS | 499 | | | | | |
| AT4G38050 | protein coding gene | | 37.5 | $7.97 \times 10^{-20}$ | $2.37 \times 10^{-18}$ | mRNA-intron | 1263 | | | | | |
| AT1G34760 | protein coding gene | *GRF11* | 37.5 | $9.09 \times 10^{-30}$ | $6.68 \times 10^{-28}$ | mRNA-exon | 1397 | | | | | |
| AT5G49030 | protein coding gene | *OVA2* | 37.4 | $2.37 \times 10^{-6}$ | $1.37 \times 10^{-5}$ | mRNA-intron | 2311 | | | | | |
| AT5G36950 | protein coding gene | *DEG1* | 37.4 | $1.37 \times 10^{-13}$ | $2.09 \times 10^{-12}$ | mRNA-intron | 882 | | | | | |
| AT1G04647 | anti sense lncRNA | | 37.3 | $3.21 \times 10^{-32}$ | $2.85 \times 10^{-30}$ | lnc RNA-promoter-TSS | -688 | | | | | |
| AT1G51690 | protein coding gene | | 37.2 | $2.07 \times 10^{-9}$ | $1.82 \times 10^{-8}$ | mRNA-intron | 2677 | | | | | |
| AT5G50990 | protein coding gene | | 37.2 | $2.55 \times 10^{-17}$ | $5.97 \times 10^{-16}$ | | -1368 | | | | | |
| AT3G04100 | protein coding gene | *AGL57* | 37.1 | $9.89 \times 10^{-26}$ | $5.04 \times 10^{-24}$ | lnc RNA-TTS | 252 | | | | | |
| AT2G20910 | pseudogene | | 37.0 | $2.93 \times 10^{-15}$ | $5.49 \times 10^{-14}$ | transcript-intron | 1041 | | | | | |
| AT2G20910 | pseudogene | | 37.0 | $2.93 \times 10^{-15}$ | $5.49 \times 10^{-14}$ | transcript-intron | 1041 | 25.5 | $1.30 \times 10^{-5}$ | $3.56 \times 10^{-4}$ | transcript-intron | 1041 |
| AT5G00920 | anti sense lncRNA | | 37.0 | $7.01 \times 10^{-16}$ | $1.41 \times 10^{-14}$ | lnc RNA-promoter-TSS | 9 | | | | | |
| AT2G18938 | protein coding gene | | 36.9 | $1.76 \times 10^{-14}$ | $2.99 \times 10^{-13}$ | mRNA-exon | 2035 | | | | | |
| AT1G63540 | protein coding gene | | 36.9 | $2.12 \times 10^{-23}$ | $8.60 \times 10^{-22}$ | mRNA-exon | 893 | | | | | |
| AT2G26350 | protein coding gene | *PEX1* | 36.8 | $1.67 \times 10^{-11}$ | $1.97 \times 10^{-10}$ | mRNA-exon | 1156 | | | | | |
| AT4G27080 | protein coding gene | *PDIL5-4* | 36.8 | $2.47 \times 10^{-18}$ | $6.32 \times 10^{-17}$ | mRNA-intron | 2579 | | | | | |
| AT1G54000 | protein coding gene | *GLL22* | 36.8 | $1.71 \times 10^{-18}$ | $4.46 \times 10^{-17}$ | mRNA-promoter-TSS | -631 | | | | | |
| AT3G13225 | protein coding gene | | 36.8 | $1.66 \times 10^{-6}$ | $9.86 \times 10^{-6}$ | mRNA-exon | 1004 | | | | | |
| AT5G40450 | protein coding gene | | 36.8 | $2.45 \times 10^{-29}$ | $1.74 \times 10^{-27}$ | mRNA-exon | 2400 | | | | | |
| AT2G14570 | transposable element gene | | 36.8 | $8.88 \times 10^{-11}$ | $9.53 \times 10^{-10}$ | mRNA-promoter-TSS | -834 | | | | | |
| AT4G30110 | protein coding gene | *HMA2* | 36.7 | $3.00 \times 10^{-16}$ | $6.24 \times 10^{-15}$ | mRNA-exon | 3336 | | | | | |
| AT5G03170 | protein coding gene | *FLA11* | 36.5 | $4.30 \times 10^{-13}$ | $6.14 \times 10^{-12}$ | | -1194 | | | | | |
| AT1G06390 | protein coding gene | *GSK1* | 36.5 | $7.05 \times 10^{-6}$ | $3.76 \times 10^{-5}$ | mRNA-intron | 2260 | | | | | |
| AT1G65740 | protein coding gene | *UCL1* | 36.4 | $1.37 \times 10^{-10}$ | $1.43 \times 10^{-9}$ | mRNA-promoter-TSS | -640 | | | | | |
| AT1G77095 | transposable element gene | | 36.4 | $4.70 \times 10^{-27}$ | $2.72 \times 10^{-25}$ | lnc RNA-TTS | 616 | | | | | |
| AT3G58100 | protein coding gene | | 36.4 | $3.74 \times 10^{-13}$ | $5.39 \times 10^{-12}$ | | -1012 | | | | | |
| AT1G55250 | protein coding gene | *HUB2* | 36.3 | $2.80 \times 10^{-10}$ | $2.79 \times 10^{-9}$ | mRNA-exon | 1137 | | | | | |
| AT4G38005 | protein coding gene | | 36.3 | $3.98 \times 10^{-18}$ | $9.94 \times 10^{-17}$ | mRNA-promoter-TSS | -947 | | | | | |

| | | | | | | | |
|---|---|---|---|---|---|---|---|
| AT4G28230 | protein coding gene | | 36.3 | $2.53 \times 10^{-10}$ | $2.53 \times 10^{-9}$ | mRNA-TTS | 1656 |
| AT2G46550 | protein coding gene | | 36.2 | $1.47 \times 10^{-10}$ | $1.52 \times 10^{-9}$ | | -3624 |
| AT5G49430 | protein coding gene | | 36.2 | $1.04 \times 10^{-14}$ | $1.82 \times 10^{-13}$ | mRNA-intron | 4203 |
| AT5G20030 | protein coding gene | | 36.2 | $3.25 \times 10^{-11}$ | $3.71 \times 10^{-10}$ | | -2149 |
| AT1G10419 | pseudogene | | 36.2 | $7.38 \times 10^{-20}$ | $2.20 \times 10^{-18}$ | transcript-TTS | 1020 |
| AT5G03535 | lncRNA | | 36.2 | $5.86 \times 10^{-9}$ | $4.85 \times 10^{-8}$ | | -2514 |
| AT4G15050 | protein coding gene | | 36.2 | $1.16 \times 10^{-15}$ | $2.26 \times 10^{-14}$ | | -1206 |
| AT4G25610 | protein coding gene | | 36.2 | $2.87 \times 10^{-8}$ | $2.19 \times 10^{-7}$ | mRNA-TTS | 2987 |
| AT2G35060 | protein coding gene | *KUP11* | 36.1 | $6.17 \times 10^{-14}$ | $9.80 \times 10^{-13}$ | mRNA-TTS | 4382 |
| AT3G29290 | protein coding gene | *emb276* | 36.1 | $8.73 \times 10^{-18}$ | $2.12 \times 10^{-16}$ | mRNA-TTS | 2640 |
| AT4G15233 | protein coding gene | *ABCG42* | 36.1 | $5.47 \times 10^{-21}$ | $1.84 \times 10^{-19}$ | mRNA-exon | 2862 |
| AT5G35420 | transposable element gene | | 36.0 | $9.63 \times 10^{-32}$ | $8.20 \times 10^{-30}$ | Intergenic | -1714 |
| AT3G50050 | protein coding gene | | 36.0 | $4.15 \times 10^{-11}$ | $4.67 \times 10^{-10}$ | mRNA-exon | 3009 |
| AT3G06930 | protein coding gene | *PRMT4B* | 36.0 | $1.73 \times 10^{-8}$ | $1.36 \times 10^{-7}$ | mRNA-intron | 1436 |
| AT5G19360 | protein coding gene | *CPK34* | 36.0 | $2.22 \times 10^{-19}$ | $6.30 \times 10^{-18}$ | mRNA-TTS | 1803 |
| AT4G36197 | pre tRNA | | 36.0 | $2.08 \times 10^{-15}$ | $3.94 \times 10^{-14}$ | mRNA-promoter-TSS | -801 |
| AT1G28005 | protein coding gene | | 36.0 | $3.20 \times 10^{-37}$ | $4.36 \times 10^{-35}$ | mRNA-exon | 224 |
| AT5G31804 | transposable element gene | | 35.9 | $2.51 \times 10^{-19}$ | $7.11 \times 10^{-18}$ | Intergenic | -5287 |
| AT4G08705 | anti sense lncRNA | | 35.9 | $7.42 \times 10^{-13}$ | $1.03 \times 10^{-11}$ | | -1844 |
| AT1G42160 | transposable element gene | | 35.9 | $4.38 \times 10^{-9}$ | $3.70 \times 10^{-8}$ | Intergenic | -3929 |
| AT1G04210 | protein coding gene | | 35.8 | $1.60 \times 10^{-14}$ | $2.72 \times 10^{-13}$ | | -1824 |
| AT1G06630 | protein coding gene | | 35.7 | $7.10 \times 10^{-8}$ | $5.07 \times 10^{-7}$ | mRNA-exon | 1112 |
| AT1G06210 | protein coding gene | | 35.7 | $6.37 \times 10^{-8}$ | $4.57 \times 10^{-7}$ | mRNA-promoter-TSS | 748 |
| AT1G72440 | protein coding gene | *EDA25* | 35.6 | $5.28 \times 10^{-19}$ | $1.44 \times 10^{-17}$ | mRNA-exon | 3084 |
| AT4G08705 | anti sense lncRNA | | 35.6 | $7.43 \times 10^{-9}$ | $6.08 \times 10^{-8}$ | | -2544 |
| AT2G34730 | protein coding gene | | 35.5 | $2.94 \times 10^{-8}$ | $2.23 \times 10^{-7}$ | mRNA-promoter-TSS | -406 |
| AT3G05270 | protein coding gene | | 35.5 | $1.15 \times 10^{-15}$ | $2.25 \times 10^{-14}$ | mRNA-exon | 2130 |
| AT5G00485 | novel transcribed region | | 35.5 | $4.93 \times 10^{-21}$ | $1.66 \times 10^{-19}$ | transcript-promoter-TSS | -264 |
| AT1G36570 | transposable element gene | | 35.5 | $5.56 \times 10^{-14}$ | $8.93 \times 10^{-13}$ | Intergenic | 6522 |
| AT1G76630 | protein coding gene | | 35.5 | $3.21 \times 10^{-50}$ | $7.61 \times 10^{-48}$ | | -1327 |
| AT2G39800 | protein coding gene | *P5CS1* | 35.5 | $5.63 \times 10^{-8}$ | $4.08 \times 10^{-7}$ | mRNA-exon | 3108 |
| AT4G10590 | protein coding gene | *UBP1* | 35.5 | $3.04 \times 10^{-8}$ | $2.30 \times 10^{-7}$ | mRNA-exon | 1697 |
| AT2G17630 | protein coding gene | | 35.4 | $4.35 \times 10^{-13}$ | $6.22 \times 10^{-12}$ | mRNA-TTS | 1903 |
| AT5G43950 | protein coding gene | | 35.4 | $9.90 \times 10^{-6}$ | $5.13 \times 10^{-5}$ | | -1383 |
| AT4G26660 | protein coding gene | | 35.4 | $2.42 \times 10^{-12}$ | $3.18 \times 10^{-11}$ | | -1248 |
| AT2G45460 | protein coding gene | | 35.4 | $9.28 \times 10^{-8}$ | $6.55 \times 10^{-7}$ | mRNA-intron | 1939 |
| AT3G05575 | anti sense lncRNA | | 35.3 | $1.53 \times 10^{-24}$ | $7.09 \times 10^{-23}$ | lnc RNA-promoter-TSS | 97 |

| | | | | | | | |
|---|---|---|---|---|---|---|---|
| AT1G13620 | protein coding gene | *RGF2* | 35.3 | $1.40 \times 10^{-11}$ | $1.66 \times 10^{-10}$ | mRNA-TTS | 1332 |
| AT2G29100 | protein coding gene | | 35.3 | $3.02 \times 10^{-9}$ | $2.59 \times 10^{-8}$ | mRNA-exon | 1361 |
| AT1G22970 | protein coding gene | | 35.3 | $7.66 \times 10^{-15}$ | $1.36 \times 10^{-13}$ | mRNA-promoter-TSS | -716 |
| AT5G12230 | protein coding gene | *MED19A* | 35.3 | $1.28 \times 10^{-16}$ | $2.78 \times 10^{-15}$ | mRNA-exon | 2034 |
| AT2G11810 | protein coding gene | *MGDC* | 35.2 | $1.72 \times 10^{-5}$ | $8.55 \times 10^{-5}$ | mRNA-TTS | 3084 |
| AT3G07780 | protein coding gene | | 35.2 | $3.95 \times 10^{-18}$ | $9.87 \times 10^{-17}$ | | -2203 |
| AT1G12360 | protein coding gene | *KEU* | 35.1 | $2.45 \times 10^{-10}$ | $2.46 \times 10^{-9}$ | mRNA-exon | 1287 |
| AT4G29730 | protein coding gene | *NFC5* | 35.1 | $1.25 \times 10^{-8}$ | $9.96 \times 10^{-8}$ | mRNA-intron | 1628 |
| AT2G24420 | protein coding gene | | 35.1 | $1.34 \times 10^{-25}$ | $6.73 \times 10^{-24}$ | mRNA-intron | 1193 |
| AT1G31935 | other RNA | | 34.9 | $8.84 \times 10^{-15}$ | $1.56 \times 10^{-13}$ | | -1331 |
| AT4G00695 | protein coding gene | | 34.9 | $7.72 \times 10^{-13}$ | $1.07 \times 10^{-11}$ | mRNA-promoter-TSS | -841 |
| AT4G30330 | protein coding gene | | 34.9 | $1.31 \times 10^{-22}$ | $5.04 \times 10^{-21}$ | mRNA-promoter-TSS | -602 |
| AT5G19485 | protein coding gene | | 34.9 | $4.86 \times 10^{-21}$ | $1.64 \times 10^{-19}$ | mRNA-intron | 1583 |
| AT1G51970 | protein coding gene | | 34.9 | $9.64 \times 10^{-6}$ | $5.02 \times 10^{-5}$ | Intergenic | -4335 |
| AT1G54923 | transposable element gene | | 34.8 | $1.39 \times 10^{-6}$ | $8.30 \times 10^{-6}$ | mRNA-exon | 2517 |
| AT1G77460 | protein coding gene | | 34.8 | $1.46 \times 10^{-6}$ | $8.69 \times 10^{-6}$ | mRNA-exon | 1909 |
| AT5G30584 | transposable element gene | | 34.8 | $5.72 \times 10^{-6}$ | $3.11 \times 10^{-5}$ | mRNA-promoter-TSS | -513 |
| AT4G05815 | anti sense lncRNA | | 34.7 | $1.33 \times 10^{-15}$ | $2.59 \times 10^{-14}$ | mRNA-TTS | 293 |
| AT5G12360 | protein coding gene | | 34.7 | $1.83 \times 10^{-9}$ | $1.63 \times 10^{-8}$ | | -2709 |
| AT4G38120 | protein coding gene | | 34.6 | $4.99 \times 10^{-11}$ | $5.54 \times 10^{-10}$ | mRNA-intron | 2556 |
| AT3G20730 | protein coding gene | | 34.5 | $1.19 \times 10^{-8}$ | $9.49 \times 10^{-8}$ | mRNA-TTS | 555 |
| AT1G04407 | lncRNA | | 34.5 | $2.97 \times 10^{-15}$ | $5.54 \times 10^{-14}$ | mRNA-exon | 1889 |
| AT5G15270 | protein coding gene | | 34.5 | $1.77 \times 10^{-17}$ | $4.19 \times 10^{-16}$ | mRNA-exon | 1536 |
| AT1G49170 | protein coding gene | | 34.5 | $1.15 \times 10^{-14}$ | $1.99 \times 10^{-13}$ | mRNA-exon | 932 |
| AT4G14300 | protein coding gene | | 34.5 | $1.79 \times 10^{-8}$ | $1.40 \times 10^{-7}$ | mRNA-exon | 1249 |
| AT4G28880 | protein coding gene | *ckl3* | 34.4 | $6.84 \times 10^{-12}$ | $8.46 \times 10^{-11}$ | mRNA-exon | 2312 |
| AT3G10330 | protein coding gene | | 34.4 | $1.58 \times 10^{-12}$ | $2.11 \times 10^{-11}$ | mRNA-intron | 1739 |
| AT4G39640 | protein coding gene | *GGT1* | 34.4 | $1.24 \times 10^{-19}$ | $3.64 \times 10^{-18}$ | mRNA-exon | 1629 |
| AT2G05670 | pseudogene | | 34.4 | $5.75 \times 10^{-9}$ | $4.78 \times 10^{-8}$ | transcript-exon | 727 |
| AT5G35420 | transposable element gene | | 34.3 | $4.52 \times 10^{-46}$ | $9.50 \times 10^{-44}$ | Intergenic | -2114 |
| AT1G77010 | protein coding gene | | 34.3 | $6.59 \times 10^{-15}$ | $1.18 \times 10^{-13}$ | mRNA-TTS | 1841 |
| AT5G00600 | novel transcribed region | | 34.2 | $1.49 \times 10^{-9}$ | $1.34 \times 10^{-8}$ | | -1309 |
| AT2G31300 | protein coding gene | | 34.2 | $3.31 \times 10^{-11}$ | $3.78 \times 10^{-10}$ | mRNA-intron | 1648 |
| AT3G06665 | pre tRNA | | 34.1 | $2.06 \times 10^{-14}$ | $3.46 \times 10^{-13}$ | | -2224 |
| AT3G46970 | protein coding gene | *PHS2* | 34.0 | $2.51 \times 10^{-16}$ | $5.28 \times 10^{-15}$ | mRNA-exon | 3382 |
| AT3G62930 | protein coding gene | | 34.0 | $4.57 \times 10^{-16}$ | $9.31 \times 10^{-15}$ | | -1224 |
| AT2G18710 | protein coding gene | *SCY1* | 33.9 | $1.08 \times 10^{-16}$ | $2.36 \times 10^{-15}$ | mRNA-exon | 1134 |

| | | | | | | | |
|---|---|---|---|---|---|---|---|
| AT1G75120 | protein coding gene | *RRA1* | 33.9 | $1.90 \times 10^{-16}$ | $4.05 \times 10^{-15}$ | mRNA-exon | 1294 |
| AT5G37560 | protein coding gene | | 33.9 | $7.94 \times 10^{-34}$ | $8.19 \times 10^{-32}$ | mRNA-exon | 165 |
| AT1G09515 | novel transcribed region | | 33.8 | $2.45 \times 10^{-13}$ | $3.62 \times 10^{-12}$ | transcript-intron | 585 |
| AT5G55100 | protein coding gene | | 33.8 | $4.76 \times 10^{-10}$ | $4.57 \times 10^{-9}$ | mRNA-exon | 1902 |
| AT4G16130 | protein coding gene | *ARA1* | 33.8 | $1.75 \times 10^{-6}$ | $1.03 \times 10^{-5}$ | mRNA-intron | 3174 |
| AT1G42980 | protein coding gene | | 33.7 | $2.59 \times 10^{-20}$ | $8.06 \times 10^{-19}$ | mRNA-intron | 1365 |
| AT3G04690 | protein coding gene | *ANX1* | 33.7 | $3.42 \times 10^{-9}$ | $2.91 \times 10^{-8}$ | mRNA-promoter-TSS | -306 |
| AT4G27040 | protein coding gene | *VPS22* | 33.6 | $4.32 \times 10^{-61}$ | $1.75 \times 10^{-58}$ | | -1112 |
| AT4G34555 | protein coding gene | | 33.6 | $1.39 \times 10^{-22}$ | $5.32 \times 10^{-21}$ | mRNA-TTS | 732 |
| AT2G32580 | protein coding gene | | 33.5 | $9.14 \times 10^{-23}$ | $3.53 \times 10^{-21}$ | mRNA-promoter-TSS | -774 |
| AT3G15640 | protein coding gene | | 33.5 | $1.10 \times 10^{-5}$ | $5.67 \times 10^{-5}$ | mRNA-intron | 1141 |
| AT1G21690 | protein coding gene | *EMB1968* | 33.5 | $4.06 \times 10^{-14}$ | $6.62 \times 10^{-13}$ | mRNA-exon | 1207 |
| AT3G20020 | protein coding gene | | 33.5 | $6.74 \times 10^{-8}$ | $4.82 \times 10^{-7}$ | mRNA-intron | 3748 |
| AT3G25110 | protein coding gene | *FaTA* | 33.4 | $1.42 \times 10^{-13}$ | $2.17 \times 10^{-12}$ | mRNA-TTS | 1374 |
| AT4G21192 | protein coding gene | | 33.4 | $4.00 \times 10^{-10}$ | $3.88 \times 10^{-9}$ | mRNA-intron | 846 |
| AT5G20030 | protein coding gene | | 33.4 | $1.61 \times 10^{-10}$ | $1.66 \times 10^{-9}$ | | -1449 |
| AT1G09900 | protein coding gene | | 33.4 | $2.67 \times 10^{-6}$ | $1.53 \times 10^{-5}$ | mRNA-TTS | 918 |
| AT5G43890 | protein coding gene | *YUC5* | 33.4 | $1.87 \times 10^{-5}$ | $9.27 \times 10^{-5}$ | | -8076 |
| AT1G07663 | anti sense lncRNA | | 33.4 | $2.03 \times 10^{-9}$ | $1.79 \times 10^{-8}$ | lnc RNA-promoter-TSS | -694 |
| AT2G16940 | protein coding gene | | 33.4 | $1.09 \times 10^{-7}$ | $7.62 \times 10^{-7}$ | mRNA-exon | 150 |
| AT3G48080 | protein coding gene | | 33.3 | $6.38 \times 10^{-10}$ | $6.02 \times 10^{-9}$ | mRNA-promoter-TSS | -861 |
| AT1G65430 | protein coding gene | *ARI8* | 33.3 | $2.47 \times 10^{-7}$ | $1.65 \times 10^{-6}$ | mRNA-intron | 887 |
| AT5G41800 | protein coding gene | | 33.3 | $1.32 \times 10^{-20}$ | $4.27 \times 10^{-19}$ | | -1912 |
| AT1G77460 | protein coding gene | | 33.3 | $9.71 \times 10^{-9}$ | $7.83 \times 10^{-8}$ | mRNA-exon | 1809 |
| AT3G08450 | lncRNA | | 33.3 | $5.34 \times 10^{-10}$ | $5.09 \times 10^{-9}$ | mRNA-exon | 2572 |
| AT1G75850 | protein coding gene | *VPS35B* | 33.3 | $2.14 \times 10^{-4}$ | $8.50 \times 10^{-4}$ | mRNA-intron | 496 |
| AT5G24660 | protein coding gene | *LSU2* | 33.3 | $1.57 \times 10^{-20}$ | $5.04 \times 10^{-19}$ | Intergenic | -1550 |
| AT3G09915 | pseudogene | | 33.3 | $1.97 \times 10^{-14}$ | $3.32 \times 10^{-13}$ | | -1265 |
| AT2G14115 | anti sense lncRNA | | 33.2 | $3.73 \times 10^{-7}$ | $2.43 \times 10^{-6}$ | mRNA-intron | 2155 |
| AT5G16680 | protein coding gene | | 33.2 | $3.99 \times 10^{-7}$ | $2.60 \times 10^{-6}$ | mRNA-exon | 803 |
| AT2G21940 | protein coding gene | *SK1* | 33.2 | $2.10 \times 10^{-13}$ | $3.13 \times 10^{-12}$ | mRNA-intron | 351 |
| AT3G07980 | protein coding gene | *MAPKKK6* | 33.2 | $2.16 \times 10^{-6}$ | $1.26 \times 10^{-5}$ | mRNA-exon | 807 |
| AT1G43700 | protein coding gene | *VIP1* | 33.1 | $2.07 \times 10^{-12}$ | $2.73 \times 10^{-11}$ | mRNA-TTS | 1720 |
| AT2G31270 | protein coding gene | *CDT1A* | 33.1 | $1.64 \times 10^{-12}$ | $2.19 \times 10^{-11}$ | mRNA-exon | 1789 |
| AT2G31590 | protein coding gene | | 33.1 | $7.51 \times 10^{-21}$ | $2.49 \times 10^{-19}$ | mRNA-TTS | 182 |
| AT3G19960 | protein coding gene | *ATM1* | 33.1 | $1.00 \times 10^{-4}$ | $4.30 \times 10^{-4}$ | mRNA-intron | 3331 |
| AT3G14120 | protein coding gene | | 33.1 | $3.63 \times 10^{-11}$ | $4.11 \times 10^{-10}$ | mRNA-promoter-TSS | 838 |

| | | | | | | | |
|---|---|---|---|---|---|---|---|
| AT1G65850 | protein coding gene | | 33.0 | $1.88 \times 10^{-28}$ | $1.21 \times 10^{-26}$ | mRNA-exon | 3017 |
| AT4G09335 | lncRNA | | 32.9 | $2.84 \times 10^{-16}$ | $5.91 \times 10^{-15}$ | | -1107 |
| AT1G05030 | protein coding gene | | 32.9 | $5.82 \times 10^{-6}$ | $3.16 \times 10^{-5}$ | mRNA-intron | 1366 |
| AT3G24880 | protein coding gene | | 32.8 | $5.19 \times 10^{-7}$ | $3.32 \times 10^{-6}$ | mRNA-exon | 2486 |
| AT5G42242 | protein coding gene | *LCR57* | 32.8 | $3.94 \times 10^{-23}$ | $1.57 \times 10^{-21}$ | mRNA-TTS | 1658 |
| AT2G33160 | protein coding gene | | 32.8 | $1.98 \times 10^{-10}$ | $2.02 \times 10^{-9}$ | mRNA-promoter-TSS | -653 |
| AT4G38932 | anti sense lncRNA | | 32.8 | $1.28 \times 10^{-36}$ | $1.61 \times 10^{-34}$ | mRNA-promoter-TSS | 546 |
| AT3G49050 | protein coding gene | | 32.8 | $2.77 \times 10^{-17}$ | $6.45 \times 10^{-16}$ | mRNA-exon | 1144 |
| AT4G27170 | protein coding gene | *SESA4* | 32.8 | $9.88 \times 10^{-9}$ | $7.95 \times 10^{-8}$ | mRNA-exon | 2455 |
| AT1G05030 | protein coding gene | | 32.7 | $1.32 \times 10^{-6}$ | $7.94 \times 10^{-6}$ | mRNA-exon | 1666 |
| AT1G73220 | protein coding gene | *Oct-01* | 32.7 | $4.70 \times 10^{-25}$ | $2.28 \times 10^{-23}$ | mRNA-TTS | 1161 |
| AT5G16280 | protein coding gene | | 32.7 | $3.67 \times 10^{-10}$ | $3.58 \times 10^{-9}$ | mRNA-intron | 3943 |
| AT4G01960 | protein coding gene | | 32.7 | $5.65 \times 10^{-20}$ | $1.72 \times 10^{-18}$ | mRNA-exon | 1087 |
| AT3G14172 | protein coding gene | | 32.7 | $6.83 \times 10^{-8}$ | $4.88 \times 10^{-7}$ | mRNA-intron | 1878 |
| AT1G67400 | protein coding gene | | 32.6 | $6.43 \times 10^{-28}$ | $4.00 \times 10^{-26}$ | mRNA-exon | 2540 |
| AT1G04437 | anti sense lncRNA | | 32.5 | $1.10 \times 10^{-6}$ | $6.74 \times 10^{-6}$ | lnc RNA-promoter-TSS | -522 |
| AT5G28150 | protein coding gene | | 32.5 | $5.12 \times 10^{-7}$ | $3.28 \times 10^{-6}$ | mRNA-exon | 942 |
| AT4G25719 | anti sense lncRNA | | 32.5 | $7.59 \times 10^{-8}$ | $5.40 \times 10^{-7}$ | lnc RNA-TTS | 584 |
| AT5G13480 | protein coding gene | *FY* | 32.5 | $4.51 \times 10^{-11}$ | $5.03 \times 10^{-10}$ | mRNA-exon | 1760 |
| AT1G16130 | protein coding gene | *WAKL2* | 32.4 | $5.73 \times 10^{-21}$ | $1.92 \times 10^{-19}$ | mRNA-exon | 1658 |
| AT1G26225 | protein coding gene | | 32.4 | $1.34 \times 10^{-12}$ | $1.81 \times 10^{-11}$ | lnc RNA-promoter-TSS | 117 |
| AT3G04810 | protein coding gene | *NEK2* | 32.4 | $9.35 \times 10^{-15}$ | $1.65 \times 10^{-13}$ | mRNA-intron | 1915 |
| AT3G05760 | protein coding gene | | 32.4 | $1.06 \times 10^{-5}$ | $5.45 \times 10^{-5}$ | mRNA-promoter-TSS | -946 |
| AT1G04170 | protein coding gene | *EIF2 GAMMA* | 32.3 | $1.41 \times 10^{-7}$ | $9.72 \times 10^{-7}$ | | -3031 |
| AT2G39080 | protein coding gene | *EMB2799* | 32.3 | $2.25 \times 10^{-18}$ | $5.76 \times 10^{-17}$ | mRNA-promoter-TSS | -429 |
| AT4G06990 | anti sense lncRNA | | 32.3 | $2.11 \times 10^{-7}$ | $1.42 \times 10^{-6}$ | lnc RNA-promoter-TSS | 26 |
| AT5G37980 | protein coding gene | | 32.3 | $3.59 \times 10^{-19}$ | $9.99 \times 10^{-18}$ | mRNA-exon | 1362 |
| AT5G02875 | anti sense lncRNA | | 32.3 | $3.97 \times 10^{-8}$ | $2.95 \times 10^{-7}$ | lnc RNA-promoter-TSS | -458 |
| AT2G08135 | antisense RNA | | 32.2 | $8.27 \times 10^{-10}$ | $7.70 \times 10^{-9}$ | antisense RNA-promoter-TSS | -791 |
| AT5G34867 | transposable element gene | | 32.2 | $2.52 \times 10^{-6}$ | $1.45 \times 10^{-5}$ | mRNA-exon | 3928 |
| AT4G35790 | protein coding gene | *PLDDELTA* | 32.2 | $6.20 \times 10^{-8}$ | $4.45 \times 10^{-7}$ | mRNA-exon | 1519 |
| AT4G12420 | protein coding gene | *SKU5* | 32.2 | $2.46 \times 10^{-6}$ | $1.42 \times 10^{-5}$ | mRNA-intron | 1666 |
| AT2G19710 | protein coding gene | | 32.2 | $4.26 \times 10^{-15}$ | $7.82 \times 10^{-14}$ | mRNA-intron | 1121 |
| AT1G34250 | transposable element gene | | 32.1 | $1.18 \times 10^{-35}$ | $1.38 \times 10^{-33}$ | Intergenic | 4183 |
| AT4G16267 | pseudogene | | 32.1 | $6.30 \times 10^{-16}$ | $1.27 \times 10^{-14}$ | transcript-promoter-TSS | -903 |
| AT4G30720 | protein coding gene | *PDE327* | 32.1 | $1.95 \times 10^{-9}$ | $1.73 \times 10^{-8}$ | mRNA-exon | 2577 |
| AT1G43980 | protein coding gene | | 32.1 | $3.49 \times 10^{-9}$ | $2.97 \times 10^{-8}$ | mRNA-exon | 1312 |

| | | | | | | | |
|---|---|---|---|---|---|---|---|
| AT4G27080 | protein coding gene | *PDIL5-4* | 32.1 | $3.30 \times 10^{-24}$ | $1.46 \times 10^{-22}$ | mRNA-intron | 1279 |
| AT1G74260 | protein coding gene | *PUR4* | 32.1 | $3.80 \times 10^{-19}$ | $1.06 \times 10^{-17}$ | mRNA-exon | 1311 |
| AT5G55280 | protein coding gene | *FTSZ1-1* | 32.0 | $7.43 \times 10^{-11}$ | $8.02 \times 10^{-10}$ | mRNA-exon | 1098 |
| AT4G20730 | transposable element gene | | 32.0 | $1.64 \times 10^{-32}$ | $1.47 \times 10^{-30}$ | mRNA-exon | 1844 |
| AT5G35700 | protein coding gene | *FIM5* | 32.0 | $2.50 \times 10^{-5}$ | $1.21 \times 10^{-4}$ | mRNA-intron | 2025 |
| AT5G16060 | protein coding gene | | 31.9 | $5.43 \times 10^{-19}$ | $1.47 \times 10^{-17}$ | mRNA-TTS | 1771 |
| AT1G67900 | protein coding gene | | 31.9 | $6.19 \times 10^{-9}$ | $5.10 \times 10^{-8}$ | mRNA-exon | 2116 |
| AT2G01350 | protein coding gene | *QPT* | 31.8 | $3.47 \times 10^{-11}$ | $3.94 \times 10^{-10}$ | mRNA-TTS | 1596 |
| AT5G30545 | transposable element gene | | 31.8 | $2.54 \times 10^{-6}$ | $1.46 \times 10^{-5}$ | mRNA-exon | 1069 |
| AT5G63460 | protein coding gene | | 31.8 | $8.63 \times 10^{-7}$ | $5.34 \times 10^{-6}$ | mRNA-intron | 1357 |
| AT1G05083 | anti sense lncRNA | | 31.8 | $9.06 \times 10^{-9}$ | $7.33 \times 10^{-8}$ | lnc RNA-promoter-TSS | -832 |
| AT5G35670 | protein coding gene | *iqd33* | 31.7 | $2.12 \times 10^{-8}$ | $1.63 \times 10^{-7}$ | mRNA-exon | 615 |
| AT1G09980 | protein coding gene | | 31.7 | $2.31 \times 10^{-16}$ | $4.89 \times 10^{-15}$ | mRNA-exon | 3563 |
| AT1G50490 | protein coding gene | *UBC2* | 31.6 | $4.08 \times 10^{-6}$ | $2.27 \times 10^{-5}$ | mRNA-TTS | 2206 |
| AT5G27750 | protein coding gene | | 31.6 | $3.19 \times 10^{-13}$ | $4.64 \times 10^{-12}$ | mRNA-exon | 814 |
| AT3G09840 | protein coding gene | *CDC48A* | 31.6 | $1.19 \times 10^{-13}$ | $1.84 \times 10^{-12}$ | | -1198 |
| AT2G24592 | lncRNA | | 31.6 | $4.06 \times 10^{-8}$ | $3.00 \times 10^{-7}$ | lnc RNA-promoter-TSS | 26 |
| AT1G27000 | protein coding gene | | 31.5 | $3.32 \times 10^{-17}$ | $7.65 \times 10^{-16}$ | mRNA-exon | 666 |
| AT1G28540 | protein coding gene | | 31.5 | $6.98 \times 10^{-10}$ | $6.55 \times 10^{-9}$ | mRNA-promoter-TSS | -966 |
| AT5G25940 | protein coding gene | | 31.5 | $3.25 \times 10^{-37}$ | $4.38 \times 10^{-35}$ | mRNA-promoter-TSS | -796 |
| AT3G08010 | protein coding gene | *ATAB2* | 31.4 | $1.51 \times 10^{-26}$ | $8.31 \times 10^{-25}$ | mRNA-TTS | 1781 |
| AT3G04670 | protein coding gene | *WRKY39* | 31.4 | $3.29 \times 10^{-6}$ | $1.85 \times 10^{-5}$ | mRNA-TTS | 1699 |
| AT1G23330 | protein coding gene | | 31.4 | $5.42 \times 10^{-24}$ | $2.36 \times 10^{-22}$ | mRNA-TTS | 3165 |
| AT3G56190 | protein coding gene | *ALPHA-SNAP2* | 31.4 | $1.43 \times 10^{-9}$ | $1.29 \times 10^{-8}$ | mRNA-intron | 1183 |
| AT2G09660 | anti sense lncRNA | | 31.4 | $4.45 \times 10^{-41}$ | $7.25 \times 10^{-39}$ | lnc RNA-promoter-TSS | -90 |
| AT1G49990 | protein coding gene | | 31.4 | $4.62 \times 10^{-9}$ | $3.90 \times 10^{-8}$ | mRNA-TTS | 1323 |
| AT1G10670 | protein coding gene | *ACLA-1* | 31.4 | $2.78 \times 10^{-9}$ | $2.41 \times 10^{-8}$ | | -1137 |
| AT4G31740 | protein coding gene | | 31.4 | $5.71 \times 10^{-14}$ | $9.15 \times 10^{-13}$ | mRNA-TTS | 873 |
| AT1G59520 | protein coding gene | | 31.3 | $1.36 \times 10^{-27}$ | $8.17 \times 10^{-26}$ | mRNA-promoter-TSS | -831 |
| AT2G42580 | protein coding gene | *TTL3* | 31.3 | $4.38 \times 10^{-22}$ | $1.63 \times 10^{-20}$ | mRNA-TTS | 2672 |
| AT1G70949 | protein coding gene | | 31.3 | $2.20 \times 10^{-16}$ | $4.66 \times 10^{-15}$ | | -1587 |
| AT4G38930 | protein coding gene | | 31.3 | $1.99 \times 10^{-6}$ | $1.16 \times 10^{-5}$ | mRNA-exon | 1218 |
| AT5G22910 | protein coding gene | *CHX9* | 31.3 | $3.12 \times 10^{-22}$ | $1.17 \times 10^{-20}$ | mRNA-exon | 1178 |
| AT3G24560 | protein coding gene | *RSY3* | 31.3 | $1.34 \times 10^{-26}$ | $7.39 \times 10^{-25}$ | mRNA-TTS | 1292 |
| AT4G12130 | protein coding gene | | 31.2 | $7.75 \times 10^{-23}$ | $3.01 \times 10^{-21}$ | mRNA-exon | 975 |
| AT5G27550 | protein coding gene | | 31.1 | $5.30 \times 10^{-8}$ | $3.86 \times 10^{-7}$ | | -1493 |
| AT4G33530 | protein coding gene | *KUP5* | 31.1 | $5.26 \times 10^{-8}$ | $3.83 \times 10^{-7}$ | mRNA-intron | 2833 |

| | | | | | | | |
|---|---|---|---|---|---|---|---|
| AT5G18410 | protein coding gene | *PIR121* | 31.1 | $2.92 \times 10^{-7}$ | $1.94 \times 10^{-6}$ | mRNA-promoter-TSS | -947 |
| AT2G03550 | protein coding gene | | 31.1 | $4.83 \times 10^{-45}$ | $9.33 \times 10^{-43}$ | mRNA-exon | 888 |
| AT3G51030 | protein coding gene | *TRX1* | 31.0 | $2.57 \times 10^{-14}$ | $4.29 \times 10^{-13}$ | mRNA-promoter-TSS | -642 |
| AT5G34863 | transposable element gene | | 31.0 | $2.02 \times 10^{-8}$ | $1.57 \times 10^{-7}$ | mRNA-exon | 1192 |
| AT1G66150 | protein coding gene | *TMK1* | 31.0 | $1.15 \times 10^{-10}$ | $1.22 \times 10^{-9}$ | mRNA-exon | 1313 |
| AT4G18810 | protein coding gene | | 31.0 | $1.49 \times 10^{-9}$ | $1.34 \times 10^{-8}$ | mRNA-exon | 1765 |
| AT3G08645 | anti sense lncRNA | | 31.0 | $7.71 \times 10^{-10}$ | $7.20 \times 10^{-9}$ | | -1734 |
| AT3G59300 | protein coding gene | | 31.0 | $4.33 \times 10^{-10}$ | $4.18 \times 10^{-9}$ | mRNA-exon | 471 |
| AT4G38600 | protein coding gene | *KAK* | 30.9 | $3.22 \times 10^{-5}$ | $1.53 \times 10^{-4}$ | mRNA-intron | 1801 |
| AT1G24420 | protein coding gene | | 30.9 | $5.67 \times 10^{-8}$ | $4.10 \times 10^{-7}$ | mRNA-TTS | 747 |
| AT5G43950 | protein coding gene | | 30.9 | $1.29 \times 10^{-6}$ | $7.77 \times 10^{-6}$ | | -1183 |
| AT1G71830 | protein coding gene | *SERK1* | 30.8 | $5.70 \times 10^{-8}$ | $4.11 \times 10^{-7}$ | | -1907 |
| AT3G48560 | protein coding gene | *CSR1* | 30.8 | $3.61 \times 10^{-11}$ | $4.09 \times 10^{-10}$ | mRNA-exon | 1298 |
| AT1G09470 | protein coding gene | | 30.8 | $5.73 \times 10^{-8}$ | $4.13 \times 10^{-7}$ | mRNA-TTS | 1646 |
| AT2G44130 | protein coding gene | | 30.7 | $2.06 \times 10^{-18}$ | $5.31 \times 10^{-17}$ | mRNA-TTS | 2241 |
| AT3G28210 | protein coding gene | *PMZ* | 30.7 | $1.81 \times 10^{-10}$ | $1.86 \times 10^{-9}$ | mRNA-TTS | 1049 |
| AT3G02290 | protein coding gene | | 30.6 | $3.36 \times 10^{-14}$ | $5.53 \times 10^{-13}$ | | -1905 |
| AT1G23870 | protein coding gene | *TPS9* | 30.6 | $3.83 \times 10^{-12}$ | $4.91 \times 10^{-11}$ | mRNA-exon | 1372 |
| AT5G49920 | protein coding gene | | 30.6 | $2.24 \times 10^{-9}$ | $1.96 \times 10^{-8}$ | | -2108 |
| AT4G38230 | protein coding gene | *CPK26* | 30.6 | $4.50 \times 10^{-6}$ | $2.48 \times 10^{-5}$ | mRNA-exon | 1131 |
| AT4G05815 | anti sense lncRNA | | 30.6 | $7.62 \times 10^{-9}$ | $6.22 \times 10^{-8}$ | | -1207 |
| AT2G30945 | other RNA | | 30.5 | $3.12 \times 10^{-11}$ | $3.58 \times 10^{-10}$ | mRNA-TTS | 379 |
| AT5G13730 | protein coding gene | *SIG4* | 30.5 | $8.40 \times 10^{-36}$ | $1.01 \times 10^{-33}$ | mRNA-TTS | 1350 |
| AT5G65110 | protein coding gene | *ACX2* | 30.5 | $2.55 \times 10^{-13}$ | $3.76 \times 10^{-12}$ | mRNA-exon | 868 |
| AT4G23540 | protein coding gene | | 30.4 | $5.86 \times 10^{-15}$ | $1.06 \times 10^{-13}$ | mRNA-exon | 935 |
| AT5G49850 | protein coding gene | | 30.4 | $4.09 \times 10^{-45}$ | $7.99 \times 10^{-43}$ | mRNA-TTS | 1634 |
| AT3G44680 | protein coding gene | *HDA9* | 30.4 | $2.00 \times 10^{-6}$ | $1.17 \times 10^{-5}$ | | -1042 |
| AT5G42230 | protein coding gene | *scpl41* | 30.3 | $3.12 \times 10^{-6}$ | $1.77 \times 10^{-5}$ | mRNA-promoter-TSS | -341 |
| AT1G36395 | transposable element gene | | 30.3 | $2.90 \times 10^{-5}$ | $1.39 \times 10^{-4}$ | mRNA-TTS | 1371 |
| AT5G30628 | transposable element gene | | 30.3 | $3.17 \times 10^{-24}$ | $1.41 \times 10^{-22}$ | mRNA-promoter-TSS | -824 |
| AT4G35800 | protein coding gene | *NRPB1* | 30.2 | $5.37 \times 10^{-7}$ | $3.43 \times 10^{-6}$ | mRNA-exon | 1777 |
| AT2G14247 | protein coding gene | | 30.2 | $1.18 \times 10^{-36}$ | $1.49 \times 10^{-34}$ | lnc RNA-TTS | 369 |
| AT5G37130 | protein coding gene | | 30.2 | $7.77 \times 10^{-7}$ | $4.83 \times 10^{-6}$ | mRNA-exon | 3303 |
| AT4G04221 | anti sense lncRNA | | 30.2 | $1.89 \times 10^{-9}$ | $1.67 \times 10^{-8}$ | mRNA-exon | 723 |
| AT2G02090 | protein coding gene | *ETL1* | 30.2 | $2.10 \times 10^{-8}$ | $1.63 \times 10^{-7}$ | mRNA-exon | 1828 |
| AT2G29030 | pre tRNA | | 30.1 | $3.24 \times 10^{-7}$ | $2.14 \times 10^{-6}$ | | -1407 |
| AT2G05180 | protein coding gene | *CYP75A6* | 30.1 | $8.47 \times 10^{-10}$ | $7.86 \times 10^{-9}$ | mRNA-promoter-TSS | -439 |

| | | | | | | | |
|---|---|---|---|---|---|---|---|
| AT1G31485 | anti sense lncRNA | | 30.1 | $1.35 \times 10^{-19}$ | $3.93 \times 10^{-18}$ | mRNA-TTS | 607 |
| AT5G35140 | transposable element gene | | 30.0 | $1.50 \times 10^{-7}$ | $1.03 \times 10^{-6}$ | mRNA-exon | 343 |
| AT5G61780 | protein coding gene | *Tudor2* | 30.0 | $7.07 \times 10^{-5}$ | $3.13 \times 10^{-4}$ | mRNA-exon | 2848 |
| AT5G53500 | protein coding gene | | 29.9 | $2.77 \times 10^{-11}$ | $3.19 \times 10^{-10}$ | mRNA-exon | 2262 |
| AT5G12990 | protein coding gene | *CLE4* | 29.9 | $3.14 \times 10^{-23}$ | $1.26 \times 10^{-21}$ | | -1167 |
| AT1G20080 | protein coding gene | *SYTB* | 29.9 | $2.85 \times 10^{-9}$ | $2.46 \times 10^{-8}$ | mRNA-intron | 1124 |
| AT4G00910 | protein coding gene | | 29.8 | $1.33 \times 10^{-21}$ | $4.71 \times 10^{-20}$ | mRNA-exon | 890 |
| AT4G14160 | protein coding gene | | 29.8 | $7.10 \times 10^{-5}$ | $3.14 \times 10^{-4}$ | mRNA-intron | 1586 |
| AT3G06530 | protein coding gene | | 29.8 | $4.37 \times 10^{-7}$ | $2.83 \times 10^{-6}$ | mRNA-exon | 3555 |
| AT5G58810 | pseudogene | | 29.8 | $6.77 \times 10^{-16}$ | $1.36 \times 10^{-14}$ | transcript-exon | 1989 |
| AT1G50490 | protein coding gene | *UBC2* | 29.8 | $1.62 \times 10^{-9}$ | $1.44 \times 10^{-8}$ | | -3494 |
| AT4G19220 | protein coding gene | | 29.8 | $7.31 \times 10^{-8}$ | $5.21 \times 10^{-7}$ | mRNA-TTS | 2070 |
| AT4G18790 | protein coding gene | *NRAMP5* | 29.8 | $5.46 \times 10^{-11}$ | $6.02 \times 10^{-10}$ | mRNA-exon | 885 |
| AT4G32170 | protein coding gene | *CYP96A2* | 29.7 | $1.25 \times 10^{-10}$ | $1.31 \times 10^{-9}$ | | -1350 |
| AT1G09093 | novel transcribed region | | 29.6 | $2.69 \times 10^{-15}$ | $5.05 \times 10^{-14}$ | transcript-promoter-TSS | -693 |
| AT3G03340 | protein coding gene | *LUC7A* | 29.6 | $2.28 \times 10^{-26}$ | $1.22 \times 10^{-24}$ | | -2154 |
| AT3G58730 | protein coding gene | | 29.6 | $8.77 \times 10^{-15}$ | $1.55 \times 10^{-13}$ | mRNA-exon | 1002 |
| AT1G35300 | transposable element gene | | 29.6 | $1.98 \times 10^{-5}$ | $9.76 \times 10^{-5}$ | mRNA-exon | 1375 |
| AT5G26670 | protein coding gene | | 29.5 | $9.89 \times 10^{-10}$ | $9.08 \times 10^{-9}$ | mRNA-TTS | 2333 |
| AT3G13670 | protein coding gene | | 29.5 | $1.17 \times 10^{-5}$ | $6.01 \times 10^{-5}$ | mRNA-intron | 1798 |
| AT2G29710 | protein coding gene | | 29.5 | $1.51 \times 10^{-30}$ | $1.19 \times 10^{-28}$ | mRNA-TTS | 1478 |
| AT5G26670 | protein coding gene | | 29.4 | $9.29 \times 10^{-14}$ | $1.44 \times 10^{-12}$ | mRNA-exon | 1833 |
| AT5G57330 | protein coding gene | | 29.4 | $4.30 \times 10^{-6}$ | $2.38 \times 10^{-5}$ | mRNA-exon | 1565 |
| AT1G67140 | protein coding gene | *SWEETIE* | 29.4 | $2.00 \times 10^{-7}$ | $1.35 \times 10^{-6}$ | mRNA-intron | 2772 |
| AT1G12750 | protein coding gene | *RBL6* | 29.4 | $5.57 \times 10^{-13}$ | $7.89 \times 10^{-12}$ | mRNA-intron | 794 |
| AT3G06930 | protein coding gene | *PRMT4B* | 29.4 | $3.71 \times 10^{-10}$ | $3.62 \times 10^{-9}$ | mRNA-exon | 1236 |
| AT2G46060 | protein coding gene | | 29.4 | $1.19 \times 10^{-10}$ | $1.25 \times 10^{-9}$ | | -1139 |
| AT1G21900 | protein coding gene | *p24delta5* | 29.3 | $1.70 \times 10^{-16}$ | $3.64 \times 10^{-15}$ | mRNA-exon | 1408 |
| AT1G18840 | protein coding gene | *IQD3* | 29.3 | $5.47 \times 10^{-12}$ | $6.87 \times 10^{-11}$ | mRNA-exon | 1859 |
| AT2G04700 | protein coding gene | | 29.3 | $2.90 \times 10^{-13}$ | $4.25 \times 10^{-12}$ | mRNA-promoter-TSS | -648 |
| AT2G38050 | protein coding gene | *DET2* | 29.2 | $5.10 \times 10^{-10}$ | $4.88 \times 10^{-9}$ | mRNA-exon | 2570 |
| AT5G07790 | protein coding gene | | 29.2 | $7.69 \times 10^{-7}$ | $4.79 \times 10^{-6}$ | mRNA-exon | 2028 |
| AT2G18938 | protein coding gene | | 29.2 | $6.05 \times 10^{-13}$ | $8.54 \times 10^{-12}$ | mRNA-exon | 1935 |
| AT1G54970 | protein coding gene | *PRP1* | 29.2 | $9.77 \times 10^{-13}$ | $1.34 \times 10^{-11}$ | | -2350 |
| AT1G70900 | protein coding gene | | 29.2 | $1.27 \times 10^{-6}$ | $7.66 \times 10^{-6}$ | mRNA-intron | 1069 |
| AT3G59040 | protein coding gene | | 29.2 | $3.17 \times 10^{-15}$ | $5.89 \times 10^{-14}$ | | -1126 |
| AT4G13120 | transposable element gene | | 29.1 | $1.17 \times 10^{-10}$ | $1.23 \times 10^{-9}$ | mRNA-exon | 1236 |

| | | | | | | | |
|---|---|---|---|---|---|---|---|
| AT5G16690 | protein coding gene | *ORC3* | 29.1 | $6.93 \times 10^{-8}$ | $4.95 \times 10^{-7}$ | mRNA-exon | 2985 |
| AT5G31804 | transposable element gene | | 29.1 | $9.19 \times 10^{-24}$ | $3.88 \times 10^{-22}$ | Intergenic | -5387 |
| AT5G61660 | protein coding gene | | 29.1 | $5.09 \times 10^{-10}$ | $4.88 \times 10^{-9}$ | mRNA-exon | 294 |
| AT3G06470 | protein coding gene | | 29.0 | $5.32 \times 10^{-10}$ | $5.08 \times 10^{-9}$ | mRNA-TTS | 2520 |
| AT2G01735 | protein coding gene | *RIE1* | 29.0 | $4.19 \times 10^{-11}$ | $4.71 \times 10^{-10}$ | mRNA-TTS | 1086 |
| AT3G11540 | protein coding gene | *SPY* | 29.0 | $1.38 \times 10^{-12}$ | $1.86 \times 10^{-11}$ | mRNA-intron | 1889 |
| AT4G06990 | anti sense lncRNA | | 28.9 | $2.10 \times 10^{-20}$ | $6.60 \times 10^{-19}$ | lnc RNA-promoter-TSS | 126 |
| AT4G23100 | protein coding gene | *GSH1* | 28.9 | $1.24 \times 10^{-6}$ | $7.51 \times 10^{-6}$ | mRNA-intron | 2448 |
| AT1G52750 | protein coding gene | | 28.9 | $2.61 \times 10^{-5}$ | $1.25 \times 10^{-4}$ | mRNA-exon | 889 |
| AT4G38600 | protein coding gene | *KAK* | 28.9 | $3.05 \times 10^{-11}$ | $3.50 \times 10^{-10}$ | mRNA-exon | 1501 |
| AT3G07160 | protein coding gene | | 28.9 | $5.55 \times 10^{-10}$ | $5.28 \times 10^{-9}$ | mRNA-intron | 1732 |
| AT5G09345 | pre tRNA | | 28.8 | $5.81 \times 10^{-9}$ | $4.82 \times 10^{-8}$ | mRNA-TTS | 799 |
| AT1G04110 | protein coding gene | *SDD1* | 28.8 | $3.00 \times 10^{-18}$ | $7.60 \times 10^{-17}$ | mRNA-exon | 958 |
| AT5G05570 | protein coding gene | | 28.8 | $2.93 \times 10^{-8}$ | $2.23 \times 10^{-7}$ | | -4330 |
| AT2G05300 | pseudogene | | 28.8 | $1.20 \times 10^{-7}$ | $8.38 \times 10^{-7}$ | mRNA-TTS | 1300 |
| AT1G09560 | protein coding gene | *GLP5* | 28.7 | $2.35 \times 10^{-13}$ | $3.49 \times 10^{-12}$ | mRNA-TTS | 2207 |
| AT4G02770 | protein coding gene | *PSAD-1* | 28.7 | $9.98 \times 10^{-8}$ | $7.02 \times 10^{-7}$ | Intergenic | -1887 |
| AT2G11700 | transposable element gene | | 28.7 | $2.00 \times 10^{-4}$ | $8.02 \times 10^{-4}$ | mRNA-exon | 466 |
| AT4G28650 | protein coding gene | | 28.7 | $3.71 \times 10^{-13}$ | $5.36 \times 10^{-12}$ | mRNA-exon | 1286 |
| AT1G30420 | protein coding gene | *ABCC11* | 28.7 | $1.22 \times 10^{-4}$ | $5.16 \times 10^{-4}$ | mRNA-intron | 2590 |
| AT4G04120 | transposable element gene | | 28.7 | $6.24 \times 10^{-16}$ | $1.26 \times 10^{-14}$ | mRNA-promoter-TSS | 807 |
| AT3G06530 | protein coding gene | | 28.6 | $6.15 \times 10^{-9}$ | $5.07 \times 10^{-8}$ | mRNA-intron | 3655 |
| AT1G73980 | protein coding gene | | 28.6 | $3.26 \times 10^{-7}$ | $2.15 \times 10^{-6}$ | mRNA-intron | 951 |
| AT4G33200 | protein coding gene | | 28.6 | $5.16 \times 10^{-10}$ | $4.94 \times 10^{-9}$ | mRNA-exon | 6496 |
| AT1G56130 | protein coding gene | | 28.6 | $7.25 \times 10^{-10}$ | $6.78 \times 10^{-9}$ | mRNA-intron | 1873 |
| AT1G59540 | protein coding gene | *ZCF125* | 28.6 | $7.15 \times 10^{-6}$ | $3.81 \times 10^{-5}$ | mRNA-exon | 2126 |
| AT3G44340 | protein coding gene | *CEF* | 28.5 | $1.67 \times 10^{-7}$ | $1.14 \times 10^{-6}$ | mRNA-intron | 6058 |
| AT3G27555 | pre tRNA | | 28.5 | $9.63 \times 10^{-15}$ | $1.69 \times 10^{-13}$ | mRNA-promoter-TSS | 823 |
| AT1G53390 | protein coding gene | *ABCG24* | 28.4 | $1.68 \times 10^{-11}$ | $1.98 \times 10^{-10}$ | mRNA-exon | 2957 |
| AT1G06613 | lncRNA | | 28.4 | $1.41 \times 10^{-6}$ | $8.44 \times 10^{-6}$ | | -1235 |
| AT5G04990 | protein coding gene | *SUN1* | 28.4 | $4.04 \times 10^{-13}$ | $5.78 \times 10^{-12}$ | mRNA-promoter-TSS | -622 |
| AT3G01380 | protein coding gene | | 28.4 | $1.20 \times 10^{-6}$ | $7.29 \times 10^{-6}$ | mRNA-intron | 5015 |
| AT1G73200 | protein coding gene | | 28.4 | $6.95 \times 10^{-9}$ | $5.71 \times 10^{-8}$ | mRNA-exon | 2067 |
| AT1G43690 | protein coding gene | | 28.4 | $2.23 \times 10^{-8}$ | $1.72 \times 10^{-7}$ | mRNA-intron | 2092 |
| AT3G09745 | anti sense lncRNA | | 28.4 | $1.67 \times 10^{-8}$ | $1.31 \times 10^{-7}$ | lnc RNA-promoter-TSS | -376 |
| AT3G09090 | protein coding gene | *DEX1* | 28.2 | $1.67 \times 10^{-5}$ | $8.36 \times 10^{-5}$ | mRNA-exon | 934 |
| AT5G39100 | protein coding gene | *GLP6* | 28.2 | $7.97 \times 10^{-6}$ | $4.19 \times 10^{-5}$ | mRNA-exon | 777 |

| | | | | | | | |
|---|---|---|---|---|---|---|---|
| AT3G49050 | protein coding gene | | 28.2 | $1.22 \times 10^{-12}$ | $1.65 \times 10^{-11}$ | mRNA-exon | 1244 |
| AT2G04840 | protein coding gene | | 28.2 | $9.05 \times 10^{-24}$ | $3.84 \times 10^{-22}$ | mRNA-TTS | 1142 |
| AT2G26000 | protein coding gene | *BRIZ2* | 28.2 | $1.35 \times 10^{-4}$ | $5.63 \times 10^{-4}$ | mRNA-exon | 1333 |
| AT5G35148 | transposable element gene | | 28.1 | $3.86 \times 10^{-9}$ | $3.28 \times 10^{-8}$ | mRNA-promoter-TSS | -459 |
| AT1G47565 | transposable element gene | | 28.1 | $1.79 \times 10^{-4}$ | $7.25 \times 10^{-4}$ | | -3271 |
| AT4G02030 | protein coding gene | | 28.1 | $2.87 \times 10^{-6}$ | $1.63 \times 10^{-5}$ | mRNA-exon | 1575 |
| AT1G67900 | protein coding gene | | 28.1 | $2.70 \times 10^{-7}$ | $1.80 \times 10^{-6}$ | mRNA-exon | 2216 |
| AT4G16250 | protein coding gene | *PHYD* | 28.1 | $6.05 \times 10^{-5}$ | $2.71 \times 10^{-4}$ | mRNA-exon | 3289 |
| AT1G07630 | protein coding gene | *PLL5* | 28.1 | $1.26 \times 10^{-8}$ | $1.00 \times 10^{-7}$ | mRNA-exon | 1736 |
| AT5G51200 | protein coding gene | *EMB3142* | 28.0 | $9.71 \times 10^{-6}$ | $5.04 \times 10^{-5}$ | mRNA-intron | 2225 |
| AT1G16300 | protein coding gene | *GAPCP-2* | 28.0 | $2.81 \times 10^{-6}$ | $1.60 \times 10^{-5}$ | mRNA-intron | 894 |
| AT3G01295 | lncRNA | | 28.0 | $6.60 \times 10^{-4}$ | $2.35 \times 10^{-3}$ | lnc RNA-promoter-TSS | -305 |
| AT4G20730 | transposable element gene | | 27.9 | $8.69 \times 10^{-25}$ | $4.15 \times 10^{-23}$ | mRNA-exon | 1744 |
| | | | 27.3 | $2.72 \times 10^{-25}$ | $1.16 \times 10^{-22}$ | mRNA-exon | 1844 |
| AT2G14247 | protein coding gene | | 27.9 | $1.03 \times 10^{-29}$ | $7.52 \times 10^{-28}$ | lnc RNA-TTS | 269 |
| AT2G31970 | protein coding gene | *RAD5* | 27.9 | $3.12 \times 10^{-6}$ | $1.77 \times 10^{-5}$ | mRNA-exon | 3120 |
| AT5G40700 | protein coding gene | | 27.9 | $2.20 \times 10^{-9}$ | $1.93 \times 10^{-8}$ | mRNA-TTS | 1162 |
| AT2G03670 | protein coding gene | *CDC48B* | 27.9 | $1.76 \times 10^{-6}$ | $1.04 \times 10^{-5}$ | mRNA-TTS | 865 |
| AT3G58430 | protein coding gene | | 27.9 | $9.70 \times 10^{-17}$ | $2.13 \times 10^{-15}$ | | -1868 |
| AT4G04895 | lncRNA | | 27.9 | $3.77 \times 10^{-5}$ | $1.76 \times 10^{-4}$ | lnc RNA-promoter-TSS | -199 |
| AT5G39510 | protein coding gene | *SGR4* | 27.9 | $7.38 \times 10^{-6}$ | $3.92 \times 10^{-5}$ | | -2042 |
| AT5G58550 | protein coding gene | *EOL2* | 27.9 | $5.11 \times 10^{-12}$ | $6.44 \times 10^{-11}$ | mRNA-exon | 1935 |
| AT5G04105 | snRNA | | 27.9 | $1.10 \times 10^{-5}$ | $5.66 \times 10^{-5}$ | snRNA-promoter-TSS | -980 |
| AT1G22740 | protein coding gene | *RABG3B* | 27.8 | $2.55 \times 10^{-6}$ | $1.47 \times 10^{-5}$ | | -1339 |
| AT1G71696 | protein coding gene | *SOL1* | 27.8 | $3.19 \times 10^{-11}$ | $3.65 \times 10^{-10}$ | mRNA-intron | 1585 |
| AT5G26594 | protein coding gene | *RR24* | 27.8 | $1.26 \times 10^{-13}$ | $1.94 \times 10^{-12}$ | | -1931 |
| AT3G29180 | protein coding gene | | 27.8 | $4.96 \times 10^{-12}$ | $6.25 \times 10^{-11}$ | mRNA-exon | 2079 |
| AT4G05915 | anti sense lncRNA | | 27.7 | $7.50 \times 10^{-6}$ | $3.97 \times 10^{-5}$ | lnc RNA-promoter-TSS | -870 |
| AT1G21170 | protein coding gene | | 27.7 | $5.61 \times 10^{-6}$ | $3.05 \times 10^{-5}$ | | -2479 |
| AT2G28640 | protein coding gene | *EXO7H5* | 27.7 | $9.91 \times 10^{-15}$ | $1.74 \times 10^{-13}$ | mRNA-exon | 1791 |
| AT3G07605 | anti sense lncRNA | | 27.7 | $3.86 \times 10^{-6}$ | $2.15 \times 10^{-5}$ | | -2940 |
| AT4G03230 | protein coding gene | | 27.7 | $7.02 \times 10^{-14}$ | $1.11 \times 10^{-12}$ | mRNA-promoter-TSS | 77 |
| AT3G58430 | protein coding gene | | 27.7 | $3.53 \times 10^{-13}$ | $5.11 \times 10^{-12}$ | | -1768 |
| AT4G11330 | protein coding gene | *MPK5* | 27.6 | $3.47 \times 10^{-10}$ | $3.40 \times 10^{-9}$ | mRNA-intron | 825 |
| AT5G20320 | protein coding gene | *DCL4* | 27.6 | $7.52 \times 10^{-6}$ | $3.97 \times 10^{-5}$ | mRNA-exon | 3915 |
| AT3G25140 | protein coding gene | *QUA1* | 27.6 | $3.10 \times 10^{-4}$ | $1.19 \times 10^{-3}$ | mRNA-exon | 1043 |
| AT2G38700 | protein coding gene | *MVD1* | 27.6 | $1.82 \times 10^{-10}$ | $1.86 \times 10^{-9}$ | mRNA-promoter-TSS | -477 |
| AT5G28937 | transposable element gene | | 27.6 | $1.76 \times 10^{-8}$ | $1.37 \times 10^{-7}$ | mRNA-promoter-TSS | -307 |

| | | | | | | | |
|---|---|---|---|---|---|---|---|
| AT5G56590 | protein coding gene | | 27.6 | $8.87 \times 10^{-9}$ | $7.19 \times 10^{-8}$ | mRNA-exon | 1387 |
| AT3G08300 | anti sense lncRNA | | 27.5 | $1.55 \times 10^{-3}$ | $5.06 \times 10^{-3}$ | | -1958 |
| AT4G31810 | protein coding gene | | 27.5 | $1.98 \times 10^{-19}$ | $5.65 \times 10^{-18}$ | | -1121 |
| AT1G79440 | protein coding gene | *ALDH5F1* | 27.5 | $8.21 \times 10^{-9}$ | $6.68 \times 10^{-8}$ | mRNA-exon | 3363 |
| AT2G28605 | protein coding gene | | 27.4 | $1.43 \times 10^{-16}$ | $3.08 \times 10^{-15}$ | mRNA-promoter-TSS | -437 |
| AT4G14970 | protein coding gene | | 27.4 | $5.72 \times 10^{-6}$ | $3.11 \times 10^{-5}$ | mRNA-exon | 2408 |
| AT1G57690 | protein coding gene | | 27.4 | $5.35 \times 10^{-14}$ | $8.61 \times 10^{-13}$ | Intergenic | -1201 |
| AT1G04700 | protein coding gene | | 27.3 | $2.90 \times 10^{-8}$ | $2.21 \times 10^{-7}$ | mRNA-exon | 1397 |
| AT2G08135 | antisense RNA | | 27.3 | $2.41 \times 10^{-6}$ | $1.40 \times 10^{-5}$ | | -1891 |
| AT5G35935 | transposable element gene | | 27.3 | $1.95 \times 10^{-10}$ | $1.99 \times 10^{-9}$ | mRNA-exon | 1740 |
| AT5G14480 | protein coding gene | | 27.2 | $1.24 \times 10^{-17}$ | $2.98 \times 10^{-16}$ | mRNA-TTS | 1777 |
| AT2G10160 | transposable element gene | | 27.2 | $1.63 \times 10^{-11}$ | $1.93 \times 10^{-10}$ | mRNA-promoter-TSS | -950 |
| AT1G60620 | protein coding gene | *RPAC43* | 27.2 | $8.92 \times 10^{-7}$ | $5.50 \times 10^{-6}$ | mRNA-exon | 987 |
| AT5G45715 | pre tRNA | | 27.2 | $1.54 \times 10^{-9}$ | $1.38 \times 10^{-8}$ | | -2000 |
| AT1G21150 | protein coding gene | | 27.2 | $9.77 \times 10^{-5}$ | $4.20 \times 10^{-4}$ | | -2381 |
| AT2G34660 | protein coding gene | *ABCC2* | 27.2 | $1.70 \times 10^{-4}$ | $6.93 \times 10^{-4}$ | mRNA-intron | 2739 |
| AT5G06155 | anti sense lncRNA | | 27.2 | $1.16 \times 10^{-6}$ | $7.05 \times 10^{-6}$ | mRNA-TTS | 128 |
| AT1G58223 | protein coding gene | | 27.2 | $3.46 \times 10^{-12}$ | $4.45 \times 10^{-11}$ | | -1032 |
| AT2G47090 | protein coding gene | | 27.2 | $4.07 \times 10^{-11}$ | $4.58 \times 10^{-10}$ | mRNA-exon | 1190 |
| AT1G09910 | protein coding gene | | 27.1 | $3.77 \times 10^{-10}$ | $3.67 \times 10^{-9}$ | mRNA-exon | 2200 |
| AT1G10910 | protein coding gene | *EMB313* | 27.1 | $1.42 \times 10^{-7}$ | $9.83 \times 10^{-7}$ | mRNA-TTS | 3716 |
| AT5G45110 | protein coding gene | *NPR3* | 27.1 | $1.49 \times 10^{-7}$ | $1.03 \times 10^{-6}$ | mRNA-exon | 1546 |
| AT5G37840 | protein coding gene | | 27.0 | $3.11 \times 10^{-17}$ | $7.19 \times 10^{-16}$ | mRNA-exon | 451 |
| AT4G34065 | pseudogene | | 27.0 | $9.50 \times 10^{-9}$ | $7.67 \times 10^{-8}$ | lnc RNA-promoter-TSS | 311 |
| AT4G19220 | protein coding gene | | 27.0 | $1.32 \times 10^{-8}$ | $1.05 \times 10^{-7}$ | mRNA-exon | 1470 |
| AT1G51480 | protein coding gene | | 27.0 | $3.70 \times 10^{-5}$ | $1.73 \times 10^{-4}$ | mRNA-TTS | 2927 |
| AT3G28345 | protein coding gene | *ABCB15* | 26.9 | $3.02 \times 10^{-10}$ | $2.99 \times 10^{-9}$ | mRNA-exon | 3391 |
| AT3G20480 | protein coding gene | *LpxK* | 26.9 | $1.32 \times 10^{-4}$ | $5.52 \times 10^{-4}$ | mRNA-intron | 3541 |
| AT1G28560 | protein coding gene | | 26.9 | $3.22 \times 10^{-10}$ | $3.18 \times 10^{-9}$ | mRNA-intron | 1193 |
| AT3G07335 | lncRNA | | 26.9 | $2.48 \times 10^{-12}$ | $3.26 \times 10^{-11}$ | lnc RNA-TTS | 682 |
| AT4G24175 | protein coding gene | | 26.9 | $2.89 \times 10^{-14}$ | $4.80 \times 10^{-13}$ | mRNA-promoter-TSS | -444 |
| AT4G15480 | protein coding gene | *UGT84A1* | 26.8 | $1.35 \times 10^{-8}$ | $1.07 \times 10^{-7}$ | mRNA-TTS | 2265 |
| AT5G24065 | pseudogene | | 26.8 | $7.30 \times 10^{-7}$ | $4.56 \times 10^{-6}$ | mRNA-promoter-TSS | 91 |
| AT2G21540 | protein coding gene | *SFH3* | 26.8 | $1.07 \times 10^{-29}$ | $7.71 \times 10^{-28}$ | mRNA-intron | 1795 |
| AT1G30430 | pre tRNA | | 26.8 | $3.47 \times 10^{-8}$ | $2.61 \times 10^{-7}$ | | -2371 |
| AT3G02750 | protein coding gene | | 26.8 | $9.95 \times 10^{-7}$ | $6.11 \times 10^{-6}$ | mRNA-exon | 1404 |
| AT2G46230 | protein coding gene | | 26.8 | $3.01 \times 10^{-7}$ | $1.99 \times 10^{-6}$ | mRNA-TTS | 908 |

| | | | | | | | |
|---|---|---|---|---|---|---|---|
| AT3G28040 | protein coding gene | | 26.8 | $1.04 \times 10^{-12}$ | $1.42 \times 10^{-11}$ | mRNA-exon | 3027 |
| AT1G30400 | protein coding gene | *ABCC1* | 26.7 | $2.18 \times 10^{-7}$ | $1.47 \times 10^{-6}$ | mRNA-exon | 2002 |
| AT1G80210 | protein coding gene | *BRCC36A* | 26.7 | $1.41 \times 10^{-6}$ | $8.42 \times 10^{-6}$ | mRNA-intron | 1304 |
| AT1G72980 | protein coding gene | *LBD7* | 26.7 | $1.23 \times 10^{-8}$ | $9.81 \times 10^{-8}$ | | -2102 |
| AT1G78922 | protein coding gene | | 26.6 | $1.31 \times 10^{-9}$ | $1.18 \times 10^{-8}$ | mRNA-TTS | 764 |
| AT2G30385 | protein coding gene | | 26.6 | $6.63 \times 10^{-10}$ | $6.24 \times 10^{-9}$ | mRNA-TTS | 954 |
| AT5G51170 | protein coding gene | | 26.6 | $8.15 \times 10^{-22}$ | $2.97 \times 10^{-20}$ | mRNA-TTS | 1348 |
| AT4G00580 | protein coding gene | | 26.6 | $1.02 \times 10^{-5}$ | $5.29 \times 10^{-5}$ | mRNA-promoter-TSS | 38 |
| AT4G00630 | protein coding gene | *KEA2* | 26.6 | $1.25 \times 10^{-5}$ | $6.38 \times 10^{-5}$ | mRNA-intron | 2380 |
| AT3G30416 | transposable element gene | | 26.6 | $3.97 \times 10^{-9}$ | $3.37 \times 10^{-8}$ | | -2156 |
| AT2G25040 | pseudogene | | 26.5 | $1.94 \times 10^{-6}$ | $1.14 \times 10^{-5}$ | mRNA-exon | 3652 |
| AT3G33080 | transposable element gene | | 26.5 | $4.69 \times 10^{-5}$ | $2.15 \times 10^{-4}$ | mRNA-promoter-TSS | 20 |
| AT1G52130 | protein coding gene | | 26.5 | $6.62 \times 10^{-8}$ | $4.74 \times 10^{-7}$ | Intergenic | -1462 |
| AT1G17350 | protein coding gene | | 26.5 | $3.81 \times 10^{-7}$ | $2.48 \times 10^{-6}$ | mRNA-intron | 1104 |
| AT1G28320 | protein coding gene | *DEG15* | 26.5 | $2.06 \times 10^{-3}$ | $6.50 \times 10^{-3}$ | mRNA-promoter-TSS | -963 |
| AT3G21890 | protein coding gene | *BBX31* | 26.5 | $1.03 \times 10^{-24}$ | $4.85 \times 10^{-23}$ | Intergenic | -1118 |
| AT3G30540 | protein coding gene | | 26.5 | $2.74 \times 10^{-7}$ | $1.82 \times 10^{-6}$ | mRNA-exon | 551 |
| AT4G04055 | anti sense lncRNA | | 26.5 | $9.91 \times 10^{-11}$ | $1.05 \times 10^{-9}$ | lnc RNA-promoter-TSS | -898 |
| AT5G38640 | protein coding gene | | 26.5 | $2.53 \times 10^{-8}$ | $1.94 \times 10^{-7}$ | mRNA-TTS | 1773 |
| AT2G09645 | anti sense lncRNA | | 26.5 | $9.65 \times 10^{-6}$ | $5.02 \times 10^{-5}$ | lnc RNA-TTS | 826 |
| AT1G52360 | protein coding gene | | 26.4 | $2.75 \times 10^{-13}$ | $4.05 \times 10^{-12}$ | mRNA-intron | 790 |
| AT4G16110 | protein coding gene | *RR2* | 26.4 | $2.01 \times 10^{-7}$ | $1.36 \times 10^{-6}$ | mRNA-exon | 2165 |
| AT2G14690 | protein coding gene | | 26.4 | $2.02 \times 10^{-10}$ | $2.05 \times 10^{-9}$ | mRNA-TTS | 2804 |
| AT1G16740 | protein coding gene | | 26.4 | $2.07 \times 10^{-7}$ | $1.40 \times 10^{-6}$ | mRNA-TTS | 1729 |
| AT5G45430 | protein coding gene | | 26.4 | $6.17 \times 10^{-10}$ | $5.83 \times 10^{-9}$ | mRNA-exon | 863 |
| AT1G64790 | protein coding gene | *ILA* | 26.4 | $1.09 \times 10^{-4}$ | $4.64 \times 10^{-4}$ | mRNA-exon | 4142 |
| AT4G32640 | protein coding gene | | 26.4 | $2.20 \times 10^{-4}$ | $8.72 \times 10^{-4}$ | mRNA-intron | 5549 |
| AT2G02210 | transposable element gene | | 26.4 | $2.02 \times 10^{-16}$ | $4.29 \times 10^{-15}$ | | -2301 |
| AT4G32215 | transposable element gene | | 26.4 | $1.74 \times 10^{-28}$ | $1.12 \times 10^{-26}$ | mRNA-TTS | 760 |
| AT3G20080 | protein coding gene | *CYP75A15* | 26.4 | $3.74 \times 10^{-7}$ | $2.44 \times 10^{-6}$ | mRNA-intron | 527 |
| AT1G08200 | protein coding gene | *AXS2* | 26.3 | $5.82 \times 10^{-23}$ | $2.30 \times 10^{-21}$ | mRNA-promoter-TSS | -507 |
| AT5G43920 | protein coding gene | | 26.3 | $5.20 \times 10^{-8}$ | $3.80 \times 10^{-7}$ | mRNA-exon | 1123 |
| AT5G26590 | transposable element gene | | 26.3 | $8.43 \times 10^{-10}$ | $7.84 \times 10^{-9}$ | mRNA-TTS | 1880 |
| AT2G45350 | protein coding gene | *CRR4* | 26.3 | $1.53 \times 10^{-28}$ | $9.94 \times 10^{-27}$ | mRNA-exon | 840 |
| AT3G55650 | protein coding gene | | 26.3 | $6.90 \times 10^{-18}$ | $1.69 \times 10^{-16}$ | mRNA-TTS | 766 |
| AT5G65445 | pre tRNA | | 26.3 | $2.76 \times 10^{-13}$ | $4.06 \times 10^{-12}$ | tRNA-TTS | 919 |
| AT2G29070 | protein coding gene | | 26.2 | $2.69 \times 10^{-9}$ | $2.33 \times 10^{-8}$ | mRNA-promoter-TSS | -497 |

| | | | | | | | |
|---|---|---|---|---|---|---|---|
| AT5G46860 | protein coding gene | *VAM3* | 26.2 | $5.48 \times 10^{-6}$ | $2.99 \times 10^{-5}$ | mRNA-TTS | 1359 |
| AT4G00720 | protein coding gene | *SK32* | 26.2 | $4.43 \times 10^{-8}$ | $3.26 \times 10^{-7}$ | mRNA-exon | 1428 |
| AT3G48190 | protein coding gene | | 26.2 | $5.44 \times 10^{-4}$ | $1.97 \times 10^{-3}$ | mRNA-intron | 6262 |
| AT3G11673 | pseudogene | | 26.2 | $1.13 \times 10^{-3}$ | $3.80 \times 10^{-3}$ | | -1304 |
| AT4G00760 | protein coding gene | *APRR8* | 26.2 | $1.94 \times 10^{-4}$ | $7.80 \times 10^{-4}$ | mRNA-promoter-TSS | -182 |
| AT3G01115 | lncRNA | | 26.2 | $7.61 \times 10^{-7}$ | $4.75 \times 10^{-6}$ | mRNA-exon | 3670 |
| AT3G07980 | protein coding gene | *MAPKKK6* | 26.2 | $6.35 \times 10^{-4}$ | $2.27 \times 10^{-3}$ | mRNA-intron | 907 |
| AT1G12150 | protein coding gene | | 26.1 | $5.34 \times 10^{-8}$ | $3.88 \times 10^{-7}$ | mRNA-TTS | 604 |
| AT2G01950 | protein coding gene | *BRL2* | 26.1 | $1.39 \times 10^{-11}$ | $1.65 \times 10^{-10}$ | mRNA-exon | 2073 |
| AT5G46070 | protein coding gene | | 26.1 | $1.45 \times 10^{-4}$ | $6.01 \times 10^{-4}$ | mRNA-intron | 2832 |
| AT5G40450 | protein coding gene | | 26.0 | $3.55 \times 10^{-16}$ | $7.31 \times 10^{-15}$ | mRNA-exon | 3900 |
| AT3G50040 | protein coding gene | | 26.0 | $5.41 \times 10^{-11}$ | $5.98 \times 10^{-10}$ | | -2876 |
| AT5G46680 | protein coding gene | | 26.0 | $3.47 \times 10^{-11}$ | $3.94 \times 10^{-10}$ | mRNA-exon | 948 |
| AT3G02725 | snoRNA | | 26.0 | $8.15 \times 10^{-7}$ | $5.06 \times 10^{-6}$ | mRNA-exon | 1361 |
| AT2G09500 | anti sense lncRNA | | 26.0 | $1.02 \times 10^{-7}$ | $7.16 \times 10^{-7}$ | | -1299 |
| AT4G12620 | protein coding gene | *ORC1B* | 26.0 | $1.14 \times 10^{-16}$ | $2.49 \times 10^{-15}$ | mRNA-exon | 1096 |
| AT2G33233 | protein coding gene | *LCR49* | 26.0 | $9.21 \times 10^{-17}$ | $2.04 \times 10^{-15}$ | | -4537 |
| AT5G61900 | protein coding gene | *BON1* | 26.0 | $7.28 \times 10^{-7}$ | $4.55 \times 10^{-6}$ | | -1710 |
| AT5G08470 | protein coding gene | *PEX1* | 26.0 | $4.32 \times 10^{-6}$ | $2.39 \times 10^{-5}$ | mRNA-intron | 3626 |
| AT3G48150 | protein coding gene | *APC8* | 26.0 | $4.08 \times 10^{-10}$ | $3.96 \times 10^{-9}$ | mRNA-intron | 518 |
| AT5G30490 | protein coding gene | | 26.0 | $3.54 \times 10^{-6}$ | $1.98 \times 10^{-5}$ | mRNA-intron | 1249 |
| AT2G07700 | transposable element gene | | 25.9 | $3.16 \times 10^{-8}$ | $2.39 \times 10^{-7}$ | Intergenic | -1150 |
| AT5G10350 | protein coding gene | | 25.9 | $8.87 \times 10^{-14}$ | $1.39 \times 10^{-12}$ | mRNA-TTS | 1600 |
| AT3G46680 | protein coding gene | | 25.9 | $5.97 \times 10^{-8}$ | $4.29 \times 10^{-7}$ | transcript-TTS | 892 |
| AT1G62250 | protein coding gene | | 25.9 | $2.29 \times 10^{-5}$ | $1.12 \times 10^{-4}$ | mRNA-promoter-TSS | -842 |
| AT5G27607 | protein coding gene | | 25.9 | $2.97 \times 10^{-8}$ | $2.25 \times 10^{-7}$ | mRNA-TTS | 1558 |
| AT1G80350 | protein coding gene | *ERH3* | 25.9 | $5.58 \times 10^{-14}$ | $8.95 \times 10^{-13}$ | mRNA-exon | 2617 |
| AT2G18340 | protein coding gene | | 25.9 | $2.66 \times 10^{-8}$ | $2.04 \times 10^{-7}$ | mRNA-TTS | 807 |
| AT3G01515 | anti sense lncRNA | | 25.9 | $5.72 \times 10^{-6}$ | $3.11 \times 10^{-5}$ | | -1078 |
| AT2G03420 | protein coding gene | | 25.9 | $9.97 \times 10^{-13}$ | $1.37 \times 10^{-11}$ | mRNA-promoter-TSS | -682 |
| AT4G32510 | protein coding gene | | 25.8 | $6.38 \times 10^{-8}$ | $4.57 \times 10^{-7}$ | mRNA-TTS | 3660 |
| AT2G34880 | protein coding gene | *MEE27* | 25.8 | $4.94 \times 10^{-7}$ | $3.18 \times 10^{-6}$ | | -1917 |
| AT2G07050 | protein coding gene | *CAS1* | 25.8 | $4.58 \times 10^{-4}$ | $1.69 \times 10^{-3}$ | mRNA-intron | 2591 |
| AT5G08210 | miRNA | | 25.8 | $1.19 \times 10^{-5}$ | $6.10 \times 10^{-5}$ | mRNA-exon | 1446 |
| AT1G36395 | transposable element gene | | 25.7 | $9.03 \times 10^{-4}$ | $3.10 \times 10^{-3}$ | mRNA-TTS | 871 |
| AT3G09200 | protein coding gene | | 25.7 | $6.56 \times 10^{-7}$ | $4.13 \times 10^{-6}$ | mRNA-promoter-TSS | -310 |
| AT3G52050 | protein coding gene | | 25.7 | $5.13 \times 10^{-7}$ | $3.28 \times 10^{-6}$ | mRNA-intron | 1258 |

| | | | | | | | |
|---|---|---|---|---|---|---|---|
| AT4G11380 | protein coding gene | | 25.7 | $4.53 \times 10^{-7}$ | $2.92 \times 10^{-6}$ | mRNA-exon | 2162 |
| AT2G36760 | protein coding gene | *UGT73C2* | 25.7 | $2.47 \times 10^{-13}$ | $3.65 \times 10^{-12}$ | mRNA-exon | 743 |
| AT1G41827 | transposable element gene | | 25.6 | $1.69 \times 10^{-4}$ | $6.90 \times 10^{-4}$ | Intergenic | 3376 |
| AT3G59530 | protein coding gene | *LAP3* | 25.6 | $2.08 \times 10^{-6}$ | $1.22 \times 10^{-5}$ | mRNA-promoter-TSS | -502 |
| AT3G25840 | protein coding gene | | 25.6 | $2.58 \times 10^{-6}$ | $1.48 \times 10^{-5}$ | mRNA-exon | 3103 |
| AT3G54290 | protein coding gene | | 25.6 | $8.04 \times 10^{-12}$ | $9.87 \times 10^{-11}$ | mRNA-exon | 5064 |
| AT5G10370 | protein coding gene | | 25.6 | $2.79 \times 10^{-7}$ | $1.85 \times 10^{-6}$ | mRNA-exon | 1702 |
| AT4G38215 | anti sense lncRNA | | 25.6 | $1.43 \times 10^{-5}$ | $7.23 \times 10^{-5}$ | | -2362 |
| AT2G27170 | protein coding gene | *TTN7* | 25.6 | $2.24 \times 10^{-3}$ | $7.00 \times 10^{-3}$ | mRNA-exon | 2634 |
| AT5G09660 | protein coding gene | *PMDH2* | 25.6 | $3.61 \times 10^{-7}$ | $2.37 \times 10^{-6}$ | mRNA-promoter-TSS | -667 |
| AT4G12900 | protein coding gene | | 25.5 | $2.20 \times 10^{-6}$ | $1.28 \times 10^{-5}$ | mRNA-promoter-TSS | -214 |
| AT5G16860 | protein coding gene | | 25.5 | $1.41 \times 10^{-4}$ | $5.88 \times 10^{-4}$ | mRNA-exon | 1417 |
| AT1G32330 | protein coding gene | *HSFA1D* | 25.5 | $2.51 \times 10^{-14}$ | $4.19 \times 10^{-13}$ | mRNA-exon | 2410 |
| AT2G08120 | anti sense lncRNA | | 25.5 | $1.60 \times 10^{-4}$ | $6.57 \times 10^{-4}$ | | -2498 |
| AT2G08295 | lncRNA | | 25.5 | $1.99 \times 10^{-10}$ | $2.03 \times 10^{-9}$ | lnc RNA-TTS | 301 |
| AT5G46380 | protein coding gene | | 25.5 | $2.22 \times 10^{-15}$ | $4.19 \times 10^{-14}$ | mRNA-intron | 1323 |
| AT1G63260 | protein coding gene | *TET1* | 25.4 | $2.70 \times 10^{-8}$ | $2.07 \times 10^{-7}$ | mRNA-intron | 1096 |
| AT2G38700 | protein coding gene | *MVD1* | 25.4 | $1.95 \times 10^{-4}$ | $7.85 \times 10^{-4}$ | | -1177 |
| AT2G20825 | protein coding gene | *ULT2* | 25.4 | $1.51 \times 10^{-9}$ | $1.36 \times 10^{-8}$ | mRNA-promoter-TSS | -408 |
| AT3G30715 | transposable element gene | | 25.4 | $1.82 \times 10^{-6}$ | $1.07 \times 10^{-5}$ | Intergenic | 7660 |
| AT1G35220 | protein coding gene | | 25.4 | $5.93 \times 10^{-5}$ | $2.67 \times 10^{-4}$ | mRNA-exon | 2150 |
| AT1G69350 | protein coding gene | | 25.4 | $1.26 \times 10^{-11}$ | $1.51 \times 10^{-10}$ | mRNA-TTS | 671 |
| AT2G04545 | anti sense lncRNA | | 25.4 | $5.20 \times 10^{-8}$ | $3.79 \times 10^{-7}$ | lnc RNA-promoter-TSS | -414 |
| AT1G40123 | transposable element gene | | 25.3 | $3.69 \times 10^{-20}$ | $1.13 \times 10^{-18}$ | Intergenic | -1408 |
| AT3G28674 | protein coding gene | | 25.3 | $1.51 \times 10^{-5}$ | $7.61 \times 10^{-5}$ | mRNA-intron | 1543 |
| AT3G41768 | rRNA | | 25.3 | | | mRNA-promoter-TSS | -226 |
| AT3G56810 | protein coding gene | | 25.3 | $6.09 \times 10^{-12}$ | $7.59 \times 10^{-11}$ | | -1177 |
| AT2G09325 | anti sense lncRNA | | 25.3 | $2.26 \times 10^{-5}$ | $1.10 \times 10^{-4}$ | mRNA-exon | 2445 |
| AT2G46920 | protein coding gene | *POL* | 25.3 | $7.74 \times 10^{-5}$ | $3.40 \times 10^{-4}$ | mRNA-exon | 1621 |
| AT3G59570 | protein coding gene | | 25.3 | $1.36 \times 10^{-19}$ | $3.94 \times 10^{-18}$ | mRNA-TTS | 4481 |
| AT2G34720 | protein coding gene | *NF-YA4* | 25.3 | $2.83 \times 10^{-6}$ | $1.62 \times 10^{-5}$ | mRNA-exon | 141 |
| AT3G01780 | protein coding gene | *TPLATE* | 25.3 | $1.71 \times 10^{-21}$ | $6.04 \times 10^{-20}$ | mRNA-exon | 1482 |
| AT4G09130 | protein coding gene | | 25.3 | $2.97 \times 10^{-10}$ | $2.95 \times 10^{-9}$ | mRNA-TTS | 376 |
| AT1G51480 | protein coding gene | | 25.3 | $1.62 \times 10^{-10}$ | $1.68 \times 10^{-9}$ | mRNA-exon | 1427 |
| AT5G50030 | protein coding gene | | 25.2 | $8.55 \times 10^{-27}$ | $4.83 \times 10^{-25}$ | mRNA-promoter-TSS | 62 |
| AT1G47550 | protein coding gene | *SEC3A* | 25.2 | $6.27 \times 10^{-5}$ | $2.80 \times 10^{-4}$ | mRNA-intron | 1129 |
| AT5G24960 | protein coding gene | *CYP71A14* | 25.2 | $1.01 \times 10^{-5}$ | $5.21 \times 10^{-5}$ | | -2530 |

| | | | | | | | |
|---|---|---|---|---|---|---|---|
| AT3G50640 | protein coding gene | | 25.2 | $2.28 \times 10^{-5}$ | $1.11 \times 10^{-4}$ | mRNA-TTS | 1380 |
| AT1G52942 | pseudogene | | 25.2 | $1.65 \times 10^{-6}$ | $9.78 \times 10^{-6}$ | | -1071 |
| AT2G04525 | anti sense lncRNA | | 25.2 | $3.64 \times 10^{-7}$ | $2.38 \times 10^{-6}$ | lnc RNA-promoter-TSS | -715 |
| AT3G20100 | protein coding gene | *CYP75A19* | 25.1 | $1.76 \times 10^{-13}$ | $2.65 \times 10^{-12}$ | mRNA-promoter-TSS | -875 |
| AT3G13080 | protein coding gene | *ABCC3* | 25.1 | $8.84 \times 10^{-11}$ | $9.48 \times 10^{-10}$ | | -3017 |
| AT2G10160 | transposable element gene | | 25.1 | $1.72 \times 10^{-5}$ | $8.55 \times 10^{-5}$ | mRNA-promoter-TSS | -750 |
| AT1G76330 | pre tRNA | | 25.1 | $3.57 \times 10^{-7}$ | $2.34 \times 10^{-6}$ | mRNA-TTS | 459 |
| AT5G40480 | protein coding gene | *EMB312* | 25.1 | $1.46 \times 10^{-4}$ | $6.07 \times 10^{-4}$ | mRNA-exon | 4815 |
| AT4G38880 | protein coding gene | *ASE3* | 25.0 | $1.71 \times 10^{-9}$ | $1.52 \times 10^{-8}$ | mRNA-TTS | 2035 |
| AT1G61040 | protein coding gene | *VIP5* | 25.0 | $2.03 \times 10^{-5}$ | $9.97 \times 10^{-5}$ | | -1753 |
| AT5G46680 | protein coding gene | | 25.0 | $5.10 \times 10^{-25}$ | $2.46 \times 10^{-23}$ | mRNA-exon | 848 |
| AT1G08750 | protein coding gene | | 25.0 | $1.27 \times 10^{-6}$ | $7.64 \times 10^{-6}$ | mRNA-intron | 1673 |
| AT2G38840 | protein coding gene | | 25.0 | $3.18 \times 10^{-8}$ | $2.40 \times 10^{-7}$ | mRNA-exon | 2757 |
| AT5G43810 | protein coding gene | *AGO1* | 25.0 | $3.60 \times 10^{-12}$ | $4.63 \times 10^{-11}$ | mRNA-exon | 3475 |
| AT3G49420 | protein coding gene | | -25.0 | $1.09 \times 10^{-5}$ | $5.61 \times 10^{-5}$ | mRNA-promoter-TSS | -479 |
| AT1G09030 | protein coding gene | *NF-YB4* | -25.0 | $3.26 \times 10^{-3}$ | $9.73 \times 10^{-3}$ | Intergenic | 1881 |
| AT3G25510 | protein coding gene | | -25.0 | $2.45 \times 10^{-10}$ | $2.46 \times 10^{-9}$ | mRNA-TTS | 401 |
| AT5G62600 | protein coding gene | *MOS14* | -25.1 | $1.04 \times 10^{-3}$ | $3.52 \times 10^{-3}$ | mRNA-intron | 4165 |
| AT3G10810 | protein coding gene | | -25.1 | $2.03 \times 10^{-5}$ | $9.97 \times 10^{-5}$ | mRNA-intron | 872 |
| AT2G37570 | protein coding gene | *SLT1* | -25.1 | $1.46 \times 10^{-6}$ | $8.69 \times 10^{-6}$ | mRNA-exon | 732 |
| AT2G25171 | miRNA | | -25.2 | $1.11 \times 10^{-3}$ | $3.73 \times 10^{-3}$ | lnc RNA-TTS | 1181 |
| AT4G05585 | transposable element gene | | -25.2 | $1.84 \times 10^{-10}$ | $1.89 \times 10^{-9}$ | mRNA-promoter-TSS | -506 |
| AT3G19520 | protein coding gene | | -25.3 | $1.32 \times 10^{-31}$ | $1.10 \times 10^{-29}$ | mRNA-promoter-TSS | 64 |
| AT2G30570 | protein coding gene | *PSBW* | -25.4 | $3.18 \times 10^{-8}$ | $2.40 \times 10^{-7}$ | | -1840 |
| AT5G57440 | protein coding gene | *GS1* | -25.4 | $1.25 \times 10^{-6}$ | $7.53 \times 10^{-6}$ | mRNA-TTS | 870 |
| AT1G06070 | protein coding gene | | -25.5 | $1.88 \times 10^{-9}$ | $1.67 \times 10^{-8}$ | mRNA-TTS | 2107 |
| AT1G74210 | protein coding gene | *GDPD5* | -25.5 | $1.82 \times 10^{-9}$ | $1.61 \times 10^{-8}$ | mRNA-exon | 937 |
| AT4G04425 | anti sense lncRNA | | -25.6 | $3.26 \times 10^{-9}$ | $2.78 \times 10^{-8}$ | lnc RNA-TTS | 951 |
| AT5G20980 | protein coding gene | *MS3* | -25.6 | $7.27 \times 10^{-6}$ | $3.86 \times 10^{-5}$ | | -1239 |
| AT1G08870 | pre tRNA | | -25.6 | $7.79 \times 10^{-5}$ | $3.42 \times 10^{-4}$ | mRNA-intron | 1557 |
| AT3G51400 | protein coding gene | | -25.6 | $8.21 \times 10^{-15}$ | $1.46 \times 10^{-13}$ | mRNA-TTS | 1204 |
| AT1G63640 | protein coding gene | | -25.6 | $1.89 \times 10^{-6}$ | $1.11 \times 10^{-5}$ | mRNA-intron | 824 |
| AT2G24205 | protein coding gene | | -25.7 | $2.98 \times 10^{-4}$ | $1.15 \times 10^{-3}$ | mRNA-exon | 103 |
| AT5G52280 | protein coding gene | | -25.7 | $7.60 \times 10^{-10}$ | $7.10 \times 10^{-9}$ | mRNA-TTS | 2530 |
| AT5G01450 | protein coding gene | *APD2* | -25.7 | $1.39 \times 10^{-18}$ | $3.65 \times 10^{-17}$ | mRNA-exon | 1010 |
| AT5G66140 | protein coding gene | *PAD2* | -25.8 | $3.86 \times 10^{-8}$ | $2.87 \times 10^{-7}$ | | -1380 |
| AT1G73920 | protein coding gene | | -25.8 | $1.00 \times 10^{-4}$ | $4.31 \times 10^{-4}$ | mRNA-intron | 2375 |

| | | | | | | | |
|---|---|---|---|---|---|---|---|
| AT1G09045 | novel transcribed region | | -25.9 | $1.37 \times 10^{-6}$ | $8.21 \times 10^{-6}$ | transcript-promoter-TSS | -128 |
| AT3G11540 | protein coding gene | *SPY* | -25.9 | $4.06 \times 10^{-5}$ | $1.89 \times 10^{-4}$ | mRNA-intron | 1989 |
| AT2G40030 | protein coding gene | *NRPD1B* | -25.9 | $1.12 \times 10^{-5}$ | $5.76 \times 10^{-5}$ | mRNA-intron | 1391 |
| AT1G71190 | protein coding gene | *SAG18* | -25.9 | $2.40 \times 10^{-12}$ | $3.16 \times 10^{-11}$ | mRNA-TTS | 859 |
| AT1G05890 | protein coding gene | *ARI5* | -25.9 | $7.54 \times 10^{-6}$ | $3.98 \times 10^{-5}$ | mRNA-exon | 1448 |
| AT2G35630 | protein coding gene | *MOR1* | -25.9 | $2.33 \times 10^{-7}$ | $1.56 \times 10^{-6}$ | mRNA-exon | 3416 |
| AT5G65300 | protein coding gene | | -26.1 | $2.03 \times 10^{-8}$ | $1.58 \times 10^{-7}$ | mRNA-exon | 2463 |
| AT1G75210 | protein coding gene | | -26.1 | $2.71 \times 10^{-4}$ | $1.06 \times 10^{-3}$ | mRNA-exon | 1914 |
| AT3G08075 | anti sense lncRNA | | -26.2 | $1.35 \times 10^{-9}$ | $1.22 \times 10^{-8}$ | lnc RNA-promoter-TSS | -317 |
| AT5G51230 | protein coding gene | *EMF2* | -26.3 | $5.73 \times 10^{-9}$ | $4.76 \times 10^{-8}$ | mRNA-intron | 3417 |
| AT4G00026 | protein coding gene | *SD3* | -26.3 | $4.22 \times 10^{-17}$ | $9.59 \times 10^{-16}$ | mRNA-promoter-TSS | 708 |
| AT3G56680 | protein coding gene | | -26.3 | $7.79 \times 10^{-9}$ | $6.35 \times 10^{-8}$ | mRNA-TTS | 1671 |
| AT3G02290 | protein coding gene | | -26.3 | $5.71 \times 10^{-9}$ | $4.75 \times 10^{-8}$ | | -1805 |
| AT1G06720 | protein coding gene | | -26.3 | $4.73 \times 10^{-4}$ | $1.74 \times 10^{-3}$ | mRNA-exon | 3072 |
| AT3G59810 | protein coding gene | *LSM6A* | -26.4 | $1.51 \times 10^{-3}$ | $4.95 \times 10^{-3}$ | mRNA-TTS | 2248 |
| AT4G24560 | protein coding gene | *UBP16* | -26.4 | $3.36 \times 10^{-5}$ | $1.59 \times 10^{-4}$ | mRNA-intron | 4125 |
| AT4G31490 | protein coding gene | | -26.4 | $1.19 \times 10^{-6}$ | $7.20 \times 10^{-6}$ | | -2095 |
| AT3G12120 | protein coding gene | *FAD2* | -26.4 | $3.63 \times 10^{-10}$ | $3.55 \times 10^{-9}$ | | -1601 |
| AT2G03855 | miRNA | | -26.4 | $1.44 \times 10^{-24}$ | $2.99 \times 10^{-21}$ | rRNA-promoter-TSS | 146 |
| AT3G03730 | protein coding gene | | -26.5 | $7.39 \times 10^{-12}$ | $9.11 \times 10^{-11}$ | mRNA-exon | 469 |
| AT1G72710 | protein coding gene | *CKL2* | -26.5 | $6.62 \times 10^{-6}$ | $3.55 \times 10^{-5}$ | | -2946 |
| AT5G10800 | protein coding gene | | -26.6 | $4.77 \times 10^{-8}$ | $3.50 \times 10^{-7}$ | mRNA-intron | 664 |
| AT3G48190 | protein coding gene | | -26.7 | $6.38 \times 10^{-5}$ | $2.85 \times 10^{-4}$ | mRNA-exon | 12462 |
| AT4G16890 | protein coding gene | *SNC1* | -26.7 | $4.25 \times 10^{-23}$ | $1.69 \times 10^{-21}$ | mRNA-exon | 304 |
| AT1G09827 | anti sense lncRNA | | -26.8 | $3.75 \times 10^{-10}$ | $3.66 \times 10^{-9}$ | | -1297 |
| AT2G01045 | protein coding gene | | -26.9 | $1.76 \times 10^{-21}$ | $6.18 \times 10^{-20}$ | Intergenic | -1083 |
| AT1G78920 | protein coding gene | *VP2* | -26.9 | $2.41 \times 10^{-3}$ | $7.46 \times 10^{-3}$ | mRNA-exon | 1271 |
| AT1G70060 | protein coding gene | *SNL4* | -27.0 | $3.75 \times 10^{-14}$ | $6.14 \times 10^{-13}$ | mRNA-exon | 3411 |
| AT1G08073 | anti sense lncRNA | | -27.1 | $1.85 \times 10^{-3}$ | $5.91 \times 10^{-3}$ | mRNA-exon | 4093 |
| AT2G26840 | protein coding gene | *MutS2* | -27.1 | $3.41 \times 10^{-9}$ | $2.91 \times 10^{-8}$ | mRNA-TTS | 2663 |
| AT1G63800 | protein coding gene | *UBC5* | -27.1 | $6.13 \times 10^{-7}$ | $3.87 \times 10^{-6}$ | | -3306 |
| AT5G18540 | protein coding gene | | -27.2 | $2.79 \times 10^{-4}$ | $1.09 \times 10^{-3}$ | | -1534 |
| AT1G77510 | protein coding gene | *PDIL1-2* | -27.2 | $1.72 \times 10^{-7}$ | $1.17 \times 10^{-6}$ | | -2597 |
| AT2G32980 | protein coding gene | *Aug-02* | -27.4 | $6.42 \times 10^{-4}$ | $2.29 \times 10^{-3}$ | mRNA-intron | 1298 |
| AT3G18485 | protein coding gene | *ILR2* | -27.4 | $7.40 \times 10^{-14}$ | $1.17 \times 10^{-12}$ | mRNA-exon | 2327 |
| AT1G79610 | protein coding gene | *NHX6* | -27.5 | $3.23 \times 10^{-4}$ | $1.24 \times 10^{-3}$ | mRNA-intron | 1550 |
| AT2G26780 | protein coding gene | | -27.5 | $9.66 \times 10^{-12}$ | $1.17 \times 10^{-10}$ | mRNA-exon | 3475 |

| | | | | | | | |
|---|---|---|---|---|---|---|---|
| AT5G38460 | protein coding gene | | -27.5 | $2.99 \times 10^{-13}$ | $4.36 \times 10^{-12}$ | mRNA-exon | 928 |
| AT5G64050 | protein coding gene | *ERS* | -27.6 | $8.98 \times 10^{-15}$ | $1.58 \times 10^{-13}$ | mRNA-intron | 1058 |
| AT1G24090 | protein coding gene | | -27.6 | $7.65 \times 10^{-26}$ | $3.94 \times 10^{-24}$ | tRNA-promoter-TSS | 608 |
| AT1G05460 | protein coding gene | *SDE3* | -27.6 | $3.74 \times 10^{-6}$ | $2.09 \times 10^{-5}$ | mRNA-exon | 1276 |
| AT1G63340 | protein coding gene | | -27.6 | $5.19 \times 10^{-13}$ | $7.36 \times 10^{-12}$ | mRNA-promoter-TSS | -732 |
| AT1G42705 | transposable element gene | | -27.6 | $3.67 \times 10^{-5}$ | $1.72 \times 10^{-4}$ | mRNA-promoter-TSS | -790 |
| AT2G16560 | transposable element gene | | -27.6 | $2.43 \times 10^{-6}$ | $1.41 \times 10^{-5}$ | mRNA-exon | 2348 |
| AT2G39810 | protein coding gene | *HOS1* | -27.7 | $3.34 \times 10^{-6}$ | $1.88 \times 10^{-5}$ | mRNA-intron | 2457 |
| AT5G07810 | protein coding gene | | -27.7 | $3.01 \times 10^{-8}$ | $2.28 \times 10^{-7}$ | mRNA-intron | 2391 |
| AT4G28470 | protein coding gene | *RPN1B* | -27.7 | $4.72 \times 10^{-5}$ | $2.16 \times 10^{-4}$ | mRNA-intron | 1631 |
| AT4G17140 | protein coding gene | | -27.8 | $2.54 \times 10^{-5}$ | $1.23 \times 10^{-4}$ | mRNA-exon | 7322 |
| AT2G27160 | protein coding gene | | -27.8 | $4.92 \times 10^{-6}$ | $2.70 \times 10^{-5}$ | | -4035 |
| AT5G11560 | protein coding gene | *catalytics* | -27.8 | $8.23 \times 10^{-12}$ | $1.01 \times 10^{-10}$ | mRNA-intron | 773 |
| AT2G31345 | protein coding gene | | -27.9 | $2.73 \times 10^{-9}$ | $2.37 \times 10^{-8}$ | mRNA-TTS | 1249 |
| AT1G49160 | protein coding gene | | -27.9 | $1.58 \times 10^{-7}$ | $1.08 \times 10^{-6}$ | mRNA-intron | 1102 |
| AT3G54750 | protein coding gene | | -27.9 | $4.32 \times 10^{-12}$ | $5.50 \times 10^{-11}$ | | -1166 |
| AT5G09680 | protein coding gene | | -27.9 | $1.09 \times 10^{-3}$ | $3.67 \times 10^{-3}$ | mRNA-promoter-TSS | -993 |
| AT4G22830 | protein coding gene | | -28.1 | $2.65 \times 10^{-6}$ | $1.52 \times 10^{-5}$ | mRNA-promoter-TSS | -860 |
| AT2G41780 | protein coding gene | | -28.2 | $7.48 \times 10^{-5}$ | $3.29 \times 10^{-4}$ | mRNA-exon | 2045 |
| AT5G48335 | protein coding gene | | -28.2 | $2.91 \times 10^{-3}$ | $8.82 \times 10^{-3}$ | mRNA-promoter-TSS | -921 |
| AT5G65570 | protein coding gene | | -28.2 | $2.31 \times 10^{-9}$ | $2.02 \times 10^{-8}$ | mRNA-exon | 1383 |
| AT3G48030 | protein coding gene | | -28.2 | $3.08 \times 10^{-10}$ | $3.05 \times 10^{-9}$ | mRNA-exon | 1103 |
| AT2G38790 | protein coding gene | | -28.4 | $1.42 \times 10^{-14}$ | $2.43 \times 10^{-13}$ | mRNA-TTS | 2013 |
| AT5G59420 | protein coding gene | *ORP3C* | -28.4 | $1.97 \times 10^{-7}$ | $1.33 \times 10^{-6}$ | mRNA-intron | 2084 |
| AT1G02580 | protein coding gene | *MEA* | -28.4 | $1.52 \times 10^{-4}$ | $6.27 \times 10^{-4}$ | mRNA-intron | 618 |
| AT1G17100 | protein coding gene | *HBP1* | -28.4 | $1.19 \times 10^{-10}$ | $1.25 \times 10^{-9}$ | mRNA-TTS | 1605 |
| AT1G10460 | protein coding gene | *GLP7* | -28.4 | $1.02 \times 10^{-4}$ | $4.35 \times 10^{-4}$ | | -1692 |
| AT5G52220 | protein coding gene | | -28.5 | $1.01 \times 10^{-8}$ | $8.15 \times 10^{-8}$ | | -1235 |
| AT2G37140 | protein coding gene | | -28.5 | $2.95 \times 10^{-13}$ | $4.31 \times 10^{-12}$ | mRNA-exon | 1988 |
| AT3G44980 | protein coding gene | | -28.6 | $5.02 \times 10^{-9}$ | $4.21 \times 10^{-8}$ | Intergenic | -1472 |
| AT5G63890 | protein coding gene | *HDH* | -28.6 | $4.71 \times 10^{-8}$ | $3.45 \times 10^{-7}$ | mRNA-intron | 999 |
| AT4G15720 | protein coding gene | | -28.6 | $3.15 \times 10^{-6}$ | $1.78 \times 10^{-5}$ | mRNA-TTS | 1391 |
| AT3G53100 | protein coding gene | | -28.7 | $4.23 \times 10^{-5}$ | $1.96 \times 10^{-4}$ | mRNA-TTS | 3133 |
| AT5G55850 | protein coding gene | *NOI* | -28.7 | $2.47 \times 10^{-3}$ | $7.64 \times 10^{-3}$ | | -1851 |
| AT4G15415 | protein coding gene | *ATB' GAMMA* | -28.8 | $5.35 \times 10^{-8}$ | $3.89 \times 10^{-7}$ | mRNA-exon | 1219 |
| AT3G06220 | protein coding gene | | -28.8 | $9.33 \times 10^{-11}$ | $9.94 \times 10^{-10}$ | mRNA-TTS | 1719 |
| AT5G08305 | protein coding gene | | -28.9 | $6.23 \times 10^{-8}$ | $4.47 \times 10^{-7}$ | mRNA-TTS | 1131 |

| | | | | | | | |
|---|---|---|---|---|---|---|---|
| AT5G61780 | protein coding gene | *Tudor2* | -29.0 | $2.80 \times 10^{-6}$ | $1.60 \times 10^{-5}$ | mRNA-exon | 2948 |
| AT3G01405 | novel transcribed region | | -29.0 | $3.17 \times 10^{-13}$ | $4.62 \times 10^{-12}$ | lnc RNA-promoter-TSS | -272 |
| AT5G21030 | protein coding gene | | -29.2 | $4.42 \times 10^{-13}$ | $6.30 \times 10^{-12}$ | | -1179 |
| AT1G31800 | protein coding gene | *CYP97A3* | -29.2 | $1.03 \times 10^{-10}$ | $1.09 \times 10^{-9}$ | mRNA-exon | 1849 |
| AT5G55670 | protein coding gene | | -29.3 | $4.47 \times 10^{-13}$ | $6.37 \times 10^{-12}$ | mRNA-intron | 2250 |
| AT3G53400 | protein coding gene | | -29.3 | $2.72 \times 10^{-8}$ | $2.08 \times 10^{-7}$ | | -2756 |
| AT2G41220 | protein coding gene | *GLU2* | -29.4 | $8.57 \times 10^{-16}$ | $1.70 \times 10^{-14}$ | mRNA-intron | 3312 |
| AT1G78880 | protein coding gene | | -29.4 | $7.15 \times 10^{-6}$ | $3.81 \times 10^{-5}$ | mRNA-TTS | 2398 |
| AT3G15610 | protein coding gene | | -29.5 | $1.55 \times 10^{-15}$ | $2.99 \times 10^{-14}$ | mRNA-promoter-TSS | -610 |
| AT1G04423 | miRNA | | -29.5 | $2.76 \times 10^{-12}$ | $3.60 \times 10^{-11}$ | mRNA-TTS | 700 |
| AT2G32110 | pre tRNA | | -29.6 | $4.28 \times 10^{-11}$ | $4.80 \times 10^{-10}$ | tRNA-TTS | 977 |
| AT1G78280 | protein coding gene | | -29.6 | $4.69 \times 10^{-10}$ | $4.51 \times 10^{-9}$ | mRNA-exon | 2896 |
| AT3G57110 | protein coding gene | | -29.6 | $8.40 \times 10^{-24}$ | $3.59 \times 10^{-22}$ | mRNA-promoter-TSS | 375 |
| AT1G35450 | pseudogene | | -29.7 | $6.68 \times 10^{-9}$ | $5.49 \times 10^{-8}$ | transcript-exon | 401 |
| AT5G05010 | protein coding gene | | -29.7 | $7.13 \times 10^{-4}$ | $2.51 \times 10^{-3}$ | mRNA-intron | 2378 |
| AT3G19040 | protein coding gene | | -29.8 | $1.41 \times 10^{-16}$ | $3.05 \times 10^{-15}$ | mRNA-exon | 2331 |
| AT3G04250 | protein coding gene | | -29.9 | $1.45 \times 10^{-8}$ | $1.15 \times 10^{-7}$ | mRNA-TTS | 401 |
| AT4G32175 | protein coding gene | | -30.0 | $7.42 \times 10^{-13}$ | $1.03 \times 10^{-11}$ | mRNA-TTS | 2817 |
| AT5G04025 | lncRNA | | -30.0 | $3.44 \times 10^{-5}$ | $1.62 \times 10^{-4}$ | Intergenic | -1776 |
| AT5G07335 | snRNA | | -30.1 | $9.24 \times 10^{-17}$ | $2.04 \times 10^{-15}$ | mRNA-exon | 1319 |
| AT1G56500 | protein coding gene | | -30.1 | $4.21 \times 10^{-4}$ | $1.57 \times 10^{-3}$ | mRNA-intron | 2107 |
| AT1G04230 | protein coding gene | | -30.2 | $1.77 \times 10^{-28}$ | $1.14 \times 10^{-26}$ | mRNA-exon | 733 |
| AT4G20520 | protein coding gene | | -30.2 | $6.69 \times 10^{-27}$ | $3.83 \times 10^{-25}$ | | -1261 |
| AT5G16070 | protein coding gene | | -30.3 | $2.56 \times 10^{-7}$ | $1.71 \times 10^{-6}$ | mRNA-intron | 1428 |
| AT3G18390 | protein coding gene | *EMB1865* | -30.3 | $6.01 \times 10^{-10}$ | $5.69 \times 10^{-9}$ | mRNA-exon | 2880 |
| AT2G40090 | protein coding gene | *ATH9* | -30.4 | $1.82 \times 10^{-6}$ | $1.07 \times 10^{-5}$ | mRNA-exon | 1335 |
| AT5G22830 | protein coding gene | *MGT1* | -30.4 | $7.18 \times 10^{-9}$ | $5.89 \times 10^{-8}$ | mRNA-intron | 1554 |
| AT2G31540 | protein coding gene | | -30.4 | $3.02 \times 10^{-9}$ | $2.59 \times 10^{-8}$ | mRNA-TTS | 2114 |
| AT4G27760 | protein coding gene | *FEY* | -30.5 | $3.06 \times 10^{-12}$ | $3.96 \times 10^{-11}$ | mRNA-exon | 892 |
| AT1G68720 | protein coding gene | *TADA* | -30.6 | $1.07 \times 10^{-6}$ | $6.53 \times 10^{-6}$ | mRNA-exon | 1459 |
| AT1G08270 | protein coding gene | | -30.7 | $5.17 \times 10^{-8}$ | $3.77 \times 10^{-7}$ | mRNA-intron | 2386 |
| AT2G04135 | transposable element gene | | -30.7 | $5.70 \times 10^{-35}$ | $6.36 \times 10^{-33}$ | mRNA-promoter-TSS | -613 |
| AT4G08065 | lncRNA | | -30.7 | $1.48 \times 10^{-15}$ | $2.86 \times 10^{-14}$ | Intergenic | -1791 |
| AT4G15885 | protein coding gene | | -30.9 | $2.34 \times 10^{-4}$ | $9.26 \times 10^{-4}$ | mRNA-exon | 2127 |
| AT1G67140 | protein coding gene | *SWEETIE* | -30.9 | $2.74 \times 10^{-6}$ | $1.57 \times 10^{-5}$ | mRNA-intron | 3972 |
| AT2G33380 | protein coding gene | *RD2* | -30.9 | $2.24 \times 10^{-12}$ | $2.96 \times 10^{-11}$ | mRNA-promoter-TSS | -596 |
| AT2G15720 | transposable element gene | | -31.0 | $3.61 \times 10^{-18}$ | $9.08 \times 10^{-17}$ | mRNA-promoter-TSS | -519 |

| | | | | | | | | | | | | |
|---|---|---|---|---|---|---|---|---|---|---|---|---|
| AT2G15720 | transposable element gene | | -31.0 | $3.61 \times 10^{-18}$ | $9.08 \times 10^{-17}$ | mRNA-promoter-TSS | -519 | -27.9 | $9.59 \times 10^{-19}$ | $2.41 \times 10^{-16}$ | mRNA-promoter-TSS | -519 |
| AT4G04370 | protein coding gene | | -31.0 | $3.05 \times 10^{-9}$ | $2.62 \times 10^{-8}$ | mRNA-exon | 970 | | | | | |
| AT2G15370 | protein coding gene | *FUT5* | -31.1 | $1.44 \times 10^{-18}$ | $3.77 \times 10^{-17}$ | mRNA-exon | 985 | | | | | |
| AT2G04135 | transposable element gene | | -31.2 | $3.76 \times 10^{-24}$ | $1.66 \times 10^{-22}$ | mRNA-promoter-TSS | -413 | | | | | |
| AT5G04025 | lncRNA | | -31.2 | $2.68 \times 10^{-6}$ | $1.54 \times 10^{-5}$ | Intergenic | -1676 | -35.1 | $4.24 \times 10^{-5}$ | $1.02 \times 10^{-3}$ | Intergenic | -1776 |
| AT1G58235 | protein coding gene | | -31.2 | $2.96 \times 10^{-12}$ | $3.85 \times 10^{-11}$ | mRNA-intron | 3915 | | | | | |
| AT5G22820 | protein coding gene | | -31.2 | $8.22 \times 10^{-7}$ | $5.10 \times 10^{-6}$ | mRNA-TTS | 2686 | | | | | |
| AT1G03820 | protein coding gene | | -31.3 | $2.18 \times 10^{-5}$ | $1.07 \times 10^{-4}$ | | -2835 | | | | | |
| AT1G49210 | protein coding gene | | -31.3 | $4.94 \times 10^{-8}$ | $3.61 \times 10^{-7}$ | mRNA-promoter-TSS | -738 | | | | | |
| AT4G08033 | transposable element gene | | -31.3 | $6.26 \times 10^{-35}$ | $6.93 \times 10^{-33}$ | Intergenic | 3943 | | | | | |
| AT2G07890 | lncRNA | | -31.4 | $6.18 \times 10^{-11}$ | $6.75 \times 10^{-10}$ | lnc RNA-TTS | 1263 | | | | | |
| AT2G23900 | protein coding gene | | -31.4 | $5.87 \times 10^{-12}$ | $7.33 \times 10^{-11}$ | | -1564 | | | | | |
| AT4G08033 | transposable element gene | | -31.4 | $1.87 \times 10^{-9}$ | $1.66 \times 10^{-8}$ | Intergenic | 4043 | | | | | |
| AT1G09077 | lncRNA | | -31.4 | $4.14 \times 10^{-6}$ | $2.29 \times 10^{-5}$ | mRNA-promoter-TSS | 645 | | | | | |
| AT1G06470 | protein coding gene | | -31.4 | $2.88 \times 10^{-10}$ | $2.86 \times 10^{-9}$ | mRNA-intron | 1722 | | | | | |
| AT4G16390 | protein coding gene | *SVR7* | -31.5 | $1.31 \times 10^{-13}$ | $2.01 \times 10^{-12}$ | mRNA-exon | 982 | | | | | |
| AT5G15170 | protein coding gene | *TDP1* | -31.5 | $1.37 \times 10^{-10}$ | $1.42 \times 10^{-9}$ | mRNA-intron | 1951 | | | | | |
| AT3G61490 | protein coding gene | | -31.6 | $5.02 \times 10^{-7}$ | $3.22 \times 10^{-6}$ | | -2810 | | | | | |
| AT2G38090 | protein coding gene | | -31.6 | $7.15 \times 10^{-5}$ | $3.16 \times 10^{-4}$ | Intergenic | -4732 | | | | | |
| AT2G01045 | protein coding gene | | -31.6 | $5.03 \times 10^{-33}$ | $4.80 \times 10^{-31}$ | Intergenic | -2383 | | | | | |
| AT4G09285 | lncRNA | | -31.7 | $1.44 \times 10^{-10}$ | $1.49 \times 10^{-9}$ | | -2654 | | | | | |
| AT5G42410 | protein coding gene | | -31.7 | $5.47 \times 10^{-8}$ | $3.97 \times 10^{-7}$ | Intergenic | -1399 | | | | | |
| AT3G26612 | other RNA | | -31.7 | $1.85 \times 10^{-7}$ | $1.26 \times 10^{-6}$ | ncRNA-exon | 1068 | | | | | |
| AT4G08033 | transposable element gene | | -31.7 | $1.42 \times 10^{-14}$ | $2.44 \times 10^{-13}$ | Intergenic | 4243 | | | | | |
| AT3G63460 | protein coding gene | | -31.8 | $6.41 \times 10^{-6}$ | $3.44 \times 10^{-5}$ | mRNA-intron | 1893 | | | | | |
| AT5G08250 | protein coding gene | | -31.9 | $2.03 \times 10^{-8}$ | $1.58 \times 10^{-7}$ | mRNA-promoter-TSS | -481 | | | | | |
| AT3G18100 | protein coding gene | *MYB4R1* | -31.9 | $1.97 \times 10^{-8}$ | $1.53 \times 10^{-7}$ | | -1173 | | | | | |
| AT1G56160 | protein coding gene | *MYB72* | -32.0 | $1.26 \times 10^{-6}$ | $7.60 \times 10^{-6}$ | Intergenic | -1333 | | | | | |
| AT3G22670 | protein coding gene | | -32.0 | $4.74 \times 10^{-15}$ | $8.68 \times 10^{-14}$ | | -2299 | | | | | |
| AT1G02850 | protein coding gene | *BGLU11* | -32.1 | $6.27 \times 10^{-5}$ | $2.80 \times 10^{-4}$ | | -1523 | | | | | |
| AT5G60730 | protein coding gene | | -32.1 | $4.91 \times 10^{-8}$ | $3.59 \times 10^{-7}$ | mRNA-TTS | 2365 | | | | | |
| AT3G14580 | protein coding gene | | -32.2 | $2.88 \times 10^{-10}$ | $2.87 \times 10^{-9}$ | | -2802 | | | | | |
| AT5G51230 | protein coding gene | *EMF2* | -32.3 | $2.07 \times 10^{-4}$ | $8.28 \times 10^{-4}$ | mRNA-intron | 3217 | | | | | |
| AT5G18650 | protein coding gene | | -32.4 | $9.46 \times 10^{-26}$ | $4.83 \times 10^{-24}$ | mRNA-TTS | 2061 | | | | | |
| AT4G15475 | protein coding gene | | -32.5 | $1.10 \times 10^{-12}$ | $1.49 \times 10^{-11}$ | mRNA-promoter-TSS | -361 | | | | | |
| AT4G08535 | protein coding gene | | -32.5 | $5.49 \times 10^{-7}$ | $3.50 \times 10^{-6}$ | mRNA-exon | 1605 | | | | | |
| AT3G29380 | protein coding gene | *pBRP2* | -32.7 | $1.44 \times 10^{-8}$ | $1.14 \times 10^{-7}$ | mRNA-exon | 273 | | | | | |

| | | | | | | | | | | | | |
|---|---|---|---|---|---|---|---|---|---|---|---|---|
| AT1G79920 | protein coding gene | *HSP7-15* | -32.7 | $4.51 \times 10^{-9}$ | $3.80 \times 10^{-8}$ | mRNA-exon | 2677 | | | | | |
| AT3G28360 | protein coding gene | *ABCB16* | -32.7 | $6.55 \times 10^{-11}$ | $7.13 \times 10^{-10}$ | Intergenic | -1390 | | | | | |
| AT1G33785 | pseudogene | | -32.7 | $6.98 \times 10^{-12}$ | $8.63 \times 10^{-11}$ | Intergenic | 3405 | | | | | |
| AT2G26790 | protein coding gene | | -32.7 | $3.38 \times 10^{-10}$ | $3.32 \times 10^{-9}$ | mRNA-intron | 8223 | | | | | |
| AT3G11910 | protein coding gene | *UBP13* | -32.7 | $4.67 \times 10^{-9}$ | $3.93 \times 10^{-8}$ | mRNA-intron | 3538 | | | | | |
| AT4G12640 | protein coding gene | | -32.7 | $7.10 \times 10^{-6}$ | $3.78 \times 10^{-5}$ | mRNA-intron | 2853 | | | | | |
| AT1G02090 | protein coding gene | *FUS5* | -32.8 | $2.17 \times 10^{-4}$ | $8.63 \times 10^{-4}$ | mRNA-intron | 5857 | | | | | |
| AT5G23110 | protein coding gene | | -32.8 | $4.61 \times 10^{-13}$ | $6.56 \times 10^{-12}$ | mRNA-exon | 4851 | | | | | |
| AT2G33580 | protein coding gene | *LYK5* | -32.8 | $1.06 \times 10^{-7}$ | $7.42 \times 10^{-7}$ | mRNA-promoter-TSS | -897 | | | | | |
| AT5G52540 | protein coding gene | | -32.9 | $4.43 \times 10^{-10}$ | $4.27 \times 10^{-9}$ | mRNA-TTS | 929 | | | | | |
| AT5G00370 | novel transcribed region | | -32.9 | $1.18 \times 10^{-6}$ | $7.20 \times 10^{-6}$ | | -1045 | | | | | |
| AT4G11220 | protein coding gene | *BTI2* | -32.9 | $2.00 \times 10^{-23}$ | $8.20 \times 10^{-22}$ | | -2615 | | | | | |
| AT1G19860 | protein coding gene | | -33.0 | $1.12 \times 10^{-7}$ | $7.83 \times 10^{-7}$ | mRNA-exon | 2101 | | | | | |
| AT3G42160 | protein coding gene | | -33.0 | $1.30 \times 10^{-9}$ | $1.18 \times 10^{-8}$ | mRNA-TTS | 1168 | | | | | |
| AT3G12380 | protein coding gene | *ARP5* | -33.0 | $6.56 \times 10^{-8}$ | $4.70 \times 10^{-7}$ | mRNA-exon | 1421 | | | | | |
| AT4G39680 | protein coding gene | | -33.1 | $1.39 \times 10^{-10}$ | $1.45 \times 10^{-9}$ | | -2443 | | | | | |
| AT2G25680 | protein coding gene | *MOT1* | -33.1 | $5.73 \times 10^{-8}$ | $4.13 \times 10^{-7}$ | Intergenic | -1347 | | | | | |
| AT5G20890 | protein coding gene | | -33.2 | $5.23 \times 10^{-8}$ | $3.81 \times 10^{-7}$ | mRNA-exon | 1118 | | | | | |
| AT1G78420 | protein coding gene | | -33.2 | $2.24 \times 10^{-10}$ | $2.26 \times 10^{-9}$ | mRNA-exon | 1618 | | | | | |
| AT3G27260 | protein coding gene | | -33.2 | $3.98 \times 10^{-13}$ | $5.71 \times 10^{-12}$ | mRNA-exon | 1006 | | | | | |
| AT5G03740 | protein coding gene | *HD2C* | -33.2 | $1.82 \times 10^{-7}$ | $1.24 \times 10^{-6}$ | mRNA-exon | 992 | | | | | |
| AT3G01045 | anti sense lncRNA | | -33.4 | $3.22 \times 10^{-6}$ | $1.82 \times 10^{-5}$ | | -1868 | | | | | |
| AT1G01920 | protein coding gene | | -33.5 | $1.65 \times 10^{-7}$ | $1.13 \times 10^{-6}$ | mRNA-promoter-TSS | 864 | | | | | |
| AT1G52905 | protein coding gene | | -33.5 | $1.08 \times 10^{-20}$ | $3.54 \times 10^{-19}$ | mRNA-promoter-TSS | -196 | | | | | |
| AT4G31760 | protein coding gene | | -33.5 | $6.50 \times 10^{-10}$ | $6.12 \times 10^{-9}$ | mRNA-promoter-TSS | -927 | | | | | |
| AT3G58810 | protein coding gene | *MTPA2* | -33.5 | $8.18 \times 10^{-6}$ | $4.30 \times 10^{-5}$ | mRNA-promoter-TSS | -215 | | | | | |
| AT5G41730 | protein coding gene | | -33.5 | $2.62 \times 10^{-17}$ | $6.11 \times 10^{-16}$ | lnc RNA-promoter-TSS | 467 | -30.4 | $1.79 \times 10^{-12}$ | $1.77 \times 10^{-10}$ | lnc_RNA-promoter-TSS | 367 |
| AT1G24320 | protein coding gene | | -33.6 | $2.03 \times 10^{-20}$ | $6.42 \times 10^{-19}$ | mRNA-exon | 2774 | | | | | |
| AT4G25340 | protein coding gene | *FKBP53* | -33.6 | $4.15 \times 10^{-10}$ | $4.02 \times 10^{-9}$ | mRNA-exon | 748 | | | | | |
| AT2G41700 | protein coding gene | *ABCA1* | -33.6 | $1.07 \times 10^{-7}$ | $7.51 \times 10^{-7}$ | mRNA-intron | 2091 | | | | | |
| AT1G04080 | protein coding gene | *PRP39a* | -33.7 | $7.80 \times 10^{-8}$ | $5.55 \times 10^{-7}$ | mRNA-exon | 1998 | | | | | |
| AT5G52530 | protein coding gene | | -33.7 | $1.16 \times 10^{-13}$ | $1.78 \times 10^{-12}$ | mRNA-exon | 1711 | | | | | |
| AT5G61780 | protein coding gene | *Tudor2* | -33.7 | $3.44 \times 10^{-10}$ | $3.37 \times 10^{-9}$ | mRNA-exon | 1348 | | | | | |
| AT4G17330 | protein coding gene | *G2484-1* | -33.9 | $2.03 \times 10^{-7}$ | $1.37 \times 10^{-6}$ | mRNA-exon | 2427 | | | | | |
| AT2G04135 | transposable element gene | | -33.9 | $5.42 \times 10^{-38}$ | $7.69 \times 10^{-36}$ | mRNA-promoter-TSS | -513 | | | | | |
| AT2G15270 | protein coding gene | | -34.0 | $8.80 \times 10^{-7}$ | $5.43 \times 10^{-6}$ | transcript-TTS | 973 | | | | | |
| AT1G12700 | protein coding gene | | -34.2 | $1.02 \times 10^{-6}$ | $6.25 \times 10^{-6}$ | mRNA-exon | 1176 | | | | | |

| | | | | | | | | | | | | |
|---|---|---|---|---|---|---|---|---|---|---|---|---|
| AT4G36940 | protein coding gene | *NAPRT1* | -34.2 | $1.35 \times 10^{-11}$ | $1.62 \times 10^{-10}$ | mRNA-exon | 1596 | | | | | |
| AT1G56580 | protein coding gene | *SVB* | -34.3 | $9.64 \times 10^{-10}$ | $8.88 \times 10^{-9}$ | mRNA-TTS | 1575 | | | | | |
| AT1G42705 | transposable element gene | | -34.3 | $4.98 \times 10^{-13}$ | $7.06 \times 10^{-12}$ | mRNA-promoter-TSS | 10 | | | | | |
| AT3G15120 | protein coding gene | | -34.4 | $3.67 \times 10^{-8}$ | $2.75 \times 10^{-7}$ | mRNA-exon | 1366 | | | | | |
| AT1G47560 | protein coding gene | *SEC3B* | -34.5 | $4.42 \times 10^{-12}$ | $5.62 \times 10^{-11}$ | | -4084 | | | | | |
| AT1G55175 | protein coding gene | | -34.5 | $4.75 \times 10^{-14}$ | $7.69 \times 10^{-13}$ | mRNA-TTS | 1886 | | | | | |
| AT2G22125 | protein coding gene | *CSI1* | -34.5 | $1.44 \times 10^{-13}$ | $2.18 \times 10^{-12}$ | mRNA-intron | 4592 | | | | | |
| AT4G05485 | lncRNA | | -34.5 | $8.44 \times 10^{-12}$ | $1.03 \times 10^{-10}$ | Intergenic | -1808 | | | | | |
| AT3G53365 | other RNA | | -34.6 | $8.89 \times 10^{-11}$ | $9.53 \times 10^{-10}$ | ncRNA-promoter-TSS | -304 | | | | | |
| AT4G04425 | anti sense lncRNA | | -34.6 | $2.79 \times 10^{-6}$ | $1.59 \times 10^{-5}$ | mRNA-intron | 1451 | | | | | |
| AT2G42640 | protein coding gene | | -34.6 | $2.70 \times 10^{-11}$ | $3.12 \times 10^{-10}$ | mRNA-exon | 1172 | | | | | |
| AT1G63340 | protein coding gene | | -34.7 | $1.57 \times 10^{-20}$ | $5.04 \times 10^{-19}$ | | -1032 | | | | | |
| AT3G26085 | protein coding gene | | -34.8 | $4.38 \times 10^{-16}$ | $8.93 \times 10^{-15}$ | mRNA-TTS | 782 | | | | | |
| AT1G15240 | protein coding gene | | -34.9 | $6.98 \times 10^{-20}$ | $2.08 \times 10^{-18}$ | mRNA-exon | 2501 | | | | | |
| AT1G08043 | anti sense lncRNA | | -34.9 | $7.05 \times 10^{-7}$ | $4.43 \times 10^{-6}$ | lnc RNA-promoter-TSS | -610 | | | | | |
| AT4G30090 | protein coding gene | | -34.9 | $3.54 \times 10^{-13}$ | $5.12 \times 10^{-12}$ | mRNA-exon | 2534 | | | | | |
| AT2G31810 | protein coding gene | | -35.0 | $9.27 \times 10^{-14}$ | $1.44 \times 10^{-12}$ | mRNA-intron | 1029 | | | | | |
| AT1G26610 | protein coding gene | | -35.4 | $6.38 \times 10^{-17}$ | $1.42 \times 10^{-15}$ | mRNA-exon | 3124 | | | | | |
| AT5G04238 | protein coding gene | | -35.4 | $1.77 \times 10^{-16}$ | $3.78 \times 10^{-15}$ | | -3040 | | | | | |
| AT5G22000 | protein coding gene | *RHF2A* | -35.5 | $9.35 \times 10^{-12}$ | $1.14 \times 10^{-10}$ | mRNA-intron | 4612 | | | | | |
| AT2G05965 | lncRNA | | -35.5 | $5.72 \times 10^{-7}$ | $3.63 \times 10^{-6}$ | lnc RNA-promoter-TSS | -344 | | | | | |
| AT3G25480 | protein coding gene | | -35.5 | $3.25 \times 10^{-10}$ | $3.20 \times 10^{-9}$ | mRNA-TTS | 1797 | | | | | |
| AT1G07990 | protein coding gene | | -35.8 | $5.95 \times 10^{-8}$ | $4.28 \times 10^{-7}$ | mRNA-intron | 2298 | | | | | |
| AT5G35600 | protein coding gene | *HDA7* | -35.8 | $1.57 \times 10^{-23}$ | $6.49 \times 10^{-22}$ | Intergenic | -2039 | | | | | |
| AT1G09195 | protein coding gene | | -35.9 | $1.81 \times 10^{-7}$ | $1.23 \times 10^{-6}$ | mRNA-intron | 1106 | | | | | |
| AT5G51690 | protein coding gene | *ACS12* | -35.9 | $2.00 \times 10^{-6}$ | $1.17 \times 10^{-5}$ | mRNA-TTS | 1499 | | | | | |
| AT1G31110 | pre tRNA | | -36.1 | $1.45 \times 10^{-9}$ | $1.30 \times 10^{-8}$ | mRNA-exon | 1738 | | | | | |
| AT3G08405 | anti sense lncRNA | | -36.2 | $3.53 \times 10^{-13}$ | $5.11 \times 10^{-12}$ | | -1620 | | | | | |
| AT2G39260 | protein coding gene | | -36.2 | $9.52 \times 10^{-8}$ | $6.72 \times 10^{-7}$ | mRNA-intron | 4161 | | | | | |
| AT3G05190 | protein coding gene | | -36.2 | $1.96 \times 10^{-8}$ | $1.53 \times 10^{-7}$ | mRNA-intron | 2095 | | | | | |
| AT4G14150 | protein coding gene | *PAKRP1* | -36.2 | $1.91 \times 10^{-5}$ | $9.43 \times 10^{-5}$ | mRNA-intron | 1963 | | | | | |
| AT1G52905 | protein coding gene | | -36.2 | $7.83 \times 10^{-34}$ | $8.13 \times 10^{-32}$ | mRNA-promoter-TSS | 4 | | | | | |
| AT1G52905 | protein coding gene | | -36.2 | $7.83 \times 10^{-34}$ | $8.13 \times 10^{-32}$ | mRNA-promoter-TSS | 4 | -27.3 | $6.00 \times 10^{-9}$ | $3.21 \times 10^{-7}$ | mRNA-promoter-TSS | 4 |
| AT4G24652 | pseudogene | | -36.3 | $1.18 \times 10^{-14}$ | $2.03 \times 10^{-13}$ | transcript-promoter-TSS | 1 | | | | | |
| AT2G30105 | protein coding gene | | -36.4 | $6.39 \times 10^{-23}$ | $2.51 \times 10^{-21}$ | mRNA-TTS | 2133 | | | | | |
| AT1G80410 | protein coding gene | *NAA15* | -36.5 | $4.01 \times 10^{-8}$ | $2.98 \times 10^{-7}$ | mRNA-exon | 1434 | | | | | |
| AT2G02180 | protein coding gene | *TOM3* | -36.5 | $1.57 \times 10^{-15}$ | $3.01 \times 10^{-14}$ | mRNA-intron | 1756 | | | | | |

| | | | | | | | |
|---|---|---|---|---|---|---|---|
| AT2G29190 | protein coding gene | *PUM2* | -36.6 | $3.48 \times 10^{-17}$ | $7.98 \times 10^{-16}$ | mRNA-exon | 1078 |
| AT3G03895 | anti sense lncRNA | | -36.7 | $4.64 \times 10^{-9}$ | $3.91 \times 10^{-8}$ | | -2049 |
| AT1G50490 | protein coding gene | *UBC2* | -36.7 | $3.95 \times 10^{-5}$ | $1.84 \times 10^{-4}$ | mRNA-TTS | 2306 |
| AT5G22765 | protein coding gene | | -36.7 | $7.54 \times 10^{-11}$ | $8.13 \times 10^{-10}$ | mRNA-exon | 3232 |
| AT3G25545 | protein coding gene | | -36.9 | $1.62 \times 10^{-40}$ | $2.50 \times 10^{-38}$ | mRNA-promoter-TSS | -994 |
| AT3G56560 | protein coding gene | *NAC65* | -36.9 | $1.86 \times 10^{-25}$ | $9.29 \times 10^{-24}$ | mRNA-intron | 647 |
| AT1G59990 | protein coding gene | *EMB318* | -37.0 | $5.98 \times 10^{-6}$ | $3.24 \times 10^{-5}$ | mRNA-exon | 1409 |
| AT5G37145 | transposable element gene | | -37.2 | $1.81 \times 10^{-16}$ | $3.88 \times 10^{-15}$ | mRNA-exon | 248 |
| AT1G79600 | protein coding gene | | -37.3 | $5.79 \times 10^{-9}$ | $4.80 \times 10^{-8}$ | | -1525 |
| AT5G13230 | protein coding gene | | -37.3 | $3.97 \times 10^{-10}$ | $3.86 \times 10^{-9}$ | mRNA-exon | 1239 |
| AT3G57300 | protein coding gene | *INO8* | -37.4 | $2.93 \times 10^{-9}$ | $2.52 \times 10^{-8}$ | mRNA-intron | 963 |
| AT4G12460 | protein coding gene | | -37.4 | $1.47 \times 10^{-14}$ | $2.50 \times 10^{-13}$ | mRNA-intron | 710 |
| AT1G12920 | protein coding gene | *ERF1-2* | -37.5 | $4.55 \times 10^{-7}$ | $2.94 \times 10^{-6}$ | | -2458 |
| AT4G08535 | protein coding gene | | -37.6 | $3.03 \times 10^{-15}$ | $5.65 \times 10^{-14}$ | mRNA-intron | 1505 |
| AT2G01045 | protein coding gene | | -37.6 | $6.88 \times 10^{-33}$ | $6.53 \times 10^{-31}$ | Intergenic | -2083 |
| AT5G04090 | protein coding gene | | -37.7 | $2.53 \times 10^{-25}$ | $1.25 \times 10^{-23}$ | | -1669 |
| AT3G26612 | other RNA | | -37.7 | $2.42 \times 10^{-9}$ | $2.11 \times 10^{-8}$ | ncRNA-exon | 568 |
| AT2G31970 | protein coding gene | *RAD5* | -37.8 | $2.79 \times 10^{-9}$ | $2.41 \times 10^{-8}$ | mRNA-intron | 3920 |
| AT1G74910 | protein coding gene | | -37.9 | $6.06 \times 10^{-14}$ | $9.65 \times 10^{-13}$ | | -1080 |
| AT3G11120 | protein coding gene | | -37.9 | $1.98 \times 10^{-6}$ | $1.16 \times 10^{-5}$ | mRNA-exon | 3431 |
| AT4G13750 | protein coding gene | *NOV* | -38.0 | $8.56 \times 10^{-12}$ | $1.05 \times 10^{-10}$ | mRNA-exon | 1287 |
| AT3G60920 | protein coding gene | | -38.0 | $2.01 \times 10^{-10}$ | $2.05 \times 10^{-9}$ | mRNA-exon | 680 |
| AT5G46920 | protein coding gene | | -38.1 | $2.54 \times 10^{-15}$ | $4.78 \times 10^{-14}$ | mRNA-exon | 1779 |
| AT1G73460 | protein coding gene | | -38.1 | $7.94 \times 10^{-14}$ | $1.25 \times 10^{-12}$ | | -1984 |
| AT1G08773 | lncRNA | | -38.1 | $7.39 \times 10^{-22}$ | $2.71 \times 10^{-20}$ | lnc RNA-TTS | 224 |
| AT2G01045 | protein coding gene | | -38.4 | $8.80 \times 10^{-57}$ | $3.11 \times 10^{-54}$ | Intergenic | -2283 |
| AT5G36270 | pseudogene | | -38.4 | $1.46 \times 10^{-14}$ | $2.50 \times 10^{-13}$ | Intergenic | -1661 |
| AT4G08033 | transposable element gene | | -38.5 | $7.22 \times 10^{-12}$ | $8.91 \times 10^{-11}$ | Intergenic | 4143 |
| AT5G12370 | protein coding gene | *SEC1* | -38.6 | $9.38 \times 10^{-14}$ | $1.46 \times 10^{-12}$ | mRNA-intron | 1924 |
| AT1G07840 | protein coding gene | | -38.8 | $1.51 \times 10^{-10}$ | $1.57 \times 10^{-9}$ | mRNA-intron | 987 |
| AT2G01045 | protein coding gene | | -38.9 | $1.08 \times 10^{-32}$ | $9.97 \times 10^{-31}$ | Intergenic | -2183 |
| AT1G50250 | protein coding gene | *FTSH1* | -38.9 | $1.29 \times 10^{-8}$ | $1.03 \times 10^{-7}$ | mRNA-intron | 4404 |
| AT2G19802 | protein coding gene | | -38.9 | $2.78 \times 10^{-16}$ | $5.80 \times 10^{-15}$ | Intergenic | 2666 |
| AT3G04585 | lncRNA | | -38.9 | $9.25 \times 10^{-11}$ | $9.87 \times 10^{-10}$ | | -2169 |
| AT3G06280 | protein coding gene | | -39.0 | $1.36 \times 10^{-6}$ | $8.18 \times 10^{-6}$ | | -2500 |
| AT2G04945 | lncRNA | | -39.1 | $2.64 \times 10^{-29}$ | $1.87 \times 10^{-27}$ | lnc RNA-promoter-TSS | -243 |
| AT3G18485 | protein coding gene | *ILR2* | -39.2 | $8.55 \times 10^{-17}$ | $1.89 \times 10^{-15}$ | mRNA-TTS | 2227 |

| | | | | | | | |
|---|---|---|---|---|---|---|---|
| AT3G01045 | anti sense lncRNA | | -39.3 | $1.09 \times 10^{-17}$ | $2.63 \times 10^{-16}$ | | -1068 |
| AT5G13010 | protein coding gene | *EMB311* | -39.3 | $1.08 \times 10^{-13}$ | $1.67 \times 10^{-12}$ | mRNA-exon | 5151 |
| AT1G16240 | protein coding gene | *SYP51* | -39.3 | $1.06 \times 10^{-6}$ | $6.49 \times 10^{-6}$ | mRNA-exon | 963 |
| AT5G61060 | protein coding gene | *HDA5* | -39.4 | $3.34 \times 10^{-14}$ | $5.51 \times 10^{-13}$ | mRNA-exon | 1440 |
| AT4G08033 | transposable element gene | | -39.4 | $1.16 \times 10^{-30}$ | $9.29 \times 10^{-29}$ | Intergenic | 4343 |
| AT3G29760 | protein coding gene | | -39.4 | $8.26 \times 10^{-9}$ | $6.72 \times 10^{-8}$ | mRNA-exon | 982 |
| AT5G67240 | protein coding gene | *SDN3* | -39.5 | $7.91 \times 10^{-18}$ | $1.93 \times 10^{-16}$ | mRNA-exon | 1355 |
| AT5G53895 | protein coding gene | | -39.6 | $1.73 \times 10^{-17}$ | $4.11 \times 10^{-16}$ | mRNA-promoter-TSS | -741 |
| AT3G18960 | protein coding gene | | -39.6 | $8.02 \times 10^{-27}$ | $4.56 \times 10^{-25}$ | mRNA-TTS | 1213 |
| AT1G18265 | protein coding gene | | -39.7 | $2.42 \times 10^{-12}$ | $3.18 \times 10^{-11}$ | | -1903 |
| AT4G23340 | protein coding gene | | -39.9 | $1.94 \times 10^{-14}$ | $3.28 \times 10^{-13}$ | mRNA-TTS | 263 |
| AT4G00155 | protein coding gene | | -40.1 | $2.80 \times 10^{-19}$ | $7.92 \times 10^{-18}$ | mRNA-promoter-TSS | -284 |
| AT1G54475 | protein coding gene | | -40.1 | $1.28 \times 10^{-16}$ | $2.77 \times 10^{-15}$ | mRNA-promoter-TSS | -46 |
| AT5G26250 | protein coding gene | | -40.5 | $4.19 \times 10^{-8}$ | $3.09 \times 10^{-7}$ | | -2907 |
| AT5G16630 | protein coding gene | *RAD4* | -40.7 | $1.92 \times 10^{-14}$ | $3.26 \times 10^{-13}$ | mRNA-exon | 1433 |
| AT3G10100 | transposable element gene | | -40.8 | $7.53 \times 10^{-44}$ | $1.36 \times 10^{-41}$ | mRNA-exon | 1873 |
| AT4G33200 | protein coding gene | | -41.1 | $1.27 \times 10^{-17}$ | $3.04 \times 10^{-16}$ | | -2770 |
| AT1G72440 | protein coding gene | *EDA25* | -41.1 | $3.92 \times 10^{-8}$ | $2.91 \times 10^{-7}$ | mRNA-exon | 2084 |
| AT5G10060 | protein coding gene | | -41.5 | $1.84 \times 10^{-8}$ | $1.44 \times 10^{-7}$ | mRNA-intron | 1449 |
| AT4G38005 | protein coding gene | | -41.5 | $3.54 \times 10^{-8}$ | $2.65 \times 10^{-7}$ | mRNA-promoter-TSS | -847 |
| AT3G55677 | protein coding gene | | -41.5 | $8.34 \times 10^{-27}$ | $4.73 \times 10^{-25}$ | mRNA-TTS | 1068 |
| AT1G78922 | protein coding gene | | -41.5 | $3.66 \times 10^{-18}$ | $9.18 \times 10^{-17}$ | mRNA-exon | 2864 |
| AT3G54360 | protein coding gene | | -41.6 | $9.60 \times 10^{-19}$ | $2.55 \times 10^{-17}$ | mRNA-intron | 2462 |
| AT1G06397 | small nucleolar RNA | | -41.9 | $3.56 \times 10^{-9}$ | $3.02 \times 10^{-8}$ | | -1330 |
| AT2G04215 | lncRNA | | -42.0 | $3.81 \times 10^{-25}$ | $1.85 \times 10^{-23}$ | lnc RNA-promoter-TSS | -749 |
| AT1G76320 | protein coding gene | *FRS4* | -42.1 | $2.89 \times 10^{-17}$ | $6.72 \times 10^{-16}$ | mRNA-exon | 1028 |
| AT4G23100 | protein coding gene | *GSH1* | -42.4 | $2.96 \times 10^{-16}$ | $6.17 \times 10^{-15}$ | mRNA-exon | 1848 |
| AT3G08520 | protein coding gene | | -42.4 | $5.55 \times 10^{-18}$ | $1.37 \times 10^{-16}$ | mRNA-exon | 2750 |
| AT2G02100 | protein coding gene | *LCR69* | -42.6 | $3.34 \times 10^{-20}$ | $1.03 \times 10^{-18}$ | | -2328 |
| AT1G48725 | protein coding gene | | -42.7 | $7.66 \times 10^{-15}$ | $1.36 \times 10^{-13}$ | mRNA-TTS | 177 |
| AT3G63140 | protein coding gene | *CSP41A* | -42.7 | $2.93 \times 10^{-7}$ | $1.95 \times 10^{-6}$ | | -1762 |
| AT2G13540 | protein coding gene | *ABH1* | -42.7 | $4.67 \times 10^{-18}$ | $1.16 \times 10^{-16}$ | mRNA-exon | 1267 |
| AT4G27090 | protein coding gene | | -42.7 | $8.88 \times 10^{-20}$ | $2.63 \times 10^{-18}$ | mRNA-intron | 2801 |
| AT1G13560 | protein coding gene | *AAPT1* | -42.9 | $1.39 \times 10^{-19}$ | $4.03 \times 10^{-18}$ | mRNA-intron | 707 |
| AT5G63920 | protein coding gene | *TOP3A* | -42.9 | $2.55 \times 10^{-7}$ | $1.71 \times 10^{-6}$ | mRNA-intron | 2432 |
| AT1G28350 | protein coding gene | | -43.1 | $6.00 \times 10^{-18}$ | $1.47 \times 10^{-16}$ | mRNA-intron | 2077 |
| AT3G27270 | protein coding gene | | -43.1 | $5.79 \times 10^{-9}$ | $4.80 \times 10^{-8}$ | | -1931 |

| | | | | | | | | | | | | |
|---|---|---|---|---|---|---|---|---|---|---|---|---|
| AT2G16670 | transposable element gene | | -43.2 | $3.25 \times 10^{-32}$ | $2.87 \times 10^{-30}$ | mRNA-exon | 3368 | | | | | |
| AT2G16670 | transposable element gene | | -43.2 | $3.25 \times 10^{-32}$ | $2.87 \times 10^{-30}$ | mRNA-exon | 3368 | -30.8 | $4.00 \times 10^{-13}$ | $4.28 \times 10^{-11}$ | mRNA-exon | 3368 |
| AT2G16905 | pseudogene | | -43.2 | $3.25 \times 10^{-32}$ | $2.87 \times 10^{-30}$ | mRNA-exon | 3368 | -29.0 | $9.54 \times 10^{-10}$ | $5.99 \times 10^{-8}$ | transcript-exon | 1034 |
| AT5G28200 | transposable element gene | | -43.2 | $1.30 \times 10^{-21}$ | $4.62 \times 10^{-20}$ | Intergenic | -2236 | | | | | |
| AT5G22010 | protein coding gene | *RFC1* | -43.2 | $3.15 \times 10^{-9}$ | $2.69 \times 10^{-8}$ | mRNA-intron | 1783 | | | | | |
| AT1G56500 | protein coding gene | | -43.2 | $1.02 \times 10^{-14}$ | $1.78 \times 10^{-13}$ | mRNA-exon | 2007 | | | | | |
| AT4G10210 | protein coding gene | | -43.2 | $4.59 \times 10^{-17}$ | $1.04 \times 10^{-15}$ | mRNA-exon | 1199 | | | | | |
| AT5G13480 | protein coding gene | *FY* | -43.3 | $8.49 \times 10^{-12}$ | $1.04 \times 10^{-10}$ | mRNA-intron | 2060 | | | | | |
| AT3G48660 | protein coding gene | | -43.4 | $1.38 \times 10^{-10}$ | $1.44 \times 10^{-9}$ | mRNA-promoter-TSS | -908 | | | | | |
| AT5G26360 | protein coding gene | | -43.6 | $2.96 \times 10^{-9}$ | $2.55 \times 10^{-8}$ | mRNA-exon | 2277 | | | | | |
| AT1G51530 | protein coding gene | | -43.7 | $4.15 \times 10^{-8}$ | $3.06 \times 10^{-7}$ | mRNA-intron | 897 | | | | | |
| AT3G57110 | protein coding gene | | -43.9 | $1.62 \times 10^{-23}$ | $6.67 \times 10^{-22}$ | mRNA-promoter-TSS | 475 | | | | | |
| AT1G55100 | transposable element gene | | -44.0 | $4.77 \times 10^{-10}$ | $4.58 \times 10^{-9}$ | mRNA-intron | 2352 | | | | | |
| AT1G65830 | pre tRNA | | -44.2 | $1.32 \times 10^{-19}$ | $3.86 \times 10^{-18}$ | tRNA-TTS | 822 | | | | | |
| AT1G79570 | protein coding gene | | -44.5 | $2.78 \times 10^{-14}$ | $4.62 \times 10^{-13}$ | mRNA-exon | 4336 | | | | | |
| AT1G42705 | transposable element gene | | -44.5 | $6.52 \times 10^{-12}$ | $8.08 \times 10^{-11}$ | mRNA-promoter-TSS | -90 | | | | | |
| AT1G48360 | protein coding gene | | -44.7 | $2.80 \times 10^{-25}$ | $1.38 \times 10^{-23}$ | mRNA-exon | 2201 | | | | | |
| AT5G39690 | protein coding gene | *NAC93* | -44.8 | $9.95 \times 10^{-16}$ | $1.97 \times 10^{-14}$ | mRNA-intron | 980 | | | | | |
| AT4G03430 | protein coding gene | *STA1* | -44.9 | $2.21 \times 10^{-10}$ | $2.23 \times 10^{-9}$ | mRNA-exon | 3312 | | | | | |
| AT1G50390 | protein coding gene | | -44.9 | $1.95 \times 10^{-13}$ | $2.92 \times 10^{-12}$ | mRNA-intron | 1826 | | | | | |
| AT1G53780 | protein coding gene | | -45.0 | $6.30 \times 10^{-12}$ | $7.85 \times 10^{-11}$ | mRNA-exon | 922 | | | | | |
| AT3G43920 | protein coding gene | *DCL3* | -45.5 | $1.67 \times 10^{-24}$ | $7.66 \times 10^{-23}$ | mRNA-exon | 3781 | | | | | |
| AT3G22790 | protein coding gene | *NET1A* | -45.7 | $2.98 \times 10^{-9}$ | $2.56 \times 10^{-8}$ | mRNA-exon | 2335 | | | | | |
| AT4G08245 | anti sense lncRNA | | -45.8 | $4.81 \times 10^{-29}$ | $3.31 \times 10^{-27}$ | | -3438 | | | | | |
| AT4G17140 | protein coding gene | | -46.1 | $1.13 \times 10^{-12}$ | $1.54 \times 10^{-11}$ | mRNA-exon | 8322 | | | | | |
| AT2G18140 | protein coding gene | | -46.1 | $9.79 \times 10^{-18}$ | $2.37 \times 10^{-16}$ | Intergenic | -1973 | | | | | |
| AT2G18140 | protein coding gene | | -46.1 | $9.79 \times 10^{-18}$ | $2.37 \times 10^{-16}$ | Intergenic | -1973 | -38.1 | $2.95 \times 10^{-16}$ | $5.09 \times 10^{-14}$ | Intergenic | -1973 |
| AT5G15400 | protein coding gene | | -46.2 | $4.01 \times 10^{-28}$ | $2.51 \times 10^{-26}$ | mRNA-promoter-TSS | -798 | | | | | |
| AT2G07585 | lncRNA | | -46.3 | $1.24 \times 10^{-15}$ | $2.41 \times 10^{-14}$ | lnc RNA-promoter-TSS | -109 | | | | | |
| AT3G56560 | protein coding gene | *NAC65* | -46.4 | $2.02 \times 10^{-30}$ | $1.55 \times 10^{-28}$ | mRNA-intron | 547 | | | | | |
| AT1G07263 | lncRNA | | -46.6 | $1.73 \times 10^{-26}$ | $9.40 \times 10^{-25}$ | lnc RNA-TTS | 167 | | | | | |
| AT1G06157 | lncRNA | | -46.7 | $5.33 \times 10^{-11}$ | $5.89 \times 10^{-10}$ | mRNA-intron | 4939 | | | | | |
| AT2G07042 | other RNA | | -46.8 | $7.29 \times 10^{-15}$ | $1.30 \times 10^{-13}$ | ncRNA-intron | 1198 | | | | | |
| AT1G42705 | transposable element gene | | -46.8 | $8.16 \times 10^{-28}$ | $5.00 \times 10^{-26}$ | mRNA-promoter-TSS | -190 | | | | | |
| AT3G44205 | transposable element gene | | -47.1 | $1.44 \times 10^{-33}$ | $1.47 \times 10^{-31}$ | mRNA-exon | 370 | | | | | |
| AT4G25235 | pseudogene | | -47.1 | $2.84 \times 10^{-7}$ | $1.89 \times 10^{-6}$ | mRNA-intron | 2249 | | | | | |
| AT1G51530 | protein coding gene | | -47.3 | $1.04 \times 10^{-32}$ | $9.75 \times 10^{-31}$ | mRNA-exon | 797 | | | | | |

| | | | | | | | | | | | | |
|---|---|---|---|---|---|---|---|---|---|---|---|---|
| AT1G51530 | protein coding gene | | -47.3 | $1.04 \times 10^{-32}$ | $9.75 \times 10^{-31}$ | mRNA-exon | 797 | -26.4 | $3.44 \times 10^{-4}$ | $6.03 \times 10^{-3}$ | mRNA-exon | 797 |
| AT1G51530 | protein coding gene | | -47.3 | $1.04 \times 10^{-32}$ | $9.75 \times 10^{-31}$ | mRNA-exon | 797 | -44.2 | $3.86 \times 10^{-27}$ | $2.27 \times 10^{-24}$ | mRNA-TTS | 697 |
| AT1G51530 | protein coding gene | | -47.3 | $1.04 \times 10^{-32}$ | $9.75 \times 10^{-31}$ | mRNA-exon | 797 | -59.4 | $6.93 \times 10^{-24}$ | $2.71 \times 10^{-21}$ | mRNA-intron | 997 |
| AT5G35410 | protein coding gene | *SOS2* | -47.4 | $2.04 \times 10^{-7}$ | $1.38 \times 10^{-6}$ | mRNA-promoter-TSS | -648 | | | | | |
| AT3G19170 | protein coding gene | *PREP1* | -47.4 | $9.64 \times 10^{-10}$ | $8.88 \times 10^{-9}$ | mRNA-intron | 3017 | | | | | |
| AT3G07150 | protein coding gene | | -47.9 | $4.15 \times 10^{-14}$ | $6.74 \times 10^{-13}$ | | -1730 | | | | | |
| AT1G06740 | protein coding gene | *MUG3* | -48.0 | $5.08 \times 10^{-12}$ | $6.40 \times 10^{-11}$ | mRNA-promoter-TSS | -886 | | | | | |
| AT2G16660 | protein coding gene | | -48.1 | $2.07 \times 10^{-14}$ | $3.49 \times 10^{-13}$ | | -3361 | | | | | |
| AT5G22220 | protein coding gene | *E2F1* | -48.2 | $8.16 \times 10^{-13}$ | $1.13 \times 10^{-11}$ | mRNA-intron | 1616 | | | | | |
| AT1G08073 | anti sense lncRNA | | -48.3 | $5.97 \times 10^{-11}$ | $6.56 \times 10^{-10}$ | lnc RNA-promoter-TSS | -507 | | | | | |
| AT1G01715 | protein coding gene | | -48.3 | $8.82 \times 10^{-16}$ | $1.75 \times 10^{-14}$ | mRNA-promoter-TSS | -982 | | | | | |
| AT5G61060 | protein coding gene | *HDA5* | -48.4 | $4.43 \times 10^{-19}$ | $1.22 \times 10^{-17}$ | mRNA-exon | 1340 | | | | | |
| AT2G31970 | protein coding gene | *RAD5* | -48.5 | $9.03 \times 10^{-10}$ | $8.36 \times 10^{-9}$ | mRNA-intron | 3020 | | | | | |
| AT3G04015 | anti sense lncRNA | | -48.6 | $2.42 \times 10^{-13}$ | $3.59 \times 10^{-12}$ | | -1220 | | | | | |
| AT1G19630 | protein coding gene | *CYP722A1* | -49.1 | $3.43 \times 10^{-19}$ | $9.56 \times 10^{-18}$ | | -1776 | | | | | |
| AT5G50420 | protein coding gene | | -49.2 | $2.69 \times 10^{-24}$ | $1.21 \times 10^{-22}$ | mRNA-exon | 1305 | | | | | |
| AT3G22690 | protein coding gene | | -49.2 | $3.68 \times 10^{-22}$ | $1.38 \times 10^{-20}$ | mRNA-exon | 2483 | | | | | |
| AT5G57280 | protein coding gene | *RID2* | -49.6 | $4.00 \times 10^{-26}$ | $2.11 \times 10^{-24}$ | mRNA-exon | 1074 | | | | | |
| AT4G04765 | lncRNA | | -49.7 | $6.19 \times 10^{-13}$ | $8.74 \times 10^{-12}$ | lnc RNA-promoter-TSS | -537 | | | | | |
| AT1G16800 | protein coding gene | | -50.5 | $1.42 \times 10^{-18}$ | $3.73 \times 10^{-17}$ | mRNA-exon | 4612 | | | | | |
| AT3G05175 | lncRNA | | -50.7 | $6.67 \times 10^{-24}$ | $2.88 \times 10^{-22}$ | Intergenic | -1422 | | | | | |
| AT4G10420 | protein coding gene | | -50.8 | $5.87 \times 10^{-38}$ | $8.27 \times 10^{-36}$ | transcript-promoter-TSS | 658 | | | | | |
| AT3G57110 | protein coding gene | | -50.9 | $2.56 \times 10^{-48}$ | $5.91 \times 10^{-46}$ | mRNA-promoter-TSS | 575 | | | | | |
| AT4G03040 | protein coding gene | | -50.9 | $5.03 \times 10^{-41}$ | $8.12 \times 10^{-39}$ | mRNA-promoter-TSS | -32 | | | | | |
| AT4G21090 | protein coding gene | *MFDX2* | -51.4 | $4.58 \times 10^{-10}$ | $4.42 \times 10^{-9}$ | | -2891 | | | | | |
| AT3G22290 | protein coding gene | | -52.4 | $5.27 \times 10^{-19}$ | $1.44 \times 10^{-17}$ | mRNA-intron | 1804 | | | | | |
| AT5G36940 | protein coding gene | *CAT3* | -52.7 | $2.56 \times 10^{-14}$ | $4.27 \times 10^{-13}$ | mRNA-exon | 1436 | | | | | |
| AT4G02465 | protein coding gene | | -53.1 | $1.09 \times 10^{-24}$ | $5.11 \times 10^{-23}$ | mRNA-exon | 2389 | | | | | |
| AT1G53165 | protein coding gene | *ATMAP4K ALPHA1* | -53.1 | $2.25 \times 10^{-7}$ | $1.51 \times 10^{-6}$ | mRNA-intron | 2547 | | | | | |
| AT4G09740 | protein coding gene | *GH9B14* | -53.2 | $4.63 \times 10^{-26}$ | $2.42 \times 10^{-24}$ | mRNA-TTS | 2252 | | | | | |
| AT1G21695 | protein coding gene | | -53.3 | $1.13 \times 10^{-9}$ | $1.03 \times 10^{-8}$ | mRNA-intron | 2189 | | | | | |
| AT3G57110 | protein coding gene | | -53.5 | $1.28 \times 10^{-25}$ | $6.45 \times 10^{-24}$ | mRNA-promoter-TSS | 275 | -28.2 | $1.34 \times 10^{-8}$ | $6.84 \times 10^{-7}$ | mRNA-promoter-TSS | 275 |
| AT3G57110 | protein coding gene | | -53.5 | $1.28 \times 10^{-25}$ | $6.45 \times 10^{-24}$ | mRNA-promoter-TSS | 275 | -32.0 | $3.71 \times 10^{-23}$ | $1.43 \times 10^{-20}$ | mRNA-promoter-TSS | 475 |
| AT1G70335 | protein coding gene | | -54.0 | $3.77 \times 10^{-18}$ | $9.44 \times 10^{-17}$ | | -1193 | | | | | |
| AT1G78280 | protein coding gene | | -54.1 | $4.22 \times 10^{-16}$ | $8.63 \times 10^{-15}$ | mRNA-intron | 2796 | | | | | |
| AT1G79960 | protein coding gene | *OFP14* | -54.4 | $5.80 \times 10^{-20}$ | $1.76 \times 10^{-18}$ | mRNA-intron | 2601 | | | | | |
| AT4G08705 | anti sense lncRNA | | -54.7 | $9.89 \times 10^{-24}$ | $4.17 \times 10^{-22}$ | | -1244 | | | | | |

| | | | | | | | | | | | | |
|---|---|---|---|---|---|---|---|---|---|---|---|---|
| AT2G26980 | protein coding gene | *CIPK3* | -54.7 | $6.34 \times 10^{-17}$ | $1.42 \times 10^{-15}$ | | -1617 | | | | | |
| AT2G21410 | protein coding gene | *VHA-A2* | -55.2 | $7.71 \times 10^{-16}$ | $1.53 \times 10^{-14}$ | mRNA-intron | 1135 | | | | | |
| AT3G08695 | snRNA | | -55.6 | $5.41 \times 10^{-10}$ | $5.15 \times 10^{-9}$ | mRNA-intron | 1478 | | | | | |
| AT3G24170 | protein coding gene | *GR1* | -55.8 | $2.72 \times 10^{-29}$ | $1.91 \times 10^{-27}$ | mRNA-exon | 3111 | | | | | |
| AT1G17760 | protein coding gene | *CSTF77* | -55.8 | $2.43 \times 10^{-13}$ | $3.60 \times 10^{-12}$ | mRNA-intron | 3966 | | | | | |
| AT5G16200 | protein coding gene | | -56.0 | $1.14 \times 10^{-32}$ | $1.04 \times 10^{-30}$ | | -1345 | | | | | |
| AT3G14890 | protein coding gene | | -56.0 | $1.67 \times 10^{-17}$ | $3.99 \times 10^{-16}$ | mRNA-exon | 2209 | | | | | |
| AT5G10235 | pre tRNA | | -56.1 | $3.62 \times 10^{-17}$ | $8.28 \times 10^{-16}$ | | -1228 | | | | | |
| AT3G19170 | protein coding gene | *PREP1* | -56.2 | $4.65 \times 10^{-18}$ | $1.16 \times 10^{-16}$ | mRNA-exon | 3617 | | | | | |
| AT3G44205 | transposable element gene | | -56.8 | $1.12 \times 10^{-12}$ | $1.52 \times 10^{-11}$ | mRNA-exon | 1870 | | | | | |
| AT2G40770 | protein coding gene | | -56.9 | $7.19 \times 10^{-14}$ | $1.13 \times 10^{-12}$ | mRNA-exon | 1764 | | | | | |
| AT4G08033 | transposable element gene | | -56.9 | $3.41 \times 10^{-37}$ | $4.57 \times 10^{-35}$ | Intergenic | 3743 | | | | | |
| AT1G42705 | transposable element gene | | -57.3 | $8.33 \times 10^{-23}$ | $3.22 \times 10^{-21}$ | mRNA-promoter-TSS | -290 | | | | | |
| AT5G63640 | protein coding gene | | -57.3 | $8.66 \times 10^{-27}$ | $4.88 \times 10^{-25}$ | mRNA-exon | 2395 | | | | | |
| AT5G20930 | protein coding gene | *TSL* | -57.4 | $2.34 \times 10^{-16}$ | $4.94 \times 10^{-15}$ | mRNA-intron | 2683 | | | | | |
| AT4G10210 | protein coding gene | | -57.8 | $1.24 \times 10^{-9}$ | $1.13 \times 10^{-8}$ | mRNA-intron | 1099 | | | | | |
| AT3G03702 | anti sense lncRNA | | -58.2 | $1.50 \times 10^{-24}$ | $6.96 \times 10^{-23}$ | mRNA-promoter-TSS | -64 | | | | | |
| AT3G03702 | anti sense lncRNA | | -58.2 | $1.50 \times 10^{-24}$ | $6.96 \times 10^{-23}$ | mRNA-promoter-TSS | -64 | -36.1 | $3.90 \times 10^{-6}$ | $1.23 \times 10^{-4}$ | mRNA-promoter-TSS | -64 |
| AT1G42705 | transposable element gene | | -58.8 | $1.57 \times 10^{-30}$ | $1.23 \times 10^{-28}$ | mRNA-promoter-TSS | -490 | | | | | |
| AT1G42705 | transposable element gene | | -58.8 | $1.57 \times 10^{-30}$ | $1.23 \times 10^{-28}$ | mRNA-promoter-TSS | -490 | -26.2 | $2.79 \times 10^{-12}$ | $2.67 \times 10^{-10}$ | mRNA-promoter-TSS | 10 |
| AT1G42705 | transposable element gene | | -58.8 | $1.57 \times 10^{-30}$ | $1.23 \times 10^{-28}$ | mRNA-promoter-TSS | -490 | -33.4 | $2.75 \times 10^{-14}$ | $3.51 \times 10^{-12}$ | mRNA-promoter-TSS | -390 |
| AT1G42705 | transposable element gene | | -58.8 | $1.57 \times 10^{-30}$ | $1.23 \times 10^{-28}$ | mRNA-promoter-TSS | -490 | -34.1 | $3.49 \times 10^{-8}$ | $1.63 \times 10^{-6}$ | mRNA-promoter-TSS | -90 |
| AT1G42705 | transposable element gene | | -58.8 | $1.57 \times 10^{-30}$ | $1.23 \times 10^{-28}$ | mRNA-promoter-TSS | -490 | -34.9 | $1.46 \times 10^{-20}$ | $4.34 \times 10^{-18}$ | mRNA-promoter-TSS | -690 |
| AT1G42705 | transposable element gene | | -58.8 | $1.57 \times 10^{-30}$ | $1.23 \times 10^{-28}$ | mRNA-promoter-TSS | -490 | -39.7 | $6.67 \times 10^{-16}$ | $1.09 \times 10^{-13}$ | mRNA-promoter-TSS | -290 |
| AT1G42705 | transposable element gene | | -58.8 | $1.57 \times 10^{-30}$ | $1.23 \times 10^{-28}$ | mRNA-promoter-TSS | -490 | -39.9 | $1.35 \times 10^{-25}$ | $5.98 \times 10^{-23}$ | mRNA-promoter-TSS | -890 |
| AT1G42705 | transposable element gene | | -58.8 | $1.57 \times 10^{-30}$ | $1.23 \times 10^{-28}$ | mRNA-promoter-TSS | -490 | -41.2 | $1.78 \times 10^{-10}$ | $1.25 \times 10^{-8}$ | mRNA-promoter-TSS | -490 |
| AT1G42705 | transposable element gene | | -58.8 | $1.57 \times 10^{-30}$ | $1.23 \times 10^{-28}$ | mRNA-promoter-TSS | -490 | -44.0 | $3.37 \times 10^{-30}$ | $2.47 \times 10^{-27}$ | mRNA-promoter-TSS | -790 |
| AT1G42705 | transposable element gene | | -58.8 | $1.57 \times 10^{-30}$ | $1.23 \times 10^{-28}$ | mRNA-promoter-TSS | -490 | -45.4 | $4.52 \times 10^{-14}$ | $5.52 \times 10^{-12}$ | mRNA-promoter-TSS | -190 |
| AT3G24120 | protein coding gene | | -59.0 | $5.97 \times 10^{-24}$ | $2.59 \times 10^{-22}$ | mRNA-intron | 1064 | | | | | |
| AT2G43490 | protein coding gene | | -60.5 | $8.85 \times 10^{-13}$ | $1.22 \times 10^{-11}$ | mRNA-exon | 1168 | | | | | |
| AT5G07970 | protein coding gene | | -60.7 | $5.13 \times 10^{-10}$ | $4.91 \times 10^{-9}$ | | -2722 | | | | | |
| AT3G10100 | transposable element gene | | -60.9 | $6.05 \times 10^{-19}$ | $5.29 \times 10^{-16}$ | mRNA-exon | 173 | | | | | |
| AT2G04945 | lncRNA | | -61.7 | $2.04 \times 10^{-33}$ | $2.04 \times 10^{-31}$ | lnc RNA-promoter-TSS | -443 | | | | | |
| AT4G32714 | protein coding gene | *SCRL25* | -62.1 | $1.14 \times 10^{-37}$ | $1.60 \times 10^{-35}$ | | -1437 | | | | | |
| AT1G44150 | pseudogene | | -63.2 | $3.15 \times 10^{-25}$ | $1.53 \times 10^{-23}$ | transcript-exon | 256 | | | | | |
| AT3G44205 | transposable element gene | | -63.3 | $1.34 \times 10^{-50}$ | $3.27 \times 10^{-48}$ | mRNA-exon | 270 | | | | | |
| AT1G04730 | protein coding gene | *CTF18* | -63.4 | $5.79 \times 10^{-22}$ | $2.13 \times 10^{-20}$ | mRNA-intron | 1151 | | | | | |

| | | | | | | | | | | | | |
|---|---|---|---|---|---|---|---|---|---|---|---|---|
| AT1G27900 | protein coding gene | | -65.9 | $7.02 \times 10^{-23}$ | $2.74 \times 10^{-21}$ | mRNA-intron | 2539 | | | | | |
| AT1G05960 | protein coding gene | | -67.3 | $7.43 \times 10^{-16}$ | $1.48 \times 10^{-14}$ | mRNA-exon | 2100 | | | | | |
| AT3G44205 | transposable element gene | | -67.5 | $2.88 \times 10^{-20}$ | $8.95 \times 10^{-19}$ | mRNA-exon | 470 | | | | | |
| AT3G44205 | transposable element gene | | -67.5 | $2.88 \times 10^{-20}$ | $8.95 \times 10^{-19}$ | mRNA-exon | 470 | -31.2 | $3.58 \times 10^{-12}$ | $3.31 \times 10^{-10}$ | mRNA-exon | 1770 |
| AT3G44205 | transposable element gene | | -67.5 | $2.88 \times 10^{-20}$ | $8.95 \times 10^{-19}$ | mRNA-exon | 470 | -38.8 | $1.26 \times 10^{-11}$ | $1.07 \times 10^{-9}$ | mRNA-exon | 470 |
| AT3G44205 | transposable element gene | | -67.5 | $2.88 \times 10^{-20}$ | $8.95 \times 10^{-19}$ | mRNA-exon | 470 | -41.6 | $4.72 \times 10^{-28}$ | $2.91 \times 10^{-25}$ | mRNA-exon | 370 |
| AT1G26130 | protein coding gene | *ALA12* | -67.8 | $9.92 \times 10^{-18}$ | $2.40 \times 10^{-16}$ | mRNA-exon | 1558 | | | | | |
| AT3G48380 | protein coding gene | | -67.9 | $9.74 \times 10^{-28}$ | $5.93 \times 10^{-26}$ | mRNA-exon | 1827 | | | | | |
| AT3G03785 | anti sense lncRNA | | -69.2 | $1.63 \times 10^{-15}$ | $3.12 \times 10^{-14}$ | | -2048 | | | | | |
| AT3G54360 | protein coding gene | *NCA1* | -75.1 | $7.06 \times 10^{-36}$ | $8.75 \times 10^{-34}$ | mRNA-intron | 2362 | | | | | |
| AT2G40730 | protein coding gene | *CTEXP* | -81.3 | $7.96 \times 10^{-33}$ | $7.51 \times 10^{-31}$ | mRNA-intron | 3357 | | | | | |
| AT4G03100 | protein coding gene | | | | | | | 51.2 | $9.00 \times 10^{-23}$ | $3.25 \times 10^{-20}$ | mRNA-promoter-TSS | -609 |
| AT1G44050 | protein coding gene | | | | | | | 36.6 | $2.99 \times 10^{-6}$ | $9.66 \times 10^{-5}$ | mRNA-TTS | 3159 |
| AT3G50630 | protein coding gene | | | | | | | 34.6 | $1.07 \times 10^{-16}$ | $1.99 \times 10^{-14}$ | | -4055 |
| AT5G34857 | transposable element gene | | | | | | | 31.9 | $1.35 \times 10^{-7}$ | $5.68 \times 10^{-6}$ | Intergenic | 2711 |
| AT2G08015 | miRNA | | | | | | | 30.5 | $1.29 \times 10^{-10}$ | $9.20 \times 10^{-9}$ | miRNA-promoter-TSS | -86 |
| AT2G08015 | miRNA | | | | | | | 29.8 | $3.69 \times 10^{-11}$ | $2.88 \times 10^{-9}$ | primary_transcript-TTS | 714 |
| AT4G03970 | transposable element gene | | | | | | | 29.2 | $4.52 \times 10^{-6}$ | $1.41 \times 10^{-4}$ | mRNA-intron | 2261 |
| AT4G08915 | lncRNA | | | | | | | 28.1 | $1.33 \times 10^{-57}$ | $3.47 \times 10^{-54}$ | lnc_RNA-promoter-TSS | -34 |
| AT4G03885 | transposable element gene | | | | | | | 28.1 | $2.78 \times 10^{-6}$ | $9.05 \times 10^{-5}$ | mRNA-exon | 1646 |
| AT5G02650 | protein coding gene | | | | | | | 28.1 | $5.61 \times 10^{-10}$ | $3.67 \times 10^{-8}$ | mRNA-promoter-TSS | -753 |
| AT5G33255 | transposable element gene | | | | | | | 27.4 | $1.12 \times 10^{-8}$ | $5.81 \times 10^{-7}$ | | -1259 |
| AT1G34390 | protein coding gene | *ARF22* | | | | | | 26.6 | $1.49 \times 10^{-4}$ | $3.00 \times 10^{-3}$ | mRNA-exon | 2246 |
| AT2G05240 | transposable element gene | | | | | | | -25.4 | $4.24 \times 10^{-6}$ | $1.33 \times 10^{-4}$ | mRNA-promoter-TSS | -782 |
| AT1G65585 | pseudogene | | | | | | | -25.7 | $2.39 \times 10^{-6}$ | $7.85 \times 10^{-5}$ | mRNA-promoter-TSS | 108 |
| AT3G11310 | protein coding gene | | | | | | | -26.2 | $9.92 \times 10^{-9}$ | $5.18 \times 10^{-7}$ | mRNA-exon | 482 |
| AT2G12440 | transposable element gene | | | | | | | -26.6 | $1.88 \times 10^{-4}$ | $3.63 \times 10^{-3}$ | mRNA-exon | 2111 |
| AT2G10740 | transposable element gene | | | | | | | -26.8 | $2.79 \times 10^{-4}$ | $5.08 \times 10^{-3}$ | mRNA-promoter-TSS | -440 |
| AT5G35965 | transposable element gene | | | | | | | -27.3 | $1.04 \times 10^{-9}$ | $6.49 \times 10^{-8}$ | mRNA-exon | 576 |
| AT2G07950 | anti sense lncRNA | | | | | | | -27.7 | $1.56 \times 10^{-17}$ | $3.33 \times 10^{-15}$ | lnc_RNA-promoter-TSS | 95 |
| AT2G04490 | transposable element gene | | | | | | | -27.8 | $6.14 \times 10^{-20}$ | $1.69 \times 10^{-17}$ | mRNA-exon | 1092 |
| AT2G24761 | pseudogene | | | | | | | -28.1 | $1.73 \times 10^{-5}$ | $4.62 \times 10^{-4}$ | lnc_RNA-promoter-TSS | 177 |
| AT4G08785 | anti sense lncRNA | | | | | | | -28.2 | $8.46 \times 10^{-10}$ | $5.34 \times 10^{-8}$ | mRNA-promoter-TSS | 54 |
| AT5G10010 | protein coding gene | | | | | | | -29.9 | $2.03 \times 10^{-16}$ | $3.55 \times 10^{-14}$ | mRNA-exon | 2123 |
| AT1G65550 | protein coding gene | | | | | | | -30.0 | $1.90 \times 10^{-10}$ | $1.32 \times 10^{-8}$ | mRNA-exon | 1349 |
| AT1G55410 | pseudogene | | | | | | | -31.8 | $5.12 \times 10^{-34}$ | $5.22 \times 10^{-31}$ | transcript-exon | 554 |
| AT2G35000 | protein coding gene | *ATL9* | | | | | | -32.0 | $2.55 \times 10^{-12}$ | $2.46 \times 10^{-10}$ | mRNA-promoter-TSS | -232 |

| | | | | | | | | | | | | |
|---|---|---|---|---|---|---|---|---|---|---|---|---|
| AT2G35000 | protein coding gene | *ATL9* | -32.5 | $2.11 \times 10^{-14}$ | $2.72 \times 10^{-12}$ | mRNA-promoter-TSS | -332 | | | | | |
| AT4G05360 | protein coding gene | | -35.5 | $3.24 \times 10^{-34}$ | $3.46 \times 10^{-31}$ | mRNA-intron | 947 | | | | | |
| AT1G76790 | protein coding gene | *IGMT5* | -36.5 | $1.02 \times 10^{-24}$ | $4.26 \times 10^{-22}$ | Intergenic | -1224 | | | | | |
| AT4G29654 | protein coding gene | | -41.4 | $3.55 \times 10^{-6}$ | $1.14 \times 10^{-4}$ | mRNA-TTS | 1203 | | | | | |
| AT2G24761 | pseudogene | | -41.4 | $1.80 \times 10^{-8}$ | $8.88 \times 10^{-7}$ | lnc_RNA-promoter-TSS | 277 | | | | | |
| AT1G44150 | pseudogene | | -41.6 | $8.95 \times 10^{-18}$ | $1.98 \times 10^{-15}$ | transcript-exon | 256 | | | | | |
| AT1G55380 | protein coding gene | | -53.0 | $2.52 \times 10^{-62}$ | $8.43 \times 10^{-59}$ | Intergenic | -1244 | | | | | |
| AT2G27229 | protein coding gene | | | | | | | 28.13 | $1.76 \times 10^{-92}$ | $1.41 \times 10^{-88}$ | mRNA-promoter-TSS | -6 |

Differential methylation during the first (D1) and second drought (D2) episodes compared between 4 drought-tolerant tolerant ecotypes with high D2/D1 induction (HIG; Na-1, Kin-0, Li-2-1, and Col-0) versus 4 drought-sensitive ecotypes with low D2/D1 induction (LIG; KNO-18, Ru3.1-31, Sq-8, and Pla-3). Differentially methylated regions (DMRs) in the CpG, CHG, and CHH sequence contexts are shown are shown as the methylation difference (MD > 25) with *q*-values (*q* < 0.01), and relative position in base pairs (bp) with respect to the transcription start site (TSS) of each gene. Minus (-) denotes a DMR upstream to the TSS, and plus (+) denotes a DMR downstream to the TSS. The relative gene expression levels in Col-0 under D1 vs control and D2 vs D1 are shown against each differentially methylated gene.

#### Table S12 Differential methylation between the second drought (D2) and the first drought (D1) in *Arabidopsis thaliana* ecotypes

| AGI code | Gene Symbol | D2 vs D1 CpG methylation difference | | | | | D2 vs D1 CHG methylation difference | | | | | D2 vs D1 CHH methylation difference | | | | |
|---|---|---|---|---|---|---|---|---|---|---|---|---|---|---|---|---|
| | | MD | *p*-value | *q*-value | Methylation position | Distance to TSS | MD | *p*-value | *q*-value | Methylation position | Distance to TSS | MD | *p*-value | *q*-value | Methylation position | Distance to TSS |
| **Tolerant (high induction) ecotypes** | | | | | | | | | | | | | | | | |
| AT4G04223 | | 33.2 | $8.21 \times 10^{-7}$ | $9.44 \times 10^{-6}$ | ncRNA-TTS | 1146 | | | | | | | | | | |
| AT2G07700 | | 30.2 | $1.48 \times 10^{-13}$ | $4.29 \times 10^{-12}$ | Intergenic | -1350 | | | | | | | | | | |
| AT3G55650 | | -26.1 | $1.07 \times 10^{-6}$ | $1.21 \times 10^{-5}$ | mRNA-TTS | 766 | | | | | | | | | | |
| AT2G08986 | | | | | | | 42.2 | | | mRNA-exon | 5493 | | | | | |
| AT4G06700 | | | | | | | 30.0 | $2.67 \times 10^{-24}$ | $5.47 \times 10^{-22}$ | mRNA-promoter-TSS | -849 | | | | | |
| AT3G06365 | | | | | | | -35.8 | $1.93 \times 10^{-54}$ | $2.27 \times 10^{-52}$ | lnc_RNA-TTS | 836 | | | | | |
| **Sensitive (low induction) ecotypes** | | | | | | | | | | | | | | | | |
| AT2G10950 | | 62.3 | $8.95 \times 10^{-24}$ | $4.15 \times 10^{-21}$ | mRNA-TTS | 2711 | | | | | | | | | | |
| AT4G36080 | *TRA1B* | 49.8 | $1.14 \times 10^{-11}$ | $1.96 \times 10^{-9}$ | mRNA-intron | 1956 | | | | | | | | | | |
| AT2G09800 | | 48.6 | $4.52 \times 10^{-33}$ | $2.95 \times 10^{-30}$ | mRNA-promoter-TSS | 98 | | | | | | | | | | |
| AT2G05110 | | 47.8 | $3.24 \times 10^{-16}$ | $1.05 \times 10^{-13}$ | mRNA-exon | 1768 | | | | | | | | | | |
| AT3G07355 | | 45.3 | $4.29 \times 10^{-11}$ | $6.73 \times 10^{-9}$ | | -1403 | | | | | | | | | | |
| AT3G18800 | | 40.8 | $3.54 \times 10^{-12}$ | $6.78 \times 10^{-10}$ | mRNA-promoter-TSS | -536 | | | | | | | | | | |
| AT2G35180 | | 40.1 | $3.87 \times 10^{-12}$ | $7.21 \times 10^{-10}$ | transcript-TTS | 588 | | | | | | | | | | |
| AT3G06520 | | 37.8 | $4.14 \times 10^{-11}$ | $6.59 \times 10^{-9}$ | | -1498 | | | | | | | | | | |
| AT1G68700 | | 36.5 | $1.18 \times 10^{-10}$ | $1.70 \times 10^{-8}$ | mRNA-intron | 1858 | | | | | | | | | | |
| AT3G56140 | | 36.4 | $4.81 \times 10^{-9}$ | $4.85 \times 10^{-7}$ | mRNA-intron | 2169 | | | | | | | | | | |
| AT5G08030 | | 35.7 | $8.44 \times 10^{-11}$ | $1.25 \times 10^{-8}$ | mRNA-TTS | 463 | | | | | | | | | | |
| AT4G00710 | *BSK3* | 35.5 | $8.77 \times 10^{-8}$ | $6.51 \times 10^{-6}$ | mRNA-TTS | 2769 | | | | | | | | | | |
| AT5G57110 | *ACA8* | 35.2 | $2.13 \times 10^{-7}$ | $1.39 \times 10^{-5}$ | mRNA-exon | 3252 | | | | | | | | | | |
| AT2G09800 | | 35.0 | $5.56 \times 10^{-12}$ | $1.02 \times 10^{-9}$ | mRNA-exon | 198 | | | | | | | | | | |
| AT1G67130 | | 34.8 | $2.47 \times 10^{-10}$ | $3.28 \times 10^{-8}$ | | -2026 | | | | | | | | | | |
| AT4G08280 | | 34.8 | $3.95 \times 10^{-29}$ | $2.29 \times 10^{-26}$ | mRNA-exon | 1729 | | | | | | | | | | |
| AT4G15230 | *ABCG30* | 34.6 | $3.40 \times 10^{-9}$ | $3.62 \times 10^{-7}$ | mRNA-exon | 3050 | | | | | | | | | | |
| AT1G13640 | | 34.6 | $4.23 \times 10^{-11}$ | $6.69 \times 10^{-9}$ | mRNA-exon | 1743 | | | | | | | | | | |
| AT3G49010 | *BBC1* | 34.6 | $3.57 \times 10^{-6}$ | $1.60 \times 10^{-4}$ | mRNA-intron | 1029 | | | | | | | | | | |
| AT5G00605 | | 34.1 | $1.02 \times 10^{-7}$ | $7.40 \times 10^{-6}$ | transcript-intron | 329 | | | | | | | | | | |
| AT1G07973 | | 33.9 | $8.14 \times 10^{-16}$ | $2.39 \times 10^{-13}$ | lnc_RNA-promoter-TSS | -49 | | | | | | | | | | |
| AT3G23330 | | 33.9 | $1.65 \times 10^{-8}$ | $1.46 \times 10^{-6}$ | mRNA-exon | 1287 | | | | | | | | | | |
| AT1G33700 | | 33.2 | $1.46 \times 10^{-10}$ | $2.05 \times 10^{-8}$ | mRNA-intron | 2132 | | | | | | | | | | |
| AT3G24820 | | 33.0 | $9.39 \times 10^{-9}$ | $8.82 \times 10^{-7}$ | mRNA-intron | 1125 | | | | | | | | | | |
| AT5G63440 | | 32.9 | $2.26 \times 10^{-6}$ | $1.09 \times 10^{-4}$ | | -1301 | | | | | | | | | | |
| AT5G63540 | *RMI1* | 32.7 | $1.79 \times 10^{-7}$ | $1.19 \times 10^{-5}$ | mRNA-intron | 2087 | | | | | | | | | | |
| AT2G33490 | | 32.5 | $6.03 \times 10^{-10}$ | $7.62 \times 10^{-8}$ | mRNA-promoter-TSS | -728 | | | | | | | | | | |
| AT2G01980 | *SOS1* | 32.5 | $5.13 \times 10^{-6}$ | $2.12 \times 10^{-4}$ | mRNA-exon | 3049 | | | | | | | | | | |
| AT3G17930 | *DAC* | 32.3 | $2.21 \times 10^{-14}$ | $5.91 \times 10^{-12}$ | mRNA-promoter-TSS | -541 | | | | | | | | | | |
| AT3G61490 | | 32.2 | $6.03 \times 10^{-12}$ | $1.08 \times 10^{-9}$ | | -1410 | | | | | | | | | | |
| AT5G53170 | *FTSH11* | 32.1 | $1.47 \times 10^{-5}$ | $5.23 \times 10^{-4}$ | mRNA-intron | 2272 | | | | | | | | | | |
| AT4G08250 | | 31.5 | $1.23 \times 10^{-15}$ | $3.56 \times 10^{-13}$ | mRNA-TTS | 1964 | | | | | | | | | | |
| AT4G34035 | | 31.2 | $1.25 \times 10^{-4}$ | $3.06 \times 10^{-3}$ | tRNA-TTS | 1027 | | | | | | | | | | |
| AT4G02075 | *PIT1* | 31.1 | $5.14 \times 10^{-7}$ | $3.02 \times 10^{-5}$ | Intergenic | -1842 | | | | | | | | | | |
| AT2G37680 | | 30.9 | $2.84 \times 10^{-4}$ | $5.81 \times 10^{-3}$ | | -3378 | | | | | | | | | | |
| AT5G59280 | *PUM16* | 30.9 | $6.43 \times 10^{-8}$ | $4.91 \times 10^{-6}$ | | -1112 | | | | | | | | | | |
| AT5G51130 | | 30.9 | $2.88 \times 10^{-7}$ | $1.82 \times 10^{-5}$ | mRNA-TTS | 1693 | | | | | | | | | | |
| AT3G17910 | *SURF1a* | 30.9 | $8.01 \times 10^{-9}$ | $7.63 \times 10^{-7}$ | | -1033 | | | | | | | | | | |
| AT4G14150 | *PAKRP1* | 30.6 | $5.66 \times 10^{-5}$ | $1.58 \times 10^{-3}$ | mRNA-intron | 1963 | | | | | | | | | | |
| AT5G27705 | | 30.6 | $2.10 \times 10^{-10}$ | $2.82 \times 10^{-8}$ | mRNA-promoter-TSS | 82 | | | | | | | | | | |
| AT3G50530 | *CRK* | 30.5 | $1.51 \times 10^{-8}$ | $1.35 \times 10^{-6}$ | mRNA-exon | 1910 | | | | | | | | | | |

| | | | | | | |
|---|---|---|---|---|---|---|
| AT5G42490 | | 30.4 | **$1.99 \times 10^{-5}$** | **$6.69 \times 10^{-4}$** | mRNA-TTS | 3289 |
| AT4G04785 | | 30.3 | **$1.26 \times 10^{-9}$** | **$1.48 \times 10^{-7}$** | lnc_RNA-promoter-TSS | 42 |
| AT5G27400 | | 30.2 | **$1.38 \times 10^{-14}$** | **$3.78 \times 10^{-12}$** | mRNA-promoter-TSS | 817 |
| AT5G27705 | | 30.2 | **$1.97 \times 10^{-10}$** | **$2.66 \times 10^{-8}$** | mRNA-promoter-TSS | 182 |
| AT1G51802 | | 30.0 | **$1.19 \times 10^{-5}$** | **$4.40 \times 10^{-4}$** | | -2668 |
| AT1G09617 | | 30.0 | **$1.65 \times 10^{-7}$** | **$1.11 \times 10^{-5}$** | lnc_RNA-promoter-TSS | -746 |
| AT5G09965 | | 29.7 | **$1.28 \times 10^{-8}$** | **$1.18 \times 10^{-6}$** | Intergenic | -1607 |
| AT5G08139 | | 29.7 | **$1.03 \times 10^{-9}$** | **$1.24 \times 10^{-7}$** | mRNA-exon | 1164 |
| AT1G49230 | | 29.7 | **$5.34 \times 10^{-13}$** | **$1.19 \times 10^{-10}$** | mRNA-exon | 557 |
| AT4G17190 | *FPS2* | 29.5 | **$1.39 \times 10^{-8}$** | **$1.26 \times 10^{-6}$** | mRNA-TTS | 1795 |
| AT5G62230 | *ERL1* | 29.5 | **$9.05 \times 10^{-6}$** | **$3.47 \times 10^{-4}$** | mRNA-exon | 2715 |
| AT1G58230 | | 29.4 | **$5.75 \times 10^{-7}$** | **$3.28 \times 10^{-5}$** | mRNA-exon | 803 |
| AT2G31265 | | 29.1 | **$1.62 \times 10^{-5}$** | **$5.62 \times 10^{-4}$** | mRNA-intron | 2685 |
| AT5G14640 | *SK13* | 29.0 | **$2.64 \times 10^{-9}$** | **$2.91 \times 10^{-7}$** | mRNA-intron | 897 |
| AT5G66300 | *NAC105* | 28.9 | **$2.91 \times 10^{-9}$** | **$3.14 \times 10^{-7}$** | mRNA-exon | 377 |
| AT4G23340 | | 28.9 | **$4.11 \times 10^{-7}$** | **$2.48 \times 10^{-5}$** | mRNA-TTS | 2163 |
| AT5G04290 | *KTF1* | 28.8 | **$8.82 \times 10^{-5}$** | **$2.29 \times 10^{-3}$** | mRNA-exon | 3182 |
| AT5G03925 | | 28.8 | **$2.43 \times 10^{-9}$** | **$2.71 \times 10^{-7}$** | lnc_RNA-promoter-TSS | -371 |
| AT2G03820 | *NMD3* | 28.6 | **$9.18 \times 10^{-23}$** | **$4.07 \times 10^{-20}$** | mRNA-TTS | 2364 |
| AT3G29290 | *emb2076* | 28.6 | **$2.08 \times 10^{-9}$** | **$2.35 \times 10^{-7}$** | Intergenic | 3640 |
| AT5G56440 | | 28.5 | **$5.57 \times 10^{-4}$** | **$9.85 \times 10^{-3}$** | mRNA-exon | 966 |
| AT2G34670 | | 28.5 | **$8.39 \times 10^{-7}$** | **$4.61 \times 10^{-5}$** | mRNA-exon | 4182 |
| AT3G43715 | | 28.5 | **$5.30 \times 10^{-7}$** | **$3.07 \times 10^{-5}$** | Intergenic | -1145 |
| AT3G30840 | | 28.4 | **$1.76 \times 10^{-9}$** | **$2.04 \times 10^{-7}$** | mRNA-promoter-TSS | -525 |
| AT5G30628 | | 28.3 | **$1.42 \times 10^{-10}$** | **$2.00 \times 10^{-8}$** | | -1024 |
| AT1G08757 | | 28.3 | **$4.35 \times 10^{-8}$** | **$3.49 \times 10^{-6}$** | Intergenic | -3120 |
| AT2G39080 | *EMB2799* | 28.2 | **$5.34 \times 10^{-11}$** | **$8.18 \times 10^{-9}$** | mRNA-intron | 1271 |
| AT4G00620 | *EMB3127* | 28.2 | **$1.39 \times 10^{-8}$** | **$1.26 \times 10^{-6}$** | | -1362 |
| AT3G62360 | | 28.2 | **$4.28 \times 10^{-8}$** | **$3.44 \times 10^{-6}$** | mRNA-exon | 1512 |
| AT3G05955 | | 28.2 | **$6.60 \times 10^{-10}$** | **$8.29 \times 10^{-8}$** | lnc_RNA-promoter-TSS | -3 |
| AT5G49420 | | 28.0 | **$6.48 \times 10^{-5}$** | **$1.77 \times 10^{-3}$** | | -2846 |
| AT5G56450 | *PM-ANT* | 28.0 | **$7.96 \times 10^{-10}$** | **$9.94 \times 10^{-8}$** | mRNA-exon | 1111 |
| AT3G19720 | *ARC5* | 27.9 | **$1.37 \times 10^{-8}$** | **$1.26 \times 10^{-6}$** | | -3361 |
| AT5G13690 | *CYL1* | 27.7 | **$1.09 \times 10^{-5}$** | **$4.12 \times 10^{-4}$** | mRNA-intron | 2897 |
| AT5G55380 | | 27.6 | **$1.18 \times 10^{-6}$** | **$6.26 \times 10^{-5}$** | | -2298 |
| AT4G09680 | *CTC1* | 27.6 | **$5.58 \times 10^{-5}$** | **$1.56 \times 10^{-3}$** | mRNA-exon | 1989 |
| AT2G06760 | | 27.4 | **$2.61 \times 10^{-5}$** | **$8.43 \times 10^{-4}$** | mRNA-promoter-TSS | -635 |
| AT1G59550 | | 27.4 | **$5.06 \times 10^{-6}$** | **$2.10 \times 10^{-4}$** | mRNA-promoter-TSS | -163 |
| AT4G34730 | | 27.0 | **$2.49 \times 10^{-7}$** | **$1.59 \times 10^{-5}$** | mRNA-promoter-TSS | -996 |
| AT5G17570 | | 26.8 | **$2.48 \times 10^{-8}$** | **$2.13 \times 10^{-6}$** | mRNA-promoter-TSS | -675 |
| AT4G24770 | *RBP31* | 26.8 | **$3.56 \times 10^{-9}$** | **$3.74 \times 10^{-7}$** | mRNA-promoter-TSS | -512 |
| AT3G22990 | | 26.4 | **$1.34 \times 10^{-6}$** | **$6.98 \times 10^{-5}$** | mRNA-promoter-TSS | 810 |
| AT3G62110 | | 26.3 | **$1.47 \times 10^{-6}$** | **$7.55 \times 10^{-5}$** | | -1530 |
| AT1G73970 | | 26.3 | **$9.27 \times 10^{-9}$** | **$8.75 \times 10^{-7}$** | mRNA-exon | 1153 |
| AT3G29380 | *pBRP2* | 26.3 | **$1.01 \times 10^{-11}$** | **$1.75 \times 10^{-9}$** | mRNA-exon | 173 |
| AT2G03670 | *CDC48B* | 26.2 | **$1.52 \times 10^{-5}$** | **$5.38 \times 10^{-4}$** | mRNA-promoter-TSS | -834 |
| AT1G11800 | | 26.0 | **$4.14 \times 10^{-6}$** | **$1.79 \times 10^{-4}$** | mRNA-exon | 1039 |
| AT3G12140 | *EML1* | 26.0 | **$2.51 \times 10^{-7}$** | **$1.61 \times 10^{-5}$** | mRNA-intron | 1765 |
| AT1G34260 | *FAB1D* | 26.0 | **$4.97 \times 10^{-5}$** | **$1.42 \times 10^{-3}$** | mRNA-promoter-TSS | 431 |
| AT5G39980 | | 25.9 | **$1.36 \times 10^{-5}$** | **$4.92 \times 10^{-4}$** | mRNA-exon | 1021 |
| AT1G65740 | *UCL1* | 25.9 | **$4.60 \times 10^{-9}$** | **$4.68 \times 10^{-7}$** | mRNA-promoter-TSS | -640 |
| AT2G32260 | *CCT1* | 25.8 | **$1.33 \times 10^{-10}$** | **$1.89 \times 10^{-8}$** | mRNA-exon | 2666 |
| AT5G08570 | | 25.7 | **$4.30 \times 10^{-5}$** | **$1.26 \times 10^{-3}$** | mRNA-exon | 1126 |
| AT4G23100 | *GSH1* | 25.7 | **$3.02 \times 10^{-7}$** | **$1.90 \times 10^{-5}$** | mRNA-exon | 1848 |
| AT3G02660 | *EMB2768* | 25.7 | **$2.28 \times 10^{-7}$** | **$1.48 \times 10^{-5}$** | mRNA-TTS | 1391 |
| AT5G13780 | | 25.7 | **$1.01 \times 10^{-5}$** | **$3.83 \times 10^{-4}$** | mRNA-TTS | 924 |
| AT1G68090 | *ANN5* | 25.5 | **$3.09 \times 10^{-10}$** | **$3.98 \times 10^{-8}$** | mRNA-TTS | 844 |
| AT5G11350 | | 25.3 | **$2.88 \times 10^{-6}$** | **$1.33 \times 10^{-4}$** | mRNA-intron | 1197 |
| AT1G18060 | | 25.3 | **$4.58 \times 10^{-5}$** | **$1.33 \times 10^{-3}$** | | -1960 |

| | | | | | | |
|---|---|---|---|---|---|---|
| AT5G35750 | *HK2* | 25.2 | **$6.56 \times 10^{-9}$** | **$6.36 \times 10^{-7}$** | Intergenic | -3446 |
| AT4G32470 | | 25.2 | **$1.07 \times 10^{-9}$** | **$1.28 \times 10^{-7}$** | mRNA-exon | 1743 |
| AT2G14605 | | -25.0 | **$3.08 \times 10^{-8}$** | **$2.57 \times 10^{-6}$** | Intergenic | 2287 |
| AT5G34853 | | -25.1 | **$1.64 \times 10^{-4}$** | **$3.78 \times 10^{-3}$** | mRNA-promoter-TSS | -327 |
| AT1G32520 | | -25.1 | **$3.73 \times 10^{-5}$** | **$1.12 \times 10^{-3}$** | mRNA-promoter-TSS | -327 |
| AT2G22730 | | -25.2 | **$6.81 \times 10^{-14}$** | **$1.63 \times 10^{-11}$** | mRNA-TTS | 346 |
| AT4G16250 | *PHYD* | -25.2 | **$5.82 \times 10^{-11}$** | **$8.85 \times 10^{-9}$** | mRNA-exon | 1389 |
| AT2G15870 | | -25.3 | **$2.42 \times 10^{-4}$** | **$5.11 \times 10^{-3}$** | mRNA-exon | 761 |
| AT2G17043 | | -25.3 | **$8.40 \times 10^{-8}$** | **$6.25 \times 10^{-6}$** | mRNA-exon | 2988 |
| AT1G28350 | | -25.4 | **$2.18 \times 10^{-4}$** | **$4.72 \times 10^{-3}$** | | -1586 |
| AT5G06600 | *UBP12* | -25.4 | **$9.40 \times 10^{-5}$** | **$2.41 \times 10^{-3}$** | mRNA-intron | 1292 |
| AT2G48060 | | -25.6 | **$1.57 \times 10^{-6}$** | **$7.98 \times 10^{-5}$** | mRNA-exon | 5966 |
| AT5G63280 | | -25.6 | **$4.70 \times 10^{-5}$** | **$1.36 \times 10^{-3}$** | mRNA-TTS | 1620 |
| AT3G07160 | | -25.7 | **$1.94 \times 10^{-4}$** | **$4.29 \times 10^{-3}$** | mRNA-intron | 3032 |
| AT1G06630 | | -25.7 | **$7.64 \times 10^{-5}$** | **$2.03 \times 10^{-3}$** | mRNA-exon | 1112 |
| AT1G53645 | | -25.7 | **$3.89 \times 10^{-5}$** | **$1.16 \times 10^{-3}$** | mRNA-promoter-TSS | -995 |
| AT1G80880 | | -25.8 | **$3.81 \times 10^{-9}$** | **$3.99 \times 10^{-7}$** | mRNA-TTS | 1870 |
| AT3G28380 | *ABCB17* | -25.8 | **$1.04 \times 10^{-12}$** | **$2.21 \times 10^{-10}$** | mRNA-TTS | 3150 |
| AT4G14240 | | -26.0 | **$1.55 \times 10^{-6}$** | **$7.90 \times 10^{-5}$** | mRNA-exon | 1412 |
| AT1G28550 | *RABA1i* | -26.0 | **$1.34 \times 10^{-6}$** | **$6.97 \times 10^{-5}$** | mRNA-promoter-TSS | -979 |
| AT1G19770 | *PUP14* | -26.2 | **$4.97 \times 10^{-6}$** | **$2.07 \times 10^{-4}$** | | -2574 |
| AT4G26610 | *D6PKL1* | -26.2 | **$4.05 \times 10^{-7}$** | **$2.45 \times 10^{-5}$** | mRNA-TTS | 2174 |
| AT2G11080 | | -26.3 | **$7.89 \times 10^{-6}$** | **$3.10 \times 10^{-4}$** | mRNA-promoter-TSS | -889 |
| AT4G32200 | *ASY2* | -26.4 | **$1.16 \times 10^{-9}$** | **$1.37 \times 10^{-7}$** | mRNA-exon | 2011 |
| AT4G00070 | | -26.4 | **$9.10 \times 10^{-5}$** | **$2.35 \times 10^{-3}$** | mRNA-intron | 4975 |
| AT1G16800 | | -26.4 | **$2.22 \times 10^{-5}$** | **$7.36 \times 10^{-4}$** | mRNA-intron | 4412 |
| AT1G31740 | *BGAL15* | -26.5 | **$2.73 \times 10^{-6}$** | **$1.27 \times 10^{-4}$** | mRNA-intron | 1988 |
| AT5G04055 | | -26.8 | **$1.55 \times 10^{-5}$** | **$5.46 \times 10^{-4}$** | lnc_RNA-promoter-TSS | -103 |
| AT1G28880 | | -26.8 | **$1.24 \times 10^{-5}$** | **$4.56 \times 10^{-4}$** | tRNA-promoter-TSS | -76 |
| AT3G18730 | *TSK* | -26.9 | **$2.59 \times 10^{-5}$** | **$8.36 \times 10^{-4}$** | mRNA-exon | 3250 |
| AT2G11970 | | -26.9 | **$2.20 \times 10^{-8}$** | **$1.89 \times 10^{-6}$** | mRNA-exon | 4052 |
| AT5G51190 | | -27.1 | **$4.47 \times 10^{-6}$** | **$1.90 \times 10^{-4}$** | mRNA-intron | 2086 |
| AT1G08857 | | -27.3 | **$5.08 \times 10^{-9}$** | **$5.09 \times 10^{-7}$** | mRNA-exon | 537 |
| AT4G36630 | *EMB2754* | -27.4 | **$3.16 \times 10^{-4}$** | **$6.33 \times 10^{-3}$** | mRNA-exon | 3175 |
| AT5G58140 | *PHOT2* | -27.5 | **$2.81 \times 10^{-4}$** | **$5.75 \times 10^{-3}$** | mRNA-exon | 929 |
| AT4G25270 | *OTP70* | -27.5 | **$2.56 \times 10^{-10}$** | **$3.37 \times 10^{-8}$** | mRNA-promoter-TSS | -741 |
| AT1G24470 | *KCR2* | -27.5 | **$1.03 \times 10^{-7}$** | **$7.49 \times 10^{-6}$** | | -1963 |
| AT3G43715 | | -27.5 | **$7.16 \times 10^{-5}$** | **$1.94 \times 10^{-3}$** | mRNA-promoter-TSS | 55 |
| AT1G01790 | *KEA1* | -27.8 | **$3.96 \times 10^{-5}$** | **$1.18 \times 10^{-3}$** | mRNA-exon | 3191 |
| AT4G38215 | | -27.9 | **$5.34 \times 10^{-8}$** | **$4.22 \times 10^{-6}$** | lnc_RNA-promoter-TSS | -962 |
| AT1G23360 | *MENG* | -28.3 | **$1.99 \times 10^{-6}$** | **$9.70 \times 10^{-5}$** | mRNA-exon | 772 |
| AT1G64340 | | -28.3 | **$5.40 \times 10^{-14}$** | **$1.37 \times 10^{-11}$** | mRNA-exon | 526 |
| AT2G25030 | | -28.4 | **$3.52 \times 10^{-6}$** | **$1.58 \times 10^{-4}$** | transcript-exon | 188 |
| AT2G35630 | *MOR1* | -28.5 | **$4.74 \times 10^{-4}$** | **$8.70 \times 10^{-3}$** | mRNA-intron | 4216 |
| AT1G19870 | *IQD32* | -28.7 | **$2.36 \times 10^{-6}$** | **$1.13 \times 10^{-4}$** | mRNA-exon | 1267 |
| AT2G31320 | *PARP1* | -28.8 | **$1.12 \times 10^{-4}$** | **$2.79 \times 10^{-3}$** | mRNA-intron | 2999 |
| AT3G56500 | | -28.9 | **$1.12 \times 10^{-4}$** | **$2.79 \times 10^{-3}$** | mRNA-TTS | 1013 |
| AT1G15480 | | -28.9 | **$2.02 \times 10^{-5}$** | **$6.75 \times 10^{-4}$** | | -1222 |
| AT2G08295 | | -29.3 | **$3.23 \times 10^{-13}$** | **$7.40 \times 10^{-11}$** | lnc_RNA-TTS | 601 |
| AT5G01455 | | -30.0 | **$1.73 \times 10^{-4}$** | **$3.93 \times 10^{-3}$** | | -2671 |
| AT1G03240 | | -30.1 | **$3.47 \times 10^{-11}$** | **$5.57 \times 10^{-9}$** | mRNA-TTS | 409 |
| AT4G05815 | | -30.3 | **$2.70 \times 10^{-9}$** | **$2.97 \times 10^{-7}$** | | -1207 |
| AT5G13050 | *5-FCL* | -30.3 | **$5.74 \times 10^{-9}$** | **$5.67 \times 10^{-7}$** | Intergenic | -1259 |
| AT4G32220 | | -30.3 | **$1.98 \times 10^{-25}$** | **$9.59 \times 10^{-23}$** | mRNA-exon | 749 |
| AT4G08028 | | -30.4 | **$4.82 \times 10^{-8}$** | **$3.84 \times 10^{-6}$** | mRNA-TTS | 321 |
| AT5G42270 | *VAR1* | -31.0 | **$4.33 \times 10^{-11}$** | **$6.73 \times 10^{-9}$** | mRNA-exon | 1376 |
| AT2G23470 | *RUS4* | -31.1 | **$2.02 \times 10^{-14}$** | **$5.46 \times 10^{-12}$** | mRNA-TTS | 2258 |
| AT1G02890 | | -31.2 | **$5.20 \times 10^{-7}$** | **$3.03 \times 10^{-5}$** | mRNA-intron | 1356 |
| AT2G06315 | | -31.5 | **$5.57 \times 10^{-14}$** | **$1.37 \times 10^{-11}$** | mRNA-promoter-TSS | 285 |

| | | | | | | | | | | | | | | | | |
|---|---|---|---|---|---|---|---|---|---|---|---|---|---|---|---|---|
| AT3G51630 | *WNK5* | -31.6 | **$4.75 \times 10^{-9}$** | **$4.81 \times 10^{-7}$** | | -2520 | | | | | | | | | | |
| AT3G15260 | | -31.6 | **$6.36 \times 10^{-14}$** | **$1.54 \times 10^{-11}$** | mRNA-promoter-TSS | -633 | | | | | | | | | | |
| AT5G63510 | *GAMMA CAL1* | -31.9 | **$1.47 \times 10^{-4}$** | **$3.49 \times 10^{-3}$** | mRNA-TTS | 1467 | | | | | | | | | | |
| AT5G05570 | | -32.7 | **$2.99 \times 10^{-6}$** | **$1.38 \times 10^{-4}$** | mRNA-exon | 3185 | | | | | | | | | | |
| AT5G60760 | | -32.8 | **$3.69 \times 10^{-16}$** | **$1.18 \times 10^{-13}$** | mRNA-exon | 1623 | | | | | | | | | | |
| AT4G16150 | | -32.9 | **$5.57 \times 10^{-14}$** | **$1.37 \times 10^{-11}$** | mRNA-intron | 4492 | | | | | | | | | | |
| AT4G02460 | *PMS1* | -33.0 | **$1.03 \times 10^{-9}$** | **$1.24 \times 10^{-7}$** | mRNA-exon | 1271 | | | | | | | | | | |
| AT5G35615 | | -33.0 | **$2.31 \times 10^{-9}$** | **$2.59 \times 10^{-7}$** | transcript-promoter-TSS | -368 | | | | | | | | | | |
| AT1G12360 | *KEU* | -33.2 | **$4.04 \times 10^{-9}$** | **$4.17 \times 10^{-7}$** | mRNA-exon | 1287 | | | | | | | | | | |
| AT1G44542 | | -33.3 | **$4.43 \times 10^{-13}$** | **$9.92 \times 10^{-11}$** | Intergenic | 3736 | | | | | | | | | | |
| AT1G79030 | | -33.4 | **$1.98 \times 10^{-5}$** | **$6.69 \times 10^{-4}$** | mRNA-exon | 727 | | | | | | | | | | |
| AT3G14140 | | -33.4 | **$3.18 \times 10^{-11}$** | **$5.15 \times 10^{-9}$** | mRNA-TTS | 1305 | | | | | | | | | | |
| AT4G08033 | | -33.7 | **$3.12 \times 10^{-11}$** | **$5.09 \times 10^{-9}$** | Intergenic | 3943 | | | | | | | | | | |
| AT2G41790 | | -33.9 | **$1.86 \times 10^{-11}$** | **$3.11 \times 10^{-9}$** | mRNA-intron | 4054 | | | | | | | | | | |
| AT1G07967 | | -34.1 | **$4.95 \times 10^{-14}$** | **$1.29 \times 10^{-11}$** | lnc_RNA-TTS | 542 | | | | | | | | | | |
| AT5G47030 | | -34.3 | **$8.83 \times 10^{-18}$** | **$3.18 \times 10^{-15}$** | | -3399 | | | | | | | | | | |
| AT4G01110 | | -34.5 | **$1.28 \times 10^{-9}$** | **$1.49 \times 10^{-7}$** | mRNA-TTS | 1609 | | | | | | | | | | |
| AT1G01960 | *EDA10* | -35.1 | **$7.32 \times 10^{-8}$** | **$5.53 \times 10^{-6}$** | mRNA-intron | 4861 | | | | | | | | | | |
| AT5G02100 | *UNE18* | -35.7 | **$5.84 \times 10^{-10}$** | **$7.43 \times 10^{-8}$** | mRNA-exon | 699 | | | | | | | | | | |
| AT1G75210 | | -36.0 | **$8.54 \times 10^{-6}$** | **$3.31 \times 10^{-4}$** | mRNA-intron | 2614 | | | | | | | | | | |
| AT5G01930 | *MAN6* | -36.2 | **$4.35 \times 10^{-5}$** | **$1.28 \times 10^{-3}$** | mRNA-TTS | 850 | | | | | | | | | | |
| AT4G27430 | *CIP7* | -36.8 | **$7.79 \times 10^{-8}$** | **$5.85 \times 10^{-6}$** | mRNA-exon | 1972 | | | | | | | | | | |
| AT5G36960 | | -36.9 | **$5.66 \times 10^{-8}$** | **$4.43 \times 10^{-6}$** | | -1256 | | | | | | | | | | |
| AT3G52957 | | -37.8 | **$1.15 \times 10^{-27}$** | **$5.98 \times 10^{-25}$** | mRNA-promoter-TSS | 127 | | | | | | | | | | |
| AT2G46550 | | -37.8 | **$1.06 \times 10^{-10}$** | **$1.55 \times 10^{-8}$** | | -3924 | | | | | | | | | | |
| AT1G51965 | *ABO5* | -38.1 | **$2.02 \times 10^{-5}$** | **$6.75 \times 10^{-4}$** | mRNA-TTS | 1744 | | | | | | | | | | |
| AT5G28680 | *ANX2* | -39.4 | **$1.65 \times 10^{-12}$** | **$3.27 \times 10^{-10}$** | mRNA-promoter-TSS | 262 | | | | | | | | | | |
| AT4G29980 | | -39.7 | **$1.74 \times 10^{-17}$** | **$5.96 \times 10^{-15}$** | | -1668 | | | | | | | | | | |
| AT4G39460 | *SAMC1* | -39.7 | **$1.61 \times 10^{-17}$** | **$5.59 \times 10^{-15}$** | mRNA-exon | 5329 | | | | | | | | | | |
| AT2G34250 | | -39.9 | **$1.41 \times 10^{-8}$** | **$1.27 \times 10^{-6}$** | mRNA-intron | 1565 | | | | | | | | | | |
| AT3G14850 | *TBL41* | -40.0 | **$5.24 \times 10^{-16}$** | **$1.63 \times 10^{-13}$** | | -3175 | | | | | | | | | | |
| AT5G46400 | *PRP39-2* | -40.7 | **$1.68 \times 10^{-10}$** | **$2.30 \times 10^{-8}$** | mRNA-promoter-TSS | -82 | | | | | | | | | | |
| AT3G54100 | | -41.6 | **$8.45 \times 10^{-29}$** | **$4.76 \times 10^{-26}$** | mRNA-exon | 1849 | | | | | | | | | | |
| AT3G60860 | *BIG2* | -42.5 | **$3.66 \times 10^{-7}$** | **$2.23 \times 10^{-5}$** | mRNA-exon | 3754 | | | | | | | | | | |
| AT1G04303 | | -47.1 | **$9.22 \times 10^{-12}$** | **$1.63 \times 10^{-9}$** | lnc_RNA-TTS | 863 | | | | | | | | | | |
| AT1G14770 | | -47.9 | **$4.81 \times 10^{-18}$** | **$1.79 \times 10^{-15}$** | mRNA-exon | 1173 | | | | | | | | | | |
| AT5G00885 | | | | | | | 32.8 | **$1.66 \times 10^{-11}$** | **$7.09 \times 10^{-9}$** | ncRNA-promoter-TSS | 92 | | | | | |
| AT5G30584 | | | | | | | 32.43 | **$8.12 \times 10^{-7}$** | **$1.32 \times 10^{-4}$** | mRNA-exon | 3987 | | | | | |
| AT2G05110 | | | | | | | 27.58 | **$8.63 \times 10^{-9}$** | **$2.45 \times 10^{-6}$** | mRNA-exon | 1768 | | | | | |
| AT2G06995 | | | | | | | 27.13 | **$1.38 \times 10^{-7}$** | **$2.80 \times 10^{-5}$** | lnc_RNA-promoter-TSS | 37 | | | | | |
| AT2G11430 | | | | | | | 27.08 | **$1.15 \times 10^{-6}$** | **$1.76 \times 10^{-4}$** | mRNA-exon | 1482 | | | | | |
| AT2G07789 | | | | | | | 26.7 | **$2.88 \times 10^{-5}$** | **$2.44 \times 10^{-3}$** | mRNA-exon | 2403 | | | | | |
| AT5G28469 | | | | | | | 26.05 | **$3.52 \times 10^{-5}$** | **$2.93 \times 10^{-3}$** | Intergenic | 1642 | | | | | |
| AT1G35860 | *TOC75-I* | | | | | | -26.94 | **$2.16 \times 10^{-5}$** | **$1.96 \times 10^{-3}$** | mRNA-exon | 999 | | | | | |
| AT1G35895 | | | | | | | -27.24 | **$1.42 \times 10^{-5}$** | **$1.38 \times 10^{-3}$** | mRNA-intron | 5174 | | | | | |
| AT4G06585 | | | | | | | -28.48 | **$2.00 \times 10^{-7}$** | **$3.89 \times 10^{-5}$** | mRNA-exon | 1081 | | | | | |
| AT1G06973 | | | | | | | -31.89 | **$1.13 \times 10^{-4}$** | **$7.11 \times 10^{-3}$** | lnc_RNA-TTS | 739 | | | | | |
| AT4G32880 | *HB-8* | | | | | | -37.36 | **$1.26 \times 10^{-7}$** | **$2.57 \times 10^{-5}$** | mRNA-promoter-TSS | -601 | | | | | |
| AT1G77880 | | | | | | | | | | | | -31 | **3.08E-38** | **$2.31 \times 10^{-35}$** | mRNA-TTS | 677 |

Differential methylation between the second drought (D2) and the first drought (D1) in 4 drought-tolerant tolerant ecotypes with high D2/D1 induction (HIG; Na-1, Kin-0, Li-2-1, and Col-0) and 4 drought-sensitive ecotypes with low D2/D1 induction (LIG; KNO-18, Ru3.1-31, Sq-8, and Pla-3). Differentially methylated regions (DMRs) in the CpG, CHG, and CHH sequence contexts are shown as the methylation difference (MD > 25) with $q$-values ($q < 0.01$), and relative position in base pairs (bp) with respect to the transcription start site (TSS) of each gene. Minus (-) denotes a DMR upstream to the TSS, and plus (+) denotes a DMR downstream to the TSS. The relative gene expression levels in Col-0 under D1 vs control and D2 vs D1 are shown against each differentially methylated gene.

### Table S13 Differential methylation of eGWAS-delineated genes under recurrent drought

| AGI code | Gene Symbol | Short Description | HIG vs LIG CpG methylation difference in D2 | | | | HIG vs LIG CHG methylation difference in D2 | | | | HIG vs LIG CHH methylation difference in D2 | | | |
|---|---|---|---|---|---|---|---|---|---|---|---|---|---|---|
| | | | MD | *p*-value | *q*-value | Methylation position (bp) | MD | *p*-value | *q*-value | Methylation position (bp) | MD | *p*-value | *q*-value | Methylation position (bp) |
| ***LKR* eGWAS** | | | | | | | | | | | | | | |
| AT2G29080 | *FTSH3* | FTSH protease 3 | **47.69** | $8.69 \times 10^{-16}$ | $1.73 \times 10^{-14}$ | Intron (1178) | | | | | | | | |
| AT2G34660 | *ABCC2* | Multidrug resistance-associated protein 2 | **27.21** | $1.70 \times 10^{-4}$ | $6.93 \times 10^{-4}$ | Intron (2739) | | | | | | | | |
| AT2G29070 | | Ubiquitin fusion degradation UFD1 family protein | **26.25** | $2.69 \times 10^{-9}$ | $2.33 \times 10^{-8}$ | Promoter (-497) | | | | | | | | |
| AT4G38062 | | Basic helix loop helix (bHLH) DNA-binding superfamily protein | 15.63 | $5.61 \times 10^{-10}$ | $5.34 \times 10^{-9}$ | Exon (449) | | | | | | | | |
| AT1G31920 | | Tetratricopeptide repeat (TPR)-like superfamily protein | 14.05 | $5.24 \times 10^{-7}$ | $3.35 \times 10^{-6}$ | Exon (806) | | | | | | | | |
| AT3G15430 | | Regulator of chromosome condensation (RCC1) family protein | 9.26 | $1.68 \times 10^{-4}$ | $6.86 \times 10^{-4}$ | Exon (1086) | | | | | | | | |
| AT3G04340 | *TSL2* | FtsH extracellular protease family | 5.91 | $3.16 \times 10^{-3}$ | $9.47 \times 10^{-3}$ | Promoter (-1301) | | | | | | | | |
| AT2G19120 | | P-loop containing nucleoside triphosphate hydrolases superfamily protein | -10.56 | $1.30 \times 10^{-3}$ | $4.32 \times 10^{-3}$ | Intron (1447) | | | | | | | | |
| AT2G19110 | *HMA4* | Arabidopsis heavy metal ATPase 4 | -11.81 | $2.90 \times 10^{-5}$ | $1.38 \times 10^{-4}$ | Exon (5914) | | | | | | | | |
| AT5G04090 | | Histidine-tRNA ligase | **-37.68** | $2.53 \times 10^{-25}$ | $1.25 \times 10^{-23}$ | Promoter (-1669) | | | | | | | | |
| AT5G16200 | | 50S ribosomal protein-like protein | **-55.96** | $1.14 \times 10^{-32}$ | $1.04 \times 10^{-40}$ | Promoter (-1345) | | | | | | | | |
| ***HIS1-3* eGWAS** | | | | | | | | | | | | | | |
| AT3G25110 | *FATA1* | Fata acyl-acp thioesterase | **33.44** | $1.42 \times 10^{-13}$ | $2.17 \times 10^{-12}$ | Exon-intron boundary (1374) | | | | | | | | |
| AT3G01780 | *TPLATE* | ARM repeat superfamily protein | **25.28** | $1.71 \times 10^{-21}$ | $6.04 \times 10^{-20}$ | Exon (1482) | | | | | | | | |
| AT5G13000 | *GSL12* | Glucan synthase-like 12 | 22.31 | $1.00 \times 10^{-3}$ | $3.41 \times 10^{-3}$ | Intron (3176) | | | | | | | | |
| AT5G08550 | *ILP1* | Increased level of polyploidy1-1d | 16.48 | $1.86 \times 10^{-8}$ | $1.45 \times 10^{-7}$ | Promoter (-1469) | | | | | | | | |
| AT5G11210 | *GLR2.5* | Glutamate receptor 2.5 | 16.26 | $6.52 \times 10^{-17}$ | $1.45 \times 10^{-15}$ | Exon (2764) | | | | | | | | |
| AT1G14230 | | GDA1/CD39 nucleoside phosphatase family protein | 14.10 | $1.16 \times 10^{-6}$ | $7.06 \times 10^{-6}$ | Promoter-TSS (-617) | | | | | | | | |
| AT5G13010 | *EMB3011* | RNA helicase family protein | 9.16 | $5.01 \times 10^{-4}$ | $1.84 \times 10^{-3}$ | Intron (2351) | | | | | | | | |
| AT2G20740 | | Tetraspanin family protein | 8.73 | $2.52 \times 10^{-4}$ | $9.89 \times 10^{-4}$ | Promoter-TSS (-61) | | | | | | | | |
| AT2G38630 | *FBIP2* | D FBS interacting protein 2 | 7.45 | $8.77 \times 10^{-5}$ | $3.82 \times 10^{-4}$ | Exon (828) | | | | | | | | |
| AT3G28480 | | Oxoglutarate/iron-dependent oxygenase | -8.50 | $3.56 \times 10^{-5}$ | $1.68 \times 10^{-4}$ | Intron (854) | | | | | | | | |
| AT3G61730 | *RMF1* | Reduced male fertility | -10.70 | $1.35 \times 10^{-3}$ | $4.47 \times 10^{-3}$ | Intron-exon boundary (624) | | | | | | | | |
| AT5G11200 | *UAP56B* | DEAD/DEAH box RNA helicase family protein | -16.96 | $3.02 \times 10^{-3}$ | $9.10 \times 10^{-3}$ | Exon (2276) | | | | | | | | |
| AT1G16800 | | P-loop containing nucleoside triphosphate hydrolases superfamily protein | -21.13 | $3.86 \times 10^{-5}$ | $1.80 \times 10^{-4}$ | Exon (3112) | | | | | | | | |
| AT4G03430 | *EMB2770* | Embryo defective 2770 | **-44.87** | $2.21 \times 10^{-10}$ | $2.23 \times 10^{-9}$ | Exon (3312) | | | | | | | | |
| AT1G14220 | | Ribonuclease T2 family protein | | | | | | | | | -13.31 | $1.14 \times 10^{-6}$ | $2.04 \times 10^{-5}$ | Promoter-TSS (-690) |
| AT4G05420 | *DDB1A* | Damaged dna binding protein 1a | | | | | -9.18 | $2.25 \times 10^{-6}$ | $7.43 \times 10^{-5}$ | Intron (2829) | | | | |
| AT4G26450 | | Hypothetical protein | | | | | | | | | -6.06 | $3.27 \times 10^{-9}$ | $1.55 \times 10^{-7}$ | Exon (479) |
| ***DREB1A* eGWAS** | | | | | | | | | | | | | | |
| AT5G01950 | | Leucine-rich repeat protein kinase family protein | **57.79** | $1.58 \times 10^{-8}$ | $1.25 \times 10^{-7}$ | Exon (1749) | | | | | | | | |
| AT5G23575 | | Transmembrane CLPTM1 family protein | **46.47** | $7.99 \times 10^{-13}$ | $1.11 \times 10^{-11}$ | Exon (3166) | | | | | | | | |
| AT1G12920 | *ERF1-2* | Eukaryotic release factor 1-2 | **44.26** | $2.15 \times 10^{-11}$ | $2.50 \times 10^{-10}$ | Upstream (-2558) | | | | | | | | |
| AT5G23580 | *CPK12* | *Arabidopsis thaliana* calmodulin-like domain protein kinase 9 | 21.59 | $5.07 \times 10^{-9}$ | $4.24 \times 10^{-8}$ | Intron-exon boundary (1368) | | | | | | | | |
| AT1G14850 | *NUP155* | Nucleoporin 155 | 17.30 | $1.15 \times 10^{-4}$ | $4.89 \times 10^{-4}$ | Exon (1208) | | | | | | | | |
| AT4G21300 | *ATDEK36* | Tetratricopeptide repeat (TPR)-like superfamily protein | 16.63 | $5.66 \times 10^{-4}$ | $2.05 \times 10^{-3}$ | Exon (1033) | | | | | | | | |
| AT5G12920 | | Transducin/WD40 repeat-like superfamily protein | 11.49 | $3.31 \times 10^{-3}$ | $9.86 \times 10^{-3}$ | Intron (1832) | | | | | | | | |

| | | | | | | | | | | | | | | |
|---|---|---|---|---|---|---|---|---|---|---|---|---|---|---|
| AT4G21310 | *DEAL2* | DUF1218 family member | 7.96 | **$2.78 \times 10^{-5}$** | **$1.33\times 10^{-4}$** | Promoter-exon boundary (334) | | | | | | | | |
| AT4G00740 | *QUA3* | Quasimodo 3 | -22.12 | **$2.19 \times 10^{-3}$** | **$6.87\times 10^{-3}$** | Intron (2233) | | | | | | | | |
| AT5G62110 | | Homeodomain-like superfamily protein | -23.37 | **$7.12 \times 10^{-13}$** | **$9.95\times 10^{-12}$** | Promoter-TSS (-964) | -8.45 | **$1.63 \times 10^{-4}$** | **$3.23\times 10^{-3}$** | Promoter-TSS (-964) | | | | |
| AT3G48030 | *ATL48* | Arabidopsis toxicos en levadura 48 | **-28.24** | **$3.08 \times 10^{-10}$** | **$3.05\times 10^{-9}$** | Exon (1103) | | | | | | | | |
| AT5G35750 | *HK2* | Histidine kinase 2 | | | | | | | | | 8.49 | **$6.41 \times 10^{-6}$** | **$8.19\times 10^{-5}$** | Promoter-TSS (-746) |
| AT5G37650 | | Hypothetical protein (DUF577) | | | | | | | | | -11.73 | **$1.11 \times 10^{-5}$** | **$1.27\times 10^{-4}$** | Promoter-TSS (-417) |

The genes identified by eGWAS of *LKR*, *HIS1-3*, and *DREB1A* having differential methylation during the second drought (D2) between 4 ecotypes with high D2/D1 induction (HIG; Na-1, Kin-0, Li-2-1, and Col-0) versus 4 ecotypes with low D2/D1 induction (LIG; KNO-18, Ru3.1-31, Sq-8, and Pla-3). Differentially methylated regions (DMRs) in the CpG, CHG, and CHH sequence contexts are shown are shown as the methylation difference (MD) with $q$-values ($q < 0.01$), and relative position in base pairs (bp) with respect to the transcription start site (TSS) of each gene. Minus (-) denotes a DMR upstream to the TSS, and plus (+) denotes a DMR downstream to the TSS. HIG: high induction group, LIG: low induction group.
